# Genomic transfers help to decipher the ancient evolution of filoviruses and interactions with vertebrate hosts

**DOI:** 10.1101/2023.11.29.569234

**Authors:** Derek J. Taylor, Max H. Barnhart

## Abstract

Although several filoviruses are dangerous human pathogens, there is conflicting evidence regarding their origins and interactions with animal hosts. Here we attempt to improve this understanding using the paleoviral record over a geological time scale, protein structure predictions, tests for evolutionary maintenance, and phylogenetic methods that alleviate sources of bias and error. We found evidence for long branch attraction bias in the L gene tree for filoviruses, and that using codon-specific models and protein structural comparisons of paleoviruses ameliorated conflict and bias. We found evidence for four ancient filoviral groups, each with extant viruses and paleoviruses with open reading frames. Furthermore, we found evidence of repeated transfers of filovirus-like elements to mouse-like rodents. A filovirus-like nucleoprotein ortholog with an open reading frame was detected in three subfamilies of spalacid rodents (present since the Miocene). These elements were unique among the detected filovirus-like paleoviruses in possessing open reading frames, expression products, and evidence for purifying selection. Our finding of structural conservation over geological time for paleoviruses informs virus and paleovirus discovery methods. Our results resolve a deep conflict in the evolutionary framework for filoviruses and reveal that genomic transfers to vertebrate hosts with potentially functional co-options have been more widespread than previously appreciated.

**Author Summary:** Filoviruses are a family of RNA viruses discovered in 1967 and notorious for spillover of the dangerous pathogens, Ebola virus and Marburg virus. However, their origins, deeper relations, diversity, and interactions with animal hosts remain controversial. Part of the confusion may be that differing rates of evolution among divergent viral lineages can create a bias termed long branch attraction (LBA). We tested for this scenario in the L protein gene sequence of filoviruses and found evidence that LBA is occurring leading to a false pairing of filovirus lineages associated with a fish and a snake. We found that using nucleotides instead of amino acids when inferring trees, paleoviral sequences with open reading frames, additional conserved genes, and comparisons of predicted protein structures can resolve the LBA. We found four major groups of filoviruses, with the paleoviral record and trees being consistent with a fish origin for the family. Moreover, we found evidence of a filovirus-like element in spalacid rodents that has been evolutionarily maintained at the open-reading frame, amino acid sequence and structural level for over 20 million years. This element was also expressed in the liver, a target of filoviral infections. We conclude that genomic interactions of filoviruses with vertebrates, including the co-option of viral genes, are more important than previously appreciated.

## Introduction

Despite the ongoing importance of RNA viruses as zoonotic human pathogens and key components of ecosystems, we know little about their pre-historic evolution and host interactions. However, recent advances in paleovirology, evolutionary genomics, structural biology and artificial intelligence are enabling rapid insights[1–5]. There is now evidence that many families of RNA viruses have long evolutionary histories of interactions with vertebrate hosts (including gene co-option) and branching orders that recapitulate those from ancient host phylogenies[6]. Still, host jumps have occurred (often from prey to predator) and the propensity to harbor viruses that jump hosts varies among host taxa[2].

One group that still has mysterious origins and host interactions is the filoviruses – a family of negative stranded unsegmented RNA genomes whose best-known members are dangerous human pathogens, Ebola virus (EBOV) and Marburg virus (MARV). Filovirus genomes contain from 6 to 10 genes with the nucleoprotein (NP) and RNA-dependent RNA polymerase (L) genes being the only apparent common homologs. In all known filoviruses, save the thamnoviruses, the first gene (3’) is NP, the second gene is the polymerase cofactor (viral protein 35, VP35) and the last gene is L[7]. For decades, filoviruses were thought to be closely related African viruses that diverged less than a few thousand years ago[8, 9]. However, the discovery of filovirus-like elements (paleoviruses) and filoviruses from several continents suggested that the group was much more diverse and ancient than previously proposed[10–13]. Indeed, fossil-calibrated orthologs from mammalian genomes indicated that the family itself is older than tens of millions of years and that host infections have occurred in marsupials and eutherians [11, 14–16]. Some of these paleoviruses have the potential to be co-opted elements that function in the host [17]. For example, an open reading frame (ORF) of a VP35-like element has been maintained by purifying selection throughout a radiation of mouse-eared bats and relatives and has a similar protein structure to that found in the extant human pathogen EBOV[18–20]. Another ORF for VP35-like elements has been identified in African spiny mice[21]. Putative filoviral co-options in marsupials lack an ORF but show tissue-specific RNA expression [11, 16]. Nevertheless, there are no known filovirus-like elements with extended ORFs, evidence for purifying selection, and evidence for expression.

There is regional evidence that rodents harbor RNA viruses with the greatest potential for host jumps and shrews harbor the greatest richness of viruses[2]. While rodents are important non-human study systems for filovirus infection[22], natural filovirus-host interactions with rodents and shrews remain poorly studied. Although the immune responses differ among host taxa, adult rodents fail to show significant disease with wild-type filoviral infections[23], suggesting independently evolved immune adaptions to filoviral infections. Indeed, the genomes of rodents and shrews have filovirus-like paleoviral elements indicating prior infections with filoviruses [11]. Taylor et al.[14] reported filovirus-like orthologs in three genomes of hamsters and voles that were phylogenetically nested inside the clade that contains human pathogens, (orthomarburgviruses and orthoebolaviruses). The genomic location of this VP35-like element was the last intron of TAX1BP1, a gene that has recently become known as a key player in the mammalian immune protection against viruses via CD4+ T cell immunity [24, 25]. Notably, the evolved protection of hamsters to EBOV requires CD4+ cells and CD4 dependent antibody responses[26]. While the location of an EBOV-like element within a key gene for the immune response in hamsters to EBOV is suggestive of a co-opted immune element, we know little about the biological implications or evolutionary maintenance of the element.

Recently, partial viral genome sequences related to MARV and to EBOV were obtained from bats in China[13, 27]. Moreover, seven genomes of filoviruses with marked sequence divergences from MARV were assembled from non-mammalian vertebrate transcriptomes (from percomorph fishes and a snake[6, 7, 28]). Shi et al. [6] found that one of the fish filoviruses (*Striavirus antennarii*, Xīlǎng virus, XILV) was basal to mammal-associated viruses even when fossil calibrated mouse/rat paleoviral elements[11] are included on the tree. This suggests that divergent fish filoviruses are far older than the tens of millions of years from known mammal calibrations. Geoghan et al.[29] later detected fragments (68-69 aa) of filovirus-like L-protein sequence in a percomorph (blue spotted goatfish) and a zeiform (John Dory fish). These grouped with HUJV (*Thamnovirus thamnaconi*, Huángjiāo virus) and are consistent with a fish origin for filoviruses. Presently, all of the known piscine filovirus-like sequences are from fishes of the acanthomorph clade which includes percomorphs and Zeiformes[30]. Many groups of fishes beyond the acanthomorph clade remain poorly sampled. The result is several deep phylogenetic gaps in potential vertebrate hosts of filoviruses. However, a divergent filovirus genome (Tapajós virus, TAPV, *Tapjovirus bothropis*) was recently assembled from the transcriptome of a common lancehead snake (*Bothrops atrox* (Linnaeus, 1758)[7]. TAPV formed a well-supported phylogenetic sister group (based on the conserved L or RDRP protein) with the fish-associated virus, XILV[7]. The L gene is essentially the standard for comparing divergent RNA viral genomes because this gene often the longest and most conserved element of viral genomes. At face value, the L tree suggests a host jump between fish and reptiles. However, for TAPV, the gene order and nucleotide sequence identity was similar to mammal-associated filoviruses[7, 21]. Moreover, gene trees (NP and VP35, but without fish-associated sequences or an outgroup) support the basal position of TAPV to one of the mammal-associated clades of paleoviruses[21]. A retrovirus-like domain in the glycoprotein gene of TAPV groups with lizards, while a similar element in mammal-associated filoviruses groups with cartilaginous fish[31]. Presently, these conflicts hinder our understanding of the deeper relations of filoviruses and the importance of host co-option of filovirus-like elements.

In this study, we examine over 400 filovirus-like paleoviruses within vertebrate genomes, seeking a deeper understanding of the ancient evolution and interactions of filoviruses with their hosts. We mitigated potential biases and artifacts, by employing simulations, codon-partitioned substitution models, taxon additions with paleoviral sequences, and comparisons of predicted protein structure distances between viruses and paleoviruses. Additionally, we scrutinized the evolutionary maintenance of putative co-opted elements in vertebrate genomes. This approach aims to inform about filovirus-vertebrate interactions over a geological time scale.

## Results

### Long branch attraction in filoviruses and its resolution

The phylogenetic analysis based on unfiltered L-protein amino acid sequences from filoviruses revealed a distinctive TAPV-XILV grouping in the phylogram, characterized by two extended branches connected by a short intermediary branch (Fig 1A). This configuration raised concerns about a potential long branch artifact (LBA). Furthermore, significant differences in amino acid composition were identified between sequences linked to fish and those associated with tetrapods (Table S1). When the phylogram was based on methods that are less susceptible to LBA[32], (i.e., nucleotides with a partitioned codon model including all or only first and second positions), TAPV grouped with the sequences from genomes with a shared architecture, the mammalian MARV-like clade, instead of with the fish-associated virus, XILV (Fig 1B; Figs S1-S2). Another method that can reduce LBA due to amino acid site heterogeneity, PMSF (Posterior Mean Site Frequency profiles analysis [33]), found the same tree as the putative LBA phylogram (Fig S3). We carried out simulations to assess if the branch lengths and amino acid compositions are sufficient to form a long branch attraction involving TAPV and XILV. In the first simulation, the clade observed from the L-protein amino acid tree (TAPV/XILV) is enforced in the constraint parameter. As expected, most of the trees (95%) estimated from simulated alignments were consistent with the constrained TAPV/XILV clade (Fig 2A). However, when simulations were constrained to favor the non-LBA grouping (TAPV/MARV-like grouping, Fig 2B), the putative LBA grouping of TAPV/XILV remained predominant, being recovered in 61% of the simulations. A further simulation with the same TAPV/MARV-like clade constraint but with terminal branches leading to TAPV and XILV shortened by approximately half, reduced the putative LBA grouping of TAPV/XILV to 6% (Fig 2C). Another approach that alleviated LBA was the addition of outgroup sequences from related families of RNA viruses (paramyxoviruses, lispiviruses, and rhabdoviruses). This approach also led to strong support for the monophyly of filoviruses (Fig. 3), and L-like paleoviral elements from Neotropical opossums being placed within the MARV-like clade.

**Fig 1.**
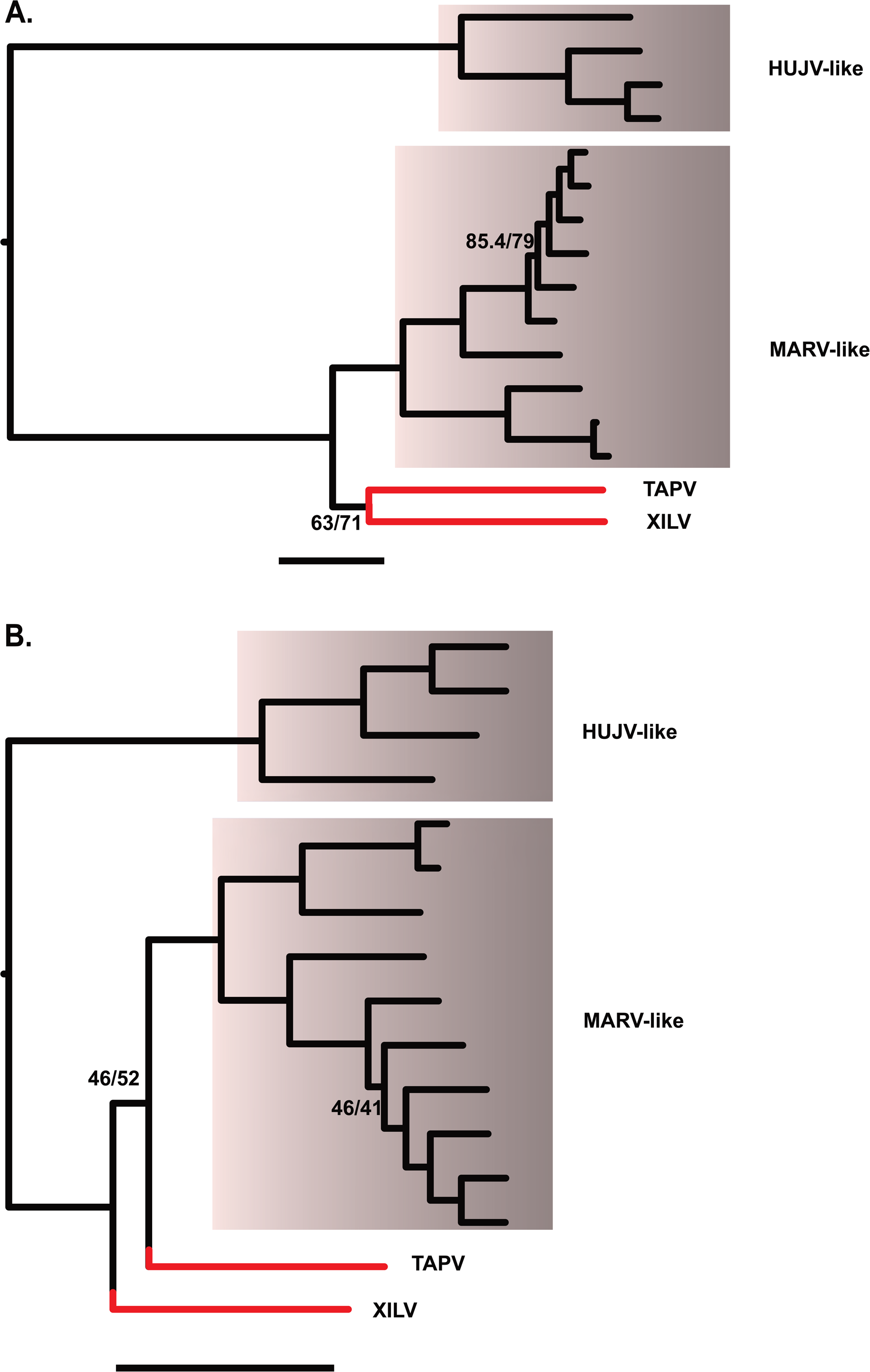
Phylogenies showing relationships of four deep lineages/clades of known filoviruses. The x-axis for each graph is proportional in length to genetic distances (the scale bar is 1 substitution/site). Acronyms are Huángjiāo virus (HUJV), Marburg virus (MARV), Tapajós virus (TAPV), and Xīlǎng virus (XILV). Note that TAPV and XILV form a long branch pair with the amino acid data. Numbers represent branches with the lowest approximate likelihood ratio tests and bootstrap values. The remaining branches had support greater than 96. A. ML tree based on the L Protein amino acid sequence alignment B. ML tree based on partitioned codon model (nucleotides) for the same alignment as in A.

**Fig 2.**
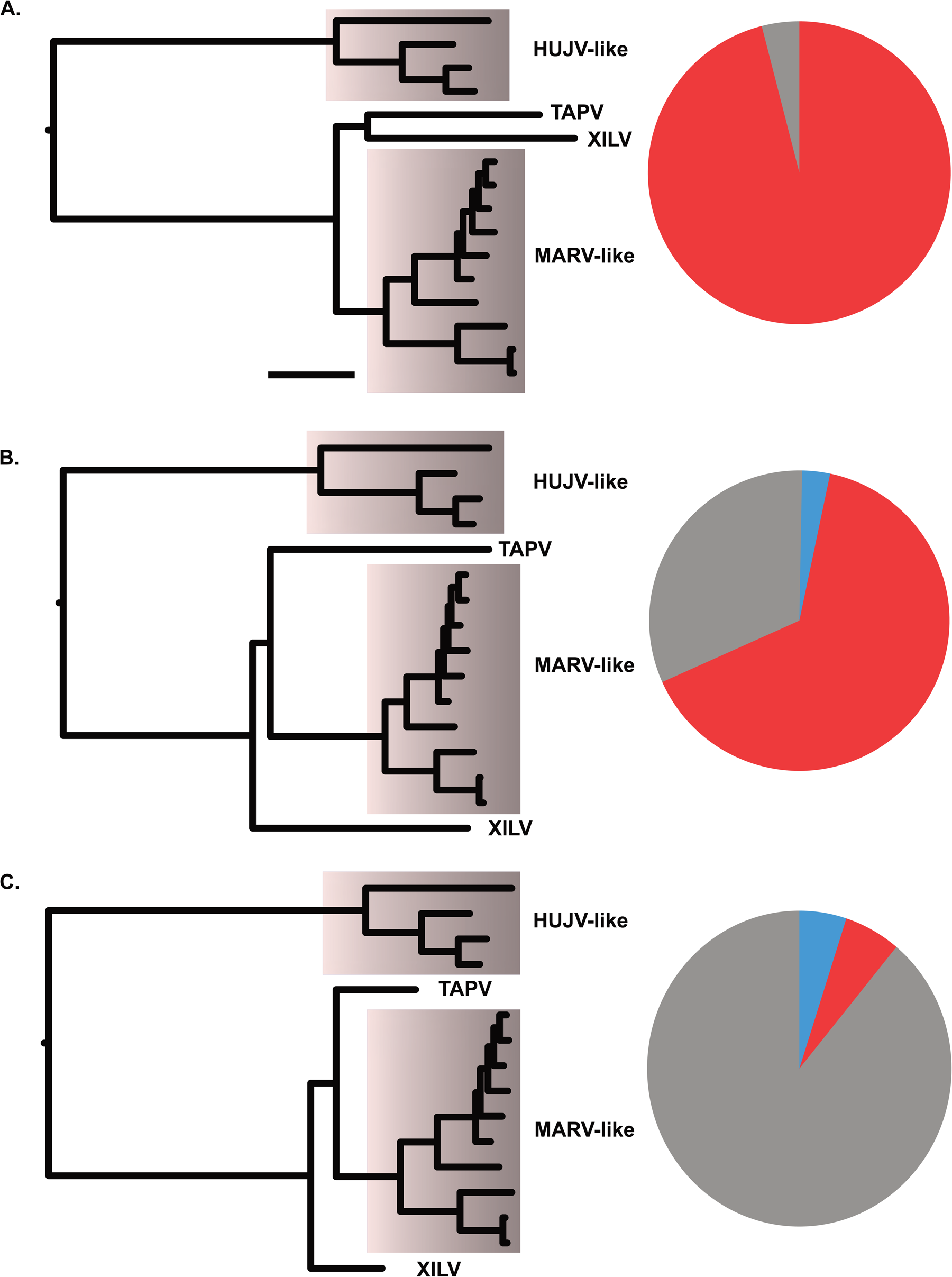
Proportions of topologies observed from maximum likelihood analysis of alignments from parametric simulations that included tree parameters (shown on left side cartoon). Acronyms are Huángjiāo virus (HUJV), Marburg virus (MARV), Tapajós virus (TAPV), and Xīlǎng virus (XILV). A) a TAPV/XILV sister group, B) a TAPV basal to MARV-like taxa with observed branch lengths, or C) TAPV basal to MARV-like taxa where the branches leading to TAPV and XILV are shortened to 0.5 in length. Red fill on the pie graphs indicates proportion of simulations with a TAPV/XILV group (putative long branch attraction), gray indicates proportion of simulations with a TAPV basal to MARV-like taxa, and blue indicates proportion of topologies observed that differ from red or gray.

**Fig 3.**
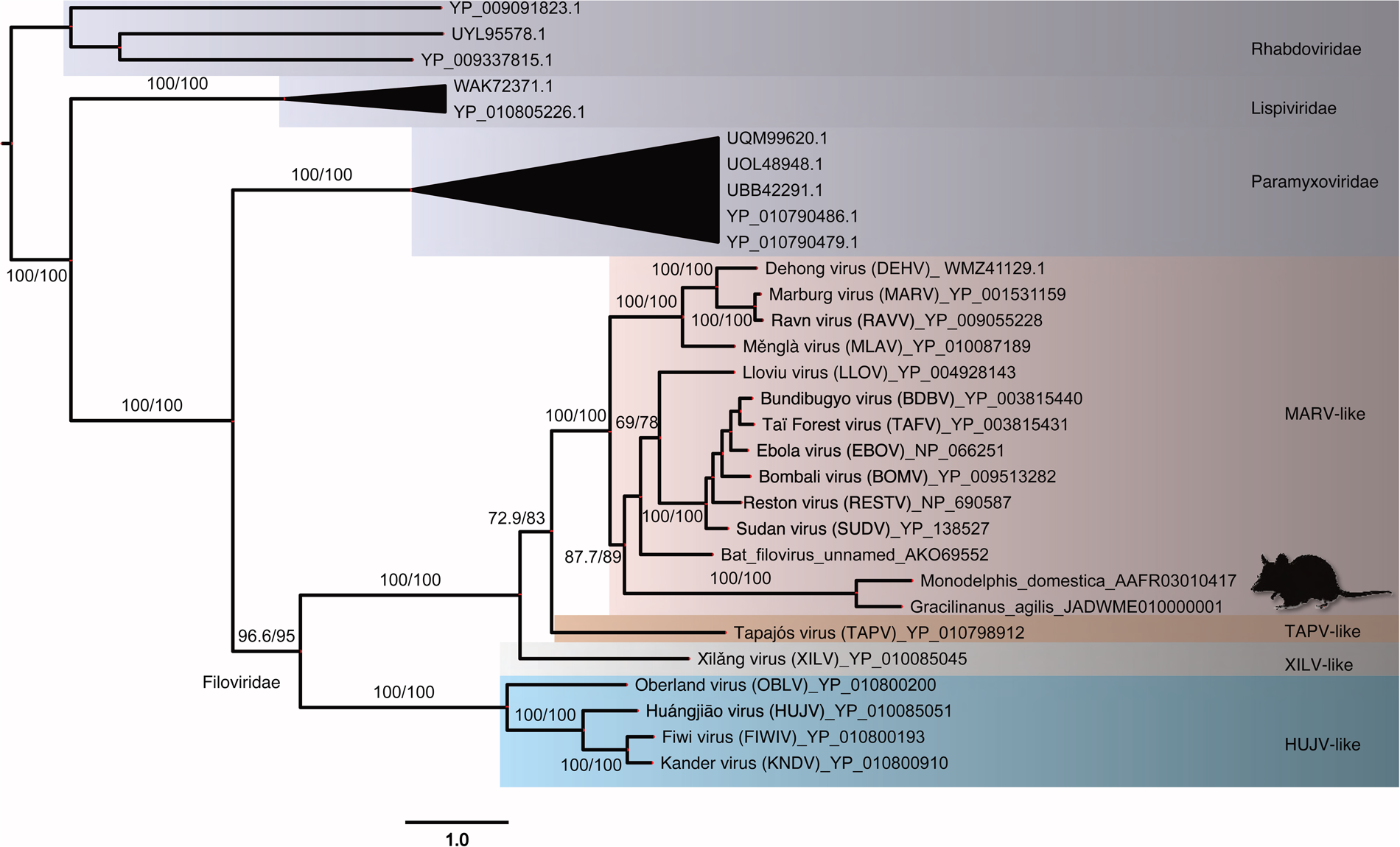
Maximum Likelihood phylogram of L-protein amino acid sequences from filoviruses and filovirus-like paleoviruses with outgroup rooting using sequences of rhabdoviruses. Numbers represent support values from approximate likelihood ratios and ultrafast bootstraps. Major clades of filoviruses are named after the original sequence of each group.

### Affiliations of a snake-associated filoviruses with paleoviruses

We then estimated phylograms from the NP and VP35 gene regions of the filovirus genome. These regions evolve at different rates than the L-protein regions and unlike the L-protein analyses, we initially added paleoviruses from vertebrate genomes with open reading frames that might “break up” long branches. With the NP gene, four deep clades were apparent that we termed HUJV-like, XILV-like, TAPV-like, and MARV-like after the first described filovirus in each clade. Notably, the putative LBA is absent in the NP data as the TAPV-like clade groups with MARV-like sequences associated with mammals with strong support values (Fig 4). Indeed, TAPV groups with open reading frame paleoviruses from three spalacid rodents (bamboo rats, mole-rats, and zokors). The XILV branch is paired with paleoviruses from the genome of the freshwater fish, *Paedocypris*. Switching the data and model for the NP gene to nucleotides with a partitioned codon models increased the support values of the TAPV/spalacid rodent grouping from 59/78 to above 96 (Figs S4-S5). The main supported topological difference between data types was the movement of the paleoviruses from *Borostomias* from the base of the HUJV-like clade (AA tree) to within the HUJV-like clade (partitioned codon model tree). When the analysis of the NP-like sequences is expanded to include paleoviruses with disrupted reading frames, TAPV groups strongly with mammalian sequences (Fig 5, S6). However, TAPV is nested within a clade of paleoviruses from shrews (*Sorex* sp.) with strong support values. We also note that NP-like paleoviruses from *Acomys* (African spiny mice) are nested within the cricetid rodent clade that is more closely related to EBOV than MARV is (Fig 5). Indeed, while several groups of mammals are represented in the paleoviruses by single paleoviruses or small monophyletic clades of paleoviruses (vespertilionid bats, a tenrec, anteaters, shrews, opossums, diprotodontid marsupials and tarsiers), filovirus-like sequences are common in every rodent suborder (save Sciuromorpha) and dispersed throughout the NP-like phylogeny (with some groups such as cricetid rodents present in several divergent phylogenetic groups).

**Fig 4.**
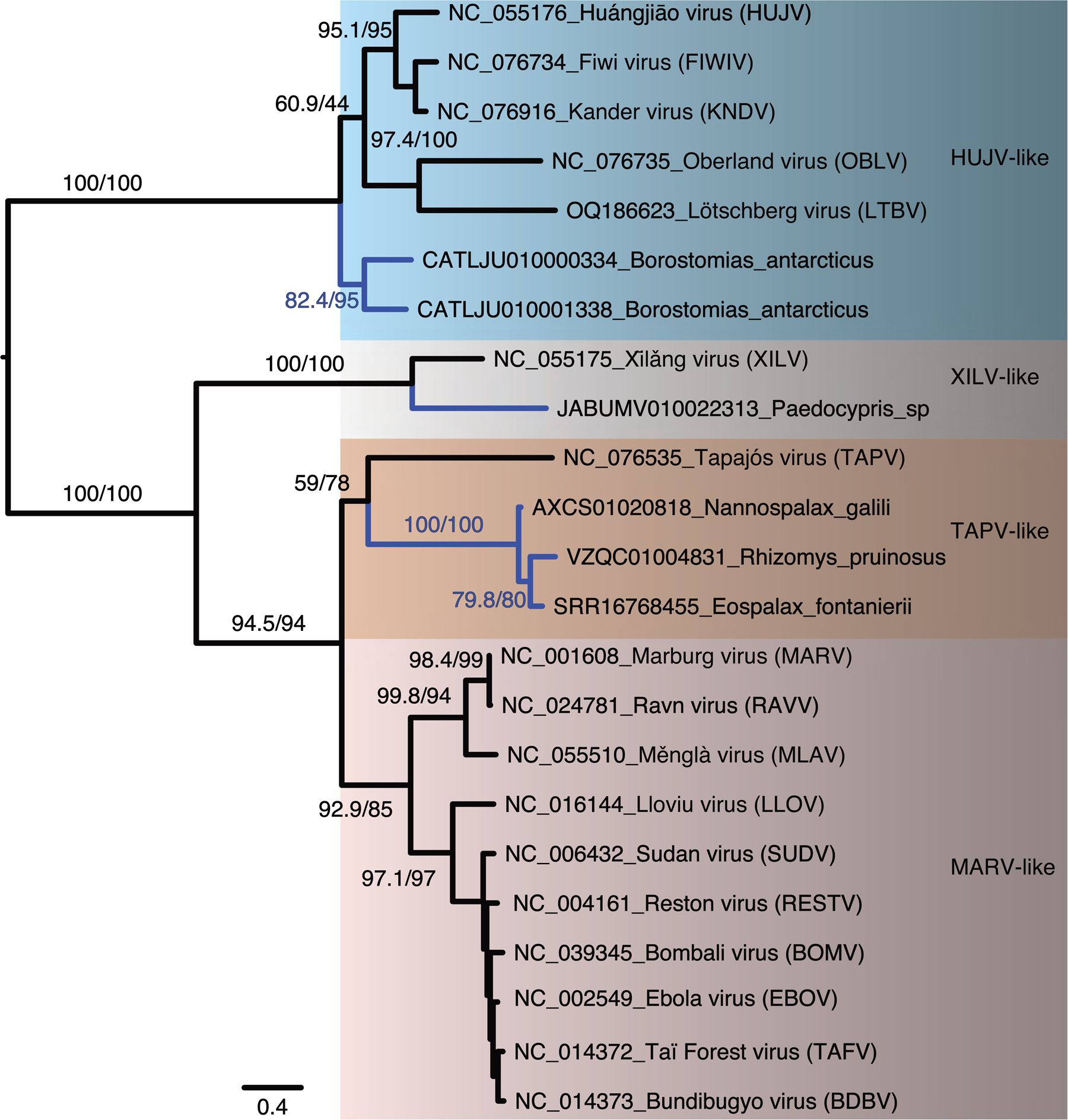
Maximum likelihood phylogram based on amino acid sequences of the nucleoprotein gene for filoviruses and filovirus-like paleoviruses (from vertebrate genomes) with open reading frames (blue lines). Four major clades are identified. Numbers represent approximate likelihood ratio test values and bootstrap values.

**Fig 5.**
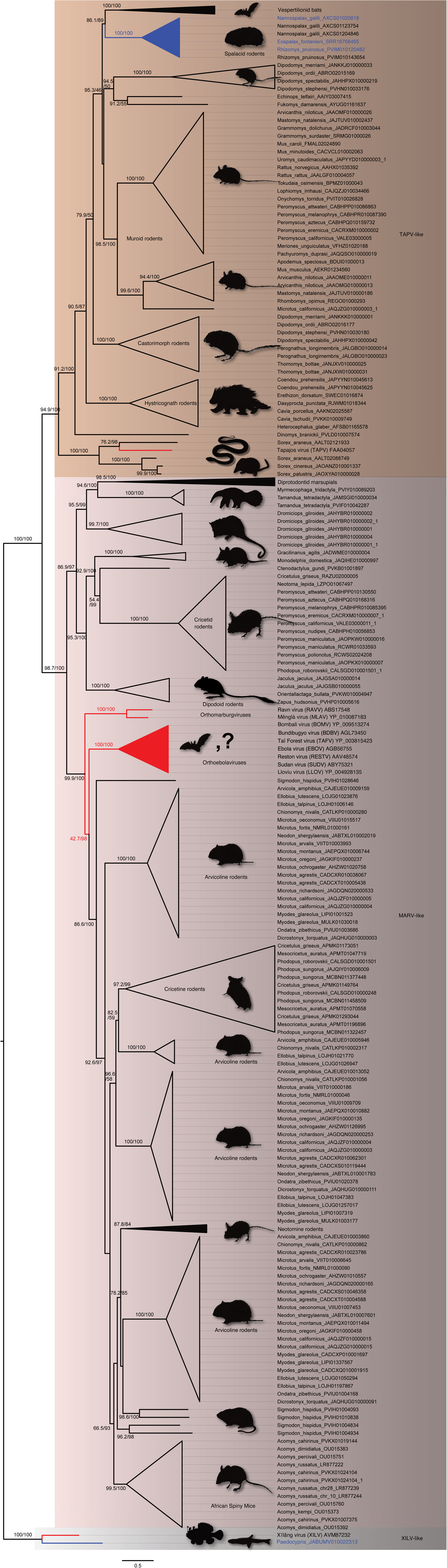
Maximum likelihood phylogram based on the amino acid sequences of the NP or nucleoprotein gene of filoviruses and the NP-like paleoviruses from vertebrates. Black shaded triangles are large clades that were collapsed to save space (see Fig S6 for details). Red lines indicate viral lineages, blue lines indicate vertebrate sequences with open reading frames and black lines indicate vertebrate paleoviral sequences that are pseudogenes. Numbers represent approximate likelihood ratio test values and bootstrap values. The scale bar is present. Tree is rooted by XILV and major clades are shown in shaded rectangles.

For the VP35 gene, the sequences from fish lack significant similarity with the VP35 of the rest of the known filoviruses. Thus, LBA involving fish-associated sequences with other filoviral sequences such as TAPV cannot be assessed. However, the TAPV VP35 does have significant similarity to mammal-associated viruses and groups with paleoviral ORF’s from vespertilionid bats (Fig S7). In the analysis without fish-associated viruses, there was no significant amino acid composition heterogeneity found. The VP35-like elements had an ORF in African spiny mice that groups more closely with EBOV than MARV does. Applying an alignment filter (clipKIT with dynamic determination of gaps) removed 8.79% of sites, increased many support values, but did not significantly alter the topology (Fig S8). Using a model that accounted for heterotachy (within site variation), altered the branch lengths but did not affect the major groupings or topological placements of paleoviruses (Fig S9). Using nt with a partitioned model had little change on the topology of the phylogeny, with the exception of the movement of the paleovirus from the bat, *Murina*, moving to a position within the paleoviruses from *Myotis* (Fig S10). When paleoviruses with disrupted reading frames are included, the topologies are similar to the ORF tree (Fig S11). As with the NP gene tree, *Acomys* VP35-like sequences are nested within cricetid rodents that are in turn more closely related to EBOV-like sequences than to MARV-like sequences. The *Acomys* sequences group with the cricetid ortholog found in the last intron of the TAX1BP1 gene. However, these ORFs in *Acomys* (CM057028.1 reference genome location: CHR3: 33310914 – 33309961) are not found in the TAX1BP1 last intron region (JAJTCZ010000003.1: CHR3:96194980), suggesting an independent insertion from the cricetids. As with the NP gene, VP35-like paleoviruses in the MARV-like clade are widespread in the genomes of diprotodont marsupials.

### Protein structure recapitulates phylogeny

Analyses of predicted protein structures revealed conservation of some NP protein structures from divergent filovirus sequences. For example, the predicted structure of NP from TAPV was more similar to that of *Nannospalax* (root mean squared deviations, RMSD= 0.916; Fig 6A), than to the predicted NP structure of XILV (RMSD = 1.587; Fig 6B). Indeed, the multidimensional scaling (MDS) analysis of pairwise distances from estimated protein structures largely recapitulated the major groupings found in the phylogenies (Fig 6C). For the NP gene, XILV grouped closely with the paleovirus from the fish *Paedocypris*, while TAPV from a snake again grouped with mammalian sequences from bats (*Myotis*) and spalacid rodents (e.g. *Nannospalax*). Predicted structures from *Acomys* and cricetid rodents grouped with structures of the MARV-like clade. The MDS plot for VP35 structures (Fig S12) was less resolved than the MDS from the NP structures. Still, in agreement with the other analyses, TAPV was most closely grouped with bat sequences, while *Acomys* grouped closely to MARV-like sequences. The predicted structure based on a paleoviral pseudogene from the hamster (*Phodopus sungorus*) was an outlier and did not group within the MARV-like clade based on sequences alone. We do note that when flexibility was permitted (FATCAT), the structure of the hamster pseudogene had significant similarity to the structure predicted from EBOV VP35(P=2.56e-13; RMSD=1.78 with 3 twists).

**Fig 6.**
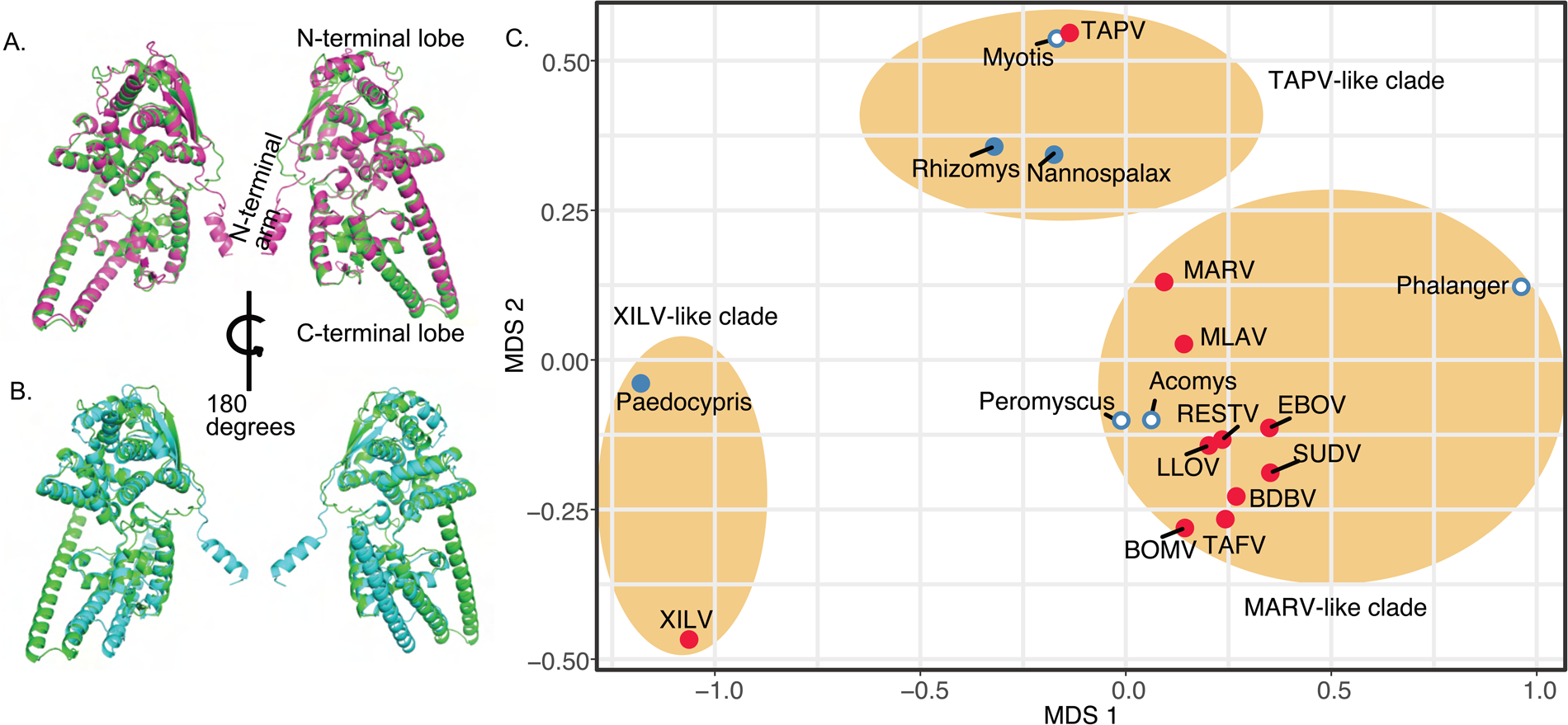
Protein Structure model divergence predicted by alphafold2 and aligned and visualized in PyMol 2.55 for the nucleoprotein gene of filoviruses and filovirus-like sequences in vertebrate genomes. A. Purple cartoon indicates the predicted the structure from the NP-like open reading frame sequence of *Nannospalax* (African mole rat; predicted template modeling score, pTM=0.83) and green cartoon represents the predicted structure of NP from TAPV (assembled from a lancehead snake, pTM=0.87) B. Blue cartoon indicates the predicted the structure from the NP-like open reading frame sequence of XILV (a fish-associated filovirus, pTM=0.77) and the green cartoon represents the predicted structure of NP from TAPV (assembled from a lancehead snake). C. Multidimensional Scaling plot of root mean squared deviations (RMSD) of predicted protein structure of the Nucleoprotein NP from filoviruses and filovirus-like sequences in vertebrates after alignment in PyMol. Ovals indicate major clades found in the phylogenetic analyses presented here. Red shaded stimuli are based on viral structures while blue stimuli are predicted from vertebrate genome sequences. Solid shading indicates extended open reading frames are present.

### Candidates for co-opted filovirus-like elements

Several new potential co-options were detected in the present study with filovirus-like elements containing open reading frames present in each major clade. For the XILV-like clade, the fish *Paedocypris*, contained open reading frames of an NP-like element. One near complete ORF was detected (JABUMV010022313.1) in the genome of a specimen of *Paedocypris* sp. from Singkep Island, Indonesia. As the contig was short (2197 bp), we could not assess the genomic context of the element. However, there are no further gene matches to XILV-like sequences on this contig or in the genome assembly, suggesting that this is ORF is from a genomic element rather than an RNA virus. RNA sequences from the sequence read archive (SRA) of two specimens of *Paedocypris* from Sarawak, Malaysia (SRX8475542-SRX8475543) had a total of 1771 matched reads to the NP element from Singkep (many reads were 100% identical). Thus, the NP-like element with the ORF has an RNA expression product in *Paedocypris*.

We also found paleoviruses with ORFs in the HUJV-clade from the Antarctic snaggletooth fish (*Borostomias antarcticus*). Fourteen NP-like elements were detected in the genome assembly (CATLJU000000000.1), with nine having extended open reading frames and a maximum length of 352 codons (Fig S13). There are two clades with one group having about 57-66% identity to sequences of the other clade.

In the TAPV-like clade, novel NP-like ORF elements were detected in spalacid rodents (mole rats, bamboo rats and zokors) and the previously described bat VP35-like orthologs with an ORF were present in assemblies from *Myotis* and *Murina* (bats). Along the branches from the common ancestor of *Murina* and *Myotis*, FEL detected 26 sites under significant pervasive purifying selection and 16 sites under diversifying positive selection. In comparison, FEL detected 39 sites under significant pervasive purifying selection and 11 sites under positive selection in an NP-like with disrupted ORFs along branches from the same common ancestor (*Murina*/*Myotis*). Both the pseudogenic elements (NP-like in bats) and the VP35 ORFs showed site-specific dN/dS distributions consistent with relaxed purifying selection and less diversifying selection. That is, a large peak of sites occurred where dN/dS < 0.5 (Figs S14A,B), with a gradual decrease in represented sites between 0.5 and 1 and beyond. Spalacid rodents also had ORF elements (NP-like). Here, FEL analysis returned 56 sites under significant purifying selection and 10 sites under positive selection. The distribution of site-specific dN/dS sites for these ORF’s was concentrated below a dN/dS of 0.5 (Fig S14C). While there are no known expression products for the bat VP35-like sequences, expression products were found for the paleoviruses with ORF’s in spalacids. The aligned ORF region for the three species (*Nannospalax galili*, *Rhizomys pruinosus*, and *Eospalax fontanierii*) was 1320 nt in length with seven apparent indels (each a multiple of three in length). The filovirus-like inserts were orthologous based on the nucleotide similarities of flanking regions of rodent contigs with elements: *N. galili* (AXCS01020818.1, 22694 nt) had 81% nt identity to a larger contig with 64% coverage from *R. pruinosus* (VZQC01004831.1). Likewise *N. galili* (AXCS01020818.1) had 83.4% nt identity with 73% coverage using a contig (SRA:SRR16768455.3388524.1) from *E. fontanierii*. 1000 parametric simulations of evolution (assuming no purifying selection to maintain codon structure, starting from an ancestral reconstructed sequence of the spalacid NP-like element, and using observed branch length parameters) yielded no cases of alignments that lacked stop codons (Fig 7). Indeed, the mean number of codons detected was 9.2 per alignment. The probability of observing no ORF disruptions for the spalacid paleoviruses under a scenario of pseudogenization is thus very low (P<0.001).

**Fig 7.**
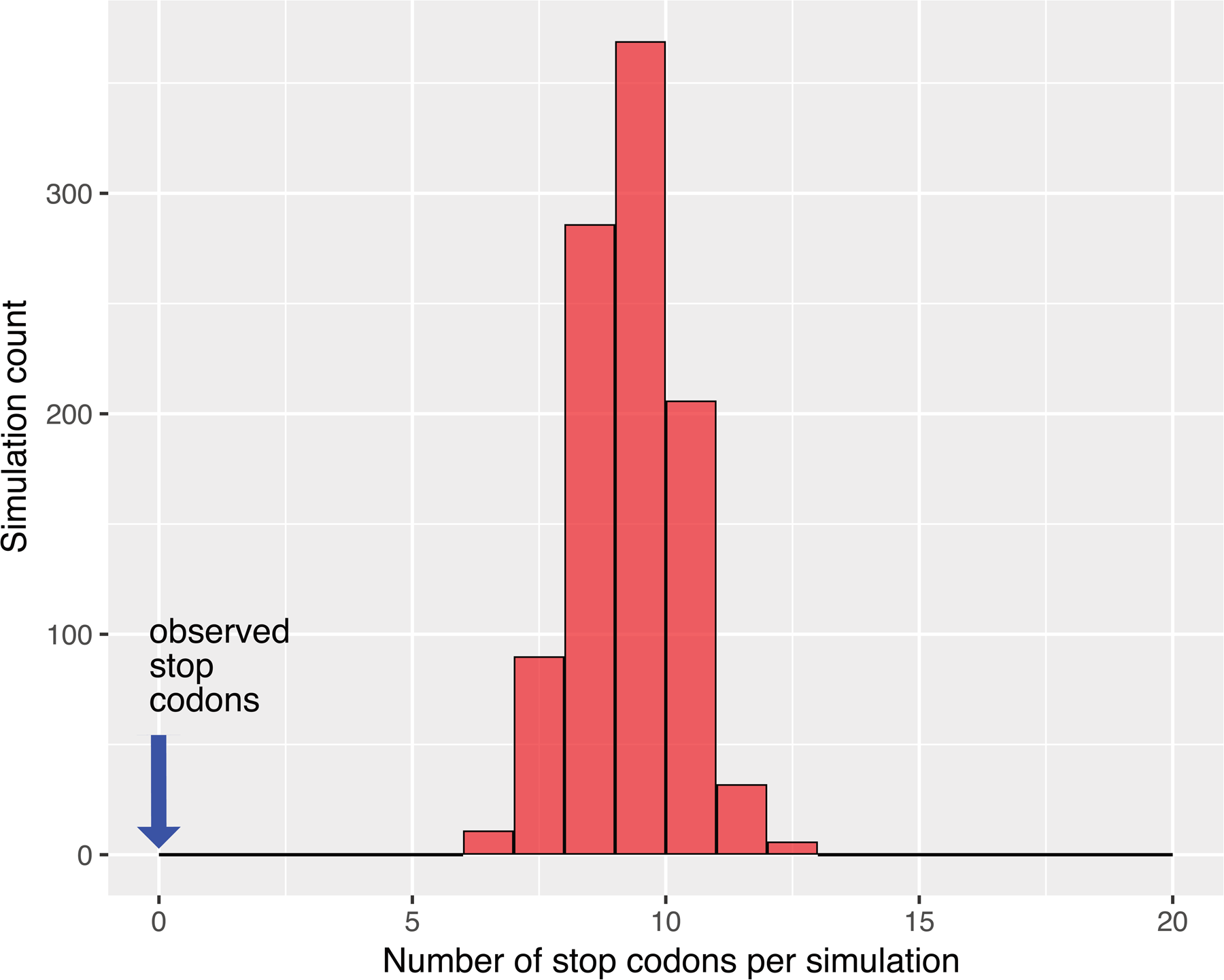
Histogram of stop codon counts from 1000 simulated alignments based on orthologous filovirus NP-like sequences of spalacid rodents. The parametric simulations used an ancestral reconstructed sequence as the starting sequence and the observed tree and substitution model values.

The spalacid filovirus-like elements also have RNA expression products. For example, searching a transcriptome project in bamboo rats (*R. pruinosus*: 270 G bases, 36 runs, SRP367919 [34]) that separated RNA into three tissue sources (liver, colon, duodenum) using BLASTn, found 60 positive read matches to the NP-like element with 42 (70%) of positive reads being from the liver (complete sequence coverage of the element was achieved). No matches were detected in the nine experiments with post-weaning 80-day old subjects (Fig S15). Another study with tissue specific RNA-seq experiments (*Eospalax fontanierii*: SRS3962618, nine experiments for each of four tissue types, 271 Gbases total) had 36 matches (maximum read depth of 11) to the NP-like element from *E*. *fontanierii* with liver experiments, 4 with brain tissue, 2 with heart and 0 with skeletal tissue. RNA-seq experiments with *Nannospalax galili* also yielded significant matches to its NP-like element. 338 matches (SRP331054, 8 tissue types) were found with top matches being skin (133) and lung (70). RNA extracted from a fourth spalacid taxon, *Tachyorcytes splendens* (SRR2141217), also has significant matches to the NP-like element from *R. pruinosus* (VZQC01004831.1), but with incomplete coverage.

The VP35-like element in the genomes of African spiny mice (*Acomys*) also had ORFs with expression products. An RNA-seq study of 6 tissue types (SRP350516) had the most matches to the VP35-like element with lung (42 reads) and brain (54 reads) tissue. Unlike with *Acomys*, the related VP35-like elements from cricetid rodents lacked an open reading frame and were present within the *TAX1BP1* gene region (Fig 8A). More specifically, these elements appeared at the same genomic location in the genomes of all examined hamsters, voles, lemmings, and musk rats (Fig 8B), suggesting long term presence in the Tax1bp1 gene (last intron).

**Fig 8.**
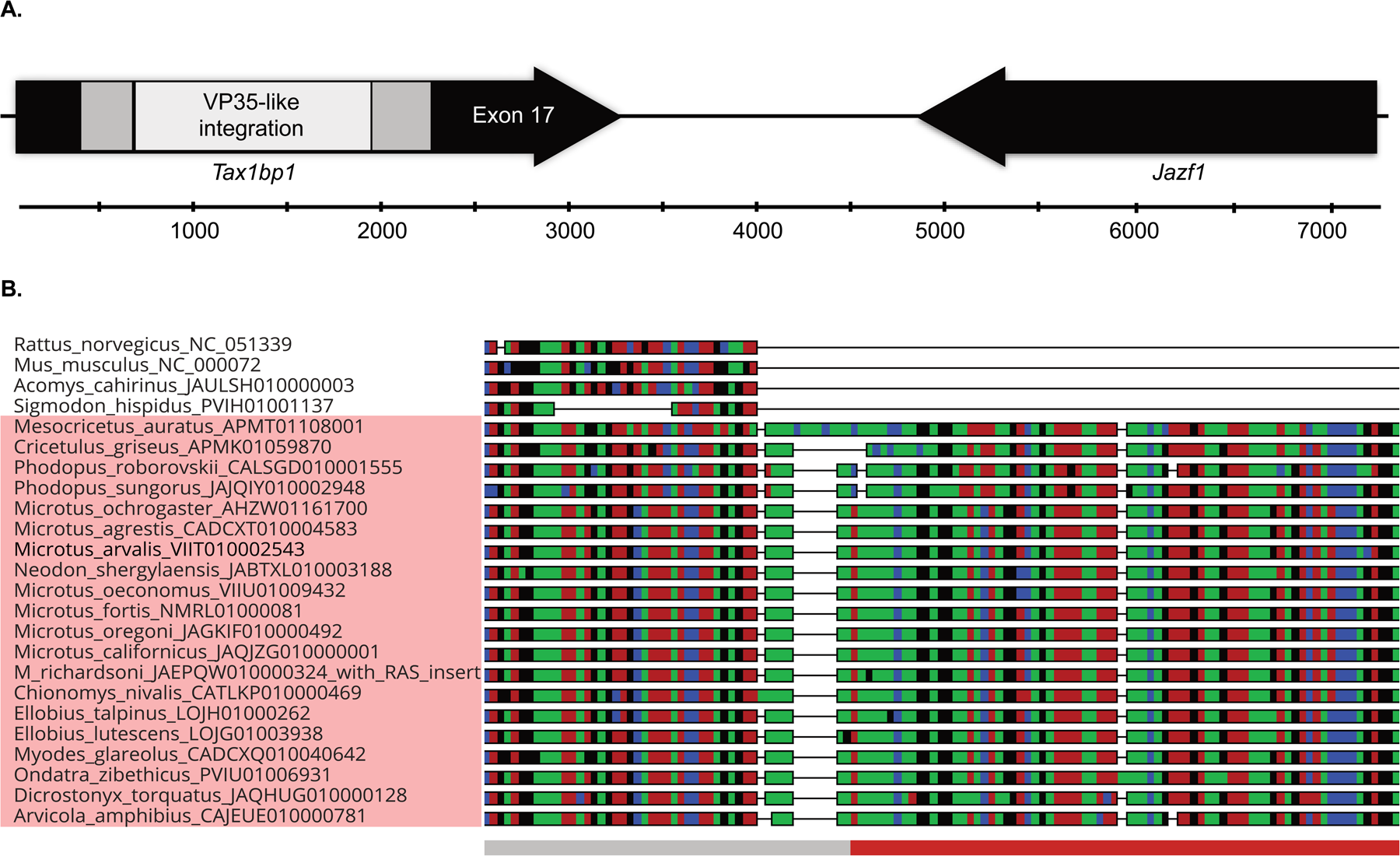
A. Cartoon of the 3’ end of the *TAX1BP1* gene showing an orthologous filovirus VP35-like element in the genomes of cricetine (hamsters) and arvicoline (voles, lemmings, muskrats) rodents. B. Alignment showing microsynteny of the 5’ end of the VP35-like element in hamsters, voles, lemmings, and muskrats (taxon names highlighted by pink rectangle). Colors in the alignment represent bases. Genomic regions of four muroid rodents that lack the insert are shown for comparison. Red bar below the alignment indicates region that has significant similarity to protein sequences of pathogenic filoviruses.

## Discussion

Our results improve our understanding of the deeper evolutionary relationships and host interactions of a group of RNA viruses with dangerous human pathogens. The finding of filovirus-like paleoviruses in fish genomes in each of the major lineages of viruses proposed to be associated with fishes (XILV and HUJV), is consistent with these viruses being piscine rather than the result of more recent jumps from terrestrial hosts. Indeed, the phylogenetic and structural associations between XILV (isolated from a marine frogfish) and the paleoviruses from *Paedocypris* sp. (a freshwater cyprinoform) suggest an ancient association between fish and the XILV-like lineage itself. The sequences from *Paedocypris* differ from other piscine filovirus-like sequences in being associated with fish from one of the early branching clades of teleosts (Otocephala-herrings, catfishes, cyprinids and others). The results also suggest that XILV-like viruses infect a broader taxonomic range of bony fish than previously thought. A lack of paleoviral orthologs from divergent fish genomes precludes timescale estimates for filovirus-like sequences of fish. However, the paleoviruses with open reading frames in both the XILV-like and HUJV-like clades for NP, suggest possible co-options of viral proteins by the host. Indeed, the existence of two divergent clades of elements in the same genome of the Antarctic snaggletooth fish (*Borostomias antarcticus*), both containing extended ORFs, is unique for filovirus-like elements.

Our results also suggest that the standard of using amino acid sequence from the RDRP (L-protein), for comparison of divergent viruses can be susceptible to long branch attraction artifacts. Our simulations reveal that XILV and TAPV are prone to LBA for the amino acid data. Using a codon-partitioned nt substitution model appeared to alleviate the LBA. We recommend that evolutionary analyses of divergent RNA viruses include codon-partitioned models. Such data uses the same alignment hypothesis as for amino acids but have three times as much characters and less susceptibility to LBA[32]. Adding divergent outgroups can reduce ingroup LBA (as occurred here with the L protein), but also introduce new sources of bias and LBA[35]. A site-specific model failed to affect the topology, suggesting that such models may not account for some LBA scenarios (such as lineage specific rate accelerations). Other lines of evidence support TAPV grouping with MARV-like sequences and not XILV. NP phylogenies (AA and partitioned nt) with outgroup taxa strongly supported a placement of TAPV with the MARV-like sequences. Surprisingly, TAPV (from a snake) groups *within* a clade of filovirus-like sequences from shrews (*Sorex* sp.) with strong support, suggesting at least one relatively recent host transfer involving snakes and shrews. Estimates of divergence based on protein structure (independent from potential biases of substitution models) recapitulate the phylogenetic associations of the NP-like sequences. Notably, TAPV structures are most similar to those predicted for paleoviruses of the same phylogenetic clade, bats and spalacid rodents. Taken together, our paleoviral evidence and simulations to address bias suggest that TAPV is more closely related to mammalian paleoviruses than to fish-associated viruses. If TAPV is a reptile-associated virus [31], then transfer to small mammals has occurred on several occasions. The nesting of TAPV within a clade of elements from shrews suggests a transfer from shrew to snake. Regardless of direction, the opportunity for prey-predator transfer of viruses between snakes and small mammals has been persistent in evolutionary time. Presently, shrews (Soricidae) overlap in geographic distribution with lancehead snakes in the northern Neotropical zone.

Our findings further establish the phylogenetic position of paleoviruses from rodents being nested inside the clade containing human filoviral pathogens. The pattern is present for both the NP and the VP35-like elements and is supported by predicted protein structures. Notably, the ortholog found in the last intron of the *TAX1BP1* locus has been present since the common ancestor of hamsters and voles (about 12 MYA). Our results expand knowledge of the presence of the ortholog from 3 species to 21 species (including the muskrat). While, this paleovirus is a pseudogene, and has suffered indels, it has likely remained present in the last intron of *TAX1BP1* since the Miocene. Rodents (hamsters, mice and guinea pigs) can be infected by filoviruses but have a natural immunity to the examined wild type strains. For hamsters, immunity to EBOV disease is imparted by an antibody response that is dependent on CD4+ T cells. One of the most important regulators of mammalian CD4+ T cell-mediated immunity to viruses is TAX1BP1. Our finding of maintenance of an EBOV-like element in hamsters and voles in the gene of a key player for hamster immunity to EBOV, suggests a co-option event that affects host immunity. Changes in intron length can affect gene expression mammals and can even coevolve in pathways where coordinated timing is relevant[36]. Understanding the details of rodent immune protection to human pathogens, such as EBOV, has direct implications for medicine and for studies that use these rodents as models for filovirus research.

While viruses of the clade that contains human pathogens are most often associated with bat hosts, isolation of infectious virus from bat hosts has occurred for only MARV and LLOV. So, there is still much unknown about natural host reservoirs for filoviruses. The presence of independently inserted EBOV-like elements suggests that rodents have had significant evolutionary interactions with EBOV-like viruses since the common ancestor of MARV and EBOV. Although we detected expression products of VP35-like elements in the transcriptomes of *Acomys*, it is unknown if proteins are produced or if the expression has phenotypic effects. The close association of *Acomys* sequences to sequences of LLOV/EBOV did not appear to be due to a bias such as lineage-specific rate differences between mammal and viral sequences. First an analysis that accounted for heterotachy did not change the phylogenetic position of the filovirus-like elements in *Acomys*. Also, the estimated protein structure distance analysis, which is unaffected by the fit of substitution models, showed a close relationship of the *Acomys* sequences with LLOV/EBOV-like sequences. Under a scenario of heterotachy, we might expect the mammalian sequences to be the outlier and basal to the most recent extant viral clades rather than nested within an extant viral clade (MARV/LLOV/EBOV) as seen here. Finally, the phylogenetic association of *Acomys* filovirus-like elements with LLOV/EBOV is found for both VP35-like genes and NP-like genes. The finding of independent paleoviruses related to EBOV in African and cricetid mouse-like rodents suggests that the clade of filoviruses containing human pathogens is more diverse (even in Africa) than presently known.

Our results show that protein structure has likely been highly conserved since fish origins (between XILV, TAPV, and MARV-like clades). Note that the structural predictions shown here from alphafold 2 have limitations as we didn’t include the disordered region for the NP sequences or consider RNA binding and interactions among viral proteins. As we used a conservative approach, we also didn’t consider structural flexibility for most comparisons. However, the close agreement of the major phylogroups with the structural divergences from viruses and paleoviruses supports the existence of evolutionary signal and overall structural conservation over geological time. Note that the crystal structure of a bat VP35-like paleovirus had strong identity to the structure of VP35 from EBOV[19], indicating the finding of general deep structural conservation of filovirus protein structures is consistent with structural biological evidence. As such, structural paleovirology has the potential to add important information for understanding ancient RNA virus evolution and detecting paleoviruses.

The NP-like ORF’s present in genomes of spalacid rodents are a unique case of a potential filoviral gene co-option with evidence of evolutionary maintenance of the ORF, amino acids, predicted protein structure and gene expression products. As the virus-like elements are flanked by significantly similar rodent sequences, we infer that the integration was present in the common ancestor of spalacids (about 28 MYA[37]). The parametric simulations and site tests for selection suggest that purifying selection is acting to maintain ORF’s and many amino acids in the spalacid NP-like orthologs. Our test of ORF maintenance was conservative as our simulations did not include indels -- a common ORF disruptor for pseudogenes. The bamboo rat and zokor (here a sister species) had a similar tissue pattern of expression with reads from the liver being numerically superior to other tissues. If the NP-like element had an anti-filoviral role in spalacid rodents, we might expect increased expression in the liver (a target of filoviral infection). While this pattern appeared to be present for bamboo rats, the number of individuals examined was low. Moreover, there are no male rats for the experiments at 80 days of age. Specific experiments with viral infection and spalacid rodents are required to further assess antiviral hypotheses.

Our results support the hypothesis of at least four ancient major clades of filoviruses -- each with extant viruses and vertebrate paleoviruses. In agreement with vertebrate evolution, the two fish-associated clades are basal to tetrapod-associated clades. For the NP-like tree, the grouping of a sequence from South American *Dromiciops gliroides* with Australian marsupials agrees with the marsupial species tree [38]. This further supports the hypothesis that a MARV-like filovirus lineage was associated with Neotropical marsupials before the Australian radiation [11, 21]. However, the monophyly of marsupial NP-like sequences is prevented by paleoviral sequences from several geographically divergent rodent groups (cricetids, gundis, jerboas) and anteaters. The presence of sequences from murid rodents (e.g. cricetids such as *Peromyscus* sp.) in several disparate positions of the NP-like tree, suggests multiple independent endogenizations and host transfer events.

The deeper evolutionary context of RNA viral pathogens is critical for understanding viral diversity, virus-host coevolution and the biology of spillover. Indeed, the grouping of TAPV within a previously known ancient clade of paleoviruses is consistent with the notion that the study of major paleoviral groupings can inform virus discovery. Importantly, our evolutionary analyses generated hypotheses about gene co-option and antiviral adaptations in hosts. The discovery of an ancient filoviral gene that appears to have been co-opted in spalacid rodents, showing differential RNA expression and signs of purifying selection, establishes an intriguing system for further study.

## Materials and methods

Filovirus and vertebrate genomic sequences were obtained from NCBI and EMBL-EBI. Filoviral taxonomy followed Biedenkopf et al. [39]. Local databases were made of genomic contigs with “significant” expect scores (E<1.0e-10) using EBOV protein sequences as queries for tBLASTn of the WGS and reference databases. Additional queries used protein sequences of Tapajós virus (TAPV), NP-like sequences (*Myotis myotis*, *Myodes*, and *Mesocricetulus*), and VP35-like sequences (*Acomys cahirensis*). Positive contigs were then searched using a tFASTy-based translation search, with the VT200 substitution matrix and the same query sequences. This approach helps to mitigate paleoviral fractioning due to frameshifts [14]. High-scoring segment pairs (HSP’s) with expect scores (E<1.0e-10) and greater than 200 amino acids were parsed from the output. We used ORF finder (https://www.ncbi.nlm.nih.gov/orffinder/) to verify the completeness of the ORF and obtain the nucleotide sequences (display ORF as nt option) when a contig had an HSP with an open reading frame. Outgroup sequences for the L protein were chosen based on sequences from three families with the highest expect values from Blastp comparisons with the L sequence of Xīlǎng virus (XILV, YP_010085045).

### Alignment

For translation alignment we used MAFFT 7 EINSI as a plugin in Aliview[40, 41]. Amino Acid alignments with >200 sequences were carried out by MAFFT using JT200 as a substitution model. Alignment trimming was carried out with either GBLOCKS[42] (least stringent parameters), ClipKIT 2.01 and the smart-gap option[43], or simply removing the disordered region of NP.

### Simulations

Parametric simulations to assess long branch attraction were carried out using the Ali-sim module in IQ-TREE[44]. Three conditions were simulated for the L-protein sequences of filoviruses. A Gblocks alignment was used to reduce indels because Ali-sim does not accommodate indel parameters. ML trees and substitution parameters were estimated for the observed topology of the L-protein gene where TAPV and XILV are sister taxa, TAPV is constrained to group with MARV-like sequences and XILV is basal (the observed NP tree). The second constraint was used in a third simulation but with the terminal branch lengths leading to TAPV and XILV shortened to 0.5. 100 simulations were carried out for each set of parameters. Four branch specific amino acid compositions (XILV, TAPV, HUJV-like clade and MARV-like clade) were included in the tree parameters and based on empirical values to simulate unequal amino acid compositions across the alignments. ML analyses of the simulated alignments for each condition were carried out in IQ-TREE using the “-S” function and a file containing boundaries of simulated alignments. Topologies were tallied and summarized in pie graphs.

Parametric simulations to test for evolutionary retention of the open reading frames were carried out using an ancestral sequence reconstruction of the NP-like elements in spalacid rodents (with a complete open reading frame) estimated using the IQ-TREE -asr option (the TAPV NP sequence was used as an outgroup). Simulations used a tree and nucleotide substitution parameters estimated for the paleoviruses in IQ-TREE. 1000 simulated alignments of 1320 nucleotides were estimated using Seq-gen 1.34[45] with the specific values: Seq-Gen-1.3.4/source/seq-gen -mGTR -r3.7022 11.3352 1.7544 3.7222 11.0280 1.0 - f0.2884,0.2339,0.2387,0.239 -l1320 -k4 -n1000 -op spalacid_np_only_for_seqgen_tree.phy. The resulting alignments in Phylip format (-op) were concatenated, translated in Aliview. We tallied stop codons and taxon-specific stop codons per simulated alignment. A histogram of the stop codons was created using ggplot2 in R.

### Phylogenetic methods

Phylogenies and substitution models were estimated using Maximum Likelihood and IQ-TREE 2.2.2.6[46]. Models partitioned by codon position used a partition file and the -p command. Trees were visualized in Figtree 1.44. Branch support was estimated by approximate likelihood ratios (aLRTs) and ultrafast bootstraps. Site-specific frequency models (PMSF using C60, the ML version of Bayesian CAT models) in IQ-TREE were used with a guide tree parameter to examine the role of site-specific frequency effects on topology. The effect of heterotachy (within-site variation) on tree topology was examined using the GHOST (General Heterogeneous evolution On a Single Topology) model implemented in IQ-TREE[47].

We used the Fixed Effects Likelihood (FEL) routine in HyPhy to assess site-specific patterns in selection[48]. Additional sequences from previous sequencing of bat VP35-like and NP-like elements [18, 19] were added to those assembled here. Test clades were selected, translation aligned in Aliview and calculations of dN\dS were made for partial NP sequences (several bat sequences were available for partial NP). NP and VP35 bat sequences were chosen to compare paleoviruses with and without an open reading frame over the same evolutionary time scale. Because FEL requires an open reading frame, we corrected stop codons and replaced disrupting indels with consensus nucleotides for paleoviruses that were pseudogenes. Kernal density estimate plots of dN\dS distributions for three clades of NP-like sequences were made in Datamonkey.

### Structural comparisons

Protein structures were predicted with Colabfold:AlphaFold2 using MMseqs2 and a PDB100 template mode[49]. Structure models with the highest per-residue estimate of confidence (pLDDT) were aligned in PyMol 2.55[50] and the average distance between atoms was calculated using pairwise root mean square deviation (RMSD). Divergent RMSD’s were tested for significant similarity using a flexible protein structure alignment algorithm in FATCAT[51]. Representative paleoviruses for major clades were used for structural analyses if they had open reading frames or, if pseudogenized, had the highest FASTA expect values compared to EBOV. A pairwise matrix of the RMSD’s was exposed to multidimensional scaling in R and the resulting values were plotted using ggplot[52] and compared with phylogenetic clade.

