## Supplemental Files for "Genomic transfers help to decipher the ancient evolution of filoviruses and interactions with vertebrate hosts"

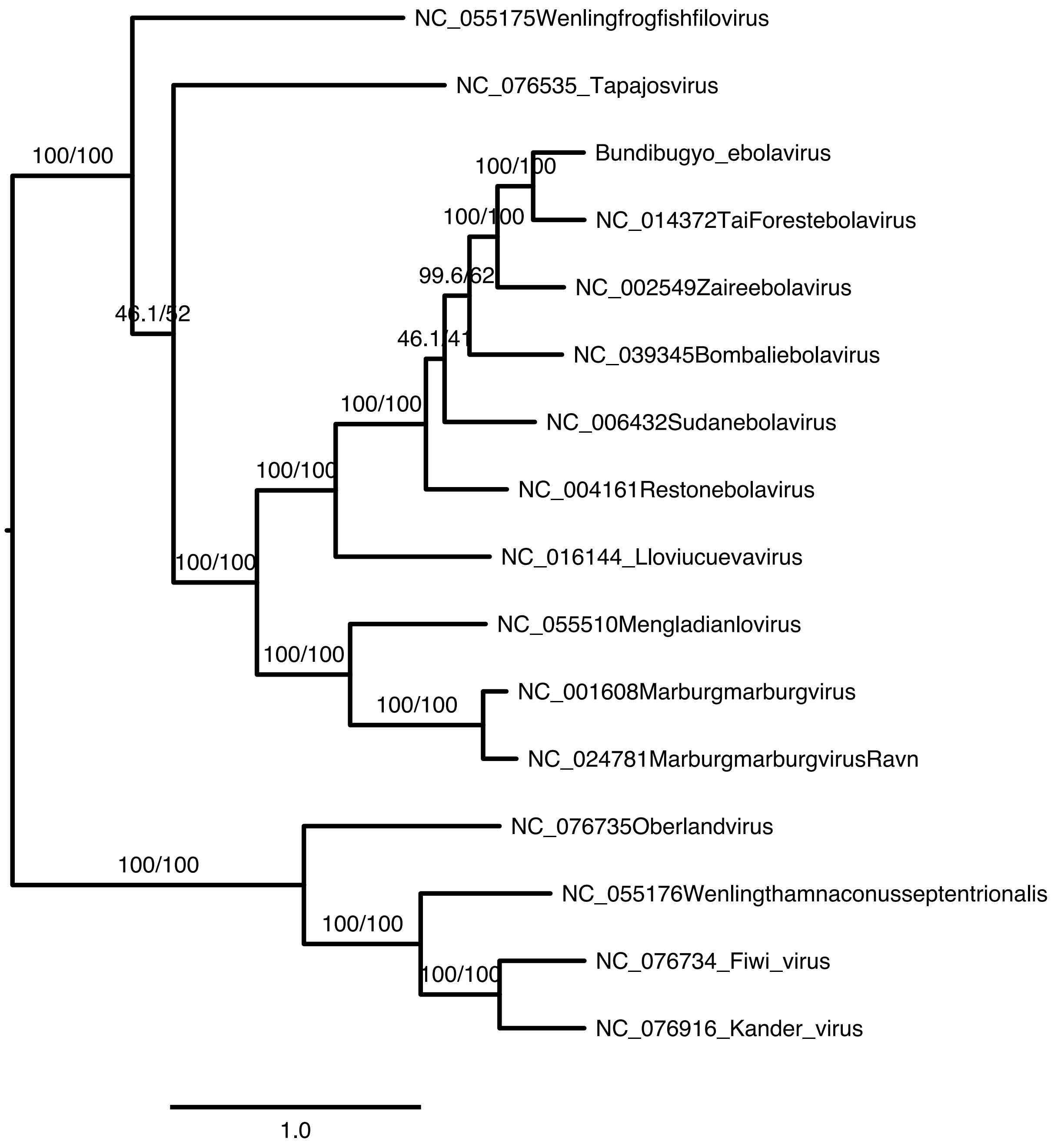


**S1 Fig. Maximum likelihood phylogram with GenBank Accession numbers based on nucleotide sequences of the L gene (RDRP) for filoviruses.** The substitution model was partitioned by three codon positions. Numbers represent approximate likelihood ratio test values and bootstrap values.

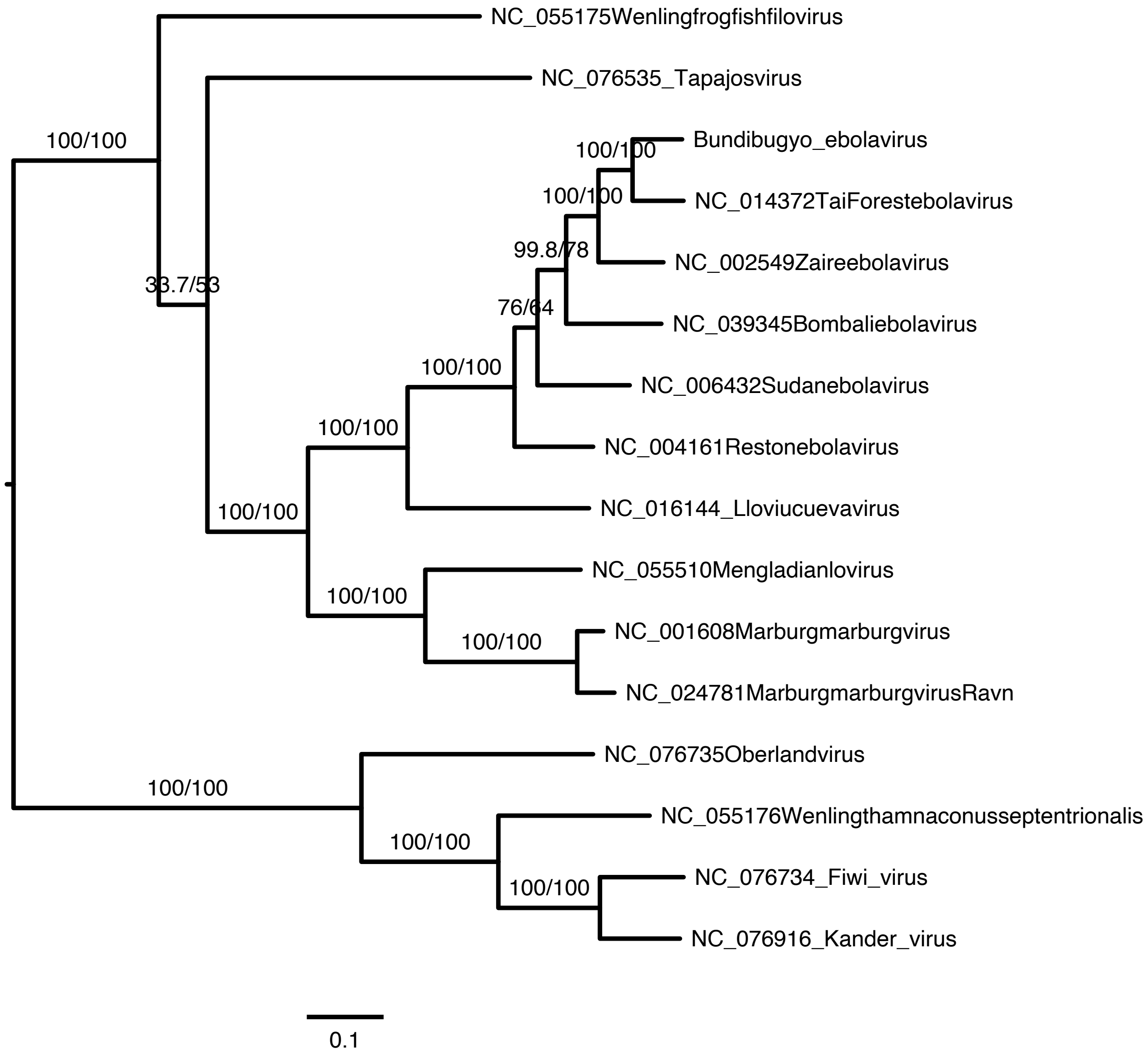


**S2 Fig. Maximum likelihood phylogram with GenBank Accession numbers based on partitioned nucleotide sequences of the L gene (RDRP) for filoviruses.** The substitution model was partitioned by the first two codon positions, while the third codon position was omitted. Numbers represent approximate likelihood ratio test values and bootstrap values.

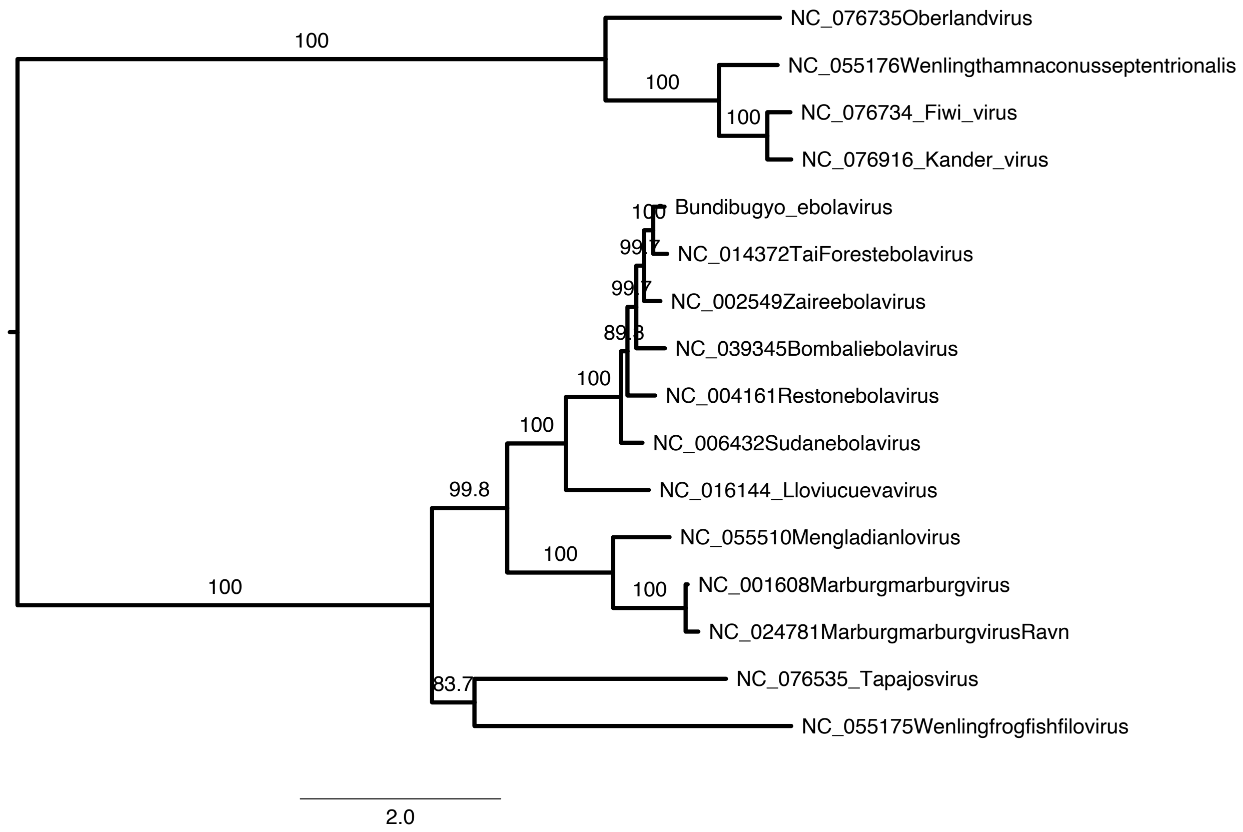


**S3 Fig. Maximum likelihood phylogram with GenBank Accession numbers based on Posterior Mean Site Frequency Profiles (PMSF) for amino acids of filoviruses.** Numbers represent approximate likelihood ratio test values and bootstrap values.

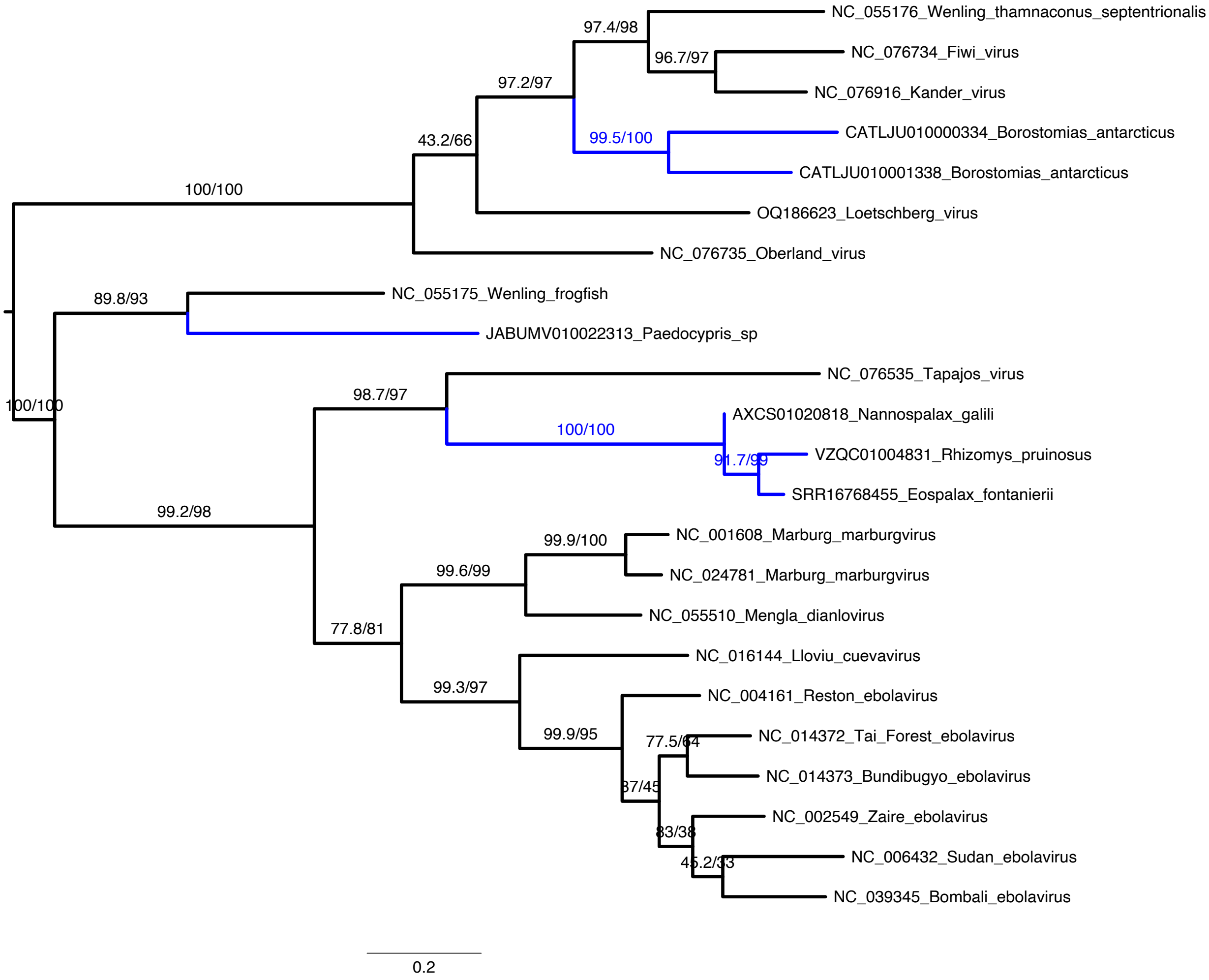


**S4 Fig. Maximum likelihood phylogram with GenBank Accession numbers based on nucleotide sequences of the nucleoprotein gene (NP) for filoviruses.** The substitution model was partitioned by the first two codon positions with third codon positions being omitted. Numbers represent approximate likelihood ratio test values and bootstrap values. Blue lines indicate branches leading to paleoviruses from vertebrate genomes with extended open reading frames.

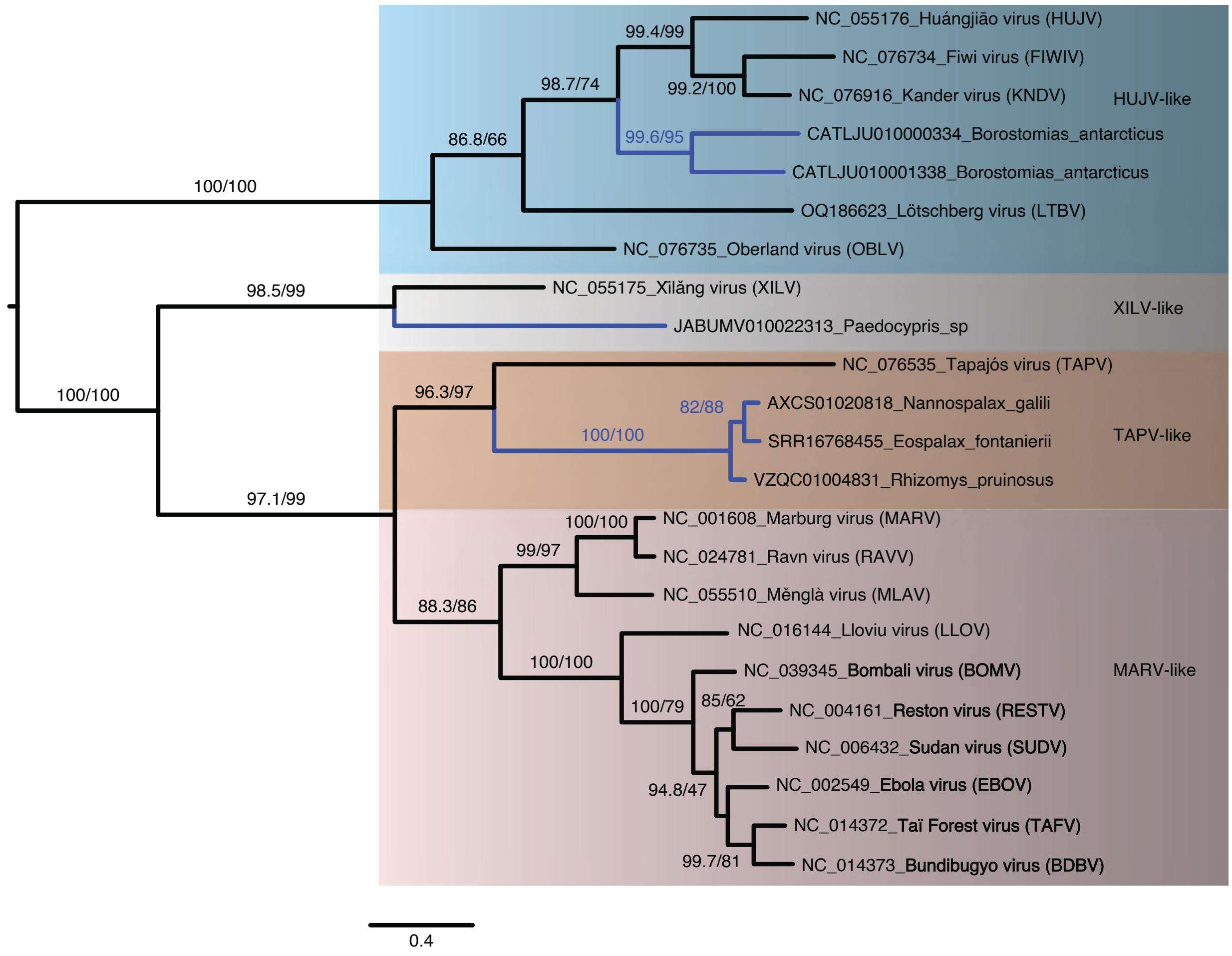


**S5 Fig. Maximum likelihood phylogram with GenBank Accession numbers based on nucleotide sequences (all codon positions) of the nucleoprotein gene (NP) for filoviruses.** The substitution model was partitioned by three codon positions. Numbers represent approximate likelihood ratio test values and bootstrap values. Blue lines indicate branches leading to paleoviruses from vertebrate genomes with extended open reading frames.

**S6 Fig. Maximum Likelihood phylogram based on the amino acid sequences of the nucleoprotein gene (NP) of filoviruses and the NP-like paleoviruses from vertebrates.** Red lines indicate viral lineages, blue lines indicate vertebrate sequences with open reading frames and black lines indicate vertebrate paleoviral sequences that have disrupted open reading frames. Numbers represent approximate likelihood ratio test values and bootstrap values. The tree is midpoint rooted and three major clades are shown in shaded rectangles.

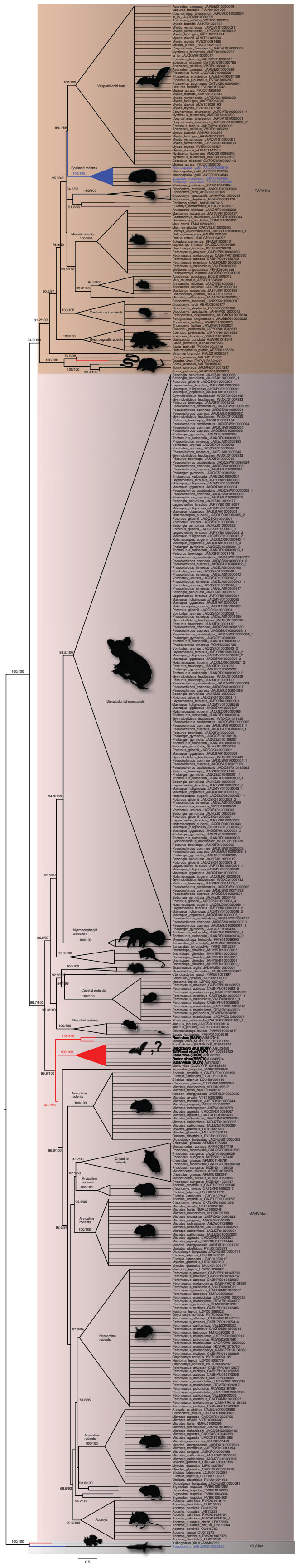

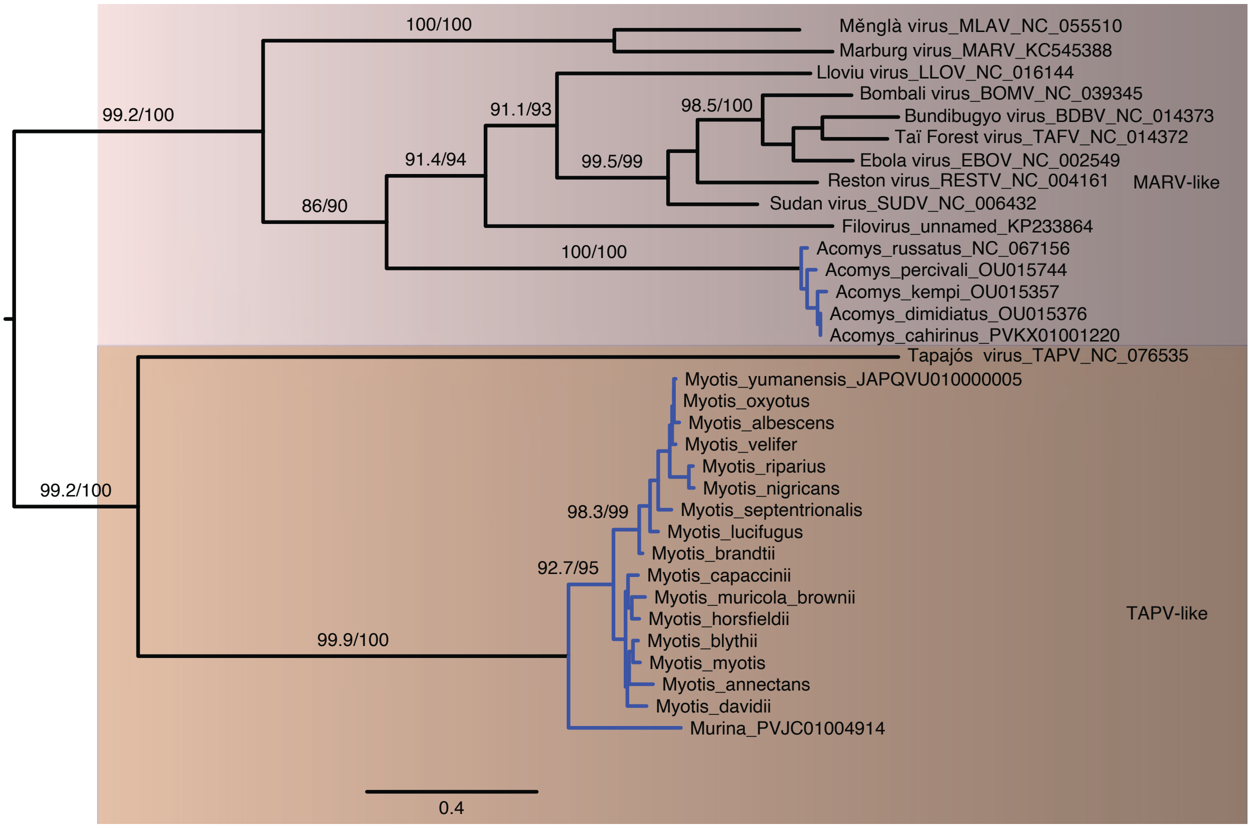


**S7 Fig. Maximum likelihood phylogram based on amino acid sequences of the VP35 gene for filoviruses and filovirus-like paleoviruses (from vertebrate genomes) with open reading frames (blue lines)**. Two major clades (MARV-like and TAPV-like) are identified and shaded. Numbers represent approximate likelihood ratio test values and bootstrap values.

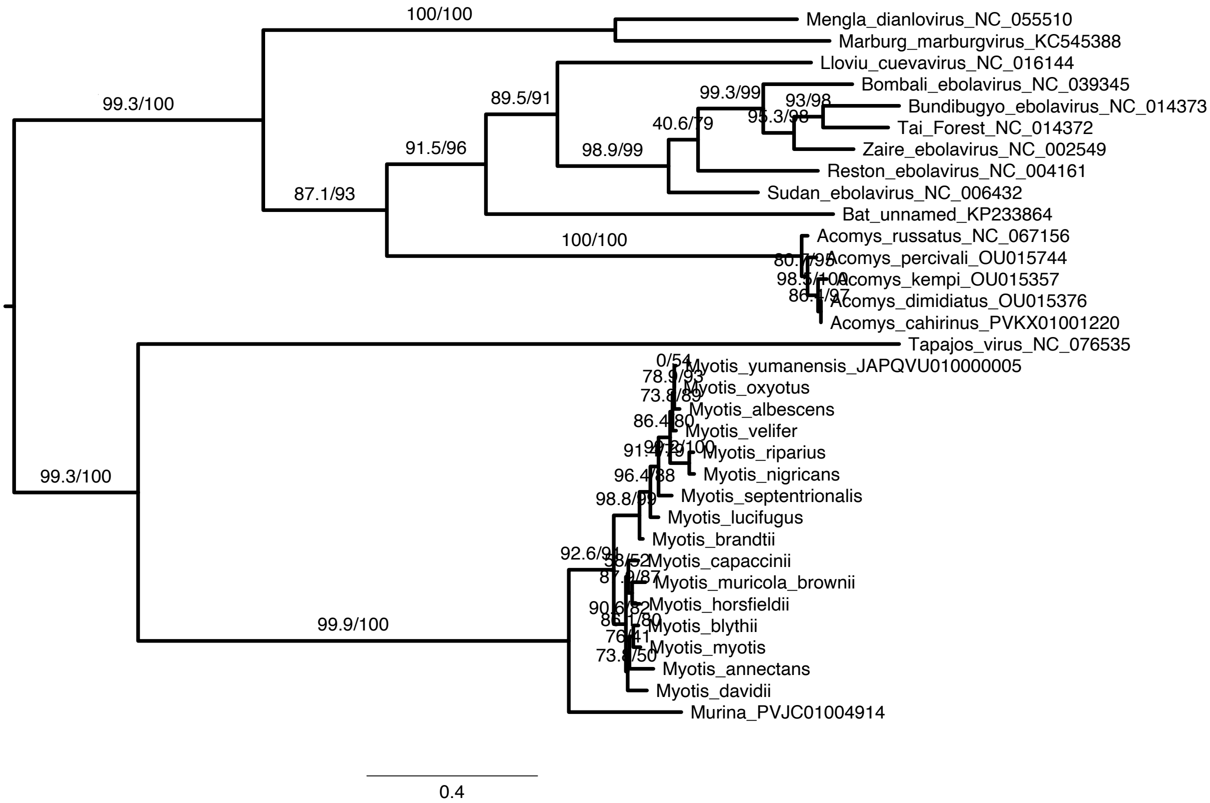


**S8 Fig. Maximum likelihood phylogram based on filtered amino acid sequences of the VP35 gene for filoviruses and filovirus-like paleoviruses (from vertebrate genomes) with open reading frames (blue lines).** The alignment was filtered using clipKIT. Two major clades (MARV-like and TAPV-like) are identified and shaded. Numbers represent approximate likelihood ratio test values and bootstrap values.

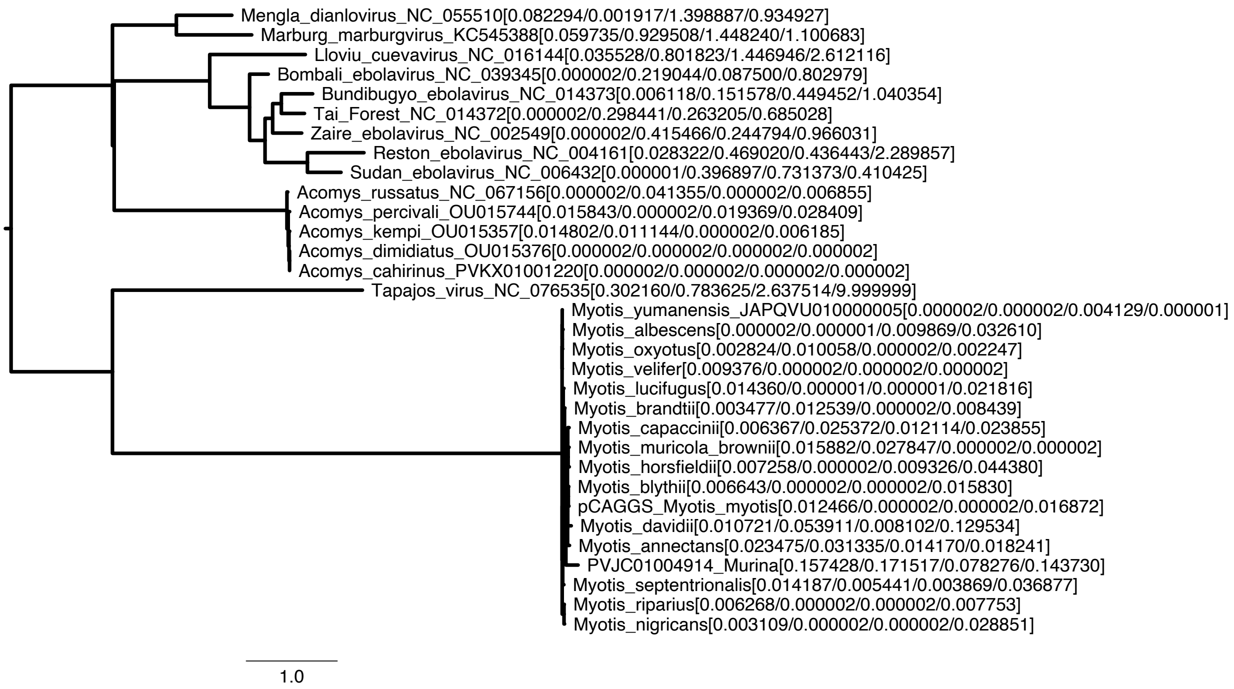


**S9 Fig. Maximum likelihood phylogram based on amino acid sequences of the VP35 gene for filoviruses and filovirus-like paleoviruses (from vertebrate genomes) with open reading frames.** The tree was midpoint rooted and based on a substitution model that specifically accounts for heterotachy (within site rate variation).

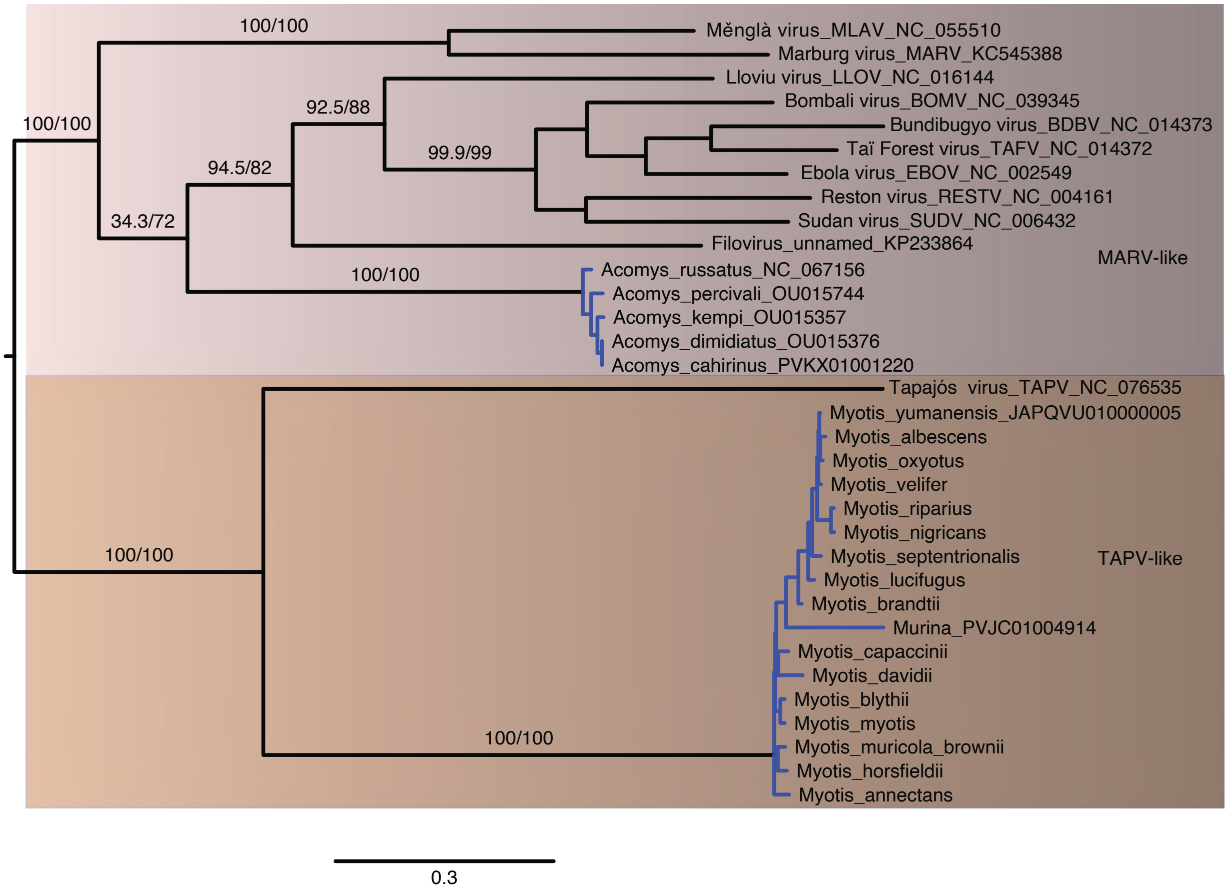


**S10 Fig. Maximum likelihood phylogram based on nucleotide sequences of the VP35 gene for filoviruses and filovirus-like paleoviruses (from vertebrate genomes) with open reading frames (blue lines).** The substitution model was partitioned by codon position. Two major clades are identified. Numbers represent approximate likelihood ratio test values and bootstrap values.

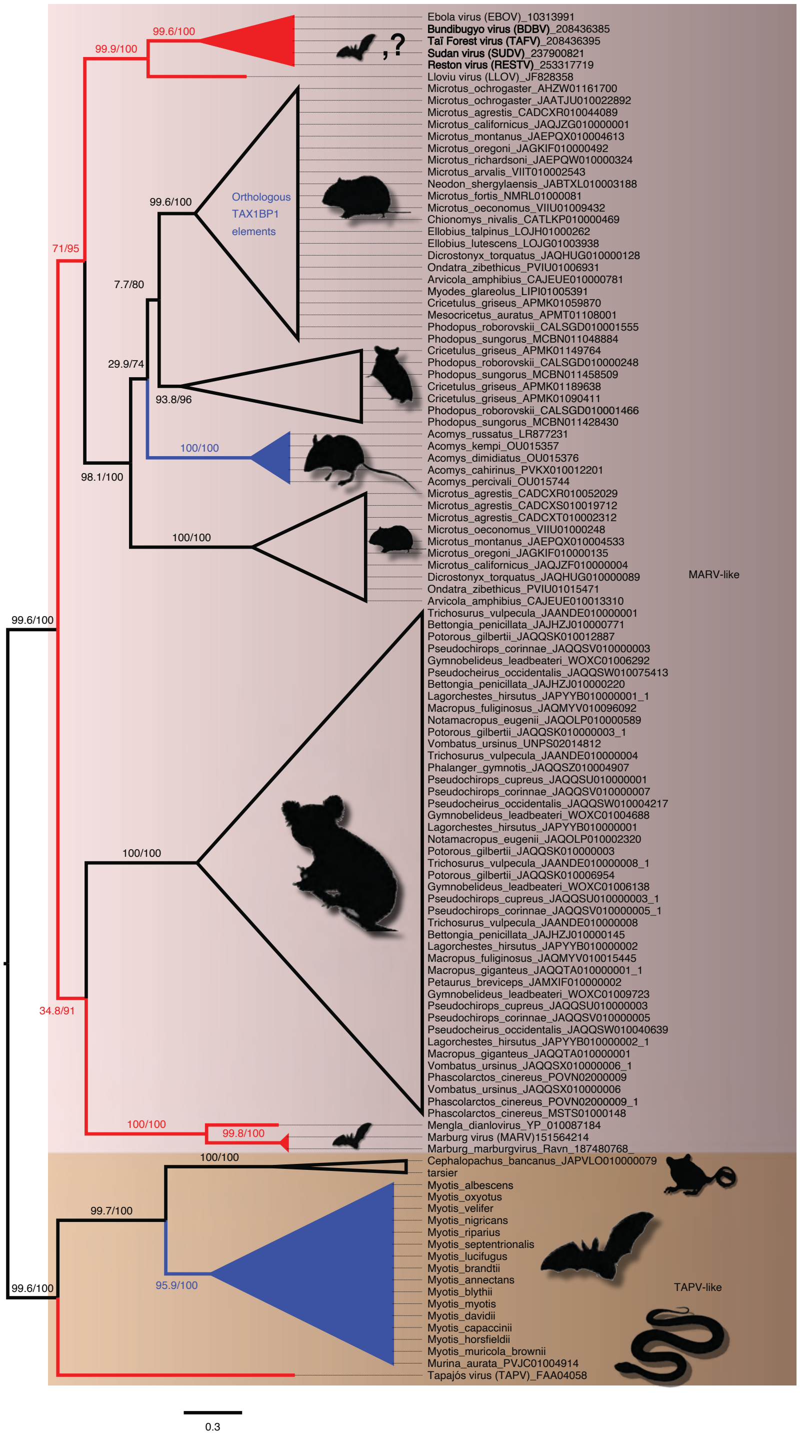


**S11 Fig. Maximum Likelihood phylogram based on the amino acid sequences of VP35 of filoviruses and the VP35-like paleoviruses from vertebrates.** Red lines indicate viral lineages, blue lines indicate vertebrate sequences with open reading frames and black lines indicate vertebrate paleoviral sequences that are pseudogenes. Numbers represent approximate likelihood ratio test values and bootstrap values. Tree is midpoint rooted and two major clades are shown in shaded rectangles.

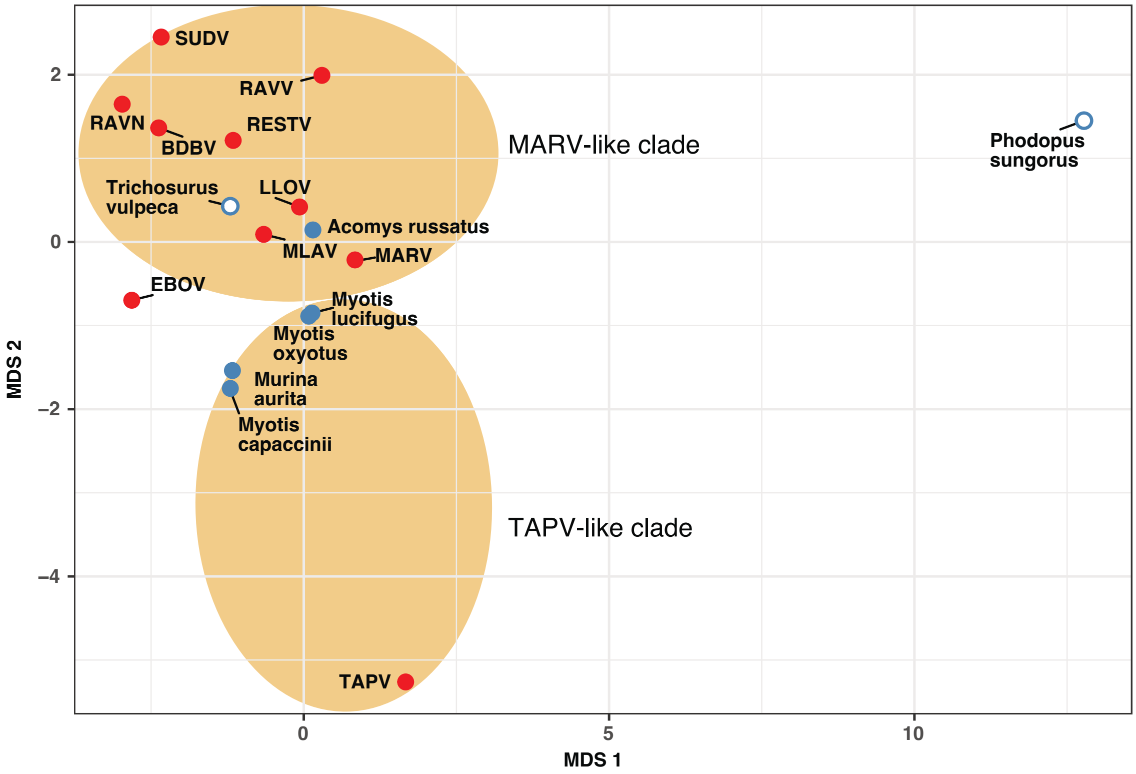


**S12 Fig. Multidimensional Scaling plot of root mean squared deviations (RMSD) of predicted protein structure of the VP35 from filoviruses and filovirus-like sequences in vertebrates after alignment in PyMol.** Ovals indicate major clades found in phylogenetic analyses. Red shaded stimuli are based on viral structures while blue stimuli are predicted from vertebrate genome sequences. Solid shading indicates open reading frames are present.

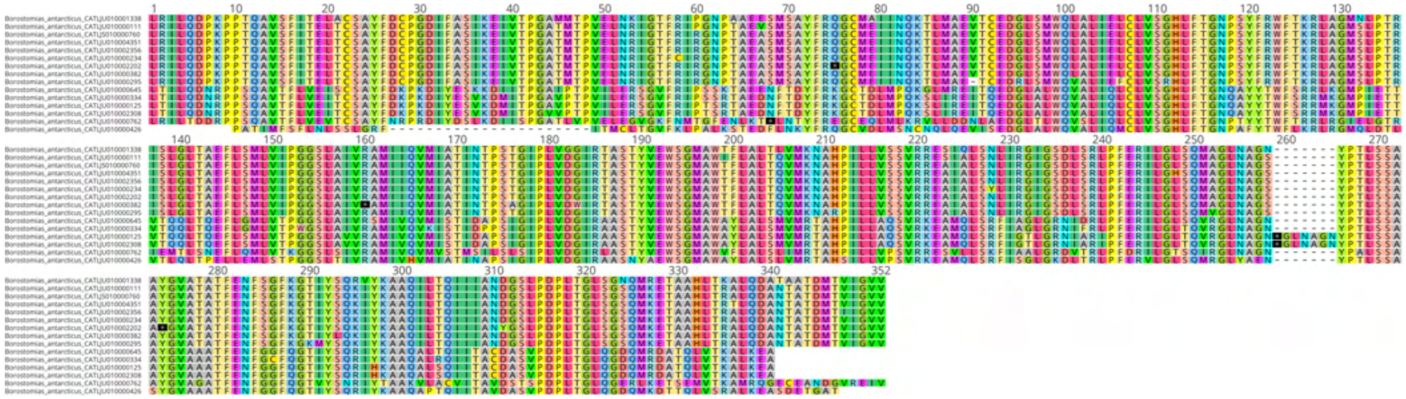

**S13 Fig. Multiple sequence alignment (amino acids) of NP-like elements from the genome of the fish, large-eye snaggletooth (*Borostomias antarcticus*).** Stop codons are depicted by an asterisk. Nine of the fifteen sequences are open reading frames.

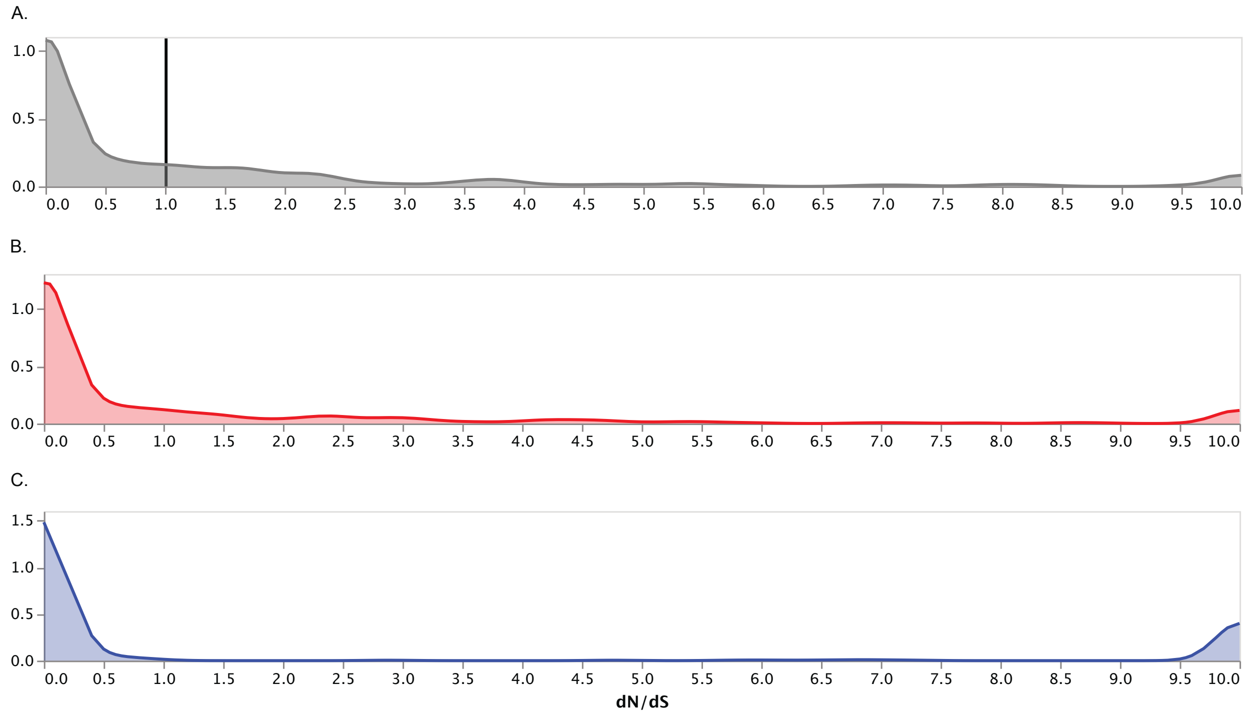


**S14 Fig. Kernal density estimates of dN/dS per site (calculated by the fixed effects likelihood method or FEL) in the alignments of filovirus-like elements of vertebrate genomes.** The black vertical bar indicates a neutral ratio. A) estimates from NP-like elements with reading frame disruptions of bats from *Murina* and *Myotis*; B) estimates from filovirus VP35-like elements (extended ORFs) in genomes of bats (*Murina* and *Myotis*); C) estimates from the filovirus NP-like elements in spalacid rodents with an open reading frame and expression products.

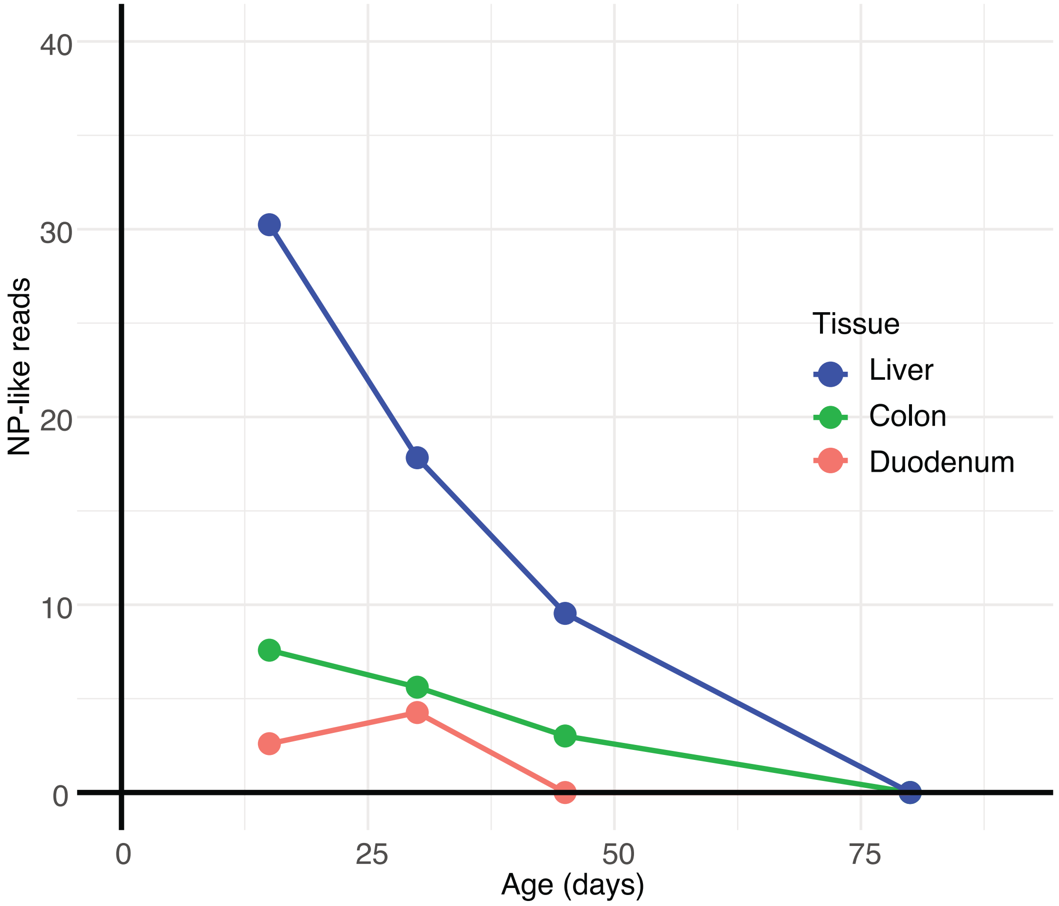


**S15 Fig. Plot showing total number of read matches (Y axis) to filovirus NP-like nucleotide sequences from RNA-seq experiments of the bamboo rat (*Rhizomys pruinosis*) from three tissue types and four ages (x-axis).** Read matches were standardized by gbases read per experiment.

Table S1. Base compositional heterogeneity test of the L protein amino acid sequences from Filoviruses.

| Taxon | Chi2 Composition test | p-value |
|---|---|---|
| 1 NC_076535_Tapajosvirus_TAPV | passed | 94.21% |
| 2 Bundibugyo_ebolavirus | passed | 66.35% |
| 3 NC_016144_Lloviucuevavirus | passed | 94.15% |
| 4 NC_014372TaiForestebolavirus | passed | 50.62% |
| 5 NC_004161Restonebolavirus | passed | 41.00% |
| 6 NC_039345Bombaliebolavirus | passed | 86.82% |
| 7 NC_002549Zaireebolavirus | passed | 69.51% |
| 8 NC_006432Sudanebolavirus | passed | 88.06% |
| 9 NC_055510Mengladianlovirus | failed | 0.00% |
| 10 NC_001608Marburgmarburgvirus | failed | 1.26% |
| 11 NC_024781MarburgmarburgvirusRavn | passed | 13.43% |
| 12 NC_055175Wenlingfrogfishfilovirus_XILV | failed | 0.37% |
| 13 NC_076735Oberlandvirus | failed | 0.12% |
| 14 NC_055176Wenlingthamnaconusseptentrionalis | failed | 0.01% |
| 15 NC_076734_Fiwi_virus | failed | 0.00% |
| 16 NC_076916_Kander_virus | failed | 0.00% |
| ** TOTAL | 7 sequences failed composition | chi2 test (p-value<5%; df=19) |

**Data S1. Pairwise root-mean-square deviations of atomic positions (RMSDs) of NP and NP-like sequences (without the terminal disordered region) from Filoviruses and vertebrate genomes.** The distance matrix was used for Fig S12.

| | EBOV | Myotis myotis | TAPV | Nannospax | Paedocypris | Peromyscus | MARV | MLAV | Acomys russatus | Phalanger | BOMV | Rhizomys | LLOV | XILV | BDBV | SUDV | TAFV | RESTV |
|---|---|---|---|---|---|---|---|---|---|---|---|---|---|---|---|---|---|---|
| EBOV | | | | | | | | | | | | | | | | | | |
| Myotis_myotis | 0.958 | | | | | | | | | | | | | | | | | |
| TAPV | 1.040 | 0.881 | | | | | | | | | | | | | | | | |
| Nannospalax | 0.857 | 0.642 | 0.916 | | | | | | | | | | | | | | | |
| Paedocypris | 1.427 | 1.168 | 1.236 | 1.217 | | | | | | | | | | | | | | |
| Peromyscus | 0.408 | 0.943 | 0.812 | 0.701 | 1.061 | | | | | | | | | | | | | |
| MARV | 0.444 | 0.549 | 0.836 | 0.787 | 1.109 | 0.549 | | | | | | | | | | | | |
| MLAV | 0.305 | 0.860 | 0.891 | 0.665 | 1.116 | 0.453 | 0.191 | | | | | | | | | | | |
| Acomys russatus | 0.446 | 1.029 | 0.969 | 0.692 | 1.094 | 0.453 | 0.585 | 0.517 | | | | | | | | | | |
| Phalanger | 0.785 | 1.258 | 1.341 | 1.136 | 2.269 | 0.844 | 0.791 | 0.797 | 0.924 | | | | | | | | | |
| BOMV | 0.045 | 0.945 | 1.069 | 0.843 | 1.230 | 0.408 | 0.440 | 0.314 | 0.462 | 0.784 | | | | | | | | |
| Rhizomys | 1.011 | 0.809 | 1.121 | 0.434 | 1.025 | 0.794 | 0.878 | 0.835 | 0.897 | 1.202 | 0.967 | | | | | | | |
| LLOV | 0.293 | 0.889 | 0.971 | 0.825 | 1.304 | 0.466 | 0.352 | 0.308 | 0.54 | 0.707 | 0.305 | 1.025 | | | | | | |
| XILV | 1.617 | 1.458 | 1.587 | 1.228 | 0.615 | 1.184 | 1.432 | 1.486 | 1.406 | 2.028 | 1.271 | 1.227 | 1.369 | | | | | |
| BDBV | 0.059 | 0.914 | 1.034 | 0.839 | 1.484 | 0.406 | 0.466 | 0.312 | 0.428 | 0.773 | 0.065 | 1.015 | 0.281 | 1.342 | | | | |
| SUDV | 0.056 | 0.972 | 1.105 | 0.854 | 1.515 | 0.405 | 0.456 | 0.307 | 0.458 | 0.793 | 0.056 | 1.009 | 0.301 | 1.529 | 0.059 | | | |
| TAFV | 0.063 | 0.926 | 1.030 | 0.833 | 1.488 | 0.402 | 0.463 | 0.300 | 0.448 | 0.788 | 0.065 | 1.022 | 0.280 | 1.263 | 0.03 | 0.058 | | |
| RESTV | 0.077 | 0.893 | 1.028 | 0.850 | 1.211 | 0.392 | 0.449 | 0.288 | 0.443 | 0.777 | 0.067 | 1.008 | 0.288 | 1.538 | 0.063 | 0.054 | 0.064 | |

**Data S2. Pairwise root-mean-square deviations of atomic positions (RMSDs) of VP35 and VP35-like sequences (without the terminal disordered region) from Filoviruses and vertebrate genomes.** The distance matrix was used for Fig 6C.

| | EBOV | Myotis luc. | TAPV | Myotis capp | RAVN | MARV | MLAV | Acomys russatus | Trichosurus | Phodopus | Myo_ox | LLOV | Murina | BDBV | SUDV | TAFV | RESTV |
|---|---|---|---|---|---|---|---|---|---|---|---|---|---|---|---|---|---|
| EBOV | | | | | | | | | | | | | | | | | |
| Myotis luc, | 1.685 | | | | | | | | | | | | | | | | |
| TAPV | 4.211 | 2.794 | | | | | | | | | | | | | | | |
| Myotis capp | 3.537 | 1.087 | 2.666 | | | | | | | | | | | | | | |
| RAVN | 1.454 | 2.453 | 7.316 | 2.402 | | | | | | | | | | | | | |
| MARV | 2.946 | 1.509 | 3.277 | 1.814 | 1.034 | | | | | | | | | | | | |
| MLAV | 1.648 | 1.869 | 4.306 | 3.050 | 1.372 | 2.407 | | | | | | | | | | | |
| Acomys russatus | 1.436 | 1.619 | 3.536 | 2.004 | 1.040 | 1.440 | 0.861 | | | | | | | | | | |
| Trichosurus | 1.740 | 2.782 | 5.455 | 3.010 | 1.937 | 1.978 | 1.658 | 1.564 | | | | | | | | | |
| Phodopus | 15.733 | 12.629 | 12.513 | 14.415 | 12.135 | 11.807 | 13.476 | 12.46 | 14.039 | | | | | | | | |
| Myotis ox | 2.372 | 1.446 | 2.978 | 2.116 | 1.769 | 1.485 | 4.979 | 4.692 | 2.936 | 12.825 | | | | | | | |
| LLOV | 0.517 | 3.580 | 3.766 | 3.329 | 1.676 | 1.934 | 1.723 | 1.096 | 1.145 | 12.694 | 4.290 | | | | | | |
| Murina | 3.045 | 1.111 | 2.854 | 1.240 | 1.716 | 1.937 | 2.210 | 1.888 | 3.016 | 14.307 | 1.612 | 2.556 | | | | | |
| BDBV | 0.735 | 1.776 | 7.328 | 3.419 | 1.888 | 2.639 | 1.607 | 1.060 | 1.573 | 15.129 | 3.344 | 0.333 | 3.236 | | | | |
| SUDV | 0.526 | 3.160 | 8.667 | 3.184 | 2.200 | 2.425 | 1.730 | 1.458 | 1.545 | 14.982 | 3.279 | 0.371 | 3.018 | 0.249 | | | |
| TAFV | 0.637 | 2.223 | 8.059 | 3.441 | 1.326 | 2.346 | 2.133 | 1.337 | 1.806 | 15.698 | 2.023 | 0.381 | 3.094 | 0.190 | 0.282 | | |
| RESTV | 0.329 | 2.006 | 6.593 | 3.267 | 2.790 | 3.334 | 2.131 | 1.184 | 1.406 | 13.836 | 1.770 | 0.360 | 3.197 | 0.353 | 0.317 | 0.274 | |

**Data S3. Multiple sequence alignment of L protein and L protein-like sequences from Filoviruses and filovirus-like elements in vertebrate genomes.** The alignment was trimmed using Clipkit 2. The data are used for the analysis in Fig. 3.

>UYL95578.1
MNLDPEEGLPRRTRHIANTHLDVPLRPGFLEEGAEYLGLRPTF-----------PRKREKSS-LDSLQPYII-----KNKPLITDYSYLFPILWREV---QNLQPTTSCQRLFMKVLSVAWGLLISQILSLASFASNLTPPE----------------------
LKHHGHLSRLYYLRYVLEGGRWSGELDHNPSAFYSED-----------------WKQTSEEVAIWNSQKRVHVYASRDFFLIA--TARDQFLITRDMFLCLSDTLSSRFLCLLAADLTDCIPSP--
SIPEYSLVYRTFDWGDDILARFGNEGYRLIGMWEPLCLGVMQKMTPDDLTDNFFWLEMSHDFLTALPGQDITRHL-----AQVSSLLEETFQQ--SPSHLAQLYGLYRLWGHPVVNSQKGISKLKVVACSPKAIDFNRAMQVACKAKEIFALRFRKTHR-
HWPKMDISELTEGSHLKKCINNWPVTIRHPDYSLLDWHYVRFEQSFEISEKVDLTAFISDKALSLTRPELKEAVESRCIGFA-----WKRSVLYQWLSTEFEGPKEFLERIGKNGFPEGENV-
VGVCPKERELKIEPRLFGLLTLSKRMYVVLTEALLAEHLLPYYEEITMMDDYLTLLKKIHRQTSDPEHTAN-------RKNLFTVLDFEKWNSNMRKEETEELFTMFDQLFGLTNVFTRSHEMFETSYLYLADGSYLPHFYSSSELR---
EDASCWTEHLGGIEGLRQKGWTIFTVALIRLIADQHDIEISLMGQGDNQVLKCSFP------
ATMTNEEIKMKHQKFLDHLFSEFGLVGPPLKKSETWTTSNLFIYGKFPVLKGVPLPMSLKRVCRMFRLSNEGFPTLESSLASIFTNVIAATLSDHHGAIPYFCGVLEAASALILGSSHSMGGQLSAVHEMSRAQKSVLEDLLTAMLIFPRSWGGFSIQMFGDLASKGFPDPATLGCS
LLKKLLPHVSGVRKMIKNMLSPRNPALSPLMLCKDPVSLNLFHPSSPSEVIRGLVSNMLYQAEKNAQIRSFLNMAFSKQD-QLADYLFQMRPLNPRVAHEVLEGTIYGRANKILTQLNKTNTMVRMA--KHTKGQLMTDRLRRAEANYFKSVLISLWTCGE---
DVWRESTCSSQWTQHLRDKGW-----KTKITGVTVASPFEAFTCHATQK-VDCLTAGHPNPDYIICRFSDR-------AHMVDARLAQDLGGIIPYLGGRTNEKVQSRGKTEMTSNPLLKIPLKLQTLVGWGVEEDSTFHRLLKRLVEAVSDWDPSVFIPEF-
SDISGSLEHRFADSSTTHGGTISILYTRPSYIYLSTDTLIKYCRGS---TNANLHFQALFSSIIYFQSLMLD--LANPLPSSLFHYHEECLQCISPIFEFFLQLPPPPSDLVPSDKENPFCWVPPKLRTARGSQYPSIDPNA---------LGEERLRRLFSYKAGGEAFRQI--
RLWAREIGDENPIARYQTSFALPWMYKCDPILLLESIAVHMTSMILRKRDLLRDTSSRRTLEAVLEGINAWPASAFSSLAPMMSRSSLVRHMVGPPYFVSPPVATPPSLKECCAAIKQAVVDILVQILLSWRTWNKEN-------------
AASLKIWSQALVVSMKSQDLGESPQDMMVQALGAIPSYEKSSFLQKFSHLTPI-------------------------------LREGVDLLSK-----------------------------------------------------------------------------------------------------------RLSEFSASQDLAMPILTTTELYETSVLALLSDH----
------------YLLETDSELLEDPVGPVYEPGDDWSLIYKPGGGATIGPYKILSLMMALRSLTFSVIASFGDGAGGFLSTLGRIYPNAKLFYNSLITPQQSLPQSLPDFVPVALLPYPDLVER-------LIRLKMTLELPSDVLLKDYPMSLRLNVDHSFDLITCDAEL-
GYENPRKRGLLLANQMLMIAH-LMNCSTLIFKTFSWDFELLDRQISILLQRFRLISVVRSKFSNQHNTELYLVCRSARLGAE-----------PRDYDQLALGRCILRLNSE---RSQLSQVSEYQSIWSSRFPTETSYLA----YRFSKESLLNKFVGYRRHFS----
TYHEVYFPETIMYHLLRTFNPV-----KGQQYKSRDRTRRNILTQGALKSSLLVFRNHRGNTQAGKILSYYPS--------VGAVLFERRERTLFPLDWGLPSEPKGVKTVFRSIWYPEHLPSLRALGAS
>YP_009337815.1
MCLDDAFSSGKGSKSYPTTHLQSAM---KCDLIKYAYGIPIGN-----------PGRFKHTD--QEIKRMFALYPLLQDYCTSLDYTKILYTLWKC--------QSSSVLHNINDALRYTHPYIGDHYSRL----------------------------------RVKNFKVARELWDKAITKSGRQRMEFTRNR----------
-----------------EERESLILPSKVNLLEIGHSANLCVLR--RKTDISIGTRDLLLMIGDLISERYLIFLSTEIAAEIGEE--QYPTSDAVKQIFDWGDSVMVEDGLAGFQLLKEFEPLVTSEMLLRTNDEFSDGAFRKSLLNELGKK-----------SVFKQQAEALI-
TILAHIPTMHHLAQIFGLYRLWGHPYVDSADGLLKIKHLGLSKKVIPNALKRELACSFKSHFCKQYRAKVG-KWPNLDTSRLPADSYLRAVISNRVLDQSNLAYKPSEWFQVKGLKTFEIDSSYNLATMLSDKALSLDRDELKYHCDRRNIGIA-----
RERRVITNWLRSTQISARELIEAVSEKG-LEKKELIIGVTPKEREMKTCPRMFSLMSLRMRMFVVAVGELLADHIVPYFPEITVGDSSIQLTRRLYDMTRSQRSATP-----
SQSVDVIMNIDFSKWNTNIRAELTNESFKFLDELFGLKQCVNYIPKMFTESLIYLADGSYLPDFDANLDVK---
KHPLAWTGHLGGLEGLNQKGWTAVTVVLIEMVARKHNIRFRLIGQGDNQVLRLTIPLVGDKFDPSGISAAKQTLKLFKASFVDEMEQAGLPVKISETWCSSKLFAYGKQHFFEGEPLAMSLKRLSRVFWLSNDLFPSLENAISTIHSNSLSACQSDLTTIVPFVVAVTETIYTILYHA
NESPGEQVTLTKSLFSLKHPVLWQLVSSIALVPRVLGGYPIQYLSSFLMRGFPDPVSESLSMLKRIIDLTDRQIKIALTSVANPSIYCSSQMIIQDPFSINALIPSSATGVLKSAVRTYLNSPGTNPFFLRFFQ-
ESESSVKPLCQALTRSTPLWPRLLHDVFDASLPGFAASVVGRIVKTTTVVMEAIKVD--GKRVEERMVYSDQNLFISIIYKLNEHRP---------TCSLEHAIDLRNNSWHR---FAPIEGVSTPHPIEYLKSVQLDY-GECRLCDLNQQPYLLVYVSPG------
KSDNPCSRMWNKLGPTTPYLGSSTSEKMRQELYTSEITDPLITRAARVLRAINWLVSESDELAQLMTRIFSCITDVAPECFFSVE-DSFSGSAEHRYQDYASFHGGLISTLYGLPSYVHMSTNQWNSYSKGA---TNHNIHFQAVLCIMQALVMFSKK--
TVSRYDSHVFHFHAVHKECIKVIQEPSITGLPGVNVILQSYPDSEYCFIKDRLIERRRDCFKSVK--------------CKKLETLPGAIMAQELTDLVLVALRETRITHIGV----
QRTAPMTWGKDADPVILIEETSLYIACSIYREINLKHVQSIQDLLSALIPLIGSLPDKAFYPLNFLFQFPESRLSLLASMYAIEGPHSFPMKDSSVAACCKRAVISVLSSRVIKQNEWLSNQIFSLLCRS------DKLSTLWESWEVLDKLICEATLSNESDFEVAIQSILTI---------------------
--------------------------------------------------------DLPLQKLSQSDRSTII------------------------LRTL---------------------------------------TFAEVNMSGDKLRPQAQVMDT-KPSLSFIQTSSVYDRLSVICPPVTWNLSASSVTPLVDMESPPR----
RSDFRSHFFRPVPLITTAHYKYLDLISEYVYNPGDMVIICGDGTGGVTTLIHRLNPNIRLYFNSLVDLEDTVAHSLTSFYPPATDYILHTKS--------IYKMRQSIEGISDITDPRFVNQLQSISGKTVRMIVCDAEGAGWTSPEKGIAIVGNLTKAAQR-
IGRDVVLVIKSYATDYRCVSVQSSILSSVFEVVKAARSAFSSYHNTEIFFIAERPTNAQQYIPILCDNSIIMPSLYQEQMKDIIKQLLNYMPPTLCDRVTTLLNSTSLSKQAMAQLEQAY----F-----------------------TQLYWGPITSRLRSRLSAIKTK-----------------
LSVMAGYTTARDVVLHIFSLLLRFKSTFADGYVMFVEECSNLTVETPFSRRGSQAYFVKISKAFSKPQTKRLFSRLGQFRNFPISQMLKDK
>YP_009091823.1
MAREGSEGRPTDERTFLARTLNSPIRALFYTIFL---GIRAVW---------------------DGFKRLLPVRT--EKGYARFSECVTYG----------MIGCDECVIDPVRELTEMQLPIKGKGSTRLRAMITE-----
RTAVPQIRVRSKILAERLGRAIRETLPAMIHHEWAFVMGKILTFMADNVGMNA--------------------------DTVEGVLSLSEVTRVSAVFNPDGLTIR--VENTGYIMTRETACMIGDIHAQFAIQYLAAYLDEVIGTR--
TSLSPAELTSLKLWGLNVLKLLGRNGYEVIACMEPIGYAVLMMRSPDPYVNDTYLNSILSEFPVD-----------SDARACVEALL-TIYMSFGTPHKVSDAFGLFRMLGHPMVDGADGIEKMRRLSKKVKIPDQSTAIDLGAIMAELFVRSFVKKHK-
RWPNCSINLPPRHPFHHARLCGYVPAETHPLNNTASWAAVEFNQEFEPPRQYNLADIIDDKSCSPNKHELYGAWMKKTAGWQ-----EQKKLILRWFTETMVKPSELLEEIDAHGFREEDKL-
IGLTPKERELKLTPRMFSLMTFKFRTYQVLTESMVADEILPHFPQITMTMSNHELTKRLISRTRPQSGGGR-------DVHITVNIDFQKWNTNMRHGLVKHVFERLDNLFGFTNLIRRTHEYFQEAKYYLAEDGTNLSFDRNGELI---
DGPYVYTGSYGGNEGLRQKPWTIVTVCGIYKVARDLKIKHQITGQGDNQVVTLIFPRELPSDPVERSKYCRDKSSQFLTRLSQYFAEVGLPVKTEETWMSSRLYAYGKRMFLEGVPLKMFLKKIGRAFALSNEFVPSLEEDLARVWSATSAAVELDLTPYVGYVLGCCLSAQAIRN
HLIYSPGTKRGAMPGL----RPTFESLVKSICWKPKAIGGWPVLMLEDLIIKGFPDPATSALAQLKSMVPYTSIDREIILSCLNLPSSVVSPSMLLKDPAAINTITTPSAGDILQEVARDYVTDYPQNPQLRAVVK-
NVKTELDTLASDLFKCEPFFPPLMSDIFSASLPAYQDRIVRKCSTTSTIRRKAAERG--SDSLLNRMKRNEINKMMLHLWATWGRSPARLDTRCLTTCTKQLAQQYRNQSW-----GKQIHGVSVGHPLELFGRITPSH-----RCLHEEDHFLQTFASEH-------
VNQVDTDITTTLGPFYPYIGSETRERAVKVRKGVYVVEPLLKPAVRLLRAINWFIPEESDASHLLSNLLASVTDINPQDHYSST-EVGGGNAVHRYSCRLSDKLSRVNNLYQLHTYLSVTTERLTKYSRGS---
KNTDAHFQSMMIYAQSRIDLILESLHTGEMVPLECHHHIECNHCIEDIPDEPITGDPAWEVKFPSSPQEPFLYIDLPVKDKLEPV--------PRMNIVRLALGPEAISELAHYFVAFRVIRAS------------
ETDVDPNDVLSWTWLSRIDPDKLVEYIVHVFASLEWHVLMSGVSVSVRDAFFKVSKRISETPLSSFYYLANLFVDPQTREALMSSKYGFSPPAETVPNANAAAAEIRRCCANSAPSI---LESALHSREVVWMPGTNN----YGDVVIWSHYIRLRFSEVKLV---------------------------
-------------------------DITRYQ----QWWRQSERDPY----------------------DLVPDMVLESDLDTLM-----------------------MRKA---------------------------------------------------RRPPLQVIRE--LDVAVINADHPAHVL-------------------------------
QNKYRKLIFREPKIITGAVYKYLSLKSE--LTEFTSAMVIGDGTGGITAAM--MADGIDVWYQTLVNYDHVTQQGLSVQAPAALDLLRGAPSGR------
LLNPGRFASFGSDLTDPRFTAYFDQYPPFKVDTLWSDAEGDFWDKPSKLNQYFENIIALRHRFVKTNGQLVVKVYLTQDTAT--TIEAFRKKLSPCAIIVSLFSTEGSTECFVLSNLIAPDTP----------------
VDLEMVENIPKLTSLVPQRTTVKCYSRRVACISKRWGLFRSPSILEVQF-------LHYITKVISDKGT---QLSLMAVADTMINSYKKAISPRRRSLHAYQHFENPAKLMLDMLTNNISDLKFMKTIMSTLGSNPSKLMTVKRT-----
QELRYNNLIYTKDAILFPNEAAKNTAPLRANMVYPVRGDLFAPPIMTLVSDE
>WAK72371.1
MNLEE--------PHLPDPHLKSPILPCTRELFVGYVDKAKPT-----------PGYIKAQV--RKLEDVVK-----NYPSLHTDTEHLYG----------AALSESCCPSMQQTLPRSYLHFLGETISDVRDLVQGLTTSE----PGHGLTFDQLLRPLNGYTELIVSAHIARTKWNKLRRAMARFRIHP------
------------------------LPRPGSLVYTICGRWKFYYTSSVVLII--IDQESYVMDYDQVLLVADTLTSRYLALLYADLCNQGYPL--QCPDPLLLSRCYEWGDELLEQYGNCAYRLIKMWEPLILGLLLQTTDPLEYSHRFYATMEDAYHQEYENVTRDRCLGACSWCQVRNLI-----
RQEPWPNISELHGIYRHWGHPTVDLKGGTLKVQSIAKEKKTLSTYTRLAIGAQFTRYLILSYIRRHG-
KWPKCRIDSSHEPKALYQAWHRVDMSDYEDNIAILDWNIIEFEKNFEFDYFEDWSQLLEDTAISYDRDHWDCTYDRTLLGYNPPPSPTLKRGILQALSTNVINIESICRRIRQRDVPHKWKI-
ITVHAKEREMKVEPRLFAMMVLEMRLYYCALEKNLAEKILEYFPQQTMTLGEIDLTQRLLTLTSRPPPGSL-------
FIPAYVCVDFEKWNLQWRLEAILMIARRCDELFGTPGLVTYAHEFFSKVMLIVGHRDIIPPGITSTNRQHPPESEYIWYDHLGGLEGICQKLWTLGTICMLLELSRRTGHRCIITGQGDNQVIKVFLPMSDSQYVSKYNAELTNQIKHVITHLTIIATELGLPIKASETWVSSKVFIYSK
DIIVDGAYAPTALKRVSRAYFDVAELFPTLESKLATIHTAGQTAAQKGDHPVISYLISLLESRIALYASFNEPTGTRLGKNLQIRT--
RQYLLEWIHFVSLLPRSVGGYPVASFLSYLYRGHPDPLTGDLNSLRLLGIKNDIFRRALAYILAELDDDPDLTMLLQDPVAVNWKLPVLTSNVVKNTLQNSVKDYAKNIDVKALFHAHWESEDQKIVETLIKADPFAVRVLNDIYMKSPAGARVGFLAMFNNMRTTQATI--
ASTQQRTMIAGIQRSE-ARWVQHLDRVYQ---AILKPLETRVCLSEEAQSLRQLSW-----RKEIVGVTVPHPVEQFLIGIKEYGEPC-----SSGYALIRYSPPD----------
DGSVPWYHRGPHAAYLGSRTQEKRVGQVIPVERSTPALAAARRLTAISSWAVSSDPVSQSTLTSLIRCRTNMPQEVLLASSGYVYSGSITHRYHDPMTSHDVLINSRPNLASHVYLSSDTLGEYARGV---RDLTMHFQGMFLLMMAYLHLYLLG-
ELNASGDVCLHAHIRCPTCIQETNDTPVSLPRA-EINYHQYEACHLIHTNALTSM-----LPDVWPTSVLLEQVIDK--DVSLTTLGSAGYVLHLDRRK-
SLATAYRGRQELAVSPGTRGLDILEFLSLGIRTIIKGLAAYTILVNAETYLTRVGKSHLNRANAVQAIVFGNPHSFWGSIRSHILLPEARVDLLDDLPTLAPSSETLKGGHQIDLTLCRLLSNEILSRLHDLEESEGNPFLDLLFFQTDEIDYLQLHRLWANSWLL--WLSTRV-
TTHTELLRARRDLPRR----------------SRDGKQLAKDLVLWY--------------------IRHIKP----------------------------VRQSQVTTIPTI---------KFTKTPPEVV---LRRC-----------------------------------------------RDMYHPSAHVESRFLNDLRSIR-
EDPGQALTCVCPLEITSIPLPIPDSSNRLDSPPTQ---VHDKGDHYFRLHGVYSTAAYKYAEIVIQEHIAAADVVVCLAEGSGGVMRYLVNKYSPNTIYFNTLISTSDFAAHRLPGYVPVELLRINYTQ---------
IPGLMTAIETGGDLTSESVVRRIFSELPSAIDLLTCDAEISGDPTPSVGGKLIKHVLWLANK-MNPTGTLIYKTFCRFPMLLTYEVSLAMSHFGKVKLVVPTYSCHKSSEIFIVATEKNEVMRAMSDEPLSSCAE-------
IIELCQHRLNPVPFQTHSRRRTMYLQNHISRYFESNLSSSMLYFGFELSPEVMINHLGLL-----------------ETMMGDYFKAYGSIREMIN------------IESLMSHWLNTRLMIMIFNLTEVLRRHTFRCGSS------------------------------VMGYSRAFFHILGHCNIKPITRG----
>YP_010805226.1
MELDE--------PHLADSHLQSPILRDEREHFVRYWDTKYAC-----------PRHITLQA--DKVKRMKVV----NVTSPRVDSEVLYG----------DILSGAMLVETTLNVEGYFQQARTQALLDVGAIVRGLGKEI----
PTLRSSLSKIYSKMTTGTPELVNLGVSREVWDKLRLNMAGCRAKPNN----------------------------PPIQGNTLIPGLGRCRYICTSGFLLVS--CRAQWWILDYEQILCLVDTILSRQLALLYISLAKDNDNI--
YVPKEDVLLAAYGWGDVLLQQLGNNAFKLIKYWESLILGNLLEQVDTWSQKSKFLMTMMADYHITFSQLTPCT---PERPCEFCQFKRTTLDKLDNPFVLSELFGVYRHWGHPTVELRAGCTKVQKIATSTKVISLDLLDEVTGQFVQYFVISYIRRHG-
HWPKVNLERLPESSHLRQLAGIEDLSEFSEAVPLKEWARLSFLKNFEFDFSEDWILMIEDKAISPYLSDWRRVYDPTLLGELVPQGESSRRVILECLRKPSVDIKKICETIMCLRVPEEWNI-
VLVHAKEREMKIDPRLFAMMVLEMRLYFCVTEHNIASSIMKYFPQQTMTLGELDLIHRLMVLTKQADGEHP-------HISININLDFQKWNLQWRYMSVRGIALKIDELFGTPGLINYTHLFFDKCHIFSSHRFSVPDGY-
QPGMEFPPTSDLHWMHHKGGFEGLRQKLWTLITLNLLIVISHRCKVRFLITGQGDNQILKVMLPLSDADYMVTYEEEILHQVDHIMNVIEDTAEKMGLPIKREETWVSHRLFAYGKELLVDGAFLPGALKKIARVYFEVNEIYPTCESKLATIHTSSQAAAQKGDNPLISYTISLME
SYYSFMKDIYYPSGPPLKLSLPNQD--
KDNILHFIRFCCLFPRALGGFPVSTLLSYLYRGHPDPVTEGLCFIKILSHEIELYANALQYLVQQPKNTIDLTMLLQDPTAINWPNPILASNIIKNLLMDTVRENTKNKDVCMLFHLNWKDEDKKFTDCLSQAVPFSPRILNDLYMKSPSGARLGFLSTFSNMRSTQAAL--
GDVKRSRAINLIRDSE-KKWILHLYLLRLRYRKKLK-PIPLPCMTELAQQLRDKSWFP---SNSVQGVTVPHPIEQFEYIIHED-----FCTKSTPYSLMIYDPEP---
QGSGQYRMPIRPWYSRGKFNPYLGSQTSEKKTGVLIPLPRSTPSIKAAQRISQIATWAFAEDKNTQDLLNTVISCRTNLNREVLLLSAGYVYGGSLTHRFNDPITSHEVLTNIRPNFLSHVTISSDTLEEYSRGA---VDVTLHFQGMFLVGGMVLHFHCLN-
EANWQHRHLVHAHVRCLKCTKEIVDTPVTLPEV-TLTYPSYQNCPLLYTNIQLSE-----YSAITPVNLIIHNAPVQL-LKNIQYTAAAACMMESNHRV-
IMSSSYRGRSELISCPGTRGLDIQEFIGITFPALLNQLAWACILDRAESVFEILKRSYQSVEEAYAVTVQRSPHSLWGHVRSHLVLPETRQWLIESARIYLPSSESWKGGPNLDLSLSRMLANKIKLVLGLLKSTSTIP---FIWFNVGTMTYKRLLVCWSNLILIRLYLQEKI--
DKHQLFCALRSKPHI----------------DPDDGSLCTKLITWL------------------SNIPHLE----------------------------ITTSELINILTATRI------CYTSEFAETY---IRKS-----------------------------------------------RDLVHPM--ITNRIYPHLGTDSPRSPSDLMFLHSPSEL----
IPAPTSRAPTKITVGRF--APSRKDHMYRLHGLYSTAPYKYYDIISTYLLSAPSYAMCLAEGSGGICRMLLDKFRCKGVYYDSLIEAADFAAHRLPGYIPPEVLLSPNRNR--------
IEGALESAEFGGDLLSLKTVRRIHRVERPPSNVITCDAEIAGDPHPVIYCRLLANVLYISSKHLHDQGILIFKTFCKFPESLCVQLSMMLRWFQNVSICVPISSSHESYEVFLVGTNPLGALKTPSREPQEFLGEDYLHMPVILDLHKYRIKQFPHQEVVMWYWKDLNNIIAGYAENN
WNSSITYFG-QLAPEVEISYESLMDC-------CDSVMQKSELLIREHIRGLGRLRELLN------------LGYLAIHWKNANIMKTILPDLPRISNNIITHSHD------------------------------FSRYGRSFHHILGHLHQQPSTRSQSNT
>UQM99620.1
MA-----GTSLSDILYPECHLNSPIVTGKLVELILITRLPNNQPLDDQTLVNNIKLNLSRGYYQRQFYHILIKKYPNLKLLRHIPYPNGNKYLFNL--------THSGIFNELKSLLSYAHSCYRKISDRLVHLVNSVESGL---YKDVDEHRKASIINLPNIM-
ESSIWYKPFLLWFTLKKEMRLCIKQSTNSNA-------------------------RGMYPNNVFDLEGL-YVQYNRNLLLIINKFTNEAHYLTFEMVLMMSDVLEGRLMIDLAMNADLRYTQF------
KTRGYLLWEFIDNLFCDLGNETYNIVALIEPLVLGFLQLHDESYLLKGAFLQFCLTEITNEFNNLGFSN---VDDHEVVINTIYSIFDLE-
DLHMIAEFFSFFRTFGHPTLEAEEAAAKVRAHMNKPKVIKFKTMMKGHALFCGTIINGFRDRHGGSWPPVIFPNHVSKRVRQLQLNSEGITDEICITEWRSFVGFKFRCFMPLTLDEDLTMYMKDKALASVCSEWDSVYPRETMGYTPPKQTTSRRLVEVFLNDDKFDPVNLIN
YVLSGEYLNDDEFNLSYSLKEKEIKKVGRLFAKMTYKMRACQVVAESLIATGVGKYFKENGMVKDEHELLKTLHKLSVSSIPKDNIKDTQYETISTFLTTDLQKFCLNWRQETTNLFAQRLNEIYGLPGFFNWLHKRLEKSVLYVADPHCPPSGDSHIPLSERENDQIFIKYPMGGIE
GYCQKLWTIITIPFLFLSAYEVGARIAAVVQGDNQAIAITKRVHPNLPFRVKKIISSQLAQKYFDRLRSNLFDIGHNLKANETIVSSHFFVYSKRIYYDGLVLSQALKPISRAVFWSETIVDETRSACSNISTAISKSIEQGYSKWIGYNINILKTIEQLVISLSFTINPSMTKDII-----
QPLLPYWLICASLIPSQLGGFNYMNMCRLYVRNIGDPVTASIADVKRMIKVRLLNKSILQKIIHQKPGDNGFLDWASDPYSVNLPHSQSISMMLKNVTSRMILQHSDNPMLKGLFHFDFEQEDRDLASFLLDRPIIIPRAAHEIMDKSLTGARQEIAGMLDSTKGLIRNSLKSGGV
KPRLTEKLSTYDYEQFRTFNNLMNNKKNDD--LITHEACSVELAILLRKRMWYSLAKGRPIYGLEVPDIIEAVNGYTLKDCEDCYYCQAQSKEYGWFFVPRNCHLDAVHEESNQMR--------
VPYFGSTTEERSEIKLSSLRSASRALKAAVRIATVYTWAYGDDDDSWAEAWYLASFRAKLTLDELKAITPISTSNNIAHRLKDKSTQMKYSGNTLNRVSRYTIISNDKLNFIINGR--
KVDTNLVYQQIMLLGLSILESRFRFQSETGKSNTVLHLHIVKNCCVIEMEDHPYTSSLLPLPNLIKVECNRLIYDETPIIDHDTEQIHKQLYNLPSLDFPRWT--
LSELNEGLAQSLAMTIVDILTKENKDHLTEFKVFSNDDDMNSLITEFMLVDPSLLTLHIGLSIAINWSYNIYYRRPEGKYQMIEYLYTIMIQSPKSYFSVLANALSHPQIFNKFWDAGY-IEPIYGPNISSQDFIRIAIDYMVLCYSTY---LNFWTNDESIQYLLTEFSEEIISKHLCMLNNL-
-----------YNQRSDMPIIRGLTSIEKCTVLTDALKNAQF--------NPGSISWNLDLLEVVAYPASLTYLRRGTIKHIR------------------------------------------------------------------LRNV--------------------------------LNRSIMY-INEVKFINQDI-NVAKGFEDQ--------------
LMGYDRGAHYFPITYLFTDDFRQLLTKQNFQPAKNRWECHIQRRIGLNSTSCYKAVEIGLYLKINLNGDRLFLGEGSGAMMTTYYAILGQALCYYNTGVFSSEVIGQRVLTLSPSEAYLVAKNNPNEMNLRKNIVPLFNGKPESTWIGNPAAFVYISNSIPHSLSMIHNDMESSFE

KDTYTIACEQCHSLALALNLGKSDSVYITKLAPRPNDYTYVLLDTFQIYYEEVFCFLPMYSNPYSSEMYIICSVPRNNSLVYPDLLMKYISGPSDPFDLQNTIINFKLTKYFERRTREKLYGDFCKSTLNNL-SNIDKILIRIGFSLNGPKLIKQMTGHDVGTG----
SQSLISSIQLLANSLVTYLDPERDKSYFFEPYPLKMDSKVKEMIETISKIIIYYLVYHQPKSPRNKYINCLRK-----NAVIPKYLLSKLKKTGYQLEWREDLDASSIKVWWKIVGYSMLI---------
>UOL48948.1
MG-----TQSTSDIIYPECHLDSPIVSGKLAELVFYTGLPSNIRLGDSTIIRNLKSNVTNGRRQNELGNVVRQNYPNIKKINHIEYPSGNYELFRK--------QSYLLTKKFTNILMYGGTCYRKITSRLIKLRKDVCTGL-YEHGEHNMHEIRSIEHLNEIM-
AGSKWFNPFCYWFTIKTEMRDLIKNSRRDFK-------------------------STKSNVLIHHVEDH-SIFLNRNLVIIINHGNYTTHYLTFEMVLLMCDVAEGRLMIDLGMSSDYRFNQL------
KPKGYKLWEFIDSQFEDLGNNTYDLVAMIEPLVLGFLQLRDPSDLLCGAFLQYCMNDLIDIYKSKGYTN---ESDLNQITDIIFDIFDID-
DIHMISEFFSFFRTFGHPTLEAWNAAEKVRSHMNKPKVVKFLTMMKGHALYCSTIINGYRDRHGGAWPPVILPAHASDRIKQAQNNSEGLTDDLVIRNWKSFSGIKFKCFMPLTLDEDLTMYMKDKALAALKSEWDSVYPKEVMSYDPPRQTSSRRLVEVFLSDRNFDPSTLI
NYVISGEYLEDNEFNISYSLKEKEIKQVGRLFAKMTYKMRACQVVAESLIATGIGKYFKENGMTKNEHDLLKTLHKLSVSSVPKDNSGSIQYETVSTFLTTDLQKFCLNWRQETTNLFAERLNEIYGLPGFFNWQHKILEKSVLYVADPHCPPNYDSHIDLNDVVNDHIYIKYPMGGI
EGYSQKLWTITTIPFLFLSAYEIGAKIAAVVQGDNQAIAITMRVHPNLPYKKKKTMCSQLAQQFFYRLRDNLGDIGHNLKANETIVSSNFFVYSKRIYYDGVVLSQSLKPLSRVVFWSETIVDETRSACSNISTAVSKSIEQGFSRWIGYCINLLKIIQQLLISLKFTINPSLTADVT-----
NPIYVYWIISAALVPSQIGGFNYMNLSRLYVRNIGDPVTASFADLKRLITAGLLNQSILQKILHQKPGQSSYLDWSSDPYSVNIPHSQSVTTMLKNVTSRTILQASDNPMLRGLFHFDFEKEDNDLAKFLLDRPVIIPRAAHEIMDKSLTGARQEIAGMLDSTKGLIRNSIRAGGVSP
GLLSKISLYDYEQFRVFNDLMKNKEEDK--LITVDACSVRLAITLRKKMWRDLAQGRQIYGLEVPDSLEVIKGYFLQECADCYYCTANQNQYGWFFCPKKCELDAVNHETNSIR--------
VPYFGSTTEERSEIKLSNVKNASRALKSAIRIATVYTWAFGDSDENWEEAWYLASFRANVTLDELKAITPISTSNNIAHRLRDKSTQMKYSSTALNRVGRYVTISNDNLNFIIDGR--
KVDTNLVYQQIMLTGLSILEEVFRFSLTTGEKNTVLHLHIQKECCVQEMTDHPYIESNSILPVLQRVEGNRLIYDSDPIIEKTKNIITQQVYKSGVLDFPRWT--
LPDLNSTLSKSLAATLIEIITKENRDHLSEFKTLSSDDDINSLITEFLLISPEEFSLDLGLYIAVNWAYDIYYRRPQGKYQMLDYISTIFSQSSRSLFSVISNAISHPKIFQKFWDSGL-VEPIYGANLSSQDFTRISIDLLIKSYQSY---LNYWLDGESIDYILSESDEEVIAKHLCMVASL---------
---YLERSTMPKILNMTSIEKCALLTDVLQQEQF--------NGIYSDWNLDPLDVINYPASITYIRRGTIKHIR--------------------------------------------------------------LRQS-------------------------------LSSEAIA-LDKLLSDRQPIFHPAIKFDEF------------------
HDNYSHFFPITALYTTDYIQLQQNPWSNKMTNYWENHISRRVGINSTSCHKALEISHYIKLEKTKPRMYLGEGSGAMMITYYIILGKTLTYYNTGVFNREVMGQRILTVHPAEALMVERQNPLELGFSRNLKTLFNGKPESSWVGSDENFAYIMSQVSHSLSLIHSDMESTPEKD
QIAVIQEQTYAMALAINLGHDKSLFITKLAPQENSYTTQFLNILTIYYEEVFGFIPASSNPYSTEFYIICSYPRVRTIVSPHIIMSTYSKPDNLIQVGNIIANFKIKTMLDKREKLKRVGDYSKSDLLGL-TKDEKMLLSIGFQLNGPKCMMQLVGYDIGSG----
EYTLIESIRTLLNNIIIAVDDDRESTPFFDPYPLRDDSKLREMMNILTKVNMLFLITENRIYHKKMTIQNLRR-----RGIFPKSILSRFSKTGLKTINILLLETAEVKVWWKCVGYTLLL---------
>UBB42291.1
MS-----GPTVADILYPECHLDSPIVTGKLYSLLEKGGFPHREELNDKTLDRNLKTNTHYVKHQVALKRKLTEKYPNVNKLVPIPYPYGNKYLFRI--------TDTEFTSDVSQILSLSSTCYRKISTRIVKLRDLTTSKL---SSEQVNHEAAMITRLPSIM-
EGGRWYDPFLFWFKCKYIMRKLIKRNTKLRY-------------------------NSYQNEAIYDTSLH-YIAVNKHICIXVNKRNMNVYYFTFELVLMLCDVLEGRMMIDLGMTCDMRFDDF------
RHRGHNLWELIDSLFEDLGNDTYDMVAMIEPLVLGFLQLKDDSVLLKGAFLQFCFTEIEGILISKGFDN---PQDIASIIESLTNIFLMK-
DIHMTGEFFSFFRTFGHPTLEAEEAADKVRAHMNKPKIVDFEIMMKGHALFCGIIINGFRERHGGAWPPHEFPIHTSNIVKSAAANNEALTHECCIQEWKSFVGFRFKCFMPLTLDEDLTMYMKDKALAAIKPEWDSVYLRENMPYPPPKQTTSRRLVEVFLNDTEFDPVNLIN
YVLSGEYLNDKEFNLSYSLKEKEIKRVGRLFAKMTYRMRACQVVGESLIATGVGQYFKENGMVKNEHELLKTLHKLSVSSVARDNITDVQYETMSTFLTTDLQKFCLNWRQETTNIFAERLDEIYGLPGFFSWLHKRLEKSTLYVADPNCPPYNETRMSIDESENKQIFIKNPMGGI
EGYCQKMWTIITIPLLFLSAYECGAKIAAVVQGDNQAIAITKRVHPNIPYKQKKFLCSQLAQQYFNRLRMNMAGIGHNLKANETIVSSHFFIYSKRIYYDGQVLSQALKPMSRCVFWSETVVDETRSACSNICTAVAKSIEQGYSRWLGYSICIFKTLQQLIISLRYTINDSMTKDII---
--
EPLLPNWILAATLLPSQLGGFNYINISRLYIRNIGDPVTASLADLKRMIKVGLLDERILQKVMHQRPGECTFLDWASDPYSININIPASQSVTIVLKNITARMILQNSRNPMLSGLFHDEFDQEDRDLARFLLDRAIIIPRAAHEIMDKSLTGARQEIAGMLDSTKGLIRNGLKAGGLRPR
LVERLSMYDYEQFRVFNNLMAVKEYSI--LITPDACAVELARRIRNVMWNHLTHGRPIYGLEVPDTIEAMNGFLIESCSDCYYCQANNTEYCWFFVPNHCELDQVQRESNSIR--------
VPYFGSTTEERSEIKLSSVRSASRALKAAIRIATVYTWAYGDTDECWEEAWYLASFRANLTLAELKAITPISTSNNIAHRLRDKSTQMKYSGNSLSRVSRYTMISNDNLNFTKDGN--
KIDTNLVYQQVMLLGLASLEDLFRFNKTTGPENTVYHLHVEENCCVIEMEDHPYVVTEEPPPVLRGVYGNRLIYDDHPLEDKEVENIYKQTYTSAILDFPRYN--
IKELNTVLAQSLSLTLIXIITRENKDHLTEFKVLANDDDVNSLITEFLLVDPSEFILYLGLAVAINWSFDVYYRRPQGKYQMVEYLSSYLRITSRSFINVFANALSHPRVFRRFWDVGL-IEPIYGPNLNTQNFNLIAIELLIKGFEVY---LDYWLDDEKAEYMLPESVQDIVSRHLCILCCL--
----------YLDREYMPKILGLTSIEKCSLLTDILQKHKY--------TPQFFTWNLDPLPVVVYPVSLTYIRRGSVKHIR--------------------------------------------------------------LRRM-------------------------------IQFNDPV-ADKVQIDPLNM-RSFVVLRNN---------------
IAGITTSFRAFQAVMLISDDFRKLQGLXEYLPTKNRWESHVHRRVGVNSTSCYKACEIGIYIIMNLKGPRLFLGEGSGAMMTTYYMLLGPAKCFYNTGVLNVGLVGQRVFQIFPSEVMLVSHNNPRDIGLEXDIKVLFNGKPECTWVGDMECFSYLMNSVXHSLALVHNDMES
SLEKSPETILQEQVHSLCIAVNLTKDDGLYVTKIAPRSGDMSNVLLNLLYHYFSEVTCFIPAYSNSCSPECYLICSGKKYHSLVYPEIIIPGLSSFTKNIVISQNILDMKFSIRKDLINKLKLYGDYMNSSLXSL-DETEKLLMSYGFQINGPKLIRQITGHDVGAG----
SINLRAYINSSVNNIINYCDTERLSSHFLEPYPLARDSKLRELIDALCKICVYILLYMKETNSRRQLINMIRG-----RSIIQDYLIKKLTKTSVNMQWSYPLPTAEVKVWWKIVGYSIIH---------
>YP_010790486.1
MNLEGITSDAVNDVLYPECHLNSPIVLNKILTYCHAGEIPSEEILLDKTCIRNYEMNMMQQRSQARSHKSYIESSVGIEHYEHIPSGFCYPDMFSA--------RDRLVSAELEKILERGHRALGKDSSKHAELVSEIYKGA-TGSEDCGNSIMSEVS-
VGAAIERSSEWWQPYLFWFTLKKHMRQTIKETPCRKD-------------------------EKTFKVKAMRGSRF-WIIHDKNMMTLIDHREYKRYHLSFELVLMFCDVIEGRLMLSIGLKLDRKIKVA------
REGLFEIMSTIDSLFPDLGHSTFKIVSLLEALTEAAIQEDDLVEELAGAFMTHCVHQLHELLETV-YDQ---ASSGRIIHSLVRAIREI--
PLMQRGEVFSLFRLFGHPNLEAKSAASKVRAHMYQAKMISPLILSKTVAVFIATIINGYRKQHGGVWPPCELPAHADDTIKDSLMRGEGLSMEDAVRYHTSFSGFKFDQFITPLLDRDLTIFLKDKALSPLRVDWNTIYHPVVAPPCKRPEPGSRRLVDVFINDEDFDPDKVLGYV
TSGEYLNDPEFNISYSLKEKETKTDGRIFAKMTYKMRAAQVAGEYLIATGVGKYFNENGMVKGEHELLKGLTRMSEACIAESNPGSHDFQTYACYLTTDIEKYCCNWRYEVMLPFTNLLDQIYGLPNFFNWQHKRLSRSTIYVADPWCPPDYANPQDLRESPDDGIFIHNPRGSI
EGYQQKIWTIISISQLHVAAVLSNTRITGMVQGDNQNIVVTKRVQKHKTLLEKREIVYREGVRYFRSLREVMAGLGHRLKEDETILSTMFFVYSKRIYIDGTVMSQSLKALTRVAIWSETLVDEIRSGCSTVSTQISKSVQQGLNAYVGYCIDKVRTIDQLLIGLKFTINNKMTVDVV-
----
DPLYQDWLASAIYVPSSLGGLSYMAHTRLFCRNIGDPLVAGIADLKRLIEFKLCDKNMLNKVFNQRHGQSTWLDWALDPYSANLPSTQSITSMLKNITARNVLKNSVNPLLAGIFHEDMDEEDAAVAQWLLDRDVIMPRTANEAIKDLPTGKRNALAGLVDTTKTVVRSALRK
NQGDKRFLHDFQYYDYKQMKALTELLSVIEYRT--AVYRESCSLELSELLRKKTWFVLAEGRPIEGLETPDPIEFSRGDILDDGHICTLCAQRNDYMTWFHIPPRCELDDVLLTTGKAR--------
SPYIGSTTEERTDIVLGRAEDMSMAALDAIHLTKTLIWAFGDDEANWRNAHQAASSRARFELEDLKALTPIATSTNIHHRMHDRFTQQTYSMAREPRVGRYLLISNDSLTAKIEGS--
ATDTNIVYQQQMLYGCALMEDLFRHCHDTGKESTTFHLHVREACCLRYQPMIDPPVEPGPIPDISVDDSNIMMYDPAPLTDKDRERLRTLLVDEAKADILTME--
TADLIELTTATSANSVSDQILELDRAVDKANVALDSPDSTNSLITEMLIMDLMLFWTMLGMNLLLKLARQIHYSRPRGISEIMLILRGLLENTTCILLQPLAHTIAHPKIFRAHLDMGL-ISSLYGHNAGSVNYISHVIDLLCRGAALF---LETVASAGDIIIILCESDDIVALRVLLIEAMI--
----------WAPLPEKYVIRGLSDKDAIDTIEALLVEKK---------SQMMVNLRRPRLFIRTYARTNTYLRRLAIKALR--------------------------------------------------------------ARGY-------------------------------IRIMKPA---------RSIINPVLRVGDITEVDVVKVRYNS--------------
------
DELKPFVYKRRNFQFAGAPIHHQERLMGINSTSAYKLYEILKLSTESYGTDTLHLAEGSGSFAALMGILEPSHKIWVNSMYLESETIPQREARYFPSEILLVDMNTGRVHNLEDRTVPLFNGNPSVTVLGKRACTDYIVEVMGTDVGVITCDLEAKKDLDLSDQVSEWVAILEISMKI
LRSNGTLICKTSLGRDNFWAWVISYVRRHFSSVDIVASSYSGPHSNEIYLICCDPSGEISSPFSI----------SLQVITDLCPEEEEILLEQVTQKILAANAAKARLAAKSLELSDILQSVGFMSNEDHCAQVLTGYDSGTL----RMNHAEIIRTYCQGIISIKDKKRDN--
YLDPFPVDSNARVAHLCDRLLRLIVTMIAMNIAKDIFSKIMKWRRT-----PIVTARLVVKRRANHTITIRW------DEIKLLMKLRGHSESQPGDFELGVA
>YP_010790479.1 L protein [Wenzhou pacific spadenose shark paramyxovirus]
MS---EESITVKELLYPEVHLDSPIVLARRDLIAFMAGLPKVIIENNTPLLFATDQYSAMQRDAAGIRRLRIH--------GTADWRKSYGALFRL--------TDTQLTRRLLKIESHAFNESLLGSDGWKRYIKMMIDLN-LCTESHRKHLVGKIPSLRARI-
ENLKQFSSSLVWLTVKQNLRKIMKESSEGRVS------------------------RFSEKPLILSGNRH-VFYLFTGVVIVHFLKTNVATMLSYEQVLMISDVTESRLSVAVSLPLMTHVRPIF-----SVFFEQVTLHFDTLFPVLGNNIYSYIKMVEALVYGGIAAKEPHPE-
SGHFLIDTLDTIIGVLEDY-----LQDDVLIKIEEFWRDWFDVL-
NVIEIAELMALYRYFGHPFLEPDKACAKVKAHMLAPKTLDRTTLEETYAMWCILLINGYMKAHQNMWPPIEIDPVICPKLHRMKTQGQAILHDEVFNLTEELMAMEWTQFEEVLLDEDLSIFMKDRAVNPPASYWDCMFYKPLLPYKPPPFPGTRRLSAHFLEDSEFTPEKYIN
YVVSGDYLADREFSISYSLKEREVNADGRIFGKMTPMMRGAQVIAEALIANGIGKYFDANGMVKSEKELLESLHRMSAASNIINTSSNAKYQTIAAFLTTDIQKYCLNFREETTNLFTYAQDQIYGLYGFFGWMHRRLKRSWIYVGDPYYPPSTVLGGRTDEPGDSEIGFWGPLG
GIEGFCQKTWTLISIASIHLAAVKVGVRVKAMIQGDNQNVAVTKRVSPLLPEEEKERIILEATYAYADQLREDLANLGLNLKRSETLISQRFFVYSKRLYHSGSLLPQALKALSRVNLWSETLLDEARTACSTIATIGSKGINQGANCHAVFLIVNWFTCRQIIHGLDYRIDSMIPKDVK
DLYFK----
PGLVRLMALLPASLGGFSYLCHSKLFVRNIGDPLVASIADCKRYVEYRLMRRTVLLAFLNQTPSNCTWLQLATDPYALNIPVTRSISVLIERITQENILASSNNPMLKDIFHQDYQQEDEQYAEFLLNREVIYPRVANALLALLPTGERRALAAILETTKTIIRQSLQRGGIDPRTFHSLV
NYESRQFTRFLDIVTSCNKQDGNIKTVDLCSLELAHFARRLSWVHLIGTRPLDGLETPDLIESTRVTYLTTSGNCQQCISGSGAYLWAHYPSETILFRNTGLIHYQR--------
DPYVGSETSERTMVSLLRMDGLSRAVKRSIRICLVYTWVFGNTDEAWENAFKLASTRSAITLEQLKILTPMPTSLNVTHRLHEQCTQMRFMSARPSVASRFITMSTDRIVTWSEVT--
NRDTNIIYQNIMLLGCSLIEDMNQGLIAYTGDPVLMHIHIQEHCCVRFLEPKPFGSEPTLPDLWKPSPSNKLIYDPKGLGETR---VQKLQGVLDQADFSNWP--
SDSIMYLGSVSVGVEIARRLIGALNDGGLRNEAIVVAPSGINLLTAFMVLDLELILKSFARELLLQLSYQITANPCQDSTDVLIRVFNLIDMLSTTVLSGIAGVINHPMILDKIVTLIP-ISDLR---SGPNFVSSCDSSLLRSIVKFYIHELDTKGYFKKVPVFISDPDY---
EANIIVSTKCKALTLYLRSR--YGNVRLEDRPIQVISQQVRSEFEDFIASAHV------LYGGRRVFETLNSLSLRVYYGSPHIVGLMAITELK--------------------------------------------------------------SRIY-------------------------------GKFMPKVT-----------
TLPRLSIRSIQEINISYTDYFS--------
CWTIRYPVGCIVFTPYDPLRMEDKDTTESVDFSYIRYRTYGCSSTSFLKLVSFFYSLNGFKVRTALCLGEGSGSIAAAIVRIWKPRTLFFNTLIKDGDYSSERVQNFVPSEYMRWWSEVNREVYYDEGFKLLTDGTEESTNLTQLRCHYFIRSTIKDLCDIITCDMEDKNLSGFTERWR
ARSLVLKTAWLLLRRRGTVIIKLYASRDNIWRFTARLAATLFNNVQIYRSMYSNPNGLEFYLILSCPKIIEEGIVSDQDESI-----------PIILRQGGFHSEDPSSLKSCSERLVPTSKLI-QAHNDFVEHLGFGPLGDQMVNRQF-----------IDKSITIVERQIKTMISQVKQTRKE--
KIMHFAIRPDRSIRDYADRLCLLQFLAK-------IYSNVVDHVPS------SLTTQVHIKQVGFREFDYQLSKPL----QKILWKIVGNITPIPREPLFSRK
>WMZ41129.1 RNA-directed RNA polymerase L [Dehong virus]
---------MSHPTQYPDARLSSPITLDHCDLFSRSLGLYSNYSHNPKIKNCRIPHHIYRLRNSNSLKSFLE-----DCSLLTIPFHSTWSYLIESLLDDAKNHNRDFQYLIPDDALNYAD----WDNDFTKNMIEEIVGVNGIFTGDTSKTEKTIIDI-----
EQNNPYFWGMLLLIHLSQIARRIKGQRGSIRSDWKFVGKAISIMGISDFILIKVPVHAILRSKNSLQEAHPAQTWYNKQDLRPYYLNDEYLISIASYESFIMIKDVFIERFNTIEISRRAWLEDPETADYPPVNLIRELYIAGDLILSQFFEDGFKLIKHLEPLCSSCIQTQGIYTPRKYWFQ
AQMIKSYYDELTQLERTYDVSDFDKQCADNFIKLIIQSKLTPQQYCELFSVQKHWGHPLLYNDVALEKVKKHAQAVKILKPSIIFETFCVFKFIIAKNHYNSQG-
SWYKTTHDLNLTPYLRSHILANSFPSQSEIYPHLWEWYYVTHDPIFSVKIISDLSIFIKDRATAVTADCWDSVFDKNVLGYNPPVRFQSKRVPEQFLTQEDFSLNQVITFAERLEYLSPDYRNFSFSFKEKELN-
TGRTFGKLPYKVRNVQTLAEALLADGLAKAFPSNMMVVTEREQKEALLHQASWHHNSAS-
VGENAIVRGASFVTDLEKYNLAFRFEFTRHFIEYCNNCYGVRNLFNWMHYLIPLCYMHVSDYYSPPYCLNENNRLNPPDCANAYHYHLGGIEGFQQKLWTCISCAQITLVELKTKLKLKSSVMGDNQCITTLSLFSIDAPHDYQEKEAELNAARVAVELAITTGFSGIFLKPEETFV
HSGFIYFGKKQYLNGVQLPQSLKTMARCGPLSDSIFDDLQGSLASIGTAFERGVSETRHILPNRWVAAFHTMISVNLLNSNHLGFPLGTSLDTLCFRKPLTYTEKVLALITPQVLGGLSFLNPEKIFYRNISDPLTSGLFQLKKALTYLN-
KEEFFYILIAKKAGQSSPSDFIMNPLGLNVPGSREIITYLRQIVRQNITLTASNKIINSLFHVGSDLEDNEVCNWLLSSDPVMSRFASDIFSRTPSGKRLQVLGYLEGTRTLLASRTLSLSSDGSMLGKLRDLTKNRWKSWFSYIDTFDEELSDALEKYNCTVDIADFLRAFSWSDILKGR
KLIGATLPCLLEQFISSWIDIDCRVASQFNCFLRFNNTYVNCAIDKKVVQAHPSQRRLTWSIGNRAPYIGSRTEDKIGYPPLKVNCPSAALKETIEMVSRLIWVTQGNANTENLLKPLITSRINLDFNVLSNFLPTHYSGNIVHRYNDQYGQHSFMANRMSNTSTRAIISTNTLGMY
AGGGQAAIDSNIIFQNTINLAVAIFDICFALKEIPQNFSAQMHLNLNK-CCTRQVPSEYLTFDTKLKVDLNKYSDNELIYDKDPLSSGIRGRSGHVEPKALNLQLNSSDIGSYDFITIAAWTLGTTIVESI---
LSDEASQSTDPISTGSTKAFVTQFLVFPIESILYAFGANLIIGLISYQQLKGVKDLSDLMYSIQINIQNLSHRSLRILQSTFRQELVLQKLSKTIPLITVVLGGSSGEKGVSDAVRLLLISSVRYFIDNIVKILLKR--------------TVTLPVWLYF-----PSEGQKIVPIIKIIKELIRLVCP------
SILCDIKVT--KDNYTIPESLWIYPSRSTK-TNHYYASLNYWRYQMRNSNLTNKQESGSIFLTEANKILSKSEQLEKCKMSTSSVGNVLTVNTANKVYKSNSEMNPVHRYEIE----QPINLDLSEQLPVKLSDRKGQIKRKTNFNPFYNAEQSNA-
LDQKTEQSFVLIRKLNDEIGPSMHPYLYITCDNFAESLKLALKNYNLDDQV--DKLIQYYKRTLL----NAIDSNNFCRFTGVVSSMHYKLYDLLPN---TPIKVATCLAEGEGGGARLLLKWKRTQSLFFNTLATDVQQEVEILSGRVIPRMLYNIEGLNTMI--
ENKALILNNLTVQETDITKMSWLNSVKQLLPNDSDLLTMDAETIEENNRIPLYEMIVK-IWDLPD-KQLPNISIIKIFVLDLEGSSYIINQALQRYSYVQLRKPYSSNARNSEWYLFCSQKRQGKIKLQYD-----------PSVFKYILSKAM------
VRQIQAIPYWLNHIEQHYKIEKHRFFLSLGFPSLELSFSHRYALSLESSQTLYGISIAITKGTKYLSKLLEDYNNKAPLLDIRNHFIHSLRGRLTKLYNDLLKLKLVLTSVNDGK-NWSYHQERLSNVHSVCYEHLLELNCNCYNRILLKLDFIKNTTTAEKKLFNRIVGYIIMFPNGINYNPK
>Orthomarburgvirus_marburgense_YP_001531159
---------MQHPTQYPDARLSSPIILDQCDLLARSLGLYSHYSHNPKLRNCRIPHHIYRLRNSTALKTFLQ-----NCSILTVPFHSIWDHILTSIQYDAINHVDDFKYLLPSELVKYAN----WDNEFLKAYLNKILGLDHVFSASARSQCEDFSP-------
KENPYYWGMLLLVHLSQLARRIKGQRGSLRSNWKFIGTDLELFGIADFVIFKVPVKTIIRNAVSLQASKPGRIWYRDQNLTPYLCDDEFIVSVASYECFIMIKDVFIERYNTWEICARAWLEDSDGADYPPLDVLGELYNQGDQIIAMYLEDGFKLIKHLEPLCVSCIQTHGIFTPRKY
WFQSQMIKSYYDELHDLNLKLQISDNKAECAQNFIKTIVQAKLTPQQYCELFSLQKHWGHPVLYNDVALDKVKKHAQSTKILKPKVMFETFCVFKFIVAKNHYHSQG-
SWYKTTHDLHLTPYLRQHIVSNSFPSQAEIYQHLWEWYFVEHEPLFSTKIISDLSIFIKDRATAVNQECWDSVFDRSVLGYNPPVRFQSKRVPEQFLGQADFSLNQILEFAEKLEYLAPSYRNFSFSLKEKELN-
IGRTFGKLPYRVRNVQTLAEALLADGLAKAFPSNMMVVTEREQKEALLHQASWHHNSAS-
IGENAIVRGASFVTDLEKYNLAFRYEFTRHFIDYCNRCYGVKNLFDWMHFLIPLCYMHVSDFYSPPHCVTEDNRNNPPDCANAYHYHLGGIEGLQQKLWTCISCAQITLVELKTKLKLKSSVMGDNQCITTLSLFPIDAPNDYQENEAELNAARVAVELAITTGYSGIFLKPEETFV
HSGFIYFGKKQYLNGVQLPQSLKTMARCGPLSDSIFDDLQGSLASIGTSFERGTSETRHIFPSRWIASFHSMLAINLLNQNHLGFPLGFNIDISCFKKPLTFSEKLIALITPQVLGGLSFLNPEKLFYRNISDPLTSGLFQLKNALEFLE-
KEELFYILISKKPGLADASDFVMNPLGLNVPGSKEIITFLRQTVRENITITSQNRIINSLFHIGSDLEDQRVCEWLLSSNPVMSRFAADIFSRTPSGKRLQVLGYLEGTRTLLASRTISLTTEGTMLMKLRELTRNRWKSWFSYIDALDDDLSESLEKFTCTVDVANFLRAYSWSDVLKG
KRLIGATLPCLLEQFEVKWINLSEDLREQFNLSSDTELNYVSCALDRKVVQKHPSVNRLAWTIGNRAPYIGSRTEDKIGYPPLRVNCPSAALKEAIEMVSRLLWVTQGTADREKLLIPLLNSRVNLDYQTVLNFLPTHYSGNIVHRYNDQYGQHSFMANRMSNTSTRAIISTNTLGK
YAGGGQAAIDSNIIFQNTINLGVAVLDIALSLAKLSSASNVTFRLMLNK-CCTRHVPSEYLYFDKPLDVDLNKYMDNELVYDNDPLCSGIKGRLGRVSRSTLTLSLNVSDIGSYDFPTIAAWTLGETIVGSI---
FSDESSQSTDPISSGCTKTFVTHFLVYPVESIFYAFGANLIVESLSLSRIKSIKNLSDLTFLISSTIRNLSHRSLRILQSTFRHELVLTRLAHHIPLISLMLGGSAGEKSSSDAVRLFLTASYQNFINNFSCLMKKG--------------QSSLPVWLYF-----PSEGQQLKPILKILQRLSDLLSP------
DKIQKRKIL--ADTCCPIGSFWVYPSKSTR-
TNHYYASLNYWRDKANKVKNTPFSHTSSVSSNQQVTNSKYIVYPENITINARTRLINYGSTALQGMDTKLSEQNLVENCPSETKHDQQMFPEDNMQTAHIHSSSPFQILIKSDFDASKNSEINNLNLTEYTLNTKLLTTPTRTEILDSPASYLYVDCSNIPSSLDPGFRSMSDQNQV
--QMLINTYKRDLH----ACFDSNQFCRFTGVVSSMHYKLYDLLPP---GKLKKAICLAEGEGSGARLLLKWKETDYLFFNTLATDSQQEAEILSGRVIPRMLYNIDRLSALL--ESRRLILNNLTIQITDITNPLWLDSVIQYLPEDSDILTMDAETTKDETREQLYKTIVN-IWTRTS-
PNIPKISIIKVFLLDYEGTLFLMKNAIQYYGQVQLKKPYSSNAKNSEWYLCCGKRRIQRLQIDFS-----------DQVGIFLICKAM------
SRQRQAIPYWLKHIEKNYPASLHEFFLTLGFPSLESSFCHRYTIPFSEGKALFHVQSYVRQGKQHLHSLMLDYENNSPLLDLRNHFICSLRGKITKYYNDILKLNLVIKAVEKGK-
NWSQLVEILPNMHSVCIVHVDHECSGCEKRLLLKLDFIRNTKIAEQKLLNRVIGYILFFPFGLFKSGS
>Orthomarburgvirus_marburgense_RAVN_YP_009055228

---------MQHPTQYPDARLSSPIILDQCDLLTRSLGLYSHYSHNPKLRNCRIPYHIYRLRNSTALKTFLQ-----NCSILTVPFHSIWDHIITSIQHDAINHINDFKYLLPSELIKYAN----WDNEFLRVFLNKILRLDHAFTNSAKLQCEDFSP-------
KENPYYWGMLLLVHLSQLARRIKGQRGSLRSNWKFIGVDLELFGIADFVIFKVPIKAIIRNATSLQASKPGKTWYRDQNLTPYLCDDEFVVSIASYECFIMIKDVFIERYNTWEICARAWVEDNEEADYPPLGILRDLYNQGDQIITMYLEDGFKLIKHLEPLCVSCIQTYGIFTPRKY
WFQSQMIKSYYDELQSLNLKLQIPDNRTECAQNFIKTIIQAKLTPQQYCELFSLQKHWGHPVLYNDVALDKVKKHAQSTKILKPKVMFETFCVFKFIVAKNHYHSQG-
SWYKTTHDLHLTPYLRQHIVSNSFPSQAEIYQHLWEWYFVEHEPLFSTKIISDLSIFIKDRATAVNRECWDSVFDRSVLGYNPPVRFQSKRVPEQFLGQADFSLNQILDFAEKLEYLAPSYRNFSFSLKEKELN-
IGRTFGKLPYRVRNVQTLAEALLADGLAKAFPSNMMVVTEREQKEALLHQASWHHNSAS-
IGENAIVRGASFVTDLEKYNLAFRYEFTRHFIDYCNRCYGVKNLFDWMHFLIPLCYMHVSDFYSPPHCVTENNRNNPPDCANAYHYHLGGIEGLQQKLWTCISCAQITLVELKTKLKLKSSVMGDNQCITTLSLFPIDAPNDYQENEAELNAARVAVELAITTGYSGIFLKPEETFV
HSGFIYFGKKQYLNGVQLPQSLKTMARCGPLSDSIFDDLQGSLASIGTSFERGASETRHIFPSRWIAAFHSMLAVNLLNQNHLGFPLGFSIDVSCFKKPLTFSEKLIALITPQVLGGLSFLNPEKLFYRNISDPLTSGLFQLRNALEFLR-
KEELFYILIAKKPGLADASDFVMNPLGLNVPGSREIITFLRQTVRENITITSQNRIINSLFHIGSDLEDQRVCEWLLSSNPVMSRFAADIFSRTPSGKRLQVLGYLEGTRTLLASRTISLTTEGTMLMRLRELTKSRWKSWFSYIDALDDDLSESLEKFICTVDVANFLRAYSWSDVLKGK
RLIGATLPCLLEQFNVKWVNLSEDLKEQFKLSSDAELNYVSCALDRKIVQKHPSDNRLAWTIGNRAPYIGSRTEDKIGYPPLRVNCPSAALKEAIEMVSRLLWVTQGTADREKLLIPLLNSRVNLDYQTVLNFLPTHYSGNIVHRYNDQYGQHSFMANRMSNTSTRAIISTNTLGK
YAGGGQAAVDSNIIFQNTINLGVAVLDITLSLSKLSSTSNVSFRLMLSK-CCTRHVPSEYLFFDKPLDVDLNKYMDNELVYDNDPLCSGIKGRLGRVSRSTLSLSLNVSDIGSYDFPTIAAWTLGETIIGSI---
FSDESSQSTDPISSGCTKTFVTHFLVYPVESIFYAFGANLIVESLSLSRINSIKSLSDLTFLISSTIRNLSHRSLRILQSTFRHELVLTRLAHHIPLISLMLGGSAGEKSSSDAVRLFLTASYQNFINNFSCLMRKN--------------QSPLPVWLYF-----PSEGQQLKPILKILQRLSCLLTT------
KKVQNHRPV--ADTCFLTDNFWVYPSKSTR-
TNHYYASLNYWRDKANKIKNTSFSHASSISSSQEVVNLKHTSRLDETPMSERAQSTNHEPTALQEVCTEYSEQDPAKSYLLELRHDLQVSSRGCLQATCPHHPSPSQTTTEPNCDAIKRSETNDPRLMSSILDMRSLKTPMRIESRNSLASYLYVDCSNISSSLDSGFRNMSDRNQ
V--QMLINTYKRDLY----TCFDSNQFCRFTGVVSSMHYKLYDLLPA---GKLGKAICLAEGEGSGARLLLKWKETDYLFFNTLATDSQQEAEILSGRVIPRMLYNIDKLSVLL--ESRKLILNNLTIQITDITNPLWLDSVIQYLPEDSDILTMDAETTKEETREQLYKTIIN-IWARTS-
PNIPKTSIIKVFLLDYGGTLFLMKNAIQYYGQVQLKKPYSSNAKNSEWYLCCGKRRVQRLRVDFP-----------DQVGIFLICKAM------
SRQRQAIPYWLKHIEKNYPASLHEFFITLGFPSLESSFCHRYTIPFTEGTALFHVQSYVRQGRQHLHSLMLDYENNSPLLDLRNHFICSLRGKITKYYNDILKLNLVVRAVERGK-
NWSQLVESLPNMHSVCITHVDHECIGCERRLLLKLDFVRNTKIAEQKLLNRVIGYILFFPFGFSRPK-
>Cuevavirus_lloviuense_YP_004928143
----------MQEAQYPDARLSSPIVLDQCDLVARACNLYSNYSYNPTLKKCRLPKHIYRLQYDIAVNEFLA-----GVPLSTLPIDYLVPLIINANTDSIKIQDQPSIKNHIKDGISFAL----HDSGFLNYYLKTIGINTMIDLGTFHNSAKKIL--------
LNERIIDEAVTWLDLVVAARRARMNRGNTRSTWYASGELIDILGYGDYIFWKIPLKLLPLNKRGVPHASAP--WYDPTTFKSCSEGSTVVVSISTVEVLLMIKDVAFSRMNTLLICKLAELEDTL-
TDYPTVESLHDLYRSGDYLISLLGSTGYDIIKYLEPLCLGRIQLLSNFTPRKTWYYSQMQVALLKDLAERTKGRELTLYQQTKIRSFHQILFRLKLRAQQWCELFSIQKHWGHPVLHSELAIAKVKKHGTCLKVLRPSIMFDTFCVFKYMVGKHHFDNTG-
HWYRVTHDKNLTPTLKTYIQKNHFPTFRDVRKFLWEFYLVQHEDLFSTKVISDLSIFIKDRATAVESICWDAVFEPNVLGYAPPMRFITKRVPEQFLAQEDFSLEAVISYAEKLQYLEPVNRNFSFSLKEKELN-
IGRTFGKLPYLTRNLQTLCEALLADGLAKAFPSNMMVVTEREQKETLLHQASWHHEQGN-
LGDHVQVRGASFVTDLEKYNLAFRFEFTRPFIEYCNHCYNIKNAFNWMHYLIPSCYMHVSDYYNPPHGLTLENRGYPPEGPSSYRGHLGGIEGLQQKLWTSISCAQISLAELRSGFKTRSAVMGDNQCITVLSVFPDHYQEDLQELEAEDNAARVAAILARITASCGIFLKPEETFV
HSGFIYFGKKQYLNGVQLPQSLKTAARMAPLSDAIFDDLQGTLASIGTAFERAIAESRHIFPIRIMAAIQSYLAVKVLQENHLGFPKNTDLGVLALGKPISAKMIRLCLSIPQVLGGLSFLNPEKCFYRNLSDPVTSGLYQVKKYLAYIG-
QGDLFPLFVAKRPGDASAIDLVLNPLGLNYPGAQDLTSFLRRIVRQSITFHSKNKLINSLFHANADLEDEYLCEWLLSSEPVMSRFAADIFSRTPSGKRLQILGYLEGTRTLLASRNLHTNSESSLLSRLRALTTKRWQLWFSYMDQYDEDLGDVIGQLTCTLDIANVLREYSWSHVL
QGRRLIGATLPCVPEQFELTWLTGDKACKHCQSKLRDKKKPYVSAALVDKIISQRPSYHRLSWTIGTLTPYIGSRTEDKIGQSAFKPRCPSAALRETIELASRILWVTQGSSQAEDIIRPFCEARINLPVQELFKLLPSHYSGNIVHRYNDQYGPRSFMANRMSNTATRIIVSTNTLGPY
SGGGQAARDSNIIFQNVINFAVAVLDIQNSFLDQPMSEFKHIHLHIKD-CCTREVPSQYLTYKANFGMNLQKYQSNELIYDKDPLRCGLTCRVSVENFFLKHHFKNSVEGELLRYPHLSAWGLANTVMTSL---
LADMNNASTDLISGNEVKSFTSQFLAYPLIGIFYSFGAITTFKLGRN-LSLQKITKSQIIYYLDSQLYNLPHRALRVFKTTFQHPAVLESLLTIDPSFSFFLGGASGGKSVADATRLFLRLAIQVFLNFISLYSQQKI------------GKTLPLWVVF-----PLEGEPDEPILEFLSELLRALSH------
PLPDQEASTTRIQPSCVFRSEQIYRSMSTM-SNFFHASLACWRDRRQGRARGSSNVKGTILLYDIKGLLSLG--WKDCSLSNAPLKNSLSSSHCSRDILEHRSRNITECPPLL----SRRTRISQESK---------------QHVRSA----PGDAPPDPTHRPGCLRLNVFKTEQD-----------------
--------ETGLSTEETIEQLLARLK----MMPDTTSYCRFTGVIASMHYKLDEVLYP--TNETPTVALLAEGEGSGALLLIRKYRARQIFFNTLATGNMVEAEVLMGESVPRMLLP---
LLPSINNKEIKIFFNTSVTMKTDITDPKWFDSIEDRLPRSLDLLVMDAETIGNSIRIPLYNAVIKCIDRDKM-QDKPKCVIVKLFLNDYLGTCWILNHITLRYNKVEIIKPLSSNARSSEWYLKAEQPVLSSENVNWE-----------GHTT-KLIKKAL------
HCQLSRQSYWVTHLMSYANLDLHREYIRMGFPPLTSVLHYRYHMVATESSPLLTIRRHTNLLLQEIRELVRDYFLSRCTRTQAFHFIKTQKGRITKLANDYLKWRLVLSALERYT-LFEQALNNIEPYIQVCILGPCNKALCSNDHLPIPLLYLTRTDQAERKLLNRLVGLVQFFPDGL--
---
>Bundibugyo_ebolavirus_YP_003815440
--------MATQHTQYPDARLSSPIVLDQCDLVTRACGLYSSYSLNPQLKNCRLPKHIYRLKFDATVTKFLS-----DVPIVTLPIDYLTPLLLRTLSGEGLCPVEPKCSQFLDEIVSYVL----QDARFLRYYFRHVGVHDDNVGKNFEPKIKALI--------
YDNEFLQQLFYWYDLAILTRRGRLNRGNNRSTWFANDDLIDILGYGDYIFWKIPLSLLSLNTEGIPHAAKD--
WYHASIFKEAVQGHTHIVSVSTADVLIMCKDIITCRFNTTLIAALANLEDSICSDYPQPETISNLYKAGDYLISILGSEGYKVIKFLEPLCLAKIQLCSNYTERKGRFLTQMHLAVNHTLEELIEGRGLKSQQDWKMREFHRILVNLKSTPQQLCELFSVQKHWGHPVLHSEKAIQKVKK
HATVIKALRPVIIFETYCVFKYSIAKHYFDSQG-SWYSVISDKHLTPGLHSYIKRNQFPPLPMIKDLLWEFYHLDHPPLFSTKIISDLSIFIKDRATAVEKTCWDAVFEPNVLGYSPPNKFSTKRVPEQFLEQENFSIDSVLTYAQRLDYLLPQYRNFSFSLKEKELN-
VGRAFGKLPYPTRNVQTLCEALLADGLAKAFPSNMMVVTEREQKESLLHQASWHHTSDD-
FGENATVRGSSFVTDLEKYNLAFRYEFTAPFIEYCNRCYGVKNLFNWMHYTIPQCYIHVSDYYNPPHGVSLENREDPPEGPSSYRGHLGGIEGLQQKLWTSISCAQISLVEIKTGFKLRSAVMGDNQCITVLSVFPLETDSNEQEHSSEDNAARVAASLAKVTSACGIFLKPDETFV
HSGFIYFGKKQYLNGVQLPQSLKTATRIAPLSDAIFDDLQGTLASIGTAFERSISETRHVYPCRVVAAFHTFFSVRILQYHHLGFNKGTDLGQLSLSKPLDFGTITLALAVPQVLGGLSFLNPEKCFYRNLGDPVTSGLFQLRTYLQMIN-
MDDLFLPLIAKNPGNCSAIDFVLNPSGLNVPGSQDLTSFLRQIVRRTITLSAKNKLINTLFHSSADLEDEMVCKWLLSSTPVMSRFAADIFSRTPSGKRLQILGYLEGTRTLLASKVINNNAETPILDRLRKITLQRWSLWFSYLDHCDQVLADALIKVSCTVDLAQILREYTWAHILEG
RQLIGATLPCMLEQFNVFWLKSYEQCPKCAKSRNPKGEPFVSIAIKKQVVSAWPNQSRLNWTIGDGVPYIGSRTEDKIGQPAIKPKCPSAALREAIELTSRLTWVTQGGANSDLLVKPFVEARVNLSVQEILQMTPSHYSGNIVHRYNDQYSPHSFMANRMSNSATRLVVSTNT
LGEFSGGGQSARDSNIIFQNVINFSVALFDLRFRNTETSSIQHNRAHLHLSQ-CCTREVPAQYLTYTSTLSLDLTRYRENELIYDNNPLKGGLNCNLSFDNPLFKGQRLNIIEEDLIRFPHLSGWELAKTIIQSI---
ISDSNNSSTDPISSGETRSFTTHFLTYPKVGLLYSFGAIVSYYLGNTIIRTKKLDLSHFMYYLTTQIHNLPHRSLRILKPTFKHVSVISRLMSIDPHFSIYIGGTAGDRGLSDATRLFLRVAISSFLQFIKKWIVEY-------------KTAIPLWVIY-----PLEGQNPDPINSFLHLIIALLQN------
ESPQNNIQFQEDRNNQQLSDNLVYMCKSTA-SNFFHASLAYWRSRHKGRPKNRSTEEQTVKPIPYDNFHSVKCASNPPSIPKSKSGTQGSSAFFEKLEYDKERELPTASTPAE----QSKTYIKALSSIYHGKTPSNAAKDDSTTSK-----GCDSKEENAVQASHRIVLPFFTLSQN------
--------------DYRTPSAKKSEYITEITKLIRQLK----AIPDTTVYCRFTGVVSSMHYKLDEVLWE--FDSFKTAVTLAEGEGSGALLLLQKYKVRTIFFNTLATEHSIEAEIVSGTTTPRMLLP---
VMAKLHDDQINVILNNSASQVTDITNPAWFTDQKSRIPTQVEIMTMDAETTENINRSKLYEAIQQLIVSHID-TRVLKIVIIKVFLSDIEGLLWLNDHLAPLFGSGYLIKPITSSPKSSEWYLCLSNFLSASRRRPHQ-----------GHATMQVIQTAL------
RLQVQRSSYWLSHLVQYADINLHLSYVNLGFPSLEKVLYHRYNLVDSRKGPLVSILYHLTHLQAEIRELVCDYNQQRQSRTQTYHFIKTTKGRITKLVNDYLKFYLVVQALKHNC-
LWQEELRTLPDLINVCNRFYHIRDCSCEDRFLIQTLYLTRMQDSEAKLMERLTGFLGLYPNGINA---
>Tai_Forest_ebolavirus_YP_003815431
--------MATQHTQYPDARLSSPIVLDQCDLVTRACGLYSAYSLNPQLKNCRLPKHIYRLKYDTTVTEFLS-----DVPVATLPADFLVPTFLRTLSGNGSCPIDPKCSQFLEEIVNYTL----QDIRFLNYYLNRAGVHNDHVDRDFGQKIRNLI--------
CDNEVLHQMFHWYDLAILARRGRLNRGNNRSTWFASDNLVDILGYGDYIFWKIPLSLLPVDTQGLPHAAKD--
WYHESVFKEAIQGHTHIVSISTADVLIMCKDIITCRFNTLLIAAVANLEDSVHSDYPLPETVSDLYKAGDYLISLLGSEGYKVIKFLEPLCLAKIQLCSNYTERKGRFLTQMHLAVNHTLEELTGSRELRPQQIRKVREFHQMLINLKATPQQLCELFSVQKHWGHPVLHSEKAIQKVKK
HATVIKALRPIIIFETYCVFKYSIAKHYFDSQG-TWYSVTSDRCLTPGLSSYIKRNQFPPLPMIKELLWEFYHLDHPPLFSTKVISDLSIFIKDRATAVEKTCWDAVFEPNVLGYNPPNKFATKRVPEQFLEQENFSIESVLHYAQRLEYLLPEYRNFSFSLKEKELN-
IGRAFGKLPYPTRNVQTLCEALLADGLAKAFPSNMMVVTEREQKESLLHQASWHHTSDD-
FGENATVRGSSFVTDLEKYNLAFRYEFTAPFIEYCNRCYGVRNLFNWMHYTIPQCYIHVSDYYNPPHGVSLENRENPPEGPSSYRGHLGGIEGLQQKLWTSISCAQISLVEIKTGFKLRSAVMGDNQCITVLSVFPLETESSEQELSSEDNAARVAASLAKVTSACGIFLKPDETFVH
SGFIYFGKKQYLNGVQLPQSLKTATRIAPLSDAIFDDLQGTLASIGTAFERSISETRHVVPCRVAAAFHTFFSVRILQYHHLGFNKGTDLGQLSLSKPLDFGTITLALAVPQVLGGLSFLNPEKCFYRNLGDPVTSGLFQLKTYLQMIH-
MDDLFLPLIAKNPGNCSAIDFVLNPSGLNVPGSQDLTSFLRQIVRRTITLSAKNKLINTLFHSSADLEDEMVCKWLLSSTPVMSRFAADIFSRTPSGKRLQILGYLEGTRTLLASKIINHNTETPILDRLRKITLQRWSLWFSYLDHCDQVLADALTQITCTVDLAQILREYTWAHILEG
RQLIGATLPCILEQLNVIWLKPYEHCPKCAKSANPKGEPFVSIAIKKHVVSAWPDQSRLSWTIGDGIPYIGSRTEDKIGQPAIKPKCPSAALREAIELTSRLTWVTQGGANSDLLVKPFIEARVNLSVQEILQMTPSHYSGNIVHRYNDQYSPHSFMANRMSNSATRLVVSTNTLGE
FSGGGQSARDSNIIFQNVINFAVALFDLRFRNVATSSIQHHRAHLHLSK-CCTREVPAQYLVYTSTLPLDLTRYRDNELIYDDNPLRGGLNCNLSFDNPLFKGQRLNIIEEDLIRLPYLSGWELAKTVIQSI---
ISDSNNSSTDPISSGETRSFTTHFLTYPKIGLLYSFGALISYYLGNTIIRTKKLTLNNFIYYLATQIHNLPHRSLRILKPTLKHASVISRLISIDSHFSIYIGGTAGDRGLSDAARLFLRTAITVFLQFVRKWIVER-------------KTAIPLWVIY-----PLEGQSPSPINSFLHHVIALLQH------
ESSHDHVCAAEAHSRVETFDNLVYMCKSTA-SNFFHASLAYWRSRSKNQDKREMTKILSLTQTEKKNSFGYTAHPESTAVLGSLQTSLAPPPSADEATYDRKNKVLKASRPGK----YSQNTTKAPPNTSCRDVSPNITGTDGCPSA-----NEGSNSNNNNLVSHRIVLPFFTLSHN--
-----------------YNERPSIRKSEGTTEIVRLTRQLR----AIPDTTIYCRFTGIVSSMHYKLDEVLWE--FDNFKSAITLAEGEGSGALLLLQKYKVETLFFNTLATEHSIEAEIISGITTPRMLLP---
IMSRFHGGQIKVTLNNSASQITDITNPSWLADQKSRIPKQVEIITMDAETTENINRSKLYEAVQQLIVSHID-PNALKVVVLKVFLSDIDGILWLNDNLTPLFGLGYLIKPITSSPKSSEWYLCLSNLLSTSRRLPHQ-----------SHTTMHVIQTAL------
QLQIQRSSYWLSHLVQYANHNLHLDYINLGFPSLERVLYHRYNLVDSQKGPLTSIVQHLAHLQTEIRELVNDYNQQRQSRTQTYHFIKTIKGRITKLVNDYLKFFLIIQALKHNC-
TWQEELRALPDLISVCTRFYHTRNCSCENRFLVQTLYLSRMQDSEIKLIDRLTGLLSLCPNGFFR---
>Zaire_ebolavirus_NP_066251
--------MATQHTQYPDARLSSPIVLDQCDLVTRACGLYSSYSLNPQLRNCKLPKHIYRLKYDVTVTKFLS-----DVPVATLPIDFIVPVLLKALSGNGFCPVEPRCQQFLDEIIKYTM----QDALFLKYYLKNVGAQEDCVDEHFQEKILSSI--------
QGNEFLHQMFFWYDLAILTRRGRLNRGNSRSTWFVHDDLIDILGYGDYVFWKIPISMLPLNTQGIPHAAMD--
WYQASVFKEAVQGHTHIVSVSTADVLIMCKDLITCRFNTTLISKIAEIEDPVCSDYPNFKIVSMLYQSGDYLLSILGSDGYKIIKFLEPLCLAKIQLCSKYTERKGRFLTQMHLAVNHTLEEITEMRALKPSQAQKIREFHRTLIRLEMTPQQLCELFSIQKHWGHPVLHSETAIQKVKKH
ATVLKALRPIVIFETYCVFKYSIAKHYFDSQG-SWYSVTSDRNLTPGLNSYIKRNQFPPLPMIKELLWEFYHLDHPPLFSTKIISDLSIFIKDRATAVERTCWDAVFEPNVLGYNPPHKFSTKRVPEQFLEQENFSIENVLSYAQKLEYLLPQYRNFSFSLKEKELN-
VGRTFGKLPYPTRNVQTLCEALLADGLAKAFPSNMMVVTEREQKESLLHQASWHHTSDD-
FGEHATVRGSSFVTDLEKYNLAFRYEFTAPFIEYCNRCYGVKNVFNWMHYTIPQCYMHVSDYYNPPHNLTLENRDNPPEGPSSYRGHMGGIEGLQQKLWTSISCAQISLVEIKTGFKLRSAVMGDNQCITVLSVFPLETDADEQEQSAEDNAARVAASLAKVTSACGIFLKPDE
TFVHSGFIYFGKKQYLNGVQLPQSLKTATRMAPLSDAIFDDLQGTLASIGTAFERSISETRHIFPCRITAAFHTFFSVRILQYHHLGFNKGFDLGQLTLGKPLDFGTISLALAVPQVLGGLSFLNPEKCFYRNLGDPVTSGLFQLKTYLRMIE-
MDDLFLPLIAKNPGNCTAIDFVLNPSGLNVPGSQDLTSFLRQIVRRTITLSAKNKLINTLFHASADFEDEMVCKWLLSSTPVMSRFAADIFSRTPSGKRLQILGYLEGTRTLLASKIINNNTETPVLDRLRKITLQRWSLWFSYLDHCDNILAEALTQITCTVDLAQILREYSWAHILEG
RPLIGATLPCMIEQFKVFWLKPYEQCPQCSNAKQPGGKPFVSVAVKKHIVSAWPNASRISWTIGDGIPYIGSRTEDKIGQPAIKPKCPSAALREAIELASRLTWVTQGSSNSDLLIKPFLEARVNLSVQEILQMTPSHYSGNIVHRYNDQYSPHSFMANRMSNSATRLIVSTNTLGE
FSGGGQSARDSNIIFQNVINYAVALFDIKFRNTEATDIQYNRAHLHLTK-CCTREVPAQYLTYTSTLDLDLTRYRENELIYDSNPLKGGLNCNISFDNPFFQGKRLNIIEDDLIRLPHLSGWELAKTIMQSI---
ISDSNNSSTDPISSGETRSFTTHFLTYPKIGLLYSFGAFVSYYLGNTILRTKKLTLDNFLYYLTTQIHNLPHRSLRILKPTFKHASVMSRLMSIDPHFSIYIGGAAGDRGLSDAARLFLRTSISSFLTFVKEWIINR-------------GTIVPLWIVY-----PLEGQNPTPVNNFLYQIVELLVH------
DSSRQQAFKTTISDHVHPHDNLVYTCKSTA-SNFFHASLAYWRSRHRNSNRKYLARDSSTGSSTNNSDGHIE-RSQEQTTRDPHDGTE------RNLVLQMSHEIKRTTIPQE----QGPSFQSFLSDSACGTANPKLNFDRSRHNVKF--HNSASKREGHQIISHRLVLPFFTLSQG-----
--------------TRQLTSSNESQTQDEISKYLRQLR----SVIDTTVYCRFTGIVSSMHYKLDEVLWE--IESFKSAVTLAEGEGAGALLLIQKYQVKTLFFNTLATESSIESEIVSGMTTPRMLLP---
VMSKFHNDQIEIILNNSASQITDITNPTWFKDQRARLPKQVEVITMDAETTENINRSKLYEAVYKLILHHID-PSVLKAVVLKVFLSDTEGMLWLNDNLAPFFATGYLIKPITSSARSSEWYLCLTNFLSTTRKMPHQ-----------NHLSKQVILTAL------
QLQIQRSPYWLSHLTQYADCELHLSYIRLGFPSLEKVLYHRYNLVDSKRGPLVSITQHLAHLRAEIRELTNDYNQQRQSRTQTYHFIRTAKGRITKLVNDYLKFFLIVQALKHNG-
TWQAEFKKLPELISVCNRFYHIRDCNCEERFLVQTLYLHRMQDSEVKLIERLTGLLSLFPDGLYRFD-
>Bombali_ebolavirus_YP_009513282
--------MASSHTQYPDARLSSPIVLDQCDLVTRACGIFSAYSLNPQLKSCRLPKHIYRLRYDPTVAQFLS-----DVPVATLPIDYILPVLLRAISEGEYCPLEPRCKQFQNEILDYTL----QDARFLEYYLRTTGSEEKEGNLTANPALKNVI--------
LKNEFLPQLYFWYDLAILARRGRLNRGNARSTWFVHDKLIDILGYGDYIFWKFPLSLLPMNENGIPHAAMD--
WYQRGLFQEAIQGYKYIVSVSTADVLIMCKDIVTCRFNTVLIAAVARIEDHVHIDYPDVSVISTLYRAGDYLLSILGSEGYKIIKFLEPSCLAKIQLCATYTESKGRFLTQMHLAVIHTLEELLNGRNLKTFQLQKIREFHQIIVRLRSTPQQFCELFSIQKHWGHPILHSERAIQKVKRHAT
IMKALRPIVIFETYCVFKYSVAKHYFDSQG-TWYSVTSDKALTPSLQSYIRRNQFPPLPMIKDLLWEFYHLDHPPLFSTKVISDLSIFIKDRATAVEQTCWDAVFEPNVLGYNPPYRFTTKRVPEQFLEQENFSIDSVLNFAQSLDYLLPQYRNFSFSLKEKELN-
IGRTFGKLPYLTRNVQTLCEALLADGLAKAFPSNMMVVTEREQKESLLHQASWHHTSDD-
FGEHATVRGSSFVTDLEKYNLAFRYEFTSPFIEYCNRCYGVRNVFNWMHFIIPQCYMHVSDYYNPPHGLSLENRETPPDGPSAYRGHLGGIEGLQQKLWTSISCSQISLVEIKTGFKLRSAVMGDNQCITVLSVFPLESDPDEQELHAEDNAARVAASLAKVTSACGIFLKPEETFV
HSGFIYFGKKQYLNGVQLPQSLKTAARMAPLSDAIFDDLQGTLASIGTAFERSISETRHILPCRIAAAFHTFFSVRILQHHHLGFNKGTDLGQLAISKPLDFGTITLALAVPQVLGGLSFLNPEKCFYRNLGDPVTSGLFQLKKYLQMLD-
MEDLFLPLIAKNPGNCSAIDFVLNPSGLNVPGSQDLTSFLRQIVRRTITLSAKNSLINTLFHASADFEDEMVCKWLLSSNPVMSRFAADIFSRTPSGKRLQILGYLEGTRTLLASKIINNDSDTPILDRLRKITIQRWGLWFSYLDQCDQALAEALDKIHCTVDLAQILREYSWAHLLEG
RRLIGATLPCMLEQFQVWWLKPYEQCPACNSLKDAGASPYVSAAIKQNIVSAWPNRARLSWTIGDGIPYIGSRTEDKIGQPAIKPRCPSAALREAIELASRLTWVTQGGANSDLLIKPFLEARVNLSVEEILQMTPSHYSGNIVHRYNDQYSPHSFMANRMSNSATRLIVSTNTL
GEFSGGGQSARDSNIIFQNVINYAVALFDIRFRNKDPSSIPFNRAHLHLTR-CCTREVPAQYLTYTTTLKLDLTRYKDNELIYDNDPLRGGLNCNLTFDHPLFKGERLNIIEEDLIRFPHLSGWELAKTILQSI---
MSDSSNSSTDPISSGETRSFTTHFLTYPIIGLLYSFGALTSFYLGGTIISTKKIKIEEFLYYLTTQIHNLPHRSLRVLKPTFKHASVIARLVDIDPHFSIYIGGSAGDRGLSDAARLFLRISIKTFIDFIQDWIIIR-------------KVATPLWIIY-----PLEGHSPHPINNYLNRVISLCEC------
QSISRKLPASNDQTGTMITQNLVYMCKSTA-SNFFHASLAYWRNRQKSQTKKRLNACATKS---HKASSQKGLTTDPFNLTTTKLHTNLTPCRWRLDMGSKDPVRSQRAIPTE----QMRSSLDNMSENTTHSTLPNNSLAESWIHPDF----
STAPDSGTRCHSRCLAVPFFAQASG-------------------RNSQNTAVSTQYISIISTLLSQIK----TMVDTTRYCRFTGIVSSMHYKLDEILCG--LGNFPTAITLAEGEGSGALLLIQKYGTRNLFFNTLATEHSIESEIVTGSTTPRMLLP---
IMERFYKDELHIILNNSASQITDITSSCWFTSQKNRLPHNVDIITMDAETIENIDRSRLYEAVKHVIVNHID-TAALRVVVLKVFLSDVKGMIWLNDNLAPYFGAGLLVKPITSNARSSEWYLCLTNFLTLHRVAPHQ-----------GHVSLQVICKAL------
HCQLRRNSYWLSHLMHYGSRDLHIEYINLGFPSLEKVLYHRYNLVDSGRGPLTSVMQHLVHLQSEIRELVSDYHQLRQSRTQTYHFIKTSKGRITKLVDDYLKYYLIIRALKHGK-
DWRHEFHQLPELVSVCKRFYHVKNCQCDDRFWVQTLYLSRMPESELKLLERVTGFLTLFPEGFQNPP-
>Reston_ebolavirus_NP_690587
--------MATQHTQYPDARLSSPIVLDQCDLVTRACGLYSSYSLNPQLRQCKLPKHIYRLKFDTIVSKFLS-----DTPVATLPIDYLVPILLRSLTGHGDRPLTPTCNQFLDEIINYTL----HDAAFLDYYLKATGAQDHLTNIATREKLKNEI--------
LNNDYVHQLFFWHDLSILARRGRLNRGNNRSTWFVHDEFIDILGYGDYIFWKIPLSLLPVTIDGVPHAATD--
WYQPTLFKESILGHSQILSVSTAEILIMCKDIITCRFNTSLIASIAKLEDVDVSDYPDPSDILKIYNAGDYVISILGSEGYKIIKYLEPLCLAKIQLCSKFTERKGRFLTQMHLSVINDLRELISNRRLKDYQQEKIRDFHKILLQLQLSPQQFCELFSVQKHWGHPILHSEKAIQKVKRHATILK

ALRPNVIFETYCVFKYNIAKHYFDSQG-TWYSVISDRNLTPGLNSFIKRNHFPSLPMIKDLLWEFYHLNHPPLFSTKVISDLSIFIKDRATAVEQTCWDAVFEPNVLGYNPPNKFSTKRVPEQFLEQEDFSIESVLNYAQELHYLLPQNRNFSFSLKEKELN-
IGRTFGKLPYLTRNVQTLCEALLADGLAKAFPSNMMVVTEREQKESLLHQASWHHTSDD-
FGENATVRGSSFVTDLEKYNLAFRYEFTAPFIEYCNHCYGVRNVFNWMHYLIPQCYMHVSDYYNPPHNVNLSNREYPPEGPSSYRGHLGGIEGLQQKLWTSISCAQISLVEIKTGFKLRSAVMGDNQCITVLSVFPLKTDPEEQEQSAEDNAARVAASLAKVTSACGIFLKPDETF
VHSGFIYFGKKQYLNGVQLPQSLKTAARMAPLSDAIFDDLQGTLASIGTAFERAISETRHILPCRIVAAFHTYFAVRILQYHHLGFNKGIDLGQLSLSKPLDYGTITLTLAVPQVLGGLSFLNPEKCFYRNFGDPVTSGLFQLRVYLEMVN-
MKDLFCPLISKNPGNCSAIDFVLNPSGLNVPGSQDLTSFLRQIVRRSITLTARNKLINTLFHASADLEDEMVCKWLLSSNPVMSRFAADIFSRTPSGKRLQILGYLEGTRTLLASKIINNNSETPVLDKLRKITLQRWNLWFSYLDHCDQLLADALQKISCTVDLAQILREYTWSHILEG
RSLIGATLPCMVEQFKVKWLGQYEPCPECLNKK--
GSNAYVSVAVKDQVVSAWPNTSRISWTIGSGVPYIGSRTEDKIGQPAIKPRCPSSALKEAIELASRLTWVTQGGSNSEQLIRPFLEARVNLSVSEVLQMTPSHYSGNIVHRYNDQYSPHSFMANRMSNTATRLIVSTNTLGEFSGGGQAARDSNIIFQNVINLAVALYDIRFRNTN
TSDIRHNRAHLHLTE-CCTKEVPAQYLTYTSALNLDLSRYRDNELIYDSNPLKGGLNCNLTIDSPLVKGPRLNMIEDDLLRFPHLSGWELAKTVVQSI---
ISDNSNSSTDPISSGETRSFTTHFLTYPQIGLLYSFGAVLCFYLGNTILWTKKLDYEQFLYYLHNQLHNLPHRALRVFKPTFKHASVMSRLMEIDSNFSIYIGGTSGDRGLSDAARLFLRTAIASFLQFLKSWIIDR-------------QKTIPLWIVY-----PLEGQQPESINEFLHKILGLLKQ---
--PKSIPKEVSIQNDGHLDLAENNYVYNSKSTA-SNFFHASLAYWRSRKSRKTQDH-----------NDFSRGDGTLTEP----VRKFSSNHQSDEKYYNVTCGKSPKPQERKDFS----NGQTMSNHRKKGKFHKWNPCKMLMESLTEGDY--FQNNTPPTDDVSSPHRLILPFFKLGNH---
-----------------NHAHDQDAQELMNQNIKQYLHQLR----SMLDTTIYCRFTGIVSSMHYKLDEVLLE--YNSFDSAITLAEGEGSGALLLLQKYSTRLLFLNTLATEHSIESEVVSGFSTPRMLLP---
IMQKVHEGQVTVILNNSASQITDITSSMWLSNQKYNLPCQVEIIMMDAETTENLNRSQLYRAVYNLILDHID-PQYLKVVVLKVFLSDIEGILWINDYLAPLFGAGYLIKPITSSARSSEWYLCLSNLISTNRRSAHQ-----------THKALGVIRDAL------
QAQVQRGVYWLSHIAQYATKNLHCEYIGLGFPSLEKVLYHRYNLVDTGLGPLSSVIRHLTNLQAEIRDLVLDYNLMRESRTQTYHFIKTAKGRITKLVNDFLKFSLIVQALKNNS-
SWYTELKKLPEVINVCNRFYHTHNCECQEKFFVQTLYLQRLRDAEIKLIERLTGLMRFYPEGLIYSNH
>Sudan_ebolavirus_YP_138527
M-------MATQHTQYPDARLSSPIVLDQCDLVTRACGLYSEYSLNPKLKTCRLPKHIYRLKYDTIVLRFIS-----DVPVATIPIDYIAPMLINVLADSKNVPLEPPCLSFLDEIVNYTV----QDAAFLNYYMNQIKTQEGVITDQLKQNIRRVI--------
HKNRYLSALFFWHDLAILTRRGRMNRGNVRSTWFVTNEVVDILGYGDYIFWKIPIALLPMNTANVPHASTD--
WYQPNIFKEAIQGHTHIISVSTAEVLIMCKDLVTSRFNTLLIAELARLEDPVSADYPLVDNIQSLYNAGDYLLSILGSEGYKIIKYLEPLCLAKIQLCSQYTERKGRFLTQMHLAVIQTLRELLLNRGLKKSQLSKIREFHQLLLRLRSTPQQLCELFSIQKHWGHPVLHSEKAIQKVKNHA
TVLKALRPIIIFETYCVFKYSVAKHFFDSQG-TWYSVISDRCLTPGLNSYIRRNQFPPLPMIKDLLWEFYHLDHPPLFSTKIISDLSIFIKDRATAVEQTCWDAVFEPNVLGYSPPYRFNTKRVPEQFLEQEDFSIESVLQYAQELRYLLPQNRNFSFSLKEKELN-
VGRTFGKLPYLTRNVQTLCEALLADGLAKAFPSNMMVVTEREQKESLLHQASWHHTSDD-
FGEHATVRGSSFVTDLEKYNLAFRYEFTAPFIKYCNQCYGVRNVFDWMHFLIPQCYMHVSDYYNPPHNVTLENREYPPEGPSAYRGHLGGIEGLQQKLWTSISCAQISLVEIKTGFKLRSAVMGDNQCITVLSVFPLESSPNEQERCAEDNAARVAASLAKVTSACGIFLKPDETF
VHSGFIYFGKKQYLNGIQLPQSLKTAARMAPLSDAIFDDLQGTLASIGTAFERSISETRHILPCRVAAAFHTYFSVRILQHHHLGFHKGSDLGQLAINKPLDFGTIALSLAVPQVLGGLSFLNPEKCLYRNLGDPVTSGLFQLKHYLSMVG-
MSDIFHALIAKSPGNCSAIDFVLNPGGLNVPGSQDLTSFLRQIVRRSITLSARNKLINTLFHASADLEDELVCKWLLSSTPVMSRFAADIFSRTPSGKRLQILGYLEGTRTLLASKMISNNAETPILERLRKITLQRWNLWFSYLDHCDPALMEAIQPIKCTVDIAQILREYSWAHILDG
RQLIGATLPCIPEQFQTTWLKPYEQCVECSSTNN--
SSPYVSVALKRNVVSAWPDASRLGWTIGDGIPYIGSRTEDKIGQPAIKPRCPSAALREAIELTSRLTWVTQGSANSDQLIRPFLEARVNLSVQEILQMTPSHYSGNIVHRYNDQYSPHSFMANRMSNTATRLMVSTNTLGEFSGGGQAARDSNIIFQNVINFAVALYDIRFRNTC
TSSIQYHRAHIHLTN-CCTREVPAQYLTYTTTLNLDLSKYRNNELIYDSDPLRGGLNCNLSIDSPLMKGPRLNIIEDDLIRLPHLSGWELAKTVLQSI---
ISDSSNSSTDPISSGETRSFTTHFLTYPKIGLLYSFGALISFYLGNTILCTKKIGLTEFLYYLQNQIHNLSHRSLRIFKPTFRHSSVMSRLMDIDPNFSIYIGGTAGDRGLSDAARLFLRIAISTFLSFVEEWVIFR-------------KANIPLWVIY-----PLEGQRSDPPGEFLNRVKSLIVG------
TEDDKNKGSILSRSGEKCSSNLVYNCKSTA-SNFFHASLAYWRGRHRPKKTIGATNATTA----PHIILPLGNSDRPPGLDLNRNNDTFIPTRIKQIV------------------QGDSRNDRTTTTRFPPKSRSTPTSATTHQGKLTHLDEDHNAKEFPSNPHRLVVPFFKLTKD-----------------
---GEYSIEPSPEESRSNIKGLLQHLR----TMVDTTIYCRFTGIVSSMHYKLDEVLWE--YNKFESAVTLAEGEGSGALLLIQKYGVKKLFLNTLATEHSIESEVISGYTTPRMLLP---
IMPKTHRGELEVILNNSASQITDITHRDWFSNQKNRIPNDADIITMDAETTENLDRSRLYEAVYTIICNHIN-PKTLKVVILKVFLSDLDGMCWINNYLAPMFGSGYLIKPITSSAKSSEWYLCLSNLLSTLRTTQHQ-----------TQANLHVVQCAL------
QQQVQRGSYWLSHLTKYTTSRLHNSYIAFGFPSLEKVLYHRYNLVDSRNGPLVSITRHLALLQTEIRELVTDYNQLRQSRTQTYHFIKTSKGRITKLVNDYLRFELVIRALKNNS-
TWHHELYLLPELIGVCHRFNHTRNCTCSERFLVQTLYLHRMSDAEIKLMDRLTSLVNMFPEGFRSSSV
>AKO69552_Bat_filovirus_unnamed
------------------------------------------------------------------------------------------------------------------------------------------------------------------------------------------------------------------------------------------------------------------
--------------------------------------------------------------------------------------------QKTWGHPVLHNELAIGKVKDHATKLKLLRPNMIFYTYCVFKYMVAKHFFETQG-
TWYPMTHDRALTPTLNTYIKRNMFPSLKNIQFVLWEFYHIRHLPIYSTKIVHDLSIFIKDRATAVQKPYWDAVFESNVLGYYPPNRFESKRVPEQFLNQDNFSIDSVLTYAENLEYLLPENRNFSFSLKEKELS-
IGRTFGKLPYLTRNVQTLCEALLADGLAKAFPSNMMVVTEREQKESLLHQASWHYNRTS-
TSENVTVRGSSFVTDLEKYNLAFRYEFTRPFIDYCNECYGVKNLFNWMHFLMPQCYIHVSDFYNPPHCLTSQNREHPPPCQNAYYGHLGGVEGLQQKLWTSISCAQIILTELRTGFKLRSAVMGDNQCITVLSLLPVGMGQSEQEVISETNAAIVATDLAYITSACGIFLKPEETFI
HSGFIYFGKKQYLNGIQLPQSIKTASRIAPLSDAIFDDLQGSLSSIGTAFEKAIAETRHLIGIRAISSFHSYLAVRLLQYHHLGFKKDTDIGLLALKKPLDYSTIVVTLSIPQVLGGLSFLNPEKCFYRNIGDPVTSGLFQLKMYLKMRQ-
MMDLYPLLISKTPGNSTALDLVLNPLGLNVPGSQDITSFLRKLVRKSITLHSKNKLINSLFHVGADSEDETICQWLLSSEPVMSRFAADIFSRTPSGKRLQILGYLEGTRTLLASKVINSDPEISVIKQLQNITIKRWAIWFNLHADIDPDLLEVVKRINCSLDLAQLLREYTWAHILKEKK
LIGATLPCLLEQFKICWMTTTSTNQVCHGCID--QSPYISGLLLNNIVCARPNDQRLQWTIGSSNPYIGSRTEDKIGQAPLRPRCPSSALREAIELASRLTWATQGSGTNADLIKPFVKARVNLEVEDIQKMLPVHYSGNI----------------------------------------------------------------
------------------------------------------------------------------------------------------------------------------------------------------------------------------------------------------------------------------------------------------------------------------
------------------------------------------------------------------------------------------------------------------------------------------------------------------------------------------------------------------------------------------------------------------
------------------------------------------------------------------------------------------------------------------------------------------------------------------------------------------------------------------------------------------------------------------
-----------------------------------------------------
>Dianlovirus_menglaense_YP_010087189
--------MNSHPTQYPDARLSSPITLDHCDLLSRSCGFYSSYSHNPKIKNCRIPSHIYRLRNSVQLKAFLS-----NCSVLTLPFSSIWETLLNQFKIHSKNYNDVLNYLKIDEATSYAN----WDQEFLQSYLKTAISDPDYLNSPKFPTDPDFAK-------
RNNEYYHALILVHHLTQLARRVKSQRGSTRNHWELTSDNFSLFGIGDFFLVKIKLPLLLKAYKSLQDAIPNRLWYGEQNLLEYYQTDEYLVSFASYESFIMLKDVIIERFNTIEIINRAKIEDQAYQDYPDLLTVLNLYKAGDNIISLSGELGFKLIKFLEPLCASCIQTVGIFTPRKYWFQS
QMIKSYNETVIEISNELKLTNSQMDDLYKFSRIIIQARYTPQQYCELFSLQKHWGHPCLHNDTALKKVRQHATSVKILKPKILFETFCTFKFIIAKHHFLAQG-
GWYKVTHNLDLTPYLKKHILMNSFPNQSEIYEHLWEWYHVKHEALYSTQIISDLSIFIKDRATAVPRECWDSVFDNSVLGYTPPLRFQTKRVPEQFLSQPDFTLSRVIEFAERMEYLDDANRNFSFSLKEKELN-
VGRTFGKLVYSVRNVQTLCESLLADGLAKAFPSNMMVVTEREQKESLLHQASWHRNSAS-
IGDNAIVRGASFVTDLEKYNLAFRYEFTKPFIDYCNQCYGVKNLFDWMHFLIPLCYMHVSDYYSPPYGLNAENRERPPECENGYYYHLGGIEGFQQKLWTCISCAQITLVELRTRLKLKSCVMGDNQCITTLSLFQMDTTPENQEKIAELNAARVAVELANITGFSGIFLKPEETFV
HSGFIYFGKKQYLNGIQLPQSLKTMARCGPLSDSIFDDLQGSLASIGTAFERGISETRHIFPSRWVAAFHSFTAVNLLNSTHLGFPANTSLDIVSLGKPLTYNEKVLALITPQVLGGLSFLNPEKIFYRNISDPLTSGLYQLRHSLMMLG-
QDSLFTILIAKIPGQADESDFVMNPLGLNVPGSRDITTYLRQIVRQNITLTSQNRVINSLFHIGADIEDKEVCKWLLSSIPVMSRFAADIFSRTPSGKRLQVLGYLEGTRTLLASKTLNLSADGTMIMKLRDLTKKRWRHWFSYLDNFDEDLSDSLDTLHCTVDIADLLRAYSWDDVL
QGKKLIGATLPCILEQFTAHWEKYDQTLYNYAAKENVKMEEYISCALDKTIVQAHPNADRLSWTIGNHAPYIGSRTEDKIGHPPYKVNCPSAAFKETVEIVSRMLWVTQGLNGKEDFIEPFVKARINLPLETLRNFLPTHYSGNIVHRYNDQYGQHSFMANRMSNSSTRMIISTN
TLGKFAGGGQAAIDSNIIFQNAINCGVAIMDIRYATDSAPSEPVKKMRLYLNQ-CCTRNVPAQYLSFSKPLSVDLSKYKENELIYDPDPLSSGIKGRTKELNPAALSLDLNRSDINSYAFITTSSWTLANTIMTSI---
IYDQSSGSTDPISTGVTKSFVTQFLIFPLDGLFYSFGAWLLQLIVPFNDLRKYQDKNGLFFQLRSSVQNFSHRSLRILQGTFKHELVIQRLAKPLPLITLLLGGAAGEKGVSDSVRLFLLASLHYFTDNFIGIINNN-------------QKDMPVWMYF-----PSEGQQLLPIVKYLTRLYLILTG-
-----YSLRKRPNK--ELIDSICNKLWIYPARSTQ-SNHYYASANFWRNKKHDVNQA---RKQNRYLSDRRVINHSEGAIDQTSSTTGGVTTDIIIPDQSNPENINSSESHKERLTTV----QTQYITSNTRLDKVKSSSINSISVKSTSDSYFNKGISSL-RERNFNEQHPSLRITKSVLGKNY--
ILIVQCSNISSCLHFKSKKQKANDQVVLNDFLHQFKQSLQ----RAPESNDYCRFTGVVSSMHYKLSDILPK---RKFNSIVCLAEGEGGGARYLLMTQDVKFLFFNTLASTTQIETEILSGRVIPRMLYGTPYLNERL--
DSGQIILNNLSSQITDITDPRWIGLVKSLLPKDIDLLTMDAETPGNFSRVPLYEVILN-LIQLYL-GHLPKVLILKIFLFDVPGTKFIISRLINYYMKIDLKKPFSSSARTSEWYLVAELPREQGVTIIIP-----------DEALDGYLITAM------
RRQIIAIPYWFSYIEKIYTRDLHLSYFKLGFPLIENAFYHRYYIPKYKDLTIYSIKRYLTDGRNYIRKLINELQSGEALTELKQHFILSSKGRVNKVYNDLIKIDLILVAVETKK-TWSQLKQQMPDFQIICKNHLIEKCCTNADQHLVKLDFLKNTTVAEEKFLNRIVGLIIMFPSGIN----
>Tapajos_virus_YP_010798912
--------MDPSEAQFPDGRLNSPIVLDAVDLFSRHLSLQSCYSKNPRLSLCEIPSHIKRQAALLYNNDLLS-----YTYPLTLPCTNLLPTIIEEFKYDESLQGDNTVKAWLYEATQYASEEVHYDHRWIQEHVNEYSTQLGITEPSLHNTDVNMVP-------
DIQRISTAFSVALHCVTGARKCAKNRSNTRCTFKHNTPTLSLCCTGDFVVFRFKFTGYTIQSPCFQAGWFS--WYPSLEPTEMISKWKYFCTIVSYDWLLMLKDIIMVRYNALVLSFLGERQDNL-
TDYPSIEAIINLFEQGDKLLEVFGNAGYTLLKYFEPLILGRIQACGEESARRMNFLYQMIYAYQAVITCISKSKQLTSLLGERKNNFLSCLLTQPLRPQQLSELFSLQKAWGHPFISSHDAISKVRVHGTTRKALHVKTVQKVFCVFKYQVAKEYFRVNK-
RWYMLLNSPTLTPHICSFARRNIFPTLSQMYPFLEEWYHVEHSAIYNTRALQDLSVFIKDRATATGWNEWDSVFSPAVLGYQPPRRSYSKRVPEQFLDQSDFSLQSVISYGENLEYLTEQQVNFSYSLKEKELS-
VGRLFGKLPYKTRNVQTLCEALLADGIAKAFPNNMMVVTEREQKQTLLSQARLHHCTAV-
EGEQVEVRGANFVADLEKYNLAFRWEFTKHFIEYCNMCYGTVNVFEWMSFLIPSCYMHVSSYYDPPYSLCLQNRLRPNSCENAYRYHCGGIEGLQQKMWTSISCAQILLVEVESGLKVKSALQGDNQCITSLTIFPKGTSAREQTTVSENNAMAIAAHLCRITAACGIFLKPEETF
VHSGFIYFGKKIYLQGIVLPQSLKTLVRMGPLAQSIYDDAQGALASIGTCAEKAYSETRHILPVRWIAHTHTWLSLRMTDAFHLGCSSLGTLTEQAIGTPLTNETLRLLTLVPNTLGGLNFLLLEKVFYRNLPDPVTSSLYQLQWRLAAIG-
KIELYPCFASPKRGHAGALDLVLNPLGLNIPGSQEISMFLRRRVRSAITLRARNKLIRTLFNPCSMSEDDDLAEWLLSSTPVMPRFAADLYERTPSGKRLKILGYLEGTKTIVASSLFSDDDEESIVAELNRLIISRWKTWLMCGCQYEEFLLDVVQKIKCTEELAQFLRLYTWSHVVGD
RGMVGSTAPCLLEQFKVYNNPANGSCIGCVNEHLSNISGHATARLSSRVSAGLPSRQRLCWTIGSYFPYLGSTTGEKIGDALFIPAMPSAPLRDAIEIYSRAGWVTQSGCGASKLVANLISSRVSLDPELIKLLSPRHHSQNILHRYHDHYSQRSFMANRISNAATRVLVSTNQMG
KFSSSSEAATDSNIIFQNVINFAVAVIDIRNRFKDLYLSIENGVHIHLTG-CCTRVLEAGKLFFQNPAPKMKINYELNELVYDEDPLKTGFTRSFGANNKHFTSQDFSDMSGEVQKFALTAPYDLTEIIIQRM---INQDQDGETP-----
EGQNYITEFLAYDFDFILYCLGYALLVGLWRVHTLRPPSTEIHIITYIGSLLSAAPYSAYSVLFVTLRHPSVYERICTMLPSISHLYGGTSTTRALADLTKKLYTEGLTRFLQVLTLCLKNP-------------ITTNPLIILF-----PAVGQTLSTWSSIMSILLPTITT------MNSAKLPEN---
ITRLLFIERIPYAVWPTSLIDFRYSALDCIREPVSVSQ-----SSNSDHHLIKRLFRQMSSPISMQKLADPNIDN-------------EGRETGNTILKEKRGIERASRNGQPEE----------------------------PTTPSLKKGTLILKIKELDQL-----------------------HPAPPVADSHSVKNLLHTVA---
-EIPDMTHFVRMTGIVSSMHYKLDEVLDP--WDHYEIAVCLAEGEGAGLRYLIRQYGINRFFFNTLAHDHQIINEVAGVALIPRMLRG---VYPRNDDLLRNGVFNNRTDHVTDIGSDYWLTQWSEHLPNQYDIITMDAESVDNSERIPLYRTAIT-LFTHKK-
AAQRCTLVVKGYLADVKGLNFLLGSLFPISTSLYLIKPISSNFGNTEWYIKAVRTHTQSSYLWNI-----TRFTFLDETL-RYCKAAL------
MMQTSEARYQRDHLVKLFNTGLHERYLTMGFKTISDGFRLRYSIERIAPNIREDIVLTTLSIKSQILKMMREDQRNKTTRFQSTHFTKGFRGQISHLVAVYCKYLLLLQGGENG--FYPPIEQIWP--
IWVCEHGVFRRSPHCRNHLLVSKDFLIVNSLAERKVLNRMSAWLYLTRKSMYTSMM
>Wenling_frogfish_filovirus_YP_010085045
MD---DYTVPKHELQFMEGRLCSPITGDSIDLVARSVGKLSNVSRNPALQTCRIPCHIYRLSRYDEVTRFLN-----GTSSRFYPFHRLFPDLLTE----ESSMSDSISVAVMQEGLDWADEEVKHSAKWLQDHTDRLLGQP-----EMTPKMSQCI--------
KESGYFIPYMCWISIINEVRRCQKHREDQGRFFSTCLGRYQLLVAGEFFFIKWTL--TPEDITAILGGTYHEELPQGENPTTFA--GTCALVAGVLDELLMVKDVVVARLNATVAARAAHQEDDL-
PDHPNPDSLQLVFEKGDDLLRLLGNEAYPIIKMFEPLMLGRAQLFGVMTERKSWFLKTMIRATNTLISGVQGSK-----EKTAGRAWLTAILSLGCRPQQMCEAFSVQKCWGHPILHAEDAIKKVQKHATVRKVISINEVLEVFCVFKYQILKHHFDTVG-
GWPTPLHETTLTPGINRFMKAHQYPTLQQAESFLYEYVHFHHGQIYSSDIAHDLRLFVKDKATACEKPYWDSSFDRNILGYDPPRQPTPRRVPEAFLAQEDFSPQEIIEYAENLGYLDKDQINFSFSLKEKELG-
IGRCFGKLPYKTRNLQTLCEGLLSGGIAHAFPNNMMVVTERELKEALLSQGHSHRNATA-
HDDNTVIRGATFVADLEKYNLAFRYELTRPFIEYCNTCYNTPNLFNWMHLLIPLCYMHVSDRFNPPHGLNETNRDNPPIGPSAYRQHAGGIEGLQQKLWTSISCSQILLTELRTGFKIESAVQGDNQSVTTKTVFTKSEPPEVQENIAEDTAAKVAAALAKSTNRIGIRLKPEETFV
HSGFIYFGKKQYLNGVLLPQSNKVFLRCGPFADSLYDDLQGMLASIGTCVERAMSETRHVLGIRGLGAFHTWVSVRNLDRHHLGVEPTQSLTVLALGKELTNSAIKRALRTPQVLGGLGFLLREKMFYRNLPDPVSSALYQMKWAAETLD-
QYEEYLAALSVAPGSATELELCSSPLSLNIDSSQDITMYLRRLVRDGITSTARNKIIKTLFHAQSADEDTAIAKFLVSSTPVFPRFASSMMELTPSGKRLSILGYLEGTKTLVSSNITRSNHAESLIAKLNDITLDRWSCWFNTDKGLLQAIAAELITERCTMVLAQTLRLYTWKDLLEGR
NMIGSTLPCMLEQFTIIQQTEGDTCSSCHDPPPDVHQFFAIGCFNHSAIVSRPDDRRIHWTLGPHLPYIGSRTEDKIGDPPVKLKRSTPPMKEAAKLMSHFQWVTGSKTEGAKYATPFVGARVALEPEVVELLAPTHFSGNICHRYADHELSKAFMANRMPNVASHAICSTNHL
GPYSGGSKEAIDTNIIFQSLINYMVAVFDLSNQNTRTEKGKRVFIHAHPRK-CCTKEVPSEYITFSGEPITLPTLYQDNIFVYDPDPLAAGFSR-WDDLPVADLLTHINQAKEDLKRTPELFAWSFAEEILKET---LLHDLSGLAN--GSKQSKGFITEFLSFPVTGILYCFGCQMLIWLSR-
LLTTRDLDECDLTYTVFTLLKSMHHGAYKTLFDTFVHPQVQERFFPVVGRVIYDVEGQAGLAARANVVRMFLMACLRHAATTITDTKD---------------RQTKVVWVGF-----PLEGETEDICREHMDRSYNIATR-----IIESDRLTAT--ANRDNFVVRSSIYLCAVTK-
TLYRFQTLSWAREIRAASHTIPKS-------------------------------------------QQPPAPKKRDKPD------------------------------------------YLGPQAATLGLRVPVMRASSG----------------------FSKDVAGLLPCNVQGLCRMIR----SASSSNHFCRWTGKHSSMHYKLDELLEG-
-KRTYQQVATLAEGEGSGARLLILKYGAHSVLFNTLANPNQETTEVHGGRVRPRMLVD---IWDRA-----NIELNNSLLQNTDLSKASWCTHAKSLKGLPVSLITMDADVKEAETSLAIADTIEQ-CLLSGF-
LSRGGDLVLKVYLDMIHVVHRYLCVALTYFGHVELRKPISSDRRSVEWYIVCQHLTDKIPTAHA------GCETWYDDTE-ALIISCM------
QRQVTHIAKWLGVLKSQEFTARHKEYHGLGFEWVSSSLFRRLGLVGDVDGWAGVLSQSIDMFSQQILRQLAAMRSARTSLNKERHFVASPLGRLSHLSYEHWKRQLLLGVTEVGE-
FTRRVMDAKNHEVYTCQHLVFTYRPHCADSVRLSSTFVSRPDIAFLKLAERFVAMTIAR---------
>Monodelphis{AAFR03010417}72745-75762
------------------------------------------------------------------------------------------------------------------------------------------------------------------------------------------------------------------------------------------------------------------
------------------------------------------------------------------------------------------------------------------------------------------------------------------------------------------------------------------------------------------------------------------
-------------------KLSLHNQTVMYHYKAC-KGESVEIRGTSFITNQEKHNLAFLYKFTCQFIEYANKCHGIINLFN-MHFLIPKHYMHINDYYNPLH---QDNCSYPPEGSSAFGDQMERKEGL-
HKLYTSINCAQITLVEVEIGIRLGSMVIKDNQHIIVLCLFPLETSPKEQEHFSEYHVSVIESWLVYITLSHGIYL--------------
KKQYLNGVLLPQSLKTISKGAPLADTIFDDLQSTLAATGISFEQDVSKTRHVVPLCAVVAYQSFMLVRINQHYHLGFPKDVDLALYILGKPLLYEELVLTLSLFVI-----FLILKKSCTNNMVDPITSSIFQLQEVL-YLN-QPDLFLLLACKTPTDAGTIDLILNPLDLNI-GSQDLISFL-
RVIQDAIISMSFNKLINFFFHSQSVLGNHEIFE-VFFSDAVMSRFTVEIFSKTLSGKWMKILSYLEGTYTLFASQALLSISDEGISSNLCNLTKNWWQA-FNYLDQIDLSLLNICKDIKSLADSAQLLRQYGW----------------------------------------------------------------
KLSRSTRLLG-----------------------------------------QSFSNF---------------DSHSSGNLAHRYNHD-SPQSFMANGMSNSSTHCYVSTNTLGKYAVGSCVKINSN-IFSNAINLCLATLDI------------------------------------------------------------------------------------------------
------------------------------------------------------------------------------------------------------------------------------------------------------------------------------------------------------------------------------------------------------------------
------------------------------------------------------------------------------------------------------------------------------------------------------------------------------------------------------------------------------------------------------------------
---------------------------------------------------------------------------------------------------------------------------------------------------------------------------------------------------------------------------------------

```
>Gracilinanus_agilis
------------------------------------------------------------------------------------------------------------------------------------------------------------------------------------------------------------------------------------------------------------------------------------------------------------------------------------------
-----------------------------------------------------------------------------------------------------------------------------------------------------------------------------------------------------------------------------------FDRYVLYSVPLKRFTSKRVPE-FLVQ-
DFSVDQIPEFAKNLGYLEPKFYNFSFSLKEKELN-YGRPISKLV--KEHIDSFCESICSN----------------REKETLINQTVMHHYKAC-EGESMEIRGTSFITNLEKYNLAFLYEFTCQFIEYANKCHTIKNLFN-MHFLIPNPYMYSSDFCNPLH----DNCSFPPEGRHAYGGHMERNKGL-
YKLWTSITCAQITLVEVEIGVR-GSAVIKYNQYITVVCLFPLGTSPKEQEHFSAYDVSIIAGWLVYITSSHGIYLKPE-------
LIDFGKKQYLSRVLLPQSLKTISKGTPLVDTIFDDLQSTLAATDISFEQVVSETSHMIPLHAVAAYYSFMLVRINQHYHLGFPEDVDLTLFVLEKPLLYEEVVLTLSIPLVLGGLNFLNPEKILYH-------------------------------CKTPSNAGTIDLGLNPLDLNI-GSQ-LLQF--
DMIQDAITIMSSNKLINSLFHPQYDLENHEIFE-LFFSDAVMSRFAVEIFSSRLSGKWMKILGYLEGTYTLFASQALNSRSGEGISSNLCNLAKHWWDG-FNYF-YIDLSLLNVCKDIKCFTDFAQLLRQYGVHPKLG--------------------------CPICKNN---
YCLANYGEGPITLVTTYHSIKDRLSWTLGIHHVYLG-LTADKIGYSPIKPRCLTICLKSG-----KIRLPVPMEAKTLKSYRTFVKARVNLAME-LEIMIPKHSSGNLVHRYNYQYGPQSFMANGMSNSSTHCYVSTNTLSKYAVGSHVAIGSN-
IFQNAINLCLATLAIKLELIRFNDHGTEFIYSELFR-FCFANL--------------------------MKPLSSDASSS----------------EWHIFKSPYQAHQDLRIKSCYSL---YSSKGQPNRGPFC-----NFITIYISMATHLQYQSFGFVLVAEVFR-------------------------------------------------------------
---------------------------------------------------------------------------------------------------------------------------------------------------------------------------------------------------------------NRYG--------------------------------------------------------------------------------------------------
-----------------------------------------------------------------------------------------------------------------------------------------------PTITRNLGNDLESSF---------------------EKARASTLSQMYL-------YMQNVRLQYYLDILFIHP-------------------------------------------------------------
---------------------------F--------------------------------------------------------------GKVTKLI----------------------------------------------------------------------------------------
>Oberland_virus_YP_010800200
MMSQGSGGTPRSHIIFPDPHLNSPIVPEFRDSILIHCGLHQHVTLSPKVKGCSNSKRSERVS--KELRRVLEV-NRQEPCDSFLPPQKTYGFVYQH-----QTDLDTPSLRRFRDSLNVSSKHMNRQLADLDPILSHLYRGA----
PESLKRVLEKTDRLKGSVQSHDEYLRRWSTIQALVDRLRLDKKEKSRS---------------------------CPVEATVLNTPD--LQFFLLEECVMVISPKSSWWTLYCYEEVLMLADVLISRANAILYLHLGGLECPT-
SRLPDTQKVVELYTFIDGLFEQWGQVVFSLVKMIEPLCVGLLLLHSPAVDYRAHFYIEQYSSLLDTLMEKGVPG---
LQAKEMGRRFIQLLVQAGPTPNHLSELHGLFRHWGHNTILIPDACASLMKHACHQKVLKLDNITRIHCTWVYLFFKHHYTKHSNTWPPCDIKPYLSPNLYRHRMQSTFPTGKEQLSYIYEWSFVEPGILYSIDCIEDPAIFIKDKSISPNRSHWDSVFSPDMLGYSPPFNTESRRLA
DRFINDPDFSPYRVIEYIESGEVYQNEEFCINLSMKERELS-
TGRAFGKLPYLEREAQALGEYLLSEGPAKAFESNTKSLDEIKLMKELAEQSSLHSDEGDITVTRGTPLSATMVTDYMKFCANFRHETCLLIERFCAKTYGLRDFFSWHHRRISDGFIYVADYHHPPSEVTQHNRSAPPPSDCLWYNHLGGIEGYKQTVWTMFCCALTHLVSLELDI
PYTAMIQGDNQIISAVVRAKEGELLEDLRIRAVGTVQTFGNRMSQIALGAGLTLKPEETFIHTACMVYSKRILHHGVDLTQATKIVCRAMPNSSAVFDDPDLQLSSISSCGSRSVARGVPIITPYLVAEHQSALAYRLSEHFHIGVPKGLARELQT--
TVSSEEELYLILNVGTALGGPSANSLLRYLVRHEPDPVTTGFYWLMLKAEANPIFFLVASGVMNKVTPSRDLLSLVKDSRAISLPVCRTVSSAVKAAVQEGLTTGAVNPMLRGLFHEHTDEEDLAYGIFLLSTTPCIPLVSSELYHSSTPGVREKLIGYVDSTKTSLASSLTRGF-
GGSMHETLQGILLLRYRYQRELASTLPQAA--NPMLRNCTLAIADHLRVTSWRHTLDAGPLEGMSEPCVFEQTTLTEVRS-ENCTLCECGRQEHLLFRVAER---------
DGQEPICFKRGLLPPYVGSETSEKLAPPLLKLRSMSAPLKSAVRICSILKWVCEPDPTCDEALRWLSSQRVNAEPEDLELLTPVSTRSNVLHRLKDSFGKHNFATNRPLNVSSQVHISTNKMGKYSGSG-GGTDCNLIFQNAKHAGLSLIDLAWG--DSAAIPIVNFHLHANG-
CCVRETPAVFIKTFNTKPPPIERIEGNPMVYDDSPLGDGGAEDMMSAEVIRRKKKGSPNS--
HEGITETCTWALSQHYYSRFKMWEEKSKGADRQMMITEEAGMNVTECIHLGCGRVFKLFCLQLLMDKVQHFILPLLIDGGGASEIMRSYWTVLPQDTLRNFDVLLTMGPIRSEVVDYCKQKGYQFRRDSTRADFSELMRFYLMYTCEDILNPDDPFLSSMTHVHVFRKEKMF
DWANSLLQMICLSDIRRYAPEVAPEDWLSLIIALQRGPSHGRSSLLAVIDEMSRGSKRCLTMKVGSLLSARVQLLNVSKWECDPAAWAREGVRDLKD--------------------------------------------------RFKEEKNDQETP----I---------------------------------------
KITMRINAGDLAHPQHALCKHGPGGSIVLNCSTINN-----------------LISISETSVLPTPKHTQSLREVFVGRTTGIATSAHYKLLEMLKYVNIPAGSNVLCLAEGSGGFAKTLLQTGKVHKLYFNTLVQWGKATPHYTGESIGSELRDSS-----------
PDSVRMLTREINDLRDQRILDVFSGC-KTRFEVITCDAEY-YIKNPSKDANLPKIVALVAAEHLTKEGCLIWKAFCDDNPTMRFCEWFLTSLFEEVRVVKPIFSSPDSSEVYWVCQKKKTCVSDLNALVKSTLDEPRYCRDTLSSSFTNDLRVIRTEAKRL-----
HRVIGDYTLRTELHRSLCYLGWEENFNQSRCALGLTNPFDGVDNDMDAMLD-------------------------------------------------------------------------------------------------------------------------------
>Wenling_thamnaconus_septentrionalis_filovirus_YP_010085051
--MESTGGVSRSQIMFPDPHLDSPILPESRDAILIICGHSDLVTKNSKVLRVGRHPRFSRVA--SKVSDILAIISTESTLLTGVKPQHMYGRIFQM-----KTGSTSKSLRRYQDAVKCAYTVEGLQLDDLQGVLSEWFRHT----
PSLQQGVMEKLKRLKESSKHETEILTRWTIWHAMVEHMRLTKKNRRRS---------------------------DPVEPVLLRTEQ--IDLYEIEEFCLVHDKQSLTWCLLNTEECLMFSDTLISRFNSIRYLRMTLAESPD-
ALVPTPEDVLSLYRMIDSWFRRLGQVTYTLVKFLEPICVGLTLMDSEIIHQRAHFLTEQVCALRDKLGSEGLSK---
SDAEFFTRSFVNKLASITGNPNKASEIHGCFRHWGHNTIIVGPAVEAIKKHACQIKALRLGTLTRVHCVWVMLVMKHHYHSHNRSWAYAKIDKYLAPHLDHHRRMSTFPSDDESLAYLHEWSHVTPGVLYSLDSIEDPRVFVKDKSVAPDKSHWDCVFDQTLLGYWPPFYNK
SRRLADCFVQDEEFSPYDLIDYVESGEVFVDPEFNINLSMKERELS-
AGRPFGKLTYKARQCQALGEYLLAEGPGHAFESNTMCHSELELMKEMATQSQVQTTSEDFIPRAGEPCGATMVTDYAKFCANFRHETCRQVALTAGKLFGLKGFFAWQHQRIASSRIYVADYHHPPTRTTEQNRLDPPDDETVWHGHTGGIEGFQQKLWTMFCCALLHLA
SMETKVKYSAMIQGDNQIITGILAATGSETRDELRIRAVDEVKRLGHRMMLLAKQIGLTLKPEETFVAGSCTMYSKRLLLHGNDLTGALKIICRSMPNSSAVFDDPDAQASSISSCGSRSVGRGVSILTSYLVAEFHVALSYKLTELYHLGIPSGLSKAAGL--
PPLNNYELYLLVNIGTALGGFSSNSLMRYLVRHEPDPVSGAIHWLFLKGRTNPDHLRTLTKVLRSVTPSKDILSLIKDPRAVSLPTCRTISSLVKHAVKEGLRETAKNPILSQLFHENADEDDLALAVFLLATEPCIPVFSSEIFSNSIAGVRDKISGFVDSTKTTLASALTRGY-
AGGMQLNIVNILIFRYRFQVSLVRELPEKL--DKRVLKCSLDIAHDLRKQSWGAILKAGTLKGDSHPTVFEQFTLSEAGP-EGCELCEGGSDVHLDMRVVPS---------
RHPGPMRYKRGGLVPYIGSTTSEKLAPPIVRVDMMSGPLKSCVRTASILAWVCEDNDRTKVAIDWAIKQRCEVESEDIKLLIPSSTNSNVHHRLHDNFGKNNFTSNRPLNFSSQVHLSTNTMGIYSGEG-GGQDGNMIFQNVKHLGLSLLDLGCG--DMTTPQSYNFHLHGIG-
CCVRTMDADFVSSFNTEPPDIDLAKGNPLIYDPNPLGGSNLEKIRMVNTNQHTKDMLKTD--
PTSRAQSATQALAQHLLSRLHVALKRTTDRGTVYLPQEDGVGNISEMIQLGPLKVFKEVGLQLLVEIFQHSIRDSILQGLGLDGGFTGFVASLPSGSFSVFNNMLTVRKIREPLRDYALKRGFSYNRDSTAHAFNELVRFCIIHLCTDTLKDPNELKNSSLVRHVFRSDRYFDWAFTL
VVWSAVRDLNQTIPNLHYRQWVDLLNRIQSSPSHGKSAVVSVAARIVSEGGFDLTCLVE--LRRHLELTNVHVWECDPAAWAREAVKQEK-----------------------------------------------------GGERTDDVACK----I--------------------------------------
IVQKLCHLGDLDSAIKTLQRLGTSRVECVHGEPLSR-----------------STNATTVFPEPTVRCTTSLREIFSIRRTGISSSAHYKILELLPLMMNKRPTRACCVAEGAGGFAKTLCSILSIEKVHYSTLVDWGKATPHYTANDVGSELRVTS------------
DPKIENLVRGPNDLRDPTTVTFLTKK-CKQVDILTCDAEM-FFKDLSRDSNLYKNIMILAANILSSTGTLIMKFFLGDTPVSCFLISAANRMFHEVFLVKPFYSSPDSSEVYLVAQSMKPEKVKVVEELDQYMDLPKF------
PIWHDDLVPVVKRRERLIDGQKIRAWEKQDLTLKLDAQLVSMGWTSNRNQILSQLGVRTDVADLHECATELLSMATGVMTGLFGSFSER-----VGRGEQKTLRRALLYLGGILL---------------TRTQLRQFLPIVVDNGRGTTPCVNVRSCDGVTY-
IDLPYLMGTGQLKTLQKLSGHF------------
>Fiwi_virus_YP_010800193
--MEGGCSLPRSSIMFPDPHLNSPIVPEARDCVFILAGYSHLISKNMKVRLVSSHPRFSRVA--GKLSEVLSLAQLEDEMLQGKKPQQIYGTIFQM-----DPTSDSKSLNRYRDAYKIARGVERRQTHDLVGVLSEWYKGA----
PSLLECMLEKVDRLLGSQKDESALMWRWAVTNAVVEHMRLTKKNRKRA---------------------------DPIQTYMLRTIW--VDFFEIEEFCCVIDRKSGVWSLLSTEEMLMFTDTLISRFNTAKYLRLSHAENKN-
AQLPNVESVMGLYELVDGWFRTLGGLTYTIIKFLEPLCVGLTLMDSPLIDQRSHFLTEQACSLVDKLADTGMSR---
AEAVMYMQSFLGALAQAAPGPNHASELHGCFRHWGHNTIIIEEAVVAIKKHAQQVKILKLDTMTRIHCTWTMLLIRHHYQTHSRSWPYVEIDRVLAPNLCHHRKMSTFPTDEQSLGHIYEWAHVRPGVLYSLDSIEDPRVFVKDKSVAPDRPFWDCAFDSVLLGYQPPFYTKS
RRLADRFVTDDEFSPYELIRFVESGEIYSDESFNINLSMKERELS-
AGRPFGKLTYKARQCQALGEYLLAEGPGKAFDSNTMCHTELQLMKEMASQSHVQTTSEEQVPSQGESCGATMVTDYSKFCANFRHETCRSVSKQCAELFGLSDFFSWQHQRIASSMIYVADYHHPPTGNNSSTREHPREDETVWYNHTGGIEGYQQKLWTMYCCALLHLA
AMETGVKYSAMIQGDNQIITGIITATGRETRLELRERAVREVKSLGHRMMVLAKQIGLTLKPEETFVAGSCTMYSKRLLLHGSDLTGALKIICRSMPNSSAVFDDPDAQASSISSCGSRSVGRGIGVLASYLLAEFHVSLALRLTELYHLGLPEGLARGSGV--
EPLTEYEMYLLCNIGTGIGGSSANSLMRYLVRHEPDPITGAIHWLYLKGKLDPNHLRTLTKIIQSVTPSRDILSLIKDPHSVSLPTCRTVASLVKSAVKDGLRDMAKNPVLKGLFHDDCDKDDLALTVFLLSTTPCIPVFSSELYQNSVCGVRDKLTGFVDSSKTTLASALTRGY-
SGGMQVNIVRVLVLRHQFQVSLVRDLPEKV--SPLVLKCSLDVAHFLREKSWGEALKAGELKGDSHPTVFEQFSLYDAGG-VGCEACERGDNTYLDLRVVES---------
RHPGPLRYKRGRIVPYVGSNTSEKLAPPILRMEMMSGPLKSCIRTASILGWVCEENPRTKIAIDWAVQQRASIDMSDLKLLTPASTSSNVYHRLHDSFGKNNFTSNRPLNFGSQIHLSTNTLGIFSGDG-GGVDGNMIFQNVKHMSISLLDMALG--DKEDVKDVNYHLHGLG-
CCVREMGADFVSSFNCDPPDINLAKGNPLIYDPSPLTGPNLERIRMVGSQGTSRDILRSG--
VADRDMSARQALGQTLLTKLHTNISRVAARGNAPVVEEDGVGNISEMCYLGPVEVMREVGLQLVSETFQHSLRDLMEQGYTLAESYRAFLASLHVGCFQGFHNMLAVKAIREPLKEFALRHGFVYSRDAPFHGFGEIIRATLFYTVEDALRNPAEMVACAANRNVFRTDKYFD
WAYTLTVWAAMRDLSCPSPVMTIREWVELLKRTQETPTHGRAAVVLVLGEMLEANGWTMEKLVS--VRRLMEGTRVSVWECDPSAWAREAVRQEK-----------------------------------------------------YGRVPENPDRASRRAC-----------------------------------------
DVRLKVNQGVLGEVMQKMSAIGDAGVVRVRAGILRR---------------------SREQVPGRVTHKKLQSYREIFSARLTGIASSAHYKFIELLRLVDHESLKTCLCVAEGAGGFARVLCELPLTQKVYYSTLIDWSRTTEHYTAADMGSELRVNA------------
HTKIVSLVRFLNDLRDLRTIQFLEER-CHSVEIVTCDAEM-FFKDLSYDMNLYKNVLCLAGSVLVPSGVFILKVFSDDEPVMQFSLSACCAMFHDVRFVKPMFSSPDSAEVYVIAQNKRPEAHQICDMIRDSLGSTYF------
PLMHQDTREVAADLSSLKELRIPKPWQDRSGVKSLTSAFIRCGWPSNRDQVLCSMGIDPKGWDDCAGGYSLISAGIQTMTACLVAQPER-----GNKHLLGSLQRGTAVLTGFLC---------------RREDLLRYLPLVGDTGE--VAQVRRKCTDGLAY-
ITMTSVVKSGILKMVQKLVGSLS-----------
>Kander_virus_YP_010800910
--MEGLASVPRSSIMFPDPHLNSPIVPEARDCVLILAGYSNLITQNPKVRLVSNHPRFSKVA--RKLSEILLLAQDEGDCLLGVKPQHLYGEIFQM-----DPQTESRSLLRYRDAYKVARGVERRQTTDLRHVLTKWYSGA----
PHLLELMLEKVERMLESRSEETNIMWRWAIINAVSEHMRMTKKNRKRS---------------------------DPVLTFKLKTTW--VDLYEIEEFTCVVSRRSGTWTLLSTEEILMFSDTLISRFNTVKYLRLTHSEATD-
ARLPTVKNVLELYHHVDGWFRPMGSTVYTIVKFLEPLCVGLTLMDSPLCEQRSHFLTEQACSLVDKLVEKHLPR---
ADAVRYMQKFLNLLAVSAPGPNEASELHGCFRHWGHNTIIIGEAVKAIKKHAQQIKMLKLDTLTRVHCVWMMLLVKHHYQTHSRSWPYVEIDRVLCPNLTHHRRMSTFPTDEQSLGFIYEWAHVKPGVLYSLDSIEDPRVFVKDKSVAPDRSHWDCAFDGSLLGYQPPYYKE
SRRLADRFVTDEGFSPYELIRFVESGEIYEDPEFNINLSMKERELS-
AGRPFGKLTYKARQCQALGEYLMAEGPGRAFESNTMCHSELELMKEMASQSHVQTTCEDQVPPQGESCGATMVTDYSKFCANFRHETCRSVAKDCARLFGLEDFFSWQHQRIASSMIYVADYHHPPSMNDQTTRKDPLEDETLWYNHTGGIEGYQQKLWTMYCCALLH
LASMETGVRYSAMIQGDNQIITGIVTATGNETRLELRERAVGEVKVLGHRMMVLAKQIGLTLKPEETFVAGSCTMYSKRLLLHGSDLTGALKIICRSMPNSSAVFDDPDAQTSSISSCGSRSVGRGVGVLGSYALSEFHVSFAIRLSELYHLGLPEGLARGSGV--
DLLSEYELYLLCNIGTGIGGVSANSLMRYLVRHEPDPVSNAIHWLYLKGGCDPNHLRTLSKIISSVQPSRDLLSLIKDPHSISLPTCRTVSSLVKTAVKDGLRETSKNPVLKGLFHDSCDQDDLAMAVFLLSTEPCIPVFSSELYQNSVCGVRDKLTGFVDSSKTTLASSLTRGY-
SGGMQVNIIRVLVLRHQFQVSLVKELPERV--NPLVLKCSLDLAHYLREKSWGDALRAGVMKGDSHPTVFEQFSLFEAGG-VGCHSCEGGNPTYLDLRIVEA---------
KHPGPLRYKRGKIIPYVGSTTSEKLTPPILRMEMMSGPLKSCVRTASVISWVCEENDRTKVAIEWAVKQRACIETSDLRLLTPASTNSNVYHRLHDNFGKNNFTSNRPLNFGSQVHLSTNTMGKFSGDG-GGVDGNMIFQNVKHLAISLLDLALG--DSSLVPDVNYHLHGIG-
CCIRELPPDFVTSFNCDPPEIDLAKGNPLIYDPSPLTGPNLDRIRMVGSGGTSRDILRSG--
GADREISARQALGQTLLAKLHVCIGKVAARGGTPAVEEDGVGNISEMCHLGPVEVFEEVGLQLITEIFQDTLKDLMLQGYTLIESYRAFLNSLHVGCFQGFHNMLTVRSIREPLKEFVLKHGFMYNSDAPFHSFGEIIRAVLFFTVDDLFRRPEALENTSRQRHVFRSDKXFDWAY
TLTVWSALKDLSGGGGVMGIGDWVELLKRTQRTPTHGRAAVILVLGEMLEGEGWGMGQLVQ--VRRLMEGTHVSVWECDPSAWAREAVRQVK----------------------------------------------------YGSSAPVCSPRDGPTI--------------------------------------
SVKEEVNKGSLSDVLRIMGEIGEPGVVKIRTGSMQR-----------------LPNPERVTGQVTHRKSTSFREIFSVRLTGVASSAHYKFCELLRLIDVQPIRRALCVAEGAGGFARVLCELPGVETVFYSTLIDWSRTTDHYTASDMGSELRVNA------------
HEKVTSLVRYPSDLRDQNTIRFLQVR-CRNIDLLSCDAEM-FFKDLPFDMNLYKNVLILAGSVLSGSGVFILKLFSSDGPAFQFVLSSCAAMFFDVEVVKPVFSSPDSTEVYILSCRLKPEVGNITALIERSLGTDSF------
PLFHRDVRGVTDDTHALREARVPKPWLKKSSVQALTQALIKCGWSKNHEQVLCSMGVDPAGWNSDNGGYTLISIGTQTMTAGLVAQPER-----DNRHLLSNLQRGMTVLTGYLC--------------KRDDLLRYLPLVGDTGE--VAQIRRKCTDGLAY-IN-----------------------------
----
```

**Data S4. Multiple sequence alignment of NP and NP-like sequences (nucleotides) from Filoviruses and open reading frame (ORF) filovirus-like elements in vertebrate genomes.** The data were used for the analysis in Fig. S5 and related to Fig. 4 (amino acids).

>NC_002549_Zaire_ebolavirus
ATGGATTCTCGTCCTCAGAAAATCTGGATGGCGCCGAGTCTCACTGAATCTGACATGGATTACCACAAGATCTTGACA--------------------GCAGGTCTG---TCCGTTCAACAG--------------------GGGATTGTTCGGCAAAGAGTCATCCCAGTG-----------
TATCAAGTAAACAATCTTGAAGAAATTTGCCAACTTATCATACAGGCCTTTGAAGCAGGTGTTGATTTTCAAGAGAGTGCGGACAGTTTCCTTCTCATGCTTTGTCTTCATCATGCGTACCAGGGAGATTACAAACTTTTCTTGGAAAGTGGCGCAGTCAAGTATTTGG
AAGGGCACGGGTTCCGTTTTGAAGTCAAGAAGCGTGATGGA------GTGAAGCGCCTTGAGGAATTGCTGCCAGCAGTATCTAGTGGAAAAAACATTAAGAGAACACTTGCTGCCATGCCGGAAGAGGAGACAACTGAAGCTAATGCCGGTCAG---------------------
---------------TTTCTCTCCTTTGCAAGTCTATTCCTTCCGAAATTGGTAGTAGGAGAAAAGGCTTGCCTTGAGAAGGTTCAAAGGCAAATTCAAGTACATGCAGAGCAAGGACTGATACAA--------------------
TATCCAACAGCTTGGCAATCAGTAGGACACATGATGGTGATTTTCCGTTTGATGCGAACAAATTTTCTGATCAAATTTCTCCTAATACACCAAGGGATGCACATGGTTGCCGGGCATGATGCCAACGATGCTGTGATTTCAAATTCAGTGGCTCAAGCTCGTTTTTCAG
GCTTATTGATTGTCAAAACAGTACTTGATCATATCCTACAAAAGACAGAACGAGGA------
GTTCGTCTCCATCCTCTTGCAAGGACCGCCAAGGTAAAAAATGAGGTGAACTCCTTTAAGGCTGCACTCAGCTCCCTGGCCAAGCATGGAGAGTATGCTCCTTTCGCCCGACTTTTGAACCTTTCTGGAGTAAATAATCTTGAGCATGGTCTTTTCCCTCAACTATCGG
CAATTGCACTCGGAGTCGCCACAGCACACGGGAGTACCCTCGCAGGAGTAAATGTTGGAGAACAGTATCAACAACTCAGAGAGGCTGCCACTGAGGCTGAGAAGCAACTC---------CAACAATATGCAGAGTCTCGCGAA------CTT---
GACCATCTTGGACTTGATGATCAGGAAAAGAAAATTCTTATGAACTTCCATCAGAAAAAGAACGAAATCAGCTTCCAGCAAACAAACGCTATGGTAACTCTAAGAAAAGAGCGCCTGGCCAAGCTGACAGAAGCTATC
>NC_004161_Reston_ebolavirus
ATGGATCGTGGGACCAGAAGAATCTGGGTGTCGCAAAATCAAGGTGATACTGATTTAGATTATCATAAAATTTTGACA--------------------GCTGGCCTT---ACTGTTCAACAG--------------------GGAATTGTCAGGCAGAAAATAATTTCTGTA-----------
TATCTTGTTGATAACTTGGAGGCTATGTGTCAATTGGTAATACAAGCCTTTGAGGCCGGAATTGATTTCCAAGAAAATGCCGACAGCTTCCTTCTGATGCTTTGCCTACATCATGCTTACCAAGGTGACTATAAATTGTTCTTGGAGAGCAATGCTGTACAGTATTTGG
AAGGTCATGGATTCAAATTTGAGCTCCGGAAGAAGGACGGT------GTCAATCGGCTCGAGGAATTGCTTCCTGCTGCAACGAGTGGAAAAAACATCAGGCGTACGTTGGCCGCACTGCCTGAAGAGGAGACTACAGAAGCAAATGCAGGGCAA---------------------
---------------TTTCTCTCATTTGCGAGTTTGTTTCTTCCCAAACTGGTTGTGGGAGAGAAGGCTTGCTTGGAAAAAGTCCAGCGACAAATTCAGGTTCATGCAGAACAGGGTTTAATTCAA--------------------
TATCCCACTGCATGGCAATCAGTTGGACACATGATGGTAATCTTCAGATTGATGAGGACTAATTTCTTGATTAAATATTTACTGATCCACCAGGGTATGCATATGGTAGCTGGCCACGATGCCAATGATGCTGTCATTGCTAATTCAGTTGCTCAGGCTCGCTTTTCAG
GACTCCTAATTGTCAAAACCGTTCTTGATCATATTCTGCAAAAAACCGACCAAGGA------
GTAAGACTTCACCCTTTGGCCCGAACAGCCAAAGTGCGTAATGAGGTTAATGCATTTAAGGCCGCCCTAAGCTCACTTGCTAAGCATGGGGAATATGCCCCTTTTGCTCGCCTTCTCAATCTCTCGGGAGTTAACAACCTAGAACATGGTCTCTACCCACAGTTATCAG
CAATTGCTCTTGGAGTTGCCACAGCACATGGTAGCACCCTTGCAGGAGTTAATGTTGGTGAGCAGTATCAGCAGCTTAGAGAGGCTGCCACTGAAGCTGAGAAGCAACTC---------CAACAATATGCTGAGTCCAGAGAA------CTC---
GACAGCCTAGGCCTGGACGATCAGGAAAGAAGAATACTAATGAACTTCCATCAGAAGAAAAACGAAATTAGTTTCCAGCAGACCAATGCAATGGTAACCCTTAGGAAAGAGCGACTGGCTAAATTAACAGAAGCTATA
>NC_006432_Sudan_ebolavirus
ATGGATAAACGGGTGAGAGGTTCATGGGCCCTGGGAGGACAATCTGAAGTTGATCTTGACTACCACAAAATATTAACA--------------------GCCGGGCTT---TCGGTCCAACAA--------------------GGGATTGTGCGACAAAGAGTCATCCCGGTA-----------
TATGTTGTGAGTGATCTTGAGGGTATTTGTCAACATATCATTCAGGCCTTTGAAGCAGGCGTAGATTTCCAAGATAATGCTGACAGCTTCCTTTTACTTTTATGTTTACATCATGCTTACCAAGGAGATCATAGGCTCTTCCTCAAAAGTGATGCAGTTCAATACTTAGA
GGGCCATGGTTTCAGGTTTGAGGTCCGAGAAAAGGAGAAT------GTGCACCGTCTGGATGAATTGTTGCCCAATGTCACCGGTGGAAAAAATCTTAGGAGAACATTGGCTGCAATGCCTGAAGAGGAGACAACAGAAGCTAATGCTGGTCAG-----------------------
-------------TTTTTATCCTTTGCCAGTTTGTTTCTACCCAAACTTGTCGTTGGGGAGAAAGCGTGTCTGGAAAAAGTACAAAGGCAGATTCAGGTCCATGCAGAACAAGGGCTCATTCAA--------------------
TATCCAACTTCCTGGCAATCAGTTGGACACATGATGGTGATCTTCCGTTTGATGAGAACAAACTTTTTAATCAAGTTCCTACTAATACATCAGGGGATGCACATGGTCGCAGGCCATGATGCGAATGACACAGTAATATCTAATTCTGTTGCCCAAGCAAGGTTCTCTG
GTCTTCTGATTGTAAAGACTGTTCTGGACCACATCCTACAAAAAACAGATCTTGGA------
GTACGACTTCATCCACTGGCCAGGACAGCAAAAGTCAAGAATGAGGTCAGTTCATTCAAGGCAGCTCTTGGCTCACTTGCCAAGCATGGAGAATATGCTCCATTTGCACGTCTCCTCAATCTTTCTGGAGTCAACAACTTGGAACATGGGCTTTATCCACAACTCTCA
GCCATTGCTTTGGGTGTTGCAACTGCCCACGGGAGCACGCTGGCTGGTGTTAATGTAGGGGAGCAATATCAGCAACTGCGTGAGGCTGCTACTGAAGCTGAAAAGCAACTC---------CAACAATATGCTGAAACACGTGAG------TTG---
GACAACCTTGGGCTTGATGAACAGGAAAAGAAGATTCTCATGAGCTTCCACCAGAAGAAGAATGAGATCAGCTTCCAGCAAACTAACGCAATGGTAACCCTGAGGAAAGAGCGGCTGGCCAAACTGACCGAAGCCATC
>NC_001608_Marburg_marburgvirus
ATG---------------------------------------------------GATTTACACAGTTTGTTGGAG--------------------TTGGGTACA---AAACCCACTGCC--------------------CCTCATGTTCGTAATAAGAAAGTGATATTA-----------
TTTGACACAAATCATCAGGTTAGTATCTGTAATCAGATAATAGATGCAATAAACTCAGGGATTGATCTTGGAGATCTCCTAGAAGGGGGTTTGCTGACGTTGTGTGTTGAGCATTACTATAATTCTGATAAGGATAAATTCAACACAAGTCCTATCGCGAAGTACTTA
CGTGATGCGGGCTATGAATTTGATGTCATCAAGAATGCAGAT------GCAACCCGCTTTCTGGATGTGATTCCTAATGAACCTCATTACAGCCCTTTAATTCTAGCCCTTAAGACATTGGAAAGTACTGAATCTCAGAGGGGGGAGAATTGGGCTC--------------------------
-----------TTTTTATCATTTTGCAGTCTTTTCCTCCCAAAACTTGTCGTCGGAGACCGAGCTAGTATCGAAAAGGCTTTAAGACAAGTAACAGTGCATCAAGAACAGGGGATCGTCACA--------------------
TACCCTAATCATTGGCTTACCACAGGCCACATGAAAGTAATTTTCGGGATTTTGAGGTCCAGCTTCATTTTAAAGTTTGTGTTGATTCATCAAGGAGTAAATTTGGTGACAGGTCATGATGCCTATGACAGTATCATTAGTAATTCAGTAGGTCAAACTAGATTCTCAG
GACTTCTTATCGTGAAAACAGTTCTCGAGTTCATCTTGCAAAAAACTGATTCAGGG------
GTGACACTACATCCTTTGGTGCGGACCTCCAAAGTAAAAAATGAAGTTGCTAGTTTCAAGCAGGCGTTGAGCAACCTAGCCCGACATGGGGAATACGCACCATTTGCACGGGTTCTGAATTTATCAGGGATTAACAACCTCGAACATGGACTCTATCCTCAGCTTTCA
GCAATTGCGCTGGGTGTGGCAACAGCACACGGCAGTACATTGGCTGGTGTCAATGTTGGCGAACAATATCAACAACTACGAGAGGCGGCACATGATGCGGAAGTAAAACTA---------CAAAGGCGACATGAACATCAGGAA------ATT---
CAAGCTATTGCCGAGGATGACGAGGAAAGGAAGATATTAGAACAATTCCACCTTCAGAAAACTGAAATCACACACAGTCAGACACTAGCCGTCCTCAGCCAGAAACGAGAAAAATTAGCTCGTCTCGCTGCAGAAATT
>NC_014372_Tai_Forest_ebolavirus
ATGGAGAGTCGGGCCCACAAAGCATGGATGACGCACACCGCATCAGGTTTCGAAACAGATTACCATAAGATTTTAACA--------------------GCAGGATTG---TCAGTCCAACAA--------------------GGCATTGTGAGACAACGGGTCATTCAAGTC-----------
CACCAGGTTACAAACCTAGAAGAAATATGCCAATTGATCATTCAAGCCTTTGAAGCTGGTGTTGATTTTCAAGAGAGTGCAGACAGTTTCTTGCTGATGCTATGTTTACATCATGCTTATCAGGGTGACTACAAGCAATTCTTGGAAAGCAATGCAGTCAAGTACCTT
GAGGGTCATGGCTTTCGCTTTGAGGTCAGGAAAAAGGAAGGA------GTCAAGCGACTCGAAGAATTGCTTCCTGCTGCATCCAGTGGCAAGAGCATCAGGAGAACACTGGCTGCAATGCCTGAAGAGGAGACAACAGAAGCAAATGCCGGACAG-------------------
-------------------TTCCTCTCTTTTGCTAGCTTATTTCTTCCTAAGCTAGTTGTCGGAGAAAAAGCCTGTCTAGAAAAGGTGCAGCGGCAAATTCAAGTTCATTCTGAGCAGGGATTGATCCAA--------------------
TACCCCACAGCCTGGCAGTCAGTTGGACACATGATGGTCATTTTCAGACTGATGAGAACAAATTTTCTAATTAAGTTCCTCCTTATACATCAAGGGATGCATATGGTAGCAGGACACGATGCTAACGATGCTGTCATCGCAAACTCTGTAGCTCAAGCACGTTTTTCAG
GATTATTGATCGTTAAAACAGTGCTAGATCACATCCTTCAGAAAACAGAGCACGGA------
GTGCGTCTTCATCCTTTGGCAAGAACTGCTAAGGTCAAGAACGAAGTAAATTCCTTTAAGGCTGCCCTTAGCTCGCTAGCACAACATGGAGAGTATGCTCCTTTTGCTCGCTTGCTGAATCTTTCTGGAGTCAACAATCTCGAGCACGGACTGTTTCCTCAGCTTTCTG
CAATTGCCCTAGGTGTCGCAACGGCACACGGCAGTACCCTGGCAGGAGTAAATGTGGGGGAACAGTATCAGCAACTACGAGAAGCAGCCACTGAGGCAGAAAAACAATTG---------CAGAAATACGCTGAATCTCGCGAG------CTT---
GACCATCTAGGTCTCGATGATCAAGAGAAGAAGATCTTGAAAGACTTCCATCAGAAGAAAAATGAAATCAGCTTCCAGCAGACAACAGCCATGGTCACACTACGGAAGGAAAGGCTAGCCAAGCTCACTGAGGCAATC
>NC_014373_Bundibugyo_ebolavirus
ATGGATCCTCGTCCAATCAGAACCTGGATGATGCATAACACATCTGAAGTTGAAGCAGACTACCATAAGATTCTAACT--------------------GCCGGATTG---TCCGTCCAGCAA--------------------GGCATTGTGAGACAAAGAATCATTCCTGTT-----------
TACCAAATCTCAAACCTGGAGGAAGTATGTCAACTCATCATACAGGCATTCGAGGCTGGCGTCGACTTCCAGGATAGTGCAGATAGCTTTTTGTTAATGCTATGTCTGCATCATGCCTATCAAGGGGATTATAAACAATTTTTGGAAAGTAATGCGGTAAAATACCTT
GAAGGTCATGGATTCCGTTTTGAGATGAAGAAAAAGGAAGGT------GTCAAGCGCCTGGAGGAACTACTCCCTGCTGCCTCGAGTGGAAAGAACATCAAGAGAACATTGGCTGCAATGCCCGAGGAGGAAACAACAGAAGCAAATGCTGGACAA-------------------
-------------------TTTCTTTCATTTGCTAGTCTGTTTCTCCCAAAATTGGTTGTCGGAGAAAAGGCCTGTCTGGAGAAGGTTCAACGACAAATCCAAGTGCACGCAGAACAAGGTCTGATTCAA--------------------
TACCCGACATCTTGGCAATCGGTGGGACATATGATGGTCATCTTCAGACTAATGCGAACCAACTTCCTGATTAAGTTCCTCCTAATACATCAAGGAATGCATATGGTTGCAGGGCATGATGCTAATGATGCCGTCATTGCCAACTCTGTAGCTCAAGCTCGTTTCTCCG
GATTGTTGATAGTCAAAACAGTGCTTGATCATATCCTCCAAAAAACAGAGCACGGA------
GTTCGCCTGCATCCCTTGGCGCGAACAGCCAAAGTCAAAAATGAGGTGAGCTCTTTTAAGGCCGCTTTAGCCTCACTAGCACAACATGGAGAATATGCCCCGTTTGCTCGTCTGCTGAATCTATCTGGGGTTAATAATCTTGAGCATGGGCTTTTCCCTCAACTTTCTG
CAATTGCTTTGGGAGTAGCAACTGCACATGGGAGCACTCTGGCTGGAGTCAATGTAGGAGAGCAATACCAACAACTGCGAGAAGCAGCCACTGAGGCCGAAAAGCAGTTG---------CAGAAATATGCTGAATCTCGTGAA------CTT---
GATCACCTAGGTCTTGATGATCAGGAAAAGAAAATCCTAAAAGACTTCCATCAGAAAAAGAATGAGATCAGCTTCCAGCAGACGACAGCCATGGTCACACTGCGGAAAGAGAGATTGGCCAAATTGACCGAAGCTATT
>NC_016144_Lloviu_cuevavirus
ATGAATCGTTATCTTGGTCATGGTACTAGAACATCTAGGGAAAATACCAACCTGAGTGAATTACATGGTATTCTAAGC--------------------CTCGGGTTA---AATGTAGATCAT--------------------ACCATTGTAAGGAAGAAGAGCATCCCGTTG-----------
TTTGAGATCGGTAACTCTGATCAAGTATGTAATTGGATAATTCAAATAATTGAGGCTGGGGTGGACCTGCAGGATGTTGCTGATAGCTTTTTAACAATGCTCTGCGTAAATCATGCTTATCAAGGAGATCCTAATTTGTTCCTTGAGAGTCCTGCAGCTCATTATTTAA
AAGGACATGGTATCCACTTCGAAATCCAGCATCGTGATAAT------GTTGATCACATCACAGATCTCCTTGGAGTAGGGAGTCGAGACAAATCACTCAGAAAAACTCTTTCTGCACTTGAATTCGAGCCTGATGGCTCAACTACGGCTGGGATG---------------------------
-----------TTTTTGTCTTTCGCAAGTTTATTTCTCCCGAAGTTAGTTGTCGGTGAACGTGCCTGCTTGGAAAAAGTGCAGCGGCAAATTCAAATTCATGCAGAACAAGGCTTAATCCAA--------------------
TATCCCACCCAATGGCAGTCCGTAGGACATATGATGGTTGTATTCAGGCTGATTCGTGTCAACTTTGTCCTCAAATTCCTCCTAGTTCACCAGGGCATGCACATGATGGCTGGGCATGATGCTAATGATGCAATCATTGCCAATTCAATATCCCAAACACGATTTTCAG
GATTGTTAATCGTGAAGACTGTCCTAGAGCATATTCTCCAAAAAACTGAGGCAGGG------
GTCCAACTTCATCCATTAGCTCGAACCTCGAAAGTCAAAGGAGAACTCCTTGCATTCAAATCAGCACTGGAAGCGTTGGCCTCACACAGAGAGTATGCACCATTTGCACGGCTCTTGAACCTCTCTGGTGTTAATAATCTAGAACATGGCCTGTATCCTCAGCTCTCTG
CCATCGCGCTTGGAGTTGCTACAGCTCATGGTAGCACACTAGCAGGAGTTAATGTGAGTGAACAATACCAGCAACTCCGAGAGGCTGCAACAGAGGCTGAGAAACAATTA---------CAACAGCACTCAGAGATGCGGGAA------TTG---
GAGACCCTTGGACTGGATGAACAAGAAAGGAAGATCCTTGCAACATTCCACTCCCGGAAAAATGAGATCAACATTCAACAGACCAGCTCGATATTAGCAATTCGTAAGGAACGACTCCGCAAGCTGACAGAGGCCCTC
>NC_024781_Marburg_marburgvirus
ATG---------------------------------------------------GATTTACATAGTTTGCTAGAA--------------------TTAGGTACA---AAACCCACAGCC--------------------CCTCATGTTCGTAATAAGAAGGTGATATTG-----------
TTTGATACAAACCATCAGGTTAGTATCTGTAACCAGATAATAGATGCAATAAACTCAGGGATCGATTTAGGAGATCTTTTGGAAGGCGGCTTGTTGACATTATGTGTTGAACATTATTACAATTCTGACAAAGATAAATTCAATACAAGTCCTATTGCAAAATATCTGC
GGGACGCGGGCTATGAGTTTGATGTCATCAAGAATCCTGAT------GCAACTCGCTTTCTAGAGGTTATTCCCAATGAACCTCATTACAGCCCTTTGATTTTGGCTCTCAAAACCTTAGAAAGCACTGAATCTCAAAGGGGGAGGATTGGGCTC---------------------------
----------TTTCTGTCATTTTGCAGTCTTTTTCTCCCGAAACTCGTTGTCGGAGACCGAGCTAGCATCGAAAAGGCCCTGAGACAAGTAACAGTGCATCAGGAACAAGGGATTGTAACA--------------------
TACCCCAATCATTGGCTCACTACAGGTCACATGAAAGTAATCTTTGGGATTTTAAGATCTAGCTTCATTTTAAAGTTTGTCTTAATCCATCAGGGAGTAAACTTGGTGACAGGTCATGATGCCTATGACAGTATCATCAGTAATTCAGTAGGACAAACTAGATTCTCAG
GGCTTCTTATTGTGAAAACAGTTCTAGAGTTCATCCTACAAAAAACTGATTCAGGG------
GTGGCATTGCATCCACTTGTGCGGACCTCAAAAGTAAAAAATGAAGTTGCAAGCTTCAAACAGGCATTGAGTAACTTAGCTCGTCATGGAGAGTACGCACCATTTGCACGGGTTTTGAATTTATCAGGGATCAATAACCTTGAGCATGGGCTCTATCCTCAGCTTTCA
GCAATTGCACTGGGCGTTGCAACAGCACATGGCAGTACATTGGCTGGTGTTAACGTTGGCGAACAATACCAACAACTGCGAGAAGCAGCACATGATGCAGAAGTAAAATTA---------CAAAGGCGACATGAACACCAGGAA------ATT---
CAGGCCATCGCCGAGGATGACGAAGAGAGAAAAATATTAGAACAGTTCCATCTCCAGAAGACTGAGATTACACACAGTCAGACATTGGCCGTCCTCAGCCAGAAACGAGAGAAACTAGCCCGTCTTGCTGCAGAAATT
>NC_039345_Bombali_ebolavirus
ATGGAGGTTCGCAATTCTCGCCAATGGACTACGCAGAGTGCATCTGACTCCAGCGTGGATTACCACAGTATCCTAACT--------------------GCAGGGTTA---TCGATGCCACAA--------------------AGTATAGTTCGCCAAAGAGTAATCCCCGTG-----------
TTTCAAATTAGCAACCTTGAAGATATTTGTCAAATGATTATACAGGCTTTTGAGGCAGGTGTAGACTTCCAGGATAGTGCTGATAGTTTTCTACTAATGCTTTGTCTACATCATGCCTACCAGGGTGATTATAAGTTGTTCCTTGAGAGTGGGGCTGTTAAATATCTTGA
AGGACATGGGTTTAGGTTTGAACCCCGCAAGAAGGACGGA------GTCAAGAGATTAGAAGAATTACTTCCGGCAGTTACAAATGGAAAGAATATAAGAAGAACTCTAGCCGCGATGCCAGAAGGTGAGACAACTGAGGCTAACGCTGGGCAG----------------------
--------------TTCCTTTCATTCGCCAGCCTATTCCTGCCGAAACTTGTTGTTGGAGAAAAGGCCTGTCTGGAGAAGGTGCAACGGCAGATCCAAGTCCATGCTGAGCAGGGATTAATCCAG--------------------
TATCCAACCTCGTGGCAATCAGTTGGGCACATGATGGTTATCTTCCGGTTGATGAGAACAAATTTCCTGATCAAGTTCTTGTTGATACATCAGGGTATGCATATGATTGCCGGACATGATGCCAACGATGCTGTTATCGCTAATTCAGTTGCTCAGGCACGCTTTTCAG
GCTTGTTGATCGTCAAGACAGTGCTGGACCACATTCTGCAGAAAACAGACCGTGGA------
GTCCGTCTTCATCCTTTAGCACGTACAGCCAAGGTCAGGAACGAGGTCCATTCGTTCAAAGCTGCTCTTAACTCCCTGGCTCAGCATGGGGAATATGCCCCCTTTGCTCGACTTTTAAACCTGTCTGGAGTAAATAACCTCGAACATGGGCTGTATCCTCAACTCTCTG
CAATTGCACTTGGTGTGGCTACTGCACACGGCAGCACTTTGGCAGGGGTGAATGTTGGTGAGCAGTACCAACAGCTCAGGGAGGCTGCCACGGAGGCTGAGAAGCAGCTG---------CAGCAATATGCTGAAACACGTGAG------CTG---
GACCACTTAGGCCTAGACACAAAAGAAAAGGAAATATTAATGAATTTCCATCAACGGAAAAATGAGATTAGTTTCCAGCAAACAAATGCAATGGTGTCCCTTCGCAAGGAGAGACTAGCAAAACTAACAGAAGCTATT
>NC_055175_Wenling_frogfish
ATGGATGCC---------------------------------------------AACCAGGTCAAGATATTCTCTGAC--------------------ATCGCTAGT---ACTCCCTCAGAC--------------------AGTTATGTCTCCAGATCCAAGCTTAGGGTC-----------
TACGCATCTAAGGCTCCTGACCTGCTCTATGGACTCCTTGTTGCTGCTCTGCTGGATCCGGAGCTTCCAAGATCAACAGCGGCAAGTGTGTTCGTCTCCCTGTTCATCGGGCATTTCTACAGAGGGGATATCACGAGTTTCTCTACATCCTCCATTGCCAAGGAACTGA
CCAAATTCGGACACAACCTCATCTACGTGGAGGTGGACATCGATGTTGCGAGCCGCACCATCATCTCG------------CCAAAGGCCTCAAGACTCAGCCAGTACATCAAAAAGGCAGAACTCCTC------ACTGACAGGACCCTAGAACGT------------------------------------
--TTCATCACATACCTCGCAGCTATGATGCCTAAGCTGATGATATCAATTGATGCTGTCCACGCAAAGGTCGTCAAGACCGATGCAGTTCTCAAGTCCCAAGGTATCCCCAGC--------------------
CTGGGTGATGTCTTGATCCACAAAGCTACAGTAACCACTGTCAGGGCACACCTCATGGACTCTTTTCCAGGAAAGTATCTCCTGATCAAACTGACCATCTCTCAGGTGTCATCAGGGTCCGTCTGTGGTGGCCAGATCAAGACACTCATCGATCAGGCCAGGATGGCA

GGGCTGGTCCTGATCAAACTTTTCACAGATCATCTGATCTCCGGGTCAGGCGGGAAAGGGAACAAGTCCTGGCATCCCATCTTAAGCCTCAAGGATCTTGAAAGAGAGGCAGGGGACATGGTGGCGGCCATAAATGGAATTAGCGCACACGGAGATCTGGCTCCA
TTCGCAAGGATCCTAGGACTGTCCGGAGTCGAAAAAGTTGAGTATGGAAAATTCCCAAAATTGGCTGCCGTCATTGTTGGGATCGGACAAGCGCACAATAGCTCTCTCAATCTCATGGTGGTAGGAGCTGCACACAAGCCCCTTTGCGAGGCAGCCAGGCTTTTTGA
GGAATCCCGC---------AAGAAAGACGGCGAGGCCAAACTCGGAGGCATG---
TTCCCCACCCAGGTTACCCAGGCGGAAAAACGTCGTCTGGCTGATTTCCACAAGGACATGACAAAGGCAACAGAAGAGTCCCTGAAAGCAGCCAAAGCAGCACAGATGAGAACACTCCAAGCGATCCGGGCAAGCGTT
>NC_055510_Mengla_dianlovirus
ATG---------------------------------------------------GATTTACACGGATTGTTAGAA---------------------TTAGGAACT---AGACCAACAGCT---------------------CCTCATGTAAGGAGTCGGAAGATTGTCATA------------
TATGAGACAGGCAATCAGATCATCATCTGTAATCAAATAATTGATGCAGTTAGTGCGGGAATTGATCTTGGTGATCTATTAGAAGGATGCTTGCTGACATTGTGCTTAGAACATTACTATGGTTCTGATAAAGACAAATTCAACAGCAGTCAAATGGCTGCATATCTC
AGAGATGCAGGTTATGACTTTGAGGTTATCAGGGCTCAGGAT------GCTAAAAAATTAGCAGACCTCATACCCAGAGAAAGTCACCTATTGAATGTGATTTCTGCCCTTGAGAACTTGGATGGCTCCGAAAAGAATAAACAAAGAGTTGGATTA------------------------
-------------TTTCTATCATTCTGCAGTCTTTTCCTCCCAAAATTGGTTGTGGGTGATAAGGCAAGCATTGAAAAGGCACTTAGACAAGTTGCAATCCATCAAGAACAAGGGATGGTTGTT---------------------
TACCCACCAACCTGGCTAACCACGGGTTTCATGAAGCTCATCTTCAGCATTGTGCGCGCGAGCTTTATTGTTAAGTTTGTGCTCATCCACCAGGGAATTAATCTTGTCACAGGTCATGATGCCTATGACAATGTCATCAGCAATTCCATAAACCAAACACGCTTCTCAG
GACTGCTGATTGTAAAGACTGTCTTGGAACATATCCTCCAAAGGACTGAAAATGGA------
GTCATTCTTCACCCTCTTGTCAGAACCTCCAAGGTGAAGAGTGAAGTTGAGAGCTTCAAAGTTGCATTGAGAGGTCTGGCAAGACACAAAGAATACGCACCTTTTGCCCGGGTATTGAACTTATCTGGGGTGAATAATCTTGAGCATGGACTTTTCCCTCAACTCTCA
GCAATTGCAATCGGGGTTGCAACAGCACATGGAAGTACATTGGCTGGGGTGAGTGTTGGAGAGCAGTATCAACAACTACGGGAAGCAGCCCATGATGCAGAACTTAAACTC---------CAGAGGCGCCGTGAACAAATGGAA------ATT---
TCATCACTCGAACTTGACCTTGAAGAACAGAAAATTCTGGAACAATTCCATCAGCAGAAAACAGAAATAACTCACACACAGACTCTTGCTGTTCTAACACAGAAGAAAGAAAAGTTAGCCAAACTTGCTGGGGAACTT
>NC_076535_Tapajos_virus
ATGTCACAA------------------------------------------TCCAGTCTTCATTCTTTTTTGAAG---------------------ACTGGAATGCCAAGCCTATCACACGCCGACAAAGTCAACAAACACCGTGATACGCGTGTTAAAATCGTAAATGTT------------
AACTTGTGTTCTAATTTACGGAGAGTGTGCTCCGAGATTATACTTTGTTTTATCGGGTCATTTGACACACACGAATTCCAAGATGAGATTGTACTTCTACTCATTGTGCAATTCTATTATGCAGGGGATGCACAAAGATTCAAATCCAGCATATTCTGCCAACAACTTCT
TGAGGCGGGGTATCAGATTAATTTTTATAGTTCCAAATTA------GAAGGGCGTCTTCTGGATTGTCTTGCTCCCCTTAAAGAAGACTCTAGATTAGCAGCTGTCCTCAAGAATATCCTCTTACCTACAGATGACTCTGACCAAACTCCAGCT---------------------------------
------CTCTTTGCATTAATGAGCATGTTCTTACCCAAGCTGATGGTTGGTGCTGCTGCCTGTGCCACCAAAGTCGGGCGCACAGTTGAAAAGTTGATTCAACAAGGAATTTGTCCC---------------------
ATCCCACCGGAATGGGTAAATACCACAGTCTTAGAATCAATATATCGAGTTATGAGAGCCTCTTATATCTTGAGGTTTGCCATAATACATCATGCTTTGGAACAAAGCCATGGATCAGATTCAAATGAGCGCATAATCCATGACTGTGTAATGCAAGCTCGTTTTGCA
GGTATGCTCCTACCAGTCACCATGCTGACTCATATCCTAGTGGAAGAACCAGAGGGT------
GTCAGACTCCACCTACTAGCAAGAAGTCAAGATATGAAGGAAGAACTGACACGGTTTCAGTTAGCTATTAGATCTCTACTTCGACACGGCAGATTAACACCTTTTGCTAGGATCCTAGGCCTTCCTGAAATGGCCCTTCTTGAACATGGACACTTCCCAAAACTTTCAG
CAATAGCCATGGGAGTTGCTTCAGTACACACTCCTACTCTAAACGCCACAGTAACTGATCCACGCTATGAGAAGATGAGGAGAGCTGCCGAAGAAGCAGAGAGAGCCCTC---------CATGATTACCAAAATGCTGAAATG------
GTGGACAACGACGCAACACTTTCTGCTGGGCAACGCGAAATACTCAAAGGTTTCCACAAGCAACGATCAGCCATCCGAGACGCCGAAACAACAGCAGCAGAGGAACTCAGGCGCGAAACCATTACAAGAGCATTAAAACAGGCC
>NC_055176_Wenling_thamnaconus_septentrionalis
ATGGCCGCA------------------------CCCAATAATGGTGGATTCGGGATCAACTTAGATGAGGTGATCCAC---CCCAGGGGTAACAATGGAGCTGGGGCAGCTGCCATAGGGATG---------------------
GCTGGGTTGGAGCGAGTCCCGATCTTCGTCCTGGCTAACCCGATGACTGGACCTGAGATGAGACTCCTGAATCTTGCCTGGGCTGTGGTCCTGGAAGACCCG---
AACCCTCCTGCCGTGATTGTTCAACTGTTGGTAGAGCTGACTTGCTCTGCCTACTATAACAATCCTAATGATCTGGTGGATCACATC---------
CTCCCCATGGTCGGTCCAGGAGGCTATGCATTGAGAGTCTCTTTCAATGGACTAGGCAGGTTCACAATCCCCGGGAGA---------
CCTGAGTTGGAGAGATTAGTGTCTGATTACATAAGACCACGTATCCAGCGAGTGATGAATGACAGGGCTTGGACCGCAAGATTGGCTCCAGGTACAGGTTTGGTGTTGTGGCAGGTGATGTTAATTCAACTGTGCTTGGCATCAGGACATCTCTTTACCGGGAATAC
AGCTCTTCAGACCTGGTTTACTCGAAGGGTGATTGCATTACAACTGAACACCAGGATTGACATCAACTTGACAGGTGCAATCTTTGCAGTACTTGTGACCCCCGGGGGCTCCCTGACCGTTGTCAGGTGCATGATTGTTCAAGTGATGATGGCAACACTGGCATCTCA
GCCTTTGGGAGTACCCCTGGTGGAAGGA---------------------ATAAGAGCTGCCACTAACTTCTCCGAGTGGGCCGGAATGGCCTGGGCTTTCCTCTCACTGACCTTAGTGATG------------------------
AGGTCTGCACATCCAATACTCCTTGTCCCGTCAGTGAGAAGGGAAGCACTGGCATTTGTGACAGCCATCCAGGGTCTCGGTGCCAATAGACTCAGACTCCCCTTCGAAAGAATCCTGGGTGCCTCACTCTCCAACGGGTTGAATGCTGGGAACTTCCCAACTCTTTCA
AGTGCTGCCTACGGGGTTGCATCGGCCACCTTCTGCAACTTTGCTCAGTTCAACGGAACAATTTACTCAATGGTTGTTTACCGAGCTGCTCAAATCCTGACAAGGGTTGTC---------ACCGCAGTTGACACAGGAGTTGTGGACCCCTTG------------------------------------
---------------------------ACTGGTCTGCAAGGACAACCGATGACCGACGCGACCAGAATGGTCACAGAGGCAATGTCTGAAGCCTCACAGGCTCAG
>AXCS01020818_Nannospalax_galili
ATG---------------------------------------------------------CATTCCTTATTGAAT---------------------GCCACATTGGGCAATGTTGAGGAT------------GATAAATGTGTACCCCCAAAGTCTAAACTGGTCAGGGTA------------
TTTACTGGCGATGGGGTTAAGCAAATAATTCAGCAGATTCTGGCCAGCTTTGAAGCTGGCCTGGAAATAGACAATCTAGCAGACCACTTCTTTGCTACCATAGTGTGCTATTACCATTACCGGAACAACCTCGATTTAATGGTAAGGAGTTCTTTTGGTCAGTATCTTG
TTGAAAACGGCTATGAATTTTCAGGATTAGAACTAGACAACAAA---AGAGAACCACTTCTTCAGCATCTTGAGGAACTGGGGGTGCCCGAAGGTCTGCTGAAAGTCCTAGGAGACCTGCCCTGTGCACCACCTCTGGGTCTGCGCCCGGCTGCA------------------------
---------------TTCCTCGTATACCTTAGTCTGTTCTTGCCCAAGTTAGTCGCTGGGACTACAGCATGCATTAAAAAAGTTGAACGGACTTACCAACAATTGGAGATCCAAGGTCTCAATATA---------------------
ATCCCCAGAGAGTGGATGACCCCTGTTATGATGAACCTTGCATATCGTGCCTTACGGCTCTGTTTCTTCATTCGATTCTCCATCATCCACCATGTGCTTCATCAACGGGCTGGAATAGATGCAGCTGACACAATTATGGCAGGAGCTGTGGAGCAGACTCGTTTTGCAG
GGCTGCTGATGGTTCGAAATGTCTTAACATACATCATTACCAAAGGGAAAGAAAAAGATGAATATCATATACACCCACTGGCTCAGGACAAAAGAATGAGGGAAGAAATAACCCGATTCGAGTACTGCATACAGAATATCCAACGGCACGGAATATATGCTCCATT
TGCACGAGTCCTGGGGCTGGCCGGAGTTTCTCAGATAGAGCATGGGCAATTCCCGCGATTATCCGCCATTGCTCTTGGAGTGTCCTCTGTCTATCAAAGCACACTCTCAGGGGTGAACATGGACACGCGCTACCAATCTCTGAAAGAAGCAGCCCATCAGGCTGAAT
TGGAGCTA---------CAAGCAATCCAGTATAAGAAGGAG------
ATCAGACAAGACACTGACTTAGACAATGATGAGAGAGTGGTCCTCCTCAAATTTGCTGAAAGAAGCGGAAATATCATCAAAGAACAGCGACAGCAGGAGAATCTTGCAAGACAAGAATTGCTCCAGAAGTTCACAGAGAAGGCC
>VZQC01004831_Rhizomys_pruinosus
ATG---------------------------------------------------------CATTCCTTGTTGAAT---------------------GGCGCATTGGGCGATGTTGAGGAC---------------------ACACCCTCTCAGTCTAAAATGGTCAGGGTA------------
TTTACTGGCGATGGGATTGAGCAAATAATCCAGCAGATCCTGGCCAGCTTTGAAGCCGGCCTGGCAATAGACAATCTGGTAGACCACTTCTTTGCTACCATCGTGTGCTATCACCATTACCGGAACAACCTTGATTTAATGGTACGGAGTTCTTTTGGTCAGTATCTTG
TTGCAAATGGATATGAATTTTCAGGAGTAGAACTACACAGCAGA---AAACAAACATTTCTTCAGCATCTCAAGGAGCTGGGGATGCCTGAAAGTACGCTGAAAGTCCTAGAAAACCTCCCCAGTCTACCTCCTCTCAGTCTACGCCCTGCTGCA---------------------------
------------TTCCTTGTATACCTTAGCCTGTTCTTGCCCAAATTAGTCACTGGTACTTCAGCAAGTATTAAAAAGGCTGAACGGACTTACCAACAACTGGAGGGCCAAGGTCTCAGTATA---------------------
ATCCCTAGCGGATGGATGAACCCCACCATGATGCACCTCACATATCGCACCTTACGCCTCTGTTTCTTCATTCGATTCTCTCTTATCCACCATGTGCTTCATCAACGGGCTGGAATAGATGCTGTTGATACAATTATGGCAGGAGCTGTGGAGCAGACTCGTTTTGCAG
GGCTGCAGATGGTTCGAAATGTCTTAACATACATCATTACCAAGGGAAATGAAAGCGATAAATATAATATACATCCACTGGCTCAGGACAACAGAATGAAAGAAGAACTAACTAGATTCGAGCACTGCGTAAAGAATATCCAACGGCACGGAATATATGCTCCGTTT
GCACGAGTCCTAGGGCTGGCCGGAGTTTCTCAGATAGAGCATGGGCAATTCCCACGATTGACGGCCATTGCTCTTGGAGTGTCCTCTGTCTATGAAAACACACTTTCAGGGGTAAATCTGGACACCCGCTACCACTATCTCAAAGAGGCAGCCCGTAAGGCTGAATT
GGAGCTA---------CAAGCAATCCAGCATAAAAAGGCG------ATCAGACAAGACACTGACTTAGATGAAGATGAGAGGGCAGTCCTCCTCAAATTTGCTGAAAGAAGTGGAAATATCAATAAAGAGCAGCATCAGGAGGCT------TTCACTAGAGAGCTGCTCCAG---
TTGACCCAGAACGAG
>SRR16768455_Eospalax_fontanierii
ATG---------------------------------------------------------CATTCCTTGTTGAAT---------------------GCCACATTGGGCAATGCTGACGAT------------GATCAATGTGCACCCCCTCAGTCTAAACTGGTCAGGGTA------------
TTTACTGGTGATGGGGTTGAGCAAATAATCCACCAGATTCTGGCTAGCTTTGAAGCTGGCCTGGCAATAGATAATATGACAGACCACTTCTTTGCTACCATTGTGTGCTATTACCATTACCGGAACAGCCTCGATTTAATGTTACGGAGTTCTTTTGGCCAATATCTTGT
TGCAAATGGATATGAATTTTCAGGATTAGAACTACACAACAGA---GGAGAATCGCTTCTTCAGCATCTCAAGGAGCTGGGCATGCCAGAAGATCAGCTGAAAGTCCTCGGGGACCTCCCCAGTCCAACACCTCTCGGTCTACGCCCCGCTGCA---------------------------
------------TTCCTCGTATACCTTAGCCTGTTCTTGCCTAAGTTAGTCACTGGGACTGTAGCATGCATTAAAAAAGTTAAACGGACTTACCAACAACTGGAGAGCCAAGGTATCAATACA---------------------
ATCCCTAGAGGGTGGATGACCCCTGTTATGATGAACCTCACATATCGTGCCTTACGCCTCTGTTTCTTCATTCGATTCTCCATCATACATCATGTGCTTCATCAACGGGCTGGAATAGATGCTGTTGACACAATTATGGCAGGAGCTGTGGAGCAGACTCGTTTTGCAG
GGCTGCTGATGGTTCGAAATATCTTAACATACATCATTACCAAGGGGAATGAAAAGGATAAATATCATATACACCCACTGGCTCAGGACAAAAGAATGAGGGAAGAGGTAACCCGATTTGAGTACTGCGTGAAGAACATCCAGCGGCACGGAATATATGCTCCCTT
TGCACGCGTCCTAGGGCTGGCGGGAGTTTCTCAGATAGAGCACGGGCAATTCCCGCGATTGACTGCCATTGCTCTCGGAGTGTCCTCTGTCTATCAGAACACACTCTTGGGGGTGAACATGGACATGCGCTACCGATCTCTCAAAGAGGCAGCCCATCAGGCTGAAT
TGGAGCTA---------CAAGCAATCCAGTGTAAGAAA---------
ATCAGACAAGACACTGACTTAGACGATGATGAGAGAGTGGTCCTCCTTAAATTTGCTGAAAGAAGCCAAAGTATCAACAAAGAACAGTGGCATCAGGAGGGGCTCGCAAGACAAGAGCCGCTCCAGAAGTTGACAGGGAGG---
>NC_076734_Fiwi_virus
ATGGCA------------------------------ACAGGGGGCGGTTTTTCGATAGACCTGGATCAAATCCTCAAT---CGGCGC---CAGCAAGGATCAGGC---AGCGGGATAGGGCAA---------------------
GCTGGTCTGGAACGTGTGCCCATATTTGTCCTGGCAAATGTGCCTACCCCGGCCGAGCTGAAGTTCCTGAGCGTGGCATGGTTGGTGGTGCTTGAAGACCCG---
GAGCCACCTGCCGTTGTTGTGAGCCTTCTGGTGGAGCTGACATGCTCCGCTTACTTTGACAATGTCACGGAGCTCCTGGATGGAATC---------
CAACCTCTGGTGGGTCCAGGGGGTTACACCTGTCGAGTGACACTCCAAGGGATCGGACAATTCCGTATACCTGGCGAG---------
GTAGAACTTGAGAGGCTGCTCTGTGCCTATTTCCGACCCAGGGCGACAGCCTCGTTAAGAAACCCGCAGAACATGGAGATGCTCAACTACAGAGGCGGTCTGACTCTCTGGCAAGTGATGCTGATTCAGCTCTGCCTGGCCAGTGGACACCTGTTCACAGGGAACC
CCTCTCTCGGCACATGGTTTAGCCGAAGGCTCTCCGGTCTCCAATTGAACACTCGGATAGTCCTGTCATTGACGCCGGCTGTGTTCAAAATGATATCAGCACCAGGCGGCTCACTCACAGTAGTGAGGGCAATGGTTGTGCAGGTGATGATGACCACGTTGGCCTCG
AGGCCCATTGGGGTGGCATTAGTCGAGGGT---------------------GTAAGGATGGCGTCGACATACTCCGAATGGGCAGGTATGGCCTGGGTGTTCCTCTCCCTTTCGTTGGTGATG------------------------
AGACCCGCCCACCCTATATTGCTTGTACCGACTGTTCGGCGTGAAGCTCTGGCCCTCTCCTCAGCGATTCAAGCTCTCGGAGCCGACAGGGACCGCCTACCCTTCGAGAGGATCCTTGGGGCATCTCAAGCCAGTGGTCTTAACGCGGGCAACTACCCGACACTCTCG
AGCGCTGCATACGGAGTAGCTGTGGTTACCTTCAGTAACTTTGGACAATACCACGGGACAGTGTACTCCAATGTGGTTTACAGGGCAGCTCAGTACCTCACGAGGATCATC---------ACTGCTGTGGACACTGGCCTACCTGATCCTCTT------------------------------------
---------------------------ACAGGTCTCCAAGGGCAGCAGATGGCCGATGCAGCCGCCCTGGTGACAACCGCCATGTCGGATGCAAGCCAGACCCAA
>NC_076735_Oberland_virus
ATGAGCTCT------------------------TCAACCCCTCCCGATATG---------GTTTCGGATCTTCTCAGC---ATGAGAGGGCTCCAA---ACTGGACCT---GAAGTGGGGGTC---------------------
AATTCGGCTCCTACACGCTATCTTTACATACTTGATGCACCCATCACAGTCCCAACTGCAGTCAGGCTCAACGCGCTTTGGGTTTTGGCTTTAGAGGATCCG---
AATCCACCTCCGGTCGTTGTGGAGCTACTAGTCCAACTCACAGCTAGTGCATTCTACTCAAACCCACTTGATCTCGTCTCCTCTACA---------
GAGGCGATGTTGGGACCAGGCAGTACGGTAATCCCTGTAGTGCTCCCCGCGTTGGCTTCCGTTGGTATCGCAAGAATG---------
CCACAAACTGAGAGGCAGATCAATCAATACTTCCAACAACTCTCAGCCCAGGTCACGAACACTCGAGGATTCGGGGCCGCCATAGCCGACGACGGAACACTTAGTGTGTGGCAGGTGATGTTGATACACCTCGTCTTGGTCAGTGGGCACTTATTCACGGGGAATC
CTCAGACTGGATCTTGGTACGCAAAAAGAAGACCAGGTCTAAACCTGAACACGAATGTCACCTTTGTGCTCTCGTCTGATTTTCTCCGAATTCTCGGAATGCCAGGGACCTCACTACCAGTCAACAGAGCTGTCATAGCAGCTGTTGCATTTGATGGTATGTCAAGTT
CCGCCATTGGCATCCCCCTAGTGGCCGGA---------------------ATATCACGAGCATCTCAGTACGTTGTGTTTGCGGGCCAAGCATGGCTGTTCCTGGCGTTAAAGTTGATCTGG------------------------
GAGGCGCGTCACCCTATCCTCCTCCTTCCCTCTGTTAGGAATGAGATGGAACAGCTTTTGCAAGCGATTGCACAGTTAGGGCCGAACCGCTTCCGTCTTCCATTCCAAAGGATCTTAGTGAGCGCTGTGGCGGACAGTTTCCAAGCAGGTCGTTATCCTCAGCTGTCT
AGTGCTGCGTATGGAGCTGCCAGAGCCTTGTTTACAAGCTTCACAGGCTTCCGAGGAGTTGCCTTCTCAGCAATCATTGAGAAGACAGCCTTCTTTCTGTCTCAGGCTCTC---------AATATGGTTGATCCGACGAAAGTTGACCCCATA------------------------------------
--------------------------ACCGGTCTCTCCGGTGATCAGCTCCAATTGGCCACTGAGAGAGTCAACCAAGCCATAACAGCAGCCTCCTCTCAAGGA
>NC_076916_Kander_virus
---------------------------------------------------------------------------------------------------------------------------------------------------------------------------------------------------------------------------------------------------------------------------------------------
---------------------------------------------------------------------------------------------------------------------------------------------------------------------------------------------------------------------------------------
ATGTTGGTCCAGCTGTGTCTCGCAAGTGGGCACCTTTTCACCGGGAACGCATCTCTGACCACATGGTTCAGTCGGCGGCTGAACGCGATGCAGTTGAACACAAGGATAACCCTGTCTCTTAACCCGGCTATCTTCAGGGTCATCGCAACCCCCGGTGGCTCACTCACA
GTGGTAAGGGCCATGATCGTACAGGTGATGATGACAACCCTGGGTTCCAGACCTGTCGGTGTAGCCTTGGTTGAGGCA---------------------GTGAGAACAGCATCAACATATGCAGAGTGGGCCGGGATGGCCTGGGTCTTCCTCTCACTCTCCCTCGTGATG---------
---------------
CGCACAGCCCACCCGATCCTGCTTGTACCATCCGTGCGCAGGGAGGCCCTAGCACTGTCAGCCGCAATTCAGGCCTTAGGTGCTGACCGGTACAGGTTGCCATTCGAGAGGATTCTAGGGGCCTCCCAAGCCTCAGGGCTCAATGCAGGGAACTACCCTACTCTCTC
CAGTGCAGCATACGGGGTTGCCGTGGTGACTTTCAGCAACTTCGGACAGTACCACGGGACGGTCTACTCAGCGGTCGTTTACAGGGCTGCGCAGTTCCTGACCCGGGTGATA---------ACAGCCGTTGACACCGGGAGCCCGGATCCACTG------------------------------
---------------------------------ACAGGCCTCCAAGGTCAGCAGATGGTGGATGCCACAGCGATGGTGACCACTGCAATGTCGGATGCAAGCCAAGGGCAA
>OQ186623_Loetschberg_virus
ATGAGC------------------------------ACCGATGATGATACC---------GTTACTGAACTTTTGTCAACCCCGAGGGGAGGGGCT---AGTGGAGTTGGTCCTGTTGGTGGT---------------------
AATGGTACGAACCCTCCTCTGGTGTTTCATTTGGCTGGACCATACTCCCAACGAACAGGAAGAATCCTGAATGCTTGCTGGCTATGCTTACTCACTGACCCG---
AACCCGCCTCCTATTGTGGTTCAAGTGTTGTCCCAAGTGACATTGAGCTGCTTTTTCAATGTCCCTGCTGATGCTATTGCTGCAACA---------
CAGGCAATTCTTGGCGAAAATGTCATGCCAATGCCTGTGGAGATGACAAGATTCGGCGAGGTCACCATCCCGAAAGAC---------
GCAGGGCTGGAGACCATGATGAACAGCTACCTGTCTCACTGGATCACTGTTGTGCAACCTAACCAAAATGACTGGGCCTGGGTTGCGTACAAGGATGGTCTTCAACTATGGCAGGTCATGCTCATCCACATTATTCTGTGCTCGGGCCACATGTTCACTGGGAACTC
AAACCTAGTGGCATGGTACCAGAGAAGGAGAACTGCACTGCTGCTCAACACCGCAGTCACTATGCAACTCTCATCAGGAGTCTTGCAAGGGCTTTCATGCACAGGCGGCTCCCTTATTGGTCTGAGAGGGTTGGTGGTTGCCTGTGTTGTGTCTGCTCAAGGGACAC
CTGCGTCTGGTTCTCTACTGATAAGCGGA---------------------GTGCTCAATGCTGCTAGCTATGTCAGATGGGCTGGAATGGCCTGGGTGTACCTGGCTTTGAACTTGGTCTTG------------------------
AGAACTCACAGCATTGTGCTGCTTTTACCCCATGTGAGGACAGAAGCTATGGCTTTCTTAACAGCAGTCAGGGCACTCGGCCCAGAGAGAACTAATCTTCAATTTGCACGAATGCTCTCATCACCTCTTGCTCACCAGTTCTCTGMAGGTTCATATCCTGCACTGTCTA

GTGCTGCATACGGTGTTGCATCTGCCACATTCTCCAGCTTCTCTGGATTTTCTGGAAGAGTCTTCTGTGAGACCATATACAAGGTTGCACGGAGGTTGACCCTGATGGTC---------ACAGCTGTCGACCTGAACATCCCGGACTCCTTG-----------------------------------------------------------------ACACAACTGAGTGGTCCAGCAATGGAGAGAGCAGTCTCACAGGTCAATGATGTCCTCAAGGAGAGCAGCGGGGGATCA
>CATLJU010000334_Borostomias_antarcticus
----------------------------------------------------------------------------------------------------------------------------------------------------------------------------------------------------TTAACCATTCTTCAGGACAAC---
AGGCCTCCTTCACAGGCAGTAACCTTCCTAGTTGAGATTACATGTTCCGCGTATTTCGACAAGCCGAAGGACATCTATGAGTCAGTA---------
AAAGACATGATCACACCAGGAGCCGTACCAACACCAGTGATTCTTGAAAGGTCCGGGGTCTTCAGAATCCCCTCCTCC---------
AGGACAGCTGAGGATAACTTTACTGATTATTTTAGAAAGGGCTGTACAGACCTAATGCCTCAAAAGAGCCTCATCAGAGAGATCACTCAGGAAGATGGTTTAGCACTTTGGCAGGTTGCTCTGATTCAACTCTGCCTTGTGTCTGGGCACTTGTTCACTGGTAACCAG
GCATATTACACCTGGTTCTCGAGGCGAATGAAGGGGATGCCAATTGAGACAACTGTGACTCAACAACTCACACAAGAGTTCCTCGGTATGTTGGTCACCCCATGGGGCTCACTTGCCATAGTCAGGGCAATGATTGTCCAGGTCAAGATCTCCACAATTGATCCACC
CTCAATAGGGATTCCACTCGTGGACGGG---------------------ATCAGAGCTGCATCAACTTATGTCGAGTGGTCTGGGATGGCCTGGGCCTACCTCGCACTAAGCATGGTGATG------------------------
AGGACAGCTCACCCTATCCTGCTTGCTCAATCCGTGAGAAAAGAGGCCATGCAACTGTCACGATTCATAGCGGGCTTGGGCAGAAACATCTTCCGACTACCATTCGAGAGGATACTTGGCCTTACTCAGGTCAGAGGGCTCAATGCAGGGAATTATCCTACCCTCTC
AAGTGCTGCCTATGGGGTAGCGGCAGCGACCTTTGAGAACTTCGGGTGCTTCCAGGGAACAATATACTCCCAGCGTATCTACAAGGCAGCCCAAGCGCTAAGACAGATCATA---------ACCGCATGTGATGCCTCTGTCCCAGACCCCCTC------------------------------------------------------------ACAGGTCTCCAGGGCGACCAAATGAGGGACGCAACACAACTGGTAACCAAGGCCTTGAAGGAG---------------
>CATLJU010001338_Borostomias_antarcticus
----------------------------------------------------------------------------------------------------------------------------------------------------------------------------------------------------CTGCGGATACTTCAAGACCCC---
AAGCCACCTACACAGGCTGTGAGCTTCATCACCGAACTCGCCTGTTCAGCATACTTTGATTGTCCAGGTGACATCTTTGCATCCATA---------
AAGGAAATAGTCACACCCGGAGCCATGATGACCCCAGTGGAGCTCAACAAAATTGGGACCTTTCGCATCCCCGGGAAT---------
CCAGCTGCTGAGGAATCAATGAGTGCCTACTTTAGACAAGGCTGCATGGCAATCATCAATCAGAAGACACTGATGGCAGAGGTGACCTGCGAGGATGGGCTCTCCATGTGGCAGTTAGCCCTTATCGAGCTGTGCCTTGTCTCGGGACATTTGTTTACTGGTAATCC
ATCTTACTTCAGGTGGTTCACCAAGAGACTGGCAGGCATGAATCTCCCAACAAGAATCAGCCTGGGACTAACTGCAGAGTTCCTCTCCATGTTGGTCATCCCAGGTGGTTCTCTCGCGATAGTTAGAGCAATGATTATCCAAGTGATGATTGCGACCATCAACACCCC
CTCCACTGGAATACCATTAGTCGGTGGG---------------------ATAAGGACCGCTTCAACCTATGTCGAGTGGTCCGGAATGGCCTGGACCTTCTTGGCCCTGACACTGGTCATG------------------------
AAAAATGCACATCCGATACTTCTAGTCTCATCAGTGCGACGTGAGTCCATCCAACTTTCAAATCTCATCCGGGGCATCGGGTCAGATCTCTCTCGTCTTCCCTTTGAAAGGATCCTTGGACTCTCACAGATGGCCGGGTTGAATGCCGGGAATTATCCAACACTCTCTA
GTGCTGCTTATGGGGTAGCCACAGCCACATTTGAGAACTTCAGTGGGTTCAAAGGAACAATCTACTCGCAGAGGGTCTACAAGGCTGCTCAAATCCTCACCCAGATTATC---------ATTGCGAACGATGGAAGTTTACCTGACCCATTG------------------------------------------------------------ACTGGGCTCTCAGGGAACCAAATGAGGGAGACTGCGGCACACCTCACCAAGGCACTGCAGGATGCAACCGCCGCAACG
>JABUMV010022313_Paedocypris_sp
ATGGACCCC---------------------------------------------AAGGACCTACCTCATCTTGCCTCA---------------------CTTAGGCCT---TCGGCTGAGTTT---------------------GCCTATACAACAAGACAGAGGCTCATACTG------------
TACACCGCTAAAGCAACCCCGTGGCTCCTGGCGTGCCTAATTTCAGTGCTTTTTGATCCGGATGCGAGCCGCCAGGAGGTGGCCTCCGCATTCTGCTCGATCTTCTTAACATACTTCCTGAATCGGCCCCTCACCGACTGCATCGAAGTACCCGCAATGGCTGAGCTCA
CTGGTCTGGGCTACGCCGTCGATGTCACGGACCTGACCCCGGGGGACCCAAAGCGAACTTTGAAGCTT------------ACCAAATCCCCCCTGATCAATAGGCACATGAGGAGTGGGGAGCTACTTGAAGTCACTGAACGCTCACTTGAGCGC---------------------------------------TTTGTCCTATATGTGGCCGCCATGCTTCCAAAGCTGATGATCTCCGCTGACACTGTGCAAATTAAGCTAGTCAAGGTCCAAAAGAACCTAAGAAGTCAATCACAGGAAATG---------------------
TTCCCGGACCTCTGGATGAACAAGGCAGTTATAACCATTGTCCGAAATCATTTCAGAGATTGCTTCCCGGGAAAATATCTCTTGATAAAGCTGACGAGCACACGGGTCTCTGGGGGGATTGCATACGGGACACAGATACACACTCTTGTGGACCAGGCCAAGATGTC
TGGGTTAGTTCTGGTCAAGCTGTTCAACGATCACCTCATCACACACGACGGGCCACATGGGAGCAAACAATGGCATCCAATCCTCTCGATGGAGACCCTGACCCCCGAGGCGGTTGCCCTAGTCAAGGCTATCAAAGCGCTCAGCGACCATGGTGACCTGGCTCCAT
ATGCCAGAATCTTGGCATTATCAGGCGTTGAACAGATCGAGTACGGGAAGTTCCCTAAGCTAGCGGCGGTTATAATCGGGATAGGGAAGGCACATAATCCAACCCTTAACATGATGACCATGGCCTCGGCGTATACGAACCTAGCGAATCTAGCCTACACCTATGA
GATCAAGCGATTGGCGTCCACCAAGCGAAACCCCGAGCGTCCACCGTTTTTG---CGTCCTCCGCAAGCGCCGAGCAGCTCCAGCAGATACATGACCTCC----------------------------------------------------------------------------------------------

**Data S5. Multiple sequence alignment of VP35 and VP35-like sequences (nucleotides) from Filoviruses and open reading frame (ORF) filovirus-like elements in vertebrate genomes.** The data were used for the analysis in Fig. S7 and related analyses.

```
>Acomys_dimidiatus_OU015376
------------------------------------------------------------
------------ATGGCAACCGGGCCAGAAAGAAGTGGTCGGATCTCAGAGCAACTGATG
ACAGGCCAGCTCTCCATCGTTGCCTTTTTTAGCTCTGGGACTGAA---------------
---GGCTCTCTGACCAGTCCTGCAATCAAAACCTTTAGGAGTGCCGAA---------GGT
CAT------------------------AGTGCTGCCTCACCCGCTGAAAATACAGTT---
------GCAGGTGAATTACAAGATACCCGAAAACAAATTCATGAGATTACAAGGAATGTG
AGTAAGGTACTT------------GCCCAACAAGAGGAACTGACCGAAACACTCAACAGC
AGTCTCAAAGATCTGAAA---------------------GCAATAGCAGCCACGGCTGAG
ATGGTGGAGGGGATCAGAAACTGTCACGCTGAGTTGCTTGCTAAGTATGATTCATTAGTA
ACGACCACGGGCAGGTTGACGGCTACAGCGGCAGCAAGCGAGGCC---------------
---------------------------------------TCCTTGGAAGAGCAAGGAAGT
GCACCCACTGTCCCCTCTTTGTATAAGGTTGCCGTCCTTGAACAACTGTACTGTGACAAA
GACTCCAGA---------GATCCA---------GATGTCAAGTCCGCCTTACATGCCCTG
GATGAGGCTTACACCTTAAGAGAGGAGACATTTGCCAAGCCTCAGATGTCAGCCAGAGAT
TTTGCCAACCTGTTATATGGTCACCTCCCGGGCAACGACACACCCTTTCACCAATTGGCC
CAGGTTATCGTCAAGATAGCCAAACGGTCAGAGGCTATCGACATCTGCGATGCAGAATTC
CAGGCAAGTTTGGCCCACGGTGCCTCCCCTCAAGAAGCCCTGGTACAAATGATTCAACAA
ATCCCCTGCTTGAAGGAGGCCAGTCCCCCAGAAGTTGAGATCAAGAGCCAAACGGAAATC
CCACAAGCCTGCCAGAAGAGCCTGCGTGCACTTCCGCCTCATCTGCGG---CTTGGGCAA
GGCTGGATCTGTGTTTATTGTCTCCCGGATGGCAAGAAATTGGGTCTGAAGGTT
>Acomys_percivali_OU015744
------------------------------------------------------------
------------ATGGCAACAGGGCCAGAAAGAAGTGGTCGGATCTCAGAGCAAATGATG
ACAGGCCAGCTCTCCATTGATGACTTTTTTAGCTCTGGGACTGAA---------------
---GGCTCTCTGACCAGTCCTGCAATCAAAACCTTTAGGAGTGCCGAA---------GGT
CAT------------------------AGGGCTGCCTCACCCTCTGAAAATACAGTT---
------GCAGGTGAATTACAAGATATCCGAAAACAAATTCATGAGATTACACGGAATGTG
AGTAAGGTACTT------------GCTCAACAAGAGGAACTGACCGAAACGCTCAACAGC
AGTCTCAAAGATCTGAAA---------------------GCAATAGCAGCCACGGCTGAG
ATGGTGGAGGGGATCAGAAACTGTCACACTGAGTTGCTTGCTAAGTATGATTCATTAGTA
ACAAGCACGGGCAGGTTGACTGCTACAGCGACAGCAAGCGAGGCC---------------
---------------------------------------TCCTTGGAAGAGCAAGGAAGT
GCACCCACTGTCCCCTCTTTGTATAAGGTTGCAGTCCTTGAACAACTGTACTGTGACAAA
GACTCCAGA---------GATCCA---------GATATTAAGTCCGCCTTACATGCCCTG
GATGAGGCTTGCACCTTAAGAGAGGAGACGTTTGCCAAGCCTCAGATGTCAGGCAGAGAT
TTTGCCAAACTGTTGTATGGTCACCTCCCGGGCAACGACACACCATTTCACCAATTGGCC
CAGGTTATCGCCAAGATAGCCAAACGGTCAGAGGCTATCGACATCTGCGATGCTGAATTC
CAGGCAAGTTTGGCCCACGGTGCCTCCCCTCAAGAAGCCCTGGTACAATTGATTCAACAA
ATCCCCTGCTTGAAGGAGGCCAGTCCCCCAGAAGTTGAGATCAAGAGCCAAACGGAAATC
CCACAAGCCTGCCAGAAGAGCCTGCGTGCACTTCCGCCCCATCTGCGG---CTTGGGCAA
GGCTGGATCTGTGTTTATTGTCTCCCGGATGGCAAGAAATTGGGTCTGAAGGTT
>Acomys_kempi_OU015357
------------------------------------------------------------
------------ATGGCAACCGGGCCAGAAAGAAGTGGTCGGATCTCAGAGCAACTGATG
ACAGGCCAGCTCTCCATCGTTGCCTTTTTTAGCTCTGGGACTGAA---------------
---GGCTCTCTGACCAGTCCTGCAATCAAAACCTTTAGGAGTGCCGAA---------GGT
CAT------------------------AGTGCTGCCTCACCCGCTGAAAATACAGTT---
------GCAGGTGAATTACAAGATACCCGAAAACAAATTCATGAAATTACAAGGAATGTG
AGTAAGGTACTT------------GCTCAACAAGAGGAACTGACCGAAACGCTCAACAGC
AGTCTCAAAGATCTGAAA---------------------GCAATAGCAGCCACGGCTGAG
ATGGTGGAGGGGATCAGAAACTGTCACGCTGAGTTGCTTGCTAAGTATGATTCATTAGTA
ATGACCACGGGCAGGTTGACGGCTACAGCGGCAGCAAGCGAGGCC---------------
---------------------------------------TCCTTGGAAGAGCAAGGAAGT
GCACCCACTGTCCCCTCTTTGTATAAGGTTGCCGTCCTTGAACAACTGTACTGTGACAAA
GACTCCAGA---------GATCCA---------GATGTCAAGACCACCTTACATGCCCTG
GATAAGGCTTACACCTTAAGAGAGGAGACATTTGCCAAGCCTCAGATGTCAGCCAGAGAT
TTTGCCAAACTGTTATATGGTCACCTCCCGGGCAACGACACACCATTTCACCAATTGGCC
CAGGTTATCGCCAAGATAGCCAAACGGTCAGAGGCTATCGACATCTGCGATGCAGAATTC
CAGGCAAGTTTGGCCCACGGTGCCTCCCCTCAAGAAGCCCTGGTACAAATGATTCAACAA
ATCCCCTGCTTGAAGGAGGCCAGTCCCCCAGAAGTTGAGATCAAGAGCCAAACGGAAATC
CCACAAGCCTGCCAGAAGAGCCTGTGTGCACTTCCGCCCCATTTGCGG---CTTGGGCAA
GGCTGGATCTGTGTTTATTGTCTCCCGGATGGCAAGAAAGTGGGTCTGAAGGTT
>Lloviu_cuevavirus_NC_016144
ATG---------------------------------------------------------
------------------------------------GACAAAGTGACGGATGCCATCATG
ACTGGGCGATTATCACTCGACACAATCCCTGGCTTTCAATGCTCTGCGCCG---------
------------ACCTCCAGGAGCAGGCCCACCGGGCGAAAGTGTCCACCGTGGGAAGGA
CCACTCATTGAGCCCAAGACTTCCGCCAACAAATCAACACAAACTGAGAACATCTAC---
------------CAAAGCGACCAGGTCCTACGCGAAATCAAACAACTGAGCTCAATTCTA
GCCCACCAAATC------------ACAACCATGGAGAGGCTAGAGGGTTCACTTCAAAGC
ATACACCAACAGCTTCAG---------------------GTGGTCGGGGGCATGGCACAT
AACCTAACATCCTTGACAACCCTGTGCACAGAAATGGTTGCCAAATATGACTTCCTTGTC
ATGACCACCGGACGAGCAACTGCAACAGCGGCGGCAACAGAGGCT---------------
---------------------------------------TACTGGAAAGAACATGGGAAA
GCCCCACCTGGCCCAGCCTTGTTCGAAGCAGATGCACTTAAGGCACGAATCCAAGAAAAT
CCCAAAAGT---------GTCCCG------ACTGATGTTAGAGAGGCCTTTGACCGTCTA
GAGAAGACTGAGGAGGTAACAGAGAGGACGTTCGGAAAGCCAACCATCTCAGCTAAGCTC
CTCAAGGAACTTCTCTATGATCATCTTCCTGGATATGGGACTGCCTTCCATCAGCTTGCC
CAAGTCGTATGCAAGGTCGGGAAAGACAATGATCTCCTGGATATCATACATGCTGAGTTC
CAGGCCAGCCTAGCAGAGGGTGACTCTCCGCAATGTGCCCTGATTCAGATCACGAAGCGA
ATCCCCATCTTCGGGGAGACACCACCTCCCGTCATCTACATAAAAACAAAACAAGAAATA
CCCAAAGCATGCCAAAAAAGCCTCCGCCCTCTTCCCCCCAATCCTAAA---ATTGAACGG
GGCTGGGTATGCCATTATGTAACAGCAGATGGAAAGGAAATGGGCCTCAAGATA
>Mengla_dianlovirus_NC_055510
ATGTGG------------------------------------------------------
---------------------GACCCTAATTACATGAAAGGTGTTACGACTGATTTAATG
ACTGGTAAAATCCCGGTTGATGCAGTTTTCAATCAGAATCCTCTA---------------
---GGATCACTATACAAAAGAAGAAAGCCAAGGACAAACCAAGCA---------------
---ATTCAGGTTGGCCCAGAGATGAAGGAAAAATCAACAAATACTGATCAGCTTGTT---
---------GGCCAAGAATACAGACTGACTGAAGAAATAAGAGATCTTTTTACTAATATG
GAGCAAAAAATGAACTCCTTGATCATCCAAAGTATGGAGAATTCAGAAAGAATCAATGCT
CTTGAACAGCAGCTGAAGGAT------------CTTGTGCCTGTAATCAAAATGGGAAAA
AACATAGAATATCTTACAAAGCAGGTGTCCGAACTACTTGCAAAATATGAGCATCTTGTT
ATCTCAACAGGGCGTACAACTGCACCGGCTGCTGCATTTGATGCT---------------
---------------------------------------TACTTGAAGGAAAATGGAAGA
CCTCCATCAAATCCGGCTATATTTAAGGATCTTGGTGTTGCATGTTCTATGACCAAACCT
```

TCAACTAAT---------GAGAGT------ACTGAAATCTCAGATGCTGGCAAAAAAGTG
TCTCGAGTTTTGGAACTAAATGAAGAAACATTTGCAAAGCCTCAGATGAATGCAAAGGAT
CTTGCTCTCCTAATATTCTCACACTTACCAGGAAATAATACTCCATTCCATATTTTGGCT
CAAGTAATTGCAAAAATTGCAGCAAAAGATGGAGAAACTGGTGCCCTACTAGACACATTC
CATCAGTATCTAAGTGAAGGGGATAATGCTCAGGCTGCTCTGACTAGGATAGTGAGACAA
GTAGGAATATTCACTGGTAGACAGCCTCCAACTCTTAAAATCAAGAGTCTTACATTGGTC
CCGAGACCTTGTCAAAAAAGCCTACGAGCTGTTCCTCCAAAACCTCAA---CTGGATAAG
GGTTGGGTTTGTATCTATGAATCAGAGGATGGTGACCGGAAAGCATTAAAAATC
>Bombali_ebolavirus_NC_039345
ATGAATAGCCAACGAATAAAAGTCATCAATGCA---------------TCATCAACACAG
CCAAAAATTCAACCGAAACATGGGCCCGATCTATCCGGATGGATTTCAGAGCAACTCATG
ACCGGGAAGATTCCAGTAAATGATATTTTTTATGACACTGATAGCATATCAAGTTACACC
GGGATTCCTCCTCTAGCTGTGAAAACCCAAAAACCTAAAAGGGTC---------------
--------------------------GACAATGACACTCAAACTGACCCCGTTTGT---
------CAACACAGCTTTGATGAGGTTGTGCAGACATTAACCTCCTTGACAACTGTGGTG
CAGCAGCAAGCC------------CTTGCAACGGAGTCTTTGGAACAACGGATTTCTAGC
TTGGAAAGCAACTTTAAA---------------------CCAGTCCTTGATATGGCGAAA
ACCATAGCCTCTCTCCAACGCGCATGTTCAGAGATGGTGGCAAAGTACGATTTATTAGTA
ATGACCACTGGACGTGCAACAGCGACTGCTGCCGCCACCGAGGCT---------------
--------------------------------------TATTGGAAAGAGCATGGTCAG
CCTCCACCTGGGCCCTCTCTTTATGAAGAGGATGCGATTAGGCATAAAATTGAAACCATG
CAAGATGTA---------GTTCCT------CAAGCAGTTCAAGAGGCCTTCAAAAATTTG
GAGAGCACAACATCCTTGACTGAGGAAAACTTTGGGAAGCCATATATTTCAGCAAAGGAT
TTACGAGATATCATGTATGACAATCTCCCAGGTTTTGGGACAGCCTTTCATCAGCTAGTG
CAGGTAATATGTAAGATTGGGAAAGACAATGGATTGCTGGATACCATCCACGCTGAATTT
CAGGCAAGCTTGGCTGATGGAGACTCTCCACAATGTGCTCTCATCCAAATCACTAAACGG
GTACCAGCCTTTCAGGAGATCCCGCCTCCCACCATCCACATCAGGTCAAGGGGAGACATC
CCTCGAGCTTGTCAAAAGAGCCTCCGCCCAGTTCCCCCTTCACCCAAG---ATTGACCGG
GGCTGGGTATGCATCTTTCAACTACAGGATGGAAAAACCTTGGGCCTCAAAATC
>Bundibugyo_ebolavirus_NC_014373
ATGACCTCTAACAGAGCAAGGGTGACTTACAAC---------------CCACCACCAACA
ACCACAGGCACACGATCGTGTGGGCCGGAACTTTCCGGGTGGATCTCTGAGCAATTGATG
ACAGGCAAGATTCCGATTACCGATATCTTCAATGAAATTGAAACCTTACCTAGTATAAGT
CCCTCGATCCACTCCAAAATCAAAACCCCAAGTGTTCAAACACGC---------------
--------------------------AGTGTCCAGACCCAAACTGACCCAAATTGT---
------AATCATGATTTTGCAGAGGTTGTGAAAATGCTAACATCTCTAACCCTTGTCGTA
CAAAAACAAACC------------CTTGCAACTGAATCACTTGAGCAACGCATTACTGAC
CTGGAAGGTAGCCTGAAA---------------------CCAGTGTCTGAGATCACCAAG
ATTGTTTCTGCACTAAATAGATCCTGTGCAGAGATGGTGGCCAAATATGATCTTCTAGTA
ATGACGACTGGTCGTGCAACTGCCACTGCTGCAGCTACTGAAGCA---------------
--------------------------------------TACTGGGCAGAACATGGACGT
CCTCCACCGGGGCCCTCATTGTACGAGGAGGATGCAATCAGGACTAAAATTGGAAAACAA
GGGGATATG---------GTACCC------AAGGAAGTGCAAGAGGCCTTCCGTAATCTG
GATAGTACTGCCCTTCTAACGGAAGAGAATTTTGGGAAACCAGACATATCCGCAAAAGAC
TTGCGCAATATCATGTATGATCACCTCCCAGGTTTTGGCACAGCATTTCATCAACTAGTG
CAAGTTATCTGCAAGTTAGGGAAGGACAATTCCTCACTTGATGTAATTCATGCAGAATTT
CAGGCCAGCCTTGCTGAAGGAGACTCTCCTCAGTGTGCCCTGATTCAGATAACCAAACGG
ATTCCTATTTTCCAAGATGCAGCACCACCCGTAATCCATATTCGGTCACGCGGTGATATA
CCAAAGGCGTGTCAAAAGAGCCTCCGCCCTGTTCCACCATCACCAAAG---ATTGATAGG
GGTTGGGTATGCATATTCCAGCTACAAGACGGAAAAACACTCGGACTCAAAATC
>Reston_ebolavirus_NC_004161
ATGTAC--------------------------------------------------AAT
AATAAATTGAAGGTATGTTCAGGCCCAGAAACGACTGGATGGATTTCTGAGCAACTTATG
ACAGGTAAGATTCCAGTAACTGATATATTCATTGATATTGATAACAAGCCAGATCAAATG
GAAGTCCGACTCAAACCATCATCAAGGAGCTCAACAAGAACTTGT---------------
--------------------------ACAAGTAGCAGTCAGACGGAGGTCAACTAT---
------GTACCTCTCCTTAAAAAGGTTGAGGATACATTAACTATGCTAGTGAATGCCACC
AGTCGTCAGAAT------------GCTGCAATCGAGGCCCTTGAAAACCGCCTCAGCACA
CTTGAGAGTAGCTTAAAG---------------------CCAATCCAAGACATGGGTAAA
GTGATTTCATCATTGAATCGCAGTTGTGCCGAAATGGTTGCAAAATATGATCTTCTAGTT
ATGACAACTGGACGGGCTACTTCAACTGCAGCTGCAGTAGATGCG---------------
--------------------------------------TATTGGAAAGAGCACAAACAG
CCACCACCAGGGCCAGCGTTGTATGAAGAGAATGCGCTTAAAGGAAAAATCGATGATCCA
AACAGCTAT---------GTACCA------GATGCTGTGCAAGAGGCTTACAAGAACCTT
GACAGTACATCGACCCTGACCGAGGAAAATTTTGGGAAACCTTATATATCTGCTAAAGAC
CTGAAGGAGATCATGTATGATCATCTACCTGGTTTTGGGACTGCCTTTCACCAACTTGTT
CAAGTGATTTGTAAAATAGGAAAGGATAACAACCTTTTGGACACAATCCATGCTGAGTTC
CAGGCAAGTCTAGCAGATGGTGACTCTCCCCAATGTGCACTCATACAGATAACCAAAAGG
GTCCCAATCTTTCAGGATGTGCCGCCCCCGATAATCCATATTAGATCCCGTGGTGACATC
CCACGAGCATGCCAAAAGAGTCTCCGACCAGCACCACCATCACCCAAA---ATTGATCGT
GGTTGGGTTTGTTTGTTTAAGATGCAAGATGGTAAAACGCTTGGACTTAAGATC
>Sudan_ebolavirus_NC_006432
ATGCAG--------------------------------------------------CAG
GATAGGACTTATAGACATCATGGACCCGAAGTGTCTGGCTGGTTTTCTGAGCAATTAATG
ACCGGCAAAATACCGCTAACAGAGGTGTTTGTTGATGTTGAAAACAAACCAAGTCCTGCC
CCGATAACCATTATTAGTAAGAATCCCAAGACAACACGTAAAAGT---------------
--------------------------GATAAGCAAGTCCAAACAGATGATGCCAGT---
------AGCTTATTGACAGAAGAAGTCAAGGCTGCCATAAATTCGGTGATATCAGCTGTG
CGTCGGCAAACC------------AATGCTATTGAATCACTAGAAGGTCGAGTAACAACT
CTTGAGGCCAGCTTAAAA---------------------CCCGTTCAAGACATGGCAAAG
ACCATATCATCCCTGAATCGCAGCTGTGCCGAAATGGTTGCAAAATACGACCTACTGGTG
ATGACCACTGGGCGAGCAACTGCCACTGCTGCAGCAACAGAAGCA---------------
--------------------------------------TATTGGAATGAACATGGACAA
GCACCTCCAGGCCCATCATTGTACGAGGATGATGCTATTAAGGCTAAATTGAAAGATCCG
AACGGGAAG---------GTTCCA------GAAAGTGTCAAACAGGCCTACATAAATCTA
GATAGCACAAGTGCCCTCAATGAGGAAAATTTCGGGCGACCTTACATTTCAGCAAAAGAT
CTCAAGGAAATCATCTATGACCATCTCCCAGGATTTGGGACAGCTTTTCATCAGTTGGTG
CAGGTTATCTGCAAAATTGGTAAGGATAATAATATCCTAGACATAATTCATGCAGAATTC
CAAGCAAGCTTGGCTGAGGGAGACTCCCCCCAGTGTGCATTAATCCAGATAACAAAACGG
ATCCCTGCTTTCCAAGATGCCTCTCCTCCAATTGTGCATATCAAGTCTCGAGGAGATATA
CCCAAAGCCTGTCAGAAAAGCCTCCGGCCGGTCCCACCGTCACCAAAG---ATCGATAGA
GGTTGGGTCTGTATTTTTCAATTCCAAGACGGGAAGGCCCTTGGGCTAAAAATA
>Tai_Forest_NC_014372
ATGATTTCCACTAGGGCTGCAGCAATCAATGAT---------------CCTTCATTACCA
ATCAGAAACCAGTGTACACGTGGCCCTGAACTATCAGGATGGATCTCCGAACAATTAATG
ACAGGCAAAATTCCGGTACATGAAATCTTCAACGACACTGAGCCCCACATAAGCTCAGGG
TCCGACTGCCTTCCCAGACCCAAAAACACGGCCCCCCGGACTCGC---------------
--------------------------AACACCCAGACACAGACCGATCCGGTTTGC---
------AATCACAATTTTGAAGACGTTACACAAGCACTAACATCATTAACCAATGTCATA
CAAAAACAGGCT------------CTTAACTTAGAGTCTCTCGAACAACGCATCATAGAT
CTAGAGAATGGCTTAAAG---------------------CCAATGTATGACATGGCTAAA
GTCATTTCTGCATTGAATAGATCTTGTGCTGAGATGGTAGCAAAATATGATCTCCTGGTG
ATGACAACTGGCCGCGCAACCGCCACCGCCGCTGCAACTGAGGCT---------------
--------------------------------------TATTGGGAGGAACATGGACAA
CCACCACCTGGACCATCACTTTATGAAGAGAGTGCGATTAGAGGCAAGATTAACAAGCAA
GAGGATAAA---------GTACCT------AAGGAAGTTCAAGAAGCTTTTCGTAATCTG

GACAGTACCAGCTCACTAACAGAAGAGAACTTTGGCAAGCCAGATATATCTGCAAAGGAC
CTACGAGACATCATGTATGACCACCTACCAGGCTTCGGTACGGCTTTTCACCAACTGGTC
CAGGTAATTTGCAAGCTAGGAAAAGACAATTCTGCATTGGACATTATTCATGCTGAGTTC
CAAGCCAGCCTTGCTGAAGGTGATTCTCCCCAATGTGCCCTGATCCAAATAACAAAACGG
ATCCCCATCTTCCAGGATGCCACTCCGCCCACAATTCACATCCGCTCTCGTGGTGACATC
CCACGTGCCTGCCAAAAAAGTCTCCGTCCAGTTCCTCCATCACCAAAA---ATAGACAGA
GGTTGGGTTTGCATTTTCCAATTGCAGGACGGGAAGACACTTGGGCTCAAGATA
>Zaire_ebolavirus_NC_002549
ATGACA---ACTAGAACAAAGGGCAGGGGCCAT---------------ACTGCGGCCACG
ACTCAAAACGACAGAATGCCAGGCCCTGAGCTTTCGGGCTGGATCTCTGAGCAGCTAATG
ACCGGAAGAATTCCTGTAAGCGACATCTTCTGTGATATTGAGAACAATCCAGGATTATGC
TACGCATCCCAAATGCAACAAACGAAGCCAAACCCGAAGACGCGC---------------
---------------------------AACAGTCAAACCCAAACGGACCCAATTTGC---
------AATCATAGTTTTGAGGAGGTAGTACAAACATTGGCTTCATTGGCTACTGTTGTG
CAACAACAAACC------------ATCGCATCAGAATCATTAGAACAACGCATTACGAGT
CTTGAGAATGGTCTAAAG---------------------CCAGTTTATGATATGGCAAAA
ACAATCTCCTCATTGAACAGGGTTTGTGCTGAGATGGTTGCAAAATATGATCTTCTGGTG
ATGACAACCGGTCGGGCAACAGCAACCGCTGCGGCAACTGAGGCT---------------
---------------------------------------TATTGGGCCGAACATGGTCAA
CCACCACCTGGACCATCACTTTATGAAGAAAGTGCGATTCGGGGTAAGATTGAATCTAGA
GATGAGACC---------GTCCCT------CAAAGTGTTAGGGAGGCATTCAACAATCTA
AACAGTACCACTTCACTAACTGAGGAAAATTTTGGGAAACCTGACATTTCGGCAAAGGAT
TTGAGAAACATTATGTATGATCACTTGCCTGGTTTTGGAACTGCTTTCCACCAATTAGTA
CAAGTGATTTGTAAATTGGGAAAAGATAGCAACTCATTGGACATCATTCATGCTGAGTTC
CAGGCCAGCCTGGCTGAAGGAGACTCTCCTCAATGTGCCCTAATTCAAATTACAAAAAGA
GTTCCAATCTTCCAAGATGCTGCTCCACCTGTCATCCACATCCGCTCTCGAGGTGACATT
CCCCGAGCTTGCCAGAAAAGCTTGCGTCCAGTCCCACCATCGCCCAAG---ATTGATCGA
GGTTGGGTATGTGTTTTTCAGCTTCAAGATGGTAAAACACTTGGACTCAAAATT
>Marburg_marburgvirus_KC545388
ATGTGG------------------------------------------------------
---------------------GACTCATCATACATGCAACAAGTCAGCGAGGGGTTGATG
ACTGGAAAAGTTCCCATAGATCAAGTGTTTGGTGCCAATCCCTTA---------------
---GAGAAGTTATACAAGAGAAGGAAACCAAAAGGCACAGTTGGA---------------
---CTACAATGTAGTCCTTGTCTAATGTCAAAGGCGACAAGCACTGATGATATTGTT---
---------TGGGACCAACTGATCGTGAAGAAAACACTAGCTGATCTACTTATACCGATA
AATAGGCAAATATCGGACATTCAAAGCACTCTAAACGAAGTAACAACAAGAGTCCATGAA
ATTGAGCGGCAATTACATGAG------------ATAACCCCAGTGTTAAAAATGGGAAGG
ACATTGGAAGCAATTTCCAAGGGGATGTCAGAAATGTTAGCCAAATATGACCACCTCGTA
ATTTCAACCGGAAGAACCACTGCACCAGCTGCTGCCTTTGATGCT---------------
---------------------------------------TACTTAAATGAGCATGGTGTC
CCTCCCCCCCAACCCGCAATTTTCAAAGATCTTGGGGTTGCTCAACAAGCTTGCAGCAAG
GGGACCATG---------GTTAAA------AATGAAACAACAGATGCAGCCGACAAGATG
TCGAAAGTTCTTGAACTCAGTGAGGAGACGTTCTCCAAGCCAAACCTTTCAGCTAAGGAT
TTAGCCCTTTTGTTGTTTACCCATCTACCCGGCAACAACACTCCATTCCATATCCTAGCT
CAAGTCCTTTCAAAAATTGCTTACAAGTCAGGAAAATCCGGAGCATTTTTGGATGCATTT
CACCAGATTCTAAGTGAAGGAGAGAATGCTCAGGCAGCATTAACTCGACTAAGCAGAACA
TTTGATGCCTTCCTCGGAGTAGTTCCTCCAGTGATAAGAGTCAAAAACTTCCAAACAGTC
CCTCGCCCATGTCAAAAGAGTCTTCGGGCTGTCCCTCCCAACCCAACA---ATTGACAAA
GGATGGGTCTGTGTTTATTCATCTGAGCAGGGTGAGACACGGGCCCTGAAAATC
>Tapajos_virus_NC_076535
ATGGCCCTTAGCCTTACCTTGGACGACAGAGATATCGACCCAGTCTCTAACCACATATCT
GCCACATCTCTATGGGATGATGCATCACAGTTCACATCTTCGTTGGCTGAACAGGTGCTC
ACAGGAAATTTAAGTTTCGATCAAATAGTGGAAGCCTCCACAAAA---------------
---ACCCAGCAACCCCCTCCGACCACCCCCAACTCAACAACCCAC------------ACC
CACTCCGATGGCTCACCAACGCCGCGCTCTCAAAACAGCAATATGAAAGAT---------
------------------------------------------GACATTAACAATATTACC
AATCAGAATATA------------GACTCATATCATATTGCACAGGGTGAGATAAGCTCC
ATTAAAGCCGAGCTGCAT---------------------GGACTGCGATCTGATCTTAAA
ATCCTCCCAGAGATTCAGAATCTCTTGTCTACACTTATTGCAGACTTTAGGACTTTGTTT
GTGATGACTGGCCCTGCGACCTCCACCATTGCGGCCGCACAAGCT---------------
---------------------------------------TATCGCGAAAAACATGGAACT
TTACCTTCAGGTCCGATTGTTGGAACGCTAACCGATCTACAATTGACCGTTGATGAAACA
CTCAAAGGTGAGTTTCATGCACCTGGCTATAAAGATACGAAGAACACACTTGTCAAAACA
GCAGCCGCTTTGAACACAGAGCCATCTCACTTTTCATCCGAACCGTCTACTGCCAAAGAA
CTTGAAGCTATATTTCAAGCACACCTTCCAGGATACCAGACACCTTTCCACATGTTAGCA
CAATCTTTAGCCAAAGTATGTTTTGTGCGAGATATGGTGGGGACAACCTTGAAAATGTTC
CACTATCAAATGGCTGCAGGCGAGTCAGCACACGCAGCTTTAATCTATCTGGCAAGAGAC
AAAGAATTCTTTCGTGAGGACACTGTTCCTAGAAAAAGAATCACCGATCAGGAGGAGATC
CCAAAAGCTTGTCGCAAGTCAATACACAGGGTGCCTGGCGGAACCGGCACCTTTGGGCAA
GGATGGGCCTTGATTTACGATGCACCCGGTGGT---GTGCTGGGAATCAAGATA
>Myotis_yumanensis_JAPQVU010000005
------------------------------------------------------------
------------------------------------------------------------
---------ATGTCCCTGGAGCAGTGCATC------------------------------
------------------------------------------------------------
---------------------------GAACAGATAAGTAAGCTCACCGATCGCTGT---
------------------GATAGAATCAAAGAAGCCATGACATCTCTTGTAAGCTGCATG
GAAAAGCAGTTT------------GTTGTAATGGATCATCTCCTAGCTGCCCTGAAGGAG
ATAAAAGCA---------GATCAAGTCGATTTTAGTCAGAGTCTTCTGTCTACGTCTTCT
AAGGTTAATCAACTAGTAGAGAATTCGTCTGAGCTTTTAGCCAAGCTTAGTTATTTGCCT
GTAATGTCAGGACCTGCAACATCCACTCTCGAGGCAGCTGGAGCA---------------
---------------------------------------AACACGCAGGAGCATAGAAGG
CCTCCCCCAGGGCCCATCCTAGCAACCCTGGAGCGACACGGAGCAAGACTGACAGATACT
CTCACTTCTGAT------ATTCCAGGGTACGTGAAGGCTGCTGAGGCCGAGAAGAAAATG
CATACTGCACAGACTGTCACTGGGGAGAGTGTCTCTCGGCTGCCTCTAGTTCCCACCGAA
TTCGTACGAGTCCTCACAAGTTATCTGCCAGGACCGCGCACTGCATTTCATGAATTAGTA
TCGGCAATCGCTTTGGTGAGCCGAGACTCTCATGATCTACAGGTAGCCATGGACCATTTC
AATCGAGAGCTAACGGATGGTTTCTCAGCTCATGCTGCCATAATATCCATCACTCGGAGA
TGTGAGTATTTTCGGAACTGCGAAGCTCCGACAATGCAGGTAACTTCGAAGAGCCAGATT
CCAGAAGCATATCATGGCAGACTTAGCGATGTACCGGAGGGTCCCAAAACCCTAGGACAA
GGATGGATATATATATATCTAACTCCTGAAGGA---AGCCTCGGGTTAAAGATT
>Myotis_lucifugus
------------------------------------------------------------
------------------------------------------------------------
---------ATGTCCCTGGAGCAGTGCATC------------------------------
------------------------------------------------------------
---------------------------GAACAGATAAGTAAGCTCACCGATCGCTGT---
------------------GATAGAATCAAAGAAGGCATGACATCTCTTGTAAGCTGCATG
GAAAAGCAGTTT------------GTTATAATGGATCATCTCGTAGCTGCCCAGATGGAG
ATAAAAGCA---------GATCAAGTCGATTTTAGTCAGAGTCTTCTGTCTACGTCTTCT
AAGGTTAATCAACTAGTAGAGAATTTGTCTGAGCTTTTAGCCAAGCTTAGTTATTTGCCT
GTAATGTCAGGACCTGCAACATCCACTCTCGAGGCAGCTGGAGCA---------------
---------------------------------------AACACGCAGGAGCATAGAAGG
CCTCCCCCAGGGCCCATCCTAGCAACCCTGGAACGACACGGAGCAAGACCCACAGATACT
CTCACTTCTGAT------ATTCCAGGGTCCGTGAAGGCTGCTGAGGCCGAGAAGAAAATG
CATACTGCACAGACTGTCCCTGGGGAGAGTGTCTCTCGGCTGCCTCTAGTTCCCACCGAA

TTCGTACGAGTCCTCACAAGTTATCTGACAGGACCGCGCACTGCATTTCATGAATTAGTA
TCGGCAATCGCTTTGGTGAGCCGAGACTCTCATGATCTACAGGTAGCCATGGACCATTTC
AATCGAGAGCTAATGGATGGTTTCTCAGCTCATGCTGCCATAATATCCATCACTCAGAGA
TGTGAGTATTTTCGGAACTGCGAAGCTCCGACAACGCAGGTAACTTCGAAGAGCCAGATT
CCACAAGCATGTCATGGCAGACTTAGGGATGTACCGGAGGGTCCCAAAACCCTAGGACGA
GGATGGGTATATATATATCTAACTCCTGAAGGA---AGCCTCGGGTTAAAGATT
>Myotis_brandtii
------------------------------------------------------------
------------------------------------------------------------
---------ATGTCCCTGGAGCAGTGCATC------------------------------
------------------------------------------------------------
---------------------------GAACAGATAAGTAAGCTCACCGATCGCTGT---
------------------GATAGAATCAAAGAAGCCATGACATCTCTTGTAAGCTGCATG
GAAAAGCAGTTT------------GTTATAATGGATCATCTCGTAGCTGCCCAGATGGAG
ATAAAAGCA---------GATCAAGTCGATTTTAGTCAGAGTCTTCTGTCTACGTCTTCT
AAGGTTAATCAACTAGTAGAGGATTTGTCTGAGCTTTTAGCCAAGCTTAGTTATTTGCCT
GTAATGTCAGGACCTGCAACATCCACTCTCGAGGCAGCTGGAGCA---------------
---------------------------------------AACACGCAGGAGCATAGAAGG
CCTCCCCCAGGGCCCATCCTAGCAACCCTGGAGCGACACGGAGCAAGACCGACAGATACT
CTCACTTCTGAT------ATTCCGGGGTATGTGGAGGCTGCTGAGGCCGAGAAGAAAATG
CATACTGCACAGACTGTCACTGGGGAGAGTGTCTCCCGGCTGCCTCTAGTTCCCACCGAA
TTCGTACGAGTCCTCACAAGTCATCTGACAGGACCCCGCACCGCATTTCATGAATTAGTA
TCAGCAATCGCTTTGGTGAGCCGAGACTCTCATGATCTACAGGTAGCCATGGACCAGTTC
AATCGAGAGCTAAAGGATGGTTCCTCAGCTCATGCTGCCATAATATCCATCACTTGGAGG
TGTGAGTATTTTCGGAACTGCGAAGCTCCGACAACGCAGGTAACTTCGAAGAGCCAGATT
CCACAAGCATGTCATGGCAGACTTAGGGATGTACCGGAGGGTCCCAAACCCCTAGGACGA
GGATGGGTATATATATATCTAACTCCTGAAGGA---AGCCTCGGGTTAAAGATT
>Myotis_septentrionalis
------------------------------------------------------------
------------------------------------------------------------
---------ATGTCCCTGGAGCAGTGCATC------------------------------
------------------------------------------------------------
---------------------------GAACAGATAAGTAAGCTCACCGATCTCTGT---
------------------GATAGAATCAAAGAAGCCATGACATCTCTTGTAAGCTGCATG
GAAAAGCAGTTT------------GTTATACTGGATCATCTCGTAGCTGCCCAGATGGAG
ATAAAAGCA---------GATCAAGTCGATTTTAGTCAGAGTCTTCTGTCTACGTCTTCT
AAGGTTAATCAACTAGTAGAGGATTTGTCTGAGCTTTTAGCCAAGCTTAGTTATTTGCCT
GTAATGTCAGGACCTGTAACATCCACTCTCCAGGCAGCTGGAGCA---------------
---------------------------------------AACACGCAGGAGCATAGAAGG
CCTCCCCCAGGGCCCATCCTAGCAACCCTGGAGCGACAAGGAGCAAGACTGACAGATACT
CTCACTTCTGAT------ATTCCAGGGTACGTGAAGGCTGCTGAGGCCGAGAAGAAAATG
CATACTGCACAGACTGTCACTGGGGAGAGTGTCTCTCGGCTGCCTCTAGTTCCCACCGAA
TTCGTACGAGTCCTCACAAGTTACCTGACAGGACCGCGCACTACATTTCATGAATTAGTG
TCGGCAATCGCTTTGGTGAGCCGAGACTCTCATGATCTACAGGTAGCCATGGACCATTTC
AATCGAGAGCTAACGGATGGTTTCTCAGCTCATGCTGCCATAATATCCATCACTCAGAGA
TGTGAGTATTTTCGGAACTGCGAAGCTCCGACAATGCAGGTAACTTCGAAGAGCCAGATT
CCAGAAGCATATCATGGCAGACTTAGGGATGTACCGGAGGGTCCCAAAACCCTAGGACAA
GGATGGGTATATATATATCTAACTCCTGAAGGA---AGCCTCGGGTTAAAGATT
>Myotis_albescens
------------------------------------------------------------
------------------------------------------------------------
---------ATGTCCCTGGAGCAGTGCATC------------------------------
------------------------------------------------------------
---------------------------GAACAGATAAGTAAGCTCACCGATCGCTGT---
------------------GATAGAGTCAAAGAAGCCATGACATCTCTTGTAAGCTGCATG
GAAAAGCAGTTT------------GTTGTAATGGATCATCTCCTAGCTGCCCTGAAGGAG
ATAAAAGCA---------GATCAAGTCGATTTTAGTCAGAGTCTTCTGTCTACGTCTTCT
ATGGTTAATCAACTAGTAGAGAATTCGTCTGAGCTTTTAGCCAAGCTTAGTTATTTGCCT
GTAATGTCAGGACCTGCAACATCCACTCTCGAGGCAGCTGGAGCA---------------
---------------------------------------AACACGCAGGAGCATAGAAGG
CCTCCCCCAGGGCCCATCCTAGCAACCCTGGAGCGACACGGAGCAAGACTGACAGATACT
CTCACTTCTGAT------ATTCCGGGGTATGTGAAGGCTGCTGAGGCCGAGAAGAAAATG
CATACTGCACAGACTGTCACTGGGGAGAGTGTCTCTGGGCTGCCTCTAGTTCCCACCGAA
TTCGTACGAGTCCTCACAAGTTATCTGCCAGGACCGCGCACTGCATTTCATGAATTAGTA
TCGGCAATCGCTTTGGTGAGCCGAGACTCTCACGATCTACAGGTAGCCATGGACCATTTC
AATCGAGAGCTAACGGATGGTTTCTCAGCTCATGCTGCCATAATATCCATCACTCGGAGA
TGTGAGTATTTTCGGAACTGCGAAGCTCCGACAATGCAGGTAACTTCGAAGAGCCAGATT
CCAGAAGCATATCATGGCAGACTTAGGGATGTACCAGAGGGTCCCAAAACCCTAGGACAA
GGATGGATATATATATATCTAACTCCTGAAGGA---AGCCTCGGGTTAAAGATT
>Myotis_oxyotus
------------------------------------------------------------
------------------------------------------------------------
---------ATGTCCCTGGAGCAGTGCATC------------------------------
------------------------------------------------------------
---------------------------GAACAGATAAGTAAGCTCACCGATCGCTGT---
------------------GATAGAATCAAAGAAGCCATGACATCTCTTGTAAGCTGCATG
GAAAAGCAGTTT------------GTTGTAATGGATCATCTCCTAGCTGCCCTGAAGGAG
ATAAAAGCA---------GATCAAGTCGATTTTAGTCAGAGTCTTCTGTCTACTTCTTCT
AAGGTTAATCAACTAGTAGAGAATTCGTCTGAGCTTTTAGCCAAGCTTAGTTATTTGCCT
GTAATGTCAGGACCTGCAACATCCACTCTCGAGGCAGCTGGAGCA---------------
---------------------------------------AACACGCAGGAGCATAGAAGG
CCTCCCCCAGGGCCCATCCTAGCAACCCTGGAGCGACACGGAGCAAGACTGACAGATACT
CTCACTTCTGAT------ATTCCAGGGTACGTGAAGGCTGCTGAGGCCGAGAAGAAAATG
CATACTGCACAGACTGTCACTGGGGAGAGTGTCTCTCGGCTGCCTCTAGTTCCCACCGAA
TTCGTACGAGTCCTCACAAGTTATCTTCCAGGACCGCGCACTGCATTTCATGAATTAGTA
TCAGCAATCGCTTTGGTGAGCCGAGACTCTCACGATCTACAGGTAGCCATGGACCATTTC
AATCGAGAGCTAACGGATGGTTTCTCAGCTCATGCTGCCATAATATCCATCACTCGGAGA
TGTGAGTATTTTCGGAACTGCGAAGCTCCGACAATGCAGGTAACTTCGAAGAGCCAGATT
CCAGAAGCATATCATGGCAGACTTAGGGATGTACCGGAGGGTCCCAAAACCCTAGGACAA
GGATGGATATATATATATCTAACTCCTGAAGGA---AGCCTCGGGTTAAAGATT
>Myotis_velifer
------------------------------------------------------------
------------------------------------------------------------
---------ATGTCCCTGGAGCAGTGCATC------------------------------
------------------------------------------------------------
---------------------------GAACAGATAAGTAAGCTCACCGATCGCTGT---
------------------GATAGAATCAAAGAAGCCATGACATCTCTTGTAAGCTGCATG
GAAAAGCAGTTT------------GTTGTAATGGATCATCTCCTAGCTGCCCTGAAGGAG
ATAAAAGCA---------GATCAAGTCGATTTTAGTCAGAGTCTTCTGTCTACGTCTTCT
AAGGTTAATCAACTAGTAGAGAATTCGTCTGAGCTTTTAGCCAAGCTTAGTTATTTGCCT
GTAATGTCAGGACCTGCAACATCCACTCTCGAGGCAGCTGGAGCA---------------
---------------------------------------AACACGCAGGAGCATAGAAGG
CCTCCCCCAGGGCCCATCCTAGCAACCCTGGAGCGACACGGAGCAAGACTGACAGATACT
CTCACTTCTGAT------ATTCCAGGGTACGTGAAGGCTGCTGAGGCCGAGAAGAAAATG
CATCCTGCACAGACTGTCACTGGGGAGAGTGTCTCTCGGCTGCCTCTAGTTCCCACCGAA
TTCGTACGAGTCCTCACAAGTTATCTGCCAGGACCGCGCACTGCATTTCATGAATTAGTA

TCGGCAATCGCTTTGGTGAGCCGAGACTCTCATGATCTACAGGTAGCCATGGACCATTTC
AATAGAGAGCTAACGGATGGTTTCTCAGTTCATGCTGCCATAATATCCATCACTCGGAGA
TGTGAGTATTTTCGGAACTGCGAAGCTCCGACAATGCAGGTAACTTCGAAGAGCCAGATT
CCAGAAGCATATCATGGCAGACTTAGGGATGTACCGGAGGGTCCCAAAACCCTAGGACAA
GGATGGGTATATATATATCTAACTCCTGAAGGA---AGCCTCGGGTTAAAGATT
>Myotis_riparius
------------------------------------------------------------
------------------------------------------------------------
---------ATGTCCCTGGAGCAGTGCATC------------------------------
------------------------------------------------------------
---------------------------GAACAGATAAGTAAGCTCACCGATCGCTGT---
------------------GATAGAATCAACGAAGCCATGACATCTCTTGTAAGCTGCATG
GAAAAGCAGTTT------------GTTGTAATGCATCATCTCCTAGCTGCCCTGAAGGAG
ATAAAAGCA---------GATCAAGTCGATTTTAGTCAGAGTCTTCTGTCTACGTCTTCT
AAGGTTAATCAACTAGTAGAGAATTCGTCTGAGCTTTTAGCCAAGCTTAGTTATTTGCCT
GTAATGTCAGGACCTGCAACATCCACTCTCGAGGCAGCTGGAGCA---------------
---------------------------------------ATCACGCAGGAGCATAGAAGG
CCTCCCCCAGGGCCCATCCTAGCAACCCTGGAGCGACATGGAGCGAGACTGACAGATACT
CTCACTTCTGAT------ATTCCAGGGTATGTGAAGTCTGCTGAGGCCGAGAAGAAAATG
CATACTGCACCGACTGTCACTGGGGAGAGTGTCTCTCGGCTGCCTCTAGTTCCCACCGAA
TTCGTCCGAGTCCTCACAAGTTATCTGATAGGACCCCACACTGCATTTCATGAATTAGTA
TCGGCAATCGCTGTGGTGAGCCGAGACTCTCATGATCTACAGGTAGCTATGGACCATTTC
AATCGAGAGCTAATGGATGGTTTCTCAGCTCATGCTGCCATAATATCCATCACTCGGAGA
TGTGAGTATTTTCGGAACTGCGAAGCTCCGACAGTGCAGGTAACTTCGAAGAGCCAGATT
CCACAAGCATGTCATGGCAGACTTAGGGATGTACCGGAGGGTCCCAAACCCCTAGGACAA
GGATGGGTATATATATATCTAACTCCTGAAGGA---AGCCTCGGGTTAAAGATT
>Myotis_nigricans
------------------------------------------------------------
------------------------------------------------------------
---------ATGTCCCTGGAGCAGTGCATC------------------------------
------------------------------------------------------------
---------------------------GAACAGATAAGTAAGCTCACCGATCGCTGT---
------------------GATAGAATCAACGAAGCCATGACATCTCTTGTAAGCTGCATG
GAAAAGCAGTTT------------GTTGTAATGCATCATCTCCTAGCTGCCCTGAAGGAG
ATAAAAGCA---------GATCAAGTCGATTTTAGTCAGAGTCTTCTGTCTACGTCTTCT
AAGGTTAATCAACTAGTAGAGAATTCGTCTGAGCTTTTAGCCAAGCTTAGTTATTTGCCT
GTAATGTCAGGACCTGCAACATCCACTCTCGAGGCAGCTGGAGCA---------------
---------------------------------------ATCACGCAGGAGCATACAAGG
CCTCCCCCAGGGCCCATCCTAGCAACCCTGGAGCGACATGGAGCAAGACTGACAGATACT
CTCACTTCTGAT------ATTCCAGGGTATGTGAACGCTGCTGAGGCCGAGAAGAAAATG
CATACTGCACTGACTGTCACTGGGGAGAGTGTGTCTCGGCTGCCTCTAGTTCCCACCGAA
TTCGTCCGAGTCCTCACAAGTTATCTGATAGGACCCCGCACTGCATTTCATGAATTAGTA
TCGGCAATCGCTGTGGTGAGCCGAGACTCTCATGATCTACAGGTAGCTATGGACCATTTC
AATCGAGAGCTAATGGATGGTTTCTCAGCTCATGCTGCCATAATATCCATCACTCGGAGA
TGTGAGTATTTTCGGAACTGCGAAGCTCCGACAGTGCAGGTAACTTCGAAGAGCCAGATT
CCACAAGCATGTCATGGCAGACTTAGGGATGTACCGGAGGGTCCCAAACCCCTAGGACAA
GGATGGGTATATATATATCTAACTCCTGAAGGA---AGCCTCGGGTTAAAGATT
>Myotis_capaccinii
------------------------------------------------------------
------------------------------------------------------------
---------ATGTCCCTGGAGCAGTGCATC------------------------------
------------------------------------------------------------
---------------------------AAACAGATAAGTCAGCTCACCGATCACTGT---
------------------GATAGAATCAAAGAAGCCATGACAACTCTTACAAGCTGCATG
GAAAAGCAGTTA------------GTTACAATGGATCATCTCCTGGCTGCCCTGATGGAG
ATAAAAGCACAACTCCCAGATCAAGTCTTTTTTAGTCAGAGTCTTCTGTCTATGTCTTCT
AAGATTAATCAACTAGTAGAGGATTCTTCTGAGCTTTTAGCCAAGCTTAGTTATTTGCCT
GTAATGTCAGGACCTGCAACATCCATTCTGGAGGCAGCTGGAGCA---------------
---------------------------------------AACACGCAGGAGCATAGAAGG
CCTCTCCCAGGGCCCATCCTAGCAACCCTGGAGCGACACGGAGCCAGACCGACAGATACT
CTCACTTCTGAT------ATTCCCGGGTATGCGGAGGCTGCTGAGGCCGAGAAGAAAGTG
CGTACTGCACAGACTCTCACTGGGGAGGGTGTCTCCCCGCTGCCTCTAGTTCCCACCGAA
TTCGTACGAGTCCTCACAAGTCATCTGACCGGACCCCGCACTGCATTTCATGAATTAGTA
TCGGCCATCGCCGTGGTGAGCCGAGACTCTCATGATCTACAGGTAGCCATGGACCAGTTC
AATCGAGAGCTAACGGACGGTTCCTCAGCTCATGCTGCCATAATATCCATCACTCGGAGG
TGTGAGCATTTTCGGAACTGCGAAGCTCCGACGACGCAGGTAACTTCGAAGAGCCAGATT
CCAAAAGCATGTCATGGCAGACTTAGGGATGTACCGGAGGGTCCCAAACCCCTAGGACGA
GGATGGGTATATATATATCTAACTCCGGAAGGA---AGCCTCGGGTTAAAGATT
>Myotis_blythii
------------------------------------------------------------
------------------------------------------------------------
---------ATGTCCCTGGAGCAGTGCATC------------------------------
------------------------------------------------------------
---------------------------AAACAGATAAGTCAGCTCGCCGATCACTGT---
------------------GATAGAATCAAAGAAGCCATGACATCTCTAACAAGCTGCATG
GAAAAGCAGTTA------------GTTACAATGGATCATCTCCTAGCTGCCCTGATGGAG
ATAAAAGCACAACTCCCAGATCAAGTCTTTTTTAGTCAGAGTCTTCTGTCTATGTCTTCT
AAGATTAATCAACTAGTAGGGGATTCTTCTGAGCTTTTAGCCAAGCTTAATTATTTGCCT
GTAATGTCAGAACCTGCAACATCCACTCTCGAGGCAGCTGGAGCA---------------
---------------------------------------AACACGCAGCAGCATAGAAGG
CCTCCCCCAGGGCCCAACCTAGCAACCCCGGAGCGACACGGAGCAAGACCGACAGATACT
CTCACTTCTGAT------ATTCCGGGGTATGCGGAGGCTGCGGAGGCCGAGAAGAAAGTG
CGCACTGCACAGACTGTCACTGGGGAGGGTGTCTCCCGGCTGCCTCTAGTTCCCACCGAA
TTCGTACGAGTCCTCACAAGTCATCTGACCGGACCCCGCACTGCATTTCACGAATTAGTG
TCGGCCATCGCTGTGGTGAGCCGAGACTCTCATGATCTACAGGTAGCCATGGACCAGTTC
AATCGAGAGCTAAGGGACGGTTCCTCAGCTCATGCTGCCATAATATCCATCACTCGGAGG
TGTGAGTATTTTCGGAACTGCGAAGCTCCGACAACGCAGGTAACTTCGAAGAGCCAGATT
CCAAAAGCATGTCATGGCAGACTTAGGGATGTACCGGAGGGTCCCAAACCCCTAGGACGA
GGATGGGTATATATATATCTAACTCCTGAAGGA---AGCCTCGGGTTAAAGATT
>Myotis_myotis
------------------------------------------------------------
------------------------------------------------------------
---------ATGTCCCTGGAGCAGTGCATC------------------------------
------------------------------------------------------------
---------------------------AAACAGATAAGTCAGCTCGCCGATCACTGT---
------------------GATAGAATCAAAGAAGCCATGACATCTCTAACAAGCTGCATG
GAAAAGCAGTTA------------GTTACAATGGATCATCTCCTAGCTGCCCTGATGGAG
ATAAAAGCACAACTCCCAGATCAAGTCTTTTTTAGTCAGAGTCTTCTGTCTATGTCTTCT
AAGATTAATCAACTAGTAGGGGATTCTTCTGAGCTTTTAGCCAAGCTTAATTATTTGCCT
GTAATGTCAGAACCTGCAACATCCACTCTCGAGGCAGCTGGAGCA---------------
---------------------------------------AACACGCAGGGGCATAGAAGG
CCTCCCCCAGGGCCCATCCTAGCAACCCTGGAGCGACACGGAGCAAGACCGACGGATACT
CTCACTTCTGAT------ATTCCGGGGTATGCGGAGGCCGCGGAGGCCGAGAAGAAAGTG
CGCGCTGCACAGACTGTCACTGCGGAGGGTGTCTCCCGGCTGCCTCTAGTTCCCACCGAA
TTCGTACGAGTCCTCACAAGTCATCTGACCGGACCCCGCACTGCATTTCACGAATTAGTA
TCGGCCATCGCTGTGGTGAGCCGAGACTCTCATGATCTACAGGTAGCCATGGACCAGTTC

AATCGAGAGCTAACGGACGGTTCCTCAGCTCATGCTGCCATAATATCCATCACTCGGAGG
TGTGAGTATTTTCGGAACTGCGAAGCTCCGACAACGCAGGTAACTTCGAAGAGCCAGATT
CCAAAAGCATGTCATGGCAGACTTAGGGATGTACCGGAGGGTCCCAAACCCCTAGGACGA
GGATGGGTATATATATATCTAACTCCTGAAGGA---AGCCTCGGGTTAAAGATT
>Myotis_davidii
------------------------------------------------------------
------------------------------------------------------------
---------ATGTCCCTGGAGCAGTGCATC------------------------------
------------------------------------------------------------
---------------------------CAACAGATAAGTCAGCTCAGCGATCACTGT---
------------------GATAGAATCAAAGAAGCCATGACATCTCTTACAAGCTGCATG
GAAAAGCAGTTA------------GTTACAATGGACCATCTCCTAGCTGCCCTGGTGGAG
ATAAAAGCACAACTCCCAGATCAAGTCTTTTTTAGTCAGAGTCTTCTGTCTATGTCTTCT
AAGATTAATCAACTAGTAGAGGATTCTTCTGAGCTTTTGGCCAAGCTTAATTATTTGCCT
GTAATGTCGGGACCTGCAACATCCACTCTCGAGGCGGCTGGAGCA---------------
---------------------------------------AACACGCAGGAGCACAGAAGG
CCTCCCCCAGGGCCCATCCTAGCAGCCCTGGAGCGACACGGAGCAAGACGGACAGATACT
CTCTCTTCTGAT------ATTCCCGGGTATGCAGAGGCTGCTGAGGCCGAGGAGAAAGTG
---------------------------------------CTGCCTCTAGTTCCCACCGAA
TTCGTACGAGTCCTCACAAGTCATCTGACCGGACCCCGCACCGCATTTCACGAATTAGTA
TCGGCCATCGCTACGGTGAGCCGAGACTCCCATGATCTACAGGTAGCCATGGACCAGTTC
AATCGAGAGCTAAGGGATGGTTCCTCAGCTCATGCTGCCATAATATCCATCACTCGGAGG
TGTGAGTATTTTCGGAACTGCGAAGCTCCGACAACGGAGGTAACTTCCAAGAGCCAGATT
CCAAAAGCCTGTCATGGCAGACTTAGGGATGCACCGGAGGGTCCCAAACCCCTGGGACGA
GGCTGGGTATATATATATCTAACTCCGGAAGGA---AGCCTCGGGTTAAAGATT
>Myotis_annectans
------------------------------------------------------------
------------------------------------------------------------
---------ATGTCCCTGGAGCAGTTCATC------------------------------
------------------------------------------------------------
---------------------------AAAGAGATAAGTCAGCTTACCGATCACTGT---
------------------GATAGAATCAAAGAAGCCATGACATCTCTTACAAGCTGCATG
GAAAAGCAATTA------------GTTACAATGGATCATCTCCTAGCTGCCCTGATGGAG
ATAAAAGCACAACTCCCAGATCAAGTCTTTTTGAGTCAGAGTCTTCTGTCTCTGTCTTCT
AAGATTAATCAACTAGTAGGGGATTCTTCTGAGCTTTTAGCCAAGCTTAATCATTTGACT
GTAATGGCAGGACCTGCAACATCCACTCTCGAGGCAGCTGGAGCA---------------
---------------------------------------AACACGCAGGAGCATAGAAGG
CCTCCCCCAGGGCCCATCCTAGCAACCCTGGAGCGACACGGAGCAAGACCGACAGATGCT
CTCACTTCTGAT------ATTCCGGGGTGTGCGGAGGCTGCTGAGGCCGAGAAGAAAGTG
CGTACTGCACAGCCTGTCCCTGGGGAGGGTGTCTCCCGGCTGCCTCTAGTCCCCACCGAA
TTCGTACAAGTCCTCACAAGTCATCTGACCGGACCCCGCACTGCATTTCATGAATTAGTA
TCGGCCGTCGCTGTGGTGAGCCGAGACTCTCATGATCTACAGGTAGCCATGGACCAGTTC
AATCGAGAGCTAAGGGACGGTTCCTCAGCTCATGCTGCCATAATATCCATCACTCGGAGG
TGTGAGTATTTTCGGAACTGCGAAGCTCCGACAACGCAGGTAACTTCGAAGAGCCAGATT
CCAAAAGCATGTCATGGCAGACTTAGGGATGTACCGGAGGGTCCCAAACCCCTAGGACGA
GGATGGGTATATATATATCTAACTCCAGAAGGA---AGCCTCGGGTTAAAGATT
>Myotis_muricola_brownii
------------------------------------------------------------
------------------------------------------------------------
---------ATGTCCCTGGAGCAGTTCATC------------------------------
------------------------------------------------------------
---------------------------CAACAGATAAGTCAGCTCACCGATCACTGT---
------------------GATAGAATCAAAGAAGCCATGACATCTCTTACAAGCTGCATG
GAAAAGCAGTTA------------GTTACAATGGATCATCTCCTCGCTGCCCTGATGGAG
ATAAAAGCACAACTCCCAGATCAAGTCTTTTTTAGTCAGAGTCTTCTGTCTATGTCTTCT
AAGATTAATCAACTAGTAGGGGATTCTTCTGAGCTTTTAGCCAAGCTTAATTATTTGCCT
GTAATGTCAGGACCTGCAACATCCACTCTGGAGGCAGCTGGAGCA---------------
---------------------------------------AACACGCAGGAGCATAGAAGG
CCTCCCCCAGGGCCCATCCTAGCAACCCTGGAGCGACACGGAGCAAGACCGACAGATACT
CTCACTTCTGAT------ATTCCGGGGTATGCGGAGGCTGCTGAGGCCGAGAAGAAAGTG
---------------------------------------CTGCCTCTAGTTCCCACCGAA
TTCGTACGAGTCCTCACAAGTCATCTGACCGGACCCCGCACCGCATTTCATGAATTAGTA
TCGGCCATCGCTATGGTGAGCCGAGACTCTCATGATCTACAGGTAGCCATGGACCATTTC
AATAGAGAGCTAATGGACGGTTCCTCAGCTCATGCTGCCATAATATCCATCACTCGGAGG
TGTGAGCATTTTCGGAACTGCGAAGCTCCGACAATGCAGGTAACTTCGAAGAGCCAGATT
CCAAAAGCATGTCATGGCAGACTTAGGGATGTACCGGAGGGTGCCAAAACCCTAGGACGA
GGATGGGTATATATATATCTAACTCCTGAAGGA---AGCCTCGGGTTAAAGATT
>Myotis_horsfieldii
------------------------------------------------------------
------------------------------------------------------------
---------ATGTCCCTGGAGCAGTTCATC------------------------------
------------------------------------------------------------
---------------------------AAACAGATAAGTCAGCTCATGGATCACTGT---
------------------GATAGAATCAAAGAAGCCATGACATCTCTTACAAGCTGCATG
GAAAAACAGTTA------------GTTACAATGGATCATCTCCTAGCTGCCCTGATGGAG
ATAAAAGCACAACTCCCAGATCAAGTCTTTTTTAGTCAGAGTCTTCTGTCTATGTCTTCT
AAGATTAATCAACTAGTAGAGAATTCTTCTGAGCTTTTAGCCAAGCTTAATTATTTGCCT
GTAATGTCAGGACCTGCAACATCCACTCTGGAGGCAGCTGGAGCA---------------
---------------------------------------AACATGCAGGAGCACAGAAGG
CCTCCCCCAGGGCCCATCCTAGCAACCCTGGAGCGACGCGGAGCAAGACCGACAGATACG
CTCACTTCTGAT------ATTCCGGGGTATGCGGAGGCTGCTGAGGCCGAGAAGAAAGTG
---------------------------------------CTGCCTCTAGTTCCCACCGAA
TTCGTACGAGTCCTCACAAGTCATCTGACCGGACCCCGCACCGCATTTCATGAATTAGTA
TCGGCCATCGCTGTGGTGAGCCGAGACTCTCATGATCTACAGGTAGCCATGGACCATTTC
AATCGAGAGCTAACGGACGGTTCCTCAGCTCATGCTGCCATAATATCCATCACTCGGAGG
TGTGAGCATTTTCGGAACTGCGAAGCTCCGACAACGCAGGTAACTTCGAAGAGCCAGATT
CCAAAAGCATGTCATGGCAGACTTAGGGATGTACCGGAGGGTCCCAAACCCCTAGGACGA
GGTTGGGTATATATATATCTAACTCCTGAAGGA---AGCCTCGGGTTAAAGATT
>Murina_PVJC01004914
------------------------------------------------------------
------------------------------------------------------------
---------ATGTCCCTGGAGCAGTGCATT------------------------------
------------------------------------------------------------
---------------------------GAACAGATAAGTCAGCTCACCGATCGCTGT---
------------------TGTAGAATCGAAGAAGCCATAACATCTCTTGGAACCTTTATG
GAAAAGCAGTCA------------GTTACAATGGATCATCTCCTAGCTGCCATCCTGGAG
ATGAAAGAACAACTCCCAGATCAAGCCTATTTTAATCAGAGTCTTCTATCTATATCTTCT
ATGATTAATCAACTAATGGTGGGTTCTTCTGAGCTTTCAGCCAAGCTTGGTTATTTGGCT
GAAATGGTAGGATCTGTAACATCCACTCTGGGCGCAGCTCCAGCATTCATAGAAGTTCCA
GCATTCACACAAGTTCAAGCATTCACACCAGCTCCAACATTCATACAGGCGCACGGAGGG
CCTCCCCAAGGGCCCATCCCAGCAACCCTGGAGTCACATGGAGCAAGACCGACAGATCCT
TCCACTTTTGAT------ATTCCGGGGCATGTGAAGGCTGCTGGGGCCGAGAAGAAAATG
CGTACTGCACAGACCGTCACTGGGGAGAATCTCTCTCAGCTGCCTCTAGATCCCACCCGA
TTCATACAAGTTCTCACAAGTCACCTGACAGGACACTGCACTGTGTTTCATGAATTAGTA
TCGGCAATCGCTAAAGTGAGCCGAGACTCTCGTGATCTACAGGTAGCCATGGACGAGTTC
AATCGGGTGCGAACGAGTGGTGCCTCAGCTCATGCGGCCATAATATCCATCACTCAGAGG

TGTGAGTATTTTCGGAACTGCGAAGCTCCGAAAATTCAGGTAACTTCTAAGAGCCAGATT
CCAAAAGCATGTCATGGCAGCCTCAGGGTTGTACCGGAGGGTCCCAAACTACTAGGACAA
GGATGGATATATATATATCTAACTCCCAAAGGA---AGCCTTGGGTTAAAGATT
>Acomys_cahirinus_PVKX01001220
------------------------------------------------------------
-----------ATGGCAACCGGGCCAGAAAGAAGTGGTCGGATCTCAGAGCAACTGATG
ACAGGCCAGCTCTCCATCGTTGCCTTTTTTAGCTCTGGGACTGAA---------------
---GGCTCTCTGACCAGTCCTGCAATCAAAACCTTTAGGAGTGCCGAA---------GGT
CAT------------------------AGTGCTGCCTCACCCGCTGAAAATACAGTT---
------GCAGGTGAATTACAAGATACCCGAAAACAAATTCATGAGATTACAAGGAATGTG
AGTAAGGTACTT------------GCCCAACAAGAGGAACTGACCGAAACACTCAACAGC
AGTCTCAAAGATCTGAAA---------------------GCAATAGCAGCCACGGCTGAG
ATGGTGGAGGGGATCAGAAACTGTCACGCTGAGTTGCTTGCTAAGTATGATTCATTAGTA
ACGACCACGGGCAGGTTGACGGCTACAGCGGCAGCAAGCGAGGCC---------------
---------------------------------------TCCTTGGAAGAGCAAGGAAGT
GCACCCACTGTCCCCTCTTTGTATAAGGTTGCCGTCCTTGAACAACTGTACTGTGACAAA
GACTCCAGA---------GATCCA---------GATGTCAAGTCCGCCTTACATGCCCTG
GATGAGGCTTACACCTTAAGAGAGGAGACATTTGCCAAGCCTCAGATGTCAGCCAGAGAT
TTTGCCAACCTGTTATATGGTCACCTCCCGGGCAACGACACACCCTTTCACCAATTGGCC
CAGGTTATCGTCAAGATAGCCAAACGGTCAGAGGCTATCGACATCTGCGATGCAGAATTC
CAGGCAAGTTTGGCCCACGGTGCCTCCCCTCAAGAAGCCCTGGTACAAATGATTCAACAA
ATCCCCTGCTTGAAGGAGGCCAGTCCCCCAGAAGTTGAGATCAAGAGCCAAACGGAAATC
CCACAAGCCTGCCAGAAGAGCCTGCGTGCACTTCCGCCTCATCTGCGG---CTTGGGCAA
GGCTGGATCTGTGTTTATTGTCTCCCGGATGGCAAGAAATTGGGTCTGAAGGTT
>Acomys_russatus_NC_067156
------------------------------------------------------------
-----------ATGGCAACAGGGCCAGAAAGAAGTGGTCGGATCTCAGAGCAACTGATG
ACAGGCCAGCTCTCCATCGATGACTTTTTTAGCTCTGGGACTGAA---------------
---GGCTTTCTGACCAGTCCTGCAATCAAAACCTTTAGGAGTGCCGAA---------GGT
CAT------------------------AGGGCTGCCTCACCCTCTGAAAATACAGTT---
------GCAGGTGAATTACAAGATATCCGTAAACAAATTCATGAGATTACAAGGAATGTG
AGCAAGGTACTT------------GCTCAGCAAGAGGAACTCACTGAAACGCTCAACAGC
AGTCTTAAAGATCTGAAA---------------------GCAATAGCAACCACGGCTGAG
ATGGTGGAGGGGATCAGAAACTGTCACGCTGAATTGCTTGCTAAGTATGATTCATTAGTA
ACGACCACGGGCAGGTTGACGGCTACAGCGGCAGCAAGCGAGGCC---------------
---------------------------------------TCCTTGGAAGAGCAAGGAAGT
GCACCCACTGTCCCCTCTTTGTATAAGGTTGCCGTCCTTGAACAACTGGACTGTGACAAA
GACTCCAGA---------GATCCA---------GATGTCAAGTCCGCCTTACATGCCCTG
GATGAGGCTTGCACCTTAAGAGAGGAGACATTTGCCAAGCCTCAGATGTCAGCCAGAGAT
TTTGCCAAACTGTTATATGGTCACCTCCCGGGCAACGACACACCATTTCACAAACTGGCC
CAGGTTATCGCCAAGATTGCCAAAAGGTCAGAGGCTATCGACATCTGCGATGCAGAATTC
CAGGCAAGTTTGGCCCACGGTGCCTCCCCTCAAGAAGCCCTGGTACAATTGATTCAGCAA
ATCCCCTGCTTGAAGGAAGGTAGCCCCCCAGAAGTTGAGATCAAGAGCCAAATGGAAATC
CCACAAGCCTGCCAGAAGAGCCTGCGTGCACTTCCGCCCCATCCGCGG---CTTGAGCAA
GGCTGGATCTGTGTTTATTGTCTCCCGGATGGCAAGAAATTGGGTCTGAAGGTT
>Bat_unnamed_KP233864
ATGGAGGACAAAATA---------------------------------TGTATTAATAAA
TCACGCAATCCTTATGGGATAGGACCTGAGACAACTGGGAGAATATCTGACGAATTAATG
ACAGGGAGAATACCAGTAGATGAAGTCTTGAATCATGATACATATTATCCATCT------
CAGGCTAGGTGGCAGGATCCTACAACTATATCAGCCTCGTCTGCC---------------
---------------AGAGAAGAGAAATCCATACTATTGAATATTCAGAAATTATTATTA
TCTCATAATGAGGGAATTAAGGGGATAGAGTTAGCTATTAAAGAGTTGTCACATAAATTA
TCTGAAATCAAT------------CTAGTTAATTCAAAATTAGAAACCCGACTCAATCAT
TTAGAAGTTCAGCTAAAT---------------------CAGATCAAGACATTAGCACCA
AACATTAATGGAATTCAACGTGCATGCTCAGAATTATTAGCTAAGTACGATTTGTTAGTA
TTATCTACGGGGAGGTCGACTTCAACTGCTGCAGCTACTGAAGCT---------------
---------------------------------------TACTGGAAACAACATGGACAG
ATGCCTCCTGGACCGTGCTTGTATGAAGAAAGTTATATTAAAGAGAAAATTAAAACTAAT
GATCCGTTT---------ATTCCC------GAACAATTGAAAGATGCTCATGAGAATTTG
ACTCGTGTTAATTCATTAACTGAAAAAACTTTTGCTAAACCTAGTTTTACAGCTAAAGAA
CTTAGAGATATGATATATGATCATCTTCCAGGATATGGGACAGCATTTCACCAATTAACA
CAGGTAATATGCAAGATTGCAAAAGATGAGGGTCAATTGGAGCAAGTTCATACAGAATTT
CAATCCTCCTTAGCTGAAGGTGATTCTCCACAAAGTGCATTAATACAATTGACTAAACGC
ATGACTATTTTCGACGGAAGATCACCTCCACTGATTCATATA------------------
------------------------------------------------------------

**Data S6. NP and NP-like sequences (amino acids) from Filoviruses and filovirus-like elements in vertebrate genomes (from tFasty results).** The data were used for the analysis in Fig. 5 and related analyses.

>PVKX01007375_Acomys_cahirinus
DVKQNKTLTNNNIEQRCTGQLLSFCSLFLPKSVVGEKACQEKASKVLHQVAIHAEQGLTKYPASWKATGFMIITFRIMWASFKFKFLLIYQGINVEAGHDADDISITNSTSQVRFSGLLIVKTMLDRILHHEGDHIMLHPLVRTFQVRAKVDSFCLTLKGLARHKEYALFALSLSGVN
KLEHGLFPQLSAIALGVAMAHGGTLAVTVGEQDQQLIKEAASEAEHKLQEHAEEQKIKSLGLHQAEIKIHSDFHRKKHEIGSQQAEIILATKHEKLQQLATAISSAAMGGLSQPANPFTSLPPQMPPTNFSYVSDYHSGNIETNLDHILGSSPRCEGKSSSGTPQKAPDDSYPDSFR
QRGRGRISNSVQPTTTSTSRFGTENSLNILPCDHSASDIFHLLPTSQSNPFSREYLFLKLASIHGQAPMFSQGHNPQQMMDNCQQFQSPFRITGTNCPDRTPAPANLSITGKLFRSLFTIPVSPSVLKLSQSEQQPSQYETTEASSLSSEQLEDVEVNYMNILDQEGVQAILLYYHTT
TGQPRNFITLFGTLLQYPTSLLGEVPIITSESCSTGNKKTVTINGQEQMWGDLTIQCRF
>OU015392_Acomys_dimidiatus
GQLLSFCSLFLPKSVVGEKACQEKASKVLHQVAIHAEQGLTKYPASWKATGFMIITFRIMWASFKFKFLLIYQGINVEAGHDADDISITNSTSQVRFSGLLIVKTMLDRILHHEGDHIMLHPLVRTFQVRAKVDSFCLTLKGLARHKEYALFALSLSGVNKLEHGLFPQLSAIALGVA
MAHGGTLAVTVGEQDQQLIKEAASEAEHKLQEHAEEQKIKSLGLHQAEIKIHSDFHRKKHEIGSQQAEIILATKHEKLQQLATAISSAAMGGLSQPANPFTSLPPQMPPTNFSYVSDYHSGNIETNLDHILGSSPRCEGKSSSGTPQKAPDDSYPDSFRQRGRGRISNSVQPTTTS
TSRFGTENSLNILPCDHSASESFTCSLHDHNPTLSHERSLVPQAGLDPWAGTHVQPGAPLTTNDGQLPTIPESLQDNWYQLVLTEPLPLPTYPLQAKNCSGPCSLFLYHPSVLKLSQSEQQPSQYETTEASSLSSEQLEDVEVNYMNILDQEGVQAILLYYHTTTGQPRNFITLFGT
LLQYPTSLLGEVPIITSESCSTGNKKTVTINGQE
>OU015373_Acomys_kempi
GQLLSFCSLFLPKSVVGEKACQEKASKVLHQVAIHAEKGLTKYPASWKATGFTIITFGIMWASFKFKFLLIYQGINVEAGHDADDISITNSTSQVRFSGLLIVKTMLDRILHHEGDHIMLHPLVRTFQVRAKVDSFCLTLKGLARHKEYAPFALSLSGVNNLEHGLFPQLSAIALGIAM
AHGGTLAVTVGEHQQLTKEAASEAEHKLQEHAEEQRIKSLGLHQAEIKIHSDFHQKKHEIGSQQAEIILATKHEKLQQLAAAISSAAMGGLSQPANPSTSLPPQMPPTNFSYVSDYHSGNIETKLDHILGSSSRCEGKSSSGTPQKAPDDLIQILDSDSEEGEGEEYQTQSNPPQRS
TSRLGTENSLNILPCDHSASESFTCSLHDHNPTLSHERSLVPQAGLDPWAGTHVQPGAPLTTNDGQLPTIPESLQDNWYQLVLTEPLPLPTYPLQAKNCSGPCSLFLYHPSVLKLSQSEQQPSQYETTEASSLSSEQLEDVEVNYMNILDQEGVQATLLYYHTTTGQPRNFITLFGT
LLQYPTSLLGEVPIITSESCSTGNKKTVTINGQE
>OU015760_Acomys_percivali
GQLLLFCSPFLPKLVVGEKACEESKVLHQVAIHAKQGLTKYPASWKATGFTIIIFGIRWASFKFKFLLIYQGTNLEAGHDADGISITNSISRVRFSGLLIVKTMLDRILHHEGDHIMFHSLVRTLQVRAKVNSFLTLKGLARHKEYAPFALSLSGVNNLEHGLFPQLSAIALGVAMAYGG
TLAVTVGEQHQQLIKEAASEAEHKLQEHAEEQRIKSLGCDQAEIKIHSEFHQKKHEIGSQQAEIILATKHEKLQQLATAISGAAMGGLSQPANPSTSLPPRMPPTNFSYVSDYHSGNIIETNLDHILGSSPRCEGKSSSGTRQKASDDSHPDSFRQRGRGRIPNVQPNTISTSRLGTE
NSINILPCDYSASESFTCSLHDHNPTLSHESIFSSLVPQAGLDPWAGTHVQPGAPLTTNDGQLPTIPESLQDNWDQLVLTEPLPLPTYPSQAKNCSGPRSLFLYHPSVLELSQSEQQPSQYETTEASSLSSEQLEDVEVNYMNILDQEGVQATLLYYHTTTGQPRNFITLSGTLLQYP
TSLLEVPIITSESCSTGNKKTVTINGQELQWGLKIQCRF
>chr_10_LR877244_Acomys_russatus
ALTNNNIEQRCTGQLLSFCSPFLPKSVVGEKACQESKVLHQVAIHAEQGLTKYPASWKATGFTIIIFEIMWASIKFKFLLIYQGINLEAGHDADDISITNSISQVFSGLLIVKTMLDHILHHEGDHIMLHPLVRTLQVRAKVDSFRLTLKGLARHKEYAPFALSLSRVNNLEHGLFPQLS
AIALGVVMAHGGTLTVIVGEQHQQLIKEAASEAEHKLQEHAEEQRIKSLGLHQAEIKIHSDFHQKKHEIGSQQAEIILATKHEKLQQLATAISGAAMGGLSQPANPSTNLPPRMPPTNFSYISDYHSGNIIETNLDHILGSSLRCEGKSSSGTPKDQMTLIQILDSDSEEGEGEEYQN
SVQPTTISTSRLGTENSLNILPCDHSASESFTCSLHDHNPTLSHESIFSSLVPQAGLDPWTGTHVQPGAPLTNEGQLLTIPESLQDNWDQLVLTEPLPLPTYPSQAKNCSGPRLLFLYHPSVLELSQSEQQPPSHETTEASSLSSEQLEDEEVNYMNILDQEGVQATLLYYHTTTGQA
CNFITLSGTLLQYPTSLLGEVPIITSESCSTGNKKTVTI
>PVKX01019144_Acomys_cahirinus
DTDLYGILNLGSQPVESFMCSSKIHALVAAEPNGLCTQIIHAMEAGTDLGDLLDSRYLLMLCLHHAYEGNWQVFEGRAAHRYLEANSLVVKALEEKNHDIHEFFPSAGWGRGTKQALCTLAIILSRDAQANSFFLPKLVGEKACLEKVLHQITIHAEQGLPQYPAGWEATGLMIT
VFGIMRASFSLKFLLIHHSINLEAGHGADDIITSSGAQACFLGLLIVKMVLAHTLHCEGNRIMLHPLVWTAGVRAKVDSLQPALKGLARHKEYAPFACILSLAGVNNLEHGLFLQSSAFALGVVTAQGSTLAGVTVGEQYQHLREAAGKAKQKLQAHAEEQEIKNFGLDQAEAKIL
SEFSKNHEIGTLADVILATKNEKLQQLAAATGGAAAGGPGPLAARSTNPSLPSGPLTSPTCSRPAQGIVERDLDYNPRISLRAGGVSREETPREDTRESYPPFQLLGSDSDGEKEEECDQCDPQQDADLDWEHSQMACHMTAPLLKLSNPSTPHSKPASWQRACFRSSSWSRP
GGGLAARNTLYGVRNQTALSNPSPHRVDGSNQLQSLCPSGYPLSGKTSPEMHCSHLHPLTPQNPPQAAETAGVNPRQLEITYMAILKQEGLAALSYNPEMSGQPHGFVTQSVRGGPWVEFHQRSDNKKIVIITGQEHTWEDMTLPAQL
>OU015751_Acomys_percivali

DTDLYGILNLGSQPVESFMCSKIHALVAAEPNGLCTQIIHAMGAGTDLGDLLDSRYLLILCLYHAYEGNQVFETRAAHWYLEANSLVVKALEEQNHDLHEFFPSAGWGRGTKQALCTLAIILSRDAQANFFLPSQIVGKKACLEKVLCQITVHAEQELPQYPAGWEAIELMITVFGI
MRASFTLKFLLIHHSINLEARHGADAIITSSVAQACFLGLLSVKMVLNHTLHCEGNRIMLHPLVRDSGVRAEVDSLQPALKGLARHKEYAPFACILSLSGVNNPEHGLFLQLSAIALRVVTAQGSTLAGVTVGEQYQHLREAAGEAKQKLQAHAEEQEIKNFGLDQAEAKILSEFNK
KHEIGTLQADVIVAIKSEKLQQLAAATGGAA
>LR877222_Acomys_russatus
DTDLYGILNLGSQPVESFMFSKIHALVAAEPNGLCTQIIHTMEAGTDLGDLLDSRYLLMLCLHHAYEGNQVFEGRAAHWYLEANSLVVKALEEKNHDLHEFFPSAGWGRGTKQALCTQESAVTHGPISFFLPKLVVSEKACLEKVLRQITIHAEQRLPQYPAGWEATGLMITVFG
IMRATFTLKFLLIHHSINLEAGHGADDIIITSSVAQACFLGLLIVKMVLDHTLHREGNRIMLHPLVRTARVRAEVDSLQPALKGLARHKEYSPFACILSLAGVNNPEHGLFLQLSAFALGVVTAQGSTLAGVTVGEQYQHLREAAGEAKQKLQAHAEEQEIKNFGLDQAEAKILSEFN
KKHEIGTLQADVILATKREKLQQLAAAAGGAAAGGPGPLAARSTNPSLPSGPPHLPRALGQHDRAIVERDQTTIHHQSEGRGSEQRGSSSGGMRESYPPFQLLGSESDGEEEEECDQCDPQQSDADSLGAEPDGLQHDTSTTEALQPLRMIHTPSLPHGNVLSASDLQADPG
QGLSWQPETPSMALGDQTALSNLSPHRVDESNQLQSLCPSDYPLSGKTSPGKHPLPPSSPAPQNPPQAAETGGVNPRQLEITYMAIVKREGLAALSYYPEMSGQPLVLSLISVRGGPWVEFHQRSVNNKIVIIIGQEHTWEDM
>OU015383_Acomys_dimidiatus
DTDLYGILNLGSQPVESFMCSSKIHALVAAEPNGLCTQIIHAMEAGTDLGDLLDSRYLLMLCLHHAYEGNWQVFEGRAAHRYLEANSLVVKALEEKNHDIHEFFPSAGWGRGTKQALCTLAIILSRDAQANSFFLPKLVGEKACLEKVLHQITIHAEQGLPQYPAGWEATGLMIT
VFGIMRASFSLKFLLIHHSINLEAGHGADDIITSSGAQACFLGLLIVKMVLAHTLHCEGNRIMLHPLVWTAGVRAKVDSLQPALKGLARHKEYAPFACILSLAGVNNLEHGLFLQSSAFALGVVTAQGSTLAGVTVGEQYQHLREAAGKAKQKLQAHAEEQEIKNFGLDQAEAKIL
SEFSKNHEIGTLADVILATKNEKLQQLAAATGGAAAGGPGPLAARSTNPSLPSGPPHLPHALGQHNRALLKGTTTIPRISLRAGGVSREEYSSGGTRESYPPFQLLGSDSDGEKEEECDQCDPQQSDADLDWEHSQMACHTDGSTTEARPTPPHDHTPSLPHGNGLSASDLQA
GPGQGVGWQPETPSMALGTRQLSPIPVPTGLMEAISYKAYALPVIRYLAKLPQKSLFPSSSPDPSKSTPSRNSRSKSQTARDHLHGHTEARRTVGSALQPEMSGQPHGFVTHVFRGGPWVEFHQRSDNKKIVIITGQEHTWED
>PVKX01024104_Acomys_cahirinus
DATEAGTDFGDLLDSCLLIVCLHHAYEGNRWVFDGSDCLSHQYLEANGLILKALENKENSDLHQLFPSAGLRCATKQTLRTSTNNNLELHTGQFLSFCSLFLPKLVVGEKVCLEKVLHQINIHAEQGLPQYPAGRKATGLMTIVSGIMQASFTFKLLFIHHGLNLEARHDAEDIVITS
SVAQARFSGLLTVKTVLDHILHHEGNHAMLHPLVRDAGVRAEVDSLQPALKGLARHKEYAPSVMSKNLEHGLYPAIALGISTAHESTLAGVTVREQYQQLREAASEAGWELQAHAEVSTRLKQRSFCEFNRKKHEFGTLQADVILATKHEKLKQLAAAIGGAAAGGPDPLTGRI
HSMPPQVASTSPTCSRPAQLLLNRTTAIHHQSEDPGYKQRGSSSGGLRESYPPFRLLGSDSEREEKEECQHLIHNMSDVDLDWEQRPLDDLPHDGSTPEALNPLPQDHTSRLPRGNVLSASDLQAGPDRGVWASRVSKQPYGIRNQTALQSLRLHRVDGSNQLQSLCPSHSP
GMHPILPSSLASQNPPEVNRQSSPPQATETAGVKSQTAGGNQDHLNEGLWAALSYYHSWIQPHGFVTQCRAQLQHPDCLLGDKAHGWSSTDDDPENKKIVIMNGCEQIWEDLALPARF
>PVKX01024104_1_Acomys_cahirinus
AGQLLVNVCLHHAYEGNRWVFDGSAACYLEANGLILKALENKENSDLHQLFPSAGLRCATKQTLRTSTNNNLELHTGQFLSFCSLFLPKLVVGEKVCLEKVLHQINIHAEQGLPQYPAGRKATGLMTIVSGIMQASFTFKLLFIHHGLNLEARHDAEDIVITSSVAQARFSGLLTVK
TVLDHILHHEGNHAMLHPLVQTLEEQRLTAFGAWPLPAIALGISTAHESTLAGVTVREQYQQLREAASEAGWELQAHAEVSTRLKQRSFCEFNRKKHEFGTLQADVILATKHEKLKQL
>chr28_LR877239_Acomys_russatus
NLHDILNLGSQLVESSVRNRKNPALVVAEPNELYTQIIHATEAGADFGDLLDSCLLTLCLHHAYEGNRWVFEGSAAHPVRYLEANGLILKALENKENSDLHQLFPSAGLGCRTKQTLRTLTNNNLELHRGQFLSFCSLFLPKLVVGEKVCLEKVLHQINIHAEQGLPQYPAGGKATG
LMTIVFGIMRASFTFKLLFIHHGLNLEGRHDAKDIVITSSVAQACFSRLLTIKMVLDHILHHEGNHTMLHPLVRDAGVRAEVDSFQPALKGLSRHKYAPYLSGVNNLEHGLFPQLHWGQSTLAGVTVGEQCQQLREAASEARWELQAHAEDQEMKNLGLDQAEAEILLEFNRK
KHEIGTLQADVILATKHEKLKQLAAAIGGAA
>LZPO01028967_Neotoma_lepida
GTQPVESLVRSQKIYSLDVPEPSELCSQIIYAIEAGTDLGNLLVSSLMTLCIQHTWGNQKAFQTSAARRYLEAHGLMLKVLDNKKSQVFLHWGQGKPHIILENIQEVGQFLXFCNLLLPKLVWERACLEKVQRQIVXHAEQGLPQYPQSQTPGFMIIMFGIMRVSFTFKFLFIYQGI
ILEAGRDANDIIITNSVSQARLSGLLIVKTVLDQLGDGGARLSPALRRQRQADLCEFEATLGYRVSSRKKGQSYTEKPRLEKRKKIKNHGLFPQLSAVALGVGTAHDGILGGVAVGEQYQQLEAAGERKLXEHTEEQEIHQLSLDQAEAKILLDFHKKHEIGNQQAEIILATKXGKLQ
LAAAISNNNQ
>LZPO01096522_Neotoma_lepida
VRNRKIYLLDVPEPSKLCLQIIHAIEAGIXLDLLDSCLMTLCIHHAYEGNQEAFETSAAHQYLEAHELMLKVLDNKNSNLSEFFXLTDPSGTRQALVALMNNLGEHQVGFLFFCSLFLPXLVVGERACLEKVQRQIVIXAEQGLTQYPSWQTPGYMITVLGIMRVNFTFKFLLIHQGI
NLEAGHDADDIIMTNTRFSGLLILKTVLDHILSKKVDHILLHPLVPTAKXRAELEGFRLALKGLVHPREYAPFARILNLSGVNNLGHGLFPAVVCNCPGSSNSMWKNIGRYHRGAVPATMRSSSTGITYLEHELFLQLSAIAWGVATAHGGTLAGITVXEQYQQLEVASEAERRLQE
HAEEKEIYQLGLDHAEAKILFDFHKKKHEIGNPQAEIILATKREKLQQLAAAISNNNQRVMLSPGSLAPPSSPSGCSGHRSPSKEHRDSERDFEDVQVQEVDDNDQSELSDQDETPMPRSLFQILDAGSESDEDEGHLAQRKITQILTRDLTDSATQPVSPDQTYFGTTLLPTKGP
MGDFLDRRXSSPALQDHGRRPNHRTTLPYSSQPTLDHHTRAPYSRPAIAHADASSTSDLSNXKQCQGRCPSPLPPNKRQPL
>PVIT010007961_Onychomys_torridus
DMGLHGILNYGTQPGRVRNRKIYSLGVPAPSKLCLQIMHAVEAGTDQDLLDSCLMTLCIQHTSKGNQKALETSAAPYLEAHGLMLKVLDNKNNNLNEFFALTGPSGRTRALVALRNNLGERGVGKFLSFCSLCLPTLVVRDRTCLEKVQRQIVIHAEQRLTQYPHSWQTPGCM
ITALGIIRVSFTFTFLFIQQGFNLEAGHNADDIITNSLSGKIFDHILGKEGDRILLQRSVKVRDEVESFRLAVKGLACHHEYAPFALEGVNNLEDGSFLQLSAAALRVATAPGGTLAGIPVGEQYQQLGQVATEAEHRMQEHVEEQEIYQLGLDHAEAQILLDFHKKKHEIGDQQAEII
LATKREKLQQLAAAISSNNQRVMLSSGSLAASAPLQAPWVALATVQPQRSTGITEASEDVEDQEMDHNDQSSLRQILDAGSESDEDEGHLPQERSKPNFSPKILQTHATQPLVSPDQAYFGTTSLPTKGQLTMGDFLDRHSNSDQSLSSPRSGAEPKNNIPLQLSAMNPPHILE
SPTQDQSPPMPIEAPPPPIYPMRSSSFGDNATISPPSPSNPFDSMGASSKRQPGHPLLSIGPTRDVTEAEDSWVNTKGFRQGCYFVIIHQGVPATLPLTLFQMCCNLQKPNCLTLQEAKAKMPSNSLHASCADGNLPSQSSSKTQEACLIFRHFLFINIIYLLAKFSYIAVL
>CABHPP010147144_Peromyscus_attwateri
DVELHGILNYGTQPGRVWNREIYLLGVPEPSDLCSKIIHAIEVGIDLDLLDSCLMTLCIQHTSEGKQKAFETSAAHYLEAHGLMLKVLDNKNSNLSEFFALTGPSRGTRALVALMNNLGELGVGQYLSFCSLCLPKLVVRDRACLEKVQRQIVIHAEQGLTQYPQSWQTPGCMITV
FGIVRVSFTFTFLFIHQGISLEAGHDADDIKEGDHILLHPLVQTVKVRDEVESFRLALKGLARHHEYAPFARRMNLSGVNNLEGGLSLQLSAIALGVATAPGGTLAGIPVGEQYHQLLQVVASGADRRLQEHVEEQEICQLGLDHAEAQILLDFHKKKHEIGNQAEIILATKCEKLQQ
LASAISNNNQRVMLSPRSLAASAPLQAPWVPRATVQPQRSTGITERDFEDVEDQEMDDNDQSELSDQDETPTPRSFFQILDAGSESDEDKGHLPQERSKPKILQTQQLSQESARTRRTSEPPRSPLKDNYGGFPRQTVESRSVPSSPGSQVEPKNNIPLQLTAMNPPHILESAT
QDQ
>CABHPQ010165414_Peromyscus_aztecus
DVELHGILNYGPQPGRVWNREIYSLGVPEPSELCSQIIHAVEVGIDLNLLDSCLMTLCIRHTSEGKQKAFETTAAHYLEAHGLMLKVLDNKNSNLSEFFALIGPSRGTRALVALMNNLGELGVGQYLSFCSFCLPKLVVRDRACLEKVQRQIVIHADQGLTQYPQSWQTPGCMITV
FGIVRVSFTFTFLFIHQGISLEAGHDADDIIITNTVSRARFSGLLIVKNVLDHILGKEGDHILLHPLVQTVKVRDEVESFRLALKGLARHHEYAPFARRMNLSGVHNLEGGLSLQLSAIALGVATAPGGTLAGIPVGEQYHQLLQVASGGDRGLQEHVEEQEIYQLGLDHAEAQILLDF
HKKKHEIGNQQAEIILATKREKLQQLASAISNNNQRVMLSPGSLAASAPLQAPWVAPATIQPQRSTEITERDFEDVEDQEMDDNHQSELSDQEEMPTPRSFFQILDAGSESDEDKGHLPQERSRTQVLTQDLTDSATQPGVSPDRRTSKRPHSPLKDNYGRFPRQTVESRLVPS
SPGSRAEPKNNIPLQLTAMNPPHILESPLKTSHRPSEAPPPPTYPMRSSFGASAAIPPLSSSNPFDSMGASSKRQP
>VALE03000023_Peromyscus_californicus
DVELHGILNYGTQPVEFGIEKYTRWVSRNGTKLCSKIIHAIEVGIDLHLLDSCLMTLCIRHTSEGKQKAFETSAAHYLEAHGLMLKVLDNKNSNLSEFFALTGPSRGTRALVALMNNLGELVGQYLSFCSLCLPKLVVRDRACLEKVQRQIVIHAEQGVTQYPQSWQTPGCMITVF
GIVRVSFTFTFLFIHQGISLEAGHDADDIIITNTVSQARFSGLLIVKNVLDHILGKEEDHILLHPLVQTVKVDEVESFRLALKGLARHHEYAPFAQRMNLSGVNNLEGLFLQLSAIALGVATAPGGTLAGIPVGEQYHQLLQVASGADRRLQEHVEEQEIYQLGLDHAEAQILLDFHKK
KHETGNQQAEIILATKREKLQQLASAISNNNQRVMLSPGSLAASEPLQAPRVAPTTVQPQRSTGITEDFEDVEGQEMDDNDQSELSDQDETPMPRSFFQILDAGSESDEDKGHLPQERSKPKFSPKILQTQQLSQESAQTGRTSEPPCSPLKYNYGRLPRQTVESRSAPSSPGSP
AEPKNNIPLQLTAMNPPHILESHTQDQSPMPRLHPPPIYPTRSSSGASAAIPPLSSSNPFDSMGASSKRASTRPPLDLDLKHWMQRLKTEVGSRQKGFGQGCYFVIIQQGVPATLPLTRFQIYCHFQEPNCLILHEAKARIPSNSLHASCADWQPSQSVLQQNSGGLFNFQALSL
LFINIISLLAKFSYIAIL
>CABHPR010130060_Peromyscus_melanophrys
DVELHGILNYGTQPGRVWNREIYLLGVPEPSDLCLKIIHAIEVGIDLDLLDSCLMTLCIQHTSEGNKKAFETSAAHYLEAHGLMLKMLDNKNSNLSEFFALTGPSQGTRALVALMNNLGELGVGQYLSFCSLCLPKLVVRDRACLEKVQRQIVIHAEQGFTQYPQSWQTPGCMIT
VFGIVRVSFTFTFLFIHQGISLEAGHDADDIIITNTRFSGLLIVKNVLDHILGKEGDRILLHPLVQTVKVRDEVESFRLALKGLARHHEYAPFALRMNLSGVNNLEGGLSLQLSAIALGVATAPSGTLAGIPVGEQYHQLLQLASGADEVARAHVEEQEIYQLGLDHAEAQILLDFHKKK
HEIGNQAEIILATKREKLQQLASAISNNNQRVMLSPGSLAASAPLPSPLGCSGHRSASKEHRDYRTRFGCRGSRDGQSKVVRPGRDAHAHVPSYQILDAGSESDEDKGRLPQERSKPKFSPKILQTQQLSQESARTGVLRNPSLPTKGQLWEISSTGSQSGSVPSSPGSRAEPKN
NIPLQLTAMNPPHILESATQDQSSPMTEVPPPPIYPMRSSSGASAAIPPPSSSNPSNSMGASSKRQPRHAL
>CABHPH010150859_Peromyscus_nudipes
DVELHGILNYGTQPGRVWNREIYLLGVPEPSDLCSKIIHAIEVGIDLDLLDSCLMTVCIQHTSEGKQKAFETSAAHYLEAHGLMLKVLDNKNSNLSEFFALTGPSRGKRALVALMNNLGELGVGQYLSFCSLCLPKLVVRDRACLEKVQRQIVIHAEQGLTQYPQSWQTPGCMITV
FGIVRVSFTFTFLFIHQGISLEAGHDADDIIITNTRFSGLLIVKNVLDHILSKEGDRILLHPLVQTVKVRDEVESFRLALKGLARHHEYAPFARRMNLWGGGVNNLGGGLSLLLSAIALGVATAPGGTLAGIPVGEQYRQLLQVASGADNRLQEHVEEQEIYQLGLDHAEAQILLDFH
KKKHEIGNQAEIILATKREKLQQLASAISNNNQRVMLSPGSLAASAPWVARATVQPQRSTGITERDFEDVEDQEMDDNDQSGVVRPGRDAHAHVPSYQILDAGSESDEDRGHLPQERSKPKFSPKILQTQQLSQESARTGVLRNPSLPTKGQLWEISSTGSQSGSVPSSPGSR
AEPKNNIPLQLTAMNPPHILESATQDQSSPMTEVPPPPIYPMRSSSGASAAIPPPSSSNPSNSMGASSKRQPRHAL
>CACRXM010000018_Peromyscus_eremicus
DVELHRILNYGTQPVEFGIEKYTRWVSWNGTELCSKIIHAIEVGIDLDLLDSCLMTLCIRHTSEGKQKAFETSAAHYLEAHGLMLKVLDNKNSNLSEFFALTGPSGTRALVALMNNLGELGVGQYLSFCSLCPPKLVVRYRACLEKVQRQIVIHAEQGLTQYPQSWQTPGCMITVF
GIVRVSFTFTFLCIHQGISLEAGHDADDIIITNTVSQARFSGLLIVKNVLDHVLGKGEDHILLHPLVQTVKVDEVESFRLALKGLARHHEYAPFAQRMNLSGVNNLEGLLLQLSAIALGVATAPGGTLAGIPVGEQYHQLLQVASGADRRLQEHVEEQEIYQSGLDHAEAQILLDFHK
KKHEIGNQQAEITLATKREKLQPLASAISNNNQRVMLSPGGLAASAPLQVPQRSTGITEDFEDVEDQEMDDNDQSELSDQDETPTPRSFFQILDAGSESDEDKGHLPQERSRTQVLTQDLTDSATQPGVSPDRAYFGTTLLPTKGQLWETSSTDSRIPISPQLSRVTGGPQEQRS
PTAHSHEPLESHTQDQSPMPRLAPPQIYPMQSSSGASAAIPPRSSSNPFDSMGASSKRASTRPPDLDRKHWMWRLKTEVGSIQKGFGQGCYFVIIQQGVPATFLTSSTCWPNFLTQY
>NMRJ02000012_Peromyscus_leucopus
NAKVSANSPLGGRSNLPNDVELHGILNYGTQPGRIWNREIYSLGVPEPSKLCSQIIHAIEVGRDLDLLESCLMTLRIRHTSEGKQKAFETSAAHYLEAHGLTLKVLDNKDSNLSEFFALTGPSRGTRALVALMNNLGELGVGQYLSFCSLCLPKLVVRDRACLEKVQRQIVIHAEQGL
TYPQSWQTPGCMITVFGIVRVSFTFTFLFIHRGISLEAGHDADDIIIINTVSQARFSGLLIVKNVLDHILGKEEDRIVLHPLVQTVKVDEAESFRLALKGLARHPEYAPFARVMNLSGVNTLEEGLFLQLSAVALGVATAPGGSLAGIPVGEQYHQLLQVASGADRRSQKHVEKQEIY
QLGLDHAEAQILLDFHKKKHEIGNQQAEIILATKREKLQHPQLKATISKGRVEPRESYSLSTPSSPLGCSDHRSASKEHRDYRMRFLRMSRIKRWDDNDQSELTGQDETPTPRSFFQILDAGSESDEDKGHLPQERSKPKFSPKILQTQQLSQESAQTGRTSEPPRSP
>JAOPKX010000017_Peromyscus_maniculatus
NTKVSANSPLGGRSNLPNDVELHGILNYGTQPGRVWNREIYSLGVPEPSKLCSQIIHAIEVGRDLDLLESCLMTLRIRHTSEGKQKAFETSADHYLEAHGLMLKVLYNKDSNLSEFFALTGPSRETRALAALMNNLGELGVGQYLSFCSLCLPKLVVRDRACLEKVQRQIVIHAEQG
LTYPQSWQTPGCMITVFGIVRVSFTFTFLFIHRGISLEAGHDADDIIITNTVSQARFSGLLIVKNVLDHILGKEEDRIVLHPLVQTVKVRDEAESFRLALKGLARHHEYAPFARRMNLSGVNNLEEGLFLQLSAVALGVATAPGGSLAGIPVGEQYHQLLQVASGADRRLQEHVEEQ
EIYQLGLDHAEAQILLDFHKKKHEIGNQQAEIILATKREKLQQLASAKSNNNQRVLLSPGSLAASAPLPSPLGCSDHRSASKEHRDYRMRFLRMSRIKRWDDNDQSELSGQDETPTPRSFFQILDAGSESDEDKGHLPQERSKPKFSPKILQTQQLSQESAQTGRTSEPPRSP
>RCWS02057567_Peromyscus_polionotus
NAKVSANSPLGGRSNLPNDVELHGILNYGTQPGRVWNREIYSLGVPELSKLCSQIIHAIEVGRDLDLLESCLMTLHIRHTSEGKQKAFETSAAHYLEAHGLTLKVLDNKDSNLSEFFALTGPSRETRALAALMNNLGELGVGQYLSSCSLCLPKLVVRDRACLEKVQRQIVIHAEQGL
TYPQSWQTPGYMITVFGIVRVSFTFTFLFIHQGISLEAGHDADDIIITNTVSQARFSGLLILKNVLDHILGKEEDRIVLHPLVQTVKVDEVESFRLALKGLAHHHEYAPFARRMNLSGVNNLEEGLFLQLSAVALGVATVPGGSLAGIPVGEQYHQLLQVASGADRRLQEHVEEQEIY
QLGLDHAEAQILLDFHKKKHEIGNQQAEIILATKREKLQQLASAKSSNNQRVVLSPGSLAASAPLPSPLGCSNHRSASKEHRDYRMRFLRMSRIKRWDDNDQSELSDQDETPTPRSFFQILDAGSESDEDKGHLPQERSKPKFSPKILQTQQLSQESAQTGRTSEPPRSP
>JAOPKW010000020_Peromyscus_maniculatus
NAKVSANSPLGGRSNLPNDVELHGILNYGTQPGRVWNREIYSLGVPEPSKLCSQIIHAIEVGRDLDLLDSCLMTLRIRHTSEGKQKAFETSAAHYLEAHGLTLRVLDNKDSNLSEFFALTGPSRGTRALAALMNNLGELGVGQYLSFCSLCLPKLVVRDRACLEKVQRQIVIHAEQG
LTYPQSWQTPGCMITVFGIVRVSFTFTFLFIHRGISLAAGHDADGIIITNTVSQARFSGLLIVKNVLDHILGKEEDRIVLHPLVQTVKVRDEVESFRLALKGLARHHEYAPFARRMNLSGVNNLEEGLFLQLSAVALGVATAPGGSLAGIPVGEQYHQLLQVASGADRRLQEHVEEQ
EIYQLGLDHAEAQILLDFHKKKHEIGNQQAEIILATKREKLQQLASAKSNNNQRVMLSPGSLAASAPLPSPLGCSDHHSASKEGLQNEIFEDVEDQEMDDNDQSELSGQDETPTPRSFFQILDAGSESDEDKGHLPQERSKPKLSPKILQTQQLSQESAQTRRTSEPPRSPLKDNM
GDFLDRPSNPDQSDRG
>RCWR01075790_Peromyscus_maniculatus
NAKVSANSPLGGRSNLPNDVELHGILNYGTQPGRVWNREIYSLGVPEPSKLCSQIIHAIEVGRDLDLLESCLMTLHIRHTSEGKQKAFETSAAHYLEAHGLTLRVLDNKDSNLSEFFALTGPSRGTRALAALMNNLGELGVGQYLSFCSLCLPKLVVRDRACLEKVQRQIVIHAEQG
LTYPQSWQTPGCMITVLRIVRVSFTFTFLFIHRGISLKAGHDADDIIITNTVSQARFSGLLIVKNVLDHILGKEEDRILLHPLVQTVKVDEAESFRLALKGLARHHEYAPFARRMNLSGVNNLEEGLFLQLSAVALGVATAPGGSLAGIPVGEQYHQLLQVASGADRRLQEHVEEQEI
YQLGLDHAEAQILLDFHKKKHEIGNQQAEIILATKREKLQQLASAKSNNNQRVLLSPGSLAASAPLQAPWVAPTTVQPRSTGITEDFGYRGSRDGYNDQSELSGQDETPTPRSFFQILDAGSESDEDKGHLPQERSKPKFSPKILQTQQLSQESAQTGRTSEPPRSPLKDNMGDF
LDRPSNPDQSDRG
>CABHPP010168786_Peromyscus_attwateri
DVELRRILNYGTQLVKSLVRSKIYSLDVPEPSELCSQIIQAIETRIDNCLMTLGIQHTYEGNQKAESAARYLEAHGLMLDNRNNSNLSEFFALVGPSGTRQALVALMNNLGERRVGQFLSFCSLFLPKLVVRERACHEKVQRQGLPQYPQSWQTPGFMVISFGIMRVSFTFKFLFH
QGINLEAGHNADDIIITNSAQARFSGLPIVKTVLDHILRKEGDHILLPLVQTVKVQAEVESFRLVLKGLAHHHEYAPFAQIRNLSGVNNLEHGLFPYLSAVALGVTTAHGILAGCHVREQYQLREAAGEAERRLQEHAEEQEIHQLGLDQAETKILLDFHKKKHAIGNQQAEIILTTK
QEKLQQLAASISNNSQRVTLSPGSFVVSASLQAPWVAPVAVQPQRSAEITDPDSEYLKGQDMEDNDPSQLSDYDSTPTQHSMFQILDAESESNEDEGHLLCEKPRPKLSPEILQTRQFNSEPAQIRYTAEPPPSLPKNHHGRSPQQTIQSQSVPNSPGSWAEPQTNEQCSPTA
HSHEPLAIILESLTQDWSSPMPTEPPLPPPTPPPPAPLIQCKAVTVASASIPHLSPSNSFSSLGASPKQETASTSPPLALDYKQLEAIKAEYQSRVNTKGIRAGLLFYYNTTGQACNFTTHSGSILQCPTSLTGTTIPAVTPRAGVGDESTVTVDNQEQAWSDLSVQSWF
>CABHPP010168787_Peromyscus_attwateri
DVELRRILNYGTQLVKSLVRSKIYSLDVPEPSELCSQIIQAIETRIDNCLMTLGIQHTYEGNQKAESAARYLEAHGLMLDNRNNSNLSEFFALVGPSGTRQALVALMNNLGERRVGQFLSFCSLFLPKLVVRERACHEKVQRGLPQYPQSWQTPGFMVISFGIMRVSFTFKFLFHQ
GINLEAGHNADDIIITNSAQARFSGLPIVKTVLDHILRKEGDHILLPLVQTVKVQAEVESFRLVLKGLAHHHEYAPFAQIRNLSGVNNLEHGLFPYLSAVALGVTTAHDGILAGCHVREQYQLREAAGEAERRLQEHAEEQEIHQLGLDQAETKILLDFHKKKHAIGNQQAEIILTTK
QEKLQQLAASISNNSQRVTLSPGSFVVSASLQAPWVAPVAVQPQRSAEITDPDSEYLKGQDMEDNDPSQLSDYDSTPTQHSMFQILDAESESNEDEGHLLCEKPRPKLSPEILQTRQFNSEPAQIRYTAEPPPSLPKNHHGRSPQQTIQSQSVPNSPGSWAEPQTNEQCSPTA
HSHEPLAIILESLTQDWSSPMPTEPPLPPPTPPPPAPLIQCKAVTVASASIPHLSPSNSFSSLGASPKQETASTSPPLALDYKQLEAIKAEYQSRVNTKGIQAGLLFYYNTTGQACNFTTHSGSILQCPTSLTGTTIPAVTPRAGVGDESTVTVDNQEQAWSDLSVQSWF
>CABHPQ010139687_Peromyscus_aztecus
WYSNVQNDVELHRILNYGTQLVKSLVGSKIYSLDVPEPSELCSQIIQAIETRIDNCLMTLGIQHTYEGNEKAESAARYLEAHGLMLDNRNNSNLSEFFALVGPSGTRQALVALMNNLGERRVGQFLSFCSLFLPKLVVGERACHEKVQRQGLPQYPQSWQTPGFMVISFGIMRV
SFTFKFLFHQGIKLEAGHNADDIIITNSAQARFSGLPIVKTVLDHILRKEGDHILLPLVQTVKVQAEVESFRLVLKGLAHHLEYAPFAQIRNLSGVNNLEHGLFPYLSAIALGVTTAHGGILTGVNVREQYQLREATGEAERRLQEHAEEQEIHQLGLDQAETKILLDFHKKKHAIGNQ
QAGIILTTKQEKLQQLAASISNNNQRVTLSPGSFVVSASLQAPWVAPVAVQPQRSAEITEPDSEDLKGQDMEDNDPSQLSDNDSTPTQHSMFQILDAESESNEDEGHLLCEKPRPLSPEILQIRQFNSEPAQIRYTAEPPPSLPKNHHGRSPQQTIQSQSVRNSPGSWAEPQTN

EQCSPTAHSHEPLATILESLTQDWSSPMPTEPPLPXXXXXXXXXXXXXXXXXXXXXXXXXXXXXXXXXXXXPAPLIQCKAVTVASASIPPLSPSNSFSSLGASPKQETASTSPPLALDYKQLEAIKAEYQSRVNTKGIRAGLLFYYNTTGQACNFTTHSGSILQYPTSLKRTTIPAMNLRAGI
GVESTVTVDNQEQAWSDLSVQSWF
>CABHPR010136284_Peromyscus_melanophrys
FFALVLCFLFIRRLVTSTDHILLPLVQTVKVQAEVESFRLVLKGLAHHHEYAPFAQIRNLSGVNNLEHGLFPYLSAVVLGVTTAHDGILAGVNVREQYQLQEAAGEAERRLQEHAEEQEIHQLGLDQAETKILLDFHKKKHAIGNQQAEIILTTKQEKLQQLAASISNNNQRVTLSPG
SLVVSASLQAPWIAPVVVQPQRSAEITEPDSEDLKGQDMEDNDPSQLSDHDSTPTQHSMFQILDAESESNEDEGHLLCEKPIPKLSPEILQTRQFNSEPAQIRYTAEPPPSLPKNHPGRSLWQTIQSQSVPNSPGSWAEPQTNEQCSPTAHSREPLAIILESLTQDWSSPMPTEP
PLPPPPPLIQCKAVTVASASIPPLSPSNPFSSLGPSPKQETASTPPPLALDHKQLEAIKAEYQSRVNTKGIWAGLLFYYNTTGRACNFTTHSGSILQCSNSLTGTTIPAVTPRAGIGDESTVTVDNQEQAWSDLSVQSWF
>VALE03000011_Peromyscus_californicus
DVELCRILNYGTQPVESLVRSKICSLDVPEPSELCSQIIHTIETRTDNCLMTLGIQHTYEGNQKAESATCYLEAHGLMLDNRNNTNLSEFFALMGPSGTRQALVALMNNHGERRVGQFLSFCSLFLPKLVVGERACLEKVQRQVLPQYPQSWQTPGFMVISFGIMRVSFTFKFLFH
QGINLEAGHNADDIIITNSAQARFSGLLIVKTVLDHILRKEGDHILLHLLVQTVKVQAEVESFRLILKGLAHHHEYAPFAQINLSGVNNLEHGLFPHLPAVALGVTTAHDGILAGVTVREQYQQLREAVGEAERRLQEHAEEQEIHQLGLDQAKAKILLDFHKKKHAIGNQQAEIILT
TKQEKLQQLAALISNNNQRVTLSPGSFVASVSLQAPWVAPVAVQPQRSAGTTEPDSEDLKGDMEDNDPSELSDHDSIPTQHSMFQILDAESESNEDEGHLLCEKPRPKLSPEILQTRQFNSEPAQIRYTAEPPPSLPKNHHGRSPRQTIQSQSVPNSPGSWAEPPTSKQCSPTA
HSHEPLATILESLTQDWSSPMPTEPPLPPPPLPSAKQSTIASASISPLSPSNPFSSMGASPKQETASTPPPLALDPKQLEAIKTEYQSRVNTKGIWAGLLFYYNTTGHACNFTTHSGSILQYPTSLTGTTIPAMTPRGGIGDESTVTVDNQEQAWSDLSVQSWF
>CACRXM010000007_Peromyscus_eremicus
DVELCRILNYGTQPIESLIRSKIYSLDVPEPSELCSQIIHAVETRMDNCLMTLGIQHTYEGNQKAESAARYLEVHRLMLDNRNNSNLNEFFASMGPSGTRQALVALMNNLGERRRVGRFLSFCTLFLPKLVVGERDCLEKVQRQVLPQYPQSWQTPGFMVISFGIMRVSFTFKFLF
HQGINLEAGHNADDIIISNKLSQARFSGLPIVETVLDHILRKEGDHILLHLLVQTVKVQAEVESFRLILKGLAHHHEYAPFAQMNLSGVNNLEHGLFPHLPAVALGVTTAHDGILAGVTIREQYQQLREAVGEAEHRLQEHAEEQEIHQLGLDQAEAKILLDFHKKKHAIGNQQAEII
LTAKQEKLQQLAVSISNNNQRVTLSPGSFVASASLQVPWVAPVAVQPQGSAGTTEPDSEDLKGQDMEDNDLSELSDHGSTPMQHSMFQILDAEFESKEDEGHLLCEKPRPKLSPEILQPDNSTASQLRSDTLQNHLPPRQRTTMGDPQGRSSNPNQSLTLQGRGQSSDQRA
MLSHSSQPTLDHHTHPRLVIAHANASFTPPLTQCEAVTIASASIPLLSPSNPFSSMGASPQQETASTLPPLALDHKQLEAIKTEYQSWVNTKGIQAGLLFYYNTTGHACNFTTHSGSILQYPTSLTGTTISVMTPRGGIGDESTYITVDNQEQAWSDLSVRSWF
>NMRJ02000021_Peromyscus_leucopus
WYSNTQKDVELRRILNYGTQPVESPVRSKIYSLDVPEPSELCSQIIQAIETRIDNCLMTLDIQHTYEGNQKGESAAHYLEAHGLMLDNRNNSNLSEFFCIDGPLRDKASLSSIDESPRVSSVGHFLSFCSLFLPKLVVGERACLEKVQRQGLPQYSLSWLVLRIMRMRFTFKFLFIKGT
HLEAGHNADDIIITNSAQARYSGLLIVKTVLDHILCQEGDHILLHLLVQTVKVQAGVESFSLVLKGLAHHHEYVPFAHIRNLSGVNNLEHGLFPHLSAVALGVTTAHDGILAGVTVREQYQQLREAVGEVERRLQEHAEEQEIHQLGLDQAEAKVLLDFHKKKHAIGNQQAEIILTT
NQEKLQQLAASISNNNQRVTLSPGGFAASASLQAPWVAPVAVQPQRRAGITEPDSEDLQGQDVEDNDPSELSDHDSTPTQHSMFQILDAESESNEDEGHFSVKNPDPNFHLRSCRPGNSTVSQLRSNTLQNHLPTKELPWEIPHSRPTNPNQSLTLQGHGQSLKPTSNALP
QLTAMNPWPPYSPSPKTGHRSCQLSLLYPTHLSSIEAVTIASASIPPLSPSNPFSSVGASPKQETASTPPPLALDHHQLEAIKTEYRSRVNTKGIQADLLFYYNTTGRACNFSTHSGSILQYPTSLTGTTIPAMMTPEGWYWKYITVDNQEQAWSDLSVQSWF
>JAOPKW010000015_Peromyscus_maniculatus
WYSSTQKDVELCRILNYGTQPVKSPVRSKIYSLDVPEPSELCSQIIQAIETRIDNCLMTLDIQHTYEGNQKAESAAHYLEAHGLMLDNRNNSNLSEFFLPLMGPSGTRQALVALMNNLRERKCVGHFLSFCSLFLPKLVVGERACLEKVQRQGLPQYPQSWLVLRIMRMRFTFKFL
FIKGTHLEAGHNADDIIITNSLRQRYSGLLIVKTVLDHILCQEGDHILLHLLVQTVKVQAEVESFRLVLKGLAHHHEYAPFAHIRNLSGVNNLEHGLFPHLSAVALGVTTAHDGILAGVTIREQYQQLREAVGEVERRLQEHAEEQEIHLGLDQAEAKVLLDFHKKKHAIGNQQAEIIL
TTNQEKLQQLAASISNNNQRVTLSPGSFAASASLQAPWVAPVAVQTQRRAGILEPDSEDLQGQDMEDNDPSELSDHDSTPTQHSMFQILDAESESNEDEGHFSVKNPDPNFHLRSCRPGNSTVSQLRSNILQNHLPPHQELPWEIPAADRPIPISPLSRVMGRESLKPASNAL
PQLTAMNPWPPYSPSPKTGHHSCQLSLLYPTHLSNDEAVTIASASIPPLSPSNPFSSVGASPKQETASTPPPLALDHQQLEAIKTEYQSRVNTKGIRAGLLFYYNTTGRACNFSTHSGSILQYPTSLTGTTMPAMTPRGGIGDESTVTVDNQEQAWSDLSVQSWF
>RCWR01034677_Peromyscus_maniculatus
WYSSTQKDVELCRILNYGTQPVKSPVRSKIYSLDVPEPSELCSQIIQAIETRIDNCLMTLDIQHTYEGNQKAESAAHYLEAHRLMLDNRNNSNLSEFFCTVGPSGTRQALVALMNNLRERRCVGHFLSFCSLFLPKLVVGERACLEKVQRQGLPQYPLSWLVLRIMRMRFTFKFLFI
KGTHLEAGHNADDIIITNSAQARYSGLLIVKTVLDHILCKEGDHILLHLLVQTVKVQAEVESFRLVLKGLAHHHEYAPFAHIRNLSGVNNLEHGLFPHLSAVALGVTTAHDGILAGVTVREQYQQLREAVGEVERRLQEHAEEQEIHQLGLDQAEAKVLLDFHKKKHAIGNQQAEII
LTTNQEKLQQLAASISNNNQRVTLSPGSFAASASLQAPWVAPVAVQPQRRAGILEPDSEDLQGQDMEDNDPSELSDHDSTPTQHSMFQILDAESESNEDEGHFSVKNPDPNFHLRSCRPGNSTVSQLRSNILQNHLPPHQELPWEIPAADRPIPISPLSRVMGRESLKPASNA
LPQLTAMNPWPPYSPSPKTGHHSCQLSLLYPTHLSNDEAVTIASASIPPLSPSNPFSSVGASPKQETASTPPPLALDHQQLEAIKTEYQSRVNTKGIRAGLLFYYNTTGRACNFSTHSGSILQYPTSLTGTTMPAMTPRGGIGDESTVTVDNQEQAWSDLSVQSWF
>RCWS02021267_Peromyscus_polionotus
NVELRRILNYGTQPVESPARSKIYSLDVPEPSELCSQIIQAIETRIDNCLMTLDIQHIYEGNQKAESAAHYLEAHGLMLNNRRNNSNLSEFFSLMGPSETRQALVALMNNLRERRCVGHFLSFCSLFLPKLVVLERACLEKVQRQGLPQYPLSWLVLRIMRMRFTFKFLFIKGTHLEAG
HNADDIIITNSAQARYSGLLIVKTVLDHILCQEGDHILPHLLVQTVKVQAEVESFRLVLKGLAHHHEYAPFAHIQNLSGVNNLEHGLFPHLSAVALGVTTAHDGILAGVTVREQYQQLRESVGEVECRLQEHAEEQEIHQLGLDQAEAKVLLDFHKKKHAIGNQAEIILTTKQEKLQ
QLAASISNNNQRVTLSPGSFAASASLQAPWVAPVAVQPQRRAGITEPDSEDLQGQDMEDNDPSELSDHDSTPTQHSMFQILDAESESNEDEGHFSVKNPDPNFHLRSCRPGNSTVSQLRSNTLQNHLPPHQRTPMGDPRSRPTNPNQSLTLQGHGQSLKPTSNALPQLTA
MNPWPPYSPSPKTGHRSSNRASFTHPLIQCEAVTIASASIPPLSPSNPFSSVGASSKQETASRNSLPLPVDHQQLENIKTEYQSRVNTKGIRAGLLFYYNTTGLACNFSTHSGTILQYPTSLTGTTMPAMTPRGGIGDESSVTVDNQEQAWSDLSMQSWFAYCYEH
>CABHPH010117406_Peromyscus_nudipes
DVELRRILNYGISTHRITSQELIYSLDVSELSELCSQIIQAIETRIDNCLMTLDSQHTYEGNQKAESTARYLEAHGLMLDNRNNSNLGEFFSLMGPSGTRQALVALMNNLGERRVGQFLSFCSLFLPKGRACLEKVQRQGLPQYSQSWQTPGFMVISFGIMRVSFTFKFLFHQGINL
EAGHNADDIIITNSAQARFSGLPIVKTVLDHILRKEGDHILLPLVQTVKVQAEVESFRLVLKGAHHHEYAPFAQIRNLSGVNNLEHGLFPYLSAVVLEVTTAHDGILAGVNVREQYQLEAAGEAERRLQEHAEELEIHQLGLDQAETKILLDFHKKKHAIGNQQAEIILTTKQEKLQQ
LAASISNNNQRVTLNPGSFVVSASLQAPWVVPVAVQPQRSVEITEPDSEDLKGQDMEDNDPSQLSDHDSTPTQHSMFQILDAESESNEDEGHFSVKNPDPNFHLRSRRSGNSTASQLRSDTLQNHLPPCQRTTMGDPRGRSSNPNQFLTLQGHGQSLKPTSNALPQLTAM
NPWPPYSLSPKTGHRPCQLSLLYRPPPPPPPPPQCKAVTVASASIPPLSRPILSVPWVLSPKQETASTPPPLALDHKQLEAIKAEYQSRVNTKGIRAGLLFYYNTTGCACNFTTHSGSILQCPTSLTGTTIPVVTPRAGIADESTVTVDNQEQAWSDLSVQSWF
>APMK01173051_Cricetulus_griseus
QPQEDLELHSLLNLGTQPVESFVRSHKIYSFGVPDSIELGSQIIYSIETRVEFIDLVDRCLLTLCILHAYKGNQEAFMTSAARRYLEKKKDNFKVLDNNSSSKLNEFLPDTGPSRRTRQALLALSDNNNGHREIEQFLSFCSLFLPKLVVEEGACLQKVQHQIVVHREQDLTQCPQGWN
TPDSRMIIFRVMKASFTFKFMLIHQGINLVAGQDPDNIIITNSVTQARFSGLLLKTVRDHTLDKDRDPATLHPLARGKVRSEVENFKLALKALACHCEYAPFARTMSLSGVNNLEHGPFSQLSAINLQVATAHGCTLAGMAVGEQQQLRCCGEAATEAERRLQKYTLNREQETSR
LGLDLTEAKILSDFHKKKHEIGNQQAEVTKWEKLQQLAADIINTNQRQPASQASPHPPILAPPQPPPAEEHPTSLGEQEADLAEQELLEHDADLEVEDDNPNGCLNMKAPRIDTPNFQSMPTLSRQRPGSRVTRDPKFPKKKNPHWPHGSPSSETIQSEGIVSSQPRSTWSSM
SGRDLNTEPCHNHHHSQDTGSLGHRFLLAIPARESLSSIPELPGENWAHSSFVEPLPPPPIYPISGNEKGVAQQPQTSQLEALRPSNVTSKLRLDTTPTKPDLGAEQLKEAKETYRVCAQGPPCRTVALLQHNLTPQHFSTRSSATLQDPKSLLGVSAPVITKEGGISGEDNSLTLDG
QKQHWSQLNMPSRF
>APMT01047719_Mesocricetus_auratus
QPQENLELHSTLNLGTQPVESFVGSHKIYSFGVPDSIELGSQIIHAIETGVEFGDLLDRRLLALCIHHAYKGNHGNFMNSATRRYLEKNGIICKVLDNNPSSNLNEFLPDTVPSRKMRQALLALANNSNGHGKIGQFLSICILFLPKLVVGEGPCLEKVQHQIVVHREEGLARCPGW
NIPGSRMIIFRVMRVSFTFKFMLIHQGINLVAGHEPDNIIITISVTQARFSGLLIKTFLDHTLGKDRDETLHPLVRDKVRSEVENFKLALKGLAHCEYAPFAIMSLSGANNLEHGATPQLSVIALVATAYGCILTDRTVGEQQLNNSYAMGSHYSRKRFAETTLNKEHETSRLGFDLTE
AKILSDFHKKKHEIGNQQAEVILVTKGEKLQQLAAGIINTNERPASQAPLPPPVSGTTPAPSRAEERPSLLDAGRTGSRSSQSKNYSYDADLEVEDNNPNKLPEHESSQDRHPKFSILDADSGGQGRQEWRLCRDPTIPKKNSHWPHGSPSRGNQTVRRHEFSTNSTSSVSGREL
NTEPCHNHHPSQDTGSLGHRFLVIPAQESLSSIPELPGETAHSSFVEAPPPPIYPISGNEKGVAQQQQNSQLEALSPSNVTSNLRLDIPPTKPGLSAEQLKEAKETHCKGAHSEGLHAGLSLYYSTTGQARFSTLSSASLQGLTSLPGVSAPVITKREAFEEDDTLTLDGQKQHWSQL
NMSSRF
>CALSGD010001501_Phodopus_roborovskii
KPQEDLELHSILNLGTQPVESFVRSQKIYSFGVPDPIDCGFTIDAVEAGAEFVDLLNRCLALCIQHASKGNQKVFINSVARHYLEKNGIICKVLDNNPISISSEFLQTQGMRQALLALADDNYGHRKTGQSLSFCSLFLPKLVVGERACLKKKMNRQIVTHRGQGLTQCPQGWNTP
GSRMTIFRVMRVMLLHQGINLVAGHDPDHIIITNSVAQARFSELLSKLFITHLEKDGDLVTTLCPLVQTIKVRSEVENFKLALKGLAQHCEYAPFAQIVNLSGVNNLEIELFPQLSTTALVATAHQCILAGVTVGGQQQLRYGEAATEAERHLQEHSKQRTRNYSTGLDLTEARIVSD
FHKKKHEIGNQQQRFLSRGGGEKLQQLAADIINIKQSQPASQTPLPPQILAPHQPPPAEEHPASLDKQEVDLAKQELLEYDADLEVEDDNPNELPRNMKAPRINIPNFQSMPMQAGIDKERDHPSLSTTRKLPKIILTGHMEAPPGNQTVRRHRAFSIQSTSSMSGRELNTEPC
HNHHHSQDHRQSWAESFWPFQLENRSSIPKLSGENWAHPSFVEPLPPPIYPISGNEKGFAQQPQTRRFEALSPSNVTSKLQLIHPQQNLTALNGLKEAKETYHECVYSEGLHAGVSLYYSTMDGLKQCQSPGPNKPAGSECPCNPKEWWHLGEDNTLTLDGKKQHWSQLN
MPSQF
>JAJQIY010006009_Phodopus_sungorus
DLELHSILNLGTQPVESFISHNIYSFGVPDPIECGFTMDAVEAGAEFVDLLNRCLTLCIQHASKGNQKVFINSAARHYLENNGIICKVLDNNPISILNEFLQTQGMRQALLALVNDNHGHRKTGQSLSFCSLFLPKLVVGERACQKKKIQCQIVAHREQGLTQCPQGWNTPGSRMII
FRVMRVSFTFKFMLLHQGINLVAGHDPDNIIITNSVAQARFSGLLSKLFITHLGKDEDRVTLRLLVQTIQVKSKEENFKLALKGLARHCEYAPFAQIVSLSGVNKLEIELFPQLSTIALVATAHGCILAGVTVGKQYTAMLWEAATEAERRLQEHSEQRTRNYSTGLDLTEARILSDFH
KKKHEIGNQQAEIILVMKGGKLQQLAADIVNINQRQPASQAPLPPRILAPHQLPQAEEHPASLDKQEGDLAKQELLEYDADLEVEDDNPNELPVHESFQDQHPKFPILDADSERASIKRGIDPSLSATPKLPKIILTGYMEAPPANQTVRRHREFSTQSTRSSMSGKELNTEPCHN
HHHSQDLGSLGQRFLAIPARGRSSIPKLPGENWAHSSFVEPLPPPIHPISGNKKGVAQQPQTSQLEALSPSNVTSKLQLDTPPTKPDVSAQWLKEAKETYRKCVRGPHAGLSLYYSTPEQP
>MCBN011377448_Phodopus_sungorus
DLELHSILNLGTQPVESFISHNIYSFGVPDPIECGFTMDAVEAGAEFVDLLNRCLTLCIQHASKGNQKVFINSAARHYLENNGIICKVLDNNPISILNEFLQTQGMRQALLALVNDNHGHRKTGQSLSFCSLFLPKLVVGEGQSKKKIQCQIVAHREQGLTQCPQGWNTPGSRMIIF
RVMRVSFTFKFMLLHQGINLVAGHDPDNIIITNSVAQARFSGLLSKLFITHLGKDEDRVTLRLLVQTIQVKSKEENFKLALKGLARHCEYAPFAQIVSLSGVNKLEIELFPQLSTIALVATAHGCILAGVTVGKQYTAMLWEAATEAERRLQEHSEQRTRNYSTGLDLTEARILSDFHK
KKHEIGNQQAEIILVMKGGKLQQLAADIVNINQRQPASQAPLPPRILAPHQLPQAEEHPASLDKQEGDLAKQELLEYDADLEVEDDNPNELPVHESFQDQHPKFPILDADSERASIKRGIDPSLSATPKLPKIILTGYMEAPPANQTVRRHREFSTQSTRSSMSGKELNTEPCHNH
HHSQDLGSLGQRFLAIPARGRSSIPKLPGENWAHSSFVEPLPPPIHPISGNKKGVAQQPQTSQLEALSPSNVTSKLQLDTPPTKPDVSAQWLKEAKETYRKCVRGPHAGLSLYYSTPEQP
>CAJEUE010009159_Arvicola_amphibius
TSTWVALWFIPCCSQFVFEERVCLEKVKHQIAIHTEQGLIRYPSWHNSGFMMIIFGIMRAMQAFTFKFMLIHHGISLVAGHNADDIIITNSMAQALFSRLLIVKSVLDRVLHKADQITLHRLIQTIKVRTEADSFKLALKGLARHGEYAPFVQILNLSGVNSLEWGIFPQLSAIASGV
VTVCGDTLGDTLAGSVGQYQQLQEATSEADRCLQEYAESYRETLSRKTKKKKKREYAEKQEICQLGLDHAEAKSLLDFRKKVSIGNQQAELVLVTKREKLQQLIAAVKNTNQRTLILQDPPAPAPRQPFPPDVYTTVHKRSEN
>CATLKP010000280_Chionomys_nivalis
TSTWICALFIPCCSQLVFGERACLEKVKHQIAIHTEQGLIRYTGWHTSGFMMIIFGIMRAMQVCTFKLMLIHHGINLVAGHDIDDIIVTNLMAQAFSRLLIVKTVLDRVLHKERDRITLHRLIQTIKVRTEVDSFKLALKGLARHEDAPFVILNLSGVNNLEHGIFPRLSAIASGVATAH
GDTLGDTSAGITVGEQRHLQEAESEAADACKNMLRATEKPCLEKPKRKKKEYAEKQEICQLGLDHAELLDFHKKKVGIGNQRAKLVLVTKQEKLQQIIVAVNNTNQRNLILQDPPAPAPRQPFPPDVYTTVHKRSEN
>VIIU01015517_Microtus_oeconomus
TSTWICALRFIPCCSELVFGERACLEKVKRQIAIHTEQGLICCPGWHTSGFMIIFGIMRAMAFTFKFMLIHHDINLVAGNDIDDIIVTNLMAQAFSRLLIVKTVLDCVLHKEWDRITLHQLIQTIKVRMEVGSFKLALKGLARHQEDAPFVILNLSGVNNLEHGIFSRLSAIASGVVTA
HGDALGDTSAGVTVGEQYRQLQEAASEAAGACKNMLRATEKPCLEKPKRKKKKEYAEKQEICQLGLDHAEAKSLLDFHKKKVGIGNQRAKLVLVTKQEKLQQLIATVNNTNQRNLNLQDPPVPAPHQPFPPDVYTIVHKRSDN
>JABTXL010002019_Neodon_shergylaensis
TSTWICALWFIPCCSQLVFGERACLEKVKHQIAIHTEQGLICCPGWHTSGFMMIIFGIMRAMQAFTFKFMLIHHDINLVAGNDIDDIIITNLMAQARFPGLLIVKTVLDCVLHKERDRITLHRLIQTIKVRTEVDSFKLALKGLARHEDAPFVILNLSGVNNLEHGIFPRLSAIASGVA
TAHGDTLGDTSAGVTVREQYRQLQEAASEAAGACKNMLRATEKPCLEKPKRKKKEYAEKQEICQLGLDHAEAKSLLDFHKKKVGIGNQQAKLVLVTKQEKLQQLIATVNNTNQRNLNLQDPPVPAPHQPFPPDVYTTVHRRSEN
>VIIT010003993_Microtus_arvalis
TSTWICALWFIPCYSQLVFGERACLEKVKCQIAIHTEQGLICCSGWHTSGFMMIIFGIMRAMQAFTFKFMLIHHGINLVAGYDAHDIIVTNLLAQARFPGLLIVRSVLDRVLHKEQDRITLHRLIQTIKVRTEVDSFKLALKGLAHHEDAPFVILSLSGVNNLEHGIVPRLSAIASGVA
TAHGDTLGDTTAGVTVGEQYRQLQEATSEAEVPARICELQRNLVSKNQKEKKKKEYAEKQEICQLGLDHAEAKSLLDFHKKKVGIGNQQAELVLVTEQEKLQQLIATVNNTNQRNLNLQDPPVPAPHQPFPPDVYTTVHKRSEN
>JAEPQX010006744_Microtus_montanus
TSTWICTLWFIPCCSQLVFGERACLEKVKRQIAIHTEQGLICCPGWHTSGFMMIIFGIMRAMQAFTFKFMLIHHGINLVAEYDADDIIVTNLMAQARFPGSLPKLFIVYCTRNGTESLSIDSRQLKLECFTTALILQSMMLMILSLPIWLKPFSRLLIAKTVLDRVLHKERDRITLHRLI
QTIKVRTEVDSFKLALKGLARHEDAPFVILNLSGVNNLEHGIFPRLSATASEVATAHGDTLGDTSAGITVGEQYRQLQEATSEAEVPARICELQRNLVSKNQKEKKKEYAEKQEICQLGLDHAEAKSLLDFHKKKVGIRNQKAKLVLVIEQEKLQQLIATVNNTNQRNLNLQDPPAP
APRPFPPDVYTTVHKRSEN
>JAGKIF010000237_Microtus_oregoni
TSTWICALWFIPCCSQLVFGERVCLEKVKHQIAIHTEQGLICCPGWHTSGFMMIIFGIMRAMQAFTFKFRLIHHGINLVAGYDADDIIVTNLMAQARFPGLLIVKTVLDRVLHKERDRITLHRLIQTIKVRTEVDSFKLALKGLARHEDAPFVILNLSGVNNLEHGIFPRLSATASEVA
TAHGDTLGDTSAGITVGEQYRQLQEATSEAEVPARICELQRNLVSKNQKEKKKEYAEKQEICQLGLDHADAKSLLDFLKKKVGIRNQQAKLVLVTEQEELQQLIATVNNTNQRNLNLQDPPVPAPCQPFPPDVYTTVHKRSEN
>AHZW01020758_Microtus_ochrogaster
TSTWICALWFIPRCSQLVFGERACLEKVKRQIVIHTEQSLICCPGWHTSGFMMIIFGIMRAMQAFTFKFMLIHHGINLVAGYDADDIIITNLMAQARFPGLIIVKTVLDRVLHKERDRITLHRLIQTIKVRTEVDSFKLALKGLARHEDAPFVILNLSGVNNLEHGIFPELSATSSGVAT
AHGDTLGDTSAGITVGEQYWQLQDTTSEAEVPARICELQRNLVSKKPKRKKKKEYAEKQEICQLGLDHAEAKSLLDFHKKVGIGNQQAKLVLVTEQEKLQLIATVNNTNQRNLNLQDPPVPAPHQPFPPDVYTTVHKRSEN
>NMRL01000161_Microtus_fortis
TSTWICALWFIPCCSELVFGERACLEKVQIAIHTEQGLICCPGWHTSGFMMIIFGIMRAMAFTFKFMLIYHDINLVAGNDIDDIIVTNLMAQAFSRLLIVKTVLDCVLHKERDQITLHQLIQTIKVRTEVGSFKLALKGLARHQEDAPFVILNLSGVNNLEHGIFPRLSAIASGIMTAH
GDALGDTSAGVTVGEQYWQLQEAASEAEVPARICELQRNLVSRNQKEKKKEYAEKQEICQLGLDHAEAKSLLDFHKKKVGIGNQRAKLVLVTKQEKLQQLIATVNNTNQRNLNLQDPPVPAPHQPFPPDVYTTVHKRSDN
>CADCXR010038067_Microtus_agrestis
TSTWICALWFIPCCSQLAFGEGACLEKVKRQIAIHTEQGLICCPVWHTSGFIMIIFGIMRAMQAFTFKFMLIHHGINLVAGYDTDDIIVTNLMAQARFSGLLIVKTVLDHVLHKERDRITLHRLIQTLKVRTEVDSFKLALKGLAHHEDAPFVILNLSGVNNLEHGIFPRLSAIASGVA
TAHGDTGDTSAGVTVGEQYRQLQEATSEADRCLQEYAESYRETLSRKTQKKKKKEYAEKQEICQLGLDHAEAKSLLDFYKKRSGIGNQQAELVLVTEQETLQQLIATVNNTNQRNLNLQDPPVPVPRQPFPPDVYTTVHKRSEN
>CADCXT010005438_Microtus_agrestis
TSTWICALWFIPCCSQLAFGERACLEKVKRQIAIHTEQGLICCPVWHTSGFIMIIFGIMRAMQAFTFKFMLIHHGINLVAGYDTDDIIVTNLMAQARFSGLLIVKTVLDHVLHKERDRITLHRLIQTLKVRTEVDSFKLALKGLAHHEDAPFVILNLSGVNNLEHGIFPRLSAIASGVAT
AHGDTGDTSAGVTVGEQYRQLQEATSEADRCLQEYAESYRETLSRKTQKKKKKEYAEKQEICQLGLDHAEAKSLLDFYKKRSGIGNQQAELVLVTEQETLQQLIATVNNTNQRNLNLQDPPVPVPRQPFPPDVYTTVHKRSEN
>JAGDQN020000533_Microtus_richardsoni
TSTWICALWFIPCCSQLVFGERACLEKVKRQIAIHTEQGLICCPGWHTSGFMMIIFGIMRAMQAFTFKFMLIHQGINLVAGYDADDIIVTNLMAQAFSRLLIVKTVLDRVLHKERDQITLHRLIQTIKVRTEVDSFKLALKGLARHEDAPFVILNLSGVNNLEHGIFPRLPAIASGVA
TAHGDTLGDTSAGVTVGEQYRQLQEATSEADRCLQEYAESYRETLSRKTKKKQKQYAEKQEICQLGPNHAEAKSLLDFHKKKVGIGNQQAKLVLVTEQEKLQQLIATVNNTNQRNLNLQDPPVPAPRQPFPPDVYTTVHKRSEN

>JAQJZF010000005_Microtus_californicus
TSTWIYALWFIPCCSQLVFGERARLEKVKRQITIHTEQGLICCPGWHTSGFMMIIFGIMRAMQAFTFKFILIHHGINLVAGYDADDITVTNLMAQAFSRLLIVKTVLDRVLHKERDRITLHRLIQTIKVRTDVDSFKLALKGLARREDAPLVILRLSGVNNLEQGTCPRLSAIASGVAT
AHGDTLGDTSAGVTVGEQCRQLQEATSEAAGACKNMLRATEKPCLEKPKKKKKEYAEKQEICLGLDHAEAKSLLDFHKKKVGIGNPQAKLVLVTEQEKLPKLIATVNNTNQRNLNLQDPPVPAPIHQPFPPDGTPPCTRDQKIGIMGALPAYPLQGRIQQVPKLTINKHLQDEE
HVLSTFISRALSHHWEHHSRTLKTSSTNITLWRGPLFFKHRRSSAQNINTPGYQCSLHGGVST
>JAQJZG010000004_Microtus_californicus
TSTWIYALWFIPCCSQLVFGERARLEKVKRQITIHTEQGLICCPGWHTSGFMMIIFGIMRAMQAFTFKFILIHHGINLVAGYDADDITVTNLMAQAFSRLLIVKTVLDRVLHKERDRITLHRLIQTIKVRTDVDSFKLALKGLARREDAPLVILRLSGVNNLEQGTCPRLSAIASGVAT
AHGDTLGDTSAGVTVGEQCRQLQEATSEADRCLQEYAESYREALSRKTKKKKKKEYAEKQEICQLGLDHAEAKSLLDFHKKKVGIGNPQAKLVLVTEQEKLPKLIATVNNTNQRNLNLQDPPVPAPIHQPFPPDGTPPCTRDQKIGIMGALPAYPLQGRIQQVPKLTINKHLQDE
EHVLSTFISRALSHHWEHHSRTLKTSSTNITLWRGPLFFKHRRSSAQNINTPGYQCSLHGGVST
>LIPI01001523_Myodes_glareolus
QLVFGERACLEKVKHQVAIHAEQGLLRYPGWHTSGLMMIIFGIMRAMQAFTFKFMLLIHHGINLVAGHDADDIIITNLMAQALFSKLLIVKTVLDRVLHKEGDQITLHRLVQTIKVRTEVDSFKLALKGLARHREDAPFVRILNRSGVNNLEHGISPQLSAIASGVATARGDTLGD
SLAGVTVGEQYQQLQEAASEADRCLQEYAXXXXXXXXXXXXXXXXX
>MULK01030018_Myodes_glareolus
QLVFGERACLEKVKHQVAIHTEQGLLRYPGWHTSGLMMIIFGIMRAMQAFTFKFMLLIHHGINLVAGHDADDIIITNLMAQALFSKLLIVKTVLDRVLHKEGDQITLHRLVQTIKVRTEVDSFKLALKGLARHREDAPFVRILNRSGVNNLEHGISPQLSAIASGVATARGDTLGDS
LAGVTVEEQYQQLQEAASEAAGACKNMLRATEKPCLEKPKRKKKECAEKQEICQLGLGHAEAMILLDFHKKKVGIGNQQAELVLVTKEKLQQLIAVVNNTNQRNLILQDPPVPAPHQPFPPDVYTTMHRRSEN
>LOJG01023876_Ellobius_lutescens
TSTWVCALWFIPCCSQLVFGERACLEKVKRQIAIHTEQGLIRYPGWHTSGFMMIIFGIMRAMQTFTFKFMLIHHGINLVAGHDDIIITNLMSQALFSRLLIVKTVLERVLHKEGDQITLHRLIQTIKVRTEVDSFKLALKGLARHQEYAPFVRILLSGVNNLKHGIFPQLSAIASGVAT
TREDTFRETLAGVTVGKHYQQLQEAASEAEVPERICREPQRNLVSKNQKEKQKKEYAEKQEICQLGLDHAEAKSLLDFHKKKVGIGNQQAELVLVTKQEKLQQLIAAVNNTNQGNLILHDPLVPAPRQPFPPDVYTTVHKRAEN
>LOJH01006146_Ellobius_talpinus
TSTWVCALWFIPCCYQLVLGERACLETVKHQIAIHTEQGLIRYPGWHTSGFMMIIFGIMKSNASFYKFMLIHHGINLIAGHDADDIIITNLTSQALFSRLLIVKTVLDRVLHKEGDRITLHRLIQTIKVRTEVDSFKLALKGLARHREYCSFCMNTLSGVNNLKHGIFPQLSAIALIGVVT
VRGDTLGETLAGVTVGEQYQQLQEAASEAEVPARICELQRNLVSKKPKRKKKEYAEKQEICQLGLDHSEAKSLLDFHKKKVGIGNQQAELVLVTKQEKLQQLIAAVNNTNQRNLILHDPPVPAPRQPFPPNVYTNMHKRAEN
>PVIU01003686_Ondatra_zibethicus
IPAPPILTLSTSTWVCCALWFIPCCSQLVFGERACLEKVKRQIAIHTEQGLIRYPQGWHTSGFMMIIFGIMRAMQAFTFKFMLIHQGINLVAGHDADDIIITNLMAQALFSRLLIVKTVLDRVLHKEGDQITLHRLIQTIKVRTEVDSFKLTSNHLHPLGQTVKVRTEIDSFKLALKGL
ARHREYAPFVRILNLSGVNNLEHGIYPSAIASGVATARGDTLAGVTVGEQYQQLQGAASEVEVPARICELRNLVSKNQKEKKEREYAEKQEICQLGLYHAEAKILLDFHEKKVGIRNQQAELVLVTKQEKLQQLIAAVNNTNQRNLIPQDPPVPAPHQPFPPDDVHHRAHQKIGI
MGHCLEHIRNSKAEDNNPNEPSGHESSHDQCPKYHILDASSSSEEERGYRSQSTAPSFLINTRWPRGSPRSETKQSEGVESSQLNLQEANQHLMHNPTPIRGPTTTAPQTESSNTPGQKLSHPLQ
>JAQHUG010000003_Dicrostonyx_torquatus
WVCALWFIPCCSQLVFGERACLEKVKHQIAIHTEQGLIRYPQGWHTSGFMIFIFGIMRAMQAFTFKFMLIHQGINLVAGHDADDIIITNLMAQALFSRLIIVKTVLERVLYKEGDRITLHQLIQTIKVRTEVDSFKLASNHSPSTGPNNGNKIDSFKLALKGLACHQEYAPFVQILNL
SGVSNLEHGIFPQLSAIALGVATARGDTLTGVTVGEQYQQLQEAASEADRCLQEYAESYRETLSQKTKKKKTYAEKQEICQLGLDHAEAKILLDFHKKVGIGNQQAELVLVTKQEKLQQLIAAVKIPTRGIYPQDPPVPAPHQPFPPDGTPPCRRDQKIDVMGCCLEHIRNSKAED
NNPNELSGHESSHDQCPKYHILGACSSSEEERAQPPSFLINTYWPRGSPCSETKQSEGAGSSQLNSQEADQHLMNNSTPVRVIMATAPRTWESSDTPGQKLSSIPTLESPSAMPESPVENWACSPCEESLTT
>marburgvirus_Ravn_151564207_Lake_Victoria
MDLHSLLELGTKPSAPHVRNKKVILFDTNHQVSICNQIIDAINSGIDLGDLLEGGLLTLCVEHYYNSDKDKFNTSPIAKYLRDAGYEFDVIKNPDATRFLEVIPNEPHYSPLILALKTLESTESQRGRIGLFLSFCSLFLPKLVVGDRASIEKALRQVTVHQEQGIVTYPNHWLTTGHMK
VIFGILRSSFILKFVLIHQGVNLVTGHDAYDSIISNSVGQTRFSGLLIVKTVLEFILQKTDSGVALHPLVRTSKVKNEVASFKQALSNLARHGEYAPFARVLNLSGINNLEHGLYPQLSAIALGVATAHGSTLAGVNVGEQYQQLREAAHDAEVKLQRRHEHQEIQAIAEDDEERKILE
QFHLQKTEITHSQTLAVLSQKREKLARLAAEIENNIAEDQGFKQSQNQVSQSFLNDPTPVEVTVQARSINRPTALPPPVDNKIEHETEEDSSSSSSFVDLNDPFALLNEDEDTLENSVMAPSTTLREPKEVSEPLRQTQDLDISQKKQGNESTDPARKQFLRYQELPPVQEDDESEY
TTDSQESDDQPGSDNEQGVDLPPPPLYAQEKRQDPIQHPAVSSQDPFGSIGDVDGDILEPIRSPSSPSAPQEDTRMGEAYELSPDFTSYEDNQQNWPQRVVTKKGRTFLYPNDLLQTSPPESLITALVEEYQNPVSAKELQADWPDMSFDERRHVAMNL
>YP_010087183_Mengla_Dianlovirus
MDLHGLLELGTRPTAPHVRSRKIVIYETGNQIIICNQIIDAVSAGIDLGDLLEGCLLTLCLEHYYGSDKDKFNSSQMAAYLRDAGYDFEVIRAQDAKKLADLIPRESHLLNVISALENLDGSEKNKQRVGLFLSFCSLFLPKLVVGDKASIEKALRQVAIHQEQGMVVYPPTWLTTGF
MKLIFSIVRASFIVKFVLIHQGINLVTGHDAYDNVISNSINQTRFSGLLIVKTVLEHILQRTENGVILHPLVRTSKVKSEVESFKVALRGLARHKEYAPFARVLNLSGVNNLEHGLFPQLSAIAIGVATAHGSTLAGVSVGEQYQQLREAAHDAELKLQRRREQMEISSLELDLEEQKILE
QFHQQKTEITHTQTLAVLTQKKEKLAKLAGELGADKLFQPVIPQPNDKPQMTTVIDPSRTSVKIQTSFLPPPSTKDNMEGQIPESPESSTISSSSSCIDPNDPFALLIDDDEQEGDFQSADEGDSQDQSAEAARQEEIRKAEQKFKGLPRTIPTKDQNPPIKPQRTNLAPVQEESESE
YTTTSDESGSEDESSDQRNIEVPPPPLYDTSTSEIGDMGAAEHTLDPFGDMPPLETDVLTPQDTATVPSAPPSSPTPSTNRQGEAQTQNGEDSSQDWPRRVKTNKGREFMFPTDLLHRTPPQVLLDALVNEYESPLSATELSDDWPEMTFEERKNVAFNL
>BH05257_Bombali_ebolavirus
MEVRNPRQWTTQSASDSSVDYHSILTAGLSMPQSIVRQRVIPVFQVSNLEDICQMIIQAFEAGVDFQDSADSFLLMLCLHHAYQGDYKLFLESGAVKYLEGHGFRFELRKKEGVKRLEELLPAVTNGKNIRRTLAAMPEGETTEANAGQFLSFASLFLPKLVVGEKACLEKVQRQ
IQVHAEQGLIQYPTSWQSVGHMMVIFRLMRTNFLIKFLLIHQGMHMIAGHDANDAVIANSVAQARFSGLLIVKTVLDHILQKTDRGVRLHPLARTAKVRNEVHSFKAALNSLAQHGEYAPFARLLNLSGVNNLEHGLYPQLSAIALGVATAHGSTLAGVNVGEQYQQLREAA
TEAEKQLQQYAETRELDHLGLDTKEKEILMNFHQRKNEISFQQTNAMVSLRKERLAKLTEAIAAASAQRERGYYDDDNEIPFPGPINDNDDQDQHVDDPTDTQDTTIPDIVVDPDDGGYRGYQIRQDDDMDAPDDLVLFDLDDNEQGPPSENARPGFTVERGPRSSGQQKS
TEQHQPPLSSDLQAPTYNHHSREHHADQGTRSHELRSVQESDLLASIPEEHHPADHTDDKASLSLPPDSESDGESSPDTDEADRTTAPPAPVYKNHKDAGPATVGETNKETDFDDNINNQPESPLNDHGIERMYRHILETQGPFDAILYYHMMKDEPVTFSTNDGKEYIYPDSL
ENEYPPWLSEKEATQSESRFINIDGQQFYWPVMNYRDKFMAILQHHS
>499104233_Bundibugyo_ebolavirus
MDPRPIRTWMMHNTSEVEADYHKILTAGLSVQQGIVRQRIIPVYQISNLEEVCQLIIQAFEAGVDFQDSADSFLLMLCLHHAYQGDYKQFLESNAVKYLEGHGFRFEMKKKEGVKRLEELLPAASSGKNIKRTLAAMPEEETTEANAGQFLSFASLFLPKLVVGEKACLEKVQRQI
QVHAEQGLIQYPTSWQSVGHMMVIFRLMRTNFLIKFLLIHQGMHMVAGHDANDAVIANSVAQARFSGLLIVKTVLDHILQKTEHGVRLHPLARTAKVKNEVSSFKAALASLAQHGEYAPFARLLNLSGVNNLEHGLFPQLSAIALGVATAHGSTLAGVNVGEQYQQLREAAT
EAEKQLQKYAESRELDHLGLDDQEKKILKDFHQKKNEISFQQTTAMVTLRKERLAKLTEAITSTSILKTGRRYDDDNDIPFPGPINDNENSGQNDDDPTDSQDTTIPDVIIDPNDGGYNNYSDYANDAASAPDDLVLFDLEDEDDADNPAQNTPEKNDRPATTKLRNGRDQDG
NQSETASPRAAPNQYRDKPMPQVQSRSENHDQTLQTQPRVLTPISEEADPSDHNDGDNESIPPLESDDEGSTDTTAAETKPATAPPAPVYRSISVDDSVPLENIPAQSNQTNNEDNVRNNAQSEQSIAEMYQHILKTQGPFDAILYYHMMKEEPIIFSTSDGKEYTYPDSLEDE
YPPWLSEKEAMNEDNRFITMDGQQFYWPVMNHRNKFMAILQHHR
>Cote_d'Ivoire_ebolavirus_302315370
MESRAHKAWMTHTASGFETDYHKILTAGLSVQQGIVRQRVIQVHQVTNLEEICQLIIQAFEAGVDFQESADSFLLMLCLHHAYQGDYKQFLESNAVKYLEGHGFRFEVRKKEGVKRLEELLPAASSGKSIRRTLAAMPEEETTEANAGQFLSFASLFLPKLVVGEKACLEKVQRQI
QVHSEQGLIQYPTAWQSVGHMMVIFRLMRTNFLIKFLLIHQGMHMVAGHDANDAVIANSVAQARFSGLLIVKTVLDHILQKTEHGVRLHPLARTAKVKNEVNSFKAALSSLAQHGEYAPFARLLNLSGVNNLEHGLFPQLSAIALGVATAHGSTLAGVNVGEQYQQLREAAT
EAEKQLQKYAESRELDHLGLDDQEKKILKDFHQKKNEISFQQTTAMVTLRKERLAKLTEAITSTSLLKTGKQYDDDNDIPFPGPINDNENSEQQDDDPTDSQDTTIPDIIVDPDDGRYNNYGDYPSETANAPEDLVLFDLEDGDEDDHRPSSSSENNNKHSLTGTDSNKTSNWNR
NPTNMPKKDSTQNNDNPAQRAQEYARDNIQDTPTPHRALTPISEETGSNGHNEDDIDSIPPLESDEENNTETTITTTKNTTAPPAPVYRSNSEKEPLPQEKSQKQPNQVSGSENTDNKPHSEQSVEEMYRHILQTQGPFDAILYYYMMTEEPIVFSTSDGKEYVVYPDSLEGEHP
PWLSEKEALNEDNRFITMDDQQFYWPVMNHRNKFMAILQHHK
>436409350_Zaire_ebolavirus
MDSRPQKVWMTPSLTESDMDYHKILTAGLSVQQGIVRQRVIPVYQVNNLEEICQLIIQAFEAGVDFQESADSFLLMLCLHHAYQGDCKLFLESGAVKYLEGHGFRFEVKKRDGVKRLEELLPAVSSGKNIKRTLAAMPEEETTEANAGQFLSFASLFLPKLVVGEKACLEKVQRQI
QVHAEQGLIQYPTAWQSVGHMMVIFRLMRTNFLIKFLLIHQGMHMVAGHDANDAVISNSVAQARFSGLLIVKTVLDHILQKTERGVRLHPLARTAKVKNEVNSFKAALSSLAKHGEYAPFARLLNLSGVNNLEHGLFPQLSAIALGVATAHGSTLAGVNVGEQYQQLREAAT
EAEKQLQQYAESRELDHLGLDDQEKKILMNFHQKKNEISFQQTNAMVTLKKERLAKLTEAITAASLPKTSGHYDDDDDIPFPGPINDDDNPGHQDDDPTDSQDTTIPDVVVDPDDGSYGEYQSYSENGMNAPDDLILFDLDEDDEDTKPVPNRSTKGGQQKNSQKGQHTEG
RQTQSRPTQNIPGPHRTIHHASAPLTDNDRRNEPSGSTSPRMLTPINEEADPLDDADDETSSLPPLESDDEEQDRDGTSNRTPTVAPPAPVYRDHSEKKELPQDERQDQDHTQEARNQDSDNTQPEHSFEEMYRHILRSQGPFDAVLYYHMMKDEPVVFSTSDGKEYTYPDS
LEEEYPPWLTEKEAMNEENRFVTLDGQQFYWPVMNHKDKFMAILQHHQ
>55247456_Reston_ebolavirus
MDRGTRRIWVSQNQGDTDLDYHKILTAVLTVQQGIVRQKIISVYLVDNLEAMCQLVIQAFEAGIDFRENADSFLLMLCLHHAYQGDYKLFLESNAVQYLEGHGFKFELRKKDGVNRLEELLPAATSGKNIRRTLAALPEEETTEANAGQFLSFASLFLPKLVVGEKACLEKVQRQI
QVHAEQGLIQYPTAWQSVGHMMVIFRLMRTNFLIKYLLIHQGMHMVAGHDANDAVIANSVAQARFSGLLIVKTVLDHILQKTDQGVRLHPLARTAKVRNEVNAFKAALSSLAKHGEYAPFARLLNLSGVNNLEHGLYPQLSAIALGVATAHGSTLAGVNVGEQYQQLREAA
TEAEKQLQQYAESRELDSLGLDDQEGRILMNFHQKKNEISFQQTNAMVTLRKERLAKLTEAITLASRPNLGSRQDDGNEIPFPGPISNNPDQDHLEDDPRDSRDTIIPNGAIDPEDGDFENYNGYHDDEVGTAGDWVLFDLDDHEDDNKAFEPQDSSPQSQREIERERLTHPP
PGNNKDDNRASDNNQQSADSEEQGGQYNWHRGPERTTANRRLSPVHEEDTLMDQGDDDPSSLPPLESDDDDASSSQQDPDYTAVAPPAPVYRSAEAHEPPHKSSNEPAETSQLNEDPDIGQSKSMQKLEETYHHLLRTQGPFEAINYYHMMKDEPVIFSTDDGKEYTYP
DSLEEAYPPWLTEKERLDKENRYIYINNQQFSWPVMSPRDKFLAILQHHQ
>165940955_Sudan_ebolavirus
MDKRVRGSWALGGQSEVDLDYHKILTAGLSVQQGIVRQRVIPVYVVNDLEGICQHIIQAFEAGVDFQDNADSFLLLLCLHHAYQGDHRLFLKSDAVQYLEGHGFRFEVREKENVHRLDELLPNVTGGKNLRRTLAAMPEEETTEANAGQFLSFASLFLPKLVVGEKACLEKVQR
QIQVHAEQGLIQYPTSWQSVGHMMVIFRLMRTNFLIKFLLIHQGMHMVAGHDANDTVISNSVAQARFSGLLIVKTVLDHILQKTDLGVRLHPLARTAKVKNEVSSFKAALGSLAKHGEYAPFARLLNLSGVNNLEHGLYPQLSAIALGVATAHGSTLAGVNVGEQYQQLREAA
TEAEKQLQQYAETRELDNLGLDEQEKKILMSFHQKKNEISFQQTNAMVTLRKERLAKLTEAITTASKIKVGDRYPDDNDIPFPGPIYDDTHPNPSDDNTDDSRDTTIPGGVVDPYDDESNNYPDYEDSAEGTTGDLDLFNLDDDDDDSRPGPPDREQNKERAARTYGLQDPTLD
GAKKVSELTPGSHQPGNLHITKSGSNTNQPQGNMSSTLHSMTPIQEESEPDDQKDNDDESLTSLDSEGDEDGESISEENTPTVAPPAPVYKDTGVDTNQQNGPSSTVDSQGSESGALPINSKKSSALEETYYHLLKTQGPFEAINYYHLMSDEPIAFSTESGKEYIFPDSLEEAYPP
WLSEKEALEKENRYLVIDGQQFFWPVMSLQDKFLAVLQHD
>355469072_Lloviu_virus
MNRYLGHGTRTSRENTNLSELHGILSLGLNVDHTIVRKKSIPLFEIGNSDQVCNWIIQIIEAGVDLQDVADSFLTMLCVNHAYQGDPNLFLESPAAHYLKGHGIHFEIQHRDNVDHITDLLGVGSRDKSLRKTLSALEFEPDGSTTAGMFLSFASLFLPKLVVGERACLEKVQRQIQI
HAEQGLIQYPTQWQSVGHMMVVFRLIRVNFVLKFLLVHQGMHMMAGHDANDAIIANSISQTRFSGLLIVKTVLEHILQKTEAGVQLHPLARTSKVKGELLAFKSALEALASHREYAPFARLLNLSGVNNLEHGLYPQLSAIALGVATAHGSTLAGVNVSEQYQQLREAATEAEK
QLQQHSEMRELETLGLDEQERKILATFHSRKNEINIQQTSSILAIRKERLRKLTEALNEEKNKNALDDEDESEEDDWSPENRGIRSNKGSTKESSSYTASRTEEDRNNYNSKDDHLSGKEQMSTQQESGADDLDLFDLDDDGDTNSQDPNNRQKQSDTQQTQESSDRSDYSRRP
AYDWPPGDRPHTTQATDEHTDLLNKDHRRNQVKPGRRGNDPRTLPLISFDDNEGEILDDKSDLPAPDTHSDPDTEESEEEHPDEELLPPAPKYNTKTSEQEPGDWKQPTSPLSTIPEEEGGHEANNDNSESDLIDQMYQHIFETEGAYAAINYYYKTTGRPVTFTSNNNHDYTF
PQDIEGLFPPWEGKENQKVAEILTNSLHETGQEWADMSAKERYLFLINN
>CAJEUE010013052_Arvicola_amphibius
FGGRSHFSFSNPRELRAAHIQLQVRNQWNPSHRAPEPTELCPQIIHAIQAGVELGDLLDSCLLTLRTQHPEGNSGNFEQRTARRYLERHGLVFHIIEKTTSNLTKFCLDTGLGSTQQALLALANNHNEHWKVGQFLSFCSLFLLLKLVVGEKACLEKVQCQITVHSEQGLIHPQGW
NTLGFMMVTSGIMRASFTFKFMLTHQGTNVEAGHDSAISISNSLAQGRCSGLLTVTTVLDHIFRKEGNQITLYSCRTVMVRAEVDSFKLALQGLAKEAPFARSLNLGVNSLEHGLFPQLSAVALGTATAHGGTLVGVTVVEQYQQLWEAASEAERRPKKHAEEQEIDQLDLDH
AEQGILVDFHTKKTNKKNPPKPHNIGHQGRDDACSKVGGKIQQLAASVHTADQGSPTPQGSSMSTPAVRPFEYFYGPRSGNQCQGARPPTPGFSCRRRSNNNHGSDVHDQDSSSQILDASSDDEYEEDYSPLIATPDFLVGPYWSRKSPVDETKQPEEKSSQNDSVDTSQQ
TPQDQAFPMRQQQQPWGCRKSPDASGQKLPPALTRENLTAIPEPGSWTYSYFTEPPPPPVYPSSDSDKETPQSPQVDPLGNTTEVPTGAATVTPNLTTEQISRTEQEHQNITANEGLWAGQLFYHRPTGRPKDFPTANGAIIQFPTSLRATAPINNQEGWHFWTDGTIAMD
GQMPWCDLDLQSRLVYI
>CATLKP010001056_Chionomys_nivalis
PSHRAPEPTELCLQIIHAKQAGVELGDLLDSCLLTLCTQYPEGNLGNFEQRAARHYLNGHRLLFHIFENKTTSNLTEFCLDTELGSTQQALLALANNHNEHWKVGQFLSFCSLFLPKLVVGERACLEKVQCQIPIHAEQGLTQHPQGWNTPGFMMMISGIMWASFTFKFMLTH
QGTNVEAGHDSAISISNPVAQGRRSGLLTVTTVLDHIFRKEGNQITLYSCRTVMVQAEVDSFKLALQELAKEYAPSARSLNLGVNSLEHGPFPQLSAVAPGTLVGVTVGEQYQQLWEAASEAERCPKKHAEEQEIDQLDLDHAEQGILVDFHTKKKTINPKTTTLDIKAEMALAA
KRGKKNPTTCSFGSHCKSREPNALPMSAPQQPAPSNTSTRGPDDQETSARERGLQHLDDLAAGGKDPIITHGSDMQDQDPSSPDLGYKLRIRRLWLTHCHSFSGWTSLVTQEPSDATRKKLPKRLSKHSADTPRSGFSYETTATTSGLQESPDASGQKLPPALTRENLAAIPEP
GQRWTHSYFIEPPPPPVYPSLDIDKETSQSPQVDPLEIPLKSPTGAPSITPNLTTEQISKMEQEYQSIMANEGLWVGQLFPHRRTGRTKDFPTSNGAILQFPTSLIGATAPINNQEGWHFGTDGTITMDGQMPLWSDLDLQSRLVYIV
>VIIT010000186_Microtus_arvalis
PSHRAPEPTELCLQIIRAIQAGVELGDLLDSCLLTLRTQHLEGNSGNFEQRAARRYLERHRLVSRILENKTTSNVTDFCPDTGLGSTQQALLTLANNHNEHWKVGRFLSSCSLFLTKLVVGERALLEKVQCQIPVHAKQGLTQHPQGWNTPGFMMVISGIMRASFTFKFMLTHQ
GTNMETGHDSAISISNPVAQGRCSGLLTVTTVLDHIFRKEGNQITLYSCRTVMVQAEVDSFKLALQELAKEYAPSTRSLNLEVNSLEHRQLSAVALGTATAHGGTLVGVTVGEQYQQLWEAASKAERRPKKHAEEQEIDQLDLDHAEQGILLDLHTKKKKKKNPKTTTLDIKVRD
DACSKEGEKIQQLAANQRSPTPQGSSMSAPQQPAPSNTSTRGPDDQETSAREGLQHLDDLAAGGKDPIITHGSDVQDQDPSSPDLRCKLRVRGLWLTHCHSFSGWTSLVTQEPSDATRKKLPKRLSGHSADTPRSGFSYETTATTSGLQESPDASGQKLPPALTRENLTAIPET
GSWTHSYFIEPPSPPVYPSSDIDKETSQSPSSRSFRNTLKSPTGAPSVTPNLTTEQISKMEQEYQNIMANEGLWAGQLFHHRTGRPKDFPTSNGATLQFPTSLTGATAPINNQEGWHFGTNGTIAMDGQMPRSDLDLQSRLVYIALIIH
>CADCXR010062301_Microtus_agrestis
PSHRAPEPTELCLQIIRAIQAGVELGDLLDSCLLTLRTQHPEGNSGNFEQRAARRYLERHRLVSHILENKTTSNMTDFCLDTGLGSTQQALLTLANNHNEHWKVGRFLSFLPIFLPKLLVGERACLEKVQCQIPVHAEQGLTQHPQGWNTPGFISGIMRASFTFQFMLTHQGTN
METGRDSAISISNPVAQGRCSGLLTVTTVLDHIFHKEGNITLYSCRTVMVQAEVDSFKLAGAGQEYAASARSLNLGVNSLEHGLSPQLSAVAPGIATAHGGTLVGVTVGEQYQQLWEAASEAERRPKKHAEEQEIDQLDLDHAEQGILVDFHTKKKNPNPKTTTLDIKAEMTLA
AKRVEKIQQLAALVHTANQGSPTPQGSSMSAPQQPAPSNTSTRGPDDQETSAREGLQHLDDLAAGGKDPIITHGSDVQDQDPSSQILDASSDDEYEEDYGSLIATPDFLVGPHWSRKGPVDETKQPEERSSQNDSVDTSQQTPQDQAFPVRQQQQPRGCRKAQMLQGKS
FHLLPGKTSLQSLEPGSWTHSYFIEPPPPPVYPSSDIDKETSQSPQVDPLEYHSPQLVLPHHSTKPEQISKMEQEYQSIMANEGLWAGQLFHHRRTGRPKDFPTSNGAIHLQFPTSLTGATAPINNQEGWHFEDRWDIAMDGQMPWSDLDLQSRLVYI
>CADCXS010119444_Microtus_agrestis
PSHRAPEPTELCLQIIRAIQAGVELGDLLDSCLLTLRTQHPEGNSGNFEQRAARRYLERHRLVSHILENKTTSNMTDFCLDTGLGSTQQALLTLANNHNEHWKVGRFLSSCSLFLPKLVVGERACLEKVQCQIPVHAEQGLTQHPQGWNTPGFISGIMRASFTFQFMLTHQGTN
METGRDSAISISNPVAQGRCSGLLTVTTVLDHIFHKEGNITLYSCRTVMVQAEVDSFKLAGAGQEYAASARTLNLGVNSLEHGLSPQLSAVAPGIATAHGGTLVGVTVGEQYQQLWEAASEAERRPKKHAEEQEIDQLDLDHAEQGILVDFHTKKKKPKPKNHNIGHQGRDDT
CSKEGEKIQQLAALVHTANQGSPTPQGSSMSAPQQPAPSNTSTRGPDDQETSAREGLQHLDDLAAGGKDPIITHGSDVQDQDPSSQILDASSDDEYEEDYGSLIATPDFLVGPHWSRKGPVDETKQPEERSSQNDSVDTSQQTPQDQAFPVRQQQQPRGCRKAQMLQGK
SFHLLPGKTSLQSLEPGSWTHSYFIEPPPPPVYPSSDIDKETSQSPQVDPLEYHSPQLVLPHHSTKPEQISKMEQEYQSIMANEGLWAGQLFHHRRTGRPKDIPTSNGAIHLQFPTSLTGATAPINNQEGWHFEDRWDIAMDGQMPWSDLDLQSRLVYI

>JAQJZF010000004_Microtus_californicus
FGGRSHFSFSNPRGLRAAHIQLQDTTSRILPHRAPEPTELCLQIIRAIQAGVELGDLLDSCLLTLCTQHPGTSGNFEQRAARRYLERHRLVFRILENKTTSNLTEFCLDTALGSTQQALLALANNHNEHWKVGQFLSSCSLFLPKLVVGERACLEKVQCQIPVHAEQDLIQHPQGLNT
PGFMMVISGITQASFTFKFMLTHQGTNVEAGHDSASSISNPVAQGRCSGLLTDKTVLDHIFRKEGNQITLYSCRTVMVQAEVDSFKLALQELGQEYAPSARSLNLGVNSLEHGLFPLSAVAPGTATAHGGTLVGVTVGEQYQQLWEAASEAERCPKKHAEEQEIDQLDLDHAE
QGILVDFHTKKKKQNKTQKPQHWTSRQRCLQQRVGKRIQQLAALVHTANQGSPTPQGSSMSAPQQPAPSNTSTRGPDDQETSAREGLQHLDDLAAGGKDPNNNPWQRCARPGSLFSDLDASSDDEYEEDYGSLIATPGFLVGPHWSRKSPVDETKQPEERSSQNDSVDT
SQHTPQDQAFPMRQQQQPRGCRKAQMLQGKSSPPALTRENLTAIPETGNWTHSYFIEPPPPPGYPSLDIDKETSQSPQVDPLEIPSPQLVLPQHPTQPEQISKMEQEYQNIMANEGLWAGQLFHHRTGRPKDFPTSNGAILQLPTSLTGATAPINNQEGWHFGTDGTIAIDG
QMLWSDLDLQSRLVYIALIIH
>JAQJZG010000003_Microtus_californicus
FGGRSHFSFSNPRGLRAAHIQLQDTTSRILPHRAPEPTELCLQIIRAIQAGVELGDLLDSCLLTLCTQHPEGTSGNFEQRAARRYLERHRLVFRILENKTTSNLTEFCLDTGLGSLTQQALLALANNHNEHWKVGQFLSSCSLFLPKLVVGERACLEKVQCQIPVHAEQDLIQHPQG
WNTPGFMMVISGITQASFTFKFMLTHQGTNVEAGHDSASSISNPVAQGRCSGLLTDKTVLDHIFRKEGNQITLYSCRTVMVQAEVDSFKLALQELGQEYAPSARSLNLGVNSLEHGLFPLSAVAPGTATAHGGTLVGVTVGEQYQQLWEAASEAERCPKKHAEEQEIDQLDL
DHAEQGILVDFHTKKKNKTKPKNHNIVTSRQRCLQQRVGKRIQQLAALVHTANQGSPTPQGSSMSAPQQPAPSNTSTRGPDDQETSAREGLQHLDDLAAGGKDPIITHGSDVQDQDPSFQILDASSDDEYEEDYGSLIATPGFLVGPHWSRKSPVDETKQPEERSSQNDSVD
TSQHTPQDQAFPMRQQQQPLGLQKSPDASGQKLPPALTRENLTAIPETENWTHSYFIEPPPPPGYPSLDIDKETSQSPQVDPLEIPSPQLVLPQHPTQPEQISKMEQEYQNIMANEGLWAGQLFHHRTGRPKDFPTSNGAILQLPTSLTGATAPINNQEGWHFGTDGTIAIDG
QMLWSDLDLQSRLVYIALIIH
>JAEPQX010010882_Microtus_montanus
FGGRSHFSFSNPRGLRAAHIQLQDTTSGILPHWAPEPTELCLQIIRAIQAGVELGDLLDSCLLTLRTQHPEGNSGNFEQRAARRYLEQHRLVFCILENKTTSNLTEFCLDTGLGSTQQALLALASNHNEHWKVGQFLSSCSLFLPKLVVGERACLEKVQCQIPVHAEQGLIQHPQG
WNTPGFMMVISGITQASFTFKFMLTHQGTNVEAGHDSASSISNPVAQGRCSGLLTVKTVLDHIFRKEGNQITLYSCRTVMVQAEVDSFKLALQELAKEYAPSARSLNLGVNSLEHGLFPQLSAVAPGTATAHGGTLVGVTVGEQYQQFQATSKAERRPKKHAEEQEIDQLDLD
HAEQGILVDFHTKKKKKKKKPKNHNIGHQGRDDACSKEGEKIQQLAALVHTANQGSPTPQGSSMSAPQQPAPSNTSTRGPDDQETSAREGLQHLDDLAAGDKDPIITHGRDVQDQDPSFQILDASSDDEYEEDNGSLIATPGFLVGPHWSRMSPVDETKQPEERSSQIRLSG
HSADTPRSSFSYETTATTSGLQESPDASGQKLPPALTRENLTAIPETGSWTHSYFIEPPPPPVYPSLDIDKETSQSPQVDPLEIPSPQLVLPQHPTQPEISKMEQEYQNIMANEGLWAGQLFHHRTGRPKDFPTSNGAILQFPTSLTGTTAPINNQEGWHFGTDGTIAMDGQML
WSDLDLQSRLVYIALIIH
>JAGDQN020000253_Microtus_richardsoni
PSHRAPEPTELCLQIIRAIQAGVELGDLLDSYLLTLRTQHPEGNSGNFEQRAARHYLEWHRLVFCILENKTTSNLTEFCLDTGLGSTQQALLALANNHNEHWKVGQFLSSCSLFLLKLVVGERACLEKVQCQIPVHTEQGLIQHPQGWNTPGFMMVISGIMQASFTFKFMLTHQ
GTNVEAGHDSASSISNPVAQGRCSGLLTVKTVLDHIFRKEGNQITLYSCRTVMVQVEVDSFKLTLQELAKEYAPSARSLNLGVNSLEHGLFPQLSAVALGTATAYGGTLVGVTVGEQYQQFQAASKAERRPKKHAEEQEIDQLDLDHAEQGILVNFHKKTKTKTTTLDIKAEMTL
AAKRVEKIQQLAALVHTANQGSPTPQGSSMSAPQQPAPSNTSTRGPDDQETSAREGLQHLDDLAAEGKDPIITHGSDVQDQDPSFQILDASSDDEYEEDYGSRHCHTPGFLVGPHWSRKSPVDETKQPEERSSQIRLSRHSADTPRSGFFYETTATTSGLQESPDASGQKLPP
ALTRENLTAIPETGSWTHSYFIEPPPPPVYPSLDIDKETSQSPQVDPLEIPSPQLVLPQHPTQPEQISKMEQEYQNVMANEGLWAGQLFHHRTGRPKDFPTSNGAILQFPTSLTGATAPINNQEGWHFGTDGTIAMDRQMLWSDLDLQSRLVYIALIIH
>AHZW01126995_Microtus_ochrogaster
PSHRAPEPTELCLQIIHAIQAGVELGDLLDSCLLTLLTQHPEGNSGNFEQRAARRYLERHRLVFCILENKTTSNLTEFCLDIGLGSTQQALLALANNHNELESGQFLSFCSLFLPKLVVGERACLEKVQCIPVHAKQGLIQHPQGWNTPGFMMVISGITQASFTFKFMLTHQGTNVE
AGHDSASSISNSVAQGRCSGLLTVKTVLDHIFRKESNQITLYSLALSTIVQAEVDSFKLALQELAKEYAQSARSLNLGVNSLEHRLFPQLSAIAPGTATAHGGTLVGVTVGEQYQQFAASKAERRPKKHAEEQEIDQLDLDHAEQGILVDFHTKKQNINPKTTTLDIKAEMTLAAKG
EKIQQLAALVHTANQGSPKPQGSSMSAPQQPAPSNTSTRGPDDQETSAREGLQHLDDLAAGDKDPIITHGSDVQDQDPSFQILDASSDDEYEEDYGSLIATPGFLVGPHWSRKSPVDETKQPEERSSQIRLSGHSADTLRSGFSYETTATTSGLQESPDASGQKLSPALTKENLT
AIPETGSWTHSYFIEPPPPPVYPSLDIDKETSQSPQVDPLEISSPQLVLPQHPTQPEQISKMEQEQNIMANEGLRAGQLFHHRTGRPKDFPTSNGAILQFPTSLTGATAPINNQEGWHFGTDGTIAIDGQMLWSDLDLQSQLVYIALIIH
>JAGKIF010000135_Microtus_oregoni
PSHRAPEPTELCLQIIRAIQAGVELGDLLDSCLLTLHTQHPEGNSGNFEQRVARRYLERHRLVFCILENKTTSNLTEFCLDTGLGSTQQALLALANNHNEHWKVGQFLSSCSLFLPKLVVGERVCLEKVQCQIPVHAEQGLIQHPQGWNTPGFMMVISGITQASFTFKFMLTHQG
TNVEAGHDSASSISNPVAQGRCSGLLTVKTVLDHIFRKEGNQITLYSCRTVMVQAEVDSFKLALQELAKEYAPSARSLNLGVNSLEHRLFPQLSVVAPGTATAHGGTLVGVTVGEQYQQFQAASKAERRPKKHAEEQEIDQLDLDHAEQGILVDFHTKKKNKTKLQHWTSRQR
RLQRGGKNPTTCSFGSHCKSREPNTSRLFYVKPPQQPAPSNTSTRGPDDQETSAREGLQHLDDLAAGGKDPIITHGRDVQDQDPSFQILDASLDDEYEEDYGSLIATPGFLVGPYWSRKSPVDETKQPEERSSQIQLSGHSADTPRSGFSYETTATTLGLQESPDASGQKLPPALAR
ENLTAIPETGSWTHSYFIEPPPPPVYPSLDIDKETSQSPQVDPLEIPSPQLVLPQHPTQPEQISKMEQEYQNIMANEGLWAGQLFHHRTGRPKDFPTSNGAILQFPTSLTGATAPINNQEGWHFGTDGTIAMDGQMLWSDLDLQSRLVYIALIIH
>NMRL01000046_Microtus_fortis
PSHRAPEPTELCLQIIRAIQAGVELGDLLDSCLLTLRTQHSEGNSGNFEQRAARYYLERHRLVSRILENKTTSNLTDFCLDTGLGSTQQALLALANNHNEHWKVGQFFLPVALFLPKLVVGERACLEKVQCQIPAHAEQGQIQHPQGWNTPGFMMVISGIRRASFIFKFMLTHQG
TNMETGHDSAISISNPVAQGRCSGLLTVTTVLDHTFRREGNQITLYSCRTVMVQAEVDSFKLALQELAKEYAPSARSLNLGVNSLEHGLFPQLSAVAPGHGGTLVGVTVGEQYQQLWETASKAERRPKKHAEEQEIDQLDFDHAEQGILVDFQKKKKKKTQNPKTTTLDIKAGR
LQQRGGKIQQLAALVHTANQGSPTPQGSSMSAPQQPAPSNTSTRGPDDQETSARDKPPTPGFSCRKRSNNNPWQRCALGSSSQILDASSDDECEEDYGSLTATPGFLVGPHWSRKSPVDETKQPEKRSSQIRLSGHSADTPRSGFAYETTVTTSGLQESPDASGQKLPPALTR
ENLTAIPETGSWTHSYFIEPPPPPVYPSLDIDKETSQSPQVDPLEIPLKSPTGAPSVTPNLTTEQIGKMEQERSIMANEGLWAGQLFHHRPTGRPKDFPTWNGAILQFPTSLTGATAPINNQEGWHFGTDGTIAMEGQMPWSDLDLQSRLVHI
>VIIU01009709_Microtus_oeconomus
PSHRAPEPTELCLQIIRAIQAGVELGDLLDSCLLTLRTQHPEGNSGNFEQRAARCYLERHRLVSCILENKTTSNLTDFCLDTGLGSTQQALLALANNHNEHWKVGQFLSSCSLFLPKLVVGERACLEKVQCQIPVHAEQSQIQHPQGWNTPGFMMVISGIRRASFIFKFMFSHQG
TNMETGHDSAIRISNPVAQGRCSGLLTVTTVLDHIFHKEGNQITLYSCRTVMVQAEVDSFKLALQELAKEYAPSARSLNLGVNSLEHGLYPQLSAVAPGTATAHGGTLVGVTVGEQYQQLWEAASKAERRPKKHAEEQEIDQLDLDHAEQGILVDFHTKKKKTKTQKPQHTSK
QDERLQQRGGKIQQLAALVHTANQGSPTPQGSSMSAPQQPAPSNTSTRGPDDQETSAREASNTWMILQEERSNNNPWQRCALGSLFSDLDASSDDECEEDYGSLIATPGFLVGPHWSRKSPVDETKQPGRKKLPKRLSGHSADTPRSGFSYETTATTSGLQESPDASGQKL
PPALTRENLTAIPESRSWTHSYFIEPPPPPVYPSLDIDKETSQSPQVDPLENHSPQLVLPQHPTQLIGKMEQEQSIMANEGLWAGQLFHHRRTGRPKDFPTSNGTILQFPTSLTGATAPINNQEGWHFGTDGTIAIDGQMPWSDLDLQSRLVHI
>PVIU01020378_Ondatra_zibethicus
PSHRAPEPTELCPQIIRAIQAGVELGDLLDSCLLTLRTQHPEGNSGNFEQRTAGRYLARHRLVFQIIENRTTSNLTEFCLDTGLGSTQQALLALANNHTEHWKVGQFLSFCSLFLPKLVVGERACLEKVQCQITVHAEQGLIQHPQGWNIPGFMMVISGIMRASFTFKFMLTHQG
TNMEAGHDSDISISNSVAQARCPGLLTVKTVLDHIFRKGGNQITLYPLVRTIMVQAEVGSFKLALQGLAKEYAPFAWILNLRVNSLEHGLFPQLSATALGTATAHGGALVGVTVGEQYQQLWEAASEAERRPKKHAEEQEIDQLDLDHAEQGILMGFHTKKTKTKKNPATLDIK
AEMMLAANLGKKKIQQLAASVHTANQGSPTPQGSSMSAPQQPAPSNTSTRAPADQETSAREQSLQHLDDLAAGGKDPIITHGSNVQDQDSSSQILDASSDDEYEEDYGSLIATPDFLVGRHWSCKSPVDETKQPEERSSQNDLVATSHQTPQDQVFPMRQQQQPRDCRK
AQMLQGKHFHLLPGNTSLQSLSPSRVGSNSSFIEPPPPPVYPSSDNNKETSQSPQVDPLEIPLKSPTDAPSITTNLTTLQISETEQEYQSIIAHEGLWAGQLFHHRRTGRPKDFPTSNGAILQFPTSPTGATAPINNQEGWHFGTDGTITMDGQMPWCDLDLQSQLVYI
>JABTXL010001783_Neodon_shergylaensis
PSHRAPEPTELCLQIIRAIQFGVELGDLLDSCLLTLHTQYPEGNSGNFEHRAARRYLERYRLVFRILENKTTSNLTEFCLDTGLGSTQQALLALANNHNEHWKVGQFLSSCSLFLPKLVIGERACVEKVQCQIPVHAEQGLTQHPQGWNTPGCMMVISGIMWASFTFKFMLTHQ
GTNMETGHDSAISISQGRCSGLLTVTTVLDHIFRKEGNQITLYSCRTVMVQAEVDSFKLALQELAKEYAPSARSLNLVNSLEHGCYAHGGTLVGVTVGEQYQQLWEAASEAERRPKKHAEEQEIDQLDLDHAEQGILVDFHKKKNKTKNPKTTTLASRQRRLQQRGGKNPTTCS
FGSHYKSRDPTPQGSPMSAPQQPAPSNTSTRGPDDQETSARESKAYQHLDDLAAGGKDPIITHGSDVQDQDPSSQILDASSDDEYEEDYGSLIATPDFLVGPHWSRKSPVDETKQPEERSSQNDSVDTSQQTPQDQAFPMRQQQPRGCRKAQMLQGKSFHLLPGKTSLQSL
SPGRVGPTSYFIEPPPPPVYPSSDIDKETSNHPKILNTTSSPLVTPNLTTEQISKMEQEYQSIMANEGLWAGQLFHHRRTGRPKDFPTSNGAILQFPTSLTGATAPINNQEGWHFGTDGTIAMDGQMPWSDLDLQSQLVYIALIIH
>LOJG01257017_Ellobius_lutescens
IPAIQAGVELGDLLDSCLMTLHTQHPEGNSGNFEQRAGRHYLERNGLVFHIIENKTTSNLTEFCLDTRLGSTQALLALANNHNVHXXXXXXXXXXXXXXXXXXXWQLKQALLALANNHNVHWKVRQFLSFCSLFLPKLVVGERACLEKVQCQITVHAEQGLIQQHPQGWNTPG
FMISGIMRASFTFKFTLTHQGTNVEAGPDSAISISNWVAQGRCSRLLTVSTVLDYIFSKKEIRSLSTLVWIVMVHAEVDSFKLALQALAKEYAPFARSLNLGVNSLEHGLFPQLSAVASGTATAHGGTLAGTVTHGVTVGEQYQQLWEAASEAERRPKKYAEEQEIDQLDLDHAE
QGILVDFHTKKKK
>LOJH01047383_Ellobius_talpinus
QHSIFNFRTQPVESLRAPEPTELCPQIIRAIQAGVELGDLLDSCLLTLRTQHPEGNSGNFEQRAGRCYLERHGLVFHIIENKTTSNLTEFCLDTGFGSTQQALLAPANNHNEHWKVGQFLCFPKLVVGERACLEKVQCQHAEQGLIQHPQGWNTPGFMISGIMRASFTFKFMLT
HQGTNVEAGHDSAISISNSVAQGRCSGLLTVSTVLEHIFCKEGNQITLYSSRIVMVQAEVDSFKLALQALAKEYAPFARSLNLGVNSLEHGLFPQLSAVASGTATAHGGTLVGVTVGEQYQQLWEAASEAERRPKHAEEQEMDQLDLDHAEQGIPVDFHTKKKNHDIGQSRQRR
LQQSGGKIQQPAASVHNANQGSPTPQGSSMSAPQQPTPSNTSTRDPNDQETSAREQGLQHRMILQEERSNNNPWQCARPSSQISNASSEDEYEEDYGSLIATPDFLVGPHWSRKSPVDETKQPEESSQNDSVDNNQQTPQDQAFHMRQQQQPQGCRKAQTPLGQKLP
PALTRGNLTAIPDPGTWTHSYFIEPPPPPVYPSSDNDKETSQSPPSNPLEIPLKSPTGAPSITPNLTTEQISKTELECQNVIAHEGLWAGQLFHQRRTGRPKDFPTLNGFYQFPTSLTGATAP
>LIPI01007319_Myodes_glareolus
TQEDLELHSILTAGLNQWDPSHRAPEPTELCPQIIRAIQAGVELGDLLDSCLLTLHTQHPEGNSGNFEKRATRHYLERHGLVFHIIENNTTSNLTEFCLDTGLGSTQALLALANNHNEHWKVGQFLSFCSLFLPKLAVGERACLEKVQCQITIHANRGLIHPQGWNTPGFMMVISG
IMRVSFTFKFMLTHQGANVEAGHDSAISISKSVAQARCSGLLTVTTVLDHIFRKKDIRSLSTLVWTVMVQAEVDSFKLALQGLAKEYAPFARSLNLGVNSLKHGLSPQSAVAQGTVTAHGGTLVGATVGEQYQKSREAASEAERRPKKHAEEQEIDQLDLDHAEQGILVDFHTH
THTQKKTHNIGHQGRDDTCSKRGGKIQQLAASVHTANQGSPYTQGSSMSAPQQPAPSNTSTSPDDQETSAREQGLQHWMILQEVKIQPHGSDIQDQDSSSQILDASSDDEYEEVYSSLVATPDFLVGPHWSCKSPVDETKQPEERSSPSDSVDSSQQTPQGQAFPMRKQQ
QPRGCRKAQAFQGESSHLLSQGKPHCNPERGRSGTHSYFTEPPPPPVYPPSDNDKEHLNHPKILKYHSPQPVLPQHPIQPEQISRTEQEYQSTIAHEGLWAGQLFHHRLTGGPKGFPTANGAILQFPTSLDEATAPINNEEWHFGTDGTIAMDGQMPWCDLDLQSRLVYI
>MULK01003177_Myodes_glareolus
TQEDLELHSILTAGLNQWDPSHRAPEPTELCPQIIRAIQAGVELGDLLDSCLLTLHTQHPEGNSGNFEKRATRHYLERHGLVFHIIENNTTSNLTEFCLDTGLGSTQALLALANNHNEHWKVGQFLSFCSLFLPKLAVGERACLEKVQCQITIHANRGLIHPQGWNTPGFMMVISG
IMRVSFTFKFMLTHQGANVEAGHDSAISISKSVAQARCSGLLTVTTVLDHIFRKKDIRSLSTLVWTVMVQAEVDSFKLALRGLAKEYAPFARSLNLGVNSLKHGLSPQSAVAQGTATAHGGTLVGATVGEQYQKSREAASEAERRPKKHAEEQEIDQLDLDHAEQGILVDFHTHT
HTHRKKPTTLDIKAEMTPAAKWGENPTTCSFGSHCKSREPNTQGSSMSAPQQPAPSNTSTSPDDQETSAREQGLQHWMILQEVKIQPHGSDIQDQDSSSQILDASSDDEYEEVYSSLVATPDFLVGPHWSCKSPVDETKQPEERSSPSDSVDSSQQTPQGQAFPMRKQQQ
PRGCRKAQAFQGESSHLLSQGKPHCNPERGQSGTHSYFTEPPPPPVYPPSDNDKEHLNHPKVLKYHSPQPVLPQHPIQPEQISRTEQEYQSTIAHEGLWAGQLFHHRLTGGPKGFPTANGAILQFPTSLDEATAPINNEEWHFGTDGTIAMDGQMPWCDLDLQSRLVYI
>JAQHUG010000111_Dicrostonyx_torquatus
FGGRSHSHSPTHKGFEQHSILNFGTQPVESFTPRPEPTELCPQTICAIAAVDLGDLLDSCLLPLRTPHPYEGNLGILSKGQPVHHRARHGLVFQIMENKTSSNLTEFCLDTGLGSIQQALLALANNHNGHWKVGQFLSFCSLFLFLPKLVVGERACLEKIQCQITVHAEQGLIQHPQ
GWKTPGFMMVISGIMRASFTFKFMLTHQGANVEAGHDSDITISNLVAQARCSGLLTVKTVLDHIFHKEGNQTTLHSCLDNNGSGRGRFQAHTAGLAKNGVPLARSLNLGVNSLEHGLFPQLSAIASGTATARGGTLVGVSVGEQYQQLWEAASEQAERRPKKHAEEQEIDQ
LDLDHAEQGILMGFHTKKKNKKQKTKKPTTLDIKAEMMLAAKWGEKSNNLQLWFTLQIKGAQHLKGSSMSAPQPASSNTPTRGPDDQEASAREQGLQHLDDLAAGGKEPMITHGSNVQDQDSSSQTLAASSDAEYEEEYDSLNATPDFLAGPHWSRKSPVDETKQPEERS
SQNDSVDTSQQTPQDQASPETTAATSGLESPDASGQKLPPALTRENLDCNPAQAELDPGKGGSFIEPPPPLVYPSSDNDKETSQSPQVDPLGNTTKSHSQPEQISETEQEYQSIIALEGLWVGQLFHYRTGRPKDFPTLNEAILQFPTSLTGATASINNQEGWHFGTDGTITMD
GQMPWCDPELQSRL
>APMK01149764_Cricetulus_griseus
DFNLHGILNLGAQPVPFVQSRKEGLYILCSTLSGLWAQIICVIEAGVELGDALDSCLLTVCVHHAYDGSAEAFMNSAAYGYLERNCTDCWLLSNSVSNIFLSFCSLFLPKLVVGEAACLEKVQCQMVIHEEQGLIYLQGQTLGFRRIFFLRLMTASFTYKFLPIHQGINMKAGHDAT
NITITNSVAQSCFSRLLIVKAVLEHILCKEETRGVVLHPLVKLVRVREVENFKVAMKGLTQHKEYTPFSRILHLAGLNNLEHGLFPSLSITALGDAIAHSCTLASVTMGEQYQRREAATEAEWRLQEHAEEKEINRLGLDKVETKILIDFHRKHHEISNQQTKRRSWIKEKKARAARCC
QAKACCQSIQSSGMPFQWPQNQAPSCQDIPTTRSSDPPSRDQSSHMVDSDQHAPTDSPQGGDPLLLFHSNSHREEYRPPNGHSTQVRSCTFRFVRQPACSSWNSRKPTRSGGRECRDRLERNVMYSDSLTNVPKILEELDVNIQSSPSHPQTSLLPIIHETSEEEKSTSSHHEF
PPPPIYPGDGVKTNDCHYQTQQPPSGNNNRGSSQYDALSQTISLPKEQIQEVERVFDRVRKEGDLHSGLVFFHNLTSMGKTFTTKTGAILPYPDCLEQLLPGWNFGEERHVYFKGTEYTWAVLATCIKF
>CALSGD010000248_Phodopus_roborovskii
DFNLHGILNLGAKPVPFVRSLKVFTKYAPRSSVLWAQIICATETGVELGDTLDSCLLTVCVHHAYDESAEAFMNSPAYGYLEKQHLLASEQQCQQQFDEFIPKMGSGGRHTHHALSTLNINTSGHQNVGPFLSFCSLFLPTLVLGEAACREKVQCQFVIHGEQGLIQYPQGWQTL
GFRRIVFLMRASFTYKFFLIYQGINMKVGHDSNDTTITNSVAQSSFSRLFIVKTVLEHTFHKKETVIPYPLVKLVPVRSEVEHFKVALKGFTQQKEYTPFARILNSAGVNNLEHGFFPLLSIIMLGDVTNHGCTLASVTVGEQYQQLREATEAKQRLQEHPEEQDINRLGLDKVETKILI
DFHRKHHEIGNQQTDTERERLEQLDAAITVASQRHGYQGIRSLGMPLQWPQKNQGPSCQDIPTTRSSDCQSGPVKPHDDSDQHATTDCPQDGDLLLLFEANSDREEDRPQKGTACRQELVKDSDLLDNQSVPSWNSRKPKKKWRTKDADDESHFENEMSCTLTSSQMSKI
LEE
>MCBN011458509_Phodopus_sungorus
DFNWHGILKLGAQPVQPFFRSLKGYTFYAPPSYVLLAQIICAIEAGVELGDTLDSCLLTVCVHHAYDESAEAFMNSPTLYLEKHFLASDQQCQQQFDEFIPELGPGDRHTCHALSTLNINTSGHQNVGPFLSFCSLFLPTLVVGEAACEKVQCQFVIHGEQGLIQYPQGWQTLGFR
RIVFLMRASFTYKFLLIYQGINMKAGHGSNDITITNSVAQSHFSRLFIVKTVLEHIFHKKETVIPHPLVKLVPVKSEVEHFKVALKGFAQQKEYTPFARILNSAGINNLEHGIFPLLSIIVLGDATAYGCTLAITVGEQYQQLRAEQRLQEHTEEQDINRLGLDNVETKILIDFHGKHHEIG
NQQTDAKRERLEQIDASVTVASQRHGQGIRSLGMPLQWPQNQGPSSHQDIPTTRSSDQSGPVKYMDDSDQHAPTDCPQDGELLLLFEANSDREEDRHRHTGNGHSTARSCRFRFVRQQSVLHDEILESPKEMEDEDADDESHFENEMSCTLTPSHMSKILEELDVNSQSSP
SYLQTSLFPTIHETSGEEKNTSSHYELPLPQFILEMEKPQLSLPGPAVPSSSNNNGSSQCDALSQTISLTKEQIQKVERVFDRQVCEGRPYSVLGFFHNLTGMSKNFTTNAGTILPHPDCLEQLLLSALQAGILVREGTSVSRDGVQLGCAHLHIVFVWILYDSTTLSSSFLHILRNN
>APMT01070558_Mesocricetus_auratus
FNLHGILNLGSQPVPFVSQKVYTFYVPHSFGLWAQIICVIEAGVELGDTLDSCLLTLCVHHAYDESAEAFMKCPAYGYLQRNRIRLLVSEQQCQQQFDVYSEIGSGRQTHRCLSTLSINTNGRQSVGPFLSFCNLFLPKLVVGKACLEKVWCIMIHREQGLIQYPTGLADLGFRRIIF
NLMRDSFTYKFLLTHQGINMKAGHDASNITITNSVAQSHFSRLLIVKTVLEHILCKEETWVVLHPLVKLVRVREVENFKIVQLCLVLTGDLISLASCLYGATSSHLGKNAQIAIMKISIPTTKKGIPGTCQIGPLPDAQDCNHQDFVLRYARGKTHLKWTEEYKQVFELERTPTLTFLTS
SNSSIFISMEERALLRVSLPKLWVPDKCQE
>CAJEUE010003860_Arvicola_amphibius
IRGCAHRFTPSKAAIGLGRSSESYLLTLCIQHAYESHSKAFETSAARRYLEAQGLVFKALDRANSSYLNEFLPHTHTGRGHGLRLTLLTLPQQPRANAGRSASSSLFCHLFLPNLVVGEKARLEKVQRQLAIHADLLQFPQNWQAPGFMVIFRIMRVSFTFKFLLIRQMNMDAGHN
ADDIIITNNFSSSGFLIVKTVLDNILWREGDQITLHPLVCIVRVHTEVENFKLALRSLAHHGAYSFALLNLSGVNNLEYDLFPQLPAIALGVATAHRGTLAGVAIGCRVYQQLREAAREAECLQDHLESRKIGRLGLGQMKAEILLDFHKKKHEMGASKSKSSLRPNFSEKTSATGSCN
SEQLSFASPIQLCPHAPPPSAPPSHQMEQPMPPGHKSISPKHPESDTKDPDEIPNLEGSEED
>CATLKP010000862_Chionomys_nivalis
LCSQIMYAIKAAIGLGRSAESCLSTLCFQHACESHSKAFVNSAARWYLEAQGLVLKALDRANRGNLNEFLPHTHTGCQHGLRLTLLTLLNNQSKCRQECQFLSFCHLFLPKLVVGEKARLEKVQRQLAIHAGPTQFPQNQAPAFMLIFRIMRAGFTFKFLLIRQRINMDTGHNVD
DIIIAKVLVAQARFSGLLIVKTVLERILRREGDQITLHSLVLIVRVHEAENFKLVLKGLACHGAYSFAWILTLSGVKNLEYGLFPQLPAIALRAATAHRGTLAGVAIGRVYQQLQEAAGGAECLQDHLEKQEIGHVGLGQVRAEILMDFHKKKHEMGVSKSKSSLPNLSKKLQPPAVIN
NLARSASPIQLRHRALPSHPIIPTDGAASAPGHKSISPKHPESDTKDPDEIPNLEGSEEDCSNFQILDAYPEGGGG
>CADCXR010023786_Microtus_agrestis

LCSQIMYAIKAAIGLGRSSESCLTLCIQHACESHSKAFETSAARQYLEAQGLVLKALDRANRGNLNEFLPCTHTGCRHGQRLTLLTLLNSQSERRQECQFLSFCHLFLPKLIVGEKARLEKVQWQLAIHAGPTPFPQNWQAPAFMLIFRIMRAGFTFKFLLIRQRINMDAGHNADN
TIITKILVAQAHFSGFLILKTVLDHILRREGDQITLHPLVLIVRVHSGEFQACSKGSLARHGAYSFAWILTLSGVKNLEYGLFPPAIALRIATAHRGTLAGVAIGRVYQQLQEVAGGVEYLQDRQIDRVGLGRRKAEILMDFHKKKHEMGVQQVEIILATKLERETSATCACNSELSSLR
SHSAVLPALPSRPIIPTDGAASAPGHKSISPKHPESDTEDHDEIPNLEGSEEDCSNFQILDAYPAGGRG
>VIIT010006645_Microtus_arvalis
RQTTVYIIKKIFKKKTYAIKVAIGLGRSSESCLSTLCIQHACESHSKAFETSAARRYLEAQGLVLKALDRANRDNLNEFLPHTHTGCRHGLRLTLLTLLNNQSEHWQECQFLSFCHLFLPKLVVGEKTRLEKVQRQLAIHTGPTQFPQNWQAPAFMLIFRIMRAGFTFKFLLIRQRINM
DAGHNMDDTIITKILVAQAHFSGFLIVKTVLDHILRREGDQITLHPLVLIVRVHSGEFQACSKGSLARHGAYSFAWILTVSGVKNLEYGLFPQLPAIALRVATAHRSTLAGVAIAEEYYQQLQEAAGGVEYLQTRKIDRVGLGQRKAEILMDFHEKKHEMGVQQVEIILATKLERETS
ATYAYNSELSSLCSHSAVPPALPSRPIIPTDGAARAPGHKLISPKHPESNTEDRDEIPNLEGSEEDCSNFQILDAFPAGGRG
>NMRL01000090_Microtus_fortis
LCSQIMYAIKAATGLGRSSESCLSTLCIQHTCESHSKAFETSEAQGLVLKALDRANRDNLNEFLPRTHTGCRHGLRLTLLTLLNNQSKHGQECQFLSFCHLFLPKLVVGEKARLEKVQRQLAIHARPTQFPQNWQAPAFMLIFRIMGAGFTFKFLLIRQRINMDAGHNADDTIITKIL
VAQARFSGFLIVKTVLDHILWREGDQITLHPLVLIVRVHSREFQACSKGSLARHGAYSFAWILTVSGVKNLEYGLFPQLPAIALRVATAHRGTLAGVAIGRVYQQLQEAAGGVEYLQDHQIDRVGLGRMKAKILMDFHKKKHEMGVQQVEIILATKLERETSATGACNSELSSLRS
HSAVPPAPPSHPIIPTDGAASAPGHKSISPKHPESDTEDCDEISNLEGSEGDCSNFQILDAYPEGGGGPNQHLPDHPFGHWHKCEVSRIVAHKNVNTQNVMCCKPQAQVEKPRMILATSGGLVPLLDQLLNILNINPQNRQPRNINTYRTKFNSPFQGTPSKSLSSGSKQFSIVH
LWDCASPPPTHTHTHVCVHTQSPDVPVLCRLCHCVIPACSQGQAVALERQYLVSMWSQGQYKANSHCCIVRFCGERPMAQGAQDARNCGWLPD
>AHZW01010557_Microtus_ochrogaster
CSQIMYAIKAAIGLRRSSESCLSTLCIQHTCESHSKAFETSTARRYLEAQGLVLKALERANSGNLNEFLPHIHTGCRHGLRLTLLTLPQQPESAVILSFCHLFSPKLVVGEKAHLEKVQRQLAIHTGPTQFPQNWQAPAFMLIFRIMRAGFTLKFLLIRQRINMDAGHNADDIIITKILV
AQARFSGFLIVRTVLDHILRRKGDQITLHPLVLIVRVHSGEFQACSKGSLARHGAYSFAWILTLSGVKNLEYGLFPQLPAITLRVATAHRGTLAGVATGRVYQQLQEAAGGAECLQDHLEKQEIGRVGLGQMKAETLMDFHKKKHEMGVQQVEIILVTKLERETSATCSCNSEQLS
SLRFSHSAAPPALPSHPIIPTDGAASAPGHKSISPKHPESDTEDRDKIRNLEGSEEDCSNFQILDTYSEGGRG
>JAGDQN020000165_Microtus_richardsoni
LCSQIMYAIKAAIGLGRSSESCLSTLCIQHACKSHSKAFETSTARRYLEAQGLVLKALDRANRGNLNEFLPHIHTGCRHGLRLTLLTLFNNQSEHQQECQFLSFCHLFLPKLVVGEKARLEKVQRQLAIHTGPTQFPQNWQAPAFMLIFRIMRAGFTLKFLLICQRINMDAGHNADD
IIITKILVAQARFSGFLIVNDLSIIILRRKGDQITLHPLFLIVRVHSGEFQACSKGSLARHGAYSFAWILTLSGVKNLEYGLFPQLPAIALRVATAHRGTLAGVATGRVYQQLQEAAGGAECLQDHLEKQEIGRMGLGQMKAEILMDFHKKKHEMGVQQVEIILATKLDRETSATGGC
NSEQLSSLRFSHSAAPPALPSHPIIPTDGAGSAPGHKSISPKHPESDTEDCDEIPNLEGSEEDCSNFQILDAYPEGGRG
>CADCXS010046358_Microtus_agrestis
LCSQIMYAIKAAIGLGRSSESCLTLCIQHACESHSKAFETSAARQYLEAQGLVLKALDRANRGNLNEFLPCTHTGCRHGQRLTLLTLLNSQSERRQECQFLSFCHLFLPKLIVGEKARLEKVQWQLAIHAGPTPFPQNWQAPAFMLIFRIMRAGFTFKFLLIRQRINMDAGHNADN
TIITKILVAQAHFSGFLILKTVLDHILRREGDQITLHPLVLIVRVHSGEFQACSKGSLARHGAYSFAWILTVSGVKNLEYGLFPPAIALRIATAHRGTLAGVAIGRVYQQLQDVAGGVEYLQDRQIDRVGLGRRKAEILMDFHKKKHEMGVSKSKSSLRPNLSEKLQPPVPVIQNLARS
ADPIQLCYLLFHPALSSQQTEQPVHQGINQYPPSIQSQYTEDHDEIPNLEGSEEDCSNFQILDAYPAGGRGPNQHLPDHPFGHWHKFEVSRIVTQKKMLTLRTKCVVKKHKYKWKCHTCLPHGGLVPLLDQLLNILNINPQNRQSRNINTYRTKFNSPFQGTPSKSLNSGSKQFS
IVPSLRLCLPTHKHTHVCVHTQSPDVPVLCHLCHCVIPACSQGQAMALEQQYLVSMWSQGQYKANSHCCIVRFCGERPTAQGAQDARNRGWLPDVRMRYLENEATCPMQMFASPPWLSR
>CADCXT010004588_Microtus_agrestis
LCSQIMYAIKAAIGLGRSSESCLTLCIQHACESHSKAFETSAARQYLEAQGLVLKALDRANRGNLNEFLPCTHTGCRHGQRLTLLTLLNSQSERRQECQFLSFCHLFLPKLIVGEKARLEKVQWQLAIHAGPTPFPQNWQAPAFMLIFRIMRAGFTFKFLLIRQRINMDAGHNADN
TIITKILVAQAHFSGFLILKTVLDHILRREGDQITLHPLVLIVRVHSGEFQACSKGSLARHGAYSFAWILTVSGVKNLEYGLFPPAIALRIATAHRGTLAGVAIAEEYQQQLQEVAGGVEYLQDRQIDRVGLGRRKAEILMDFHKKKHEMGVSKSKSSLRPNLSEKLQPPVPVIQNLAR
SADPIQLCYLLFHPALSSQQTEQPVHQGINQYPPSIQSQYTEDHDEIPNLEGSEEDCSNFQILDAYPAGGRGPNQHLPDHPFGHWHKFEVSRIVTQKKMLTLRTKCVVKKHKYKWKCHTCLPHGGLVPLLDQLLNILNINPQNRQSRNINTYRTKFNSPFQGTPSKSLNSGSKQF
SIVPSLRLCLPTHKHTHVCVHTQSPDVPVLCHLCHCVIPACSQGQAMALEQQYLVSMWSQGQYKANSHCCIVRFCGERPTAQGAQDARNRGWLPDVRMRYLENEATCPMQMFASPPWLSR
>VIIU01007453_Microtus_oeconomus
LCSQIMYAIKAAIGIRELLVALCIQHACESHSKAFETSAARRYLEAQGLVLKALDRANRGNLNEFLPRTHTGCRHGLRLTLLTLLNNQSEHGQECQFLSFCHLFFLPKLVVGEKARLEKVQRQLAIHAGPTQFPQNWQAPAFMLIFRIMGAGFTFKFLLIRQRINMDAGHNVDDTII
TKILVAQARFSGFLVVFFFFLFFLFFFSFSFFETGFLCGFGACPGSGFLVVKTVLDHILRREGDQITLHPLVLIVRVHSGEFQACSKGSLARHGAYSFAWILTISGVKNLEYGLFPQLPAIALRVATAHRGTLAGVAIGRVYQQLQEAAGGVEYLQDHQIDRVGLGRMKAEILMDFHKK
KHEMGVSKSKSSLRPNLSEKLQPRVPVIQNLARSANPIQLCHLLFHPTLSSQQTEQPVHQDINQYPPSIQSQYTEDRDEIPNLEGSEEDCSNFQILDAYPEGGGGPNQHLPDHPFGHWHKFEVSRIVTQKKCHSELSNVLLKKTQAQVEKPRMILATSGGLVPLLDQLLNILNINP
QNRQSRNINTYRTKFNSPFQGTPSKSLSSGSKQFSIVHLDCAFPPHTHTHVCVHTQSPDVPVLCHLCHCVIPACSQGQAVALERQYLVSMWSQGQYKANSHCCIVRFCGERPMAQGAQDARNCGWLPDVRVRYLENGATCPMQMFAFLPWLSR
>JAEPQX010011494_Microtus_montanus
TLLNNQSEHRQECQFLSFCHFFFLPKLVVGEKARLEKVQRQLVIHTGPTQFPQNWQAPAFMLIFRIMRAGFTLKFLLIRQRINMDAGHNADDIIITKILVAQARFSGFLIVKTVLDHILRRKGDQITLHPLVLIVRVHSGEFQACSKGSLARQGAYSFAWILTLSGVKNLEYGLFPQLP
AIALRVATAHRGTLAGVNCRVYQQLQEAAGGAECLQDHLEKQEIGRVGLGQMKAEILMDFHKKKHEMGVSKSKSSLRPNLSEKLQPPAAVIQNNLARSASPIQLRHLLFHPTLSSQQTEQAVHQGINQYPPSIQNQYTEDRDEIPNLEGSEEDCSNFQILDAYPEGGRGPNQH
LPDHPPLATGISLKSLESHK
>JAQJZF010000015_Microtus_californicus
ILNCGTQPVESFIRSRKIYSSDGGVCSQIIYTIKAAIGLGRSSESCLSTLCIQHACESHSKAFETSTARRYLEAQGLVLKALDRANRGNLNEFLPHIHTGCQHGLRLTLLTLLNNQSEHRQECQFLSFFFFFYLFIYLLKISASSPPPPPISLPLPRSSLPPTLFLPKLVVGEKARLEKVQRQLAI
HTGPTQFPQNWQAPAFMLIFRIMRAGFTLKFLLIRQRINMDAGHNADDIIITKILVAQARFSGFLIVKTVLDHILQREGDQITLHPLVLIVRVHSGEFQACSKGSLARHGAYSFAWILTLSGVKNLEYGLFPQLPAIALRVATAHRGTLAGVATGRVYQQLQEAAGGAECLDHLEK
QEIGRVGLGQMKVEILMDFHKKKHEMGVSKSKSSLRPNLSEKLQPPASVIQNNLACSASPIQLRPLLFHPTLSSQQTEQPVHQGINQYPPSIQSQYTEDRDEIPNLEGSEEDCSNFILDAYPEGGRGP
>JAGKIF010000458_Microtus_oregoni
LCSQIMCAIKAAIGLGRSSESCLSTLCVQHACESHSKAFETSTARRYLEAQGLVLKALDRANRGNLNEFLPHIHTGCRIGLRLTLLTLLNNQSEHRQECQFLPFCHLFLPKLVVGEKARLEKVQRQLAIHTGPTQFPQNWQAPAFMLIFRIMRARFTLKFLLIHQRINMDAGHNADD
IIITKILVAQARFSGFLIVKTVLDHILQRKGDQITLHPLVLIVRVQSGEFQACSKGSLARHGAYSFAWILTLSGVENLEYGLFPQLPAITLRVATAHRGTLAGVATAKEQYQQLQEAAGGAECLQDHLEKQEIGRVGLGQMKAEILMDFHKKKHEMGVSKSKSSLRPNLSEKLQPPA
AVIQNNLARSASPIQLRHLLFHPTLSSQQTEQPVHQGINQYPPSIQNQYTEDRDEIPNLEGSEEDCSNFQILDAYPEGGRGP
>JAQJZG010000015_Microtus_californicus
ILNCGTQPVESFIRSRKIYSSDGGVCSQIIYAIKAAIGLGRSSESCLSTLCIQHTCESHSKAFETSTVRQYLEAQGLVLKALDRANRGNLNEFLPHIHTGCQHGLRLTLLTLLNNQSEHRQECQFLSFFFFLSFFLSLFIYLFIYLFIKDFCLLPTTASHFPPPPPIKSPSAHLFLPKLVVGEKARL
EKVQRQLAIHTGPTQFPQNWQAPAFMLIFRIMRAGFTLKFLLISQRINMDAGHNADDIIITKILVAQARFSGFLIVKTVLDHILQREGDQITLHPLVLIVRVHSGEFQACSKGSLARHGAYSFAWILTLSGVKNLEYGLFPQLPAIALRVATAHRGTLAGVATGRVYQQLQEAAGGA
ECLDHLEKQEIGRVGLGQMKVEILMDFHKKKHEMGVSKSKSSLRPNLSEKLQPPASVIQNNLACSASPIQLRHLLFHPTLSSQQTEQPVHQGINQYPPSIQSQYTEDRDEIPNLEGSEEDCSNSLNTCLSRGWQRTQPTPAPPLWPLAVSLNSHTHKKKMLTLRTKCVVKKRKHK
WKSHTYLPLVALSLLLDQCPWCTSCIFILTLKKDSQETIHITNRTKFNSPFQGTPSKSLSSGSRQFSIVRLDCAFPPPTHTHTGVCAHTVQPRCSSALPFVSLCHPSLQSRPKWQCLVSMWSQGQYKANSHCSILRFCGERPMAQGAQDARKRGWLPDVRVRYLENGTTCPMQ
MFASPPWLSR
>LOJG01050294_Ellobius_lutescens
LCSQIIYAIKAAIGLGRSSESCLLTLCIQHAYESHSKAFETSAARRYLEAQGLVLKALDRANSGNLNEFLPHTHTGHGHGLRLTLLTFLNNESERRQEGQFLSFHLFLPKLVVSEKVHLEKVQRQLAIHTGPTQFPQNWQAPGFMLIFRIMRASCFTFKFLLIHQRINMDAGHNADDII
ITKILVAQAHFSGFLIVKTVLDHILGREGDQLTLHPLVRIVRVHNEVENFKLALKGLARHGAYSFAWILTVSGVNNLEHDLLPQLPAIALGVTTAHRSTLAGVAIGRVYQQLREAASKAECLQDHLEKQEIGCLGLGQMKAEFLMDFHKNKHEMGVVSKSKSSLRPNFNEKLQPLA
TAIQNNLAHPASPIQLCPPAPPTSAPPSHPIIPTDGAASVPGHKSISPKHPESDTEDPDEILNLEGSEEDCSNFQI
>LOJH01197867_Ellobius_talpinus
LCSQIIYAIKAAIGLGRSSESCLLTLCIQHAYESHSKAFETSAAKRYFEAQGLVLKALDRANSGNLNEFLPHTHTGHGHGLRILLLTLLNNQSERRQEGQFLSFCHLFLPKLVVRRHCLEKVQRQLAIHTGPTQFPQNWQAPGFMLIFRIMRASCFTFKFLLIHQRINMDAGHNADD
MIITKILVAQAHFSGFLIVKTVLDHILWRGDQLTLHPLVRIVRVHTEVENFKLALKGLACHGAYSFAWVVTLSGVNNLEYGLFPQLPSIALGVATAHRGTLAGVAIGRVYQQLPEAASKAERCLQDHLEKQEIGRLELGQMKAEILMDFHKKKHEMGVSKSKSSLRPNFSEKLQPL
AAAIQNNLARPASPIQLCPPAPPTSAPPSHPIIPTDGAASVPGHKSISPKHPESDTEDPDEIPNLEGSEEDCSNFQILGGGGPNQHLPGPPLSTGISLKSLESQKKVNTQNVMCCLKTQAQVEKPTVTLATSGGLVPSARSVPMSHQLLNILNINPQKNXXXXXXXXXXXXXXXXX
XXXXXXXXXXXXXXXASKSLSSG
>CADCXP010001697_Myodes_glareolus
MAWNRLVSNYGTQPVESFIRSRTIYSFDVGLRFISGVCSQIIYAIKAATGLGRSSESSLSTLGIQHTYQSHSKACETSAARRYLEAWGLVPKALDGANSGNLNEFLPHTHTGRGHGLRLTLLTLLNNQSERRQECQFLSFCHLFLPKLVVGEKARLEKVQQQQLAIYAEPTQFPQNQ
TPGFMLIFTIMRASFTFKFLLIRQRINMDAGHNADIIITKILVAQAHFSGFLIVKTVLDHILWREGYQITLRPLVCTVKVHTEVENFKLALEGLVHHGAYSFCLMDTSSRVNILEYGPFPQLPSIALGVETSHRGTLAGVAIAKELYQQLWGGRAAIIEAECLQDHLEKQEMGRLGLGQ
MKAEILMDFHKKKHEMGVSKSKSSLRPNFSEKLQTLAAAIQNNQARSPSRYSAVPTCSAPAPLHPTLSSQQMEQPEHQGVNQYPPNIQSQYAEDPDEIPNLEGAEEDGSNFQILDAYPEGGGGP
>LIPI01337567_Myodes_glareolus
YQSHSKACETSAARRYLEAWGLVPKALDGANSGNLNEFLPHTHTGSGHGLRLTLLTLLNNQSERRQECQFLSFCHLFLPKLVVGEKARLEKVQQQQLAIYAEPTQFPQNQTPGFMLIFTIMRASFTFKFLLIRQRINMDAGHNADIIITKILVAQAHFSGFLIVKTVLDHILWREGY
QITLRPLVCTVKV
>CADCXQ010001915_Myodes_glareolus
MAWNRLVSNYGTQPVESFIRSRTIYSFDVGLRFISGVCSQIIYAIKAATGLWRSSESCLPTLGIQHAYQSHSKACETSAARRYLEAWGLVPQALDGANSGNLNEFLPHTHTGSGHGLRLTLLTLLNNQSERRQECQFLSFCHLFLPKLVVGEKARLEKVQQQLAAYAEPTQFPQNQ
TPGFMLIFTIMRASFTFKFLLIRQRINMDAGHNADIIITKILVAQAHFSGFLIVKTVLDHILWREGYQITLRPLVCTVKVHTEVENFRLALEGLVHPGAYSFAWIPHLGVNILEYGPFPQLPAIALGVETSHRGTLAGVAIAKELYQQLRERAAIIEAECLQDHLEKQEMGRLGLGQMK
AEILMDFHKKKHEMGVSKSKSSLRPNFSEKLQTLAAAIQNNQARSPSRYSAVPTCSAPAPLHPTLSSQQMEQPEHQGVNQYPPNIQSPAAEDPDEIPNLEGAEEDGSNFQILDAYPEGGGGGPNQHTCLTTPLATGISLKSLESHTHTKKKNANTQNVVCCKNTSISGKARVILAT
SGSLVPSARSVPVSHQLLNILNINPQNRQSKNHKDGTITNRTKFNSPFQGTASKSISSGSRQLSTAHLDCASLAPPPPDVCVHTQCSPDVPVLSHLCHCVLPTCNQGQAVALEMAVSGLHVESRSIQSKFSLLYCGERPTAQGAQDTRNCWGWRPDVHVRYLEHGATCPMQM
PTSPPWLSR
>JABTXL010007601_Neodon_shergylaensis
LCSQIMYAIKAVIGLGRSSESCLSTLCIQHACESHSKALETSAARRYLEAQGLVLKALDRANRGNLNEFLPRTHTGCRHGLRLTLLTLLNNQSEHWQECQFLSFCHLFLPKLVVGEKARLEKVQQQLAIHAGPTQFPQNQAPAFMLIFRIMRAGFTFKFLLIRQRINMDAGHNTDD
TIITKILVARARFSGFLIVKTVLDHILQREGDQITLHPLVLIVRVHSGEFQACAKGSLAHHGAYSFAILTVSGVKNLEYGLFPQLPAIALRVATAHRSTLAGVAIGRVYQQLQEAAGVVEYLQDHQIGLVGLGPMKAKILMDFHKKKHEMGVSKSKSSLRPNLSEKLSHRRLFRTTLAP
HLPFSATCSSIPPYHPNRRSSQCTRATNIPQASRVRYRGDEIPNLEGSEEDCSTFQILDAYPEGGRGP
>PVIU01004168_Ondatra_zibethicus
NVDSPGILNYGTQPVESFIRSRKIYLFDVGLRFIRGVCSQIICAIKAATGLWRSSESCLLTLCIQHAYESHSKAFKTSAARRYLEAQGLVLQALDRANCSNLNEFPHTHTHTGPGHGLRLTLLTLLNNQSDRANAGQEHQFLSFCHLFLPKLVAGCLEKVQQQLAIHAGPTQFPQNW
QTPGLLLIFRIMRASFTFNCLLIRQRINMDAGHTVDDIIIIAQARFSGFLVVKTVLDHILQRGGDQTTLHPLVRIVRVHTEVENFKLALKGLARHGVYPFAWILTLSVKSIINMLPAIALGVATAHRGPCLGRSDQQLWEAASEAECLQDHLEKQEIGRLGLGQMKAEILMDFHKEK
HEMGLGKSKSPLQPNFSEKLQPLVAAIQNNLAHSASPIQLSPPAPPPSAPPSHPIIPTDGAASAPGRGSISPKHPESDSEDPDEIPNLDGSDCSNFQILDAYPESGGGPNQHPPDHPFGHRHVSLNSHKKIKCHSELSNMLLKNTSTSKPKRDIS
>JAQHUG010000091_Dicrostonyx_torquatus
YLKLWHTTCVIHPEPRDSLISCWAQIIYQRLCSQVIYAITAAIGLGRSERCLLTPCIQHAENHSKAFETSAARQYLEAQELVLEALDRANSSNLNEFLPHTHTGPGLTLLNHQSKRRKECQFLSFCHLFLPKLVVGEKACLEKVRQLAIHAGPIQCPQNWQTPGFTLIFRIMRASFTFKF
LLIHQRINMDAGHNADDIIITKILVFQAHSSGFLIVKTVLDHILRRERDQITLHPVLHVVRVHTKVENFKLALKGLAHHGAFLCLDTNLICVNNLEYGLFPQLPAIALGVATAHGTLAGVTIAEEYNQQLREAECLQDHLEKPEISRLGLSQMRAEILMDFHRKKHEMGVQQVEIILAT
KLQPLVAAIQNNLAHSASPLPLSPPAPPPSAPQSHPILPRDGAASAPRHGSISPKHPESDSEDPDEIPNLEGSEEDCSNFQIFD
>APMK01293044_Cricetulus_griseus
VCLFGSPESSELCSQKIYTLEAGVEFGDSLDCCVTLCCIHHAYEGNQEAFRGSPECEYLERNGIPCEILNHSTSNNLDSSLIPLLVGGQSMLSRLPKMPMSTGKENHSSHGQVGAFLSFCLFLPKLEVGAACLEKVQRQIVIHREQSWITSGFMIIIFRLMRASFTFKFMIHQVINLTV
KHDPNDNSVTKARFSGLLIVKAVLDHILHKRDRVTLHPLVRTIRVRSEVESFKPALEGLAGHYVPFAQILNLSGVNNLELGLYPQLSVIALGVATAHGCTLVGITVGEQYQQLREAATEAEKRLQEYAEEPEINLELDKTETKILAEFHKKKQRLGYQEAEEILSAKQEKLEQLSGAIGS
ANQRHPGSHSPPFRSKGSGPHDNTEDKSTEPKPDLDDKRGPRDSGANEQP
>MCBN011322457_Phodopus_sungorus
EFSELRSQIIYTTGIEFGDLDYCLTLCIHCAYEGSQEAFMGSPACRYLGRNVIACKTLNHSTTNKLGEFIPDTSPCWRTKHALALAQNANEQGKVLFLSFCSLFLPKLVVGEAACLEKVQREIVHREQGLFQYPQCWATYGFKIIIFRLMRASFTFQFMLIHQVISLIADHDPNDIIITQA
RFSGFLIIKAVLDHILPKGDRVTLHPLVRSIRVRSEVESFKLALKGLTGHGEYAPLVHILNLSGVSNLEIGLFPQLSAIAGTATGHGCSLVGITVGENYQGLREPATEAERRLQEYAEEQEINPSGLDKTDTKILVDFYKKKHEIGDVEEILRAKHEKLKQLAAAISAANQRQLAIRGIQGA
MIRPLTKTIWEINPQNQSLIWMTRKDQVIAAPTDNTQQQFPILADDSGSNDDQDHGSPNSVPSFPQGSPLASREPLNEGRFSEDLEDSQPELFQGSRYLMTRMASLSALSSYPSRALENLNDLDAQQYLASARGSLPSIQEVPGKELPPTSHLPVYLTQHDRIGIGQSVASQWT
IIDSTPQAFAQLSGAQQARHSLTPEQLQEVEGAFNSCVEGKSFYAGLTFYHISIRLTKEFAIKTG
>APMT01196896_Mesocricetus_auratus
LCSQSIYAAEAGVEFEDSLDCCLTLCIHHAYKGNQEAFMGSPECEYLERNGTAFEILSHSTRNNLAEFIPDTSSCQRTKHALQALVQNASERRQKNHFSHRQVGPFLSFCSLFLPKLAVGKAARKPQHQVVVHREQGLSQYSQSWTASGFMIIIFRLMRASFTFKFLLLHQSYHLIA
DHAPNDIVITAQARLSGLLIVKVLDHILHKRDRVILHPLVGIIRVRSEVESFKLALEGLDGHGEYAPFARILNLSGVSSLELGQFPTEAERSLQEYPEEQEVNPGLDKTETKILVDFHRRKTRGLVTSRQRRSKLEQLAAAIGAANQVATSHRGHPRSHSPPFARKGHQATLRDNMED
KPQSQSLIWMTREDQATAVPVINPQRGDACYKQRPFPILADDFSSDEGRDHGSLNSVPSFLQGPLGPHGSPMGATSPKTLKSH
>PVIT010050105_Onychomys_torridus
NMEFHGINYGLSHSLVHNRKIYSLYVPEPSELCSQIIHAVEAGIELGGLMDSCLMTPCIQHTHEGNQKAFETSVAGRSLEAHGLMLKVLDTNSNLSEVLHQLVLVGGQDKHWVALMDNAREHQRVGQSLSFYSLFLPKLVVGEKACPEKVQRQIVIRAEQGLINLPQGWQTPG
FMITLFGIRRLSFTCKFLLTHQRISLEAGHAADTIITNSVSQTSVSGLLTVKTVLDHILHKVGHQILLHPLVTLSFQPALKGLAYHEEVPPFAQISNLSGVSSLELGLFLRLSNTALVATAHGGTLARLTVGEPYQQLREAAGEAERRLQLRSRENLGLDHAEAKILLDFHQKQYETGNQ
QAQIILAMKREKLQHLAPVKGSHVGHTPPQHHLKPNRCPRYTTQPPSTGITERDFEDADSLEGDDNDRGELSDRKGTPAPRSLSQILDTESESEDGRRTRGPGETHTKAPNCGPTDLKPNMEHIQDGPTLEQPPKTFSGNFPGSCQNPSSLA
>LZPO01008779_Neotoma_lepida

RTTQAPWQSNTQHHFELHQVLNYGTQPVESLVRNRWIYALEVPGPGELCWQIILXWVDLGELIDSYILLTCVQHTHKGNQKAFETSAAHKYLVAHGLXLTVFDNKANSNRLNSSTTGPGHKRQVPEMPITDNMRERRPVGQFLTFCSLFLPKLVVGEKACLEKKVQRQIVTHA
EQGLMQYPPNWQAPGFLTQASSIRRELPPLIHQGINLEAGNDADX
>PVIT010005287_Onychomys_torridus
NTQNDFGLHWILNYDTQPVEPFVRNRQIYALEVPDPGKLCLTHTVEYGVDLRDLINVCLLMCIQHMYEGNQKTIESSAARKYLVANRLMLKVQKANSSLIEFFTTGPGRKTRQALMPIMDNMGKCRPVGQFLSFSSLFLPKLVVGEKVCLEKVQRQIIYAEQGLTQYPTPPNW
QALVLTHIFSIMRVSFTFKFLLIHPQGINLEAGRDADIIITNSVVQARFLGLLIMKTVLKKGDRVVLHPLVTLKIRAEVDNFRFALKGLAQHDYVPSAQILTFSGVNNLEHGLFPQHWAIALRVSTVQGHALAGVTVGELYQQLCKAASKAKDEAARAEGQAIHQLGLDCTNAEILLD
FHQKTNRQYQQAKVILATKHEKPPKLAATITRTGQRVTWTLKVHLKLPRWIPVALRLPHPLKEQDFSEVAPQEETDEMDNLSKGSKASASQHLLILDAESADGERGNKSNVDKTGNFPRLRPSQGEAAGYARTKTRHSATTPLAVRTAILQSPCPITCWLPIEFSGAPNGPGSM
NTHPPTMPRNPPPQDQRQQTLDCLSPVWANPPTSNLSSNTNVDMYQPSPVYNRVPEFNRTLQRGTTLALEPLKTDSQQDPIKSEE
>CABHPP010142577_Peromyscus_attwateri
RPAWQSNTQNEFGLYPILNHGTQPVEPLVRHRQIYALEVPDPGRCCACTHTAESGVDLRDLINSCLLMGIQHMYEGNQKAFESIAAHVVANGLMLKVFDNKAKSNIIEFFTTGPGLRRDKPSCPSITWRKFLSFSRLFLPKLIVGENVCLEKVKQIYTEQGLMQYPPKLASSQLDI
VLSIMRVSFTFKFLLIHQGINLEADRDADNIVTNSVAQARFFGLLIIKMVLRLDPLVQTVKTRAEVDNLRFALKGLARHEYVPFAQILNLSGVNNMEYGLFPQHWAIALRVATAHGRALAGVTVGELYQQLHEAASQGQHVRAEGQAIHQLGLNCTKAEILLDFHQKNNMKSA
NQQAEVILATRREKLPKLAAAITRTGQRVTWTPRVHPTLSPPQAIQADTSGTQAAPPPQITEQDFSEVAPQEETDDEVDNLSQGSEVSASQHLEQILDAESGDESEVNKSNVDKTGNSPDFDPPRGASGYARTKTPHSAATPLAVGAAISRPCPIAGQFPIESPGAPNRPGKMN
THPLSTMPMDTLPQDQRQQLRIACRQSGPTLPHLQSILPNTNVDISAFPSPNRVPEFNRTLQGGTTLALEPLKTNSQQDPIKSEE
>CABHPH010136995_Peromyscus_nudipes
RPPWQSNTQNEFGLHPILNHGTQPVEPLVRHRRIYALVPDPGRCCACTHTVEYGVDLRDLINSCLLMGIQHMYEGNQKAFESIAAHVVANGLMLKVFDNKAKSNLIEFFTTGPGLRRDKPSCPSITWRKFLSFSSLFLPKLVVGEKVCLEKVQRQIYTEQGLTQYPSKLASSQLDI
VFSIMRASFTFKFLVIHQGINLEADRDADNIVTNSVTQARFFGLLIRKTVLKKGDRVVLHPLVQTVKTRAEVDNLRFALKGLAWHEYVPFAQILNLSGVNNLEYGLFPQHWAIALRVATAHGHALAGVTVGELYQQLHKAASQGQHVRAEGQEIHQLGLNCTKAEILLDFHQKN
NMKSASQQAEVILATRCVKLPTVGSHNHQDRPKGNLDLRVHPTFSPPQDIQADTSGTQAAPPPQITEQDFSEVAPQEETDDEVDNLSEGSEVSASQLLEQDSQCQVCQERGNKSIVDKTGNHSLTSTLPGEQLGTEYAPRHPILQQLHLLSEQQSHEDPVQSLEASSPNPLEH
QTD
>CABHPQ010172256_Peromyscus_aztecus
RPPWQSNTQNDFGLHPILNHGTQPVEPLVRHRRIYALEVPDPGRIVHCTHTVESGVNLRDLINSCLLMGIQHMYEGNQKAFESIAARVVANRLMLKIFDNKAKSNLIEFFTTGHDHKTRQALMSIMNYMEKCRPVGQFLSFSSLFLPKLVVGEKVCLEKVQRQIYAEQGLTQYP
QNWQALSLTVFSIMRASFTFKFLLIHQGINLEADRDADNIVTNSVAQARFFGLLIMKTVLKKGDRVVLHLLVQTVKTRAEVDNLRFALKGLAQHEYAPFAQILNLSAVNNLEYGLFPQHWAIALRVATGHVLAGVTVGELYQQLHEAASQGQHVRAEGQAIHQLGLNCTKAEIL
LDFHKKKHEISNQQAEVILTTRLEKLPTVGSHNHQDRPKSNLDSKGPSYFVSTSSYPGRYQWHSGCPTPSNNRARLFSCPSRRNCNGLICPKEAKSLPHSDLEQILDAESADGERGNKSNVDKTGNSPDFDPPRGAAGYVRTKTPHSAATPLAVGAAISRPCPITGQIPIESPGAP
NRPGSMNTHPLSTMPMDPQPQDQRQQLRIARHQSGPTLPHLQSIFPNTNVDISAFPCPNRVPEFNRTLQGGTTLALEPLKTNSQQDPIKSEE
>NMRJ02000008_Peromyscus_leucopus
WQSNTQSDFGLHPILNYGTQPVEPLERHRQIYALEVPDPGRCCACTHTVESGVDLRDLINSCLLMGIQHMYEGNQKAFESRAALIAANRLMLKVFDNKAKSNLIEFFTTGPGHKTRQALMSIMENMGKCRPVGHFLSFSSLFLPKLVVGGNVCFEEVQRQIIYAEQGLMQYPQ
NWQALCLTHVFSIMRASFTFKFLLIHQGINLEADRDANNIVTNSVPKPSFLGLLIMKMDLKKGELGVVFHPLVQTVKTRTEVDNFRFALKGLARHEYAPFAQILNLSGVTNLEHGLFPQHWAIALGVATAHGHALAGVTVGELYQHYREAVSQGQHVRAEGQAIHQLGLNCTK
AEILLDFHQEKNMKSATNRPRILATKREKLPKLAAAITRTGQRVTVDSERPSYFVSTSSYPGRYQWHSSCPTPQITEQDFSEVASQEETDDETDNLSEGSEVSTSQYLEQILDAESVDESEDIRQMLTRPETDPQTSTRPRGAAGYARTKTPHSAAILVKTRPIIGRLSIEFPGAPNRS
GSTLILFPTMPMDPRLPQDQRQQHPDCLLPVQANPPPPPIYPPNTNVDIYQPSPVPKGSPSSTGLSREEPPLNPSKPTA
>JAOPKW010000083_Peromyscus_maniculatus
NTQNDFGLHPILNYGTQPVEPLVRHRRIYALEVPDPGRCCACTHTVESGVDLRDLINSCLLVGIQHMYEGNQKAFESSAAPVAANGLMLKVFDNKAKSNLIEFFTTGPGHKTRQALMSIMENMGKCRPVGQFLSFSSLFLPKLVVGGKVCFEEVQRQIYAEQGLTQYPQNWQ
ALCLTHVFSIMRASFTFKFLLIHQGINPEADRDADNIVTNSVPKPSFLGLLIMKIDLKKGELGVVLHPLVQTVKTRTEVDNFRFALKGLARHEYAPFAQILNLSGVTNLEHGLFPQHWAIALGVATAHGHVLAGVTVGELYQKLCEAVSQGQHVRAEGQAIHQLGLNCTKAEILLDF
HQEKNMKSATNRPRILATKREKLPKLAAAITRTGQRVTWTLSVHPTLSPPQATQADTSRHSSCPTPQITEQDFSEVAPQEETDDETDNLSEGSISLYLTVSRANSRCVCERGHKSNVDKTGNRSPDFDPPRGAAGYARTKTPHSAAILVKNPSSCLRTAPHRISWSTKQILRKLNT
HPLPTVPMDSLPQDQRQQLQIACRQSEPTLPYLQSIHPNTNVDIYQPSPVPKGSPSSTGPSREEPPLWNPSKPTANKTQSSLRNKVWKTSSMQHHCYFSSALPLVLGGHSAETSKRVVP
>RCWR01024917_Peromyscus_maniculatus
GTQPVEPLVRHRQIYALEVPDPGRCCACTHTVESGVDLRDLINSCLLVGIQHMYEGNQKAFESSAALVAANGLMLKVFDNKAKSNLIEFFTTGPGHKTQALMSIMENMGKYWPVGQFLSFSSLFLPKLVLEEVQRQIIYVEQGLTQYPQNWQALGLTHVFSIMRASFTFKFLLI
HQGINQAADRDADNIVTNSVPNPGFLGLLIMKMDLKKGELGVVLHPLVQTVKTRTEVDNFRFALKGLARHEYAPFVQILNLSGVTNLEHGLFPQHWTIALGVATAHGHELAGVTVGLYQQLYEAVSQGQHVRAEGQAIHQLGLNCTQAEILLDFHQEKNMKSATNRPMSIL
ATKREKLPKLAAAITRTGQRVTWTLSVHPTLSPPQATQADTSRHSSCPTPQITEQGFSEVAPQEETDDETDNLSEGSISLYLTVSRANSRCVCERGHKANVDKTGNRSPDFNPPRGAAGYARTKTPHSAAILVKNPSSCLRTAPHRISWSTKQILRKLNTHPLPTVPMDSLPQDHR
QQLQIACRQSEPTLPHLQSIHPNTNVDIYQPSPVPKGSPSSTGPSREEPPLWNPSKPTANKTQSSLRNKVWKTSSMQHHCYFSSALPLVLGGHSAETSKRVVP
>JAOPKX010000018_Peromyscus_maniculatus
GTQPVEPLVRHRQIYALEVPDPGRCCACTHTVESGVDLRDLINSCLLVGIQHMYEGYQKAFESSAAPVAANGLMLKVFDNKAKSNLIEFFTTGPGHKTRQALMSIMENMGKCRPVGQFLSFSSLFLPKLVVGGKVCFEEVQRQIYAEQGLTQYPQNWQALCLTHVFSIMRASF
TFKFLLIHQGINLEADRDANNIVTNSVPKPSFLGLLNMKMDLKKGELGVVLHPLVQTVKTRTEVDNFRFALKGLARHEYAPFAQIFNLSGVTNLEHGLFPQHWTIALGVATAHGHALAGVTVGELYQQLYEAVSQGQHVRAEGQAIHQLGLNCTQAEILLDFHQEKNMKSATN
RPRSILATKREKLPKLAAAITRTGQRVTVDSEGPSYFVSTSSYPGRYQWHSSCPTPQITEQGFSEVAPQEETDDETDNLSEGSKVSTSQYLEQILDAESGERGHKSNVDKTGNQSPDFDPPRGAAGYARTKTPHSAAILVKNPSSCLRTAPHRISWSTKQILRKLNTHPLPTVPMDS
LPQDQRQQLQIACRQSEPTLPHLQSIHPNTNVDIYQPSPVPKGSPSSTGPSREEPPLWNPSKPTA
>RCWS02197384_Peromyscus_polionotus
IRPPWQSNTQNDFGLHPILNYGTQPVEPLVRHRRIYALEVPDPGCCACTHTVESGVDLRDLINSCLLMGIQHMYEGNQKAFESSAAPVAANGLMLKVFDNKAKSNLVEFFTTGPGHKMRQALMSIMENMGKCRPVGQFLSFSSLFLPKLVVWGKVCFEEVQRQIYAEQGLT
QYPQNWQALGLTHVFSIMRASFTFKFLLIHQGINQAADRDADNIVTNSVPNPGFLGLLIMKMDLKKGELGVVLHPLVQTVKTRTEVDNFRFALKGLARHEYAPFAQIFNLSGVTNLEHGLFPQHWTIVLGVATAHGHALAGVTVGELYQQLYEAVSQGQHVRAEGQAIHQLG
LNCTQAEILLDFHQEKNMKSATNRPRSILATKREKLPTVGSCNHQDRPKDNLDSEGPSYFVSTSSYPGRYQWHSSCPTPQITEQGFSEVAPQEETDDETDNLSEGSKVSTSQYLEQILDAESGERGHKSNVDKTGNQSPDFDPPRGAAGYARTKTPHSAAILVKNPSSCLRMAP
HRISWSTKQILRKLNTHPLPTVPMDSLPQDQRQQLQIACRQSEPTLPHLQSIHPNTNVDIYQPSPVPKGSPSSTGPSREEPPLWNPSKPTANKTQSSLRNKVWKTSSMQHHCYFSSALPLVLGGHSAETSKRVVP
>CABHPR010156129_Peromyscus_melanophrys
RPPWQSNTQNDFGLHPILNHGTQPVEPLVRHRWIYALEVPDPGPCCACTHTVESGVDLRDLINSCLLMGIQHMYEGNQKAFESSAAHIVANRLMLKVFDNKAKSNLIEFFTTGPGHKTRQTLMSIMDYMEKCRPVGQFLSFSSLFLPKLAVGEKVCLEKVQRQIYTEQGLMQ
YPQNWQALSLAHVFSIMRASFTFKFLLIQNPGINLEADCDADNIITNSVAQARFLGLLIMKTVLKKGSQVVLHPLVQTVKTRAEVDNFRFALNSCRHEYAPFAQILNLSGVNNLEDGLFSQHLAIALGVVTAHGHVLAGVNIGELCQQLREAASKAKEIAVRAEGQAIHQLGLNCT
KAEILLDFHQKKKPNRQYQQAEVILATRCEKLPKLAAAITRTGQRVTWAPRVHPTLSPPQAIQADTSGTQAAPPPSNNR
>CABHPH010163383_Peromyscus_nudipes
RPPWQSNTQNDFGLHPILNHGTQPVEPLVRHRWIYALEVSDPGPCCACTHTVESGVDLRDLINSCLLMGIRHMYEGNQKAFESSAAHKWPTGCKYLTIKQSNLIEFFTTGPGHKTRQTLMSIMDYMEKCRPVGQFLSFSSLFLPKLAVGEKVCLEKVQRQIYTEQGLMQYPQN
WQALSLAHVFSIMRASFTFKFLLIQNPGINLEADCEADNIITNSVAQARFLGLLIMKTVLKKGELGVVLHPLVQTVKTRAEVDNFRFALKGLARHEYAPFAQILNLSGVNNLEHGLFSQHLAIALGVVTAHGHVLAGVNIGELCQQLREAASKAKEIAVRTEGQAIHQLGLNCTKAE
ILLDFHQKKKTMKSANQQAEVILATRRENLPKLAATITRTGQRVTWAPRVHPTLSPPQAIQAGTSGTQAAPPPSNNRARLFSSPQEETDAEVDNLSQGSKVSASQHLEQILDVESAHESEEISQMLTRPETDPQTSTLPGEHTPRHPILQQLHLLSEQQSHEDPVHIAGQLPIESP
GAPNRPGSMNTHPLSTMPMDPLPQDQKEATLDGLSPGQARPGQANPPPPPVYPPTLMLTYQPSPVPTGPLSSTGLSREEPPLWNPSKPTA
>VALE03000002_Peromyscus_californicus
RPPWQSNTQNNFGLHPISNYGTQPVEPLVRHKADLCSGSARSWASCACTHTVESGVDLRDLINSCLLMGIQHMYEGNQKAFESSAAHVAANGLMLKVFDNKAKSNLIELFTTGPGHKMRQALMSIMDNMGKCWPVGQFLSCSSLFLPKLVVGEKVCLEKVQRQIYAEQGL
MQYPPPQNWQALSLTHIFSIMRVSFTFKFLLIHQGTNLEADRDAYNIITNSMAQARFLGLLIMKTDLQKVDRVVLHPLVQTEVDNFRFTLKGLARHEYAPFAQILNLSGVNNLEHGLFPQHWAIALGVVTAHGHTLAGVTVGELYQQLREAASPGQQHVRAEGQAIHQLGLN
CTKAEILLDFHQKKNEIGNQQAKVILATKCGKLSTVGSHIHQDRLKANLDSEGPSYFVSTSSYPGRYQWHSSCPTPSKNRADFSEVVPQETDEEMDNLSEGSKVSASQHLEQILDAESANESEEISQTLTSPETDPTLTLLGKQLGTEHTPRHLILQQSHEDPVQSLEVSSPNPLEHQ
TDPEEAEHSSSSHHAKERLTTTPGSEVATLDCPSPVWANPPPPPIYLPNTNVDISAFPSPNRALSSTGLSREELPLWNPSKPTA
>CACRXM010000023_Peromyscus_eremicus
LGNQTPKTTLDYCPILNYGTQPVEPLVRHRQIYALEVPDPGRTVHCTHTVESGVDLRDFIDGDSAHVEGNQKAFESSAALQRPTGCKYLTIKQSNLIELFTTGPGHKMRQALMSIMDNMGKCWPVGQFLSFSSLFLPKLVVGEKVCLEKVQRQIYAEQGLMQYPPPKKKNWQ
ALGLTHVFSIMRVSFTFKFGGPPKKKNWQALGLTHVFSIMRVSFTFKFLLIHQGINLEADRDAYNIITYSMTQARFFGTFDHVRRTSRKGDRVVLHPLVQTVKTRTEVDNFRFTLKGLAWHEYAPFAQILNLSGVNNLEHGLFPQHWAIALGVATAHGHTLAGVTVEELYQQLH
EAASQGQQHVRAEGQAIHQLGLNCTKAEILLDFHQKKNEIGNRPRILATKHEKLPKLAAAITRIGQRLTWTLRVHPTLSLPQATQADTSGTQAALPPQIIEQDFSEVVPQEETDEEMDNLSEGSKVSASPASRANSCRVCRERNKSNIDKTGNRSPDLTPGEQLGTEYVPRYLILQ
QSHEDPVQSLEVSSQNPLEHQTDPEEAEHSSSSHHAKELLTTTPGSEVATPDCLSPVWANPPPPPIYPSQHCHISLPQSQQGPVQQDSPGSEPPLWNPSKPTANKIQSSLRNKVWKTSSVQHHCYFSSALPLVLGAGHSAETSPRQL
>CAJEUE010005946_Arvicola_amphibius
IRPGGTQVLLTLMGNNNGHRKAGQVISFCSLFLPKLVVGERASVRMRRSNAVHAEQGHVILVLFLESGLAKFMLLHQGINLEAWHGAVIITNTVAQAFSVKTVLDHIRRSIIHPLVHMSKVKPEVDTFEVHQQEYGPFADPLSGVSDLEDGLLPQLPAIALGLAEAHGGTTLAEN
GVGGKHQLCMTCSLSERHQQEHAEEEISQLGFDYAKTKILDFHKRKLDIGTQQAALILTTKREKLKQFTVAVATSQTWSPQDHPVPTLPQFHLPDAHPAVLSDQEIFWSKILSFVITEYNPDEPSGYDGQRDGPTYRLLDAGLGSEDYHSHSTTPSFRDPSWPHGTPIDQTIGSSE
CSQPSLLEAGSEVGQRSQVPSDIDHHTSRIENVSTSHQKLPSSSIESLWQLPSCLMKSGSTLPSNHFCLPSTQHPMPKEVAQWSQAKPPEMLHSPNTTSRLLPGAISNSTQPKRGTFKVEEAYLYCVHNQGLHAGLLYYHTTLDEPKNFSNQSSGLFIYPVSLAGEIVAIITPNGW
QLREDSTITISGQGQLWCNLNMQSHL
>CATLKP010002317_Chionomys_nivalis
NQLYPDIGPGRGTQVLLTLTGNNNGHRKAGQVLSFCSLFLPKLVVGERASVHREGTMLSILNKAWNSGFMIFGIMKTSKFMLLQEGINLEAHGAVSNTVAQAFSVKTVPVTYKGDIILHPLVHMSKVKPEVDNFKAHHQEYGTFADTLSGVSDLKDGLLPQLPAIALGLSEPHG
GTTLAEIGMGEQHQEVASQAKRRQQEHAEEQEISQLGLDHAKTILDFHQKKLDIGTQAALILTTKREKLQQFTVTVVKTSQTVTQDHPVPTLAQFHLPDVHP
>LOJH01021770_Ellobius_talpinus
EFYPDIGPGGTQVLLTLMCNNNGYRKGGQVLSFCSLFLPKLVVGERASVLEKVQCQIAVHTEQGLEFWFYDFNHENAKFMLLHQGINLEAWRGAVIITDTVAQARFSVKTVLVIYKRDQIILHPLVHMSKVKPEVDNFARHEYGPFADTLSGVSDLKDGLLPQLIALGLAEAHRG
TTLAGTGMGEQYQLREAASQAERCPQEHAEEQEISQLGLDHAKTKILDFHKRNFDIGTQKTALILTTKGEKRQQFTVAVATTSQILVTQDHPLPTLPQFHLPDAHPVVLSDQEINILEQDHELICNLDIENPDEPSGYDGQQDQGPTYRLLDAGSSSEDYHSTYYKLSDDPSWPH
GTPSSRDQTIERSERSQPSLLEAGQAARSQVPSQMTIALLEFGKISVLVIKSFPLVQFERAFGNCQAASMGPLSLHRTTSASHLPNAQCQKEVTQWSQAKPSEMHHSPNTTSRLLPGESNNANLNEAFKVEEAYLNCVHNQSLHAGLLYYTNGRPKNFSTQSSGFFIYPVSLAGEI
VAIITQNGWQLREDSTITTIGQEQLCNLNMQSHF
>LOJG01026947_Ellobius_lutescens
EFYSDIGPGGTQVLLTLMDNDNGHRKAGQVLSFCSLFLPKLVVGERASILEKVQCQIAVRTEQGLELWFYDFGIMRTSYSLLLHQGIVIITNTVAQARFSVKLFWSYIKEHIILHRLVHTSKVKPEVDNFKARLQEYAPFARYSVGVSDLRDGLFPQLSCVALGLAEAHRGTTLAEIGV
GEQYQLREAGSQAERRQQEHAEEQEISQLGLDHDKTKILDFHKRNRDIGTQDCNFSRGRKSNSLQLLPVKLGHSGSSIAHIASVSPSRCTPCSAKPGNKYSGARSADLSRHKQSRTIGYDGQQDQGPTLLDAGSSSEDYHSALQAFCDPIWPRGTPSSETKQLEGQSVQPSFLEA
GQATRSQVPSHMTTTLLEFGKISTLDQKLPSSSIEPLAIAKLPDEAGSTLPSNHFCLPSTQCPMPKRNDPVVTSQTSDAAFSQHNFAATRCISKQRPTMALKVEEAYLNCVHNQSLHARLLYYTTGRPKNFSTQSNGLFIYPVSLAGEIAITQNGCQLREDSTITIIGEQLWCNLNM
QSHF
>JAJHZJ010000066_Bettongia_penicillata
HNLLEVGIIKQSMQGQRILIFYAKKDLNCLCCRIICVFGARVDLEDHLDHCLLTFLICYYYDGDIKKFLDSTVVQYLSEHGYEVHKLGGDKTTSLGKYLDRKLSKSTITPRNPHGEKLIFSFCSLFLPKFLGETCIQVLCQVWVHVEDLYEFPQRWVTTASIKIIFKIMHQNFLPKFINIHQ
GIHQQAGHDAYDVIVAASVAQACFSGLLIDKTVLEHIF
>JAJHZJ010000066_1_Bettongia_penicillata
HNLLEVGIIKQSMQGQRILIFYAKKDLNCLCCRIICVFGARVDLEDHLDHCLLTFLICYYYDGDIKKFLDSTVVQYLSEHGYEVHKLGGDKTTSLGKYLDRKLSKSTITPRNPHGEKLIFSFCSLFLPKFLGETCIQVLCQVWVHVEDLYEFPQRWVTTASIKIIFKIMHQNFLPKFINIHQ
GIHQQAGHDAYDVIVAASVAQACFSGLLIDKTVLEHIFFFFIFIMHFHRILTSEFSSHLASFHLKTSCILIPFSQSALPSITSLPSLIPIFSPFLG
>JAQQSK010000304_Potorous_gilbertii
VRTKKLIFYAKRDLNCLGCRIIYVFGAGVDLEDHLDHCLLILLICYYYDGDIKKFLYSTVVQYLSEHSYEVHKLGGDNTTSLGKYLNRKLSKSTITPSNPQGEKLVFSFCSLFLPKFLGEACIQVLCQVWVHVEGLSEFPQMWVTTASSKIIFKIMHQNFLPKFINIHQGLHQQAGQDSY
DAIVAASVAQVCFVELLIAKTVLEH
>JAPYYB010000006_Lagorchestes_hirsutus
HNLLEVGIIKQSIARTRKLNSMLKNLKCLCCRIIYVFGAGVDLEDHLDHCLLTLLICYYYDRDIKKFLDSTVIQYLSEHGYEVHQVGGDKTASLGKYLDANYNPQLQQGIQAVPQGETSQGKVGLSFCSLFLPKFLGEACNQKVLCQVWIHVEGLSEFPQIWASSANIKIIFKIMHQNF
LPKFTNIHQGLHEQAGHDAYDAILPASVAQACFAGLLIAKSVLEHI
>JAQMYV010000001_Macropus_fuliginosus
HNLLEVGIIKQSMKEQGNSSMPKNLKCCCRIIYVFGAGVDLEDHLDHCLLTLLICYYYDGDIKKFLDSTVIQYLSEHGYEVHKLGGDKTASLGKYLDANYQNPQLQQAIQAVPQEETGQGKVGLSFCSLFLPKFLGEACNEKVLCQVWVHVEGLSEFPQIWVTSANIKIIFKTMHQ
NFLPKFTNIHQGLHQQAGHDAYDAILPASLAQACFAGLLVAKSVLEHIFFFLVFIIHFHKVLSSKFSPHLSSLHLKTSCILTTPSPNLLSFLSHPSLPLSAFSL
>JAQQTA010000001_Macropus_giganteus
HNLLEVGIIKQSMKEQGNSSMPKNLKCCCRIIYVFGAGVDLEDHLDHCLLTLLICYYYDGDIKKFLDSTVIQYLSEHGYEVHKLGGDKTASLGKYLDANYQNPQLQQAIQAVPQEETGQGKVGLSFCSLFLPKFLGEACSEKVLCQVWVHVEGLSEFPQIWVTSANIKIIFKTMHQN
FLPKFTNIRQGLHQQAGHDAYDAILPASLAQACFAGLLIAKSVLEHIFFFLVFIIHFHKVLSSKFSPHLSSLHLKTSCILTTPSPNLLSFLSHPSLPLSAFSL
>WOXC01003109_Gymnobelideus_leadbeateri
HNLLEFELNITKYARARKLIFYATKNLNCLCCRIFDVIEAGVDLGDHLDHCLLTLLICYSFEGDNKKFQHSPVAQYLSEHGYGHQLDGDKTASLGKYLDTNNQSPQFQQAIQAVPQGETSKRRVGLFISFCSLFLPELLGWAACLEVLCQVRVHVEQGLAEFPQIWVTTATMKIMV
KIMHQRFLLKFITVHLGLHQAEHDAYVKIIATSVAKIRFVGLLIVKTVLEHILHFTELYPLTWGRSLQNELYNLHLALKEISKHVEFMPQMLLNLARGNHLEQRQYPQLSAICFGVTSVHECTLMEIKLDDRYQTLQEAAHKAESKLNKLRPKKYIQNIYSEKDKNI
>WOXC01007833_Gymnobelideus_leadbeateri

HDLLEFGLNIQTKYARARKLIFYATKNLNCLCCRIIDVIEARVDLGDHLDHCLLTLLICYSYEGDIKKFQDSPIVQYLSEHGYKFINWVRSLTASLGKYLDTNNQSPQFQQAIQAVPQGETSKRRVGLFISFCSLFLPELLGGGACLEVLCQVRVHVEQGLAEFPQIWVTTATMKIIFKIS
NFLLKFITIHLGLHQQAEHDAYVKIIATSVAKIHFVGLLIVKTVLEHILHFTEDGVELHPLTWGRSLQNELYNLHLAIKEISKRVEFMPQMLLSLARGNHLEQSQYPQLSAVHECTLMEIKLDDRYQTLEAAHKAESKLNKLRPKKYIQNIYSEEDKKSIASKVKASKKNMNWSQAME
ELKKD
>JAMXIF010001212_Petaurus_breviceps
HDLLEVGLNIQTKYARARKLIFSATKNLNCLCCRIIDVIEAGVDLGDHLDHCLLTLLICYYYEGDVKKFQDSQYLSEHRYEVHQLGGDKTASLGKYLDPNNQSPQFQQAIQAVPQGETSKRRVGLFISFCSLFLPKLLGGEACIEVLCQVRVHVEQGLAEFPQIWVTTATMKIIFKMM
HQHFLLKFIIIHLGLHQQAEHDAYDKIIATSVAKIHFVGLLIVKTVLEYIWHFTEDGVELYPLIWGKSLQNELYNLHLAIKEISKRMEFMPQMLLYLARDNHLEQGQYPQLSATSVAVASVHECTLMEVKLDDRYQTLEAAHKAESKNKLRPKKYIQNIYSEEDKKV
>JAQQSW010006939_Pseudocheirus_occidentalis
HDLLEVGLNIQTKYARTRRSIYATKDLNCLCCKFKLFKLLCIHVIEAGVDLGDHLDCCLLTLLICYYYEEDIKKFQDSTVAQYLSEHGYEVHQLDGDKTASLGKYLDTKNQSPQFQAIQAVPQGETSKRRIGLLISFCSLFLPKLLGGEACIEQVLCQVQVHVDQGLAEFPQTWVTTAT
MKIIFKILRVKFIIIYQGLYQQTEHEAYDTIIAASVAQACFEGLFVKTVLEHILHFTENGELRFHSLVLGRSLQNELDSLHRVIKEISKHRIYAPYAQLLNLARGNHLEQGQYPQLLAIAIGVASVTEHTLMEVKLDDRFRTLEAAQKAESKLNKYTDSRYIQNTYSEEDNEVKAPAYKASK
KSMNWSQAMEELRKDFENRSKRSTEKLGREMRMMQENNEKMSQQLAKGGPKNAEENNDLKNRQGAARWLSELSTGPESGGPEFRSSLRHLRQNPCDLGQVTPQLSYLPLSKKKKNTKPNGKKGSESQGEESLKKQTWSDGKDDAKVHRKLLENNGANAKWISMRNQ
EIIKQNTSSFKKLKKAAERENRNKNPPVTSKRSQIENSQEYSQIPEFPGQEENIASRQKKIQVLWNHNQDNTGFSSLYIKESEDLEYDVLGYQRNNENKKLLTENQVSMRGNKTFSVIAEFHAFLKEKSKIENLTFKYKTEEKHGKVNRKEKYKGLIK
>JAQQSV010000001_Pseudochirops_corinnae
HDLLEVGLNIMKYARARRLIFYVTKDLNCLCCRIIHVIEAGVDLGDHLDHCLLTLLICYYYEDIKKFQDSPVTQYLTEHGYEVHQLGGDKTASLCKYLDTNNQSPQFQAIQAVPQAETSKRKIGLFISFCSLFLPKLLSGEQACIEQVLRQVQFHVDQGLAEFPQTWVTTATMKIIFKI
MRIKFIIIYQGLHQQAEHDAYDTIIAASVAQARFAGLLIVKTVLEHILYFTEDGAKLHPLVPGRSLQNELDNLHRAIKEISKHRIYAPYAQLLNLARGSHLKQQYPQSSATAIGLASVHECTLMEVNTGQVVTLEAAQKAESKLNKLRLKRYIQNTYSEEDSEVKLLNTKPPRKIIGLRP
WKSSKRILKIKEKGKIEKRNESDARKSKMSQQFAKGGPKKYRKCLKKTRGSVAQIEHQPWSQEDLSSDPASDTHKTSHVTLDKSLKPNCPTFPFPKKKQKKNTRPNGKRNQKASEEKKALKSR
>JAQQSU010000002_Pseudochirops_cupreus
VSLHDLLEVGLNIQTKYARARRLIFYVTKDLNYLCCRIIHVNEAGVDLGYHLDHCLLTLLICYYYEGDIKKFQDSPVAQYLTEHGYEVHQLGGDKTASLRKYLDTNNQSPQFQAIQAVPQAETSKRRIGLFISFCSLFLPKLLGGERACIEQVLRQVQFHVDQGLAEFPQTWVTTATM
KIIFKIMRIKFIIIYQGLHQQAEHDAYDTIIAASVAQARFAGLLIVKTVLEHILYFTEDGAKLHPLVPGRSLQNELDNLHRAIKEISKHRIYAPYAQLLNLARGNHLKQQYPQLSATAIGLASVHECTLMEVKLDDRTLEAAQKAESKLNKLRLKRYIQNTYSEEDSEVKAPEYKASKKNIN
WSAMEELKKD
>JAQQSZ010000004_Phalanger_gymnotis
HDLLEVGLNIQSIARARTLIFYATKDLNCLCCRIIDVIEAVSLDHLDHCLLTPLICYYYEGDIKKFQDSPVAQYLSEHGYEVHQLGGDKTANLGKYLDANNQSPQLQQAIQAVPQGETSKGGVGLFISFSSLFLPKLLGGEQVCIEQVLCQVRVHVEQGLTEFPQTWVTTAIMKINFKI
MHQHFLLKFIIIHQELHQQAGHDATDTIIATSVAQARFAGLLIVKTVLEYILQLTEDGVELHPLARGRDLQKKLDSLHRAIKEISKHGIYALYALLNLAGDNHLQGLYSQLSAIATGVVSAHECTLMGVKLDDRYQALREAPTEAEKKLSKLRFERYIQNICSKEDNKVKAPTSKASKKN
MNCQAMGDLKKD
>JAANDE010000003_Trichosurus_vulpecula
HDLLEVGLNIQTKYARAKKLIFYATKDLNCLCYRIIDVIEAGASLDHLDHCLLTLLICYYEGDIKKFQDSPVAQYLSEHGYEVHQLGGDKTANLGKYLDANNQSPQLQQAIAVPQGEMSKGRVGLFISFSSLFLPKLLGTSLHTSLTSSLSPCGTRPHIPSDMGYRCHHEKSWSFHFVL
PVSPKIVEQVCIEQVLHQVRVHVEQGLTEFPQTWVTTATMKIIFKIMCQHFLLKFIIIHHELHQQAGHDATDKIIATSVAQARFAGLLVVKTVLEYILQLTEDGVELHPLAQGRDLQKKLDSLHRAIKERSKHGIYAPCALLNLAGGNHLEGLYPQLSAIAIGVASAHECTLMGVKLD
DRYQALREAAQKAERKLSKLRFERYIQNIYSEEDNKVKAPTYKASKKNMNWSQAMGDLKKD
>JAOEJA010000383_Phascolarctos_cinereus
HDLLKVGLNIQTKYVRARKLIFYAAKDLNCLCCRIIDVIEAGVNLGDHLDHCLLTLLICYYYEGDIKKFQDCPVTQYLSEHGYEVHQLCGDRTASLGKYLDANNHNPQLQQAIQAVPQGEMNKRRIGLFISFCSLFFQKLVVGECACIEEVLHQVQVHVEGLAEFPQIWVTIVTMKII
FKIMCQCFLLKFIIIHQGLHQQARHDASDIIIATSVAACFAELLIVQTVLDYVLQCTEHGVELHPLAWDRGVQNELDSFCQAVKQISKHGIYAPYAQLLKLAGVSQLKYGLYPQLSAIAIGIASAREKTLIGVKLDDRYQAIREAAYEAESKLSKLRCERYINIYSGEDNKVKVPTSKLSKK
NMNWSQAMEELRKD
>JAQQSX010000002_Vombatus_ursinus
HDLLMVGLNIQPKYVRARKLIFHAAEDLNCLCCRVIDVIEAGVNLGDHLDHCLLTLLICYYYEGDIRKFQDSPVAQYLSEHGYEVHQLCGDQTASLGKYLDANNHGPQLQQAIQAVPQAETNKRRTGLFISFCSLFLKLVVGERACTEEVFHQVRVHMEQGLAEFPQTWVTTAT
MKIIFKIMSSFLLKFIIIHQGLHQQATHDASDIIIATSVAHALFAGLLIVQTVLGYVLQCTEHGVELHPLTWDRVIQNELDNFRQAVKQISKHGIYAPYAQLLKLAGVNHLEYGLYPQLSAIAVDIASAHERTLIGVKLDDRYQALEAAYEAESKLRCERYIQNIYSEEDNKVKAPTSKAS
KKNMNWSQALEELRKD
>JAQQSX010000004_Vombatus_ursinus
HNLLEVGLNIQTKYARARKLIFYATKDLNCSCCRIIDVIEAGVNLGDHLDHCLLTLLICYYCEGDIKKFQDSLVAQCSKYEYEVHQLAGDQTARLGKYLDAIIGHNIQQATQAVGQGEINKGRIGIFISFCSLFLPKLVVGEVCTEEVLHQVWVHVEQGLPELPQTWVTTATLKIIFKM
MRSMFFFLLKFIIMHQGLHQQVRHDASDTIIATSVAQARFVGLLIVKTVLGHILQCTEHGVQLHPLALGGGLNEVDSFRAAKQVSKHGMYVPYVRLLNLAWVKGLEYGLYPQLSAFAIGVASAHDRTLIEAKLDDRYQTLEAAHETEKQVKQVKMREIHTNHIPEEDNKVKAPT
SKASKKNTNWSQAMKELKKD
>JAOEJA010000643_Phascolarctos_cinereus
HDLVEVGLNIQTKMARARKLIFYATKDLNCSCCRIIDMTEAGVNLGNHLDHFLLTLLICYYYERDIKKFQDRLVAKYSKHGYEVHQMGGDQTASLGKYLDAIIIHNYRKQLSALPQGGTNKGRIDIFISFCSLFLPKLVVGEVCTKGLLHQVWDNVEQGLAELPQTWVTTATMKIIF
KMTYQSFFLLKFIIVHQGLHWQARHDASNITIATSFSQACFAGLLIVKTILGHVLQQTEHAVELHRLALGEGLQNEVDNFCAVNQVSKHGMYVPYAQLLNLAVNQSEYLYPQLSAFAKGVASAHEKTLIQAKLDDRYQTLETAHEAESKLRCKRYIQTIYHNKTTKSKLLHLKASKK
NMNWSAMEELKKDFENQIREVEENWEEKLMQENDENESIAWRRPQNTEENNTLKNDTKQKRSKKPIRRTIPQKGEWVKWKRRSKSSLK
>WOXC01003232_Gymnobelideus_leadbeateri
LEVGLHIQIKYARARKFIFYATKDLNSLWCRIIDVIEAGFDLGSHLNHCLLTLLVCYYYRDIKFQDSGSPEHGHEVYQLDGDKTANLGKYVYANDQSPQLQQAIQAVLQGEMSKGRVGLFILFCSLFIPKLLGRERVCIEQISHQVWVHVEQGLAEPPQIRVTPATMKTIFKIMCQHF
LLKFIIVHQGLHQQAGHDAYDTIITTSVAQACFVALLIVKTVLESILQFTEDGVELHPLAQGSSLQNELDSLHAIKEISKHRIYAPYALLNLASDNHLEHRLYPQLSTIVIGIASVQEHTLMGVNLDVRYKALEAVHKGESNSKLRLERYIQNLYLKKDNKVKASKKNMNSQSMEELKKD
FENQVREVEEKLGREMRVMQENHEKQINSLLKETQKNTDENNILKKTRTNDKRGPKSQEEECLKSRGGQMENEVNLTEENNSLKIRMEQMDPNDSMRNQEIIKQNQDIFQEISRNTTQRVKKLKESTNHLLKISNKFPGILPNSRVPRSRRKYCKQPERNNSS
>JAMXIF010001174_Petaurus_breviceps
HDQLEVGLNIQIKYARARKLIFYATKHLNCLWCRIIDVIEAGVDLGDHLGHCLLTLLICYYYERDIKFQDSAVAQYLSKHGHKVYQLDGDKTANLGKYLDANDQSPFKQAIQEVSQGEMSKGRAGLFILFCSLFLPKLLSREPVCIEQVLHQVWVHVEQGRGEPPQTWVTNATMK
TIFKIMCQHFLLKFIIVRQGLHQQAGHDACYTVIATSVAQACFAGLLIVKTVLGHTLQFTEDGVELYHLAGSCLQNELDSLHAIKEISKHGIYVPNALLNLASGNHLEQGLYPLSTTAIDVASVQEHTLMRVKLDDRYLREATHKGESKSKRLERHIQNLHLKKRQQ
>JAQQSW010000004_Pseudocheirus_occidentalis
LEVGLTIQIKYAKERKLIFYATKDLNCSCCRIIDVIAGVDLGHHLDHCLLSLLVCYYYKRDINFQAGAQYLSKHEYEVYQLGRDKTVKLSGYLDANNQSPQLQQEIAVSQGEMSKGRVGLFISFCSQFLPKLLGRERVCIEQILHQVWIHVEQGLAESPQTWVTTATMKTIFKIMCQH
FFLKFIIVYQVLHQQAVHDAYDTIIATPVAQACFAGLLIVKTVLECILQFTEDRVELYPLAQGSSLQNELDSFHLAIKEISKHGIYAPYTFLNLGSGNHLEQGLYPQLSTTAIGIASVQEYTLNGVKLDLRYKVLDADPRGESKSKFTLERYIQNVYLKKDNKVKAPSSKASKKSMNSQA
MEELKKDR
>JAQQSV010000004_Pseudochirops_corinnae
QYQYKHGHEVSQLGGDKTANLGKYLDASNQSPLQQDIAVSQGEMSKGRVGLFISFCSLFLPKLLGREGVCIEQVLHQVWIHAEQGIAESSQTWVTTATMKTIFKIMCRAFFFLKFIIVYQILHQQAVHDAYDTIIATSVAQACFVGPLIVKTVLECILQFTEDRVELYPLAQGSSLQ
NELDSFHRAIKEISKHGIYAPYAFLNLGSGNHLEQGLSPQLSTIAIGFASVQKHTLMGAKLDVRYKALEAAHRVKASEANDLRDTHKTYTRKTTKAKLLHPKPLRKVIDLKPWKSSKKDFENQVREVEEKLGREMRVMQNNEKQINSLLKKTQKNTDENNILKNRLDQMTKEVQK
ANEEKNAEAEL
>JAQQSU010000003_Pseudochirops_cupreus
LKVGLNIQMKYAKARKLIFYATKDLNCLCCRIINVIEVGVDLRDHLDHYLLTLLVCYYVRDIKFLIQLVPQYQYKHGHEVSQLGGDKTANLGKYLDANNQSLLQQEIAVSQGEMSKGRVGLFISFCSLFLPKLLGREGVCIEQVLHQVWVHVEQGIVGFNPQTWVTTATMKTIFKII
CQHFFLKFITVYQILYQQAIHDAYDTIIATSVAQACFAGLLIVKTVLECILQFTEDRVELYPLAQGSSLQNELDSFHQAIKEISKHGIYAPYAFLNLGKCNHLEQGLYSQLSTIAIGFASVQEHTLNGAKLDVRYKALEAAHRVKASEANDLKDTHKTYTRKTTKTKLLHPKPLRKVIDLKA
WKSSKKDFENQVRELEEKLGREMRVMQNNEKQINSLLKKTQKSTDENNILKNRLDQMTKEVQKPNEEKNAEAEL
>WOXC01008483_Gymnobelideus_leadbeateri
HHHYTHDYGLFEHIRSKLIFYVTRDLDLLCCQIIDVIEAYVNLGGHLDNCLLTVLINYYYEGNIKNYDSPVALDTVDLSQFLDANNQNLPLQQVIQAVPQGELTKGRTGFFISFFSLLLPKLGENTCIEKVIRQIQVQSEQDLIEFPQTWTSTTNMKIIFVSMQQSFLLKVIIIHQGIHQA
KHDAADSIIVTSAVQICFNALLIIKTLLDHILLCTKDRIELHPHEAESKL
>JAQQSV010000004_1_Pseudochirops_corinnae
NNIFEVGLNIQTKAVRKLIFYVTKELDLLCYQIIDVIEADVNLGGHLNNCLLTAVINYYDEGDKTDSPVAQYLSEHGYSIHQFSDDTVDLSQLLDANNQNPPLQQAIQAVPQGELTKGRIGFFISLCISFIPKAYCGENTCIEKVIRQIQIHFEQNLIEFPQTWNLTANIKIFVSMRQSFLL
KVNIIQGIHQAEHDAADSVIVTSPAQIFFDSLLIIKTVLDHILLCTKNGIELHP
>JAQQSU010027108_Pseudochirops_cupreus
NNIFEVGLNIQTKYAVRKLIFYVTKELDLLCYQIIDVIEADVNLGGHLNNCLLTAVINYYYEGDIKNYDSPVAQYLSEHGYSIHQFSDDTVDLSQLLDANNQNTPLQQAIQAVPQGELTKGRIGFFISLCSLLLPKLIVGETCVEKVIRQIQVHSEQNLIEFPQTWNLTANMKIFVSMRQ
SFL
>JAQQSW010185563_Pseudocheirus_occidentalis
VFEVGLNIQTKYAVRKLIFIYVTKELDLLYYQVIDVIEANVNLRGHLNNLLTTVINYYYEGDIKNYDSPVVQYLSAHQLSDDTIDLNQLLDANNQNPPLQQAIQAVPQGELTKGRIGFFISFCSLLLPKLVVGEACIEKVIRQIQVHSEQDLIELPQTTSTANMKIIFISMQQSFLLKVSIIY
QGIHQAEHDAADSIIVISAAKYRFEGLLIMKTVLDHILLSTKDGIGLHP
>JAQQSZ010000001_1_Phalanger_gymnotis
HDVLEVELNIQTKCAVRKLIVFTTKDLDLLCHHIIVVIEAGVNLGDHPDNCLLTILINYYEGNIKKYNDSPIAQYFSEHGYSIHQLSRDDTVDLSQFLNANNQNHPLQRAIWTVPQGELTKVDFFFISFCSLFLLKLVMGKHACIEKGVGQIQVHSKQGFIEFPQTWTSKATVKMIFTSI
WQSFLLKFIIIHQGIHQQAGHDVTIIATLVTQAHFAGLLVVKTTFDHILLGTKDGIEFHPLDGRSLQNELDSFCAIKEISRQAISPYARLLNLAGVNLEHGLYLQFSTIAVGVAFDHGSLSGVNLDERYQALRESAHEAEMSLLGHLDMLEVKQHSGLEKRKKRSSQELRQQKNNISD
QKMEVAKAGLDLLNKLKVLAPKLTGEYKKSTSLVQHKIEGSGETSTHPKPELQLIRPASIDSSTSLVTSLKPPSINFQPSNTAIPEKLFYSPSGWKGRRNTSEFHPVFEKRGLPISRYSRRTKGAHRKARRNRICRRHGYWGWSATNSTKSAYPPQQKLSRPQSPPGLVSQDLI
>JAANDE010000002_1_Trichosurus_vulpecula
RDVLEFGLNITKHAVRKLIVFATKDLDLLCHHIIAVIEAGINLGDHLDNCLLTILINYYKGNIKKYNDSPIAQYLSEHGYSIHQLSREDTRRSFLDANNQNHPLQQAIRTVPQGELTKVDFFFISFCSLFLLKLVMGKHACIKKGMGQIQVHSRQGFIEFPQTWTSKATTKIIFTSIWQSFL
LKFIIIHQGIHQQAGHDVTIIATSVTQARLAGLLVVKTAFDHILLGTKEGIEFHPLAGRSLQNELDSFCAIKEISRNAIKSPYARLLSLAGVNLEHGLYPQFSTIAIGVAFAYGSSFGVNLDERYQALRESAHEAEMSLLGHLNMLEVKQHSGLEKRKKRSSQEFYPQKNNISKHQKREV
AKVGLDLLNKLKVELAPKLTEHEKSTSLVQHKIEGTGETSTHPKQELQLIRPASINSSTSPGDITQATIHKFPTINTAIPEKLFYPPSGWKGRRNTSEFHQVFEERALPISRYSRRTKGAHRKARRRYRICCRGYWGWSDTNSPKPKNLHTPPTKALLKASIPSRTDEPR
>JAOEJA010000399_Phascolarctos_cinereus
HDILEVGMNMQTKYAQVRKIFYATKDLDLLCCQIIVVIEAGVILGDHLDNCLLTLLINYYYEGNIKKYNDSPVAQYLSEHAYSIHQLSDNTVDLNQFLDANTQNPTLAIQAVPQGELTKERIAFFISFCSLFLPKLVKGEHAKIEKVTCIWVCSQQGLIEFSQTWTLIATMKITYTGVW
QSFFLKFIIIHGIHMTGHDATDSIIATLVTQACFIFQLGLLVVKTVLIRPQTLDTYLWDPGQVTPHCLPPPKKKTVLDHILLHTKDAQLHPLVQGRSLQNELDSFQHAVKKISRHGIYSHARFLNFAGVSQLENRLYPQLFAITIGVASVHGNTLIGVNLDERYQALRESAHEAETEMLE
VKHLRLGKKEEEILSEFHQQKNEIDKHSKGKLQARQDLLNKLKVQLAPKLTGEHEKSPSL
>MSTS01000053_Phascolarctos_cinereus
HDILEVGMNMQTKYAQVRKIFYATKDLDLLCCQIIVVIEAGVILGDHLDNCLLTLLINYYYEGNIKKYNDSPVAQYLSEHAYSIHQLSDNTVDLNQFLDANTQNPTLAIQAVPQGEITKERIAFFISFCSLFLPKLVKGEHAKIEKVTCIWVCSQQGLIEFSQTWTLIATMKITYTGVW
QSFFLKFIIIHGIHMTGHDATDSIIATLVTQACFIFQLGLLVVKTVLKSGLRHLTLTSYGTLGKSLNPIAYPPKKKTVLDHILLHTKDAQLHPLVQGRSLQNELDSFQHAVKKISRHGIYSHARFLNFAGVSQLENRLYPQLFAITIGVASVHGNTLIGVNLDERYQALRESAHEAETEML
EVKHLGLGKKEEEILSEFHQQKNEIDKHSKGKLQARQDLLNKLKVQLAPKLTGEHEKSPSL
>JAQQSX010000003_Vombatus_ursinus
HDILEVGLDIQTKYAQVRKLIFYVTKDLALLCHQIIDVIEANVILGDHLDNCLLTLLINYYYEDYIKKYSDSPVAQYLSEHGYSIHPLSDDTVDLNQFLDANTQNPTLQAIQAVPQELTKERIGFFISFPFASLFLPKLVMEEHAYIEKVMHIWIHSQEGLIEFSQTWTSTVTTKITYTSMR
QSFFLKFIIIHQGIHQTGHDAVDSIIATLVVQACFVFQLGSLVVKTILKSGLRHLTLTSYVTLGKSLNPIAPKKKKTVLDHILLHTKDAQLHPLCKAGLQNELDGFQHAVKKITRHGIYSPYARFLNFAGVNQLENGLCSQLFAIAIGVASVHGNTLIRVNLDERYQALRESAHEVENQE
MLEVKHLRLDKKEEEILSEFHQQKNEIDKHQQREVATRQDLLDKLKVQLAPRLTGGHEKSPSL
>WOXC01012125_Gymnobelideus_leadbeateri
HDVLEVGLNIQMKYARVRIVDFFATKDLDLLCYIIDVIARGTTFGKCLDNCLLTLLVIYYYEGDIKKYSGSPGLTQYLSEHGYSIHQLNCNDTTDLNQFLDANNKNPPQQAIQAVPQGEFTKGRIGFFISFCSLFLSKLVMGEACIEKVVYQIQVHSEQVLVVFTNWTSTATLKIIFAIM
WQNFPLKFAIIHQGIHQQAEHDASDTVIATAVAQACFAGLLIVKTVLDHILLHTKDGVELCFLARSRGLQNKLDGFQHAVKEISRNAIHIQKKELRNLN
>JAQQSW010000005_Pseudocheirus_occidentalis
YDVLVGLNIQTKYVRVRIVDFYSTKDLDLLCCIIDVIAAGINLGNCLDNCLLTLLINYYYEGDIKKYSGSPGLTQYLLEHGYSIHQLSDDATDLNQFLDANNKNPPQQQAIEAVTQGEFTKERIGFFICFCSFFLSELVMGEACIEKITCQIRVHSEQVLAEFLQTWTTTTMKIIFAIVWNF
LLKFVIIHQGIHQQAEHDALDLIIATSVAQACLAGLLIVKTVLDHILLCTKDRVELYPLARGRVFQNLDSFQYAVKEISRNAIHIQKEWSLNADGSILFVRSLFAQFCFFCLVVHSIGYNSLQHDCVNMFHANVCRAYIRSARCLGEGEEEEWGENLQLKLVELNVVNKKLIFKYRNKK
KTSTHRIYRLLNLAGVNQLDLLDKFKVQLALKLTGEHEKSPSL
>JAQQSV010000001_1_Pseudochirops_corinnae
YDVLEFGLNIQTKYARVRIVDFYSTKDPDLLCCIIDVIAAGINLGNCLDNCLLTLLINYYYEGDIRKYSGSPGLTQYLSEHGYSIHQLNRDDTADLNQFLDANNKNPPQEQAVQAVTQGEFTKGRIGFFISFWHFFLSKLVMGEACIEKVMCQIQVHSEQVLAEFLQTWTTTTMKIIF
AIMWQNFLLKFVIIHQGIHQQAEHDASDLIIATSVAQACLAGLLIVKTVLDHILLCTKDGVELYLLARGRVLQNELDSFQYAVKEIFRNAVHIRKRLWSLNADGSILFVLSLFCVLFCSFLWFILLVIILCSVTNVMCFMMYLSLYIACCVGEGEREMGRKFETKKLVELNVDKQKINFF
KNTEIEKKKISIHRIYRLLNLAGVNQLDLLDKLKVQLAPKLTGEHEKSPSLFQQNIDVTRENHNMPHQNSSLDLQVSIHLSLVTSFKPPSTNFQLTPLSLRSYSTLQMAGEEGIPLSPTDLDYFREEPPKLADILEDQMQLTACKCEEGTDPVMDTEDSGGAPPPDVPPEPKNLHTSQ
QQCSPQCQSPPGP

>JAQQSU010000002_1_Pseudochirops_cupreus
YDVLEVGLNIQTKYARVRIVDFYSTKDLDLLCCIIDVIAAGINLGNCLDNCLLTLLINYYCEGDIRKYSGSPGLTQYLSEHGYPIDQLNQDDTADLNQFLDANNKNPPQQQAVQAVTQGEFTKGRIGFFISFWRFFLSELVMGEACIEKVTCQIQVHSELAEFSQTWTTTTMKIIFAI
MWQNFLLKFVIIHQGIHQQAEHDASDLIIATSVAQACLAGLLIVKTVLDHILQCTKDGVELYLLAQGRVLQNELDSFQYAVKEISRN
>JAQQSZ010106136_Phalanger_gymnotis
HDLLEVGLNFQTKYTWVRIVDFYATKNLDLLCCIIDVIEEAGINLGDHLDNYLTLLINYYYEGDIKKYDSPVARYLSEHRYPVPQLSDDTTDRGQFLDANDQNPQQATQAVPQREFTKGRKDFFISFCSLFLPKLVVRKQACLEKVMHQIPVHSEQGLAECFPQNQASSASMKIIFTI
MQPSFLLKFVIIHQGIHQQTDSVTATLAAQACFASLLIVKTFLDHILLCTKDRVEYGIELYPLARGRGLQNELDGFRRAVKGISRHETYTLCARLL
>JAQQSZ010108307_Phalanger_gymnotis
HDLLEVGLNFQTKYTWVRIVDFYATKNLDLLCCIIDVIEEAGINLGDHLDNYLTLLINYYYEGDIKKYDSPVARYLSEHGYPVPQLSDDTTDRGQFLDANDQNPQQATQAVPQREFTKGRKDFFISFCSLFLPKLVVRKQACLEKLMHQIPVHSEQGLAECFHKIWASSTSMKIIFTI
MQPSFLLKFVIIHQGIHQQTDSVTATLAAQTCFASLLIVKTFLDHILLCTKDRVEYGIELYPLARGRGLQNELDGFR
>JAANDE010000005_Trichosurus_vulpecula
HDLLEVGLNFQTKYAWVRIVDFYAAKNLDLLCCIIDVIEEAGINLGDHLDNCLTLLINYYYEGDIKKYDSPVAQYLSEHGYPVPQLSRDDTTDRGQFLDANDQNPQQAMQAVPQREFTKGRKDFFISFCSLFLPKLVVGKQACLEKVMHQIPVHSEQGLAECFHKIWASSASMKII
FTIMRPSFLLKFVIIHQGIHQQAGHDASDSIIATLAAQACFASLLIVKTFPDHILLHQRTELSYRIELYSLARGRGLQNELDGFQHAVRGRRHERHETYTPYARLLNPARVNHLDLLDKLKSAVSPQIAGEHEKSLSLIQQNIDVLEETTIHPESEFQ
>JAJHZJ010000336_Bettongia_penicillata
YDILKIGLNIQTKYSQVRALIYATKDLDLLCTLSFEEDVTLGDHLDNCVLTLWTNYYHEGDWTTITPAQYLSERRYSFHQLSQDDTANLSQFLDTNNYNPPYSRQLQAVPQGELTKGRTGLFVAFCSLFLPKLFAGECACKEKAMCQICVHEEQGLTELPRTWTSKTTRKIIFTIMW
QGFLLKFILIYQEGHDVADSIIATLVAQARFAGLLIIKTMLDHILQSNKDGIELHPLAMGLQNELDAFWCTVKKISRHGIYSPYASLLNLAGLNQLEQLPAIGVASTHGSTLIGVNLGERYQGLRELAREIESNRWCEMLEVKHLKLAKEDEEIISEFLPQKNEINVHQQRETAKAGHN
LLEKLKVQLAPKPTGEQEE
>JAPYYB010000002_Lagorchestes_hirsutus
ITYDTMDSLYDILKIGLNIQTKYSQVRELIYATKDLDLLRYQSFEEDVNLGDHLDNCVLTLWTNYYEGDRQQSNPAQYLSEHRYSIHQLSQDNTANLSQFLDTNNYNPPLQAIQAIPQGELTKGRTGLFVALCSLFLPKVIVGKYTCKEKAMCQICVHAEQGPTEFPQTWTSKTTRK
IIFTIMQQVFLLKFILTYHGGHDVADSITATSVAQAHFAGLLIIKTMLDHILQSNQDGVELHPLVWAYRMNIPSSALPKKISRHGIYSPYASLLNLAGLNQLEHGLYTQLPAIAIGVVFTHGSTLIGVNLGERYQGLRELARETEQALRREMLEVKHVKLAKEDEEIISEFLHQKNEINIH
QQRETSKARQNLLEKLKVQLAPNPTGEQEE
>JAQOLO010000043_Notamacropus_eugenii
ITYDTMDSLYDMLKIGLNIQTKYSQVRELIYATKDLALLHYQSFEEDVNLGDHLDNCVLTLWTNYYEGDRQQSTPAQYLSEHRYSIHQLSQDNTANLSQFLDTNNYNPPLQAIQAIPQGKLTKGRTGLFVALCSLFLPKVIVEKYTCKEKAMCQICVHAEQGLTEFPQTWTSKITR
NIIFTIMRQVFLLKFILTYHGGHDVADSITATSVAQAHFAGLLIIKTTLDHILQSNKDGVELHPLVWAYRMNMPSGALPKKISRHGIYSPYASLLNLAGLNQLEHGLYTQLPAIAIGVASTHGSTLIGVNQGERYQGLRELAHETEQALRREMLEVKHVKLAKEDEEVISEFLQQKNEI
NIHQQRETSKARQNLLEKLKVQLAPKPTGEQEE
>JAQMYV010000001_1_Macropus_fuliginosus
DSRKKTSSSHDTMDSLYDILKIGLNIQTKYSQVRELIYATKDLDLLRYHIIEEDVNLGDHLDNCVITLWTNYYEGDRQQSTPAQYLSEHRYSIHQLSQDNTANLSQFLGTNNCNPPLQAIQAVPQGELTKGRTGLFVAPCNLFLPKLIVGKHTCKEKAMCQICVHAEQGLTEFPQTC
TSKTTRKIIFTIMRQVFLLKFILIYHGGHDVADSITATSVAQAHFAGLLIIKTTLDHILQSNKDGVELHPLAMGLQNELDAFRCAAKKISRHGIYSPYASLLNLAGLNQLHRLYTQLPAIAIGVASTHGSTLIGVNLGKRYQGLRELARETESKHWQREMLEVKHVKLAQEDEEIISEFLQ
QKNEINIHQQRETGKARQNLLEKLKVQLAPKPTGEQEESIISSTKNVTGGNHDSPQQNYNSDLQALKHPTSLVTPSRSPSTNLQTFKSLFQKSFSTLPIAGNEDEVSLIDLDKYFKDKPLQLAGIPEEIPPNVRQTPNPTADTEDSGGVQSPEVPPEHKGLHHLSATASSNTPYPCRT
NEPS
>JAQQTA010000001_2_Macropus_giganteus
DSRKKTSSSHDTMDSLYDILKIGLNIQTKYSQVRELIYATKDLDLLRYHIIEEDVNLGDHLDNCALTLWTNYYEGDRQQSTPAQYLSEHRYSIHQLSQDNTANLSQFLDTNNCNPPLQAIQAVPQGELTKGRTGLFVAPCSLFLPKLIVGKHTCKEKAMCQICVHAEQGLTEFPQT
WTSKTTRKIIFTIMRQVFLLKFILIYHGGHDVADSITATSVAQAHFAGLLIIKTTLDHILQSNKDGVELHPLAMGLQNELDAFRCAAKKISRHGIYSPYASLLNLAGLNQLEHRLYTQLPAIAIGVASTHGSTLIGVNLGKRYQGLRELARETESKHWQHEMLEVKHVKFAQEDEEIISE
FLQQKNEINIHQQRETGKARQNLLEKLKVQLAPKPTGEQEESIISSTKNVTGGNHDSPQQNYNSDLQALKHPTSLVTPSRSPSTNLQTFKSLFQKSFATLPIAGNEDEVSLIDLDKYFKDKPLQLAGIPEEIPPNVRQTPNPTADTEDSGGVQSPEVPPEHKGLHHLSATASSNTPYP
CRTNEPS
>JAQQSK010000001_Potorous_gilbertii
ITKVNTSRRRISSNTFNTLKVKIKINIIITYDTMDSLYDILKKGLNIQTKYLQVRESSMLPKIISYAAISFEEDVNLVDKLLPEQTTITPAQYLSEHRYSLHQLSQDDTANLSQLQSPLQAIQAVPQGELTKGRTGLFAFCSLFLPRLIVGEYTCKKCNGQICVHAEQGLTEFPQTWTSKATR
KIIFPIMWQGFLLKFILIYHGGHNVADSITDTSVAQARFAGLLVINTMLDHILQSNKDGVELHPLVWAAQNELDAFRYTVKKISRHGIYSPYASLLHLAGLNQLEHGLYTQLPAIGVASTHGSTLIGVNLGERYQGLRELARETESKHRWQMLKVKHLKLAKEDEEIISEFFQQKNEI
NIHQQRETAKARQNLLEKLKVQLAPKPTGEQEE
>JAQQSZ010000003_Phalanger_gymnotis
HDILKVGLNIQTKYAQVRELIFYASDLDPLGCQIINVIEAGMNLRDCLDNCLLTLLTNYYYEGDGRDSLVAQYLSEHRYSVHQLSQAGTEDLSQFLDSSNYNPRLQQAIQAVPQGELTKGRIGLFVAFCSLFLPKLIVGELACIEKVMGQILHSDQGLTEFPQTWISIGHENLFTIMPQ
GFLLKFVLIHQGIHQQAGHDAADSIITTSVAQAHFAGLLIVKSMLDHILLLTKDGAELHPLAGRGLQNELDGFRHPIKETPWDRIYSPYARLLNLAGVNQLDHRFMQLSVIAIGVASTYGSTLIRVNLDKKYQALRESAHEAESKLRWYEMLEVKHLKLAKEDEEVILEFHQQKNEIS
VHQQREVAKARQDLLEKLKVQLAPKHTGEQKKSPFIGSTKNVTRGDHDSPNRITISQTHKLNINLSGDDHLGHHPPIFNYYCRYSMKSYSTLLIAGSEDKGTTPEPCLRVFGTPPTGRHTRRMKGTHSLDARKTPTPNADLEDSEGAQPPKVLSTQRSTHPSLTASSKTPIFSKTNQ
PSSNAGIRNKILKLCRSFTPLQNWQKLIPDHRHFGFSRHDDFRALEGGLPSQGWYPSRHVPTAKGPVTWIDLSNDLRYKALMKGPKTFDEKKKQKTFPSRRKNSSRP
>JAANDE010000008_Trichosurus_vulpecula
ISLHDILKVGLNIQTKYAEVRELIFYATKDLDPLGCQIINVIEAGMNLGDRLDNCLLTLLTNNYYEGDGQYLVAQYLSEHRYSIHQLSQDGTADLSQFLDASNYNPRLQQAIQAVPQGELTKGRIGLFVAFCSLFLPKLIVGELACIEKVMGQIVHSDQGLTEFQQAWISTATMKIIFTI
RQQGFLLKFVLIHQGIQQQAGHDVADSIITTSVAQAHFAGLLIVKTVLDHILLLTKDGVELHPLADRGLQNELDGFRHPIKETFWDRIYSPYARLLNLAGANQLDHRFTQLSVTAIGVASTHGSTLIRVNLDERYQALRESAHEAKSKLRCEMLEVKHLKLAKEDEEIISEFHQQKNEI
SIHQREVAKARQDLLEKLKVQLAPKHTGEQKKSPFIGSTKNVTRGDRDSPNRITISRTHKLNINLSGDTTATTHQSSIITTFIPEKLLYLPIAGSEDKGTTPEPCLRVFGNRSNWQAHHKNERNSLVLDARRHPSPMQTLKILKVPNHRKVPPESKGLHTPYQHLQTPLSSPRPTSQSL
LKLES
>JAJHZJ010000509_Bettongia_penicillata
HDFLEVRLKIQTKYATARKLIFDATIDLDLLCCKITDVLEAGVNLGDHLDHCLLTLLIYYCYEGDIKKCHNSPVVSYLLEHRYAVHQLGEDRTDLSFLDASNQVPLLQQAIAVPQEELNRQIHFCSLFLTKLVVEKLACMEKLLCIWVHSEEDLVEFPQTWVSTAIMKIIFTMMQQSSLL
KFVIIHQGLHQQAGHDASDSIIATSVAQACFSQLLIVKTVLDHILLHTKHGAELHPLAANLSFLHAVKEISKHGIYIPLYARLLNLAEVNQFEHRLQLQLAAIAVGVASAHGSTLTGNVDERYQALRQTAHEAESKLRQCREILEVNSLGLEKKKEDILTEFRVPSEEECNQQTSTKGAE
DAGTTEKPKRQLAPRLTGKRERSISGPAYLNGDNTSPHQSITLTDKTLGECQDIIPKDFSHITIRKYSHFTPISLESQSTLQVTDSEEKGTSLSPDLDKYFRESPSQLADMPEDQIELTNCEFEEDTDPIMGTEDPGSALPFEFPPEPRNSHTSPTMAASSLSLLRSAAPLNRNQQKS
>JAPYYB010000004_Lagorchestes_hirsutus
HDFLEVRLKIQTKYARARKLIFYATKDLDLLCCKITDVVEVRVNLGDHLDHCLLTLLIYYYHEGDIKKCHDSPVVQHLLKHRYAVPNCLMGKTELTWASSSIQAIRTLYYNKLFKQSQEELNRQKILCSLFLTKLAVRKLACMEKLSHQIWVHSEQELVEFPQTWVSTDPMKIIFTMM
QQSFLLKFVIIHQGLHQQDGQDASDSIIATSVAQACFASLLIVKTVLDHILLHTKLGAELHPLAAELESFVHAVKEISKHGIYMPLYARLLNFAEVNQLEHRLPHLAAIAIGVASAHGSTLTGVNVDERYQALQQTAHEAESKLRQCREILEVKSLGLEKKRRILIEFHQKKNAISKHQQ
KVLKTKKELLRK
>JAQMYV010000005_Macropus_fuliginosus
HDFLEVRLKMQTKYARARKLIFYATKDLDLLWCKITDVVEARVNLGSSSRPFLLTLLIYYYYEGDIKKCHDSPVVQHLLKHRYAVHQFGGEDRANLGKFLDASNRNPLLQQAIQAVPRGTKTKNFCSLFLTKLAVRKLACMEKLSHQIWVHSEQELVEFPQTWAPTATMKIIFTTM
QQSFLLKFLLQGLHQQDGQDASDSIIATSVAQACFSQLLIVETVLDHILLHTKLGAELHPLAAELESFLHVVKEISKHGIYMPLYARLLNFAEVNQLEHRLPQLVAIAIGVASAHGSTLTGVNVDERYQALQTAHEAESKLRQCEILEIKSLGLEKKRRILTEFHQKKNAISKHQQKVLKT
KQELLRK
>JAQQTA010000005_Macropus_giganteus
HDFLEVRLKMQTKYARARKLIFYATKDLDLLWCKITDVVEARVNLGSSSRPFLLTLLIYYYYEGDIKKCHDSPVVQHLLKHRYAVHQFGGEDRANLGKFLDASNRNPLLQQAIQAVPRGTKTKNFCSLFLTKLAVRKLACMEKLSHQIWVHSEQELVEFPQTWAPTATMKIIFTTM
QQSFLLKFLLQGLHQQDGQDASDSIIATSVAQACFSQLLIVETVLDHILLHTKLGAELHPLAAELESFLHVVKEISKHGIYMPLYARLLNFAEVNQLEHRLPQLAAIAIGVASAHGSTLTGVNVDERYQALQTAHEAESKLRQCEILEIKSLGLEKKRRILTEFHQKKNAISKHQQKVLKT
KQELLRK
>JAQOLO010000057_Notamacropus_eugenii
HDFLEVRLKIQTKYARARKLIFYATKDLDLLCCKITDVAEAGVSLGDHLDHCLLTLLIYYYYEGDIKKCQDSPVVQHLLKHRYAVHQFGGEDRANLDKFLNASNQNPLLQQAIQAVPERNIDKKIPCSLFLTKLAVRKLACMEKLSHIWVHSEQELVEFLQTWVSTATTKIIFTMMQQ
SFLLKFVIIHQGLHQQGGQDASDSITATSVAQACFSQLLIVKTVLDHILLHTKLGTELHPLAAELESFLHAVKEISKHGIYMPLYAQLLNFAEVNLEHRLLPQLAAIAIGVASAHGSTLTGVNVDERYQALQTAHEAESKLRQCREILEVSLGLEKKRRILTEFHQKKNAISKHQQKVLKT
KQELLRK
>JAQQSK010000003_Potorous_gilbertii
HDFLEVRLKIQTKYARARKLIFYATKDLDLLCCKITDVVEAGVNLGDHLDHCLLTLLIYYYYEGDIKKCHNSPVVQHLLEHRYAVHQLGGEDRTDLSKFLDASNQNPLLQAIQAVPQEETKTKNSLCGLFLTKLVVEKLACMEKLLHIVHSEQDLVEFPQTWVSTAVMKITFTMMQQ
SFLLKFVIIHQGLHQQAGHDASDSISATSVAQACFSQLLIVKTVLDHILLHAKHGAELHPLTEFPTWSKGDLKAWNLYALYAWLLNLAEVNQLEHRLPRLAAIAIGVASAHGSTLTGVNVGKRYQALCAHEGESKLRQCREISEVKSLGLEKKEDFLTEFHQKKNAISKHQQKVLKT
KQELLRNSNWHLAPRLTGKQEKGPSLAQHNIDLLGDNTSPPNLLLIKLGSAKTFESPKTSLTLPSINIHYFTPIPLESQSTLQMTGSEEKGTSLSPDLDKYFRESPSQLADIPEDQIEPTNCELEEDTDPIMGTEDPGSTLPFEFPPEPRNSHTSPTMTVSTVSISLGANEPKPHTGINNR
>JAQQSX010000003_1_Vombatus_ursinus
HDSLEVKLNIQTKYARARKLASYATEDLDFLCIRVIDVIEASVNLGDDLYHCLLTLLICHYYESDIKKNHDIPVAQYLSKHGYGVHQLGEDDTTGLGKFFDASNQNPLLQKAIQAVLGELNRGRIGFFISFCSLFLPKLVVGEACIEKILCRIWVHSEQGPAEFLQTWVSTATMKITFIM
MWQSFLLKFVIIQQGLCQQAVHDAAVSIIATSVAACFARLLTVKTVLDGTVAQWIEHWHCSQEDLSLNLASDTLHLCQTQTLCDPGQVTPHCLAKKKKNKQNKTILDYILLCTKHEVELYPLAQGRSLQSKLNSVHMVKEISKHGIYSLYAQLLNFAGVNQLEHRLYPQLSAIAIG
VASTHGSTLIEVNLEKYQALQESAPQAESKLRQCEMLEVKSLGPEKEEEETLTEFHQKNAISKHQQKEVLKAKQELLKKLTTAVSPKMDRRTRQSPSLAQQNIDLLGDNKTHPKLELQLIKPSSAKTFESPVTPFGKPIHPPNNRGVNTPLSPANLDEYFKEGPSLLVDIPEYQTEPTD
CECEEDTDPTMGIGDLGGAPPFEVPVEPRNSPQPQQRHSPQPQSPESPYQTGISNRNPELHRDTSPIQD
>WOXC01007696_Gymnobelideus_leadbeateri
HDLLEFGLNIHTKYARARKLIFYASKDTDLLCCKIIQMWAGVNLGGHLDHCLLTLLIYYFYEGSINKHDSPVAQNLSEHGYAVSIHLGGDDTADLGKFLVPSNQNPLLQQAIQAVPGELNRGRTGLFISFCSLFLPKLVVAEQTCIKKVLLQIQVHSEQGLAEFPQTVSTATMKIIFKW
QNFLLKFIIHQGLHQARHDTSDSVNATSVAQVHFARLLIVKTVLDHILLCTGRRVEFHPLAGISFQNELDNFQCVAKEISKHAIYVFYAQLLNIPGVNLELCLQLSAIASAHRSTFIRVNLDERYQVLQESAHEVESKLRQHREMLEVKSLGLEKEKDILTEFHQQKNEISKHQKEVLKT
SRNYENQISPKTDRKTRERSISDPAYLTRRYNSSQTRTLNNLQGMPRHLNLQLLSDLPSTNIHFTPIPLESQSTLQVTGSEDGTYSRPVDFDEYFRKGPSQLADIPEDQDQTNRLMKGHPNRGYGPRCSTLQGSPTQKLTYPPPQHSPQPQSHQKPMSQGPTKLESIVETPSYIETL
SKYNDTKKPL
>JAMXIF010001182_Petaurus_breviceps
HDLLEFGLNIHTKYARARKLMFYASKDADLLCCKIIDVVETGVNLGVHLDHCLLTLLIYYYEGRIKKCHNSPVAQHVSEHGYAVSTNLGGDDTADLDKFLVASNQNPLLQQAIQAVPGKLNRGRRGLFISFCSLFLPKLAVAERTCIKKVLLQIQAHSEQGLAEFPQTISTATMKIIFK
RQSFLLKFVIIHQGLLQAGHDASDSINATSVAQVHFARLLIVKTVLDHILLCTKHRVEFHPLASGISLQNELDNFQCVAKEISKHAIYALYAQLLNLPGVNQLEHRLYLQLSAIAIGVASAHRSTFIRVNLNERYQVLQESAHEVESKLRQHQEMLEVKSLGLGKKDILTEFHQQKNEV
SKHQKEVLKTKQELLRKSNLAPRLTGKQEKGPSLVQHNIDLLGDNTTHPKLELLTDKTFRECQDIIPKDFSQITIHKYSLVHTHTLGKPIHPPSDRQRRHILETNRPVFQRRTLSVSRYSRKSNRTNRLMKGHWPNHGYGPWRCSAHSKFPLNPETHMPPQQQHSPQPQSHQKPI
SQGPTKLESTVETPSYIETLSKYNETRKPPYLTTFNGNPLVFPDLLISEELWKDVYLAQD
>JAQQSV010001530_Pseudochirops_corinnae
ICRMYFHCYKLNITHRLTTADLDHCLLTLLNYYYYGSIKKYHNSPVAQNLSQHGYRCQFITLVGMILLTWVKFLDVSNQNPLLQQAIQALPEELNRGRTGLFILFCSIFLPKLLFLPKLVVGERTCIKKVLLQIRVHSEQILAEFPQMWSLNCYYEAGHDASGAINATPVAACFARLLIV
KTVLDHILLCAKHGARVYPLAGISLQNELDSFQCVAKEISKHGIYALYAQLLNLPGINQLKHRLYLQLLAIAIGVASAHRSTLVRVNLDERYQVLQESAHEVESKLRQHREMLEVKSLGLEKKDILTEFHKQDEISKHQQKEVLNTKQELLRKSNCSPKTDRRTRERSISIPAYLTRNYNS
SQTRTLTDKTFRKCQDIIPNDFSQITIHEYSLLHTHTLGKPILLQVKGREEGTSLRLINLDEYFREGPSQLADIPDQIEPTDFECEKDTDPIMDIKDPGGALPFEVPPETHILPPQQQHSPQPQSHQKPMSQGPTKLESIAETSNYRDIEQIQDKTSKSNHFQWQPLGIPLAFRGTLER
>JAQQSU010000003_1_Pseudochirops_cupreus
ICRMYFHCYKLNITHRLTTADLDHCLLTLLIYYYYEGIKKYHTSPVAQNLSQHGYRCQFITLVGMILLTWVKFLDVSNQNPLLQQAIQALPEELNRGRRGLFILFCSIFLPKLLFLPKLVVGETCIKKVLLQIQVHSEQVLAEFPQIWSLNCYYEAGHDASDTINATSVAARFERLLIVKTV
LDPILLCTKHGVRVYPLAGISLQNELDSFQCVAKEISKHGIYALYAQLLNLPGVNQLKHRLYLQLLAIAIGVASAHGSTLVGVNLDERYQVLQESAHKVESKLRQHREMLEVKSLGLEKKDILTEFHKQNEISKHQQKEVLKTKQELLRK
>JAOEJA010001095_Phascolarctos_cinereus
HDLLEVKLNIQAKYSRARKLVCYATEDLDLLCFRVIDVMEARVNLGDHLDHCLLILLICYYYEGAFKKYHDSTVAQYLSEHGYVVHLGGDDITGLGKFLDASNQNHLLQKAIQVVPQGELNRGRIGFFISFCSLLPKLVVGEACMSYRQIVHSEQGLTEFLQTWVSTATMMIIFTM
MWQSFLLKFVIIRQGLHQQAGHDAAESIIATSVAPARFVRLIVKTVLDGSVKQWIEHQHYQEYLSSNLASDACHYQTQTLCDSGQVTPHCLAKKKNCFGLYPAVTKHEVELHPLAGRSLQSKLNSLQHMVKEISKHGIYSPYAQLLNFAGFNQLEHRLHKLSAIAIGIASSHASTFI
GLSLERYQALQESAPQDKSTLRQCEMLEVKSLGLEKEEEILTEFHQQKSAISKHQQKEVLKMKQEQLEKLKLQLAPRLKGEKDKSPSLAQQNIDLLGDNKTHPKLELQIIKPSSAKKFDSPVTPLKLPPPDIHSFTPIPLKSQSTFQLTGNEEAHPVQPILMNISRKAPLYSTFQNIKLNP
QIMNVKRTLTRPWHLGPRRSLHSSSVPPESRNSPHPQRHSPQLQSPQESMSQGPTKLESVIETLSHIEA
>JAQQSZ010000001_Phalanger_gymnotis
LEVGLNIQTKYARARKLILYATKDLDLYCRVIVVEAGVNLGDHLDHCLLTFIIYYYHEGNIKKLSDSTVAPHFSEHKYAVYQLGGDDTARLGEVLDARNQNPLLQKAIQAVPQGKLNRGRIALFISFCSLFLSKLVIGEACIEKVLCQIQVHSEQGLAEFLETWVLTATMKITFTMMW
QSFLLMFVIIQGLHQQVGHDALHSITATSVAQECSVALLIVKTILDHILLSKKHGVQLHPLARGISLQSKLDSFCAAKEISKHGIYAPFVLLDLAGVNQLEYEPYLQLSATTIGVASPHGSTLNGVNLHERYQDLESAHEAESKLRQSETSRVKSLGLEKEEDILTEFHQQNTFSKHQQK
EMLKNKRELLKNQTAVSPKTDRTKERCSGETKYLTRRQYNLCQTRTLTDKTFRECQDIIPNAFSQITIQEYSLLHNYTLGKPIHPPND
>JAANDE010000002_Trichosurus_vulpecula
LEVGLNITKYARARKLILYATKDLDLLYCRVIEWYEAGVNLGDHLDHCLLTFIIYYYHEGNIKKLSDSIVAPHLSKHRYAVYRLGGDNTARLGEVLDARNQNPLLQKAIQAVPQEENRGRIALFISFCSLFLSKLVIGEACIKKVLCQIQVHSEQGLAEFPETWVLTATMKIIFTTMWQS
FLLMFVIVHQGLHQQVGHDALHSITATSVAQECFAALLIVKTILDHILLSKKHRVELHPLARGISLQSKLDSFCAAKEISKHGIYAPYVQLLDLAGVNQLEYELYLQLSAITISVVSPHGSTLIGVNLDERYQALRKSAHEAESKLRQSRETSRVKSLGLEKEEDILTEFHRQENTFSKHQQ

KEMLKSKWELLKQTAISPKTDRTKERCSDEAYLTRRQYNLSQTRTLTDKTFRECQDIIPNAFSQITIQEYSLLHNHTLGEQSTLQVTGSKEGTPLSPFNLDEYFREGPSISRHSRRPNQTFRMMLRTLTESWVLRTSEVLCPSKIILNPKSHILLQQQHSPQPQSPQKPKSQGPTKPETI
VETPSYIEALSKYNEAGKPL
>JAJHZJ010000583_Bettongia_penicillata
QKVWKLTSLIISMGSLHDLLGVGLNIQTRYARARKLIFITKDLNCVCCRVIDVIELIEITTIDLLTLLICYYHEEDIKKLQDSPVAQYLSEHGYEVHTWANTMQKKFLWQQAIAVPQEETNKGGVGLLISFCSPFLPKLVVGEVCIEKREVQVHLEGLTEFSQTISTTTMIISCRHKAWSTS
PFRLRQGSTENELDSFQAIKHIYAPYVRLLNFAGAGREVNQLEHRLYPQLSAIAIGVASAHGSTLIGVNLDDSYQALRESAHECQEILKVKNLGPGKDEIILMEFHQKNTISKQQKRAKELLEKLKQQLAPGFTSRGSSPTRQRNIDVLGDNSLCPNLELQLIKPSSAKTLDSRNCILPS
SHPFLESQSTLPACAMRRKTLLVWQILEYFKERPQQIPEAKVNPMVGGHQLIYRQFGTQGCTALSSSTQKLTHHQTTPYNPHPPKSQKALSRLEPIKE
>JAQQSK010000001_1_Potorous_gilbertii
QKVWKLTSLIISVESLHDLLGVGLNIQTRYVRVRKLIFITKDLNCLCRRVIDVIEIEITTSDILTLLICCYHEGDIKIFQDSPVAQYLSEHGYEVHQLGRARLPDLGKYLDANKVLWQQAIAVPQGETKEAVGLLISFCSPLLPKLVVGEACIEKREVQVHLEGLTEFPQTWFLLPGLYPVGT
KHGVELHHLGKGRGLNELDSFQASKHIYAPYVLLNFAGASGGEHRLYPQLLAIAIGVASAHGSTLIGVNLDDRYQALRESAAHECQEILKVKNLGPGKDEIILMEFHQKNTISKQQKRAKELLEKLKQQLAPGFTSRGSSTHQHNIDVLGDNSLRPKLELQLIKPSSAKTLDSRNYTL
PSSHPFLESQSTLPACAMRRKTLLVWQILEYFKESPQRIPEVPGSQGDPIDGLRGTHSSTTGSLELPFEVPLEPKNSTHHQTTPYNRHFPKSQKGLSRLEPIKE
>JAPYYB010000002_2_Lagorchestes_hirsutus
LLEVGLNIQTRYARARKLIFITKDLNCLCCRVISVIELIEITYTIDLLTLLICYYHEGDIKKFQDSLVAQYLSEHGYEVHQREGIDCQPGQIPRCKQSKFLWQQAIAVPRRETNKGGVGLLISFCSPFLPKLVVREQACIEKREVQVYLEGLTEFLQTWSLYYHHDYILRTKHGVELCHLGKG
RGLQNELDSFQAIKALNYAPYVRLVNFAGVVRGENVNQLEHRLYSKLSAIAIGIASAHGSTLIGVNLDDRYQALRESAHECQAISKVKNLWPGKDEIILMEFHQQKNTISKQQKRAKELLEKLETALGPWIDKQTGKF
>JAQMYV010000001_2_Macropus_fuliginosus
QKVWKLTSLVISMESLLDLLEVGLNIQTRYARARKLIFITKDLNCLCCRVISVIGVNLGDNLDHYLLTLLICYYYEGDIKKFQDSPVAQYLYEHGYEVHQRDGIDCQPGQIPKCKQSKFLWQQAIAVPQRGTNKGGVGLLISSFLPKLVVGEQACIEKREVQVHLEGLTEFPQTWFLLP
PLYPVGTKHGVELHHLGKGRGLQNELDSFQAIKHIYAPYVLVNFAGVVRGENVNQLEHRLYSKLSAIAIGIASAHGSTLIGVNLDDRYQALRESAHECQEISKVKNLWPGKDEIILMEFHQQKNTISRQQKRAKELLEKLETALGPWIDKQTGKF
>JAQQTA010000001_3_Macropus_giganteus
QKVWKLTSLVISMESLLDLLEVGLNIQTRYARARKLIFITKDLNCLCCRVISVIELIEITTIIYLLICYYYEGDIKKFQDSPVAQYLYERGYEVHQRDGIDCQPGQIPRCKQSKFLWQQAIAVPQRGTNKGGVGLLISFCSPFLPKLVVGEQACIEKREVQVHLEGLTEFPQTWFLLPPYLV
GTKHGVELHHLGKGRGLQNELDSFQAIKHIYAPYVLVNFAGVVRGENVNQLEHRLYSKLSAIAIGIASAHGSTLIGVNLDDRYQALRESAHECQEISKVKNLWPGKDEIILMEFHQQKNTISRQQKRAKELLEKLETALGPWIDKQTGKF
>JAQOLO010000043_1_Notamacropus_eugenii
QKVWKLTSLLVWSHYYDLLEVGLNIQTRYARARKLIFITKDLNCLCCRVINVIGVNLGDNLDCCLLTLLICYYHEGDIKKFQDSPVAQYLSEHGYEVHQREGIDCQPGQIPRCKQSKFLWQQAIAVPQRETNKGGVGLIISFCSPFLPKLVVGEACIEKREVQVHLDGLTEFPQTWFL
LPPLYPVGTKHGVELHHLGKGRGLQNELDSFQAIKHIYAPYVRLVNFAGGGGRMSTNNIDLYSKLSAIAIGIASAHGSTLIGVNLDHSYQALRESAHECQEISKVKNLWPGKDEIILMEFHQQKNTISKQQKRAKELLEKLETALGPWIDKQTGKF
>JAMXIF010001178_Petaurus_breviceps
LLEVGLNIQTRYTRERKLICVTKDLNCLYCRLLIFESGVNLEDNLNHCLLLLICYYHEGDIKKFQDRPVAQCVSEHGYEVCQLGGDKTASLDKHLDADNKSPLLQQAIAVSQGETSKAVVGFFISFCSLFLPKLLVREQAFIEKVLCQVQVHLEGLAEFPQTWCFYHHHDHILRSKYEVE
LHPLVQGRQGSPERAKFNKPSKQGIYAPYVRFLNFAGINQLEHGLYTQFSAIAIGIASSHRSEPQIGVNLDDRYQTLKESAHKAESKLRHCQSLEVKSCGLGKAENILMEFYQQKNIISNNKKVMRAKQELLEKLKQQLAPGLTSGEGSPPSLQQNINLLGENSVLNKSLTKHSTKTF
DSSV
>JAQQSW010049037_Pseudocheirus_occidentalis
LLEVGLNIQTKYARERKLICVTKDLNYLCCRVTYVIESGVNLGDNIKHCLLTICWYRDGDIKMFQDSPVAQYVSEHGYEVCQLGGDKTASLNKNLDADNKSPLLQQAIQAVSQGERNKGIVGLFIFCTLFLSKLLVREQAFIEKVLCQVQVHLEQGLDEFPQTWVSITTMVISYRHK
AWGTLLFSPRAGISRMSLTAFNKPSKHRIYAPYVFLNFAGINHLEHGLYPQFSAIAIGIASAHGSTLIGVNLDDRYQTLKESAHETDSKLRQSCQQLESMMLEVKSLGLGKAENILMEFYQQKNTISIQQKEVMRAKQELLEKLKQQLAIDKQRGKSSSISAEHFTWRLTQSTRALV
NKNSTKTFDSSITPSKSTSETIYSLALTPLESQSTLPAVGLEAEGTPLSSADCDEYFRERYQQTPEIQGSQADPVDCLFGGHLYYGT
>JAQQSV010000001_2_Pseudochirops_corinnae
LLEVGLNIQTRYTRERKLICVTKDLNYLCCRVTDVIESGVNLGDNLNHCLLTLLICYYHEGGIKKFQDSPVAQYVSEHGYEVCQLEGDETASLDKNLDADNKSPLLQKAIQAVSRGETNKGIVGLFIFCSLFLPKLLVREQAFIEKVLCQVQVHLEQGLAEFPQTWVSTTTMIISYTKHG
VELHPLCKVGRGLQNELNSFQQTIKHGIYAPYVFLNFAGINQLECRLYLQFSAIAIGITAHGSTLIGVNLDDRYQTLKESAHEADSKLKSSCKQLESVILEVKSLGLGKAENILMEFCQQKNTINKQRQKEVMRAKQELLEKLKQQLAPGLTSREGGPFSLQQNIDLLGDNSLSPKQEL
LIKHSTKTFDSSVTPAKSTSETVYSLALAPLESQSTLPAVGLEEEGTPLSSAHCDEYFRERPQQTPEIQGSQADPRTLSIRGGHQLSYGTWGCIALSSSPQLTHSQLTYPTNPYPPRNQVKVSQDCEPIKESPSNTEAL
>JAQQSU010000002_2_Pseudochirops_cupreus
LLEVGLNIQTKYTRERKLICVTKDLNYLCCRVTDVIESGVNLGDNLNHCLLLLICYYCEGGIKKFQDSPVAQYVSGHGYEVCQLEGDKTASLDKNLDADNKSPLLQKAIQAVPQGETNKGIAGLFIFCSLFLPKLLVRERAFIEKVLCRVQVHLEQGLAEFPQTWVSTTTMIISYTKHG
VELHPLCRVGRGLQNELDSFEQTIKHGIYAPYVFLNFAGINQLECRLYLQLSAIAIGITAHGSTLIGVNLDDRYQTVKESAHEADSKLRQSCKQLESVISEVKSLGLGKAENILTEFCQQKNTINKQLQKEVMRAKQELHEKLKQQLAPGLTSREGSPFSLQQNIDLLGDNSVLNKNFS
NTKDTKTFDSSVTPAKSTSETIYSLALTPLESQSTLPAVGLEEEGTPLSSAHCDEHFRERPQQTPEIQGSQADPRTLSIRGEHQLGYGTWGCIALSSSLQLTRSQLTYPTNPCPPR
>JAQQSZ010000004_2_Phalanger_gymnotis
HDLLQVGLNIQTKYARIRKLIYVTKDLNCLCIDVIESGVNLGDNLDHCLLTLLIDSIYHEVDIKNFRDSPVAQYLSEHGCEGTSAGKGNCQPGQMPRCKQSPLLQQAITVPQGQTNKGRVGLFISLCSLFLPKLVVGEACIEKVLRQVKVHLEQGLAEFPQTWISTLLDHILQRTKHG
VELHPLVQGKGFQNKLDSFQSIKHGIYASYVRLLNFGGVNHLEHRLYPQLSVIAIDVASTHGSTLIGVNLDDKYQDLRSAYGAESKLRHCPEIVEIKSLGLRKDENILTEFYQQKNTMSKQQQKEVLRAKQELLEKLKQHLA
>JAANDE010000003_1_Trichosurus_vulpecula
HDLLEVGLNIQTKYARARKLIYVTKDLNCLCIDVIESGVNLGDNLDHCLLTLLICYCHEGDIKNFQDSPVAQYLSEYGCEVHQREGMKRQPGQLPRCKQSPLLQQAIAVPQGETNKGRVGLFISFCSLFLPKLVVGEACIEKVLGQVEVHLEQGLAEFTQTWISTGYLDHILQRTKHG
VELHPLVQGRGLQNKLDSFQAIKHGIYASYVRLLNFGGVNQLEYELYPQLSVIAMCVALTHGSTLIGVNLDDKYQALRSAHGAESKLRHCPEILEIKSLGLGKDKNILMEFYQQKNTISKQQQKEVLRAKQELLEKLKQQLA
>JAOEJA010000042_Phascolarctos_cinereus
AGLNTQTKYARARKLIFYVTIFSRVIDVTEAGVNLEGHLDHCLLTLLIYYYFEDDMEKYQDSPVAQYPSQRGYGVHQLGGDDTAGLGKFLEASNLSPLLQQAVPQGELNGGRIGPFISFCSLFLPKLAVGERGCIEKVLCQIRVHSKQGLPEFPQTSVSIATTQIIFTITQQSFLLEFVTI
HQGLHQPTRHDASDSIIATSAAQARFAGLIVKTVLGQVLHHTKPEAELHPLAGRSLQSGLDSLRRAVKEISKHRIYAPYARLESLWGVNQLEQLSVIAIGVTSAHRSTLIRVNLDERHQALRESAHEAESRPETLEVKSLGLEKEEEIFAEFHQQKNAISKNQQEEVVKAKQEFLKLKL
QLAPRLTGEQEKSPSLP
>JAQQSX010000002_1_Vombatus_ursinus
GLNTQTKYARARKLIFATKDLDLLCCRFTDVTAAGVNRGDHLDHCLLTLCMEKYQESPVAQHPSDHGYGDHQLRGDDTAGLGKLLDASNSSPRLQQAVPRGNTEARTGPFVSFCSLFLPKLAGGEQGCVEKVLTPNRVHSKQGLAEFAQTCVSIAAMKTIFTIRQQSFLLEFVIT
HQGLHQQTGHDAPDSITATSVAQECFVGLLIVKTVLGHVLHHTKPEAELHPLARGRSLQRELDSFRRAVKEISKHGIYAPYARLNLTGVNQLQQLSVMATGVASAHRSTPPRENLDEDAAHEAESRLKLCQEMLEVKSPGLEKEEEIFAEFHQQKNAICKNQQKEGVKAKQELR
SPRLTGEQEKSPSLPQQNTDLLGDSILRPKLELQLPKPSSAITFDPAVTSIKFPCTDIHSLIPTPLESQFSLQATGKEEGTSLSPDDLEEYFREGSSQPADIPEAQTNPAEREFEEDTDPPTVICPSRFLPRSETHISPNKVAPQAKTMPNKNQKPRAT
>JAOEJA010000517_Phascolarctos_cinereus
DNRSIEIYIIITAIFMDSLHDLLEVGLNIQTKYDRARKQIFYATKDLDLLCRIIDWIEAGVSLGDHLDHCLLTLLIYYYYEGEFNKYQDSPGDQYLSELGSSVHHLGGDDTAGLGRFLDASNRRPLLQQAIQEVPQGELNRERIGLFISFCSLFLLKLVVGEQACIEKVLPQIVHSEQGLAEF
PNWVSTATMKIIFTIKRSFLIKFVIIHHRLHLTGHDASNSVIAASVDQSVDFAGLLIVKTILDHILLVELHPLAELDSFQCKGVKEISKYGIYAPYARLLNLAGVNQLEHGLYPLLTAVAIGAASAHGSTLKQVNLDERYQALQESDHEAESRRRRREMLEVKSLGLEKKEEKIPSEFHQQ
KNAISKHQQKEVLKAKQELLEKLKLHLAPRLTGEQKKSPSLPNKTLIYSETTPSPNNSNSPEHQDTSPSDFHQITIPDIHSFTLIPSEGQPVLQTTDDEEEGTPLSTADLDEYFREGPSQLANIPGDQADPTNYKFEEDTDLTTGIKNPGGTLAFEVPPEPRNSHPHPTQTSSTVSTPSR
TEEPRPCQTGINNRNPKLHRGSVNTMRLENL
>JAOEJA010000643_1_Phascolarctos_cinereus
ENPNQFHYKTFNKHQNLHYNQAYNYGFTTLIGWVEYARARKLIFHDTQDIDPLCCRVTDVIEAGIDLGDHLDQFLTLLIFCYYEGDIKKYLESPVAQYLSKCGHEGHQFGGDDTAGLRFLAANNQNPPLQQAIQGVPHEELNRGRIGLFISFCSLFLPKLVVGERACIEKVQHQIRV
HSEQGLAEFPQTWVSTMIILTTMSGFLLKSVITHQGLHQHAGHTSDSMTATSVAQLHFAGLLIVKTILDHILHQTKGVEPVPEGGVSKVNLIAFWAVKEISKHGLYAPCARLLNLAMVNQMEHKLYPIISHSTSVMGVAASAHGRTLIRVNFDERYLALWESAHEAETKLSQHRE
ILEVKSLALEKEEELLAEFHQKNAISIHQQKEVLKTKQELLEKLKLPLAPRLTGEREKSPSLPQQYTNLLGDNDNPSQTRTPIDKAFKPFDIPVTSITLPSTDVHFFAPISLESQSTPQATGNEEEGTPLSPDYLDEYFRTSSPTPVNRYFRKSNRSHALMRRHQPNQRHWGSWRSLPN
PGRHKRPNNSTSPRPQTPQEPTSQVPIKLESTTETPSYLEALHQYNE
>JAQQSX010000004_1_Vombatus_ursinus
NLHYNQAYNYGFTTLTGWVEYARARKLIFHDTQDLDPLCSRVTDITEARVDLADHFNHGLVTLLIFYYYEGNIKKYLQSPVARYLLKHGHEAHQLGGDDTARLGKILDANNQNPLLQQAIQAVPQEESNRGRSGLFISFCSLSLPKLVVGERACIEKVRRLIRVYSEQGLAELPQTW
VSTMIVFTTTRQSFLLKFVIILQGLHQPAGHDASDSIIATSVARGRFAGLLIVKTVLDPILLRTKQGVELHPLAGRSLQSELDSFSMAVKEISKHGIYASCALLNLAKVNQLEHGLYPQFSIPLVSMGVAASAHGRTLIRVNLDERYLALWELAHEAESKLKQHREILEVKSLGLEKEEEIL
AEFHQQKNAISKHQQKCVLSRQERMAEAQPAVSLPRLTGGQEKSPSFPQQYTNLLGDNITHPKLELQLIKPSSAKPFNIPVTSITLPSTDVHFYHSHLKANPHPRQQAMRKKALPSPVYLDEYFRPPHPSQEIKKIPQTVNGKKPSTQSRTLGILGDPGGKLKRCSQTRTSHMSQQ
QHSPQLQTPLDQ
>JAJHZJ010000285_Bettongia_penicillata
KINIIITLMTTDSLHEIFEFELNIQIKYAVRNLIFIYMSDLDLLCHQIIDVIEAGISLRNHLDNCLLAICSYYYEGDKKVYDSPAVQYLSEDGYSLYQLSQDNAVNFSLPDANNQNPPLQTMQVVPQGELTKGRIGLFISISSLFLLKLNMGKHASIEKIMLQIWVHSKQGFAGFPQTWTL
TAIMKIIFTNMCQSFLFKFIIHQGIHQQVGDDTIVIATLVAQVHLTCLFIVKTVLDHILLQTKDGIELHPLVPGRSLQNELDSFCAVKEISRHRIYSPARLSRVNQLEHGLYPQLSAIPKGVAFAHGNTLIGVNLDERYQALRESVHKAENKFWHQEMLEVKQLKLGKEEEEILSEFHKK
EISKHQKREVAKAQQHLLDKLKVQLAPKLTAEHEKSPSLGQPKKSKCGKPQLTLDQNSSSDLQTP
>JAPYYB010000004_1_Lagorchestes_hirsutus
HEIFEFELNIQTKCAVRNLIFYTSKDLDLLCQQIIDVIEAGISLGNHLDNCLLTICNYDYEGNKKVLPSIQHLAEHSYSIHQLSQDNTVNFSLPDANNQNPPLQAMQTVPQGELTKGRIGLFISISSLFLLKLVVGKHASVEKVMLQIWVHSKQGFVGFPQTWTLTAIMKIIFTNMCQS
LFFKYIIHQGIHQQAGHDTIVIATLVAQAHLACLFIVKTVPDHILLQTKDGIELHPLAPGRSLQNELDSFCAVKEISRHRIYSPAKLLSLSGVNQLEHGRYPQLSAIPKGVAFAHESTLVGVNLDERYQALRQSVHKAENKLWYQKMLEVKQLKFGKEEEILSEFHKKEISKHQKREVAK
AQQYLFDKLKVQLAPRLTAEHEKSPSLGQQKNQSIRGNHNSPTRTPVDQTCKQQHIYP
>JAQMYV010000005_1_Macropus_fuliginosus
HEIFEFELNVQPKYAVRNLIFYTSKDLDLLCHQIIDVIEAGISLGNHLDNCLLTIYYDYEGDKKVYDSPAVHHLSEHGYSIHQLSQDNTVNFSLPDANNQNPPLQAIQAVPQGELTKGRIGLFISISSLFLLKLVVGKHASIEKVMLQIWVHSKQGFVGFPQTWTLTAIMKIIFTNMCQ
SLFFKYIIHQGIHQQAGHDTIVIATLVAQVHLACLFIVKTVPDHILLQTKDGIELHPLAPGRSLQNELGSFCAVKEISRHRIYSPAKLLSLSGVNQLEHGRYPQLSAIPKGVAFVHESTLIGVNLDERHQALRQSVHKAENKLWHQKMLEVKQLKFGKEEEILSEFHNRKRRISKHQKRE
VTKAQQHLFDKLKVQLAPRLTAEHEKSPSLGQQKNQSIRGNHNSPTRTPVDQTCKQQHIYLSVDITQATIHKFPTINTSISEKLFYSPSDWKGRRNPLNVIDLDEYFRGEPSQVGIPEESELTDSECEIDTGPFVDTEDAGSVQPNEIPPNRKILHNPQPKCFQRSQSPPGPTNPYFI
>JAQQTA010000005_1_Macropus_giganteus
HEIFEFELNVQPKYAVRNLIFYTSKDLDLLCHQIIDVIEAGISLGNHLDNCLLTIYYDYEGDKKVYDSPAVHHLSEHGYSIHQLSQDNTVNFSLPDANNQNPPLQAIQAVPQGELTKGRIGLFISISSLFLLKLVVGKHASIEKVMLQIWVHSKQGFVGFPQTWTSTAIMKIIFTNMCQ
SLFFKYIIHQGIHQQAGHDTIVIATLVAQVHLACLFIVKTVPDHILLQTKDGIELHPLAPGRSLQNELGSFCAVKEISRHRIYSPAKLLSLSGVNQLEHGRYPQLSAIPKGVAFVHESTLIGVNLDERYQALRQSVHKAENKLWHQKMLEVKQLKFGKEEEILSEFHKNEGLVNIRKGRL
QKLNNIYLINSKCSLPDQQNMRKAHPGNKKNQSIRGNQNSPTRTPVDQTCKQQHIYLSVDITQATIHKFPTINTSISEKLFYSPSDWKGRRNPLNVIDLDEYFRGEPSQVGIPEESELTDSECEIDTGPFVDTADAGSVQPNEIPPNRKILHNPQPKCFQRSQSPPGPTNPYFI
>JAQOLO010000057_1_Notamacropus_eugenii
HEIFEFELNIQPKYAVRNLIFYTSKDLDLLCHQIIDVLEAGISLRNHLDNCLLTICIHYYEGDKKVYDSPGVQHLSHGYSIHHLSQDNTVNFSLPDANNQNPPLQAMQAVPQGELTKGRIGIFISISSLFLLKLVMGKHASIEKVMLQIWVHSNQGFVGFPQTWTLTAIVKIIFTNMCQ
SLFFKYIIHQGIHQQARNDTIVIATLVAQVHLACLFIVKSVPDHILLQTKDGIELHPLAPGRSLQNELDSFCAVKEISRHRIYSPAKLLSLSGVNQLEHGRYPQLSAIPKGVAFIHESTLIGVNLDERYQALRQSVHIAENKLWHQKMLEVKQLKFGKKKEILSEFHKKEALVNIRKGRLQ
KLNNIYLINSKCSLPDQQNMRKAHPSNKKIRVLGETTTYPRPELQLIRPANSSTSTSLLTSLKPLSTNFQPYNTSISEKLFYSPSDWKGRRNPLNVIDLDECFRGDPSQVGIPEESKLTDSECEIDTDPFVDTEDTGSVQPTEIPPTKNLHNPQPKCFQRSQSPPGPTNPYFI
>JAQQSK010000003_2_Potorous_gilbertii
QKICSKFDFFYTSKDLDLLCHQITDVIEAGINLGNHLDNHLLTICNYHYEGDKKVYDSPVVQYLPEHGYSIHQLSQDNTVNFSLPTLQAMQAVPQGDLTEGRIGLFISISSLFLLKFVVGKHASIEKVMLQIWVHPKQGFVGFSQTWTLTAIMKIIFTNMCQSFLLKFIIHQIHQQAG
HDTTIIATLVIQVHLACLFIVKTVLDHILLQTKDGIELYPLAPGRSLQNELDSFCAIKEISRHGIYSPARLLSLSGVNQLEHGLRPQLSVIPKGAAFAHESILIGVNLDERHQALRESVHKAENKLWHQEMLEVKFKLGKEEEKILSEFHKKEILVNIRKGRLQKL
>WOXC01004209_Gymnobelideus_leadbeateri
KKYHDSPMVQLLSEHEYAVHQLGGVDNTSLHKFLYASNHNPLPQQTIQTIPQGELNGRTGLFLQKLVVGETCIEKVLQIQVNSEQGIAEFPQTWLSTAIFTIMRRNFLLQFVIIHYILYTIISTPSKSWAGCNRFKIATSVAQAHFAILLIVKIVLDHMLLCTKHGVELYPARRSVQSKL
HSCHVAKEISKHGIHGPCAFMNLAGVNQLEHGLYLQLPAIAIGVASVPESTLTGVNLDKRYQVFRESTHEVESKLRQHETLEVKSLRLDKKEILTDFHQKNAISKHQQKKVLKTKQELEKFKLQLAPRLAGEQDKSTSL
>JAMXIF010001117_Petaurus_breviceps
ILMNCMHYSHIKKYHDSPMVQHLSEHEYAVHHLVRVDTTSLDKFLYASNQNSVAQQTIQAVPQRELNRGRISLFISFYSLFVPKLVVGEAYIEKVLCQIQINSEQGLAEFPQTWLSTATRKIIFKMMQQNFLLQFVIIHGLHQQARQGATDIIATSVAKACFALLLIAKIVLDHMLLC
TKHGVEFYPCGQGQSKLHSCCHVAKEISKHGIHGPCAFMNLAGVNQLEHGLNPQLPAIAIGVASNLGNTLIGVNLSKRYQVLQESAHETESKLRQLEILEVKSLRLEKEEILIDFHQKNAISKHQQKEVLKAKQKLLDKL
>JAQQSW010000008_Pseudocheirus_occidentalis
ILMNCMHHSHIKKYHENPMVQYLSEHTYAVHQLGGVDTTSLDKFLYASNQNTLPQQAIQAVPQGELNRGRIGLSIFYSLFLPKLIVGKQACIEKVLCQIQVNSEQCLAEFPQTWISTATRKIIFTMMLQNFLLQFLFVHGLHQQTGHGATDTIITTSVAQACFAILFIVKIVLDHML
LCTKHRVELYSLAQGKSLQSKLHSCHVVKEISKHGIHGPHAFINLAGINQLEHGLYHQLSAIGVASAHRSTLIGVNCRETSHERYQVLQESAHETESKLRQYEMLEIKSLRLEKEKEIFTEFHQQKNQISKHQQKEVLKAKQELLEKLKSQLAPT
>JAQQSV010000007_Pseudochirops_corinnae
ENPMVWYLSEHEYAVHQLGGVDTTSLNKFLYASNQSPLPQAVPQGELNRGRIGLFISFYSFFLPKLVVGKQACIEKVLCQIQVHSEQGLAEFPQTWISTATRKIISAMMQQNFLLQFVIIHGLHQQTGHGATDSIIATSVVQECFAILFIVKIVLDHMLLCTTHGVELFSLAQGKSLR
KLHSCHVVNEISKHGIHGPRAFMNIAGVNQLEHELYPPAITIGVALAHRSTLIGVNLDERYQVLQESVHETESKLRQHEMLEVKSLRLKKEEEILTEFHQQKNAIRKHQQKEVLKAKQELLEKLKLQLGTRLAGEQNPSLAQQNIDLFGDNTVHPKPGTLANKTFEQDMIPSDISQI
TTHKYSLFYFHTFGKLNPPYDYVMRRKANPVLILVNISRKAPLSRHFRRSNTHGLMSGHLHSHGYGPPRCCAFEVPPEPRNSHTSQQQYSPQLHSPQESMSHQTGINNRNSKVHRGTSPIQEETSVSNYLQNPLVFPDMPISKE
>JAQQSU010004289_Pseudochirops_cupreus

ENPMVQYLSEHEYAVHQLGGVDSTSLNKFLYASNQSPLPQAVPQGELNRGRIGLLISFYSLFLPKLVVGKQACIEKVLCQIQVHSEQGLAEFPQTWISTATRKIIFPMMQQNFLLQFVIIHGLHQQTGHGATDSIIATSVAQECFAILFIIKIVLDHTLLCTKHGVELFSLAQGKSLRKL
HSCHVVNEISKHGIHGPHAFMNIAGVNQLEHGLYPPAITIGVALAHRSTLIGVNLDKRYQVLQELAHETENKLRQHEMLEELRLKKEEEILTEFHQQKNAIRKHQQKEVLKAKQELLEKLKLQLGTRLAGEQNPSLAQQNIDLFGDNTVHPKPEPLANKTFEQDMIPSDISQITIHR
YSLFYFHTFGKPIYPMTNKEGRQTHESSSWIFQGRPLSVSRHFRRNTHGLMSGHLHNHGQGPPRCYAFEVPPEPRNSHTSQQRYSPQLHSPQESMSHQTGINNRKSKV
>JAQQSZ010000005_Phalanger_gymnotis
RGFRQVRKLTFNATTDLDLLCCQIINLITADVNLGDHLDNCLTLLINYYYDGNIKNYNESPVAQYLSEHRYSIYHLNQDDTANLSQFLDVISHSPLQKAIQAVPQREHTKRRIGLFVAFCNLFLPKIVGEHARIEKMMRQIRVHSEQGLTEYPQTWASRGTMKIIFTIMQQPFLLKFFF
IIHQGLYQLAGHNAGELMITTVAQACFAGELIVKMLLDHILLHTSDGVELHPLAQGRGLRNELNGFQCAVKKISRSEIYSSYARSLNLAQVNQLKHRLYPLSATAIGVVSIHGEHTLIGVNLDETYQALRESARKAKSKLQRCEMLEVKLAKEDEEIISEFHQQKDEISICQQKEFSKA
RQDILEKLEVQLVLEPTGEEESKPFLGLTKLLPSATLW
>JAJHZJ010000112_Bettongia_penicillata
HDTLEAGLNIRMEYVLVRQLNYATKDLHLLCYQVINLIKAGIDLDNFLLTLSITYFYGDIEKYNESPVTPYLSELRYTIHHFSQDNRAKLSQFLEVIITIARTIAVQAVPLRELTKRSIDLFVTFCSLFLLNIVVGQHAYSEKVKHQMIHSEGLIELPQTWASTATMKIIFTIMWQSFPLKIIII
HQGLYQEASHNARSIITTLAQACFAGLFIIKIMLDSILLHSTDETEIHSSRQGRTLQNEVFSFFTIKEISRGIYSTYARLLNFAGANQLKHNLYEQLGNAIDVASTHGSILIGVNLDKKYQELRESACEAESKLWFELLKVKNFKLSKEDEDIIVFHQQKDEITIYQQNEILKLKKDLLEKFKE
QLAPKLIGQQRIPSSVQQITELPGESMGNP
>JAQQSK010000003_3_Potorous_gilbertii
HDTLEVGLNIRMEYVLVRQLNATKDLHLLCYQVINMIKAGYNLDNCLLTLSITYFYGDIEKHNKSPVTPYISEFRYSIHHFSQDNRANLSQFLEVIDHSCIHNSVQAVPQRELTKRRTDLFVTFCSLFLLNIAVGEHAYSKKVRHQIIHSEDLIELPQTLTSTATMKIIFTIMCQSFPLRIIII
HQGLYQEESHNVRYSIITTLTQVCFVLFIIKIMLGNILLHSTGETEIHSPGHGRTFQNEVGGFCTIKEIFGICSTYARLLNFAGVNQLEYNLYPQLVNAIGCATHGSTLIGKNLDKKYHVLRESACEAESKFWFELLEVKNFKLSKEEKILYVFHQQKDEINIYQQNEVLKLKKDLLEKLKG
QLA
>JAPYYB010000001_1_Lagorchestes_hirsutus
YGDIEKHNESPVTPYLSEHRYSIHHFSQDNRAKLSQFLEVNNHNCTHNSVQAVPQGELTKRRIDFFVTFCGLFLLKIVVGEHVYSEKVMNIIHSEQILIEFPQTWTSTATMKIIFTKMQNFPLKIIIIHQGLYQEASHNAIYSIITTLVQACFAGLFIIKIMLDHILLHSTERQMHYSGHG
RTLRNQVDGFQCTIKEISRCGYIPHARLLNFAGLNQLEHELYPQLSVNAIDVASTHGSTLIGVNLDEKYQVLKKSACEAESKLWFELLDVKIFKLSKEDEDIISVFHQKDEINIYSFKKKKIYLKNLRANSPPNSGNNKESHPWCNKELSYQGRLWLIPDQNYNLDQALKHLLLHSDHH
PLIFHQYNLYS
>JAQMYV010000006_Macropus_fuliginosus
YGDIEKHNESPVTPYLSEHRYSIHHFSQDNRAKLSQFLEVNITIAVQAVPQGELTKRRIDFFVTFCGLFLLKTVVGEHVYSEKVMHIIHSEQVLIEFPQTWTSTATMKIIFTIMWQSFPLKIIIIHQGLYQEASHNATYSIITTLVQACFAGLFIIKIMLDHILLHSTDETEIHYSGHGRTLR
NQVDGFQCTIKEISRCGYIPHARLLNFAGLNQLEHELYPQLSVNAIDVASTYGSTLIGVNLDEKYQVLRKSAYEAESKLWFELLDVKNFKLSKEDEDIISVFHQKDEINIYQQNEVSKNKNKKIYLKNLRANSPPNSDNNKESHPWYNKELSYQGRL
>JAQQTA010000006_Macropus_giganteus
YGDIEKHNERPVTPYLSEHRYSIHHFSQDNRAKLSQFLEVNITIAVQAVPQGELTKRRIDFFVTFCGLFLLKTVVGEHVYSEKVMHYIIHSEQVLIEFPQTWTSTATMKIIFTIMWQSFPLKIIIIHQGLYQEASHNATYSIITTLVQACFAGLFIIKIMLDHILLHSTDETEIHYSGHGRTL
RNHVGGFQCTVKEISRCGYIPHARLLNFAGLNQLEHELYPQLSVNAIDVASTYGSTLIGVNLDEKYQVLRKSACEAESKLWFELLDVKNFKLSKEDEDIISVFHQKDEINIYQQNEVSKNKKKRFT
>JAQOLP010000809_Notamacropus_eugenii
YGDIGKHNESPVTPYLSEHRYSIHHFSQDNRAKLSQFLEVNITIAVQAFPQRELTKRRIDFFVTFCSLFLLKTVVGEHVYSEKVMHIIHSEQVLIEFPQTWTSTATMKIIFTIMWQSFPLKIIIIHQGLYQEASHNATYSIITTLVQACFAGLFILKIMLDHILLHSTDETEIHYSGHGRTLR
NQVDGFCTIKEISRCGYIPHARLLNFAGLNQLEHELYPQLSVNAIDVASTYGSTLIGVNLDEKYQVLKKSACEAESKLWFELLDVKNFKLSKEDEDIISVFHQKDEINIYQQNEVSKNKKKRFTKTGL
>WOXC01006732_Gymnobelideus_leadbeateri
KNLDLQCCQILNMIKADVNLADHLDNFFLLTLLIHYHYEDDIKKYDENPYLSEHRYSICHLSQDDKANLSQFLDVINSQAPLAIQSVPQGEFTKSRTDLFVAFYILFLLKIMEGEHTCIKKTMHQIVHSEKGLTLFPQTWISTTSKKSYSQHSVAQIEHQLWIQGDLSSNPTSDTPLTHC
AMLDKLNPNCLDFIPPKIKNKIIYVYFKNRKKILFTIMWQCFSLKFILIHQGICQQVDHNAADTITTTLGPQACFAGLLIIKTMLDHILLHIIDGVELHCSACGRDFQNEIDAFQCAIKEIPRQGIYSPYARLLNLARINQLEYGLYTLSATTIGVPSPHGSTFIGINLDERYQAFRDSAHEA
ESKLWCEMLEVSHLKLEKEDEIIEFHQQKDEINIPQKKEISKARDLKKLKVHLAPKTHRRT
>JAQQSW010088965_Pseudocheirus_occidentalis
QDDTANLSQFLDVNYYNPPLDAIQSISGQLTKRRMNLFVAFCNLFLLKIIEGEHSCIEKVMHQIVYSGLTLIPQTWTSINHYQIIFTGQLGGIVNIAALESEGSGFKPDLRQLAINYLCDLGQVTTPIVLPSLLQKNKIIHVYFKKKSHSPSCGSFPLNFILIHQDLHQQLDHNAADTIITT
LVAQDCFSSLLFIKNMLDNILFHIIYGVELHQAACGRAFQNELDGFQCTIKEISRQGIDSPYARFLNMARVNPLEHKLHPLSATIVGVPSPHGSICIGINLDDRYQALRKSAHEAESKLLFEMLEIKHLKLAKEDKEVISEFHQQKDEINTHQQKEISKARD
>JAQQSV010074762_Pseudochirops_corinnae
VGNRKLIFYATKDLDLQCCQVLTMIKAHVNLADHLDNFLLTLLINYHYEDDIKKCNESLYLSEHTYSIYHLSQDDTANLSQFLDVNNCNPPPHYKRTIQSISRQIIKRRIDLFVAFCSLFLLKIIEGEHTCIEKVMHQIVYSEGLTFNPQTWTSINHYKIIFTGQLGGTVNRALALESEGPG
FKSDLRHLAINLCDLGQVTTSIALPSPLQNNKIIYVYFLKKKENHIHIMWQSFPLNFIRIHQGLHQLVDYNVEDTIITTSVSEARFAGLLIIKNMLDNILHIIYGVELHHSVCGRAFQNELDGFQCAIKEISKQGIDSLYARFLNMSRVNQLERKLYPLSATAVGVPSLHGSILMGIILEERY
QAFRELAHEAESKLWCEMRRRKHLKLAKEDKEIIEFHQQKDEINIHQKESSKARDLLEKTLSLIGPKTHRRT
>JAQQSU010020571_Pseudochirops_cupreus
NRKLIFYAAKDLDLQCCQLLTMIKAHVNLADHLDNFLLTLLINYHYDNDMKKCNESPYLSEHRYSIYHLSQYDTQTLVNFWMIIATPPPHHYKRTIQSISRQIIKRRIDLFVPFCSLFLLKIIEGEHTCIKKVMHQIVYSEQGLTFIPQTWTSTSTTKSFLQDSVVQIEHWLWSQKDLGS
NLTSDTWQLASCVTLDKSQPQLHCLHPSKNKIKEYKYIFKKKENHIHIMWQSFPNFILIHQGLHQLVDYNAEDTIITTSVSEAHFAGLLIIKNMLDNILLHIIYGVELHHSVCGKAFQNELDGFQCAIKEISRQRIDSLYARLLNMSRVNQLDHKLYLQLSATAVGMPSLHGSILMGIN
LDERHQALRELAHEAESKLCCEMVEVNHLKLAKEDKEIISEFHQQKDEINIHQKESSKARDLLEKTSIIGPKTHRRT
>POVN02000726_Phascolarctos_cinereus
HDLLEVGLNIQIKYARARKLIFYATKDLSLLRCVIDVIEAGVNMGDHLDHFLLTLLIYYYYYEKDIKKYHDSLVAQYLIEHEYAVPQLSGDYTVRWESFLMAIKTPCCHRLFKLFHKGNSELGLFFSFCSLFFPKLVVGEQACIERVLRQVQIHTEQGLAEYLQDLKTEPQTWVSTVTMKI
VFTMMQSFLLKFVIMHQELHQAGHDASIIATSVAQACFANLLIVKTVLDHILLRTKNEVEFHPSAGRGFQSELDGFQHTIKEISKHGIYAPYAQLPNLAGVNQLEHELYLQLSAIAIGVDLAHASTLTGMNLDKRYQDLECRVWAHEVEQVKTSSNVRSEEEEILTEFHQQKDAISK
HQQKEVLKAKQEILDKIT
>JAQQSX010000003_2_Vombatus_ursinus
HDLLEVGLNYIKYTRARKLIFYPIKNLLCCVIDVIEAGVNMGDHLDHFLLTLLIYYYYKGDIKKYHDSLVAQHLAQHEYRSPPIRGHHGESFLMAIKIPCCNRLFKLFSKGNRKNVFISFCSLFVPKLVAGEQACIEKVLRQVRVHSEQGLARFLQNSAKSQNMRPTTWVSTVTMKIVF
TMQSFLLKFVTMHQKLHQAGHDASDIIATSVVQAHFTSLLIVKTVLDHILLRTKNKVEFQPLAWGRSFQSELDCFQHTIKEISKHGVHAPYAPLPNLAGVNLEHELYPQLAIAVDVALAHGSTLIRVNLDKRYQDLECRVWAHEVENKLRQHGEEEKILREFHQKNAISKLQQKEV
LKAKQEILDKI
>JAJHZJ010000220_Bettongia_penicillata
QLWIHYMTYLRSDTFKILISLTCARKLIFYTTKNLDLLCSVIVTIDARIRKDLGNHLDHCLLTSIYYYYEGDINKYHDSPVAQYLSEHGHTVHPLGGGDIAGLGKCLGANNQNFLLQAMQVVPQGELSRERIGLFISFCSLFLPKLVAGEDWTENVLQIWVHCEGLAEFLQTWISTSA
MKIIFTMIWQNFLFKFVITHQGLHQQAGSDAADSIIVTSVAEACFTRLFIVKFLGSYAIARNIGLRFKPLTQGRSFKSEIEFKVKDISKHIYAPDVQPLNLAGVSLGHELYLQPLPIASGFASANGSIFTEVNLDERYQALQESAHEAESKLRQYLEMLDIKYLGLKKGEKILTEFHQENAI
RKHQQKEVLKTKQELLKKLKLLAPRLAGEQDKNPSLAQENIDLLRDNDNPSKNNSSGDHQVPNHLTPQLLSNYHSWTFILLLPYLGKPIHALSNRQKGRHTIESSQSWIFQGRPLSISRRSRRSNTQRLMWRGHPGHGHGPYGSQPCGVLPEPRNSRAPKQHSPPQSSQDPMS
EDTNKTGINNRNPQRHRGPSPVLD
>JAPYYB010000001_Lagorchestes_hirsutus
HDLLEVLSIQNKHARSRITDFYTTKYLDLLYGVIDTIEARIRKDLGNHLDHCLLALSIYYYYEGNINKYRDSAVAQYLSEHWHTVHQLGGDDITGLGKFPGASNQNFLLQATQVVPQVKISRERISLFISFCSLFLTKLVVGEDWTENVLFQIWVHCEGLAEFPHTWISAAAMKIIFTM
MWQNFLFKFVITYQGLHQQAGHDAADSIIATSVVQACFTRLLIVTLLDHILLTKYGVEVTLDPRKESQKQTEFKVKDILKHIYAPDVQALNLAGVNHLEHELYLLSLAIAIGFASAHGSTFTEVNLDERYQALRESAHEAKSKLRQCLEMLDIKYLGLEKEEEILTEFHQEKNAISKHQQ
KEVLKVKQELFEKLKLQLAPLAGEQDNSSLAQENIDLLGHNDNPSQTTAPANKTIKCQIILPSDFSQITIHGYSFFYFHIMGSQSMLVIDNEKEGTPLSLVSFDGYFRKSPSQLADIPGRSNTQRLLMWRGHPGHGHGPWRYTTLWGSSTQDSHVPKQHSPPRSSQDPMSQDTIK
LESTVETPSYIEALHQYHE
>JAQMYV010029555_Macropus_fuliginosus
HDLLEVLNIQNKHARARITDFYTTKDLDLLCRVIDTIEARIRKDLGNHLDHCLLALSIYYYYEGNINKYHGSAVAQYLSEHKHTVHQLGGDDITGLGKFLGASNQNFLLQAMQVVPQEELSRERIGLFISFCRLFLTKLIVGEDQTENVLCQIWVHCEGLAEFPQTWISAAAMKIIFT
MMWQNFLFKFVVTYQGLHQQAGHDAADSIIATSVVQACFTRLLIVKLLDHILLTKGLRFKPLTQGRSLKSKLNEFKVKDILKHIYAPDIQPLNLVGVNHLGHELYLQSLAIAIGFVSAHGSTFTEVNLNEKYQALRESAREAKSKLRQCLEMLDIKYFGLEKEEEILTEFHQENNAISK
HQQKEVLKVKQELLEKLKLQLTPLAGEQDNPFSGPRKYLTWTQYNPSQTTAPANKIIKCQIILPSDFSQITIHGYSFFYFHIMGSQSMLVIDNEKEGTPLSLVSFDGYFRESPSQLADIPEDQTESK
>JAQQTA010000127_Macropus_giganteus
EIYIIIKCTVVDMLHDLLEVLNIQNKHARARITDFYTTKDLDLLCRVIDTIEARIRKDLGNHLDHCLLALSIYYYYEGNINKYHGSAVAQYLSEHEHTVHQLGGDDITGLGKFLGASNQNFLLQAMQVVPQEELSRERIGLFISFCSLFLTKLIVGEDQTENVLCQIWVHCEGLEEFPQT
WISAAAMKIIFTMMWQNFLFKFVVTYQGLHQQAGHDAADSIIATSVVQACFTRLLIVKLLESYPIREQSIGLRFKPLTQGRSLKSKLNEFKVKDILKHIYAPDIQPLNLAGVNHLGHELYLQSLAIAFGFVSAHGSTFTEVNLNEKYQALRESAREAKSKLRQCLEMLDIKYFGLEKEE
EILTEFHQENNAISKHQQKEVLKVKQELLEKLKLQLTPLAGEQDNPSVAQENIDLLGHNDNPSQTTAPANKIIKCQIILPSDFSQITIHGYSFFYFHIMGSQSMLVIDNEKEGTPLSLVSFDGYFRESPSQLADIPEDQTESKDCGCGEDTDEAMGIRDPGGTPPYGAPPDPRNSHAP
EQHSAPQSSQDPMSHATIKLESTVETPSYIEALHQYSE
>JAQOLO010000065_Notamacropus_eugenii
HDLLEVLNIQNKYARARITDFYTTKDLDLLCGVTDTIEARIRKDLGNHLDHCLLALSIYYYYEGNINKYHDSPVAQYLSEHQHTVHQLGGDDITGLGKFPGASNQNFLLQAMQVVPQVELSRERISLFISFCSLFLTKLVVGEDWNENVLYQIWVHCEGLAEFPQTWISAAAMKIIFT
MMWQNFLFKFVITYQGLHPQAGHDAANSVIATSVVQACFTRLLIVKLLDHILLCTKGLRFKPLTQGRSLKSKLNEFKVKDILKHIYAPDVQPLNLAGVNHLGHELYLQLLAIAIGFASAHGSTFTEVNFDERYQALRKSAHEAKSKLRQCLEMLDIKYLGLEKEEEILTEFHQEKNAIS
KHQQKEVLKVKQELVEKLKLQLAPRLAGEQDNPSLAKENIDLLGHNDNPSQTTAPANKTIKCQIILPSDFSQITIYGYSFFYFRIMGSQSMLVIDNEKEGTPLSLVSFDGYFRESPSQLADIPEDQTEPKDCECGEDTDQAMGIRDPGGTPPYGAPPEPRKSHAPKQHSPPQSSQD
PMSQDTIKLESTVETPSYIEALHQYSE
>JAQQSK010000003_1_Potorous_gilbertii
ISLICARKLIFYTTKDLDLLCGIIDIIDARIRKDLGNYPGHCLLILSIYYYYEGDINKYHDSPVAQYLSEHGHTVHQLGGGDIAGLGKCLGANNQNFLLQAMQVVPQGELSRERIGLFISFCSLFLPKLVVGKDWTESVLCQIWVHCEGLPEFLQTWISTAAMKIIFTMMWQNFLFKFVI
IHQGLHQQAGHDAADSIIITSVAQACFIRLLIVKLLGSCPIVHKIGSLNPPKEGISKQIEFKVKDILKHIYAPNVQPLNLAGVNHLGHELYLQSLPIAIGFASAHGSTCTEVNLDERYQALRESAHEAKSKLRQCLEMLDIKYLGLKKEEEILTEFYQEKNAISKHQPKEVLKAKQELLKKLK
LLAPLAGEQDENPSMTQENIDLLGDNDNPSQTTVTANKTIKCQIILPSDFSQITIHRYSFFYFHILGSQSMLVMDNEKEGTPWSPVNFDGYFMESPSQLAGHSRRSNTQRLMWRGHPGHGHWGPWRHITLWGSPEPRNSQAPKQHSPPESSDPMSQDTNKLESTVETPSYI
EALPPIQDWEISVSNYLEKPPGIPHAYFRTLEFIQLQMVSQQFCTNYYGALFRMELSNNLRSFYERPLEYLIKNKKERKSETKIHSYCNPEGRKDSGCNCEPSSLLSFPW
>JAANDE010000004_Trichosurus_vulpecula
RKLIFYATKDLDLLCGKVVDVIEARVNLGDHLDHCLLTLWVYYYYEGDIKKYHDSAVTQYLSEHGYAVQQFGGDDTAGLGKFLDASNQNSLSQQVVQAVPQGEQSRGRVGLFIPFSSPKIGCSMSLIEKVLCQIWVHPKQALAEFPQTWVSTATMKIIFTTMWNFLLFVIIHQGF
YQQAGHSAADSIIATSVGQECFVKSVLDYMLWHTKYGVEIYPLVGRSLSKLDSFQYLVKGISKHGIYARYVQLLNLAAVNQLESGLCPQLLAIAIGVASAHRSSLIGVNLDEKYQPLQESVHETESKLRQCLEMLKFKCLGLKKEEEILTQFHQQKNTINKYQQKEELKAEKLLEKLKL
QAPRLSGEQDKSPSLAQQNIDLVEDNPTHPKLELQLIKPSEGQYSGVHSDFSQIISHGYSIFYSHAFGKPTHPPSNRQGGRYTLGSSSIFQGSPLLADIPEDETKLIHCECEEDTDMGIWDPGGAPPFKVLPVPMNLHTPPKQYSPHPQSPEPM
>JAOEJA010000168_Phascolarctos_cinereus
HDLLEVGLDTQQHTRARARELIFYPTKELDQSWCWVLDVIEAGVSLGDYLDSLLMDFICYSYGSDIDKYQVAQYLSAHRLGGDDTSGLGKYLDANNQSLQLQRAAQAVPRGETGIGLVISFCSLFLPKSVVGEACFEKVLCQVVCWEQGLAEFPQSWASTVTTKIIFTQMRQSFL
LKSVLVLHQAGRDAWDSIIATICFVKTVPDHFLSRGTKLHPLAWGGGLQSALDRFPGAVKDISRHGMYAPYAHLLNLARVNQLEYGLYLQSSAIVIGVASAHGSTLLGVNLDDRYQTESAHEAESKLRLPEMLEVKSLGLKKILVEFHEQKNAVSKRQQK
>JAQQSX010000006_Vombatus_ursinus
RKFWVSIKIYIVINHVMDSLHDLLEVGLDTQQNTRARELIFYATKDLDQLRWVLDVIEAGVNLGDYLDHCLPTLCCYSCEGDIDKYRDSPVVQYLSKHGYSVHQLGEDDTSGLGKYLDANYLSPQLQQAIQAVPQGETGIGLFISFYSLFLPKLVVGEACIERVLCQVQVHSGLPEFP
QTWVSSITMKLTFTMISFLLKFVIMHQGLQQQAGRDASDSIIAKFAGLLIVKIVLDHILSRTKQGVELHQLARGRGLQSELDCFPGAVEETSRHGIYALLSLAGVNQLECGLYPQWSAISIGVASAHGSTLMGCESQIPTKSAQEAESKLRLLGKCYKLRVLGEKKILVEFHQKNAVS
KYPQKEGGTGITEAPRTMEEKVLLYMRKTLTYSETIHPILSNFNSLVQGYLAPQLQSNRLQTFNPSMPFPQKAYPFFQQAYEEEGTPLSPADLDEDLERRPQPERSSRKSDRTYRWRVRGQRLNCRHECSLRSFPNAATYTLCVGHSALPLTNPRRDEP
>WOXC01009333_Gymnobelideus_leadbeateri
LRTLTVGQAYIEKALHKDQVHIEQGVAEFPPTWVSTASMKITLSIIHQCFLFKIFIIHQGLHQLGGNGISDTIIATSVAQACFPGLLIVKTVLDYTLQCTSYGVELHFLAQDRGMESELDYFQAIKEISRHGINASDVWLLNLNGVNQLQNGLFSKINSPLGVASAHGNTLIGLNLDIRH
GELLESAHEAETKLRYHLEMEEVSFGLGKDKKIYIIMEFHQKRTTSRQQQK
>JAMXIF010000043_Petaurus_breviceps
HDLLEVGLNITKYARELKIIFYAAKDLNHLCCRIINMLRAGISIGDNLDHCLLTLLTYYYYERTYKKFDSPVAQYLSEHGFEVHQLGGDDIANLHMDLDPTIKACNCSKKSRLSCPNVKWEIYSRRISLFVLFCSLFLQKLVVGEQAYIEKALHQDQVHKEQGVAEFPQTWVSNASMKI
ALSMIHQCFLFKFVIILQGLHQHDGNGIYDTITDTSVAQVCFAGLLIVKTVLDHTLQRTSHGVELHFLAQDRGMQSELHYFQAMKEISKHGINTSDAWLLNLNGDNQLQHGLFSKMSAFATGVASAHGNTLIGLNLDIRYQALLASAHEAESKLRYHLEMQEVKSFGLEERKKYII
MEFHQKSTTSRQQQK
>JAQQSW010086624_Pseudocheirus_occidentalis
LRRLRTTDIEQRVSQFPQTCVSNASVKITFSMIHQSILFKFVIIYQGLHQYSGNGISDTIIATSVAQACFAGLLIVETVLDHTLQCTSHGVKLYFLAQGTGIQSGLDYIQWAIKKISKHGIKASDAWLLNLTGVNQLQHGLFPKLSAIAIGFASVHGNTPIVVNLDRTYQALLQSAHGAE
SKLGYRPEMQEVKSFGLWKDEKYIMMEFHWQKSTISRQQQKEEGKAKQELLEKL
>JAQQSV010000003_Pseudochirops_corinnae
YYVVITCEIGLNVTKYAKAKLIFYAAKYLNHLCCRIINMFKGGINVGDNPDHCLLTLLICYCYEGTYKKFDSPVAQYPLEHGFEVHLLSEDDIANLGMDLDANNESLLQQEIQAVLQGEIYSGRINLFILFCSLFLPKLVVGEQAYIEKVLHQDQVHMEGVSEFPQTWVSIASMKITFS
MIHQSFLFKFVIIHQGLHQHSGNGISDTIIAASVAQTCFAGLLIVKTVLDHILQHTSQGAELYFLAQGRGIQSDLDYFQWAIKEVSKHGINASNVWLLNLTGVNQLQHGLFPKLSTIAILGVASAHGNTLIGVNLDRRYQALLESAHEAESKLRYHPEMQEVKSSGLGKDEKYIIMEF
HQKSTITKQHLKEELKAKQELLEKLKLQLGPEVIEEQEGNFLLPQKNVSLQQQQDPEHSVHRCVDTIYQQKITPDLFGGTEEGLYGFAHITWVTGLGGNPRARSTSIAEFVEAAEGRPPSQSTGQKRGSVVTHRPEHRQKRSKPV
>JAQQSU010000001_Pseudochirops_cupreus

LPHNLCYYVVITCEIGLNVTKYAKAKLIFYAAKYLNHLCCRIINMFKGGINVGDNPDHCLLTLLICYCYEGTYKKFDSPVAQYLSEHGSEVHLLSEDDIANLGMDLDANNQSLLQQEIWAVLQGEIYSGRINLFILFCSLFLPKLVVGEQAYIEKVLHQDQVHMEWKITFSMIHQSFLF
KFLNIIHQGLHQHSGNGISDTIIAASVAQTCFAGLLIVKTVLDHILQHTIQGAELYFLAQGRGISELDYFQWAIKEVSKHEINASNILLNLTGVNQLQHGLFPKLSTIAILGVASAHGNTLTGVNLDRRYQALLESAHEAESKLRYHPEMQEVKSSGLGKDEKYIIMEFHQKSTTTKQH
HKEELKAKQELLEKLKLLLGPEIIEEQEGSLLLPQKNVSLQQQQDPEHSVHRCVDTIYQQKITGISLVALRKDSIVRVFDVISHWGCTTTAWGRGRGPRMSQRRHDHADVSLSSGRYVSGSPGDDGTLRRESKARVKGPLTCSSPPFLYTLQTSLPFARVLGALWGVVLQASLEAG
DNRSLRGGKQLEEVSNRIGSPESKLGDALQLAPQTGILFRXXXXXXXXXXXXXXXXXXXXXXXXXXXXXXXXXXXXXXX
>JAPYYB010000003_Lagorchestes_hirsutus
LEAELNMQTKYTKARKLIFYATKDLNSLCCMIVGMVEAGINLGDNLNHCLLTALICYYYESNIKNFDSPVAQHLSEDKYVVHQLGRDDTDNLDMYLDVNCQSPQLQQAITVLQREIRSRRIGLFILFCSLLFSKLVVGEGTTLTKFYIKRRSLKQGLVEFPKTVSAAAMKIIFTMMYP
SFLLKFVVIHHGLHQAGHGPCETLITIISCSGRFAGLLIGKTVLDHILQCTSHVIELHLLFSRGMQSELECFQWSVKEMIKHEIYTPWLLDLAEDNQLEHGLFSKLSATAISVASVHGSTLTGVNLDNRYQALLESAQEKLRHCAEMQEVKSLGPGKDDKNIIMQFYQKSNISNNKKN
>JAQMYV010000003_Macropus_fuliginosus
KARKLIFYATKDLNSLCCMIIGMIEAGINLGDNLNHCLLTALICYSYESNIKKFDSPGAQHLSEDKYVVHQLGRDDTDNLDIYLDVNCQSPQLQQAITVLQRKIHRRRIGLFILFCSHLFSKLVVGEGTTLTKFYIKRRSLKQGLVEFPKTVSAAAMKIIFTIMYPSFLLKFVIIHHELHQQ
AGHAPCETIITIISCSGRFAGLLIGKTVLDHILQCTSHVIELHLLFGRSIQSELEYFLWSVKEMIKHGIYTPWLLDLAEDNQLEHGLFSKLSATALSVASVHGSTLTGVNLDNRYQALLESAQEKLRHCAEMQEVKSLGPGKDDKNIIMQFYQKSNIS
>JAQQTA010000004_Macropus_giganteus
KARKLIFYATKDLNSLCCMIIGMAEAGINLGDNLNHCLLTALICYSYESNIKKFDSPGAQHLSEDKYVVHQLGRDDTDNFDIYLDVNCQSPQLQQAITVLQRKIHSRRIGLFILFCSLLFSKLVVGEGTTLTKFYIKRRSLKQGLVEFPKTVSAAAMKIIFTIMYPSFLLKFVIIHHELHQQ
AGHAPCEIIITIISCSGRFAGLLIGKTVLDHILQCTSHVIELHLLFGRSIQSELECFLWSVKEMIKHGIYTPWLLDLAEDNQLEHGLFSKLSATAISVASVHGSTLTGVNLDNRYQALLDQALLESAQEKLRHCAEMQEVKSLGPGKDDKNIIMQFYQKSNIS
>JAQQSW010000005_1_Pseudocheirus_occidentalis
LNQSPQLQQEIKTVPQGEMHSRVILFILFCGLFLPKLAVGEQVCIEKILRQVHVHMEQGLVKPQTWVSTMKIIFTMMCVKAFYLIHQGLDQQTGHHASDRIIVTSVAQAPFVGLIIKTVLDHVWQSTSHRVELHSLMSRGIIELDISKTKHRIYLPYRLLKLAGVNQLEYRLYKQLST
AIGMASTPGSTLTEVNLDDRYQCTIREAAYEAEYKLRHYQEMQEFQSLRLRKDEDRIVEFHQQKDHLAGNNKNKYRKLNRNHFYTVTHHMYDDQLKT
>JAQQSV010000008_Pseudochirops_corinnae
HDLLEVGINMQSIARTRKFIFYAIKDLSHLYCRSINMVKVGINLGDNLNHYLTLLICYYYKGDDKMFQEESSCQYLSEQEYAVHGRDDTASLGKYLDANNQNPQLQQEIKAVSQEEMHSWRVILFILFCRLFLPKLAVGEQVCSEKILHQVPVHSEQGLAKYPQIWVSIATTKITFT
MMCQSFLLTFIHGLDQQTGHYASDRIIATSAAQGLFVGLLIIKTVLDHILQPSHRVELHRLISRGIRIELDCFAVKKISKHRIYLSYAWLLNLAGVNQLEHGLYPQLSVTAIGVASAPGSTLTEVNLDDRYQALEAAHEAENKLRHYREMQECKSLGLRKDDIIVEFHQKKNTISRQQQ
KEVLQAKQESLKI
>JAQQSU010009076_Pseudochirops_cupreus
QELLEVGINMQSIARARKFIFYAIKDLNHLYCRSINMVKVVYFRRSQPLSTLLICYYYEGDDKIFQEESGCQYLSEQEYAVHGRDDTASFGKYLDANNQNPQSQQEIKAVPQGEMHSRVILFILFCRLFLPKLAVGEQVCIEKILHQVRVHSEQGLAKYPQIWASIATMKITFTMMC
QSFLLTFIHGLDEQTGHYASDRIIATSAAQAPFVGLLIIKTVLDHILQPSHRVELHPLISRGIRIELDCFAVKNFKHKIYSSYAWLLNLSGVNQLEHGLYPQLSVTAIGVASASGSTLTEVNLDDRYQALEAAHEAENKLRHYEMQKCTSLGLRKDDIIVEFHQKKNTISRQQQKEVLQA
KQEITLKDQVAPGS
>JAQQSX010000006_1_Vombatus_ursinus
HDLLEVELNIQTKYARARKLIFYGTKELNHLSCRVINTVEAGINLGDDLDQCLLTLLICYYYEGNVKTIQESPISQYLSEHGIFNPLTRRDDTASLGKCLDANNSLQLQQAIQAEPQGEMHSRRVSLFISFCSLFPKSVVGERACIEKVLHQVQVHTDQELGEFPSTATMKIIFTICQSFLL
KFVITHQGFPQAGHDASDKMITTSVAQTHFVGLLIAKTVPDHILQHTSHPLAGKVQSELDCFQAIKEISKHELYAPYAWLLNLAGVNQMEHGLLVQLSAIAIGGTSAHRTTFIGVNLDDSYQVLGEAAHEAENKLRHRRKMQEVKSLGLRKDEDIIVEFHQQKNTISKQQTKEVLI
AKQELLLKLQLQPGSRLIGQEASPSSPQQNVNLLGDNSPHHQVKASADKTPKHGMLQCQSYQGNSFNNRLFLSHPDQSVHLPGNKLKGGYSLKSGSGQIHIQGRFTPSNRGTGVPNPQRQSKRRGHLNHMESR
>JAQQSZ010003561_Phalanger_gymnotis
FLASEMFWFCLLYGKSVIFDSELRQHIHSHQHCEYCIGDTKAKNYPKLWFPLLQKVIFTMIYIFILKFVITHQGLHQQARNSASDTIIATSVAQVHFTGYLILKTVLNHILRHKNHGVRVQHLAQGRGIQSKLDYFQQAGKEISKHGIYASDAWLLNLTGVNQLQHGLFPKLSTIAIGV
ASVHRSTLIGVNLDNNYQQLLESDHKAESKLKHHPKTQEVKRPAKDKTKKILWNFTSKKTTISRQQQREELKAKELPEKLKLQLNPRVIEEQEG
>JAANDE010000001_Trichosurus_vulpecula
FCLLYGKSVIFDSELCQHIHNHQCCEYCIGDTKAKNYPKLWFPLLHMKIIFTMIYTFLLKFVITHQGLHQQARNGASDKIIATSVAQARFTEYLILKTVLNHILQHTNHGVRVQHLAQGRGIQNKLDYFEQAGKEISKHGIYASGAQLLNLTGVNQLQHGLFHKLSTIATGVASVHRS
TLIGSNLDNTYQAVLESDYKAESKLKHHPEMQEVKKLLGQGKTNIIAEFHQKNTISSQQKEELKAKELLEKLKLQLNPGVIEEQEG
>JAJHZJ010000229_Bettongia_penicillata
STSYSTHLYDLLEVGLNIEQNMLRLESFFYAPKDVYLLYCIIDVIEAGVNLGNHLGNCLLTLLINVLGRQKIQYPSGSVSFEHGYSIRQLSRDDSTDLSHFPDANNQNPATGHTGIPQGEFTKGRLGYFISFASFFLPNLVEEQVSKALQSFHKVVFQQLPWKIIFTVMLQSFLLKFVIH
QGIHQAGHDASDSIIATLVSQGRFASLLIAKTILDHILLCTNDRAELHTLADRGLQNEQDSFQHAKSISRHEIYALFARLLDLAGVNLD
>JAQQTA010000003_Macropus_giganteus
YDLLEVGLSIQTKYAVRKLIFYATKDIYLLYRIIDVTEAGVNLGNHLDNCLLTLLVNYCEEDVKKYGESGSVSFEHGYSIRQLSRDDTTDLSHFPDANNQNPPLQQAITVPQGEFTKGRLGYFISSCSFFLPKLVEEHSKQGLAELSQTCTSTATRKIIFTVILQSFLLKFVIIHQGIHQAGH
DASDSIIANVGCRFASLLIARTILDHIVLCTNDRAELYTLAGRGLQNEQDSFQRAKSISRHEIYALFARLLDLAGVNLD
>JAQQSK010000002_Potorous_gilbertii
YDLLEVGLNIQTKYAQVRKLIFIYATKDVYLLYCIIDVIEAGVSLGNHLDNCLLTLLINYCEGDIKKYNDSPVAQYLSGEHGYSIRQLSRDDSTDLSHFPDANNKNPPLQAIALPQGEFTKGSLGYFISFCSFFLPNFVEEQVSKALQSFHKVVFQQLPWKIIFTIMLQSFLLKFFIIHQGIH
QTGHDASDSIIATLLAQGRFASNLLITKTILDHILLCTNDRAELYTLVLGRGLQNEQDSFQCAKSISRHEIYALFARLLDLAGVNLD
>JAPYYB010000004_2_Lagorchestes_hirsutus
LEVGLNMQTKYARVGKLIFYTTKDLDLLYCRVIICKKKPGINLGDQLNHCLLTLLKYYYEGDIKNYHYSPVVQYLVQRGYAVYQLSEDNSARLEKVSCEQSKAPALQQAIVVPQGELNKGRTGLLISFCIPLLPKLVVMDWAYIKKVLCQIIHYEQGLAEFLQTWISTVMSVAQACFA
GLLLLKTVLGHILLRTKNGVELYPLAWGRGLQSELDSFLAVKQITKYGIYSPYALLSLAGVNQLDQGLYLQLSAFPIGVTSAQGSMLIRVNLDERYQALKVAHEVKNKLRQGGEMLEVKNLGLSKEKVKKLLMEFYLQKN
>JAQMYV010000005_2_Macropus_fuliginosus
TKHQNLHYYTHKNISMDSLRILEVGLNMQTKYARVEKSIFYATKDLDLLYCRVINVKEVGINLGDQLNHCLLTLKYYYEGDIKNYHYSPVAQYLLERGYAVYQLSEDNSARLEKVSCEQSKSLALQQAMVVPQGELNKGRTGLFISFCIPLLPKLVVMDWACIKKVLCQIIHYEQSLA
EFLQTWISTVMSVAQACFAELLVLRTVLGHILLRTKNRVELYPLAWGRGLQSELDSFLVVKQITKYGIYTPYALLSLAGVNQLEQGLYLQLSAFPIGVTSAQGSMLIRVNLDERYQALRVAHEAKNKLRQGGEMLEVKNLGLSKEKVKKLLMEFYQKN
>JAQQTA010000005_2_Macropus_giganteus
TKHQNLHYYTHKNISMDSLRILEVGLNMQTKYARVEKSIFYATKDLDLLYCRVINVKEVGIILGDQLNHCLLTLKYYYEGDIKNYHYSPVAQYLLERGYAVYQLSEDNSARLEKVSCEQSKSPALQQAMVVPQGELNKGRTGLFISFCIPLLPKLVVMDWACIKKVLCQIIHYEQSLAE
FLQTWISTVMSVAQACFAELLVLKTVLGHILLRTKNRVELYPLAWGRGLQSELDSFLVVKQITKYGIYTPYALLSLAGVNQLEQGLYLQLSAFPIGVTSAQGSMLIRVNLDERYQALRVAHEAKNKLRQGGEMLEVKNLGLSKEKVKKLLMEFYQKN
>JAQOLO010000057_2_Notamacropus_eugenii
LEVGLNMQTKYARVGKLIFYATKDLDLLYCRVINVKEVGINLGDQLNHCLLTLKYYYEGDIKNYYYSPVVQCLVEYGYAVYQLSEDNSARLEKVSCEQSKSPALQQAMVVPQGELNKGRTGLFISFCIPLLPKLVVMDWACIKKVLCQIQIHYEQSLAEFLQTWISTVMLVAQACFA
ELLAVKTLGHILGHILLRTKTRVELYSLAGRGLQSELDSLLVVKQITKYGIYTPYALLSLAGVNQSTEQVLYLQLSAFPIGVTSAQGSMLIRVNLDERYQALRVGSEAKNKLRQDGEMLEVKNLGLSKEKVKKLLMEFYQKN
>JAMXIF010001253_Petaurus_breviceps
HDLFEVGLNIQTKYARARKLIFYATKDLDLYCRVINVIEGVNLRDHLDHCLLTLLIYYYEGDIKNYHDIPVAYLAEHGYVVHQLSGYDTTDLGKFLDVSISKSPTLQQVIQAFPQGKLNREKIGFFISFCSLFLPKLNVEEECTKKLVGYLAQLTLNKVLQNSHKVSTAPMKIVFTMMQS
FLVKFVLGLHQDAHDASDSITVMSVAQACFARLLIVKTILEHILLRTKNGAEFHPLAHGRGLQSEFDNFLVVKELIKHGIYARYVRLLSLAGANQIQLYPLSDIAIGITPSTHGSMLIGVNLDERYQALQELTHEVENKLRQCENEVKNLGIGKEEEEIITELEFHQQKNTFIKHQQKEVS
TLRLQRLSKKYSIKLNCNWPQNLQEKKRKVHLAQQNYQLTRTYNLSTRTPTDKTFKCQYITSSDMSQIPSANFLLFHPHTFEKSIYPFSDRQRRSTLKPSPYISKKASLSRHFGGSNPPHRLMRRHPNHGCGPWGCSAPRFHLNPETHILPENVSGEDFPEIVS
>JAJHZJ010000149_Bettongia_penicillata
KIIHINIPSKIMDTLHNILEVGLNIQHQIFPSTVICVQASKDLDYLCLLIDVIVVRVTLEDYFGYCLLTLLIDYSLEGDIKYIMKSCIIALGHDYLVYPHSPDDALEFNQFLVINNHHLPLLQAIQAVFQGELMTGRTGLFTAYCRLFLPKLVIGQACIEKLTHQICWHLEQDLSEFPESWTTT
ATMKIIFTVMRQSFLLKFIIIHLGVHQQAGHDAANSIAISVAQAQFTGLIIVKTVLNHILLNKDGVELHSLARGKSLQNELDSFAIREVTKYGIYSHYARLLNLLGINQLEHGLYPQLSAIALAVALANGSTVVRVNLVKRQALREAVHERDEAIISEFHQQKEEISINQQRVVKLKQE
>JAPYYB010000001_2_Lagorchestes_hirsutus
KIIHIDIPPKTMDTLHNILEVGLNMQTKYSQVWFVFYASKDLDYLCLLIDVIVVCLTLEDYYGCCLFILLIVYFEGDIKYIQSHASELLEHDYSVHPHSPDDALEFNQFLDINNHNLPLQAIQAVFQGELMTGRIGLFIAYCRLFLPKLVIGACIEKLTCQICQHLEQGLTEFPERWTTTAT
MKIIFTVMRQSFLLKFIIIHLGVHQQAGHDATNSIIAISVSQAQFAGLIIVKTVLHHILLNKGGVELHSLARGKSLQNELDGFRAIKEVTKYGIYSHYARLLNLLGINQLEHGLYPQLSAIALAVALANGSTVVGVNLVKRQALREAVHEAESKLKKYHERLQALPLQKRMKKLFLEFHQ
QKEEISINQQRVVKSKQE
>JAQMYV010000006_2_Macropus_fuliginosus
KIIHIDIPPKTMDTLHNILEVGLNMQTKYSQVRFVFYASKDLDYLCLLIDVIVVCVTLEDYYGFCLFTLLIDYFEGDIKYITKACIELLEHDYSVHPHSPDDALEFNQFLDINNDNLPLQQPIQAVFQGELMTGRIGLFIAYCRLFLPKLVIGACIEKLTCQICQHLEQGLAEFPERWTTTAT
MKIIFTVMRQSFLLKFIIIHLGVPQQAGHDAANSIIAISVAQAQFAGLIIVKTVLHHILLNKGGVELHSLARGKSLQNELDGFRAIKEVTKYGIYSHYARLLNLLGINHLEHGLYPQLSAIALAVALANDSTVVGVNLVKRQQALREAVHEAESELRKYHERLQALPLQKRMKKLFLEFH
QQKEEISINQQRVVKSKQE
>JAQQTA010000006_1_Macropus_giganteus
KIIHIDIPPKTMDTLHNILEVGLNMQTKYSQVRFVFYASKDLDYLCLLIDVIVVCVTLEDYYGFCLFTLLIDYFEGDIKYITKACIELLEHDYSVHPHSPDDALEFNQFLDINNDNLPLQQPIQAVFQGELMTGRIGLFIAYCRLFLPKLVNGACIEKLTCQICRHLEQGLTEFPERWTTTAT
MKIIFTVMRQSFLLKFIIIHLGVPQQAGHDAANSIIAISVAQAQFAGLIIVKTVLHHILLNKGGVELHSLARGKSLQNELDGFRAIKEVTKYGIYSYYARLLNLLGINQLEHGLYPQLSAIALAVALANGSTVVGVNLVKRQQALREAVHEAESELRNYHERLQALPLQKRMKKLFLEFH
QQKEEISINQQRVVKSKPE
>JAQOLO010000065_1_Notamacropus_eugenii
KIIHIDILPKTMDTLHNILGLNLQAKYSQARFAFYASKDLDYLCLFLIDVIVVCLTLEDYYGCCLFTLLIDYFEGDIKYITKACIVVLGHDYSVYPHSPDGVLEFNQFLDINNHNLPSQGAIQAVFQGELMTGRISLFIAYCRLFLPKLVIGDCIEKLTCQICRHLEQGLTEFPERWTTTATM
KIIFTVMRSFLLKFIIIHLGVPQQAGHDAANSIIAISVAQAQFTGLIIVKTVLHHILLKKGGVELHSLARGKSLQNELDGFRAIKEVTKYEIYSHYARLLNLLGINQLEHGLYPQLSAIALAVALANGSTVVGVNLVKRQALREAVHEAESKLRKYHERLQRLTLAKEDEEIISEFHQQKEEIS
INQQRVVKSEQE
>JAQQSK010000003_4_Potorous_gilbertii
KIIHIDIPPKTMDTLHNILEVGLNMQTKYSQVRFVFYASKDLDDICLLIDVIVVCVTLEDFLVVVYSHSPDDALEFNQFLDINNHHLPLQQAIQAAFQGEHITGRIGLFIAYCRLFLPKLVIGACIEKLTRQICWHLEQGLNEFPERTTTATMKIIFTVMRSFPLKFIIIHVEVHQQAGHD
ATNSIIGISVTQAQFAGLIIVKTVLHILLGNKGGVELHSLARGKSLQNELDGFAIKEVTKYGIYSHYARLFNLLGINQLEHGLYPQLSAIALAVALANGSPVVGVNLVKRQALREAVHEAERKFRKHHELLQVLPLQEEDEEIISEFHQQKEEISINQPSVVKLKQE
>JAQQSW010004217_Pseudocheirus_occidentalis
DDYSIQDHGFITQHIGSGIKHTYIYTIMETCLLCLQRSRLPCLLAHFHGESGGLSWLLTLLIDFYFESDIKTYNENQAAQFLEHGYSVYEHSPDDALELHQSLDINNHNLPFQAIQTVRQGELTMERIGLFIAFCSLSLHKFVIRELAYIEKLMNQICVYLQALTEFSESWTTTGSVKLIFT
VIRKDFLLKFIIIHQGLHQKAGHDSVDSIMANSLAQTQYTGVLIVKTILDHILLQTRDEVELHSLGRGRRLQNEMHGFQAIKEVRKFGTYLPYAKFLNIWVVNQLEHGLYSQVLAIALGVPSAHDSTLFGINLANTYQALRVVVHKAESKLWKHCELLDHLKLTKEDKGIISEFHQEEE
INKKKLAKIVEARQDLLEKLKVLALKSI
>JAQQSV010000007_1_Pseudochirops_corinnae
IHTKYVQLKLVFYASKDLSYLCCKLIDVIEAGKNLEDYLGNCLLTLLIDYYFEGDIKKYNENQAAQFLKHGFQYMSIAIKNHNLPLQAIQAVFQGELTMERIGLFIAFCKLFLPKLVNGELACIEKLMQQICVHFKQALTEFPETTTTGSMKITFTVIRQDILLKFIIVHQGLHEKAGHDSA
DSIIGNSVAQAQFTGILIVKTVLDHTLLQTKDEVELYPLARGRRLQNELHGFQAIKEVSKHGIYPYARFFNLLGVNQLEYGLYSQVLATALGVPSAHDSTLVGSSTGKTLEAPIARHLKLTKEDEEIISEFHQEEEINNNQQRMKARQDLLEKLKEHLALKSI
>JAQQSU010000001_1_Pseudochirops_cupreus
NILEVELNIHTKYTKYINIHTKYVQLKLVFYASKDLSYLCCKLIDVIEAGKNLEDYLGNCLLTLLIDYYFEGDIKKYNENQAAKFLKHGYSVHGIAINNHNLPLQAIQAVLQGELTMERTGLFIAFCNLFLPKLVTGELACIEKLMQQIWVHLEQALTEFPETTTTGSMKIIFIVMRQDML
KFIIIHQGLHEKAGHDSAHSIIGNSVAQAQFTGILIVKTVLDHTLLQTKDEDELYPLARGRRLQNELHGFRQAIKEVSKHGIYPYARFFNLSVVNQLEYGLYSQVLATALGVPSAHGSTLVGSSTGSKTLQAPIARHLKLTKEDEEIISEFHQQEEEINNNQQRVKARQDLLEKLKEHLA
LKSINNQDNSRVWGHEKHQERSWSTPKENYNLGLEVLRHQLLSLLSDHPPLAFHPCHPSFKEEITYSKVEVNKKEHLLALQLMGSLRKKNSSGNRNIRRKQSSDSERGGHLHTHSRSRWCQDIYSSTRNQRLTNFSASVSERLSSPSGQKS
>JAQQSZ010004907_Phalanger_gymnotis
SKSKRWMLTFHPRSWIHYTTYWKGINIQTKYARVQKCVFASKDLDYLCCLLDVGHESGGLSLFTHIIEGDKKYNGSQIAEYLLEHGYSVHPHSPDNAPELRQSLDINNCNLSLQHTIAVLQGELTTGRISVFIAFCSLFLPKFVLGEQAHIEKLLCQICVDLEQGLTEFPETWTTTTMK
IIFTVMRQSFLLNFIIIHYWLHQAGHDATSSIIANSVAQACLAGLLIVKTVVDHILLTKDGVELHPLASGRSLQNELDGVRATKEVTKHGIYSPYAKLLNLSGVNQLNHGLYPQLSAIALGVASIHSSTLFGVNLDKRYQAFREAAHEAESKLGKCHELLEILNLNLQKRMKKSMNFHLQK
EEIGVVKAKQDLLEKLKVQLGPKPINEQDKSPILGQLRVGTRRDYSPPQNRITTADCGFAINSFNPFSQTTDHWPSIHATSLWRRSSNSLAKQKKRGQHLTALQIMCSLRRIPPQLRAILEEQKESSDSEHEEDTDRIVVPRYLSSIGTQRPPNSSATVPAKTFFFLRAKKSRPPIASN
KIPLYRCFKLFQELGGALSRAQALDQEDLTSNPASDIWHTYLCDLEQVTPQLSCLPLSKIKKYFNKTCKAFYLTAHTGKSFVFPDMMTSE
>JAANDE010000004_1_Trichosurus_vulpecula
FVVSSLTLGMNLGAYRDNYLLTLCEGDKKCSASQVAEYLLEHGYSVHPYSPVDALELRQFLDINNCNLLLQHAIQAVLQGELTTGRISVFIAFCSLFLPKFVLGEQAHIEKLLCQICVDLEQGLTEFPETWTTASMKIIFTVMRQSFLLIFIIIHHRLHQQAGHDATSSIIANSVAQAWF
SGLLIVETFLDLILLTKNGIELYPLASGSSLQNELDGFRATEEVTEHGIYSPYSRLLNLSGVNQLDHGLYPQLSAITLGVSSTQSGTLVGVNLDKRYQALREAAHEAESKLRKCHKLLEVLNLAKEDEEMISEFHLQKKEINTNQLKVVKAKQDLLEKLRVQLAPKPINDQDKSPILGQLR
VEAPG
>JAANDE010000004_2_Trichosurus_vulpecula
SKSKSRWMFISHPRSWIHYHNILKGELTYRLNMPEYKNMSSRHSKTTTFVVSSLTLGMNLGAYRDNYLLTLCEGDKKCSASQVAEYLLEHGYSVHPYSPVDALELRQFLDINNCNLLLQHAIQAVLQGELTTGRISVFIAFCSLFLPKFVLGEQAHIEKLLCQICVDLEQGLTEFPET
WTTASMKIIFTVMRQSF
>JAQQSW010000004_1_Pseudocheirus_occidentalis

PKVTWLVLCVSEAGFEHQVLLTPGLVVYSLCHLATPRKLIFYAADLGLLCCKIVDVVEQGQPRRSSRVDHCLLTLLIYYYYEGIKKYHNSPMVQNLSEHGYAVSSSPWGYCPGLAPCKQSKSSIQQAIQAVPEELNRGKTGLFISFRSLFLPKPVIGEQTCINKVLLQIWFHSEQGLAE
FPQIVSTATMKIILLLYTGLHQAGHDALEAINATSVAQACFARLLTVKTALDHILLCTKHGVECHPSAGISLQNELDSFQCVAKEILQHGIYALYAQILNLPGVNQSEHRLYPQLSATAIRVASAHRSTLIRVNLDERYQVLQESAQEVESKLRQHQEMLEVKSLGLVRKDILTEFHQQ
NGISKHQQKEVLKTKQELLRKSNCSPQDQENKRKVHLYSSIILTYEVIQLIPNNSDKTFKECQDIIPNDFSQITIHEYSLLYTHALEKPIHPPSEMARKKLHPGDQSTLTSISEKASLSQIFQKIKSNLHIVNREKDNEPIMDIKDPGGSLPFKVPPRNSHTPPQQQHSPQPPSHQKPMG
QGPTKLSIAETPSYIETLSKYNETR
>JAJHZJ010000177_Bettongia_penicillata
QRIWVFYATKDLNHSCRIIDLVRVGLRRESPLFVHSLDLLPLSNVKIFQESSAHYLSEYRHTVSGGDDTASLGKYLDANNNLQLQEIQVVQQREMHTWTVSLFMFCILFLLKLVVEEQACVEKVFHQVQVTTEQDLAKFPQTWSATMKIIFRMMRHQSFLKCVIHQGLQQARHN
ASDTVIVISVAQAHFVGLLIVTTSLDHILQCTSHRIELILRQRYPEDCFHRAIVKEIFKHGIYAPLNPPYTQLKYAEVNQLEHELYPRLSATTSKRYLCFHRYVLGQAAHKVQSKLRHYQESKKLSNRIRKDIIMKFHQQKKHHYQARTQKHNLNRNYLKNKTHLSPAVVTEEG
>JAPYYB010019217_Lagorchestes_hirsutus
ISGLNIQSILRARNLIFYATKDLNHSCYRLREFTVRLANLGDNHCLLTLLICYYYEGNLKIFQESPAAHYLSEHRYTVYYWWGYCQLGQVSCLKSTITAGNPSCPAKGNALESQFIHVCSLFLLKLVVGQACTVKAFHQVQVAKEQDLAKFPQAWCTMKIIFRTMCLSFLRKFVTYQG
LYQRVRDDASDAVIATSVAQAHFVGLLTVTTSLDHTLQCRSQGLSLAPRQRYPEDCFHRAVVKEIFKHGIYTPYAQLLNYAEVNQLEHELYPLFATISMCCLCFHESTLVGANLDYGYVLQAAHKVQSKLRHYQETRENCLGMRKDEKDTIMKFHQQKNTISSQQQRGIIT
>JAQMYV010043230_Macropus_fuliginosus
HDLFEVGLNIQSIQRARNLIFYATKDLKHSCYRLIDLGKVGFRDNHCLLTLLICYYYEGNVKIFQESPAAHYLSEHRYTVYYWWGYCQLGQVSCLKSTITAGNPSCPAKGNALESQFIHVCSLFLLKLVVYEQACTEKAFHQVQVATEQDLAKTPPSYECTMKITFRTLCQSFLRKFVT
YQGLHQRARHDASDIIATSVAQAHFVGLLTVTTSLDHTLQCTSRGLSLAPRQRYPEYNCFHRAVVKEIFKHGIYTPYAQLLNYAEVNQLEHELYPLFTTNSMCCLCFHESTLVGVNLDYRYVLQAAHKVQSKLRHYIKKRENCLGMRKDEKDTIMKFHQQKNTISTQQQRGIITK
>JAQQTA010000003_1_Macropus_giganteus
HDLFEVGLNISKGIQRARNLIFYATKDLKLSCYRLIDLGKVGFRDNHCLLTLLICYYYEGNVKIFQESPAAHYLSEHRYTVYYWWGYCQLGQVSCLKSTITAGNPSCPAKGNALESQFIHVCSLFLLKLVVYEQACTEKAFHQVQVATEQDLAKFPQAWCTMKIIFRTLCQSFLRKFV
MYQGLHQRARQDASDTVIATSVAQAHFVGLTITTSLDHALQCTSRGLSLAPRQRYPEYNCFHRVVVKEVFKHGIYTPYAQLLNYAEVNQLEHELYPLFTTNSMCCLCFHESTLVGVNLDYRYVLQAAHKVQSKLRHYIKKRENCLGMRKDEKDTIMKFHQQKNTISTQQRGIITK
>JAQOLO010000065_2_Notamacropus_eugenii
HDLFEVGLNIQSIQRARNLIFYATKDLKHSCYRLIDLGKVGFRDNHCLLTLLTCYYYEGNVKIFQESPAAHYLSEHRYKSTISGGDDTASLVKYRDANNNPQLQKEIASCPAKRNALESQFIHVCSLFLLKLVVYEQACTEKVFHQVQVATEQDLAKFPQAWCTMKIIFRMMCSFLKF
VMYQGLHQRARHDASDAVIATSVTQAHFVGLLTVTTSPDHTLQCTSQKMELSPEAEVSRIKLFYRAVVKEIFKHGIYTPYAQLLNYAEVNQLEHELYPL
>JAQQSK010000004_Potorous_gilbertii
RNLISYSTKDLNHSCRIIDLVKVGLRENLNHCLFTLLICYRYGGNVKIFQEDSSCPLSFRTTHSHSGGDDTASLGKYLDVNNNPQLQEMPSCPSKEMHSIVSLFMFCNLFLLKLVVEEQACIEKVSHQVQVTSEQDLAKFPQTWCIFGMICQSFLLKFVIYQGLHPQARYDASDTVIA
TLVIQAHFVGLLIVTTSIDHILQCTSHRIEFTPRQRLSRVRLFYRVVVKEIFKHGIYAPYSQLLNYAEVNQLEHELYPLSATTSVLPPLPESTLVGVNLGYRYVLQAATHKVQSKLRHYQEMQEIKYLELGKMKYIIMKFHQQKNTITRQQQKEVLKSKQELLKK
>WOXC01005789_Gymnobelideus_leadbeateri
ITYQTMDSFHDILKVRLNIQTKYARQKAIFYDNQDQDPLCCQAIHVIEAGVTLGDTDNYLLTLLTNYYYEGDEQPNGSIPEHRYSIHQVSNNTANLSQFLDTIITVPHLQKVIQAVPQGELTKGTGLFVAFYSLFLSKILARECTGNLEKAMLQIWVHSEQGLTEFPQAPSMATIKIIF
TIMQQGFLLKFVLIHWGIHQQAGHDVADSIIATLVAQACFAGLLTVKTVLDLALTCLDRGLLSRVEVLGCVTSLPEHVEASERAPNYLQQLRKPQFGTKLPCLVVESYCGKKLKSQVIEWQHEQAKEASKQRNPTLYSYFGDKKGQSMEEEDNQGTPAPKASKKNTSWLQATEE
LKMDFENQIKEVEEKLGREMRMMQENYEMRVNNLTEENNTLKNRLTQMAKEAQKDNEEKNALKSICQI
>JAMXIF010000002_Petaurus_breviceps
ITYKTMDSFHDILKVRLNIQTYAKVRKVIFYDTKDQDPLCCQAIPIIEAGVTLGDHLDNYLLTLLTNYYYEGDEQPNGAIPEHRYSIYHVSQNNTANLSQFLNTNNYSPPLKQVIQAVPQEELTKGTGLFVAFYSLFLSKILVRESQVIKAMLQICVHSEQGLTEIPQTWSSTHTIIIFTIM
RQGFLLKFVLIHWGIHQQADHDVADSIIAILVAQACFAGLLTVKTVLDLALTCLDRGLWSRVEVLGCVTSLPKGVESFREGTQLIAPAAEKTPVFGTKLPSLVGRVSYCGKKLKSQVIEWQHEQVKEAKQTEQSNFLLLFWGQKGSKHGTGGRQQSQNSCTKSFQEKYELVTGHR
GRLRRILKNKEKWRKNWEEKELQENYEMRVNNLTEENNTLKNRLTQGRPGSTASKSTGSSVTMTWRSNPRLRHLAPTQKHLAVPASYSTPIAGSQVAQQVRALAPVLGPGDRIPGSDTQHHPQKLSVPASYLTPIAKKKRLTQMAKEAHKGNEEKNALKSRICQMEKESQKL
TEENHSSRIRAQMEASEFMRNRDTIKQIQTNEKIEGNVKNCARSHFTPHRWSAISTGIGQGPAETVEWLQQPHWYTQDGSYRLLDLLYE
>JAQQSV010000005_Pseudochirops_corinnae
ISYETMDSFHDTLKVRLNIQTKYIQVRKVIFYDTKEQDPLCCIIHLTEVGVTLGDHLDNYLLTLLTNYYYEGDEQPNGSIPEHRYLIHQLSQNNIVNLSQFLDTNNYSPPLQVIKAVPQGEFTKGKNGLFVAFYNLFLSKIIVRESQVIKVMLQIWVHSEQGLTEFPQTWPSMATIKIIFT
IMWQGFLLKSVLIHQGVHQQAGHDAADSIIATLVAQAHCAGLLIKTVLDHTLLHTKDGVRLYPLAWDRGLKNELDGCSSNHTGIPRMAASTGSICFTRKIVVGVNNL
>JAQQSU010000003_2_Pseudochirops_cupreus
ISYETMDSFHDTLKVRLNIQTKYIQVRKVIFYDIKEQDSLCCQIIHLIEVGVTLGDHLDNYLLTLLTNYYYEGDEQRPSGSIPEHRYLIHQLSQNNTVNLSQFLDTNNYSPPLQVTQAVPQGEFAKGKTGLFVAFYNLFLSKIIVRESQVIKVMLQIWVHSEQGLTEFPQTWPSVAAIKII
FTIMRQAFLLKSVLIHQGVHQQAGHDAADSIIATLVAQAHFAGLLIKTVLDHTLLHTKDGVRLYPLAWDRGLQNELDGCSSNHTGIPRMAASTGSICFTRKIVVGVNNLFKLHSLVQGKSQHPELQRKNREIKTNREGSDCLNQLYTLHIYIYIYIYIYIYSESPSESQDPSHKRAHKA
KYQLRTYIHFSEAENITILLTQCRNTLTKGVKAFLQASIPASFPWAPREAYWAGKILNSINITMVSGVLSEKKISRHGIYSPCQTSKPGGKPTEHRLYPQLVCNCNRCGFHPWEHTYRSKPGEIPSSQGVSTQSEQVPVSPRNAGSQTSQACKGRRKHLRVPPAEEDQHNIGKGETA
>JAQQSZ010059767_Phalanger_gymnotis
QGKPNKGRIGLFALFCRLFLPKLDRGILPQMQVHSEQSLAELPQTGVSTATMKSVFTMTQNFKFVLIHQGLHQQAGACNQTDSITAMSVDQACFIRLLLKKTVLEHILLRQRTGLIDLHPLAQESGLQSELDSFCTIKEIIKHEIYAPYALLLSLAGVNQLEHGYTRQLSATATGVTS
VHGNMLTGVDLDERHQSSPKNGPKLTSNNIEKCQVKNLGLGKEEEVLTEFQQQKNTFSKQKCWKAKQEILDKIKLQLAPRLTGKKGKGPSLARQNIDFPGDNKVRPRLKLQLIKPSSANTFGSPATCLKLLSNFLSFTLIPLDKVNLPFQQVVKKVYPAQLTSMIISKKAPLSRHLG
DSNPPHRMIKHPNHRCPWGCTAPWGSTTKKLILHSNDSLRSPKNQQSKVLPNCNQQKKFQSTKHCDGTIRPGKPLYRDYLQWQSRGIPYAHQER
>JAANDE010000002_2_Trichosurus_vulpecula
LQHTIQGKPNKERIGLFASFCRLFLPKLDGGVLHQMQVHFEQCLAELPQTGVSTATMKIVFTIVQYFLLKFVLIHQELHQQAGEHDASDYITAMSVARVHFVGLLLKTVLEHILLRTKNGVELHPLARKRGLQSELAGFQFTVKDIMKHEIYAPYALLLSLAGVNQLEHGYTCQLSAI
ATGVTSVHGNMLIGVDLDRYQGSLRDAPTEAMNELRQHQESVKNLGLGKEEQILSEFQQQKHVFSKHKEKEGLEGAGNTNTAIGPKTDRKNRDKSIPCP
>JAHYBR010000002_Dromiciops_gliroides
NLHGLLEIGLNVSQCYSAERLVYFSTSAYIQLCKDATEARVDLGENTDNCLLSLFIGYFYEGPALTFKKGAVSQYLAKYGFSIKETPKDIERKRKRGIKPIDFIAKQEDAPQVIRALRVIDSSAILKSRTQTFIEFTSFFLPKLVVSERACLEKVRQVQVHFEQGLTEFPQTWVAPGTMRVI
FLARQCFILKFTLIHQGLHGQSGHDATDSIISNFVAQARFARLLIVKTVLEFILQKTTDGAELHPLAKDRALASKLAAFRWSVKGIAVHDIYAPYVHVLNLLGVNHLEHGLYPQLCAITIVAAVHRSTLIGINIHELYQALREATHEAEHQLQQRGEEEEIAQLGLKREEAASFTKFNQN
KKIHASQQEAMRAKEELWKKLEKHLRQGEDHQETNPFQGAYRQDPGKNLRENDSFFHICFDLADQNILRLLIPGPRILDSRTPSQESESNGDTCPPQPPQGDSDNSESEEAALIGRSAGPFLWGQGQPCSKRAQPIESQVPFDPRSILPEPGLRPLTPTPFLPPGGVGEYTACSG
QSLTPIKEAVGQLGGTVYKAPALDSGGPEFKSSLRHLTLTSCVTLGKSLNPHCPLKNKKQTKKEAVYHEIQAEHKKTRVNIPFMFR
>JAHYBR010000004_Dromiciops_gliroides
KQIMNFHPCIIHRPNANQLVQALDAIPQGTINRSVGLFTVFSSLFFPKLSVGEKAYINKVQQQVQVHLEQGLLEFPKALTVPDTMKIIFTLMRCFIFKFIIIHQGIHQYIGYDANDSIISNSVSQACFAELLIIKTVFEFILQRTHNGIELQPKAQTLASELSPFRRAAKRVSSHGIYAPYTH
ILNLAGVNLEHELPQLSAMAIGVATIHHSTLISVSLDEQRQPGGPVGKAPALDSGGPEFKSGLRPLTLTSCVTLGKSLNSNCLTTKTNKQTKNLDEQYQALRETVHQAEYQLQRKEFLEIKQMDLVKEEEEIIVTFHKHKCDIHAQQQKEMVSSKQELLDKLKSQLA
>JAHYBR010000002_1_Dromiciops_gliroides
LCCSIIDVTEAGLDLGDNTDSCVLVLLIAYFYEGSITKFKGALSLYLTQHRFTIKETFPKRESRKEDLGLTDFIAMQGNTPHVVCALQAIDQGNISKSIRLFIAFTGLFLPKLVLGEACLEKVQKQIQVHLNKGLAEFPHIWVAPRTMKVIFSLVQQCFILKFTLITKVSAVRHGATDSIISN
SVAQARFAGLLIVKTIGIYSLKNNQSELSNSQHNCSKKITNGGQLGGTADKAPALDSGGPEFKSELRHLTLTSCVTLGKSLNPHCLGKNNKNNNKTTTNGVESCPLAGEVLASELAAFCQAVKGIVTHGISSSYARIFDLSSVNQLEYGLYPQLSIIAIGVAAVHGSTLVEVNIDKALQ
DAAHKAERNLQEKREEDKVTLGLKKDEEIIKNFNRNEWDIYATQQKAISRAKEELLRKFKEQLGHQDDQDLIHPPASRLQPSPHWGLKDKDQHHGSPPESTNFTIPKLLIPSPSNRGKYISAKEGTSFPQAAADGSEPEEANHSWVPRQSLFRGKGQCGQIASWIRRSTKSLCLLI
PIRFYQNPRLLGLKSFHSPQTPGPSGEILLDPEQLLMPIPEENTQETQAEYMKTKVIVSFQTFKGTTVEYTNTVKSIVGRAISCPRRKCRDHFNPTGSSSHGMSSQTRSYLNISQDEQSFNLYKKKSKGPLKAVLF
>JAMXIF010001129_Petaurus_breviceps
YMSDLKACPILSCFPLTKQIQEPVLADTLGKPGCLLLQGDIKIYDSPVAQYLSEHGYSIHFSDDTVDLSQFVDANNQNPPLQQAIQAVPQGELTKGKIGFFISLCSLLLPKLGENICIEEVIRQIQVHSEQDLIEFPQTWTSTANMKIIFISMWQSFLLKVIIFFLFGGQLGLSNLPKVTQL
VFCLFYFFLLHVGQIQTQVFLSPGPALYSLHYPPLKVIIIHQEIQAENDAADSIIATSAAQICFNGLLIIKTILDHILLRTKDRIELHPHGTESKL
>JAMXIF010001117_1_Petaurus_breviceps
DSKLKCILISSTCKIFMDSLNDILDIGLNIKMEYVIRNLIFDATKDLDLQFCQILNMIKAVNLADGLDNFLLTLLMHYHFKDDIKYDGAARWLSEEHQPQCRDDLRFESGLRYLAPHPEMTGLVTLSKLLNPNCFKKKERMVKIHTFWSTDIQSVTLFRITRMIITHPLIISHTMQFLRG
NSQKGGYAGLLIIKIMLDYILLHTIDRVELHLSACGKGFQNEVDGFPCAIKEIRRQGIYSPYAKLLNLARVNQLDHGLYPLSATALGVPSPNESTLIEIILDARYQDLRESVHEAESKHWCAMLEVNHLNLEKEDEDIIEFYQKKDEINIHEQKEVSRARDLLEK
>PVIH01004834_Sigmodon_hispidus
KQLLGIWELLVQQKRLTTILNLLKTLFKTAVYTLNVPDPNQLCFQIMLAIEAGIDLHTLLDSCLLTLCTQHTYDGNYKLLEKSADHYVESKGLMLKLIDSKNSSNLIDLYSAVGPTQGTFGIATLANNHEEHQKVGQFLSFCASLFLPNSSCKDKLSYRIRIIQYPQGWLTPGFMMVVF
GITRVWVSFTFKFLLIHHRINLRAEHYADNTVITNSVAQLRFSGLLIVKTVLDHSLWREEDHITLHPSGCSFKVWPEAENFKLVLKSLAHHGTMCPFAWILNLSGVNNLKHEQFSQLSVIVLGVATAHKTLAGVTVKENQLREAKLQELGEEEIGRLVLDQDEAQIFKDFHKQKYEI
GYKRSPGLLSVKNCSNAKWLEATATTSFCCPEFPPPGHITLP
>JAHYBR010000001_Dromiciops_gliroides
MIIYLIEAGVDWGDNADNCILVLTLQISSPSIFYEGSITKFTEGAISLCLAQHGFNQRSYRMRKRKKDSRKEDLRLTVLSQCKEIPIMLSMNYKPNLAIFEEFIVFMGLFLSKLVIGERERERERERDKESTYLEKVQQQVQVQFEQRLAKFSHTWVTPQTKEVIKEVILSMVWKSFILE
FTLIHRGLHQGHDVTDSIISKSVAQARFAGLLTVKIVLECILQKTTNGVELYPLIRSLGIVKIGSFSPNCKRIATYGIYSPYAQILNLSGNLEHLYPQLSTVPIGVAAVYGSTLVVVNINERYQALQKAAHAEYKYKKKREMDEFRLKKDEEKIIKHFNKSKWDIHATQQIAISRTKEE
>PVIY010089203_Myrmecophaga_tridactyla
TQIKLDMDDLIAYEGMKLHSRFSRTRKPMFYHVSNRIDNYCDCEYISVISMTVSILTDSNLLVTLLVTYYYDKQIDQFTKSTTVKHCSENGYTIVRPSYHRCKLNFSLTKIIKLHTLCSLSAMPADSCFLAFCSLFLPKRIEGDCLENVTMQVQVYSEQGLLECPKTWMAAATTKMIFII
MRQNLIKFVLIHQGLHQQTGHDAAYSTAANSVFQAQFASWLIVVLQRGQNSVELHPLASHDLSSARMMFRKARKEISRHGIYAPHACLISAGVNQVEHGLYTQCSATKTGVASAHSTLVGVNTDERHQPLRAAHEAEAQLKRNRDLREAQNTNLPMKRTSFSNKMFLASIQ
YQQQKGHPGKRSSRMTK
>JAMSGI010000034_Tamandua_tetradactyla
IIDADETGIHFSDQLDDFSILTLLVTYTDKQIDQFTKTTTVKHCSENGYTIVHPSTTDSTNLIFHPKSNSTPCTAVPRLPICAFVAFCSLFLPKCIEGDCLENVTMQVQVYSEQGLLECPKTWMAAAITKMIFIIMRQNLIKFVLIHQGLHQQTGHDAAYSIATNSVFQAQCASWLIVVL
QRGQNSVELHPLASHDPSSVPVMFRKARKEISRHGIYAPHACLISAGVNVEHGLYPQRSATTTGVASAQGSTLVGVNTDERHQPLRAAHEAEAQLKRNRDLREAQNTNLPMKRTSLSFQ
>PVIF010042287_Tamandua_tetradactyla
IIDADETGIHFSDQLDDFSILTLLVTYTDKQIDQFTKTTTVKHCSENGYTIVHPSTTDSTNLIFHPKSNSTPCTAVPRLPICAFVAFCSLFLPKCIEGDCLENVTMQVQVYSEQGLLECPKTWMAAAITKMIFIIMRQNLIKFVLIHQGLHQQTGHDAAYSIATNSVFQAQFASWLQYV
LQRGQNSVELHPLASHDLSSVPVMFRKARKEISRHGIYALHACLISAGVNVEHGLYPQRSATTTGVASAQGSTLVGVNTDERHQPLRAAHEAEAQLKRNRDLREAQNTNLPMKRTSLSFQ
>JADWME010000004_Gracilinanus_agilis
LDSVQFLSSSLWATRGIVTQVTIPVFFIFIKHLEIHKIQLYACGSGLLSHGKMELGSIRAFQESTVAQYLINNGFTILQSIGVSRISNYLIFLKDGYSLSRSLPSSSNCVNGKQHQLIQIRTFSSILQLIPLHQIAHLKQRMTQPTPWFIPGPRRLEFFFPCAVVLHLKLIHQGHQLSEHDSR
HYQISNSVAQPQFAGMPIIKTVFDYILRKTDKGALRPLTLLRLQSESEAFKKAVKTTAKHGIYVPSAQLLSLAGTSPQLSAIALGSAASHGSTVVGVNVEEQYQALREAVHGEERSQATGSTNPLKRTGLPTLLRDIQSEPKLTVTKNQINF
>JAQIHE010000997_Monodelphis_domestica
LSSSNIEDLRVILTGLNPDVRTRNLVLRQLSVQIQLCCQITDAPEAGIDLGECLKSALLIYFQYYFESDIRAFQEGTVAQYLTNDGFTILQSTGGGISRLSDFLKDDTVSVGLCQALATVSMGHSISFRLGLLLAYYSLFLSHQTQVHSKQGMSQLPTTWFIPGPARLECFFSCAVILHLK
LIHQGLHQLSEHDTADSIIISHSVAQARFAGMPIVKTVSDYILRKTDKGAELHPLTQLLSPQSELEAFKKAVKTTAKHGIAPTQLLSLAGINYLEHDLYPSLALGLVTSHGSTLEGVNVEEQYQALEAAHGKGVRLQFQHEIQEVRSLGLNSNEAEIVLEFHEHKSSIKYKQLEMIAIQQ
EMFEKLTKRVGQAIRTSPPTFAPGHLAEHQDFVPQPGSAWLSIRPENPPSQKNRAPHNTQGHPIKLTQVD
>PVIH01004093_Sigmodon_hispidus
RQALLAGEPGIDQRDLLDNCLMTLCIHYRKSFKTNAACQYLEVHGLILEALDNKNDLSDFLIYMRRPETRQALMALTNNLYYHPLYKLFTVFLPKIMVCEHGCLKKIQRQIAVYTNRLSQPQTTPGFMIIIFLIMRVSFTLKFLFTHQGINLEGHDGDDVTNSVSQARFMGLLIVRIA
LDHILSKDNDIILDHVVGLVNVPKKVSFRIALHCAYAPFAKIQNISWVNHVENGYPSLPPVLGVATTYREMAGVIKNEYQQLGEAASEAAVTGLYQSKGNQLGLEPAEVKILLDFHKKPRIGSKQDQIILAKSMRNSNIRQQLATKGQGQCDKVLSLGQFPWNNP
>PVIH01004934_Sigmodon_hispidus
DVDPLGILSDTQPAETLVQIRKLFSFHVPDPSLCSQIILAKKAEVDLGELLHTPLLTIVLDHTYKENHKAFEKGAVQHYLLTQGLMLKLMDKGNISNLSDFCPLDQPKKLREALTVLANNLEEQWRVGQFLFCSLFLFKLVVGSLLREIQRQRTFHAEQGLVQYPQYWHIFCLWNST
NELYHQVFFLIHQEINLEAWHNADNIITTSIPQARFSRLLTINMALDYVLPRKSNHISLHSLVRSIKVWEVVESFRLVLKGLAHHQQSTRFTGILNLSGFNKHGLFLQLSAVALVAHGGILARVSVVNEQYQLRKAVSKAERKPQEHLKEQEMRQLGLDKTKARILHDFHKKYKTGY
QAEVNLATKHEKLQQLAEKLARAAKDRCQSWPLLMQEHLCFPLAITWPTVSEQRESVIREVCEAIENVSPEEDVQSESSEPASPSVQHSEFPVLDTDLVEDKEDNFFLEKLDVLVACDASGDLVDDLHSAELDDIRATTNCQCGSFSPLNCLSNTYLSVPNTTDSSTNEAISYCLER
KDPHYPREPRRSRSMMPTSGPPLYHDLAQSRKPLTVWLLPYWHHSPLSHQNQPLKNLNEDHSYAGSFRNDNGSILRLNSRVSLHGLLDGLKYNCSFNKKNPHNFINHNGSAFYQSLERTTVEYLLRDGNLEHSSQYAINSQWLTWNLIMDAR
>JAJGSA010000014_Jaculus_jaculus
VDAVEAGIDLGRHLDGCLLTIVKYYFEGDVKRFQRGSATDYRREHGIIILQKSRSAFSRLDSVVMEGPSSSQRSQAFTSVAQGQASPSLLGQFLAHCSLFLPKLVVGDHACFEKATRQIRLCAQSLEFPKTRTLKAVCSLLRRSFVVFNRGLDQVHCTGWSRLSHHLCSRFTGMLM
VEMVLRFILLETDIGIVELHYSQALRNELDSFYKAVKEIEKHGTYVPYACLLNLMGINHLKHELFAQLVALGVAAVRRCTLAAVKVRERDQPLREAAAAAETQHGNFRRKNLGIEQEKAETILKFYGQKRREVTLQQQRALTAVRQKKAELTRGLNS
>JAJGSB010000055_Jaculus_jaculus
VDAVEAGIDLGRHLDGCLLTIVKYYFEGDVKRFQRGSATDYRREHGIIILQKSRSPLSRLDSVVMEGPSSSQRSQAFTSVAQGQASPSLLGQFLAHCSLFLPKLVVGDHACFEKATRQIWLCAQSLEFPKTWTLKAVCSLLRRSFGFNSTRGWTRFTAQDGADSVIIYAHFSGMLM
VEMVLRFILLGIVELQYSQALRNELDSFYKAVKEIEKHGTYVPYACLLNLMGINHLKHELFAQLVALGVAAVRRCTLVAVKVREQDQPLREAAAAAETQHGNFRRKNLGIEQEKAETILKFYGQKRREVTLQQQRALTAVRQKKAELTRILNSQDHPDNGERRAQLPPMKIWHD
PSGAPQFTGPDTKVTACPGAWTVTSERRNQCLPAQDSFQHEGKPCDVNIFSDVFPQGSMSHIPAPSSTLREETGPCPMKKVRVGPGASPKESLVEPQMQQSSFHSPTSSPCIIPFLNSPRTEKEHRMGIGPPALLKLFTVEDDQETVKANKVLQSGSYCWKSPNQEQLVGAD
GMNRGWTSPPGPHTGGQQEQKNVPNTGMGKLAITPISPPPTIHCLQEMLIPKSPSAGNLAFRTPKLMGDTIKPLYWVKFKTQGKDFVDYPNW
>PVKW010004947_Orientallactaga_bullata

HDLLSVGLSLESSISRHGMKTCSLQQPVFEAPINAIEAGTDWRDHLEGCLLTICIKYYFEGDLKRFQQSSATAYLREHGFTILQKTGSASLNLATLRRAPRPVQRSQALFCGQEMSPFLLGHFPAYCSLFLAKLIVGDQACFERAMRHIRVCAEQGLEFPEQWTLKAAACTLMRRCF
GCEFVLIHQGLEQVAARDRADPVTMHGHFAGMLVKQIRERSHLLKARRLKKNELASFCKALKAIEKHGIYVSYGWLLNVTAISHKRRLDPQLLPVALGVATVYCTLGTVKSHERCQPLREAVREAENTTCELGDVENLGLEQEGEDTVLKFCRQKHGEEASSDNKKLQQSAEGV
EKLRRLNSQGLMPSSLTGARHPGMEAPSTIAPIENMTSSFRLTPLHANTEMSACPGAWFPDLKQDAESVPSSTGSVSAKDGVGEPFDINSVSDFFPQGPTSHAPAPSSAPEEEIESCQMKKVKVGPGANPTESWLVGPGIHIDHPSTSPQDHPASLPSTPPNTRRVQDGNRP
>PVHP010005616_Zapus_hudsonius
YRALSASVSRHKHENLFPMAALSLGSSVLKFVQRNEIDRLDGCLLTICIKDYFKGDLKCFSKAKYLTERGFTILQKTGGPTQLGQAFNSVPQGQRSAFRLGKSLAYCSPFLPKLLCGSSCFEKVKWQIRVHANQGLAFPKKWAMAGDLKVVFTLVRCGFVLKFTLTHQGLCQDDSL
SLPAHFAGTQMVEMAFRYILWVEAELHPFARKPGIQIELGSFCQAVKGIRTHGFCAPSARLLNLAGISHLKHRLSPQLSMALGVATVHKHLSGVRNHESYQHIGEAAQEAEMQREHQEVKELEIQQEKEDVILEFHKQKHRISHQQEEIIAAKQKREKLKWELNSQGSI
>JAHYBR010000001_1_Dromiciops_gliroides
RDSTENCILTLLVGSLNNGSIEEFPGGDVARYLDRVLRVNLIKIFMLIPSKLKHQTTPQISPQLACNRILNQINLFFPKLVAERVCLEKVQQIQLHLEQGLPEFPQACTGTMTIILTLTRKCFVFKLTIVQQGFHQQMGHNATGLIVANSVSQTLVVLLIMKTEDVLEFILRKIPHDIELYT
LAKVRTLASKLAPFRKAVMKVSKHEVCIPYVHILNLAGFNQLEHGLYPQLSAIAILVATVHSCTLGVSPSEWYQTLQEAAECQQGTHLKSKWGWERKRKHWHKCNIHTQQQEMVRARQEILSQLRIRWNMNTIPTEAAERQPLNAQLSKPGKTNHTNNSCQPDPTYQGSPLP
D
>PVIH01028646_Sigmodon_hispidus
FIMMQNTVSTIQRQNAVHAQQGHITPYPKSPTSGFVITFFRGSLSFIFNSLLIHQGIQSITGHDSDDVIPPVFQTRCSGILIVKSVYFLQSGLKLHLTLFRLSFKKSFKTSLRRLGFHCAYALFARILKLCGVNNVEHGWFLHPSATALGVATAHGEPLAGVTICEGYRQSWRETASEFE
HKLQEHSKVQDLNYRPPSKLILLDFHKMQNRKTRRMRSFLPQSTRIFNNQVHTAS
>PVKB01001897_Ctenodactylus_gundi
LYIKYYFEGDIKTFSSTFVKFFTDHRFMEHTIVNALEKAIFVCLILLEKTQEGRHSSQVLSMFFIGTSKLRLGKYIAQSLFPPKLVIGDQACFGKVLWTYVHVEQGFSQFPTAALPGILKVVFIVMRQSFPFKFILMNQGLHKSGHNAADSTIDNEAYEHFSEILIIKSVLDYIQCNVIETH
PLTRRQRLKNVNSFKKATQTTARHDAYALCARLMTVSGIGHFEHDLYPQLCVINLDEGTAQCSLLTDVKVEEQCQVLREAAHKAEAKLQACQEFQEVVKINSTKNTRKLCIHKQDTIKEKQQQDTSSTNLEMEKLKEFNLIHDSDNSSTTLSHDYSIPWKVRGPQENLPESESSS
>JAAGEI010000019_Aeorestes_cinereus
MYNYLTLGQFVLIKSEESRDTIINGSIEQSQFSGFLMVRNVLIYIITNEGGEIKIHLLSANKGKKMAHFAYCVRNLQKHRIYAPFARVLGFPGVTQIEHGLFPWLSAIALGVSSVYQSTLAGVNIDTQYQFLKEAAHQAELELVIHNQREIASG
>PVJN01001708_Lasiurus_borealis
LAKHLNSRQFPQSNNAYCIPNSTFDMYNYLALGQFVLIKSEESRDTIIMGAIEQSRFSGFLMVRNVLIYIITNEGMEIKIHLLSTRKKMAHFAYCVRNLQKHRIYAPFAQVLGFPGVTQIEHGLFPWLSAITLGVSSVYQSTLAGVNTDTQYQFLKEAAHQAELELVIHNQREIAQSR
DLTIEEGGSR
>VMDP01037289_Antrozous_pallidus
LIKSKGSRDAIIAGAVEQSRFSGLLMVKNVLTYIITNEGGEIKIHPLARDKRTREEMARFEYCMRNLQKHGIYATFALVLGLPGVTQIEHGQSPRLSAIALGVPVYQSTPAGVSTDTQYQSLKEAAHQAELEPQAIPNQRE
>JAJQQW010000005_Ia_io
LIKSEGSRDTIIAGAVEQSRFSGLLMVRNVLTYIITNEGGEIKIHPLARDKTREEMARFEYCVRNLQKHGIALFALVLGLPGVTQIEHGQYPRLSAIALGVSSHQSTLAGVHIDTRYQSLKEAAHQAELELQDIHNQRETARSG
>ANKR01209731_Myotis_brandtii
RIKSVKVLDRRGIETRVSQIIECYEASISVDDVCGHLFATLICYHHSGGDVGLMVRSPFGQYLAADGYEVSSVPQELHTQPLYKITCVMGCMERLVNITKDLHPPGPLGVPAAPFMAHLSFFFPTLVTGKGACFQKVEKVFQQLGNQGIKVIARNWMSPTVMKMACRITRHCFIR
GFCLIYHALYHRAGTDAVDAIIVGAVEQSRFSGLLMVKNVLTYIITNGGGEIKIYPLARDKRKREETTRFESCVRNIQKHGIYVPFARVLGLPGMTHIEHGQFPHLSALALGVSSVYQSMLAGVNIDTRYQALKAAAHQAELELQSIHNRREIAQDKDLTAEEREVLFTFNKGNESTR
TTVSVRTKARWSFSKRLATGKNVREMNVKKRRYASSSDESFRPLSPTPPGPPIVSGTTHPEIPAWRQLPQGVLRESSSPGPRPIATPPPPSADTMPELEDLEPPPISIDHSLSSGIGHPSLVSSGDLDELESIQEYDNEGTGRDQRSIPEGFEGKETPEEPREVHSESPSSGTAVNNTQ
TQYMDLSKEGWQILRIPVASISLTTDEEASIMFSTTNRPSNRKDASIYE
>AAPE02051704_Myotis_lucifugus
LKMHSILDAALGLLDSIQPIPPRIKSVKVLDGRGIETRVSQIIACYEASISVDDVCGHLFATLICYYYSGGDVGLMVRSPSGQYLAADGYEVSSVPQELHTQPLYKITCSVGCMERGNITKGLPPPGPLGGPAAPFMAHLSFFFPTLVTGKGACFQKVEKAFQQLGNQGIKVIARNW
MSPTVMKMACRTLRHCFIRGFCLIYHALYHRAGTDAVDAIISGSLEQSRFSGLLMVKNVLTYIITNGGGEIKIYPLAGDKRTREETTRFESCVRNIQKHGIYVPFARVLGLPGMTHIEHGQFPHLSALALGVSSVYQSMLAGVNIDTRYQALKAAAHRAELELQSIHNRREIAQDKD
LTAEEREVLFTFNKGDESVRTTVSVRTKARWSFSERLATGKNVREMSVKKRRMLHRAMKVSALSRPHRQGPPYSQEPPYPEIPAWRQLPQGVLRESSPPGPRPIATPPPPSADTVPELEDLEPPPISIDHYLSSGIGPPSLVSSGDPDELESIQEDGSEVTGRDQHSIPKGFEGKET
PEEPREVHSESPSSGT
>JAPQVT010000021_Myotis_yumanensis
SIQPITAHIKSVKVLDGRGIETRVRQIIACYEASISVDDVCGHLFATSICYYHSGGDVGLMVGSPFGQYLAADEYEVSSVPQELHTLTPLNNMCDGLYGKVSEHHQGSPSPRPPRGPAAPFMAHLSFFFPTLVTGKGACFQKVEKAFQHLGNQGIKVIERNWMSPTVMKIACCTL
CHCFIRGFCWIYHALYLRAGTDAVDAIISGSLEQSRFSGLLMVKNVLTYIITNGGGEIKIYPLAQDKTREETTRFESCVRNIQKRGIYVPFARVLGLPGMTHIEHWQFPHLSALALGVSSVYQSMLAGVNIDTRYQALKAAAHQAELELQSIHNRREIAQDKDLTAEEREVLFSFNKG
NESIRTTVSVRTKARSFSERLATGKNVRVDERVEKEDVSSSDESFSPLSPTPPGPPIFSGTTLP
>JAPQVU010000012_Myotis_yumanensis
SIQPITAHIKSVKVLDGRGIETRVRQIIACYEASISIDDVCGHLFATSICYYHSGGDVGLMVGSPFGQYLAADEYEVSSVPQELHTLTPLNNMCDGLYGKVSEHHQGSPSPRPPRGPAAPFMAHLSFFFPTLVTGKGACFQKVEKAFQQLGNQGIKVIERNWMSPTVMKIACCTLC
HCFIRGFCWIYHALYLRAGTDAVDAIISGSLEQSRFSGLLMVKNVLTYIITNGGGEIKIYPLAQDKTREETTRFESCVRNIQKHGIYVPFARVLGLPGMTHIEHWQFPHLSALALGVSSVYQSMLAGVNIDTRYQALKAAAHQAELELQSIHNRREIAQDKDLTAEEREVLFSFNKGN
ESIRTTVRVRTKARSFSERLATGKNVRVDERVEKEDVSSSDESFSPLSPTPPGPPIFSGTTLP
>ALWT01149584_Myotis_davidii
KSVKVFDGRGIETRVSQIIKCYEASISVDDVCGHLFATLICYYHSGGELGLVVRSPFVHFLWQTGMKFLVCLKSYIHNLLYKITYAMGCMERLVDITKDLPPPGPLGGPAAPFMAHLSLFFPTLVTGKGACSQKVEKAFQQLGSQGIKVIARNWMSPTVMKIACRTLHHCFIMGFC
LIYHALYHRAGTDAVDAIIVGAAEQSRFSGLLMVKNVLTYIITNGGGEIKIYPLAREKRTREMTRFESCVRDIQKHGIYVPSERVLGIPGMTHIEHGQFPHLSALALGVSSVYQSMLAGVNIDTRYQARKAAAHQAELELQSIHNRREIVQDKDLTTEESEAPFAFNKGNESIRTTGS
KAQVELLKKISDREVKQSAASSRKGGCFIERKFPPSLAHTARAPHILRNHPTPKYQRGDSSRGVLESSPPGPRPITTPPPPSADTMPKLEDLETPPVSIDHSLSSGIGHPGWAGSGDPDELESLQEDDSEATVGDQHSIPEDFEGKETPEEPWEVHSESPRSVT
>PVIZ010081685_Myotis_myotis
KSVKVFDGRGIETRVSQIIECYEASISVDDVCGHLFATLICYYHSGGDFGLVVRSPFGQFLWQMGMKFLVCLKSYIHNPFIKNMCDGLYGKVSEHHQGSPCPRSPRGTCCSLVVHLSLFFPTLVTGKGACSQKVEKAFQQLGDQGIKVIARNWMSPTVMKIACHTLHHCFIMGF
CLIYHALYHRAGTDAVDAIIVGAVEQSRFLGLLMVKNVLRYIITNGGGEIKIYPLARDKRTRGEMTRFESCVRDIQKHGIYVPSARVLGLPRMTHIEHGQFPHLSALVLGVSSVYQSMLAGVNIDTRYQALKAAAHQAELELQSIHNRREIVQDKDLTTEESEVLFTFNKGNESIRTT
GSKAQVELLKKISDREVKLSPRVSVKKRRMLHGARKVSAFSRPHRQGPPYSQEPPYPEDTSVETTPSGVLRESSPPGPRPITTPPPPSADTMPELEDLEPPPISIDHSLSSGIHPGWVGSGDPDELESIQEDDSEATGGDQHSIPGDFEGKETPEEPWEVHSGSPSSVTAVNNTQT
QYMDLSED
>JAPDVT010000001_Corynorhinus_townsendii
LKDPIKGYILEEHEISFSIKMHSILDAALGPLDSSQPVPPRINSIKVLNGRGIEARISQIIACFEAGVGMDDIEHLYATLICYCHYRGDLNLMKSPLGQYLIADGYEFSTTPLDVHTQPLHRIMRAIGCSVALVQIIRDLPVPNPVSKRAASFMAYLNLFLPKLVTGGSACSQKVEKAYQKL
ENRGINVTSRDWMAPVVMRIAYRTLRHCFVIRFSLICHELHQQAGTGAGKTIIVGAAEQPRYSGLLVVKNVLTYMITNEEGDIKIHPPAQDKWTREELARLEYCVRNLQEHGICVPLGRVLGLPGVTQIEHGQFPHLSAIALGVALVYQSTLAGVSTEPRYPSLKEAAHQAELELQ
SIRNTRELPAAEISPSNNEGYYLSSLGEVKKINLDQRREDRARAGLLRRLTEREIGPSPQDEHRLGRGEMPPSDGRSHSLSPQPPVSYQPHGSPYSDLP
>JAPDVU010000001_2_Corynorhinus_townsendii
LKDPIKGYILEEHEISFSIKMHSILDAALGPLDSSQPVPPRINSIKVLNGRGIEAHISQIIACFEAGVGMDDIEHLYATLICYCHYRGDLNLMKSPLGQYLIADGYEFSTTPLDVHTQPLHRIMRAIGCSVALVQIIRDLPVPNPVSKRAASFMAYLNLFLPKLVTGGSACSQKVEKAYQK
LENRGINVTSRDWMAPVVMRIAYRTLRHCFVIRFSLICHELHQQAGTGAGKTIVVGAAEQPRYSGLLVVKNVLTYMITNEEGDIKIHPPAQDKWTREELARLEYCVRNLQEHGICVPLGRVLGLPGVTQIEHGQFPHLSAIALGVALVYQSTLAGVSTEPRYPSLKEAAHQAELEL
QSIRNTRELPAAEISPSNNERYYLSSLGEVKKINLDQRREDRARAGLLRRLTEREIGPSPQDEHRLGRGEMPPSDGRSHSLSPQPPVSYQPHGSPYSDLP
>JANZWY010000002_Eptesicus_fuscus
QIIACFEAGVSIDDECEHHVCHIDLLLPLSPFGQYLVADGYEFSTVPPDLHTRPLHRTMRATGCSEALVQIIRDLPAPNPVGKRAASFMASLSLFLPKLVTGGSACSRKVEEAYQQLEDRGIHVISRNWVAPVGMRIAYRTLRHCFTRRFSLIHHELHQRAGTGAVKTVIAGAVEQP
RYSGLLMVKKMFHVITNEEGDIKIHPAARDKRTREALARLEYCVRNLQKHGICAPFARVLGLPGVTPIEHGQFPRLSAIALGVASVYQSTLAGVSIDPRYQSLKEAAHQAELGLQSIRNTRKLPEAEISPSKNEGYCLSSLGEVKKINQDQRQREGRARAELLRRLAEGEMGPSPQD
EHGLGRGEMPPSHRHWHSPSPQPPVSYQPHGSPCSDLPDLETCPPAVNWGILSTNTSPGNTTTALQGYHACSRRLTSRVDINDYHDSSGHRGPGGPEDSVELASIQEEDSDIINGEQYSIPEFFLG
>CATOCW010000113_Eptesicus_nilssonii
SLFLPKLVTGGSACSQKVEKAYQQLEDRGIHVISRDWMAPVVMRIAYRTLRHCFIVRFSLISHELHQRAGTGAVKTIRAGAVEQPRYSGLLMVKTVLAYIITNEEGDIKIHPPARDKRTREALARFEYCVRNLQKHDICAPFARVLGLPGVTQIEHGQYPPLSAIALGVASVYQSTLA
AVSIDPGYQSLKEAAHQAELELQSIRNTRKLPEVGDLAVEERGVLLKFTWRRKKINQDQRQQEDRARAELLRRLTEREIGPIPQDEHRLGQGEMPPSHKHWHSPSPQPTASYQPHGSPYSDLP
>ANKR01300264_Myotis_brandtii
SSSIKMHSILDAASGPLYSSQPIPPRIKSIKVLDGRGIEAGIGQIIACSEAGVSIDDICEHLYATLICYCHSRGDLNLMVKSLFGQYLVADGYEFSTMPPDLYTQPLHWIIRAIGCSEALVIIRDLPVPNPVGEGAAPFMAYLSLFLPKLVTGESACSQKEEKAYRQLENRGINVISREWM
APVVMRMAYRTLGHCSIIRFSLIYHELHQRTGTGAVKTIVAGAVEQSRFSGLLMVKNVLTIITNKEGDIRIHPLARDKWTRGELACFAYCVRNLQKHGIDAPFARGLWLPGVTQMEHGPFHRLSAIALAGTSVYSTLAGVNIDPRHQSLKEAAHPAELKLQSIHNARKLPEVEISS
WKNERDYLSLLGEVKKINQDQRQQEERAGAELLGKLTEREIGPSPQDEHRQGRGEMPPSDGSPWSPSPQPPVSYRPPGSPYSGLPIWRCVHQQLIGESSPPTPRPVRTPQPPSRGTSLLSRTLNLLGLISMITSKTTPDIAPG
>PVIZ010018957_Myotis_myotis
SSSIKMHSILDAASGPPYSSQPIPPRIESIKVLDGRGMEAGVGQIIAFGSVSIDDTCEQLYATLICYCHSRGDLNLMVKSPFGQYLIADGYEFSTMPPDLHTQPLHRIMAIGYSEALVQIIRDLPAPNPVGERAASFMAYVSLFLPKLVTEESACSQKVEKAYRQLENRGINVISRERKA
PVVMRMAYRTLGHCSIMRFSLVYRELHQRAGTGAVKTIIAEAVEQSRFSGLLMVKNVLTIITNKEGDIRIHPLARDKRTRGELDSFAYCVRNLQKHGINAPFARVLGLPGVTQMEHGPFHRLSAIALGVSSVYQSTLAGVNIDPQHQSLKEAAHPVEPGLQSIRNARKLPKAEISP
WKNERDYFSSLREVKKINQDQRQQEERAGAELLRRLTERETGPGPQDERLLGRGETPPSVSPSPQPPVSYRPPGPPIQAYQ
>JAPQVT010000001_Myotis_yumanensis
SSSIKMHSILDAASGSLYSSQPIPPRIESIKVLDGRGIEAGVRQIIACSEVGVSIDDICEQLYATLICYCHSRGDLNLMVKSPFGQYLVADGYEFSTEPPDLHTQPLHRIMRAIGCSEALIQIIRDPPVPNPVGERAASFMAYLSLFLPKLVTGESACSQKVEKAHRQLENRAMNVISRE
WKAPIVMRMAYRTLGHCSIIRFSLIYHELHQRAGTGAVKTIVAGSVEQSRFSGLLMVKNVLTIITNKEGDIRIHPLANKRTRGELARFAYCVRNLQKHASYAPFARGLGLPGVTQMENGPFPRLSAIALGVASVYQSTLAGVNIDPWHQSLKEAAHPAELELQSIRNARKLPEVEIS
PWKNERDYLSSLREVKKINQDQRQQEERAGAELLRRLTEREIGPSPQDERRQGRGEMPPSDGSPQSPSPQPPVSYRPPGSHYSGLPIWRRVHQQLIGKSSTPKPHPVRTPRLPSSDTMPALEDSEPPRVDINDYLQDSSGHSPRTRSSVELAIIQEEESKIINGEQHSIPEFFGKNR
RATRGASRAIGPGYITTLEHVRCTRFFADDRTNGSKDTRGSVTPSGVEKCHGTDS
>JAPQVU010000003_Myotis_yumanensis
SSSIKMHSILDAASGSLYSSQPIPPRIESIKVLDGRGIEAGVRQIIACSEVGVSIDDICEQLYATLICYCHSRGDLNLMVKSPFGQYLVADGYEFSTEPPDLHTQPLHRIMRAIGWSEALIQIIRDPPVPNPVGERAASFMAYLSLFLPKLVTGESACSQKVEKAHRQLENRAINVISREW
KAPIVMRMAYRTLGHCSIIRFSLIYHELHQRAGTGAVKTIVAGSVEQSRFSGLLMVKNVLTIITNKEGDIRIHPLANKRTRGELARFAYCVRNLQKHASYAPFARGLGLPGVTQMENGPFPRLSAIALGVASVYQSTLAGVNIDPWHQSLKEAAHPAELELQSIRNARKLPEVEISP
WKNERDYLSSLREVKKINQDQRQQEERAGAELLRRLTEREIGPSPQDERRQGRGEMPPSDGSPQSPSPQPPVSYRPPGSHYSGLPIWRRVHQQLIGKSSTPKPHPVRTPRLPSSDTMPALEDSEPPRVDINDYLQDSSGHSPRTRSSVELAIIQEEESKIINGEQHSIPEFFGKNRR
ATRGASRAIGPGYITTLEHVRCTRFFADDRTNGSKDTRGVTPSGVEKCHGTDS
>AAPE02007767_Myotis_lucifugus
SSSIKMHSILDTVSEPLYSSQPIPPHIESIKVLDGRGIEAGVGQIIACSEAGVSIDDICEQLYATLICYCHSRGDLNLMVKSPFGQYLVADGYEFSIMPPDLHTQPLHWIMRAICCSEALVQIIRDLPVPNPVGEGAAPFMTYLSLFLPKLVTGESACSQKLEKAHRQLENRGINVISRE
WKVPVVMRMAYRTLVIIRFSLIYHELHQRAGTGAVKTIVAGAVEQSRFSGLLMVKNVLTIITNKEGDIRIHPLARDKRTRGELALFAYCVRNLQKHGIDAPFARGLRLPGVTQMEHGPFPRLSTIALGGTSVYQSTLAGVNIDPRHQSLKEAAHPAELGLQSIRNARKLPEAEISPW
NNKRDTVHSEKWKKSIKIKDSRKRGLRAELLGRLTEREIGPSPQDERRQGRGETPPSDGSPQSASPQPPVSYRLPGSPYSGLPIWDVSTSSLGNPLHQHLARLEHHDRPPGTPCLLSRTLNLLGLISMITSKTAPDIAPG
>PVJC01085686_Murina_aurata
HSILNDALGPMDSVQFIPPRVNNIKVLDGKGLEARVHQIIECCEAGASVHAREHLLATLCCYHYRGDFGSMIRSPFGQYLAADGYELSSKSAGTTRAALTPDRAAIGCMEGFVPAIKDLPPPKLLGKCAAFFMARVSLFLPKLVTGEGACAPKVEKAFPQLDTQGISIVSRSWMSP
TVMRFAHQTLGLCLTILSSLIYHGTPHQRAGSDALDTVMAGAVEQSHFSGLSMASNVLPYIVTKEGGAIEIHPAARRRRTKEEMARFESCVPNIQTHGIYAPFARVLGPPGVAQIEHGPFPRRPAIALKIPTVSHSTPAGVNTDTWYQAL
>ANKR01230743_Myotis_brandtii
GMTLLTMHSILNAELGPMNVQFIPPRIKSIKVLVGKGLERMVRQIIECCEAGMSVDDACEHLFATLLCYYHYGGDLGVMMRSPFGQYLAADRYERSSISQELHEHPLHRLMRAIGCMEGFVRVIKDLPPPKLLGKRAAFFMAYLSLFLPKLVTGEGACAPKVEKAFQQLETQGIN
IISRSRMSPMIMRIAYRTLGHCFIVRFSLIYHVLHQRAGTDAVDTVMAGAVEQSRFSGLFMAKNLLTYIITNEGGAIKIHPLARDRWTKEEMTPCESCVRNIANHGIYAPFARVFGLPGVAQIRHRPFPRLPAIALGVSAVHQSTPVGVNTDTRYQAFKETAHLAQLEQGILERQEI
IQDKDLSAEECEVIITFKKRSERINQTQRQQEAKARSELLKDIMDRVSQIKSIKIQQQQEQEERLPRSESPHLPTPPAPCPSCYQGPPPPEVPTWRPPPSRYIGESPPRILHSAALPPPSSRGLKPESKEPQLPPIDVFDFLDSEYEHQAPVGSGEPGELDSMPEESSDILSREPHSIPQG
FEGEEILDEPPDEEPNSPTHSPAVLSCSL
>JAPQVU010000020_Myotis_yumanensis
GMTLLTMHSVLNAELGSMNSVQFSPPRIKSIKVLDGKGLERRVRQIIECCEAGMSVDDVCEHLFPTLLCYYHYRGDLGVMMMSPFGQYLAADGYELSSISQRTTEHPLYRLMRAIGCMEGFVVIKDLPPPKLLGKCAAFFMAYLSLFLPKLVTGEGAYVPKVEKAFQQLETQGIN
IISRSRMSPMVMRIAYQTLGHFIICFSLIYHVHPIRDRGLVDTVMAGAVEQSRFSGLFMAKNLLTYIITNEGGAIKIHPLAWDRWTKEEMTPCEFCVRSIQNHGIYAPFARVLRLPGVAQIRHGLFPRLPAIALGVSAVHQSTLSGVNTDTRYQAFKETAHLAQLEQGILERQEIIQD
KDLSAEEREVITTFKKRSERTNQTQRQQEATACSELLKDIMDRAAKLSRIKIQQQQEQEERLPRSESPHLPTPPAPCPSCYQGSPPPEVP
>AAPE02014310_Myotis_lucifugus
GMTLLTMHSILNAELGPMNLQFIPPRIKSIKVLVGKGLERMVRQIIECCEAGMSVNDFCEHLFATLLCYYHYRGDLGVMMRSPFGQYLAADGYEHSSISQELHEHPLHRLMRAIGCMEGFVVIKDLPPPKLLGKRAAFFMAHLSLFLPKLVTGEGACAPKVEKAFQQLEIQGINII
SRSQMSPMVMRIAYRTFGHCFIIRFSLIYHVLPSESGDAVDTVMASAVEQSRFSGLFMAKNLLTYIITNEGGAIKIHPLAQDRWTKEEMTSCESCVRNIPNHGIYAPFARVLGLPGVDQIRHGPFPHLRAIALGVSAVHQALQRVSTDTRYQAFKETAHLAQLEQGFLERQEIIQD
KDLSAEERGVIITFKKRSERINQTQRQQEAKACSEVLKDIMDRAAKLSPQNPTTAGTRRGLPRSENPHLPTPPAPCRSCYQGPPPPEVPTWRPLPSRYIGESPPRILHSAALPPPSPRGLEPESKEPQLSPIDVFDFLDGEYEHQAPVGSGEPGELDSMPEESSDILSREPLGTSLHSP
SFEGEEILDEPPDEEPNSLTELRPNS

>ALWT01026193_Myotis_davidii
HSILNAALRPMDSVQFIPPHIKSIKVPDGKGLERKVRQIIECCEVGMSVDDAGEHLVAMLLCYYHYRGDYLGLMMSSPFGQYLTADGYELSSMSEPHEQCLHRVMRAIGCMEGFVVFKDLPPQSLENNAAFFMAYLSLFLPKLVTGEGTCAPKVEKAFQQLETQGINIISRSRM
SPMVIRIAYRILGHCFIIRFSLVYHVLHQRAGTDAVDTVMAGAVEQSRFSGLLMAKSVLTYIITNEGGAIKRHPPAQDRQTKEEMARFESCVRNIPNHGIYAPFARVLGLPGVARIEHGPFPHLPAIALGVSAVHQSTPAGVNTDTQYQALKEAAHQVELEQGILKRWEIIQDKAL
SAEEREVIITFKKRSERINQTQRQWEAKAHSELLRDITDRAAKLSPQNPTTAGTRREVTPQESSLTHSCTTPVSLSRTPPTVPTWRPPPSRSIGESPPHILHSAALPPPSPRGLEPESKEPELPPIGVFDFLDSEFEHQAPVGSGEPGELDSMSEEGSDTPQPPQPGTSLHSRRFRRED
SP
>PVIZ010007745_Myotis_myotis
HSILNAALGPMDSVQFIPPHTKSIKVLDGKGLERKVRQIIECCEAGMSVDDAGEHLVAMLLCYYHYRGDYLGLMMRSPFGQYLAADGYELSSVSQELHEHPLHRLMCAIGCMEGFVVIKDLPPQSRENNAAFFMAYLSLFLPKLVTGEGACAPKVEKAFQQLETQGINIISRSRIS
PMVMRIAYRTLGHCFIIRFSLVYHVLHQRAGTDVVDTVMAGAVEQSYFSGLLMAKNVLTYIITKEGGAIKRHPPARDGQTKEEMARFESCVRNIQNHGIYAPFARVLGLPGVAQIEHGPFPRLPAIALGVSAVHQSTPAGVNTDTRYQALKEAAHQAELEQGILKRREIIQDKDL
SAEERELIITFKKRSERTNQTQRQREAKARSELLKDIMDRAAKLSPQNPTTAGTRREVTPQWSPHSPTPPAQRPSPYQGPPPTEVPTWRPPPSRSIGESPPRILHSAALPPPSPRGLEPEFKEPQLPPIDVFDFLDSEYEHQAPVGYGEPGELDSMPEESSDILSWEPHSIPQGFEGK
EILDEPPDEEPNSPTSPSSLTELRP
>AXCS01020818_Nannospalax_galili
KSKLVRVFTGDGVKQIIQQILASFEAGLEIDNLADHFFATIVCYYHYRNNLDLMVRSSFGQYLVENGYEFSGLELDNKREPLLQHLEELGVPEGLLKVLGDLPCAPPLGLRPAAFLVYLSLFLPKLVAGTTACIKKVERTYQQLEIQGLNIIPREWMTPVMMNLAYRALRLCFFIRFSII
HHVLHQRAGIDAADTIMAGAVEQTRFAGLLMVRNVLTYIITKGKEKDEYHIHPLAQDKRMREEITRFEYCIQNIQRHGIYAPFARVLGLAGVSQIEHGQFPRLSAIALGVSSVYQSTLSGVNMDTRYQSLKEAAHQAELELQAIQYKKEIRQDTDLDNDERVVLLKFAERSGNIIKE
QRQQENLARQELLQKFTEKAETS
>SRR16768455_Eospalax_fontanierii
MHSLLNATLGNADDDQCAPPQSKLVRVFTGDGVEQIIHQILASFEAGLAIDNMTDHFFATIVCYYHYRNSLDLMLRSSFGQYLVANGYEFSGLELHNRGESLLQHLKELGMPEDQLKVLGDLPSPTPLGLRPAAFLVYLSLFLPKLVTGTVACIKKVKRTYQQLESQGINTIPRGW
MTPVMMNLTYRALRLCFFIRFSIIHHVLHQRAGIDAVDTIMAGAVEQTRFAGLLMVRNILTYIITKGNEKDKYHIHPLAQDKRMREEVTRFEYCVKNIQRHGIYAPFARVLGLAGVSQIEHGQFPRLTAIALGVSSVYQNTLLGVNMDMRYRSLKEAAHQAELELQAIQCKKIRQ
DTDLDDDERVVLLKFAERSQSINKEQWHQEGLARQEPLQKLTGREGGTSSQGAGPTEPSVVRSGLPGDCESISNTERLDSEKGQPRATSPQLHHTPELTSAEPAVASEPKNQPTSRLPGNTSPGSCAEELVFNGDATLPSDIASRDSSEEEPPQSDQLSDSQADFSTAGPDLLAIP
EGLEPDQPVLPTRDSEEVLPGEAPAVFPGH
>PVIM010120482_Rhizomys_pruinosus
HSLLNGALGDVEDTPSQSKMVRVFTGDGIEQIIQQILASFEAGLAIDNLVDHFFATIVCYHHYRNNLDLMVRSSFGQYLVANGYEFSGVELHSRKQTFLQHLKELGMPESTLKVLENLPSLPPLSLRPAAFLVYLSLFLPKLVTGTSASIKKAERTYQQLEGQGLSIIPSGWMNPTM
MHLTYRTLRLCFFIRFSLIHHVLHQRAGIDAVDTIMAGAVEQTRFAGLQMVRNVLTYIITKGNESDKYNIHPLAQDNRMKEELTRFEHCVKNIQRHGIYAPFARVLGLAGVSQIEHGQFPRLTAIALGVSSVYENTLSGVNLDTRYHYLKEAARKAELELQAIQHKKAIRQDTDLDE
DERAVLLKFAERSGNINKEQHQEAFTRELLQLTQNEAGTCLQGVGPAEP
>AXCS01123754_Nannospalax_galili
QVLGLAGVSQIEHGQFPRLSVIALGVSSVYQSTPSGVNMDTHYQSLKDAAHQAELELQAIHYKKIRRDTDLDNDERVVLKFAERSGNIIKEQQEEDLGRQELLRKFTEKAETSLQGARAAEPSVARTELLGNLESIPSTERLDNQERQPQATNLPFTAPLEQHLKSQPAASKPKNQ
QTSRLRGSVVEKLAFYIDALLPSNNESGDSSEESQSNQLSDSKEDFSTAGPDQSEIPGELEPSQPDQLTPITEEVPLGEGPGVFPSHLGTMSPMGNLQPPPGEKPSCQQGSGTMKEGHLPIQGASKTQKPRN
>AXCS01204846_Nannospalax_galili
KSKLIRVFTGHGVEQIIQQILGSFEAGLEIDNLADHSFATIVCYYRHRNSFDLMVRSSFGQYLLKHGYKFSRELDNQREPLLRHLWDLGVPEVLLKVLGDLPCAPPLGLRPAAFLVYLSLFLPTLVARTTACVENVEWTYQQLESQGLSIIPREWMTPVMMNLARARQLCLFIQFSLT
HHVLHQRAGIDVVDTITAGAVEQTPFAGLLMVRNVLTHIITKEKGKEEYHIHPLAQDKRMREEVISFEGIQNIQRHRIYALFAQVLGLAGVSQIEHGQFPRLSVIALGVSSLYQSTPSGVNMDTHYSLKDAAHQAELELQAIHYKKIRRDTDLDNDERVVLKFAERSGNIIKEQQEE
DLGRQELLRKFTEKAETSLQGARAAEPSVARTELLGNLESIPSKERLDNQERQPQATNLPFTAPLEQHLKSQPAASKPKNQQTSRLRGSVVEKLAFYIDALLPSNNESGDSSEESQSNQLSDSKEDFSTAGPDQSEIPGELEPSQPDQLTPITEEVPLGEGPGVFPSHLGTMSPMGN
LQPPPGEKPSCQQGSGTVKEGHLPIQGASKTQKPRN
>PVJC01021378_Murina_aurata
QLIPPHIKSVKVLDRRGIQTRVGQTIECYEAGISVNDVCEHLFATLICYYHYGGVLGLMVRSPFRQIFSNVTQELHTQPLYKITCVMGCMEGLVNIIKDLPPAGPLGRPAASFMAYLSLFSPLLVTGEGACSQKVEKAFQLGIKGINMTAINWMSPIVMKIAYRTLHHCFIMQFCLIY
HGVYQRAGTDAVDAIIAGAVEQSRFLGLLMVKNVLTYITTNGGGEIKIHSLAQDKRAREEMTHFESWVRNIQKHGIYAPFARVLGLPGVTLEHRQFPHLSALALGVSSVCQSTIARVNIDTRYQALKEAAHQDEFKLSIHNRREIAQDKDLTAEEHEVLFTFNKKSERIHQDQRQC
EDKAQVEFIKKITDRVSETESARDERQVEKEDASSSKKSFHTHSLTPQGPPYSQEPSYLEIPVWRQLPQGVLRGSSPPVPRPMATPPPSSAHTMPELEDQNLHPLALTILSLVNGHTVSVGSGDPDELQRAYKKTIMKPPNGDQHSIPKGFEGEETPEEPWEVHSESPSSGT
>JAPDVU010000004_Corynorhinus_townsendii
HFPQSYNAYCIPNLTFDMYNNFALGQFVLIKSEGSRDTIISGAIEQSQFSGLLMVRNVLTYMITKGWGDKNRDKQTREEMSIEYCVRNLQKDRIYAPFALVLGLPGVPNIEPGQFPRFSVIALGVSLVYQSTLARVNIDTRYQPLKEAAHQAELELQAIHNQREI
>JAPDVT010000001_1_Corynorhinus_townsendii
ECEAGISVDDACGHLFATLICYYRYLGDLGLTMRSPFGQYRAADGYDFSSMSQERPEQPFQRLMRAIGCVEGFVCEIEDLPPAKPLGKRAAFFTAYLNVFLPKLVTGEGACAQTVREGLSTVGNTGINVISRSWMSPMAVRIAYRTLRPCFITRFSLIYHLLHRRAGTDAADTIMA
GAVEQSCFGRLMVKNVLTYVMTNEGGAIKIHPPARDRRTKEEMAPFEYCVGNIQKHKIYAPFARVRRLPGVAQIELGQLPHLFVVALGVSTVYRSTRAGVNTDTRYRALREAAQTELEQLHQRRDISQDKNLSAEEREVMITFQKRSERINPTQRQQEGKARSQLLKHILDRAA
KSPQIQTAGARRVGHRHSESPHSPTPPAQRPSHDQGPPPPEAPTGRPPPPRSMGESPPRILHLAALPPPSSRGLEPESKEPPPPIDLLDFLDGKYDHQAPVGEPEELDSTLEKWRDSLSCEPHSIPGGCEGEEILQPPPMKNLNPQLTAQQFNG
>VMDP01006281_Antrozous_pallidus
YILEEISSSIKMHSILYAALGPLDSRQPISSRIKSIKVLSGRGIEAHITSAKSLVLRRVLAMMYVSICMPHSAIRLRGDLNLMVKSSFGQYLVADRYFSTMPPDLHTQPLQKIRLAIGCSEALVQIFRDLPAPNPVGKGAGLFLAYLSLFLPKFVTEGSACSQKVERTYQQVEDGTNVISR
NWMAPVGMTIAYCTLRHCFIIHFSLIYHELQAGTGAVKTIIAKAVEQPRYAGLLMVKNVLTYIITNEEGDIKIHPPANKRTREELAHFEYCVRNLKHGIYALFSQVIGLPRVTQIEHGQFSHLSAIALGVALHFGLSSTLAGVSIDAQYQSLKEAAHQAELELQSYPYKFRKLPEAKISQ
WKNEGYYLNSLGKVKKEINQDQREQEDRAQAELLRRLT
>JAPDVT010000005_Corynorhinus_townsendii
VRLKFISRDALDKHPRSFPGIYLGHTIIAGAVEQSKFSGLLMVKYVLTYIINNEGGGIKIHPLADKTREYFEYCIRNIKKHGSYMPFARVLGLPGVSQIEHGALPSLSAIALGVSSVYQSTLAGVNIDTRYQALKAAAYQAELELQRIHNKREIAQDKDLTAEEHEVLFTFSKRSKRIHQD
QRQHEDKARVELLEKITDREVELNPKDERQVEKEDASSNGKSFCPPSPTPSGPPYSQEPSCP
>JAJQQW010000017_Ia_io
LKFISRNSLDKHHRSFPGIYLGHTIIAGAVEQSRFSGPLMVKNVLIYIINNEGGGIKIHPLAQDKTREEMTHFEYVRNIQKHGSYMPFSRVPGLPGALEHGQFPSLSAIALGVSSVYQSKLAGLNTPYQALKAAAHQAELELQSIHNKREIAQDKDLTAEEREVLFTFSKRRKKIHQDQ
RQREDKARVELLKKITDREVELNPKDENQIEKEDASSSDKSFCPPSPTPSGPPYSEPSCP
>JANZWY010000012_Eptesicus_fuscus
WSTSNYLNVPTPARLKFISRNSLDKHPRSFPGIYLGHSIIAGTVEQSRFSGLLMVKNVLTYIINNEGGGIKMYPLAQDKTREEMTHFEYVRNFQKHGSDTPFSRVLGLPGATQIEHWQFPSLSAIALGVSSVYQSKLAGLNTPYQALKAAAHQAELELQSIHNKREIAQDKDLTAEE
REVLFTFSKRSEKKSIRIKGSVRTKLRVELLKKITDGEVELNPKDESQIEKEDASSSDKSFHPPLPTPSGPPYSEPSSSRNTSM
>CATOCW010000765_Eptesicus_nilssonii
FGPRPALGPACVEPLHLLECPNTCAVKVYLKELLRTSQIPFPGIYLGHSIIAGTVEQSRFSGLLMVKNVLTYIINNEGGGIKIHPLAQDKTREEMTHFEYVRNIQKHGSYTPFSRVLGLPGATQIEHGQFPSLSAIALGVSSVYQSKLAGLNTPYQALKEAAHQAELELQSIHNNKREIA
QDKDLTAEEREVLFTFSKRSEKKSIRIKGSVRTKLRVELLKKITGREVELNPKDESQIEKEDASSSDKSFHPPLPTPSGPPYSNHPVPKYQYGDNS
>VMDQ010002737_Nycticeius_humeralis
HGLYLGHTIIVGAVEQSRFSGLLIVKNVLTYIINNEGVKRYTHPLAQDKRMREEMTHFEYCVRNKKHGSYMPFARVIGLPGAIQIEHGQSPSLSAIARGVSSVYQSTLAGVNIDTEYQALKAAAHQAELELQSIYNKREIAQDKDLTAEELSVKGVKEFHQDQRQCEDKALVELLKKI
TDKEVELNLKDESQIEEDPSSSDKSFRPPATTLSGPPSQESSCPEIPLWRQLPQGVL
>ALWT01147311_Myotis_davidii
GVSQIIACFEAGVSIDDICEQVYATLICYCHSRGDLNLMVKSPLGQYLIADGCECSTMPPDLHTQPLRRIMAIGYSEDLVQIIRDLPGPNPVGKRAASFMAYVSLFLPKLVTGESACSQKVEKAYRQENRGINVISGEPKAPVVMRMAFRTLGHRSIMRFSLVYRELHQRAGTGAL
KTIIAEAVEQSRFSGLLMVKNVLTLQTRKGHQIHPLADKRTRGELARFAYCVRNLQKHGIDAPFARGLGLPGVTQMEHGPFPRLSAIALGVASVYQSTLAGVNIDPRHQSLKEAAHPAELGLQSIRNARKLPEAEIAPWKNERDYLSSLGEVKKSTRSKTAGREGWGRTIEKTYRK
ENRPKPAGTSTRQRRDASKRISITTATSILSASWSHYSGLPIWRRVHQQLIGESSPPTPRSVRTSPPPSRDTMPALKDSEPPRVDIDDYLQDSSETQTPGRPEDSVELASIQEEESIINEEQHSIPEFFEGERTAEEPQGGHPEPLGLDIFEQPYESMSDTQELLQMAGPMVAE
>PVJC01036705_Murina_aurata
SSSIEVHSILDAALGPLDSSPIPPPIRSIQVVDGRGKEAGVGQITACFEAGVSTDDICEHLYDTLVCYCHSRGDHQLMGKSPFGPYLVVGGYEFPKMPLDLYSQPLHRIMRATDGSEALAQIIADLPVPNPVGMRAASFMAYVSLFLPELVTGESACSQERPANNRISVISREWRTC
GDENRYRTRHHCFTRRFSLIYQELHQRPGTGASKTIVAGAVEPSRFSGLLVVKNVLTYTMTNKEGDIKIHPLAGDKQTREELARFAYGVRNLQKHDICAPFARVLGLPGVTQGGHGPFLRVSALALGVASSTEAHSQGGDIDPRHQSPKEVAHHAELELRGYRSARKLPAVGDLA
VEEREGLLRFPRRSEKINQDQRQHESAGAELSRRLPEREIGPSPHHEHPLGRGDQSPRSPSPPPESYRPHGSPYSGLPVWRHVHQIGESSPPTPGRVRTPPPSRDTTCSGAQETEPGLQTDENVESLYLPVSAPSSGSPSEKQSRPSAEFNIRGQL
>VMDQ010169293_Nycticeius_humeralis
VSQIIECYEASIKMMPGNDLFETLIHYYHYLGDLDLMIRSPFGQYLVADGYEFSSMSQEPLQRVMRAIGCMGRALHKWRKRAAFFMACLHFFLPKWVTGGRACTQKGEKAFRLETQGSNIASRSWMSPMVTGISSRCFITCFSLTDHVLHRKAGPDAVDMILAGAVEQSRFS
GLLTVKKCVHMHHDKRRGITHPPARDRRKKEGMDRLAYCVQNIPKHGELAPFARVRRLPGVAQVELGQFPHLFAIALGVSAVNQSTGAGVNVDTGYQALKGEAAHQAELEGQGLHEREIIQDKALSAEEREVIITFKKRSQ
>VMDP01004410_Antrozous_pallidus
LSKGMPIKHPGICLGHTIIVGAVEQRFSGLLLVKKKKNVLTYIINNEGGGIKIHPLAQDKTGGETTPFEHCVRNIQKHGSYTPFASVLGLPGATQIEHGQFPSLSALALLGVFSGYQSQLAGVNIGTPRALKAAAHQAEFRITGHPQQREIAQDKGFTAEEREVLFSFQKKRSERIHQ
DQRQLAVRTKALGGASQIKITDREVELNPKDEHQIEKEGASSSDKACPPSLAHTIKAFLFGTLRSRNTGMETTPSRSITRMISSLRLGQLPPPSADTRPELEDLEPPTHQHGLSLSWYQAPSSDHSGDQDELESIQEDDSDATDGDQLSVPVGFESKETPAGPREPHSRVSKQRHC
RLTHPQTPQHGPSRVI
>JANKKK010000001_Dipodomys_merriami
HSIPKAELAQIQLVPPCVMGACILCGKGIGLRGAICFEAGLVPDIFPFFATLLCYYYKGDAALMRKSLFGTHLEGEFGELFAAQDDPFHRVVSLVCPEGTIEIIKLPPPALHGSHIPAFMVHLTLFPPKLVMGEAACSQKVEPSYQQLEAWGYHNPSSWMVANAIRSVYRAVRLCYLI
CFCLIDHVLHQSARTDEVDDIIARSVEQSHFFCFFLVCSVLTYIIAGESGTIKIHPLARDRWVKEEVDHFEHCVNNLHKHGIYTHFPFLGLGVFQIEHGQFPCLSMIALGVLSVYQSTLAGVNADARYQALKKVAHKAVLKLQACNDKKEIQGEGGSVIEPCMLETLGSIPQHHISKK
KKVGSGAVAQVVERPAEELRDSAQDQSSGPMTGPKKKKQTNKEGNSNLPRSISRTSDHLSFREGSNQTTKKEDQAGLELLKRLSEQEVGLPYRGVLGSRPDQDGDTADGSLGSPVLPHSPVPSRDRGILVSSAEWPQLGTPRAELPTSNHELDQPPKQLSTIDEEEPNASPANI
DEYLPSETEQNDDKSASGLDLDSTTEVSQLKRLDPGDQSIPEAFESTPLPSVPKAKSKQVVKTSLGSPVWQDLIAVTSGMVLKQLPEYVKSGNPQYTMEGKTLYTYPHSFNLQAWEAWFFE
>ABRO02016177_Dipodomys_ordii
SYEAGLVPDIFPFFATLVCYYYKGDAALMRKSLFGIHLEGEFWELFAAQDDPFHRVVSLVCPEGTIEIIKQLPPPALHGSHIPAFMVYLTLFPPKLVMGEAACSPKVEPSYQQLEAWGYHNPSSWMVANAIRSVYRAVRLCYLIRFCLIDHVLHQSARTDETDDDIIARSVEQSHFF
GLFMVCSVLTYIITEKSGTIKIHPLARDRWVKEEADHFEHCVNNLQKHGIYAPFPFLGLGVFQTEHVQFPCLSMIALAGVNADARYQALKEVAHKAVLKLQALQCKKEIETCQDLSPEEQVTILAFAKDSDQIKLEQRKRTGRAGIVEEAFRTGSRSTLGSRPDQDGDTADGSLGSP
VLPHSPVPSRDRGILVSSAEWPQLGTPRAELPTSNHELDQPPKQLSTIDEEEPNASPANIDEYVPSETEQDDDKSASGLDLDSTTEVSQLKRLDPGDPSIPEAFESTPLPSVPKAKGKQVMSENIFGVSCVARLDSSDLIRD
>JAHHPX010000042_Dipodomys_spectabilis
HSILKAELAQIQLVQPCVMSPCILCGKGIELRAICFETGLVPDISPFFLQLSYVITYKGDAALMRKSLFGIHLEGEFGELFAAQDDRLHRVVSLVCPEGNIEIIKQLPPPALHGGHIPAFMVYLTLFPPKLVMGEAACSQKVEHPINSWKHGGINIIPQGWMVANAIRSVYRAVRLCYLI
RFCLIYHVLHSARTDEVDDIIARSVEQSHFFCLFMVCSVLTYIIAEESGTIKIHPLARDRWVKEEADHFEHCVNNLQKHGIYAPFPFLGLGIFQIEHVQFPCLSMIALAGVNADTRYQALKEVAHKAVLKLQALQYKKEIQTCQDLSPEEQVTILAFAKDSDQIKLEQ
>PVHN010030180_Dipodomys_stephensi
HSILKAELAQIQLVPPCVMSACILCGKGIELRAICYEAGLVPDIFPFFATLLCYYYKGDAALMRKSLFGTHLEGWSFENYLLRQDAPFHRVVSLVCPEGTIEIIKQLPPPALHGSHIPAFMVYLTLFPPKLVMGEAACSQKVEPSYQQLKARGYHNPSRLQVANAIRSVYRAVRLCYLIR
FCLIDHVLHQSARTDEVDDIIARSVEQSHFFGLFMVCSVLTYIITEESGTIKIHPLARDRWVKEEADHFEHCVNNLQKHGIYAPFPFLGLGVFQIEHVQFPCLSMIALAGVNAYARYQVLKEVAHKAVLKLQALQYKTEIQTCQDLSPEEQVTILAFAKDSDQIKLEQRKRIRQGWNC
RGFQNRKVYLIDGVLGSRADQDGDTADGSLGSPVLPHSPVPSRDRGILVSSAEWPQLGTPRAELPTSNHELDQPPKQLSTIDEEEPNASPANIDEYFPSETEQDDDKSASGLDLDSTMEVSQLKRLDPEIHHPSIVPK
>VMDQ010068370_Nycticeius_humeralis
ELEARVGQIIACFEAGVSIDDVREHLYAAVICYYHDRGHLHLIVKSPFGQYLIADGDEFSTVPPDLNTQPLHRYEPLVAWLGSKSIRDLLAPNPVGKRAASFRAFLSLFLPKLVTGGSASSQKVEKAYQQLEDRGINVISRDCMAPVEMRIAYGTIHHCFIIRFSPMVGKLRLTSHMRL
FGPLSVALPQNTTAWPRLFRSGIRTKFKFKNFGSQKKFQSLYCYLALLTHQFADHCFSLICHELHQQAGSGAGKTVVAGAVESQYSGLLMVKNVLTYMITNEEGDIKLHPLAQGKTKNVLACFEYCVRNLQKHGMTQIEPGKFPSLSAIALGVASSHHSSFKGVSIESYQSLKEAA
HQAELEPQSYPPHKEITSGVLAVREQRVLLKFTRSEKNQSRSRTRGRQGLGRN
>VMDQ010167862_Nycticeius_humeralis
FEAGVSIDDVREHLYAAVICYYRDRGHLNLIVKSPFGQYLIADGDEFPTVPPDLHTQPLHRYEPLVAWLGSNIIRDLPAPNPVGKRAASFRAFLSLFLPKLVTGGSASSQKVEKAYQQLEDRGINVISRDCMAPVEMRIAYGTIHHCFIIRFSPMVGKLRLTSYMRLFGPLSMALPQ
NTTAWPRLFRSGIRTKFKFKNFGSQKKFQSLYCYLALLTHQFADHCFSLICQELHQQAGSGAGKTVVAGAVESQYSGLLMVKNVLTYMITNEEGDIKLHPLAQGKTKKVLACFEYCVRNLQKHGMTQIEPGQFPSLSAIALGVASSHHSSFKGVSIEPYQSLKEAAHQAELEPQSY
PPHKEITSGVLAVREQRVLLKFTLSEKNQSRSRTRGRQGLGRN
>JANKKJ010000033_Dipodomys_merriami
YATIICCHYYHGDTTLMVKSQFSQYLTSDGYEFADMLRELTGAHTLGVLGNGMYTEWLRKVVKDLPQSEPLGGRGAAFMVSLSLFLLKLVTRESAFTGRLKRHTNNWKIKGLISYHPWIKIVFPALQCCLLILFCLVYHTLLQQAWTVDTIIAGAVKQSSFSRLLMISNVVTYIITEH
NAKLNIHPLAQDKQIWEEISQFEYCVKNLCHRIYVPFILGLSGVSQIKDGQFPRLSAIALGVAVVYQNTLGRVNIDRRCQSLKEAACQAEIELQRFISRKMSREITSCLGNLLSQMQMYNHPPLVQADPAQKAHFEPGKTQEGTHNDRLLPSTWVLPHKKDDQTPWPPNSSGSV
HNTCQPTHKSPSDDQREDIND
>ABRO02015169_Dipodomys_ordii

YVTIICCHYYHGDTTLMVKSQFSQYLTSDGYEFADMLRELQEHHSLEEFWAMGCIQSGSVKVVKDLPQSEPLGGRGAAFMVSLSLFLLKLVTRESAFTRRFKRHTNNWKSKGLISEYHSIAIKIVFPALQCCLLILSCLVYHTLLQQAWTVDTIIAGAVKQSSFSRLLMIRNVVTYIITE
HNAKLNVHPLAQDKQIWEEISQFEYCVKNLCHRIYVPFILGLSGVSQIKDGQFHRLSAIALGVAVVYQNTLGGVNIDRRCQSLKEACQAEIELQRFTSRKMSREITSCLGNLLSQMRMYSHPPLVQADPAQKAHFEPDKTQESTHNDRLLRSETWVLPHKKDDQTPWPPNSS
GFRHNTCQPTHKSPSDDQREDIND
>JAHHPX010000219_Dipodomys_spectabilis
YATIICCHYYHGDTTLMVKSQFSQYLTSDGYEFADMLRELQEHHSLRCSGLMGCIQSGSVKVVKDLPQSEPLGGRGAAFMVSLSLFLLKLVTRESAFTRRLKRHTNNWKIKGLISYHPWQIKIVFPALQGCLLILFCLVYHTLLQQAWTVDTIIAGAVKQSSFSRLLMIRNVVTYIITE
HNAKLNIHPLAQDKQIWEEISQFEYCVKNLCHRIYVPFILGLSGVSQIKDGQFHRLSALTFGVAVVYQNTLGGVNIDRRCQSLKEACQAEIELQRFTSRKMSREITSCLGNLLSQMWMYNHPPLVQLTQHKKLISRSQARLKKVHITKTSTILNTGSSTQEGPNTIATKFLRVRQP
THKSPSDDQHEDIND
>PVHN010033176_Dipodomys_stephensi
YATIICCHYYHGDKTLMVKSQFSQYLTSDGYEFADMLRELQEHHSLEEFWAMGCIQSGSVKVVKDLPQSEPLGGRGAAFMVSLSLFLLKLVTRESACHSKVEKAYQQLENQGINIIISPIAIKIVFPALQCCLLILFCLVYHTLLQQAWTVDTIIAGAVKQSSFSRLLMIRNVVRYIITE
HNAKLNIHPLAQDKQIWEEISQFEYCVKNLCHRIYVPFILGLSGVHKLKDGQFPRLSAIALGVAVVYQNTLGGVNIDRRCQSLKEACQAEIELQRFTSRKMSREITCDYPKCGCTTTLPLFKLTQHKKLISRSQARLKKVHITKTSTILNMGSSTQEGPNTMAIKFLRVRHNTCQPT
HKSPSDDQREDIND
>PVIM010143654_Rhizomys_pruinosus
FFIRFSLIHYVLHQRAGAVEQSRFAGLQMVQHFLTYIITKGNENDKYNIHPLAQDNRMKRELTRFEHCIKNIQWHGIYIPLARVLGLAGVSQIEHGQFSLTAIALRVSCIYENTLSGVNLDTRYHYLKQAALKVELELQATQHKKAIRQDTNLDEDERAVLLKFAERSENINKEQHQE
AFSRLLLQLMQSEAGTCLQGVGPAERLSGNSTPRPVDPQCGQIGEGGPASSQQSPTSPHPGLMPEEPVVASTQKNQEKSMLLGGAAEELFY
>JAAGEI010000389_Aeorestes_cinereus
LGNNQFSGLLMVKNNLTYLINNEGGGIKIHPLAQDKMREEMTHFEYCLRNIQKHGSYTLFARVLGLPRSTQIEQGQFPSLSAIALGVSSVYQNPLAGVNIDQYQDEDLTTEEREVLFTFSKRKESIRIKGMRTKPGGFSKRLPTGKWNIQKMRVKYKRRTLHQEICSALTHPHRQG
PHIPRNHSVPKYQYGDRFPQGVESSPGPWLIATSPPSPDTMPKLEDIEPPPISIDDCLSSGIRHPVSVNSGDQDELESIQQDDSETIGGHQHSIPEGFEGKETPEKPQEVHSES
>PVJN01000748_Lasiurus_borealis
LGSNQFSGLLMVKKNDLTYLINSEGGGIKIHPLAQDKTREEMTHFEYCLRNIQKHGSYTLFARVLGLPRSTQIEQGQFPSLSAIALGVCSVYQSLLAGVNIDTPYQDKNLTAEECEVLFTFSKRKESNRIKGNDEDKARLELLKKITDREVELNPKDESQIKEDTSSSDMFCPSSPTPSG
PPYSQEPFCPKIPVWRQFPQGVESSPGAWPIATSPPSPDTMPKLEDIEPPPISIDDCLSSGIRYPVSVNSGDQDELESIQQDDSEATGGDQHSIPEGFEGKETSEKPQEVHSES
>JACAGB010000036_Pipistrellus_kuhlii
MSQHLRSFPGIYLGHTIIVGAVEQSRFSGLLIVKKKINKNKNVLTYIIHILLTMKGGIKIHPLAQDKRTREEMTHFEYCIRNTQKHGSYMPSARVLGLSGATQNEPSLSAIALGVSSVYQSGSALAGVNIDTRYQALTAAAHQVQWAEAATNRGWHGLRHPIRITEHPQQKRNSPG
QRSHLRTGTIYFQKEKDRPIRIKGSVRTKPRWSFSERLP
>CAJEUF010007138_Pipistrellus_pipistrellus
VRNTSPLHRPRPTLPAPGQYHRDPIKPVPPASGTCMSNKVLCDLQLHMFHVVFLRQIIEAFCCLKLKLKLLIILALSLPLVLLVSSLFPSSSMSQYLRSFPGIYLGHTIIVGAVEQSRFSGLLMVKKKKNVLTYIINKKGENKEIKIHPLAQDKRTREELTHFEYCIRNTQKHGSYMPFARV
LGLSGATQNEPSLSAIALGVSSVYQSGGALAGVNIDTRYQALTAAAHQVQWAEAATNRGWHGLRHPIRITEHPQQKRNSPGPRSHLRTGTIYFQKEKDRSIRIKGSVRTKPRWSFSERLPTGKWNIKMSIKERRTLPSLLVSALPRPHRQGPHILRNHPVPKYQYGDNSLKAYYE
SDLLLCLGQLPPPSADAMPELENLEPPHISIDNCLSSGIRHPVSVSSGDQDELESIKGGDSEATCRDHSIPEGFEGNETPGEPREVHLESPSSGTAVQ
>PVIQ01006209_Pipistrellus_pipistrellus
VRNTSPLHRPRPALPAPGQHHRDPIKPVPPASGTCVSNKVLLFATAYVSCGIIPLADNSILRTLKIEIEASDHFGLAYPWSCLCRVASSLRSSMSQHLRSFPGIYLGHTIIVGAVEQSRFSGLLMVKKKNKKMSHILLTMKGENKEIKIHPLAQDKRTREEMTHFEYCIRNTQKHGFY
MPFARVLGLSGATQNEPSLSAIALGVSSVYQSGGALAGVNIDTRYQALTAAAHQVQWAEAEQTGAGMAGIQLELQSIHNKRETAQDQDLTSEEREVPFTFSKRSERIDPSGSKAVGQSPGGASQKDYRQGSGTESKRASNRKGGHFITLMSFCPPSPTLSGPPYSQNHPVPKY
QYGDNSLKAYYESDLLLRLGQLPPPSADAMPELENLEPPHISIDNCLSSGIRHPVSVSSGDQDELESIKGGDSEATCRDHSIPEGFEGNETPGEPREVHLESPSSGTAVQ
>CATLKD010000094_Pipistrellus_pygmaeus
VRNTSPLHRPRPALPAPGQHHRDPIKPVPPASGTCVSNKVLLFATAYVSCGIIPPADNSILRTLKIEIEASDHFGLAYPWSCLCRVASSLRSSMSQHLRSFPGIYLGHTIIVGAVEQSRFSGLLMVKKKKKKKKECLHILLTMKGENKEIKIHPLAQDKRTREEMTHFEYCIRNTQKHGS
YMPFARVLGLSGATQNEPSLSAIALGVSSVYQSGGALAGVNIDTRYQALTAAAHQVQWAEAEQTGAGMAGIQLELQSIHNKRETAQDQDLTSEEREVPFTFSKRSERIDPSGSKAVGQSPGGASQKDYRQGSGTESKRASNRKGGHFITLMSFCPPSPTLSGPPYSQNHPVPK
YQYGDNSLKAYYESDLLLRLGQLPPPSADAMPELENLEPPHISIDNCLSSGIRHPVSVSSGDQDELESIKGGDSEATCRDHSIPEGFEGNETPGEPREVHLESPSSGTAVQ
>JALGBO010000014_Perognathus_longimembris
YVAFHPAELAHIQLLPRVKSACILCGKGVELRVSYIIPHFEARLVPHSSMSVFLHLACYCYQGDAAALMRKSAVGYLGKQQGYKFSELCAAQDDPFHRVVQSLGCPEGTSEVIKQLSPLPSWQPRDSLYGLLLSLFLPKLVMGEAACSPAVEETNEQLGAQGINVKPSSWIVANAI
RSVYRAMICFCLIYQVLHPSAGNDAVDSISARSVEQSRFSGFLMVGDVPIYITTEESDTIKIYPLAGDSRVKEQLSTHREICAPFARVLDLSGVFQTEHGQFPQLSALALEVSAVYQSILADARYQALKEAADRGEYGLVGECLPSMHEALGLDSQHHIHRKSQKWHCGSSGAKRSQ
GQCSGPEFKPQDWQKKKKEAAHKVEFKLQALQHKKKIPTCQDLSPEEPVPSRGWGVLVSSAWNCLQAESRITYLQPTTSTSTTQLLMKPHAPSSNTHEYFPGEAEQDDDRSAGGLDLYSTTEDALARLGLRDQPIPEASESFSLPSIPKAK
>JALGBO010000023_Perognathus_longimembris
HSILTAGLAALEVHPAERVILIGKGLELRSSYTISSFEAEVMPDSLYNHLFVTLICQYYYKGDADFMKRTTLGTYPAQQGFEFSELSVAQQDDPSQRIPEVVPKVLGSDNYLPLIPSMIVAFLSLFFPKLLTGEAICTQKVERFFQLGEAGTNMTPHNWIRTYATPLYKAIQQCYLICFCL
IYQTLGTASIDSITARVVEQSQFSGLFTVRTVLMYIVTEEEGGLRIHPRNELDHFGYCVQDIQQHRIYAPFTCILNLPGVHQIEPRQFPWLPAIALGVSSAYQSILVGVNTDIRHQALKEVAPKAEVKLQVIQSKEVLTSQDLTQGEQEVLLKFRQTQDQRKRQDQEGVELLNKLSEV
RGSPPERRNGHSAGSGWWHYSFPLYPANPPCQGVRTSQLHPPATGHSGPQMLEHQHPAKGQTNLLSIYLLLQRRTLMGYPINIDELLPVEGNPNESVSAGGPDLESTKSPKDPQLGRSKIEDSSIPAVLEAPSPPPHPPHLHRPLSN
>JANJXV010000025_Thomomys_bottae
IHFILKSGLALEEVQLVPPKLRGVILCGKVLELRTSYIMSSFEAEVMPDNLCNHFFGTLICQCWDADFMKSSTLGTYLVQGYELSENPVAQKDDPFHTIVSLGYRKGAIGVIGQLLSLVPSMIMACLSFFLPKLVTEEATCTKVQHLEAQGINVIPCNWIAGNTSPAYLGKCYLICICPI
YCMLHLNSGLLYLIAPGLLTIHNVLTYIVMEEESELGIHPLARDKQSRNELKCFGYHVQESQQHSIYPPFAQILNLPGVHQIEYGRFPWLSAVTLGVPSAYQSTLDGVNMTWYQALKEVAHNAEVKLQKNRTFSLNLPVIEQTAMNRRITLLMRGLDFQQRQAVPRGSPLCPTNL
PHQEMRTSSCISPGTGHSGSQKPQASTPSQPDQTPTYGTCQYTPAWRTILTNIFPKTHSLEDPRLTDHPFLTPLRASIFIVSTSQLDSKKSSGHLPSPQGNI
>JANJXW010000031_Thomomys_bottae
IHFILKSGLALEEVQLVPPKLRGVILCGKVLELRTSYIMSSFEAEVMPDNLCNHFFGTLICQCWDADFMKSSTLGTYLVQGYELSENPVAQKDDPFHTIVSLGYRKGAIGVIGQLLSLVPSMIMACLSFFLPKLVTEEATCTKVQHLEAQGINVIPCNWIAGNTSPAYLGKCYLICICPI
YCMLHLNSGVALLIAPGLLTIHNVLTYIVMEEESELGIHPLARDKQSRNELKCFGYHVQESQQHSIYPPFAQILNLPGVHQIEYGRFPWLSAVTLGVPSAYQSTLDGVNMTWYQALKEVAHNAEVKLQKNRTFSLNLPVIEQTAMNRRITLLMGDWTFSSVRLSRLGVPALSHQP
TPPRNEDIQLHLTWDWPQLGFPKAQASTPSQPDQTPTYGTCQYTPAWRTILTNIFPKTHSLEDPRLTMPSISDTFEGPPYFRAQASDLIQKSPQATSQVLREIQEPMETLPQEEALTYPHSFNTRK
>JAPYYN010045613_Coendou_prehensilis
QIVLLQDTTDTIIAGSVEQSRFSGLIMVYNIVHLYCRVDSRIQVHPIAKGPATREELTQLCCIGLLQRHGYMPFAHLLGLPGIAQIERGQFPQLLAIALGVSSLYQSTLVPLNARYSALRQAAYNAELKLQELKFKREIQINPELSTEEKEVLLDVARTKEGINEQRNRKTRHAEICLKGW
PDNKQTPMPGTFTTPQAVPWADWGDSSRALVASSYSAYNDETCHCPYSTSTSSTNYSLQMTGNFHSMPPLDESDAEDEPKPKSTKPTNQSTVNDLAVAMKDMFATDDDAEVDQPSRVSDPDIHAGDPTVLGTYPGAPCNSYTTLPPIKEET
>JAPYYN010045625_Coendou_prehensilis
QIVLLQDTTDTVIAGSVEQSRFSGLIMVYNTFIYIVVNVDSGIQVHPIAKGPATREELTQFRCIALLQRHGIYAPFAHLLGLPGIAQIERGQFPQLLAIALGVSSLYQSTLVPLNARYSALRQAAYNVELKLQELKFKREIQINPELSTEEKEVLLEVTRTKEGINREQRNRKTRHAEICLK
GWPDNKQMPMLAHSLHHKQYLGQIGVLQGSGGQLQLCIQHETCHRPYSTSTSSTNYPSNEQGISTPMPPLDESDAEDEPKPKSTKPTNQSTVNDLAVAMKDMFATDDDAEVDQPSSVSDADNHAGDPTVLGTYPGAPCNSYTTLPPIKEET
>SWEC01016874_Erethizon_dorsatum
QIVLLQDTTDTVIAGSVEQSRFSGLIMVYNMFIYIVVSVDSGTQVHPIAKGPATREELTQFRCIGLLQRHGIYAPFAHLLGLPGIAQIERGQFPQLLAIALGVSSLYQSTLVPLNTHYSALRQAAYNAELKLQELKFKREIQINPELSTEEKEVLLEVARTKEGINREQRNRKTRHAEICLK
GWPDNKQMPMLAHSLHHKQYLGQIGVLQGSGGQLQLCIQHETCHRPYSTSTSSTSSPSNEQGISTPMPPLDESDAEDEPKPKSTKPTDQRTVNDHAVAMKDMFATDDDAEVDQPSSVSDPDNRAGDPTVLGTYPGAPCNSYTTLPPIKEET
>RJWM01018344_Dasyprocta_punctata
GPLCKLFQQGRSLWISKIVLHAACHTFTRLGPVHKSYCSTDTIVAGSVEQSHFSGLFVVHHVFTYIVVNGDSRIQVHPTAKIPALWEQLTFEHGIRFQQRRRIYAPFAHLFGLPGIARIEHGQFSQLLAISLEVSSPYQSTLAATDARYSALWEQSTNTELKLQELEYKREMQNNPEL
SDEEKAVLLEFSQNKEGINPEQRDKKI
>AFSB01165578_Heterocephalus_glaber
HLSGDILSREKHACSKSHPGIRVHSIAKGSAVQEELTRLEHCIRSLQRHVIYAPFVHLLGPGTAQLEHGLFPQLSAIALGVSSVYQSTLVGVNTDACSLALREAANSSELKQELKFKREMQNKPELSAEEKEVLLEFAQTKEGINQEQRRQEDKTLRLLKKLMARQQADAIPGTSTTL
QGTTQADGAPIGLWWPAEATVHTTPRLATAPTPLAPARPINPPTEQGISTPMPHLEESDAEDKSKPKLTTCTNQSTVDYLANAREDDDEDKMEDQPSSVSDPDGCARDLTTTPEEFHATPTPALPP
>JAAOMF010000026_Arvicanthis_niloticus
LFALFFCFCFSRQGFSVPGCPGTHSVDQAGLELRNPPASASQVLGLKACATTAWQHSFQNTMSTLGCSIVKNLPSALTLERRASPFIAHLSSFLPKLVTGESTCTQVEKTQQFENQGVNIITWMSAVITSITYQTVRHSFLIRFSQVYHVLHQAGIEACFKAGAAETRISAFLIVRNV
LIISNEDGRMIYLLARRRSMYKWQEIYAPFHCVLGLPGVSQIEYGPFLLSAIGFGVSSVLSEYSSSCQVWSQYLKIAFQSELELQAFQHGWEIPTASILLKSKKCYNFLQPVKNINWEQQQEGIMATKEFLRLSEEPGRAGSIMEQEGQRQNKGSVWSTDILPAPWQHQQLWSRA
IKGLAIMCLSLRTKSAEGPRHTCVKEDTDIETRSYSTEDLQVNIGDYLSNESGRPR
>JADRCF010003044_Grammomys_dolichurus
LILKAAQGTRDHNDSICPRTKAVHDLARKETETRVGNIIIYYLIVACFEAGTTIDNICKSLYFTLICFFHYYRDMNRNAFKNYLMSVGVVIDSIPFNKILAYWVAQLRRIYPVHPLERRASSFIAHLSSFLPKLVIGGRTCTQLEKTQQFENQGINIITWMSAVITIITYQTVRHSFLIRFSIY
HVLHYQAGIDAEDIIMAGAAETRISVLLIVRNVLIITNEDSRMIYQLARSVQFGYCVRSVQWQEIYAPFDYVLGLPGVSQIEYGPFLLSPIAFGVSSLYQNTVAVVNMSQYLKEVAFQSELELQVFQHGWEIPTASIWLLKNKKCYNFLQSVR
>SRMG01000026_Grammomys_surdaster
LILKAAQGTRDHNDSICPRTKAVHDLARKETETRVGNIIIYYLIVACFEAGTTIDNICKSLYFTLICFFHYYRDMNRNAFKNYLMSVGVVIDSIPFNKILAYWVAQLRRIYPVHPLERRASSFIAHLSSFLPKLVIGGRTCTQLEKTQFENQGINIITWMSAVITIITYQTVRHSFLIRFSIYH
VLHYQAGIDAEDIIMAGAAETRISVLLIVRNVLIITNEDSRMIYQLARSVQFGYCVRSVQWQEIYAPFDYVLGLPGVSQIEYGPFLLSPIAFGVSSLYQNTVAVVNMSQYLKEVAFQSELELQVFQHGWEIPTASIWLLKNKKCYNFLQSVKNISWEQQQEGMATKEFLKLNEETG
RAGSIMEQEEQRQSKESVWTDILPTPWHQQLWSRVIMGLVPMCLSLRTRPAEGPRYCTCQRRPEIETRSYSTEDLQVNIDEYLSNES
>JAJTUV010002437_Mastomys_natalensis
LILNAAQGTRDHNDSIHPRTKAVHVLARKETETRVGYIIIYYLIVACFEVEIQLIIFEIIYFTLICFFHYYRDMNRSTFKNYLMSVGFVFDSIPFNKILAYWVVSIVKDLPSDPLERRASPFVAHLSSFLAKLVIGESTCTVEKTQEFENQGINIITWMSAVIISITYQTVRHSFLIRFSIYHVLH
QAGSEDIIKAGAAETRISALLIVRNVLIITNEDGRMIYRQEEMSMFGYCVTNVQQEIYAPLIRVLGLPGVTQIEYGPFLLSAIGFGVSSVYQSTVAVVNMSQHLKEVAFQSELELQVFQHGWEIPTASIWLLKSKKCYNFLQSVKNISWEQQQEGMATKKFLRLSEEMEELVPSWS
RRNRGRVRSQCANKLIYSLPLGSMSSSPVIEQGHMGLATMCRLPLRTKPAEGSRHACVKGDTEIERRSYSTEDLQVNMDDYLMNLRRPSWKIRKSRIWCARSNREDKESDIPIPEATG
>FMAL02024890_Mus_caroli
IIMDSTLNVVQGTRDHNDSIHLRTKVVHVLARKETETRVRIYYKIYLHIVVCFEAMVAIDNTCSLYVTLICFFHCRDMNRSAFKNYLLAVGFVIDSILIQNTMSIQIVKGLSSALTLERRVSAFMACLSSFLPKLVIGKNTYTQVEKTQQFENHGISIISWMSAVIIRVTYQTVRHCFLIRL
SIYDVLHQQAGIDAEDIMMAGAAELGLLIARNVLMHIITDEEGRIQMYLLARKRSMLGYCVKSMHWKEICALFGCVLYLPGISQIEYGQILLPTTGFGMSTVYQSILGSQYLREVAFLSELELLVFQYREIPTASIWLLKNEKWYNFLQLVKNINREQQQEGMARKEFLRLTEETGR
AGSIMEQEEQRSSKESVRETDILLTPQQ
>CACVCL010002063_Mus_minutoides
IIINSTLNVAQGTRDHNDYIILRTKVVHVLARKETETRVRIYYNIYYLYILVCFEAMIAIENICNHLLPSASFIVIEIMNRSAFKNYLMAVGFVIDSIPFHKILWSIEIVKGLPSALTLERRVSAFMACLSSFLPKLVTGKNTCTQVEKTQRFENQGINIIPWMSAVIIRVTYQTVRHCFLIRFSI
YDVLHQQAGIDAEDIMMAGAAESRISALLIVRNVLMYIIANEDGRMQMYLLAKEMSMFGYCVKSVQQEIYAPFDYVLYLPGLSQIEYGQILLSTTGFGMPSVYESTLAFVKCVKSISLPFYLNWNCWFFYIDERFTSASIWLLKNEKRHNFLQSVKNINWEQQQEGMARKEFLR
LTEETGRAGSIMDQEEQRPSKESVWESDYTPYPLAAPAVALVIGQGHHRSGLPYVCPFKEQKLYKGPRHGGIKGDTEIVITSYSTEDLQVNINDYLSRESGRDGRTGRLGESVWCAKSNSEDKESESPIPEATG
>AAHX01035392_Rattus_norvegicus
IIMNSTLNTAQGTGDHNDSIQTRTKAVHGLARKEIEARVRVYYNICYLYITACFEAGIAIDNICDHYLTLICFFHYYRDMDRSAFKNYLTAVGFVIDNIPFSNAINILGCPVVKGPTALTLEKKASVSMACLSSLLPKLAIEESTYTQVETPQFENPGINIIPWMSAIIIRVTYQTLRHCFLN
RFSIYHVLHEQAGIDSEDIVMARAAETRISDLLIVRNVLMYIITNEDGRMQMYLLARRNVHGGYCVKSMQWQEMYAPFHCVLGLPGISQIEYGQFLGLSTIAFGVSSVYQSTLAVVSVSQSLKEVAFQPDLELQSFQHRWEIHTTSIWLLKNEKFYKFLQSLKNINWGQEQQEA
MGRKFLGLTECETGRAGSIMEQEEQIQSYESVWKSDYSPYPAAPAVTEQGHHGPGHQVSLPLKTKSLRRAKTCRCQGRPEIETRSSTEDLQINIDDYLSRESAGDSQAGRFNLSQSPLVQHKTDHCLQLLKKKIVDVTWVPAFTSQECTLSLKGGYLITGSTIAYQRPTSWFTASE
NSNLSRKLQLFGL
>JAALGF010004057_Rattus_rattus
IIMNSILNTAQGTRDHNDSIQTRTKAVHVLARKEIEARVRVYYNIYYLYITACFEAGIAIDNICDHLFTLICFFHYYRDMDRSAFKNYLMAVGFVIESIPFLNAINILGCPIVKGSPSALTLEKKASVSMACLSSLLPKLAIEESTYTQVETKSQFENSGINIIPWMSASIIRVTYQTLRHCFL
NRFSIYHVLHQQAGIDPEDIVMARAAETRISDLLIVRNVLMYIITNEDGRMQMYLLARRNVHGGYCVKSVQWQEIYAPFHCVLGLPGVSQIEYGQFLGLSTIAFGVSSVYQSTLAVVSVSQSLKEVAFQPDLELQVFNIDGRFTLQVSGCRMKNFIESFCNHEHQLGTRTAREFL
GLTECETGRAGSIMEQEEQIQSYESVWKSDYSPYPSAAPAVALVTEQGYHGPGHQVSLPLKNKSHRR
>JAPYYD010000003_1_Uromys_caudimaculatus
IVMNSILNVAKETRDYNDSIHPRTQAIHVLAREEMEAITYHIVVCFEAAIAIDNICNLYVTLICFFHYYRDVNRSAFKYYLMAVGFVIDSIPFYKIPAYNWKYLPSARTLERRGSAFMACLSSFLPKLITGESTYTQVEKTQQFENQGVNIIPWISAVIIQVTYQTVRHSLLIGFLNYHVLH
QQAGIDAGNIILAGSAETRVSGLLIVRNVLMYIITKEDGRMQMYREEMSMFGYCMQWREIYAPVDCVLGLSGISQIEYGQFLLSAIAFGMSSVFQSTLAVVNVSQSLREVAFQSGTAGLFNIDGRFTAASILLLKNEKCCNFLQSVENINWEQEQESMARREFLRLTEESGSIME
QEEQMQRKESVWESDYTPYPLAAPAVALVIMQGHHGSGHHVPLPLKNKSHRR
>PVIT010026828_Onychomys_torridus
IALGIRVHNDLILSRTKMVHVLDREALEALVWHILIYIKYIIACFESGIAIDDICKHLMILSICFFHYHGDMNMSAFKNYLMAVDFVINSIPVNVQGLSFYKILDILGCPVVKGLLTVLTLGRASAFVACLSLFLPKLVTGENICTQVEKTYQQFENQGIDIIPWMSAVNIISYQTLRHWVL
IRFSLIYHALHQAGIDAEDTIMAGTSEARISGLLTVRNVLMYIITNGDGRVQIHLLARRNVHVLICVRSMHEIYAPFDCVLDLPGVSQIEHRQFLLSAIAFQVSYVDQSTLAVVQCLKSIPGGCLSSCNLELQDFQYRWVIQNCKCLSDEEFLLKISAVSENINGEQ
>CABHPP010086863_Peromyscus_attwateri

EVRYLVIMHSTLNIALGIRDHNDLILPRTKMVHVLDRKELEARVWYIIIYIKYIVAYFGSEIAIDDIRKHFILLQCCFFHYHRDMNMSAFKNYLMAVDFAINSIPVNVQGLSFYKILDIFGCPVVKGLFTVLTLGRASAFMACLSLFLPKLVTGESICTQVEKTHQQFENKGIDIIPWMSA
VTIKISYQTLTHWFLIRFSLIYHGLHQAGIDAEDTIMTGAAEARISGLLTVRNVLMYIIINGDGRVQVHLLAKRNVHVWICVRSVRWHEIYVPFDCVLGLPGVSQIEHGRFLLSAMAFGVSSVDQSTLAVVQCVKSFPGGCLSSCNLELQDFQYRWVIHNCKGLADEEFLLKISAVS
ENINWEQCSKKICLEKQFLWLTE
>CABHPQ010159732_Peromyscus_aztecus
DSEENLDEVRYLVIMHSTLNIALGIRDHNDLILPRTKMVHVLDRKELEARVWYIIIYIKYIIACFGSEIAIDDIHKHFMLLQCCFFHYHRDMNMSAFKNYLMAVDFMINSIPVNVQGLSFYKILAYLVAQLLTVLTLGRASAFVACLSLFLPKLVTGESICTQVEKTYQQFENQGIDIIPW
MSAVTIKISYQTLTHWFLIRFSLIYHGLHQAGIDAEDTIMTGAAEARISGLLTVRNVLMYIIINGDGRVQVHLLAKRNVHVWICVRSVWHEIYVPFDCVLGLPGVSQIEHGQFLLSAIAFGVSSVDQSTLAVVQCVKSFPGGCLSSCNLELQDFQYRWVIHNCKGLTDEEFLLKFSA
VSENINWEQCSKKICLEKQFLWLTE
>CACRXM010000002_Peromyscus_eremicus
KYIVVCFGSGIAIDDIRKHFMLLQCCFFHYHRDMNTSAFKNYLMAVDFVINSIPVNVQGLSFYKILAYLVAQLRVYSLLILGRASASVACLSLFLPKLVTGESICTQVEKTYQQFENQGIDIIPMSAVTIKISYQTLTHWFLIMFSLIYHGLHQAGIDAEDTIMTGATEARISGLLTVRNV
LMYIITNGDGRVQMHLLARRNVHVWICVRSMWHEVYAPFDCVLVLPGVSQIEHGQFLCLSAIAFGVSSVDQSTLAVVQCVKSFPGGCLSSCNLELQNFQHRWVIHNCKGLADEEFLLKFSAISENINWEQCSKKVCLEKQFLWLTE
>VALE03000005_Peromyscus_californicus
KYIVACFGSGIAIDDIHKHFMLLQCCFFRYHRDMNTSAFKNYLMTVDFVINSIPANVQGLSFYKILAYLVAQLRIYSLLTLGRASAFMACLSLFLPKLVTGESICTQVEKTYQQFEHQGIGIIPMSAVTIKISYRLLHMFLIMFSLIYHGLHQAGIDAEDTIMTGAAEARISGLLTVRSVL
MYIITNGDSRVQMHLLARRNVHVWICVRSMCWHGIYAPLDLLGLPEVSQIEHGQFLLSPIAFGVSSVDQSTLAIVQCVKSFPGGCLSGCNLELQDFQHRWVIHNCKGLADEEFLLKFSAISENINEQCSKKVCLQKFLWLTE
>CABHPR010087390_Peromyscus_melanophrys
RLRSQITGHNHSALNIALGIRDHNDLILPRTKMVHVLDRKELEARVWYIIIYIKYIIACFGSGIVIDDIRKHFMLLPCASFTIIGYEHECLQKLSDDSLCGHQHSCKCSRAFFLQNTMSILGCPVVKGLLTVLTLGRASVFVACLSLFLPKPVTGESICTQVEKTYQQFENQGIDIIPWMSA
VTIKISYQTLTHWFLIRFSLIYHGLHQAGIDAEDTIMTGATEARISGLLTVRNVLMYIITNGDGRVQVHLLARRNVRVWICVRSMRWHEIYVPFDCVLGLPGVSQIEHGRFLLSAMAFGVSSVDQSTLAVVQCVKPFPGGCLSSCNLELQNFQYRWMIHNCKGLADEEFLLKFSA
VSENINWEQDKKVCLEKFLWLTE
>CAJQZJ010034466_Lophiomys_imhausi
VIMHPILKVAVGIRDHNDSIPPRIQGTKVFHVLARKVRGSGLYIIYIVACLEAEIGTDNICNHFMLLSYVCFFYHRAMNRSVFKNYLMAVGFVLTVLLTKPLAHWGAWLRVYSVHLEAVFMAYSLFLPKLIIGESTCTQVGKTYQFETQGINISPWMSAIIIRITYQTLRRCFLIRFSTY
HGQNQAGTDAEDTIMAGAAEAGISGLLTVRNVLMYIILSEDSRIQMHRQEEMSMFGYCVRIVQWQEIYAPFDCVLGVPGVSQIEHGFLLCAIPLGCHLYIGAHLLSVSQSLREAAFKTELELDFHYSGRLITARILLKNEKFYNFLKFVRTSTGNNDSKKVCKRIPVRVTEETGRAG
STCSRKSR
>BPMZ01000043_Tokudaia_osimensis
GGLVGYQREERPLVLERLDATVHVLARKHTEYIIIYDLYIIACFEASIAIANICNHFITLICFFHYYVMNRSTFINYLMAVEFVIEVFLLQNTMSTQTVKGLPSALTLERKASAFMACLSSFLPKLVIGEITCTQEEKTYQFENQGINIIPWMSAAIIRVTYKTLKHCFLIRLSTIMSCTNMLV
YADDIIMARARISGLLIVRNVLMYIITNEDVGCKCVCQEEMSMFGYCVKSVQWQEIYAPFDCVLGLPGVSQIVCGQFLLSTIGFGVSSVLSEYISCCQVCERQSLREVAFQSLELQVFQYRWEIHKLKSEKSKTLLKFSAISENINWEQQQESMARKEFLRL
>VFHZ01020188_Meriones_unguiculatus
VIIHSVLNVALGSRDHNDSDLARTKAVHVLARKELEAGEILSFTLYIVACFEAAFATDNICNHFVTLTYTLHYYRDMNKCVFKNYLMAVGFVADRIPFSRILAVAQLRVFPEHPLGRRTSVFMTYLSLPPTLVIGESPCTQVEKTYRENQDMRISPWISAIIIITYQTLRHCFLIRFSIYQV
LYQQAGIDSEDTIIAGVSKARISGLLIVRKVLIYIITNGENNANASVRQEVSVFGYCVVCRGMRFVPFDCVLGLPGVSQLEHGHFSLSVIAFGVSSIRVHLLSNVSPSLKETAFQAELELQGLQYRWDSPTVRIWLLKSENVCCSFLQSVKNINWEQQQQQEGLTRKAFLRLTEETEEL
VPRWSRKGKCRVCHQDRNRLRHSLGSLSSSPGDRAGTSWP
>JAQQSO010000019_Pachyuromys_duprasi
HSVLNGALGIRDHNDSDLARTTVVHVLARKELEARVGNIIYSYILACFETEIATDNTCNHFVTLTYTFHYYRDMNKCVFKNYLMAVGLVVDRLPFYRILDMIGCPVVKGLFRALTLGRRTSVFMTYLSLLLPTLVIGESPTQVEKTYCENQDMLISPWISAIIIITTRLLRHCFLIRFLLNY
QVLYQQAGIDSEDTIIAGVSKARISGMLIVRNVLMHIITNGDSNANASFSRSVHVWTLCCVQWHEIYAPFVCVLGLPEVSQIEHGHFLLSVIAFGVSSVYQSTLAIVNVSPSLKETAFQAELELQGLQYSWGIDNCKDLAIKEKILLQFSAVSENLNWEQDGLTGKAFLGEPSEKLEE
LAPWGSRKSKCRVWSYRTDYTLYPLAASAVALVIQQGHHGNDHNVPLPFEEQKQWKGQDMQVRDTLKVIMYRGSSRKYLFVYQIREDGKGGRGQLRVWCARPNSEGKELDISIPGAMGPLSEPCRRRGHKGVPAFTSECKRSLKYEHLTGSIITQRSIPLTESENYNPSRKLQ
LFCLLGELATRDPGSRPSLSTRGSCEHNGLKLDDAPAILSP
>AALT02121933_Sorex_araneus
DVVPIFSASFASTLLPTLVVGKAARESKAKRVLERLRAPEMNLIPGKWSLGVKVQSLYRLLHCSYAFRLAVIHHAPEHVSGTDQVERMTLDRVTRVRYSGMNISTHILAENDHGVTFTFSMELPRSWQPAPFARILGFPEMSLLGHGQFPKLSAIALGTASAPLLSGTVTDPQLRRL
RRGTRLASFPLGPQLHDQATRHRVQTQEPATIQDQVTGLQGCDHCHQLNPSSPGHEQLVQEIKPKEAHDETPSQPPGTTSASQSADRTLSEDNDNTAATQ
>snake_FAA04057_Tapajos_virus
MSQSSLHSFLKTGMPSLSHADKVNKHRDTRVKIVNVNLCSNLRRVCSEIILCFIGSFDTHEFQDEIVLLLIVQFYYAGDAQRFKSSIFCQQLLEAGYQINFYSSKLEGRLLDCLAPLKEDSRLAAVLKNILLPTDDSDQTPALFALMSMFLPKLMVGAAACATKVGRTVEKLIQQGICPI
PPEWVNTTVLESIYRVMRASYILRFAIIHHALEQSHGSDSNERIIHDCVMQARFAGMLLPVTMLTHILVEEPEGVRLHLLARSQDMKEELTRFQLAIRSLLRHGRLTPFARILGLPEMALLEHGHFPKLSAIAMGVASVHTPTLNATVTDPRYEKMRRAAEEAERALHDYQNAEM
VDNDATLSAGQREILKGFHKQRSAIRDAETTAAEELRRETITRALKQATNPTNAPATKISQKPTTATQTATSPPKKSATTHLPPTTITFLPVPDTPERESMTPRTEVSEWSNLPDISAYFANPSGLGSIQEEEETIQHTAHPSNLDNTNTTGGDHPRLDISSPLSDVLPRSNSSFEDLLI
LEDDLPLPPPPSPPPQASENDQYIEALNLIGQHLSGFPEEIAKKFICDHQDLIDLILSSQPGLDSIGKPYQLSIDGGEMTSDQVNMLLSFLKITSLDVAIANKHDRLLSFPSAFEDRNAWDFLIPATGSTPLGQVQLQNGSLARVEDLTQKGIMIYWILGERSK
>AAKN02025587_Cavia_porcellus
LYKLFHRTSLIYKIVLHHACHIFSKLGPIHKSYCSKILLTSCEISGSHSSGLFMVYSAFTFIIVSVGSRMKCIQVHFIAKSPILREELSQFKHCIQFLERHGIYIPFAHLLGLPGIAQLEQGQFSQLLTIALMVSSPYQSTLICATVAHHYNTLGKQPVTLNRLQEKLQREIQNHPELSEEKKVVL
LVFANKEGTNLEYKDREDKTHRFSGEANDQTTSQCQCWYIHHTTSSNLGRLECSSRDSGGELQLQHVPQETCNYLSSTSTSSTNYPIEQGFPTPVPPLEGSDAEDKPKPLDKTHKSEHSGSSNAMRHIFAENDEAKVKDQPSNVSDPDSHAGHPTIIPGEFHATPTCTFIPVKDET
>PVKK010009749_Cavia_tschudii
LYKLFHRTSLIYKIVLHHACHIFSKLGPIHKSYCSKILLTSCEISGSHSSGLFMVYSAFTFIIVSVGSRMKCIQVHFIAKSPILREELSQFKHCIQFLERHGIYIPFAHLLGLPGIAQLEQGQFSQLLTIALMVSSPYQSTLICATVAHHYNTLGKQPVTLNRLQEKLQREIQNHPELSEEKKVVL
LVFANKEGTNLEYKDREDKTHRFSGEANDQTTSQCQCWYIHHTTSSNLGRLECSSRDSGGELQLQHVPQETCNYLSSTSTSSTNYPIEQGFPTPVPPLEGSDAEDKPKPLDKTHKSEHSGSSNAMRHIFAENDEAKVKDQPSNVSDPDSHAGHPTIIPGEFHATPTCTFIPVKDET
>BDUI01000013_Apodemus_speciosus
SIKIKMTDFMLFECSAGTRDHNDPVPPRTKEVHVVAGRGQEIRSEHSLACGEVGLEVCSLCNHFHAVLICYCAADGFTISIVTIDTQSLPFYKIIRTLDCLDGVTCVIRDLPPHPPLEMKGSALMAHQSLFSPKLVTRENVYTYKVENTHQQLKNWGTSIIVWEISAFRSTSQTLLPYF
SLTHLLFQAATGCGRDFTAGTVKQARPSREGKLQIHPLAPDRMRVQMSRFKGHGENSQQGTNVPFARVIGLPRGIMDTGQFLTLSTIALEGSSMYQSTLQSVNTDVCYQPLKEVVLHQAKLELDSNIGRKFTIAETWPSQGKTNPMPFLHIRNISLEQQEDTDRKRFLHKLRRIL
QGQSTGVVVNGHAGPAPVGGNSSGPGLTVAPNHQATAYHVFTLAKSRSPGHFDDKRDPTTFHSRPSDSDSFCLLENEGPLQLEEGNILSGHLSRAGGKERSRQSIPEAV
>AEKR01234560_Mus_musculus
TRSIKMKSTDSLLFECSAIPPRTKEVHIVARRAGLRQLSCGEGLTICALCNYFHAVLICYWQQMASLVLTLDAQSLPFYKIIRMLGYLAGVTRVIRGLPAEPTLGMKASAIMAHLSLLFSKLVTRENVYTKVENTYQQLKTGPSIIVWEWISAIRSTSQMLLPYFSLTHAHLQAAAGCG
RHFKAGTVKQARPSSLLMVRNAPTYIITQENGVTIYPLAPDRMRVEMSGFEYCRQELTVPFTCVLGLPRGITNTGQFPTLSAIALTVSSTYQSTLTSVNTDVCHQPLEVVHQVKLERDSSIVRKFTIAETWPSQGKTNSIVSCDIRNISPEQEQEEWIGSEFLYKLTENLVGPVYRWN
SRQLRTPWPSSHQQQQWSARLPVTPKSQASAYHGLPLPRAGVQDTSMTREIQPYSTQDPQINSDSFLFIEDEGPLVLEEEGNVLSAPLTKAGGEAERTRQSIPEAVEDPAQEKHYPQFLKRQGLRSIGSQYLHTMEYRRTLKQIAGSLGPRGGGGGSSLVKGTHS
>JAAOME010000011_Arvicanthis_niloticus
QPPPRTKEVQVVAGRSWGSRAEHSSACGQGLRPDAICNHLHAVLICYCAADGFIICDNYYRCSEPYFLNKRTLDSLGGVTCVIKNLPAKPPLGMKDSAIMAHQHLFFSKLVTRRSRVWGWIYDAIRSTSQTLLPYFSLTHVLHQAATGCGRPFTAQTVKQARLSGILMVRNVPK
HIITQENGKLQIYPLAPDRMRVEMSRFYYGKNLQHGTYVFFASVLGLPRGITDRKGQFLTLSVIALGVSSMYQSRLVSVNTEVCYQPLKEVVYQARLEFQDSNIDRKFTIAETPSQWKGKSYLSFLHIKNISLEQQEDMDRKQFLKLRRLWWGRSTGRVAVNGCSGPTPVGGRRR
AGGSQWPERSTHCLPCVYPAKSRSPGHSDYKRDP
>JAAOMG010000013_Arvicanthis_niloticus
PPRTKEVQVVAGRSWGSRAEHSSACGQGLRPDAICNHLHAVLICYCAADGFIICDNYYRCSEPYFLNKRTLDSLGGVTCVIKNLPAKPPLGIKDSAIMAHQHLFFSKLVTRRNRVWRMDICHKYRPRHCFLIRFSLTHVLHQAATGCGRPFTAQTVKQARLSGRLMVRNVPKHII
TQENGKLQIYPLAPDKKSEMSRFYYGKNLQRGTYVFFASVLGLPRGITDRKGQFLTLSVIALGVSSMYQSRLVSVNTEVCYQPLKEVVYQARLEFQDSNIHRKFTIAETPSQWKDKSYLSFLHIKNISLEQQEDMDRKQFLKLRRLWWGRSTGRVAVNGCSGPTPVGGRRRAGG
SQWPERSTHCLPCVYPAKSRSPGHSNYKRDP
>JAJTUV010000186_Mastomys_natalensis
LNAALGLEYHNDPTSPRTKEVHVVAGRGGAGLSTACGEGLSVCAICNHLHAVLICYCAADGFVICDNYYRCSEPYFLSKAWSELLHVIEDLPAKPSLGMKDSAILTHHLFSPSLPGGTFEDGYMPLEVPPRHCFLIRFALTHVLHQAATGCGRHITAETVKKARLSGILMVRNVPKYI
ITQENGKLQIYPLAPDRMRVEMSRFHCGKNLQRHGTSVPFVSVLGLPRGITDRTGQFLTLSVINLGVSSMYQSILVSVNTDVCYPLKEVVYQAELEFQDSNIDRKFTTAETWPSQWKDKPYLSFLHIRSISLEQQEDMDRKQFLHKLRKLWWGWSTGRVAVNGCAGPTPVGG
DSRVGGSQWHRTINSLLTMCLSCQEQESRTLRLQKEILPHSTQDPQINSNSFLFTENGEASQTRGRLMSVLLRQRQVVKRKNQTI
>REGO01000293_Rhombomys_opimus
LLTLLECSAWTKDHNDAIPPRTKEVHTLVGKGLGARAEHIIASFQVGLETDGICDHFKAFICYRHQRGFIISRLLTIRGLPFYKILRTLGCPERVTWVIKVFPAEPPLGMKVSAFMAYQSLFFLITRENACIWKVEKKSQQLKNWGTNVILWKWIFAIRITCSIFRHCSFSFLSSMHQEA
GVHAVRQFTGGAIKQARPSGLLMVRNSPTYIITKENGKLQIGPLAQEKMRVEMPRFQYYAKNQDHRIHVFCRVIGLPGLSQTEEGAFPQLPATALGVSPRHQSPPARFSTDVCYHPLKEAALQAELELGSNTDRKFTIAESPAEPAQERQVLRFPLTIRDISGEQQREVMDIKELL
HKLRERGILGQQRSSSQRKDRTSTTAVGGSGRDCSRAGGTEPLHWPSMPKSRRPAQLDGKGNTRVETNTFPTEDPQKILSSGHGSPLELEEEGNSPSAHVSKGRRRERTRQSVPEAADPAQGKHHPQFLNWKRGGPDLLGSRYVLSWTQSACCTL
>JAQJZG010000003_1_Microtus_californicus
TQLSYRFSLIDHVLHQQAGTHTVDSSLVGAVKQDRFSGLLAVRNVPRYIVTEENGKLQIHPLARDNVRVEISRFRILCQELQRHGDLCPFAQVLGLPGSSQIDHEQFPRLSDVTLGCHPCVRVQWQVSRHICHQSLKDAGHQVSELQDSRKARNLQQQRLGTAKTRQILRKFSPS
RTSNRDDKRKTQMRKELLCKLTGGKAVGTRSADRAAVTTSIDGSSGGPSGHGVTELTTLTLCGGYSYADFSISGMSYSSEMEGTLGDPDLEAGRQAFDLDHGVEGTALIWATPAAGSLYKDSGRGKVCSSPACPPPVSTSIPSLALESASSGFQHTQKTSDTQP
>AAIY03007415_Echinops_telfairi
MHSILIVALGCFWDSRIPPRIKNKVLYGEGTEAWINQILSRSWDWIAVDDTCDHLYVTLICFSHSQGDVTLMAGNAPWQFLVLDGYEFQSVPHDSLEEPLHLPRALECPEGLVQVIKDLPTQAHLATFIAYLTLILPKLLTGAMACTQEVESSFQQLENEGISIIRGWRIQIPVTMRI
AYRPLCQCSLRFCLVHHALRQQAGTVGVDTIIASAVEQSGFSGLLMVQNVLTFLTYSIAHKNGNLKIHPLAQRKGRKERTHFECIHNLQRQPRHLRSFASALGLPGVTQTERGKFLRLSVIALTCADARYPTRQNLSDDWEIRQNNDSSEAEEAILRKWVDKINQAQWEQDHAQ
LNLLRQLAKGEINRAGPWGSWPRSSKRWRVLVVYHLGPVSPTPQPGSQQTQGPGWQTPPPPPLVNLQAETRRSGHEHKGHPYHRKGGPGVRPRIFSRRKTPSSRKLQGRVSGVDPFGATSGRLHSQQSVSWRER
>AYUG01161637_Fukomys_damarensis
MGRRSVASHALGYKRSLLLPLYAPATSLLGCACVLLQETCLVNISWCLSALFLQQTESGVCHHFYATLVCHCHHQGELKDGEDYLTETAISSLHHKNIFRVSGYIELPEVWIFRCDQSSPGPPVFPQWPGAFIAYLSLYLPKLVTREDAFLWKVEQSFQQLECQGISIINREWLSATV
VQVVYGTVHHCFIIHFSLVFHTLHDNNTIIAGSVEQSQFSGLLMLCNVLTYIVVNSNSGTQIHPTAQGSAIQGEPTQPEHRCRSLQKHVVYASLYVSLGYQLEHGQLLPWGVSSVYPSTLVGANTDACSLALREAANSAELKLQEYEIQEIRNNPQGQCGPLSLLKPSKAPSKSRED
RKTRHTEICSRGWPDNKQPPTLAHPLRCKEQPQAAGAPVGLQWPAEVTDAYNTETSHCPYSTSTSSTNYSTDRMTGDDSHSNSSPKGIRCRGRAQSQVDR
>JAMXIF010000058_Petaurus_breviceps
QIQVHSEQVLAEFLQTWTSTATVKIIFALMWQSFPLKFVIIHQQAEHDASDSIIAAAIAARFASLLIVKTILDHILLCTKDRVELYPLARGRGLRNKLDGFQHAVKEISRNAIHTPERELWSLNADRSILFVFFLLFCFVLFLLSCGSFHWLFFAALICKYFSCECICRAYIRLHAVLGREERK
MGRKFGTQKLVELKVVNKKLIFKKYRNKKKISTHGIYRLLNLAGMNQLDLLDRSKLTEEHEKSPSRIQQNIDVLGKTTICPKPELQLINPAGINTSISLVTSFKPPYTNFQLTPLALKSYTTLQMAGEEEISLSPTDFHDYFREEPSHLAGILEDQMELTNCECEEDTDPVTDIEDPGGAL
HTCSPTKPTYLPTTAFSTTPTPPGPISQGSIKLESTAKFPSYVGPLHHNETGKALYLTTFTGKPLVFPDTMISDEHSGKK
>RAZU02000005_Cricetulus_griseus
ITDCLQARTYLGEIDNCVLMLYVEHYFKKDIKASQASTFAYNIDHGFRYLSKDRRRYCVIPSDREGENMRQEIRQLASVPCGVSKMRLGLFLVYYSFFIPKLVMGDRACLEILHRKLQVHREQWLIKFPAWSLGLLGISFLIHGSFASKSIFIHQSNHPSGHAATEIIIKHLVVQAKFSG
MLRRVCSLIISFVVQKGLASTSQSSACLCFLVPILKAHTTVVIFISLLTNHIQYNFILLLAIGESLQKYSFEHLSTLLITVVILLQSQLCLYFLDTDSYHKEYLWKSTRKAKPSTLHHVECLNPQRKKKEVIVHFGERKTQGPF
>LZPO01067497_Neotoma_lepida
GTDLEEEIDNCLLKLNVEYYYQKDIKGFQAITFVKYLIDHGFDIFQRIGEDNPSXVGVIHIDSAGENLRQEIRQLVLCLVGKLAKLGLFLVYYSFFLPLVMGDGACLEILHPSFSSRTGLIIFPXAWSLGRTLGIFSILIHGNFAFKFILIHQSSHPSGHDTSEITTNILVAQAKFAGMLCLVFL
DYIFCWTEKGXEVHPLVRSQSI
>CABHPP010130550_Peromyscus_attwateri
ITDCLEARTDLGEIYNCLLTLYVEYNFKKYIRDFQVSIFVEYPIDHGFDIFQRMGRDNPSLVDVIRVDSVGENLRQEIRQLVLCLVGKLAKLGYFLVYYSFFLPKLVMGDRACLEILHPSFSSRTGLIIFPASLGRTLGNLLLFGTWKFCFKFILIHQSNHPSGHDATEIIANNLVAQAKFA
GMFSVCILDYIFCWTEKGVEVPPLVR
>CACRXM010000007_1_Peromyscus_eremicus
ITDCLEARTDLGDIYNCLLMLYVEYNFKKYIRDFQVSIFVEYPIDHGFDIFQRMGEDNPSLVDVIRVDSVSENLRQEIRQLVLCLVGKLAKLGLFLVYYSFFLPKLVMGDRACLEILHPSFSSRTGLIIFPATLGNLLLFGTWKFCFKFILIHQSNHPSRHDATEIIANNLVAQAKFAGMLS
VCILDYIFCWTEKEVEVPPLVRSQSL
>CABHPR010085395_Peromyscus_melanophrys
ITDCLEARTDLGEIYNCLLTLYVEYNFEKYIRDFQVSIFVEYPIDHGFDIFQRMGEDNPSLVDVIRVDSVGENLRQEIRQLVLCLVGKLAKLGLFLVYYSFFLPKLVMGDIACLEILHPSFSSRTGLIIFPASLGRTLGNLLLFGTWKFCFKFILIHQSNHPSGHDATEIIANNLVAQAKFAG
MLSVCILDYIFCWTEKGVEVPPLVRSQSL
>VALE03000011_1_Peromyscus_californicus
ITDCLEARTDLGEIYNCLLTLYVEYNFKKYIRNFQVSIFVEYPIDHGFDIFQRMEEDNPSLVDVIRVDSVSENLRQEIRQLVLCLVGKLAKLGLFLVYYSFFLPKLVMGDRACLEILHPSFSSRTGLIIFPATLGNLRLFGTWKFCFKFILIHQSNHPSRHDATEIIANNLVAQAKFAGMLSV
STFLDYIFCWTEKGVEVPPLVRSQSL
>CABHPQ010168316_Peromyscus_aztecus

ITDCLEARTDLGEIYNCLLTLYVEYNFKKYIRDFQVSIFVEYPIDHGFDFFQRMGEDNPSLVDVICVDSVGKNLRQEIRQLVLCLVGKLAKLGLFLVYYSFFLPKLVMGDRACLEILHPSFSSRTGLIIFPASLGRTIGNLLLFGTWKFCFKFILIHQSNHPSGHDATEIIIKHLVAQAKFAG
MLSVCILDYIFCWTEKGVEVPPLVSSQSLGTGGWRDASTISPVALAEDQSFVCLVFVCFRECLTVTWPAWNLLCRPGWPQIHGALPASVS
>CABHPH010056853_Peromyscus_nudipes
LLMLYVEYNFKKYIRDFQVSIFVEYPIDHGFDIFQRMGEDNPSLVDVIRVDSVGENLRQEIRQLVLCLVGKLAKLGLFLVYYSFFLPKLVMGDRTCLEILHPSFSSRTGLIIFPASLGRTLGNLLLFGTWKFCFKFILIHQSNHPSGHDATEIIANNSVAQAKFAGMLSVCILDIFCWTGK
GVEVPPLVRSQSL
>JAOPKW010000016_Peromyscus_maniculatus
HCLEARTDLGEIYNCLLMLYVEYNLKKCIRDFQASIFVKYPIDHGFDIFQRMGEDNPSLVDVIRIDSAGENLREIRQLVLCLVSKLAKLGLFLVYYGFFLPKLVMGDRACLEILHPSFSSRTGLIIFPAWSLGRTLGNLLLFGTWKFCFKFILIHQSNHPSGHDASEIIANNLAAQAKFAG
MLSVCSLDYIFCWTEKGVERLHSLVRSQSL
>JAOPKX010000007_Peromyscus_maniculatus
ITDCLEVRTDLGEIYNCLLMLNVEYNFKKCIRDFQASIFVKYPIDHGFDIFQRMGEDNPSLVDVIRIDSAGKNLRQEIRQLVLCLVSKLAKLGLFLVYYCFFLPKLVMGDRACLEILHPSFSSRTGLIIFPAWSLGRTLGNLLLFGTWKFCFKFILIHQSNHPSGHDATEIIANNLAAQAKF
AGMLSVCSLDYIFCWSEKGVERLHSLVRSQSL
>RCWR01033593_Peromyscus_maniculatus
ITDCLEARTDLGEIYNCLLMLYVEYNFKKCIRDFQASIFVKYSIDHGFDIFQRMGEDNPSLVDVIRIDSAGENLRQEIRQLVLCLVSKLAKLGLFLVYYGFFLPKLVMGDRACLEILHPSFSSRTGLIIFPAWSLGRTLGNLLLFGTWKFCFKFILIHQSNHPSGHDATEIIANNLAAQAKF
AGMLSVCSLDYIFCWTEKGVERLHSLVRS
>RCWS02024208_Peromyscus_polionotus
ITDCLEARTDLGEIYNCLLMLYVEYNFKKCIRDFQASIFVKYSIDHGFDIFQRMGEDNPSLVDVIRINSAGENLRQEIRQLVLCLVSKLAKLGLFLVYYGFFLPKLVMGDRACLEILHPSFSSRTGLIIFPAWSLGRTLGNLLLFGTWKFCFKFILIHQSNHPSGHDATEIIANNLAAQAKF
AGMLSVCSLDYIFCWTEKGVERLHSLVRSQSLGTGGWRDALMISPVAPAEDWSLVCLVCVCLREYLTVTWP
>CALSGD010001501_1_Phodopus_roborovskii
CQITDCLEARTDLGEINKFVLMLYVEHYFKKKKKKKKKKRHAFQASMFAKYTVDHGFDIFRMGEDIPNMVYLIHTDRECETEYAISTVLCGVSKMRSGLFLVYYSFFLPKLVMGDRVYLEIIAQAVVHKQWIIVFPAWSLSRTMGIFYLIHGSFAFRFVLIHQSNHPSGHAATEIIIKH
LAAQAKFPGMLSVCSLDYIFCCTEKGVEVHPLVRSQNL
>PVIH01010838_Sigmodon_hispidus
ESLIRSRIIFPLDEHEPGKLCLQIIHAIEAVITQGDLLDSCPITSCVHTYGRTHRVFQQSAAHCYIEVPGLMLEALDYKHGDLSDFSPIRSRTSPRQALILLNKEPDHSSGQVLFAVFLKLYFLRSALNRPQRQTAIHAKWRTPYSQCWLTGFKMIFCEIRILSFPFKFLLIRHIHDTVMQIV
SQAKFFQLPIIWTALSHILSKEGDPIILDHAVRTVHVPERIEIFRLALKSLTFFGAYALFARMLNLSVNNVENEVFPQLTAIALDVAPAGGGTLAGVIIGEGYQQLGEAAREAHRVQEHATGRENQIGFDSAEGKILLDFHK
>AVM87232_Xilang_striavirus
MDANQVKIFSDIASTPSDSYVSRSKLRVYASKAPDLLYGLLVAALLDPELPRSTAASVFVSLFIGHFYRGDITSFSTSSIAKELTKFGHNLIYVEVDIDVASRTIISPKASRLSQYIKKAELLTDRTLERFITYLAAMMPKLMISIDAVHAKVVKTDAVLKSQGIPSLGDVLIHKATVTTVRAH
LMDSFPGKYLLIKLTISQVSSGSVCGGQIKTLIDQARMAGLVLIKLFTDHLISGSGGKGNKSWHPILSLKDLEREAGDMVAAINGISAHGDLAPFARILGLSGVEKVEYGKFPKLAAVIVGIGQAHNSSLNLMVVGAAHKPLCEAARLFEESRKKDGEAKLGGMFPTQVTQAEKRRL
ADFHKDMTKATEESLKAAKAAQMRTLQAIRASVGGPGSAVGQPKMVTFEDERDEEHHVGNLTVRRRHREYDSDDYSYSIDQAEYGFDPPEEDLDDEVDDEDEEDEEVGPPPPPGESRPPGDGPTWTPPPPATGGPSAPSRDVNRNQIPPPASGFNCVEVAGHFGLQAGN
KRYTAQDYTAWDKEDLPKDVADFQQGSQILTTHGALIAGVEADCKSIRCIFDKKPFTFPQDYRNKDLLTMMFPKTDKYKKYCLLYGTRELSVLRVNNTTLHYALFNGSRDIQAGLLKGQSYYGTAGKEILT
>JABUMV010022313_Paedocypris
MDPKDLPHLASLRPSAEFAYTTRQRLILYTAKATPWLLACLISVLFDPDASRQEVASAFCSIFLTYFLNRPLTDCIEVPAMAELTGLGYAVDVTDLTPGDPKRTLKLTKSPLINRHMRSGELLEVTERSLERFVLYVAAMLPKLMISADTVQIKLVKVQKNLRSQSQEMFPDLWMNK
AVITIVRNHFRDCFPGKYLLIKLTSTRVSGGIAYGTQIHTLVDQAKMSGLVLVKLFNDHLITHDGPHGSKQWHPILSMETLTPEAVALVKAIKALSDHGDLAPYARILALSGVEQIEYGKFPKLAAVIIGIGKAHNPTLNMMTMASAYTNLANLAYTYEIKRLASTKRNPERPPFLRPP
QAPSSSSRYMTSRPVWPSSLRRTYRE
>AALT02088749_Sorex_araneus
SNRTTNQKDIRKKILQVIIYSITHSCDVIVTWFTGAFDPGEALNTFVITLMLGTKKSMRASNFLQHLQQAGFQLEVVSFHQNWTLVDCLQKCVRNQNIIVAAERIRTNFHGKDVVLIFFAFTSMMLPKLVVGEAASEIKVKIIIERLRAQGTNPIPDRWLQGVTTQDLCRSLQCYIF
RFAIIHHALEQCTDQVEQIVLDCATQARYSGIFPVSILTYILIENDHGVILHLLKSHHLKEGLSQFLSAIHSLHHGQLAPFACILLFPEMPLLEHRQFPKLSVIALGTASVHPTLCGAATDPRNDHEIDNNNEDFSQNLFKSPHQRGECKCMVGTTGGSPPAEGRLSFQLETQLRNMP
PQPSSPSTKISPNSVEAPGTPKPSQGRKQTAFQITKKPMINMPELDALFAPEEQSLKRIPEDPVTESQGSEASHKLNTCSPGHISDDVCDGATPPLAPEQPLGPSGRESKKEGSQDENQLSLTIGHHPPIRRPKPEASAIQELQALLGEIRTEKVGHATKNTQYFLVDGRQLPLEKTT
PGLYNALDVEASPTDGSQRCSNVLTTLLDPMDIKHD
>JAOANZ010001337_Sorex_cinereus
SNRTTNQKDIRKKILQVILCLSLHICDVIDCHLVTGAFDPGEALETFIITLMLGTKRRMRASNFLHLQQAGYQLEVVSLHQNWTPVDCLQKCVRDQNIIMATERIRTNFQGKDMVPIFFAFTSMLLPNLVVGEATSELKAKRIIERLQAQEINPIPEEWLLGVTIQGLCHSLCYIFRFAI
IHHALEHGRQTKWRELYTVSRVTQAWYLGIFPVNILIYILVENDHGVILHLLTQSHDLKKGLSKFLSAIHSLHHGQLAPSAQILGFPTMPLLKHGHFPKLLSVIALGIASVHQPTLCGTATDPRNDHDIYNNKDLTSIEREIRKNFHRTCSNLHAQAAESDKMHGEELLEALHPQNEV
DFFPAGNSTPQPGDTIPCPSSKALDHQPDKVQIQWRLLEPAQNSPKAGTDCFPDHKEANDEYAALDETQVLFVPEEQRLERISEDPVAELHVSEAYHKLNICSPGHASDDVYDGSTPPPALEQPLEQSGQESKKEGSQDENQLSLTIDHHLQSEDQNTEASAIQELQVLLGEIRT
EKVGHAAMNTQYQFLVNGRQLSLEKTTLGLSALWIGTNRFSERCSNILTISLDPMDIKHWDPFLPENSHVRAGYVVLEKGELTCSHYVSVEAHTGLWIKATHS
>JAOXYA010000028_Sorex_palustris
SNRTTNQKDIRKKILQVILCLSLHICDATVTCVTGAFDPGEALETFIITLMLGTKRRMRASNFLHLQQAGYQLEVVSLHQNWTLVDCLRKCVRDQNIIMAAERIRTNFQGKDMVPIFFAFTSMLLPNLVVGEATSELKTKRIIERLQVQEIYPIPEEWLLGVTIQGLCHSLCYIFRFAII
HHALEHVRQTKWRELYTVSCVTQARYLGIFPVNILTYILVENDHGIILHLLAQSHDLKEGLSKFLSAIHSLHHGQLAPFARILGFPVMPLLKHGHFPKLLSVIALGIASVHQPTLCGTATDPRNDRDTDNNKDLTSIEREIRKNFHRTCSNFHAQEAESAKNAWWELLEALHPRNEVD
FFPAGNSTPQLGDTTPHPSSKALDHQPEKSMIQWRLLEPAQNSPKARTDCFPDHKEANDEYAALDALFVPEEQSLERIPEDPVAELHVSEAYHKLRICSPGHASDDVCDGSTPPPALEQPLGQSGQESKKEGSQDENQLSLKLTNTTLQSEDQNIKASAIQELQALLGEIRTEKVG
HAAMNTQYQFLGDGRHLSLEKTTLGLSDTLDMAPTDSHERCSNISLDPMDIKHWDPFLPENSHVGAGYTVLEKGELT
>PVLD010007574_Dinomys_branickii
IAPRSKTIVLEGAFLYASMTEIIACFRASLTIDKLRGQFYDTLACYYHHQGDPSIIKSLFGEHLTWKGFLCTTEPHLGEQLSWIARSMGWTDVIRALLMHTSAHCAGAFMPYPSLYPPKLVIGKQACTQKVDCSFQEEHQDIHLRNWEQLPTVLVPVGYWEMYHCFLIWFSLVFYT
LHHQLGTVVTIIAGHIEQCLLSRMLMVHNTMSLVESEDNQTSRTLYLDHSYDLTSILKCLECLQLSLKSLLKPRIHMTSYMIYQTGTNRSPKISEVCYSLDLAKQPPLKTARVQSWPSKASIREFGSRFFLQLMGLRSHKWKATSNTCQYLNGNSFLHGFYKQVFDTVEPLNLLKLTP
QCGRPTFVQHETTSLSSCHFASRHSNSYCLLVRRILRSPKSPSCPSMSIPYSPSLADPRAMITHHGSPFAPKCQLCCAPFISSSLSLHSLLPHFPPHFPSNLYILSEARHRDSKCLVLLDNTKTISKQKIDIGKQEENLHGPILQVLSFYACSKRARCQQMLAQQ
